# Adipocytes regulate fibroblast function, and their loss contributes to fibroblast dysfunction in inflammatory diseases

**DOI:** 10.1101/2023.05.16.540975

**Authors:** Heather J. Faust, Tan-Yun Cheng, Ilya Korsunsky, Gerald F.M. Watts, Shani T. Gal-Oz, William Trim, Kurt Kongthong, Anna Helena Jonsson, Daimon P. Simmons, Fan Zhang, Robert Padera, Susan Chubinskaya, Accelerating Medicines Partnership Program: Rheumatoid Arthritis and Systemic Lupus Erythematosus (AMP RA/SLE) Network, Kevin Wei, Soumya Raychaudhuri, Lydia Lynch, D. Branch Moody, Michael B. Brenner

## Abstract

Fibroblasts play critical roles in tissue homeostasis, but in pathologic states can drive fibrosis, inflammation, and tissue destruction. In the joint synovium, fibroblasts provide homeostatic maintenance and lubrication. Little is known about what regulates the homeostatic functions of fibroblasts in healthy conditions. We performed RNA sequencing of healthy human synovial tissue and identified a fibroblast gene expression program characterized by enhanced fatty acid metabolism and lipid transport. We found that fat-conditioned media reproduces key aspects of the lipid-related gene signature in cultured fibroblasts. Fractionation and mass spectrometry identified cortisol in driving the healthy fibroblast phenotype, confirmed using glucocorticoid receptor gene (*NR3C1*) deleted cells. Depletion of synovial adipocytes in mice resulted in loss of the healthy fibroblast phenotype and revealed adipocytes as a major contributor to active cortisol generation via *Hsd11*β*1* expression. Cortisol signaling in fibroblasts mitigated matrix remodeling induced by TNFα- and TGFβ, while stimulation with these cytokines repressed cortisol signaling and adipogenesis. Together, these findings demonstrate the importance of adipocytes and cortisol signaling in driving the healthy synovial fibroblast state that is lost in disease.

## INTRODUCTION

In rheumatoid arthritis (RA), inflammation drives synovial fibroblasts to proliferate and mediate inflammation and cartilage and bone destruction in the joint [1, 2]. In healthy people, the synovium is composed of lining and sublining layers largely composed of fibroblasts [3]. The healthy synovium often displays marked adiposity, with the sublining, especially in villus areas, intermixed with adipocytes [3]. Knee joint synovium is also immediately adjacent to and interconnected with the infrapatellar fat pad. Adipocytes are similarly intermixed with fibroblasts in adipose tissues. During osteoarthritis (OA) and RA, the normally abundant synovial adipocytes largely disappear and are replaced by inflammatory and fibrotic tissue [4, 5]. These findings suggest a high level of communication between the adipocytes and fibroblasts in healthy conditions, but little is known about if adipocytes influence fibroblasts and what role they play in fibroblast function [6].

Adipose depots contain a specific cellular landscape that is tuned to support adipose tissue function. For example, adipose stromal cells secrete IL-33 that supports T regulatory cell survival and contributes to an anti-inflammatory homeostatic state [7]. Given the adipose-like state of the healthy synovium, we asked if adipocytes might drive a distinct phenotype of fibroblasts in the healthy synovium and how that might change when adipocytes are lost in pathologic states. To assess this, we performed single cell RNA sequencing on healthy and diseased synovium and defined transcriptomic differences between healthy and inflamed synovial fibroblasts. Among the changes detected was a program characterized by genes involving fatty acid metabolism, lipid transport, and metal ion homeostasis. Remarkably, adipocyte-conditioned media was able induce these healthy fibroblast programs in cultured RA synovial fibroblasts, and fractionation of adipose-derived lipids and mass spectrometry analysis identified cortisol and glucocorticoid signaling as the primary pathway required for maintaining the healthy synovial fibroblast program. These same programs were evident in general in fibroblasts from classical visceral and subcutaneous adipose compartments suggesting that the adipose environment is a key driver of healthy fibroblast physiology which is lost in pathologic states.

## RESULTS

### Single cell RNA sequencing reveals distinct healthy and diseased synovial fibroblast populations

We collected healthy synovial samples from knees of ten subjects undergoing autopsy who lacked a diagnosis of arthritis, history of autoimmune disease, or recorded traumatic injury to the knee (table S1, cohort 1). We stained healthy synovium with hematoxylin and eosin, demonstrating that the synovial lining layer is two to three cells thick and is adjacent to a sublining region composed of loose matrix with significant adiposity. In contrast, synovium from OA and RA subjects exhibit immune cell infiltration, fibroblast hyperplasia, and reduced adiposity (Fig. 1a, fig. S1a). Oil red O staining confirmed the presence of lipid-filled adipocytes ∼50-80uM in diameter in healthy synovium (Fig. 1b, fig. S1b). We quantified total neutral lipids in CD45- stromal synovial cells by flow cytometry with LipidTOX staining (Fig. 1b, right). Lipid content was highest in fibroblasts from healthy subjects (85%) versus RA samples (48%), suggesting that RA fibroblasts reside in a lipid-depleted environment.

**Figure 1.**
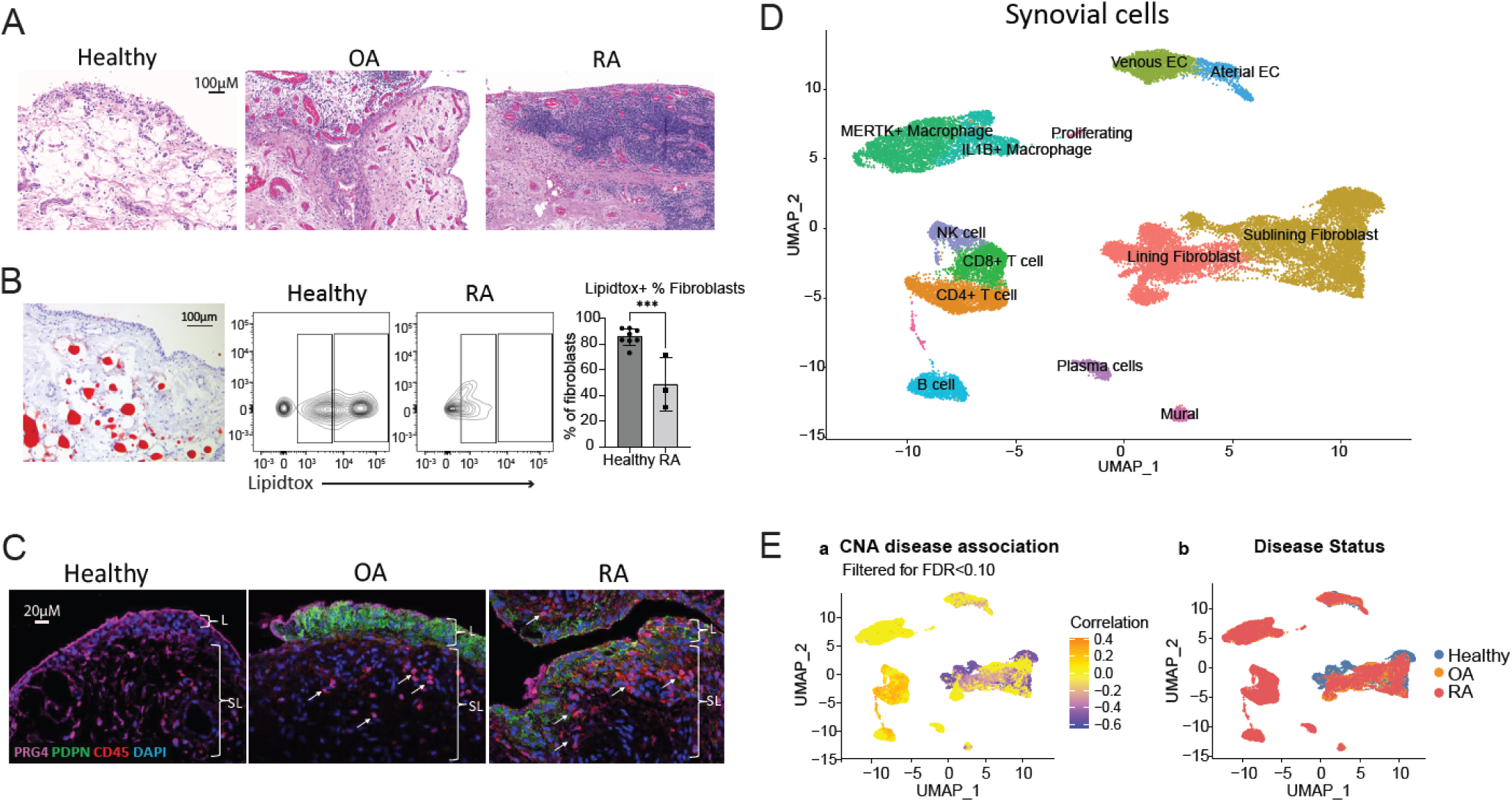
Single cell RNA sequencing reveals distinct healthy and diseased cell populations in synovial tissue. (**A**) H&E staining of synovial tissue sections highlighting broad histological differences among healthy, OA, and RA tissue. (**B**) Left: Oil Red O staining of healthy synovium. Right: LipidTox neutral lipid staining of healthy and RA fibroblasts (live, CD45-, CD31- CD146- cells). CD36 is on the y-axis. (**C**) Immunofluorescence staining for PRG4, CD45, PDPN, and DAPI in healthy, OA, and RA synovium. (**D**) Synovial cells from healthy, OA, and naïve RA donors were harmonized and clustered into a single UMAP. (**E**) UMAP projection from (A) colored by correlation with arthritis (orange) or health (purple) using covarying neighborhood analysis (CNA). All data are presented as mean ± standard deviation.

Next, we performed immunofluorescence staining on 3 healthy, OA, and RA samples, which revealed a smaller number of CD45^+^ cells in healthy synovium compared to OA and RA samples (Fig. 1c). Staining for the synovial lining marker lubricin (PRG4) showed that healthy synovium has a strong and well-defined border of PRG4 staining in the lining, which appears more irregular in OA and RA lining. The stromal cell marker podoplanin (PDPN), which is increased by TNFα exposure, is low to absent in healthy tissue, whereas OA and RA tissue have marked staining (Fig. 1c).

By applying single cell RNA sequencing to 10 healthy synovial samples, we obtained data on 8,687 cells that passed quality control, including 7,607 fibroblasts, 893 endothelial cells and pericytes, and 187 immune cells. We used the Harmony algorithm to integrate data from these 10 healthy synovial samples with 9 OA and 28 treatment-naïve RA synovial samples obtained through the Accelerating Medicines Partnership: RA/SLE consortium [8]. Clustering resulted in 13 cell states: lining and sublining fibroblasts, CD4 and CD8 T cells, NK cells, B cells and plasma cells, *IL1*β+ and *MERTK*+ macrophages, mural cells, venous, and arterial endothelial cells, and proliferating T and B cells (Fig. 1d, fig. S1c). Healthy synovium contained a mean of 83% fibroblasts, 1% T cells, 10.5% ECs, 0.2% B cells, and 1.6% macrophages on average. The proportion of T cells differed substantially among OA (8.7%), RA (30.2%) and healthy samples (1%). Similarly, B cells were 4.8%, 6.7% and 0.2% in OA, RA and healthy samples, respectively, and macrophages were 26%, 24% and 1.6%, respectively (fig. S1d). We also implemented covarying neighborhood analysis (CNA) to identify groups of cells that covary in abundance between healthy and diseased synovium [9]. Many cell populations are transcriptionally different and distinctly associated either with disease or healthy state (global p value=0.001) (Fig. 1e).

### Fine grained analysis defines synovial fibroblast ground states

We performed fine-grained clustering of healthy fibroblasts to identify fibroblast states in homeostasis. Seven clusters were defined including: *PRG4*+, *PLIN2*+, *DKK3*+, *CXCL12*+, *CD34*+, *APOD*+, and *VCAM1*+, each named by a characteristic highly expressed gene (Fig. 2a- c). *PRG4*+ and *VCAM1*+ clusters included fibroblasts enriched for lining markers, including *PRG4*, *HBEGF*, *CRTAC1*, *FN1*, *HAS1*, *HTRA1*, *TIMP1*, *CLU*, and *IGFBP5* (fig. S2a). The *VCAM1*+ cluster contained fibroblasts that also express genes downstream of TNFα signaling, including *VCAM1*, *TNFAIP2*, *TNFAIP6*, and *NFKBIA*. *DKK3*+ fibroblasts (expressing *IGBP6*, *ASPN*, *COMP*), *CXCL12*+ fibroblasts (expressing *SFRP1*, *CHI3L1*, *CHI3L2*), and *CD34*+ fibroblasts (expressing *MFAP5*, *FBLN2*, *FBN1*, *APOE*) all appear to be similar to previously defined synovial sublining fibroblast clusters, as suggested by the ratio of odds of mapping an RA or OA fibroblast cluster to a defined healthy fibroblast cluster (fig. S2b, [8]). The *APOD*+ (apolipoprotein D) cluster is enriched for genes that modulate growth factors and insulin signaling (*IGF1*, *IGFBP3*, *IGFBP7*), an adipokine gene (*RARRES2*), lipid transport gene (*APOD*) and monocyte attraction through *CXCL14*. Many highly expressed genes are secreted factors. The *APOD*+ cluster is also enriched for metallothioneins (*MT1X*, *MT1A*, *MT1M*, *MT1E*, *MT1G*), which are cysteine-rich proteins that are involved in homeostatic regulation of the storage and transport of metals, and protection against oxidative stress [10]. The *PLIN2*+ (perilipin 2) cluster expresses genes involved in regulating lipid homeostasis and metabolism (*PLIN2*, *HILPDA*, *ADM*), as well as matrix degradation (*MMP2*, *CTSK*).

**Figure 2.**
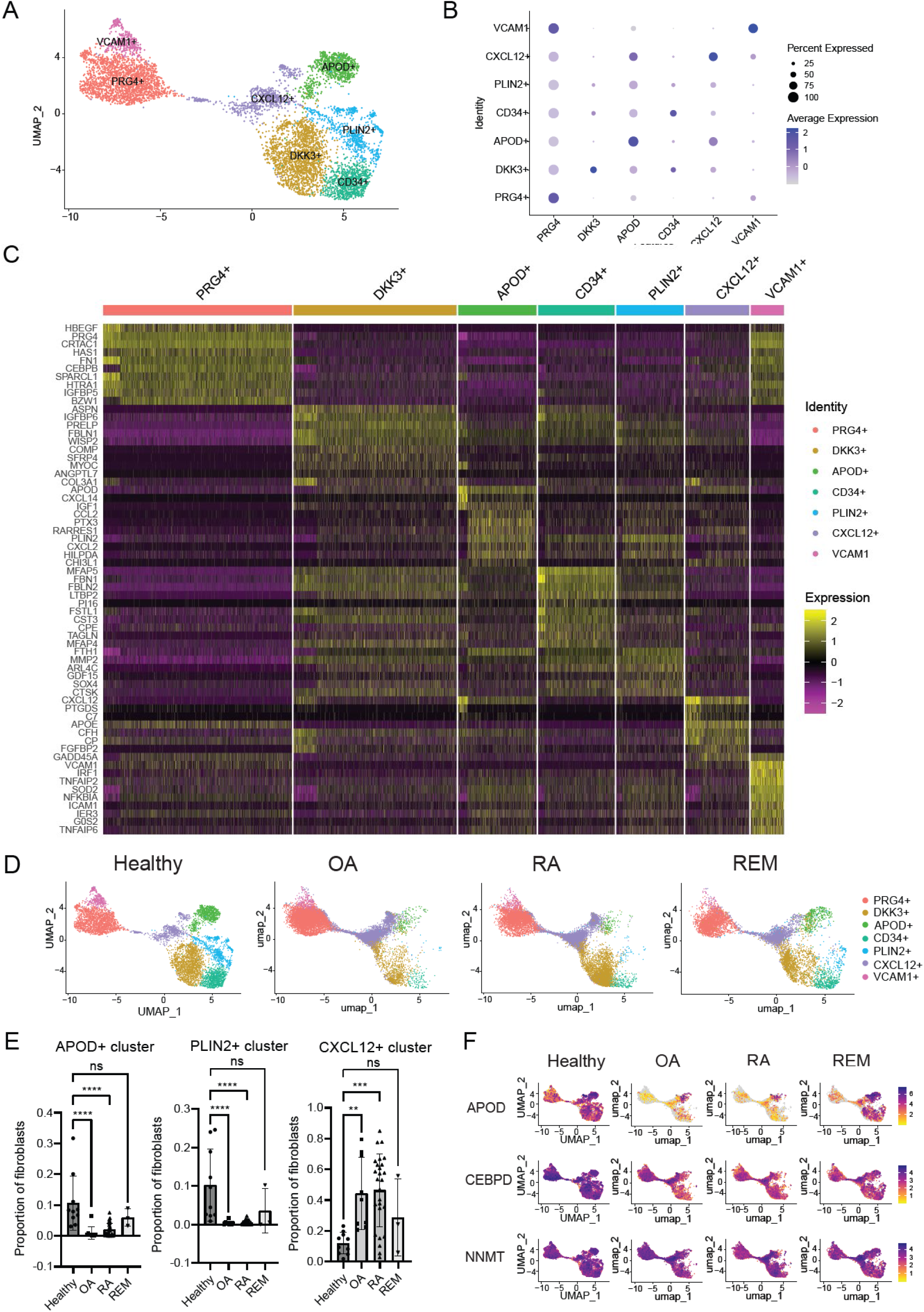
Synovial fibroblast signatures during homeostasis. (**A**) Fine clustering analysis on healthy synovial fibroblasts defines 7 distinct clusters. (**B**) Top markers of each fibroblast cluster. (**C**) Heatmap of the top 10 DEGs per cluster. (**D**) Symphony mapping of OA, naïve RA, and remission (REM) fibroblasts to healthy synovial fibroblast reference. (**E**) Quantification of fibroblast proportions mapping to each cluster. Statistical comparisons: all groups were compared to healthy. (**F**) Healthy synovium is enriched in APOD, CEBPD, and NNMT expression compared to OA and RA fibroblasts, and is partially restored in remission fibroblasts.

After identification of these healthy fibroblast clusters, we used the Symphony algorithm to map fibroblasts from OA, treatment-naïve RA, and RA remission samples [11] onto the healthy donor dataset (Fig. 2d). Strikingly, both RA and OA samples were significantly depleted in the blue colored *PLIN2*+ cluster and the emerald green *APOD*+ cluster, and significantly over-mapped to the purple *CXCL12*+ cluster, suggesting that *PLIN2*+ and *APOD*+ fibroblasts are lost or significantly changed in disease, while *CXCL12*+ fibroblasts are expanded (Fig. 2e, fig. S2c).

Further, differential gene expression analysis of all sublining fibroblasts revealed that healthy fibroblasts globally upregulate 540 and 571 genes and downregulate 953 and 882 genes compared to RA and OA fibroblasts, respectively. Many top upregulated genes were involved in metabolism, including apolipoproteins (*APOD)*, lipid droplet associated proteins (*PLIN2)*, progesterone induced genes (*DEPP1)*, metallothioneins (*MT1X*, *MT1E*, *MT2A)*, hormones (*ADM)*, transcription factors (*CEBPD*, *ZBTB16)*, and methyltransferases (*NNMT)* (Fig. 2f, fig. S2d, data files S1 and 2). Notably, APOD, CEBPD, and NNMT are highly expressed in healthy synovial fibroblasts, but they are reduced in OA and RA, and partially restored in remission (Fig. 2f).

### Healthy fibroblast signature can be induced by fat-conditioned media

Healthy fibroblasts express a gene signature enriched in metabolic pathways compared to OA and RA fibroblasts. Genes upregulated in this signature include *APOD*, *PLIN2*, *DEPP1*, *ADH1B*, *MT1X*, *CEBPD*, and *NNMT* (data files S1 and 2). Of these genes, we selected three of the most strongly upregulated genes compared to OA and RA fibroblasts to represent the healthy fibroblast gene signature: *APOD* (Apolipoprotein D), *NNMT* (Nicotinamide N-methyltransferase), and *CEBPD* (CCAAT/enhancer binding protein D). APOD is an apolipoprotein which is present in HDL and involved in cholesterol homeostasis; NNMT is expressed in adipose tissue and methylates nicotinamide, thereby determining the availability of methyl groups for regulating gene expression and metabolism; and CEBPD is an early preadipocyte transcription factor.

Due to their lipid-centric functions, we tested if coculture of RA synovial fibroblasts with oleate and palmitate, two of the most abundant fatty acids, might regenerate the lipid-related healthy fibroblast phenotype [12]. Oleate and palmitate failed to upregulate *APOD,* but they modestly upregulated *NNMT* and *CEBPD* (fig. S3a), suggesting that fatty acids are not likely to be the primary inducers of the healthy fibroblast phenotype. To more broadly test substances released by adipose tissue, we generated abdominal or synovial fat-conditioned media. Synovial fat-conditioned media induced robust expression of *APOD*, *NNMT* and *CEBPD,* with *APOD* expressed 30-fold, *NNMT* 3.5-fold, and *CEBPD* 4-fold over basal levels (Fig. 3a). Abdominal fat-conditioned media was also able to upregulate these genes to similar levels (fig. S3b). Next, we pursued isolating the active molecule within adipose tissue that is responsible for the induction of these genes.

**Figure 3.**
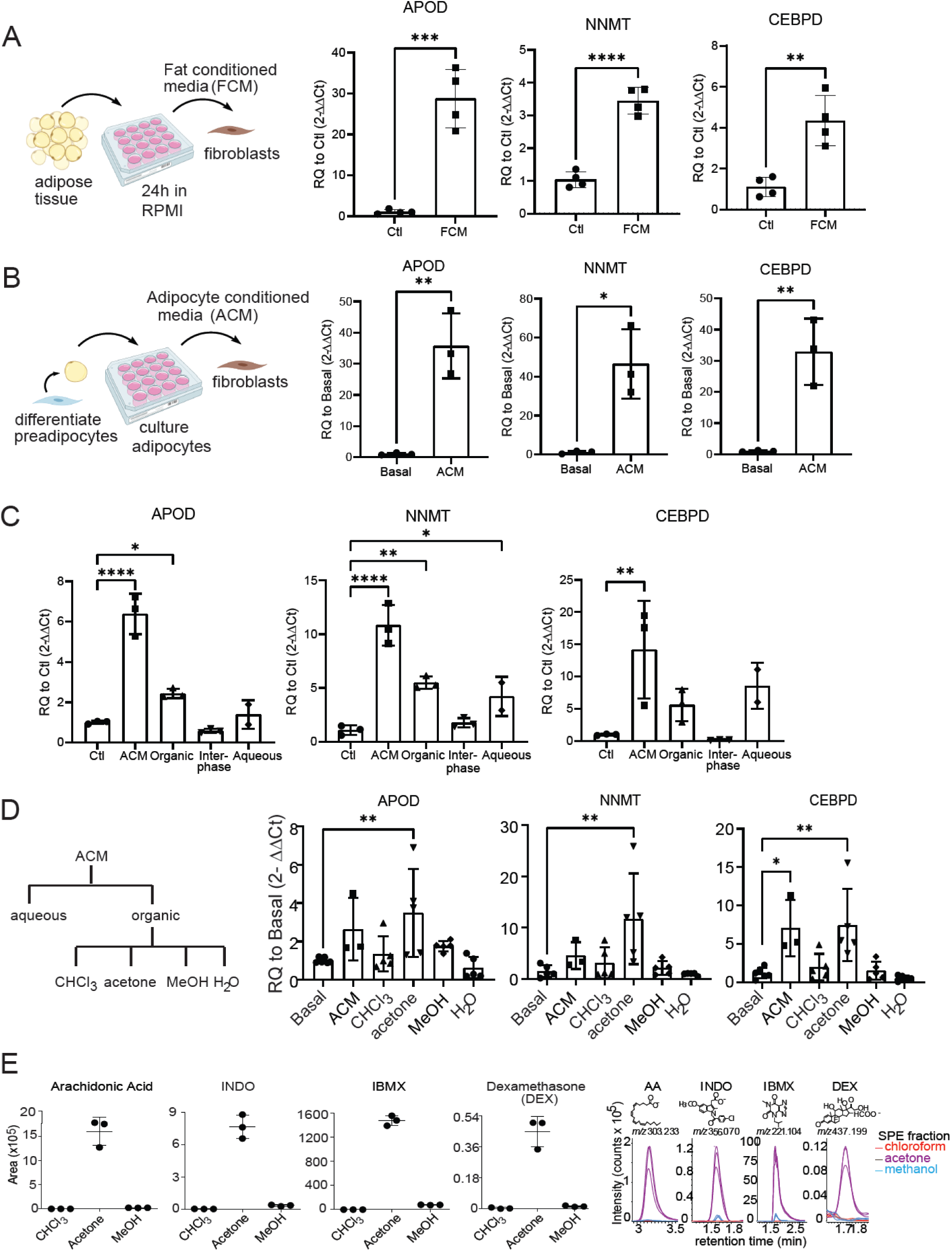
Search for active molecule driving healthy fibroblast cell state. (**A**) Left: Schematic illustrating generation and application of fat-conditioned media (FCM). Right: FCM induces healthy fibroblast signature. (**B**) Left: Schematic illustrating generation and application of adipocyte conditioned media (ACM). Right: ACM induces the healthy fibroblast gene signature. (**C**) Bligh and dyer separation of adipocyte conditioned media (ACM). (**D**) Left: Schematic detailing fractionation approach. ACM was separated using the Bligh and Dyer method into aqueous and organic phases. Then, the organic phase was taken for solid phase separation and eluted based on polarity using chloroform, acetone, methanol, and water. Right: Testing activity of each fraction. C and D statistical comparisons: all groups were compared to basal/ctl. (**E**) MS intensity values were shown as the ion chromatogram areas extracted at m/z 303.233 ([M-H]-) for arachidonic acid (AA), m/z 356.070 ([M-H]-) for indomethacin (INDO), m/z 221.104 ([M- H]-) for isobutylmethylxanthine (IBMX), and m/z 437.198 ([M+COO]-) for dexamethasone in all three fractions.

### Search for active molecules using fractionation and lipidomics

Since fat-conditioned media might contain factors from any adipose tissue cell or component, investigated the effects of conditioned media generated from cultured adipocytes on synovial fibroblasts. We cultured pre-adipocytes in adipocyte differentiation media for 10 days and applied that media onto fibroblasts. This strongly induced fibroblast expression of *APOD (35- fold)*, *NNMT (45-fold)*, and *CEBPD* (35-fold) expression (Fig. 3b). This result, together with those using fat-conditioned media (Fig 3a), suggested that adipocytes were the likely source of the active factor.

Next, we carried out a Bligh and Dyer extraction of conditioned media, which separated lipid and non-lipid material into organic and aqueous phases, respectively. Bioactivity was found in both phases (Fig. 3c). However, upon a second round of extraction from the aqueous phase, most activity extracted into the organic fraction (fig. S3c). We then further separated the organic extracts by the solid phase extraction (SPE) method to elute lipids based on their polarity using chloroform (most apolar), acetone, methanol, and water (most polar). Bioactivity was contained in the acetone fraction (Fig.3d). We analyzed the lipid contents of acetone fractions by reverse phase high performance liquid chromatography (HPLC) coupled with Electrospray Ionization (ESI)-Quadruple-Time-of-Flight (QTof) mass spectrometry (MS). We used the unbiased software based (XCMS) lipidomic method ([13]) to identify target ions with at least 10-fold higher intensity and corrected P value <0.05 in the stimulatory acetone fraction compared to the non-stimulatory chloroform fraction (fig S3d left). There were 779 ions that met these criteria. Among them, we focused on the ions with highest fold change and intensity (fig. S3d, right). By plotting *m/z* value versus retention time (fig. S3d right), three clusters were formed based on their elution pattern and mass profile. Eight ions were identified as fatty acids based on matching their mass to known compounds.

Two other ion groups with similar retention times (1.6-1.8 min) represented alternate ion adducts and multimers of two underlying unknown structures of known mass (fig. S3d, green and pink). The low mass and very high intensity of these ions prompted us to consider three chemical additives, indomethacin (INDO), isobutylmethylxanthine (IBMX), and dexamethasone (DEX), which are supplemented in the adipocyte differentiation media. Indeed, the molecule present as 4 ions at 1.6 min, whose deprotonated ion [M-H]^-^ at *m/z* 356.071 matched the expected mass of INDO. Next, the molecule present as 7 ions at 1.8 min was solved as isobutylmethylxanthine (IBMX). However, the 3^rd^ chemical additive, dexamethasone, was not extracted by automated peak-picking software, which might be due to its low intensity. We therefore manually searched the ion chromatogram, which yielded a chromatogram peak that matched the expected DEX ([M+COO]^-^ m/z 437.198). DEX was highly enriched in the acetone fraction compared to the chloroform or methanol fractions (Fig. 3E). Manual inspection of HPLC-MS chromatograms for arachidonic acid ([M-H]^-^ m/z 303.233), INDO ([M-H]^-^ m/z 356.070), and IBMX ([M-H]^-^ m/z 221.104) confirmed the unbiased lipidomic results (Fig. 3E, right).

Since the mass spectrometry analysis implicated that activity co-purified with several of the adipocyte differentiation media additives, we tested the media directly without one factor at a time. This revealed that dexamethasone, a glucocorticoid, accounted for the dominant activity (Fig. 4a). IBMX had the second largest impact on activity, which may be due to its role in raising intracellular cAMP levels which sensitizes cells to glucocorticoid signaling [14].

**Figure 4.**
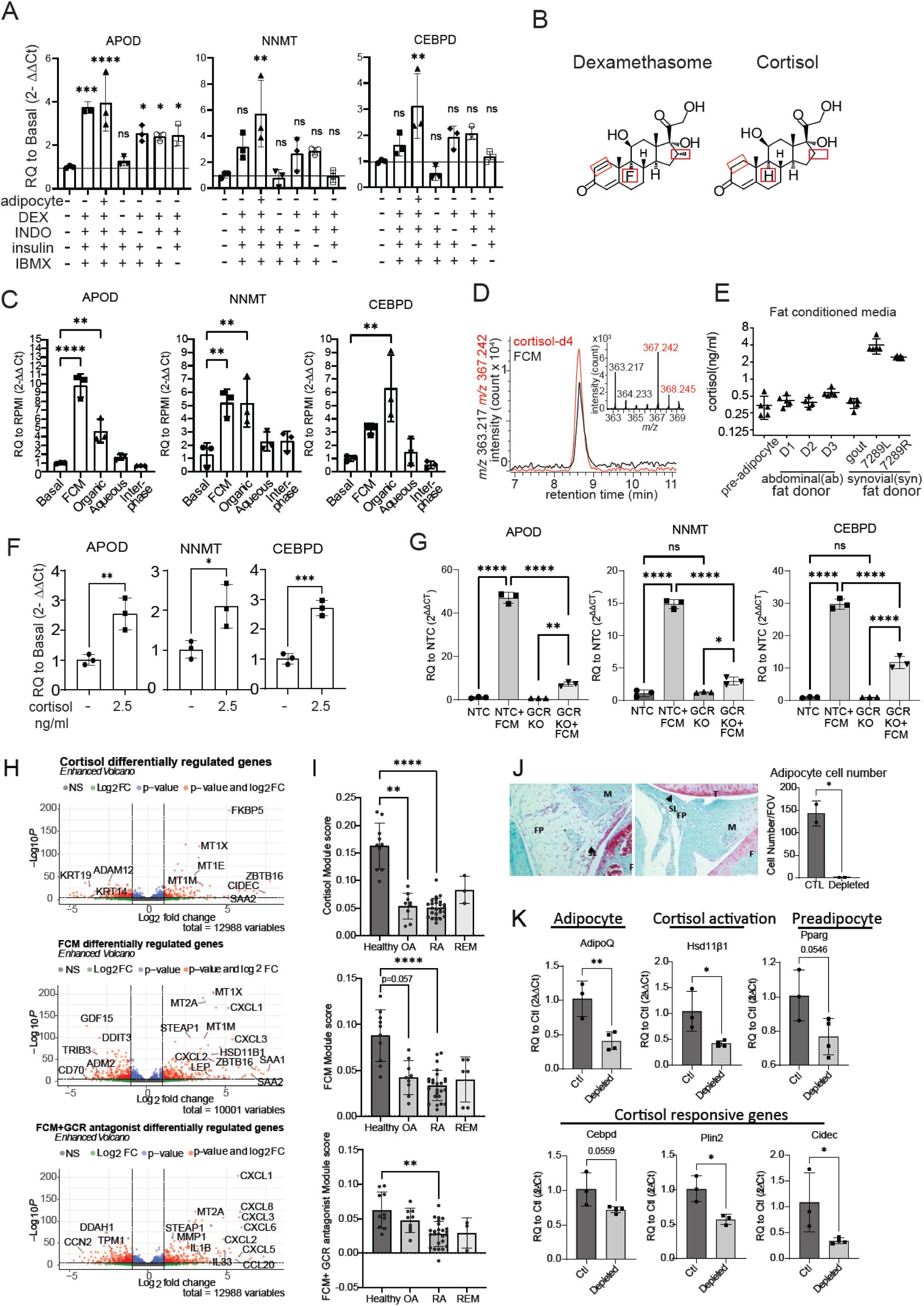
Glucocorticoid signaling is sufficient and necessary for healthy fibroblast phenotype. (**A**) ACM media was tested in the presence or absence of adipocytes and the most enriched species: dexamethasone (DEX), indomethacin (INDO), insulin, or 3-isobutyl-1-methylxanthine (IBMX). (**B**) Dexamethasone is a synthetic derivative of the naturally occurring glucocorticoid cortisol. Chemical structures are shown with changes boxed in red. (**C**) Bligh and dyer separation of fat-conditioned media (FCM), the active molecule is in organic and aqueous fractions of FCM. (**D**) FCM contains cortisol, as measured by the positive mode mass spectrometry using cordisol-d4 as the internal standard. (**E**) Approximate cortisol concentration in different FCM donors based on quantification of mass spectrometry data. (**F**) Cortisol, at biologically meaningful concentrations, is sufficient to induce healthy gene signature. (**G**) CRISPR cas-9 knockdown of NR3C1, the gene encoding the glucocorticoid receptor (GCR), resulted in dramatic reduction of fibroblasts to induce APOD, NNMT, and CEBPD expression in response to FCM. Non-targeting control (NTC) was used as a control. Two-way ANOVA was performed to calculate statistical significance. (**H**) Volcano plots of genes up and down-regulated by cortisol, FCM, and FCM+ the GCR antagonist mifepristone found by unbiased bulk RNA sequencing. (**I**) Application of module scores to single cell pseudobulk data (reads collapsed over patient). A Kruskal Wallis test was used to calculate significance in panel I. REM=remission. (**J**) Adipocyte depletion over 8wks in knee joints of AdipoQ cre+ iDTR+ mice. (**K**) Whole joint qPCR following adipocyte depletion. Ctl= AdipoQ cre- iDTR-, Depleted= AdipoQ cre+ iDTR+, in both cases the joint taken for qPCR was injected intra-articularly with diphtheria toxin over a time course of 8 weeks. A, C and I statistical comparisons: all groups were compared to basal/healthy.

Since dexamethasone is a synthetic steroid, this finding led us to consider if a natural glucocorticoid might be responsible for the activity of the fat-conditioned media (Fig. 4b). To address this, we similarly carried out the Bligh and Dyer fractionation method on fat-conditioned media, and we observed activity in the organic phase, where we expect cortisol, a nonpolar steroid, to localize (Fig. 4c). We then further separated the organic extracts by the solid phase extraction (SPE) method to elute lipids based on their polarity. Testing each fraction revealed that activity was primarily contained in the acetone fraction (fig. S4a). Next, we developed a one-step HPLC-MS method to directly measure the cortisol concentration by mixing fat-conditioned media with known concentrations of deuterated cortisol (cortisol-d4) in methanol. We found that cortisol is present in synovial and abdominal-derived fat-conditioned media at concentrations ranging from 0.5-4 ng/mL depending on the donor (Fig. 4d-e). Thus, we tested if cortisol alone is sufficient to induce APOD, NNMT and CEBPD at its physiological levels in fat-conditioned media, 2.5ng/mL. Indeed, like the synthetic additive dexamethasone, we found that physiologic concentrations of the endogenous glucocorticoid cortisol drive expression of APOD, NNMT and CEBPD (Fig 4f).

### Fat-conditioned media activity is dependent on glucocorticoid signaling

Given this cortisol activity, we next tested if glucocorticoid signaling or biologically unrelated pathways controlled the fat-conditioned media-induced gene response by using a glucocorticoid receptor antagonist (mifepristone). Mifepristone completely blocked induction of *APOD* and *CEBPD* by fat-conditioned media (fig. S4b). Using a second approach we deleted the glucocorticoid receptor by applying CRISPR guide RNA directed at *NR3C1*, which reduced *NR3C1* gene expression in fibroblasts by ∼83% (fig. S5a). When we applied cortisol to *NR3C1* deleted cells, the cells did not upregulate *APOD*, *NNMT*, and *CEBPD* (fig. S5b). We then applied fat-conditioned media and found that *NR3C1* deletion profoundly reduced the response to fat-conditioned media, evidenced by markedly decreased upregulation of *APOD* (from 47-fold to 7- fold), *NNMT* (from 15-fold to 3-fold), and *CEBPD* (from 30-fold to 11-fold), confirming that most of the activity from fat-conditioned media occurs via glucocorticoid signaling (Fig. 4G).

To further characterize the role of cortisol, we tested the activity of other related steroids and their pathways. Although progesterone can bind to the glucocorticoid receptor, we found that progesterone did not upregulate *APOD*, *NNMT*, or *CEBPD* (fig. S6a). Aldosterone is a strong mineralocorticoid receptor agonist and can also signal through the glucocorticoid receptor *NR3C1*, and it induced expression of *APOD*, *NNMT*, and *CEBPD* at 1uM (fig. S6a). Thus, we measured aldosterone levels in fat-conditioned media by ELISA. Unlike cortisol, which was present at the ng/mL level, aldosterone was present at less than 20 pg/mL (fig. S6b) and adding 10-50 pg/mL lacked activity (fig. S6b), suggesting that aldosterone is not the active molecule signaling through *NR3C1*.

Finally, we tested if fat-conditioned media enhances conversion of inactive cortisone to cortisol via the enzyme 11β-HSD1. We added metyrapone, a competitive 11β-HSD1 inhibitor, to fat-conditioned media, but it did not impact the phenotype (fig. S6c), suggesting that activation of cortisol is not induced by another factor within the fat-conditioned media.

### RNA sequencing confirms glucocorticoid signaling as a regulator of fibroblast gene expression

We performed bulk RNA sequencing on fibroblasts treated with synovial-derived fat-conditioned media, agonists and inhibitors of glucocorticoids and cytokines for 4 or 22hrs. Principal components analysis revealed that the larger variance in gene expression is at 22hrs (fig. S7a).

Focusing on the gene changes at 22hrs, we confirmed that *APOD*, *NNMT*, and *CEBPD* were upregulated by fat-conditioned media while mifepristone downregulated their expression (fig. S7b). Differential gene expression analysis of fibroblasts stimulated with either cortisol, fat-conditioned media, or fat-conditioned media + glucocorticoid receptor (GCR) antagonist for 22hrs was used to create gene lists defining activation scores (Data file S3), which were then applied to the fibroblasts from the single cell RNA sequencing dataset. Healthy fibroblasts had higher fat-conditioned media and cortisol activation scores compared to fibroblasts from OA, RA, and remission subjects. However, mifepristone greatly reduced the fat-conditioned media activation score in healthy fibroblasts, indicating that glucocorticoid signaling is a global driver of healthy fibroblast gene expression (Fig. 4h-i). Separately, OA and RA fibroblasts had higher TGFB signaling compared to healthy and remission fibroblasts (fig. S7c).

### Synovial adipose tissue is a source of glucocorticoids

Cortisol mostly circulates through the body in its inactive form, cortisone. Cortisone is primarily converted to active cortisol in tissues via the enzyme 11β-HSD1. Adipose tissue has a high level of 11β-HSD1 activity compared to many other tissues. Thus, we created a mouse with inducible adipocyte depletion by crossing C57BL/6-*Gt(ROSA)26Sor^tm1(HBEGF)Awai^*/J (iDTR) mice to B6;FVB-Tg(Adipoq-cre)1Evdr/J (ADIPOQ-cre) mice. We injected diphtheria toxin intra-articularly to locally deplete joint adipose tissue over the course of 8 weeks. The quantity of intra-articular adipose tissue was significantly depleted compared to iDTR- AdipoQ-cre- control mice injected with diphtheria toxin, as assessed histologically (150 adipocytes per FOV vs 0) and by gene expression of *AdipoQ* (Fig. 4j).

Concomitant with adipocyte depletion, we observed a reduction in expression of *Hsd11*β*1* (>50%), indicating that joint adipocytes contribute significantly to the generation of active cortisol via expression of this enzyme. Additionally, cortisol responsive genes *Cebpd*, *Plin2*, and *Cidec*, and the preadipocyte marker *Pparg* were all downregulated in adipocyte depleted joint tissues (Fig. 4k). These results are consistent with the conclusion that adipocytes are a key source activating cortisol in the synovium.

### Examination of exogenous human steroid usage on fibroblast phenotype

Thus far, the data collected on healthy, RA, OA, and remission human synovial tissues were all from donors with no reported steroid use. To further our findings, we collected and ran single cell RNA sequencing on 6 additional healthy synovial tissues from patients with reported steroid use prior to autopsy sample collection (table S1, cohort 2). Five of six of these patients received a short course of steroids during their stay at the hospital, shortly before tissue collection. All together, we sequenced a total of 19,378 synovial cells from 16 healthy donors. When steroid exposed donors were analyzed and compared to non-steroid exposed donors, broad cell types, fibroblast cluster identities, top differentially expressed genes relative to OA and RA fibroblasts, and cortisol activation scores were similar (fig. S8a-i), suggesting that healthy fibroblasts are already steroid activated by their adipose rich microenvironment and that donor steroid usage does not further enhance cortisol activation. Additionally, analysis of two separate mouse RNA sequencing datasets comparing healthy versus serum transfer arthritis or hTNFtg arthritic synovial tissue revealed that healthy mouse synovium has a higher cortisol activation score compared to arthritic synovium, independently supporting our findings in humans (fig. S9a-b) [15, 16].

The additional samples also allowed a more fine-grained analysis of healthy synovial immune cell populations. We mapped healthy synovial immune cells onto the RA and OA cell states defined by single cell RNA sequencing from Zhang et al using the Symphony algorithm (data file S4) [8]. Most healthy macrophages mapped to either M0 (*MERTK*+ *SELENOP*+ *LYVE1*+), M1 (*MERTK*+ *SELENOP*+ *LYVE1*-), M2 (*MERTK*+ *S100A8*+), M4 (*SPP1*+), M5 (*C1QA*+), or M7 (*IL1*β+ *FCN1*+) clusters. Healthy T cells mapped primarily to *CD4*+ *IL7R*+*CCR5*+, *CD4*+ *IL7R*+, *CD4*+ *CD161*+ memory, or to *CD8*+ *GZMB*+ TEMRA T cell clusters. These mapped cell states suggest that tissue resident macrophages and T cells are present in healthy synovium. Many inflammatory immune cells defined by Zhang et. al., such as *STAT1*+ *CXCL10*+ macrophages and *CD4*+ Tfh/Tph cells, were not present in healthy synovium [8]. Additionally, less than 10 cells mapped to any given B cell cluster, suggesting this is not a typical cell observed in synovium at homeostasis.

### Healthy synovial fibroblasts share similarity with adipose tissue fibroblasts

Next, we asked if adipocytes control fibroblast phenotype in classical adipose depots by comparing healthy synovial tissue and adipose tissue derived fibroblasts. We performed single cell RNA sequencing on classical visceral (VAT) and subcutaneous (SAT) adipose tissue depots from five obese but metabolically healthy donors (Table S2). We used the Harmony algorithm to integrate these fibroblasts with *PDGFRα*+ non-mesothelial cells from two other adipose single cell RNA sequencing datasets [17, 18]. Altogether, we analyzed 21,856 *PDGFRα*+ cells from 23 donors. These adipose stromal cells revealed four main clusters: one marked by *DPP4*+ expression and high expression of “universal progenitor” markers (*CD55*, *PI16,* and *CD34*) and shown to reside in interstitial regions of murine adipose tissue [19, 20] (Fig. 5a). Two of the other clusters represented committed adipocyte progenitor populations, marked by high expression of preadipocyte transcription factors and high cortisol scoring (Fig. 5b). These preadipocyte populations expressed markers consistent with different levels of preadipocyte commitment: *CEBPD*+ early preadipocytes, and the late preadipocyte marker *FABP4*+ (fig. S10a, Data file S5 a-d) [21].

**Figure 5.**
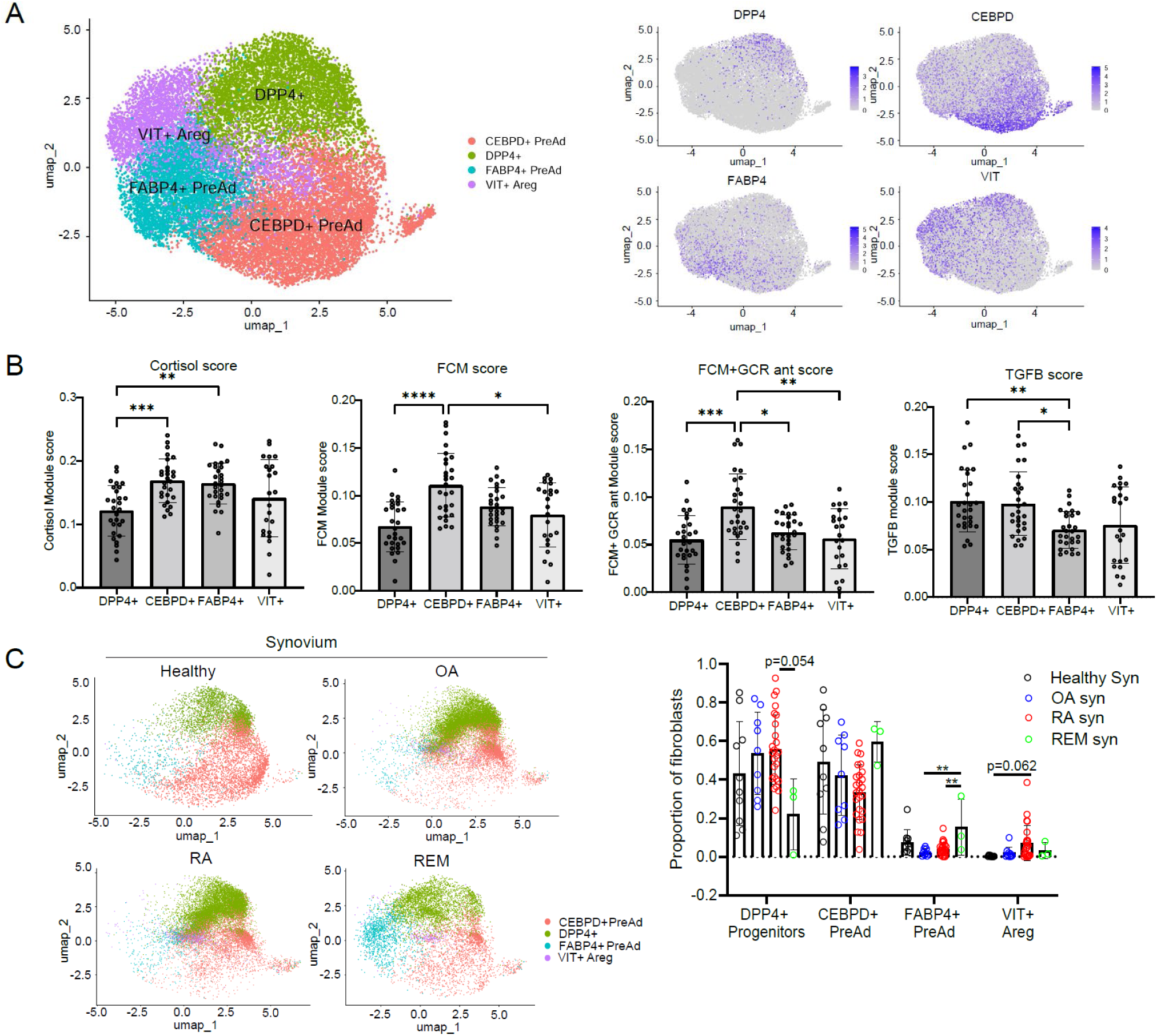
The synovium shares properties with adipose stromal cell populations. (**A**) UMAP of PDGFRα+ stromal cells from human visceral and subcutaneous depots (VAT and SAT). (**B**) Bulk RNA sequencing defined cortisol, FCM, FCM+GCR ant, and TGFB scores applied to adipose clusters. A Kruskal Wallis test was used to calculate significance. (**C**) Symphony UMAP with adipose stromal cells as a reference, quantification of mapping on right. B and C statistical comparisons: all groups were compared to each other.

Fast gene set enrichment analysis (FGSEA) revealed that the two preadipocyte populations were positively enriched for adipogenesis, supporting their identities as preadipocytes (fig. S11a). *CEBPD*+ preadipocytes were enriched for glycolysis, oxidative phosphorylation, and fatty acid synthesis, as well as scoring higher for fat-conditioned media activation, suggesting that they directly encounter fatty acids and are highly metabolically active. *FABP4*+ preadipocytes are quiescent by comparison and are enriched in few pathways (fig. S11a). Interestingly, we found a fourth cluster, *VIT*+ adipogenic regulatory cells (Aregs), which was almost exclusively found in adipose depots from patients with BMIs under 40, and scarce in patients with BMIs from 40-60, suggesting that this population may regulate adipocyte differentiation (fig. S11b). This cluster is negatively enriched for genes involved in adipogenesis and expresses high levels of *VIT* (encoding the ECM protein VITRIN), which has been shown to actively suppress adipogenesis [22].

Since adiposity in the synovium is lost in RA and OA, we asked if the cell states in healthy synovial fibroblasts and adipose tissue are lost in RA and OA. We found that healthy synovial fibroblasts and remission fibroblasts mapped to DPP4+ progenitors and *CEBPD*+ and *FABP4*+ preadipocytes. In contrast, OA and RA fibroblasts mapped more to the universal *DPP4*+ universal progenitor population, with very few cells mapped to preadipocyte states (Fig. 5c).

Additionally, *DPP4*+ universal progenitors displayed the lowest fat-conditioned media and cortisol activation, suggesting that adipocyte proximity and cortisol signaling are important in preadipocyte commitment (Fig. 5b). The diminished mapping of OA and RA fibroblasts to pre- adipocytes, and enhanced mapping to universal progenitors and VIT+ Aregs compared to healthy synovial fibroblasts, suggests that the loss of synovial adiposity changed their fundamental cell state.

### Adipose-derived factors modulate core properties of synovial fibroblast biology ECM remodeling and fibrosis

Bulk RNA sequencing of TGFβ or TNFα +/- fat-conditioned media stimulated fibroblasts identified pathways in which fat-conditioned media counteracts cytokine-induced gene expression changes. Surprisingly, fat-conditioned media was able to ameliorate both TGFβ and TNFα induced extracellular matrix remodeling, including decreasing TNFα*-*induced matrix metalloproteinase expression (*MMP1*, *MMP2*, *MMP3*, *MMP9*, *MMP13*) and TGFβ induced pro-fibrotic collagen expression (fig. S12a-d, Data files S6, 7). In 2D cultured fibroblasts, fat-conditioned media and cortisol decreased *MMP3* upregulation by TNFα (14-fold down to 2.3 and 1.6-fold, respectively Fig.6a, right), supporting the bulk RNA sequencing findings.

In 3D synovial fibroblast organoids, we observed that stimulation with TNFα and IFNγ lead to significant fibroblast tunneling through the extracellular matrix and increased PDPN expression. Strikingly, these cytokine-induced changes are abrogated by the addition of fat-conditioned media or cortisol (Fig. 6a, left, fig. S12e). For TGFβ, we found that stimulation for 21 days increased COL1A1 deposition, particularly at the edges of the organoids. Importantly, the addition of cortisol decreased COL1A1 deposition back to basal levels (from 10 to 5 AU) (Fig. 6b, fig. S12f), confirming the anti-fibrotic role of cortisol signaling in synovial fibroblasts.

**Figure 6.**
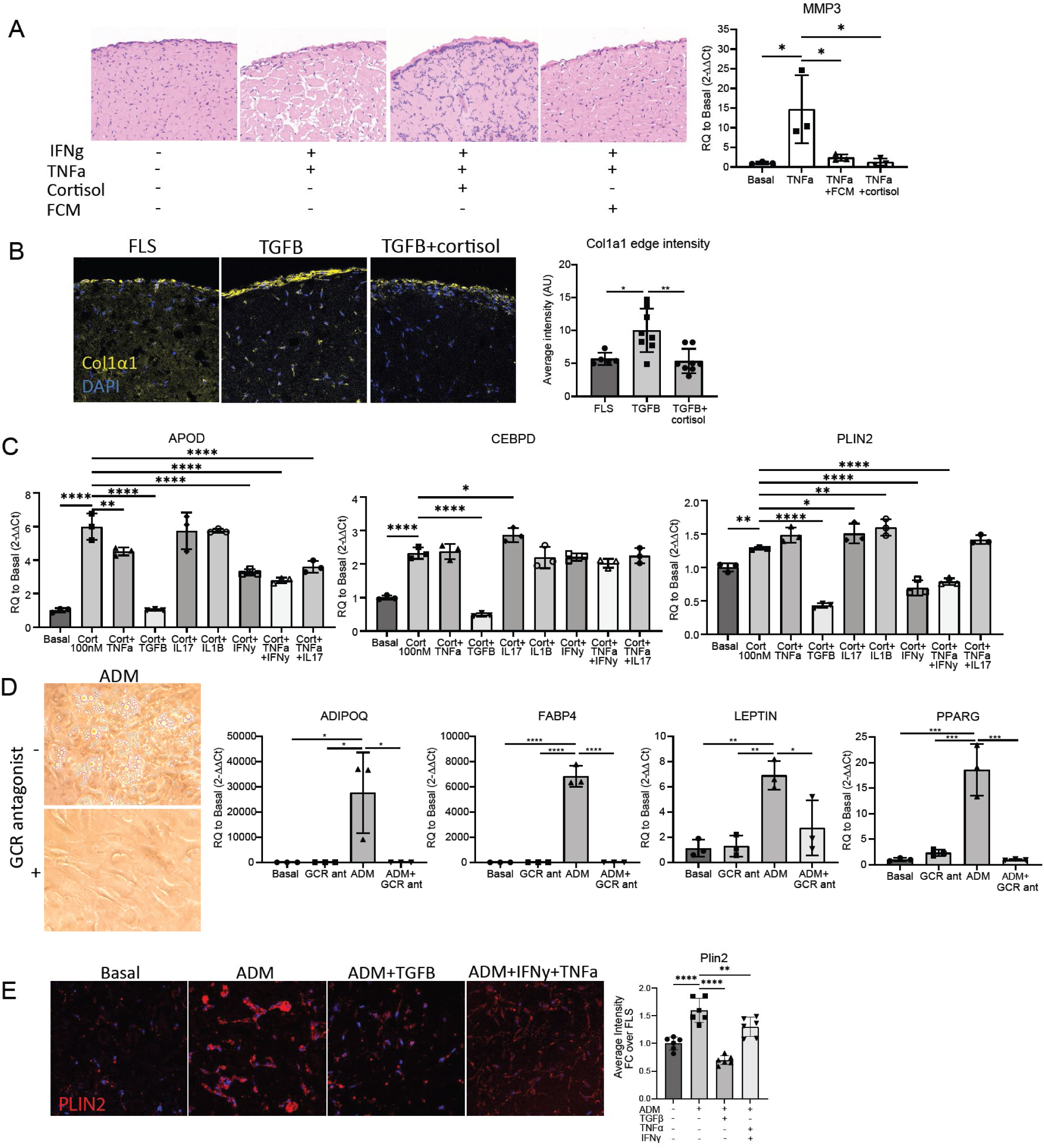
Functional effect of cortisol on fibroblasts. (**A**) FCM and cortisol (1uM) protect against TNFa (2ng/mL) and IFNy (25ng/mL) induced matrix remodeling in long term micromass culture (day 17). Cortisol and FCM were not added until day 7 of culture, TNFa and IFNy were added at day 0. (**B**) Cortisol (1uM) prevents TGFβ (10ng/mL) induced fibrosis as measured by collagen 1a1 immunostaining (Day 21). ImageJ quantification of COL1A1 staining on the right. (**C**) Cortisol (100nM) was applied to cells along with cytokines and assessed for APOD, CEBPD, and PLIN2 expression after 24hrs. Concentrations used were: TNFa (2ng/mL), IFNy (25ng/mL), TGFβ (10ng/mL), IL1β (2pg/mL) and IL17 (10ng/mL). Statistical comparisons: all groups were compared to the cortisol group. (**D**) Ability to enter adipogenic programs. Adipocyte differentiation media (ADM) was applied with or without 10uM mifepristone (GCR antagonist). Day 9 qPCR, day 19 images (cultured until day 28). Two-way ANOVA was performed to calculate statistical significance. (**E**) PLIN2 staining of micromass sections. Micromasses were treated with basal media, ADM, or ADM+ cytokines (TGFβ at 10ng/mL or TNFa at 2ng/mL+ IFNy at 25ng/mL). ImageJ quantification of PLIN2 staining is on the right. Statistical comparisons: all groups were compared to the ADM group. A, B: statistical comparisons: all groups were compared to each other.

### Inflammation

In cultured synovial fibroblasts, TNFα upregulated *IL6* 103-fold and this was significantly downregulated by adding fat-conditioned media (16-fold) or cortisol (14-fold). Similarly, TNFα upregulated *IL1*β 152-fold and this effect was significantly downregulated by adding fat-conditioned media (12.7-fold) or by cortisol (8.8-fold) (fig. S12g).

In our bulk RNA sequencing data, TGFβ stimulation significantly decreased enrichment of lipid and glucocorticoid biosynthetic pathways in fibroblasts. This led us to consider that TGFβ may counterbalance the effects of fat-conditioned media and glucocorticoids. Thus, we tested TGFβ and other cytokines and cytokine combinations for repressing the fibroblast response to cortisol (Fig. 6c). Strikingly, TGFβ significantly downregulated *APOD*, *CEBPD* and *PLIN2* expression to equal to or lower than baseline levels. In addition, IFNγ decreased *APOD* and *PLIN2* expression by 40%. Thus, the homeostatic state induced by fat-conditioned media and glucocorticoids is largely reversed by profibrotic and inflammatory cytokines.

### Blocking glucocorticoid signaling prevents adipogenesis from fibroblasts

We wondered how important glucocorticoid signaling is to adipogenesis and how the presence of cytokines would impact this process. Adipogenic medium was sufficient to induce adipogenesis in cultured fibroblasts, inducing expression of *PPARG*, adipocyte specific genes, and lipid droplet accumulation after 8-30 days (Fig. 6d). However, the addition of mifepristone completely abolished the upregulation of adipocyte specific genes, and fibroblasts exhibited no lipid accumulation. *PPARG* expression was reduced from 18-fold to 1-fold*; ADIPOQ* changed from 27,000-fold to 4-fold; *FABP4* dropped from 6,800-fold to 3.5-fold, and *LEPTIN* changed from 7- fold to 3-fold after mifepristone treatment. Thus, glucocorticoid signaling is important for synovial fibroblast adipogenesis, as has been shown in fibroblasts from other tissues [23].

After confirming that TGFβ and IFNγ signaling pathways are upregulated in OA and RA fibroblasts, and can suppress cortisol signaling, we investigated whether these cytokines could suppress synovial fibroblast adipogenesis. We cultured synovial fibroblasts in 3D organoids with adipocyte differentiation media (ADM) for 21 days with or without TGFβ or IFNγ+ TNFα.

Adipocyte differentiation media induced adipocyte differentiation as measured by strong PLIN2 staining, which defines lipid droplets, as well as by cell morphology; fibroblasts become rounder and less elongated (Fig. 6e). The addition of TGFβ or IFNγ+ TNFα diminished the adipocyte-like morphology as well as total PLIN2 staining, confirming the role of these cytokines in suppressing adipogenesis as well as cortisol signaling.

## DISCUSSION

Fibroblasts are targets of intense study given their importance in tissue pathology. They mediate fibrosis and inflammation in many disorders or inhibit immune responses in the tumor stroma [19, 24]. Many studies on RA have focused on fibroblasts as they mediate both inflammation and degradation of joint tissues in arthritis [25]. Yet, most studies highlight pathological fibroblast states that relate to tissue damage, while few studies have sought to understand key drivers of fibroblast homeostasis. Here, using the synovium as an example, we have identified the importance of adipocytes in driving the homeostatic phenotype of fibroblasts mediated by the activation of the endogenous glucocorticoid cortisol.

We show that adipocytes regulate synovial fibroblast function by expressing *Hsd11*β*1* to generate active cortisol, which acts on resident fibroblasts in the synovium. 11β-HSD1 has been shown to suppress synovitis and joint destruction in the TNF-tg mouse model of arthritis, and cortisol has been used as a first line treatment for managing RA inflammation for decades; but thought to act mainly on infiltrating immune cells to achieve its anti-inflammatory effect [26]. Koenen recently detailed that glucocorticoid signaling in stromal cells, not immune cells, is essential for the therapeutic effect of dexamethasone in serum transfer induced arthritis [27]. Here, we found that cortisol plays a key role in maintaining healthy fibroblast functions and can mitigate the inflammatory, ECM remodeling, and fibrotic effects of TNFα and TGFβ on synovial fibroblasts.

Conversely, at a ten-fold higher ratio of cytokine to cortisol, we found that TGFβ and IFNγ can overcome cortisol signaling in fibroblasts, suggesting that higher levels of inflammation and TGFβ, as observed in RA and OA, overwhelm homeostatic cortisol signaling. Inhibition of cortisol signaling likely further contributes to synovial inflammation and lipodystrophy of the synovium, leading to fibrosis. IFNγ and TGFβ were also found to blunt adipocyte differentiation, which could be a second *in vivo* mechanism of cytokine suppression of homeostatic cortisol signaling by suppressing the major source of 11β-HSD1 in the joint- adipocytes.

We found that a significant number of healthy synovial fibroblasts mapped to committed preadipocyte populations from classical adipose tissue depots. This finding, together with the high cortisol signaling in this population, suggests that the healthy synovium contains preadipocytes that rely on cortisol to maintain their preadipocyte state. This conclusion was also supported by our *in vitro* experiments which showed that blocking the glucocorticoid receptor effectively blocked adipogenesis of synovial derived fibroblasts. Additionally, RA synovial fibroblasts had enhanced mapping to adipose tissue VIT+ Aregs, suggesting that a subset of RA fibroblasts actively suppress adipogenesis.

A limitation of this study is that the healthy human synovial tissue was collected was collected in a different manner from the OA and RA synovial tissues. Whereas OA and RA tissues were collected through needle biopsy or synovectomy, healthy tissue was collected via manual dissection of whole synovium post-mortem. This could contribute to some of the cell proportion and phenotype differences observed. We have endeavored to overcome this limitation by collecting tissues from several different sources with collection times as short as six hours post-mortem. Additionally, we included mouse experiments depleting intra-articular fat in addition to computational analysis of two mouse RNA sequencing datasets to confirm the major findings of our study where the practical limitations of working with human tissues are not present.

In conclusion, this work has identified adipose tissue and cortisol signaling as important contributors to healthy synovial and adipose tissue fibroblast function and identity which are lost in disease. Cortisol plays a critical role in multiple facets of fibroblast biology, including metabolism, inflammation, and ECM homeostasis, which were previously under-appreciated.

We introduce the concept that the loss of adiposity contributes importantly to loss of cortisol signaling and the subsequent development of pathologic fibroblast states in adipose rich tissues such as the synovium.

## MATERIALS AND METHODS

### Single cell RNA-sequencing

16 synovial samples were obtained from human donors with no history of arthritis or autoimmune disease. Cells were isolated via disaggregation (see methods) and loaded onto a single lane (Chromium chip, 10X genomics) followed by encapsulation in a lipid droplet (Single Cell 3’ kit, 10X Genomics) followed by cDNA and library generation according to the manufacturer’s protocol. Cells were stained with cell-hashing antibodies (TotalSeq, BioLegend) before cell capture. cDNA libraries were sequenced to an average of 55,000 reads per cell using Illumina Nextseq 500. scRNA-seq. Reads were processed with Cell Ranger v3.1, which demultiplexed cells from different samples and quantified transcript counts per putative cell. Quantification was performed using the STAR aligner against the GRCh38-3.0.0 transcriptome.

### Bulk RNA-sequencing

Three fibroblast lines consented under institutional review board (IRB) number 2014P002558, titled Profiling of cell subsets in human disease, were plated at 10k cells/well and allowed to rest for 3 days prior to stimulation for 4hrs or 20hrs. Cells were stimulated with FCM, cortisol (100nM), TGFβ (10ng/mL), TNFα (5ng/mL), FCM+ GCR antagonist (mifepristone, 10uM), FCM+ TGFβ, and FCM+ TNFα. Cells were harvested by rinsing with PBS and then applying TCL buffer with 1% beta-mercaptoethanol. Full-length cDNA and sequencing libraries were performed using Illumina Smart-seq2 protocol [28].

Libraries were sequenced on MiSeq from Illumina to generate 35 base paired-end reads. Reads were mapped to the GRCh38.93 transcriptome using kallisto 0.42.4 and transcriptional levels of genes were quantified with the log2(TPM + 1) (transcripts per kilobase million) metric.

### Cell culture

Human synovial fibroblasts were cultured in DMEM supplemented with 5% fetal bovine serum, 1% penn strep, 1% L-glutamine, 1% non-essential amino acids, 2% essential amino acids, and 0.5% beta-mercaptoethanol. Human visceral pre-adipocytes were purchased from Lonza (PT-5005) and cultured according to the manufacturer’s instructions in PGM-2TM Preadipocyte Growth Medium-2 BulletKitTM from Lonza (PT-8002). Once cells were expanded, they were seeded for adipocyte differentiation following the manufacturer’s instructions. After 10 days in adipocyte differentiation media (ADM), media was harvested for downstream applications.

### CRISPR-Cas9 gene deletion

Alt-R® CRISPR-Cas9 sgRNA of the following sequence was directed towards *NR3C1* and was designed in ChopChop (purchased from IDT): mA*mU*mG* rArCrU rArCrG rCrUrC rArArC rArUrG rUrUrG rUrUrU rUrArG rArGrC rUrArG rArArA rUrArG rCrArA rGrUrU rArArA rArUrA rArGrG rCrUrA rGrUrC rCrGrU rUrArU rCrArA rCrUrU rGrArA rArArA rGrUrG rGrCrA rCrCrG rArGrU rCrGrG rUrGrC mU*mU*mU* rU. The primer coordinates are as follows: Left primer: chr5:143300617-143300639, Right primer: chr5:143300397-143300419. Primer sequences: Left primer: CTGGTGTCACTGTTGGAGGTTA, Right primer: GGGCTCACGATGATATAAAAGC. The sequence targets the following nucleotide sequence of the *NR3C1* coding region for deletion: ATGACTACGCTCAACATGTTAGG. sgRNA was applied to cells using the following protocol: 250k cells were resuspended in 17uL of P2 solution. sgRNA was resuspended in IFTE buffer to 80uM concentration and mixed 1:1 with cas9 and incubated for 15 minutes to allow RNP complex formation. The cell suspension and sgRNA+cas9 mixture were then combined and then underwent nucleofection program EN 150. After, pre-warmed cell media was added to each well and let sit for 15 minutes at room temperature before adding to a cell culture flask.

Fibroblasts were plated and allowed to rest for three weeks prior to validation of gene deletion and use for experiments.

### In vitro stimulation experiments

Fibroblasts were plated at 40k cells per well in a 12 well plate and allowed to rest for four days prior to stimulation for 24 hours. The following chemicals were purchased from Sigma Aldrich: cortisol was purchased as Hydrocortisone (H0888-1G), Aldosterone (A9477-5MG), Progesterone (P8783-1G), the glucocorticoid receptor antagonist Mifepristone (M8046-100MG) was used at 10μM, the mineralocorticoid receptor antagonist spironolactone (S3378-1G) was used at 1μM, the 11β-Hydroxysteroid dehydrogenase type 1 competitive inhibitor Metyrapone (M2696-10MG) was used at 100μM. The following chemicals were purchased from Peprotech: TGFβ (100-21-2UG) used at 10ng/mL, TNFα (300-01A-10UG) used at 5ng/mL, and IFNγ (300-02-20UG) used at 25ng/mL. Arachidonic acid (AA, Cat# U-71- A) was purchased from Nu-Chek Prep. Triacylglycerol (TAG, Cat# T7140) and monoacylglycerol (MAG, Cat# M2015) were purchased from Sigma. β-Glucosylceramide (β- GlcCer, Cat# 860539) and ceramide (Cer, Cat# 860518) were purchased from Avanti Polar Lipids. Sodium palmitate (Cat# P9767-5G) and sodium oleate (Cat# O7501-250MG) were purchased from Sigma, dissolved in methanol to 200µM, and then diluted to specified concentrations in culture media.

### Synovial tissue collection

Healthy synovial samples collected through three different sources: seven are from the national Disease Research Interchange (NDRI), four are from Rush University, and five are from BWH autopsy department. Patient tissues were excluded for collection if they had a diagnosis of lupus, type 1 diabetes, psoriasis, rheumatoid arthritis, psoriatic arthritis, spondyloarthritis, Crohn’s disease, Sjogren syndrome, or osteoarthritis.

Collection and sequencing of these tissues was performed under institutional review board (IRB) number 2002P000127 titled Pathways of Antigen Presentation by CDI. Additionally, tissue was obtained from Rush University through the materials transfer agreement 2020A004824 titled Molecular profiling of synovium. We thank the Brigham Autopsy department for collecting post-mortem synovial tissues for our lab. We thank Rush University for collecting post-mortem synovial tissues for our lab. Samples obtained from Rush also went through Collin’s Grading of cartilage as an additional measure determine joint health. Upon receipt of synovial tissues, synovium was dissected away from adipose tissue as much as possible. Synovium was saved in Crystor CS10 freezing media (Sigma) by incubating on ice for 10 minutes prior to freezing. Synovial adipose tissue was minced into approximately 5mm pieces and used to generate fat-conditioned media.

### Fat-conditioned media

Fat-conditioned media was generated from adipose tissue sourced from either healthy joints (syn fat donor) or abdominal adipose tissue. Briefly, adipose tissue was minced into approximately 5mm pieces and cultured in RPMI for 24hrs before harvesting the media. The conditioned media was then passed through a 70µm filter and then spun down at 1500 rpm for five minutes to remove cells.

### Lipid extraction and Fractionation

Conditioned media was fractionated into aqueous, organic, and an interphase layer using the Bligh and Dyer method. Briefly, a 1:2 solution of chloroform:methanol was added to conditioned media at a ratio of 3.75:1, respectively. The sample was vortexed and then 1.25mL of chloroform and 1.25 mL of water were added. The sample was vortexed again and then centrifuged at 2200rpm for 15 minutes. The aqueous and organic phases were separated and kept for further analyses. Separation of the organic fraction using solid phase extraction columns (Supelclean™ LC-Si SPE Tube, Sigma) based on elution with solvents with increasing polarity: chloroform, acetone, methanol, and water fraction were used serially.

### Direct cortisol quantification by HPLC-MS

To directly quantify cortisol, conditioned media was mixed with varying concentrations of cortisol-d4 (methanol solution) in a 1:1 (v/v) ratio. The mixture was then subjected to the reverse phase HPLC-MS analysis using the Agilent Poroshell column and the Q-Tof instrument as described above. The mobile phase A was methanol/water 20/80 (v/v), supplemented with 2 mM ammonium formate; the mobile phase B was methanol/water 80/20 (v/v), supplemented with 2 mM ammonium formate. 10 µl of mixed sample were injected and the flow rate was 0.15 ml/min with the binary gradient: 0–2 min, 50% A; 2–10 min, from 50% A to 100% B; 10–15 min, 100% B; 15–17 min, from 100% B to 50% A; and 17–20 min, 50% A. The unknown cortisol concentrations in the condition media were determined by the chromatogram area of the non-deuterated natural molecule compared to the known concentration of cortisol-d4 internal standard in 4-5 experiments.

### Gene expression analysis

RNA was extracted from fibroblasts using TRIzol reagent (Life Technologies) as per the manufacturer’s instructions. The aqueous phase was mixed 1:1 with 70% ethanol and applied to a minikit spin column and washed with RW1 and RPE buffer (Qiagen). cDNA was synthesized using the QuantiTect Rev. Transcription Kit following the manufacturer’s protocol. Quantitative real-time polymerase chain reaction (qRT-PCR) was carried out using SYBR Green primers and an Agilent AriaMx Real-Time PCR system. Relative gene expression was calculated by the ΔΔCt method. The ΔCt was calculated using the reference genes β2-microglobulin (*B2m*) and β-actin (*Actb*). ΔΔCt was calculated relative to the basal control group. Human and mouse primer sequences are listed in supplemental materials.

### Mouse experiments

All animal studies were performed with approval by the institutional animal care and use committee (IACUC) of the Brigham and Women’s Hospital. C57BL/6- *Gt(ROSA)26Sor^tm1(HBEGF)Awai^*/J (iDTR) and B6;FVB-Tg(Adipoq-cre)1Evdr/J (ADIPOQ-cre) mice were purchased from Jackson labs. Diphtheria toxin was injected intra-articularly at 50ng/injection in a 10uL volume into 1 knee joint/mouse at day 0, 4, 8, 14, 21, 28, 35, 42 and 49 prior to sacrifice at day 56. Investigators were single blinded while performing injections. No statistical method was used to pre-determine sample size.

### RNA-seq quality control and pre-processing

After filtering out low-quality cells (<200 and >2,800 unique genes, >25% mitochondrial reads), 19,378 cells from primary tissue were further analyzed. With these high-quality cells, we used the Seurat package (version 4.0.1) in R (version 4.1.1) to perform normalization, find variable features, perform principal components analysis, clustering and dimensional reduction using TSNE [29]. We corrected for donor-specific effects using Harmony with default parameters [30].

### Building and mapping to references

We used the buildReferenceFromSeuratObj() function from the Symphony package to build integrated reference atlases for the global and cell-type specific atlases from the Harmony objects [31]. To find concordance between cell types defined by Zhang et al and this study, we used the Symphony mapQuery() function to map the 19,378 scRNA-seq healthy synovial cells onto cells defined by Zhang et al [8]. We also mapped fibroblasts from Zhang et al onto a reference atlas defined by healthy synovial fibroblasts. We predicted reference cell types and states for the query cells using the knnPredict() function with k=5.

### Covarying neighborhood analysis (CNA)

We used the rcna R package from github to calculate cell neighborhoods which were negatively associated with healthy synovial cells. The association. Seurat function was used to perform association testing and compute neighborhood-level FDRs (false discovery rate).

### Bulk RNA sequencing calculation of rescued pathways

To identify pathways which were returned to basal levels by FCM, we ranked genes according to the following calculation for gene set enrichment analysis, with TNFα and FCM+TNFα as an example: “Statistic =MIN(ABS(Fold change difference*),ABS(TNFα fold change over basal)) x SIGN(ABS(TNFα fold change over basal)-ABS(TNFα +FCM fold change over basal)).” * Fold change difference is the fold change of TNFα over basal minus the fold change of TNFα+FCM over basal.

### Ingenuity Pathway Analysis

A set of differentially regulated genes among all sublining fibroblasts between healthy and RA patients, and between remission RA and RA patients, was generated in R. This gene list was then entered into IPA as a core analysis with an expression log ratio cutoff of 1.5 and false discovery rate cut off of 0.05. This left us with over 200 genes which were run in each data set.

### Subcutaneous and omental adipose tissue collection

For single cell RNA sequencing of adipose depots in figure 5, all patients undergoing voluntary bariatric surgery provided written informed consent. This study was carried out in accordance with the declaration of Helsinki and was reviewed and approved by the ethics committees of St. Vincent’s University Hospital and Trinity College Dublin, and Brigham and Women’s Hospital. This study was approved under IRB 2019P001128 entitled ’METABOLIC AND IMMUNOLOGICAL LINKS BETWEEN OBESITY, SYSTEMIC INFLAMMATION, TYPE 2 DIABETES MELLITUS AND NON-ALCOHOLIC FATTY LIVER DISEASE.

### Abdominal adipose tissue collection

The Human Skin Disease Resource Center of Brigham and Women’s Hospital and Harvard Medical School collected anonymous, discarded adult skin plus fat samples removed as part of cosmetic surgery procedures and provided them to our lab. The Human Skin Disease Resource Center is supported in part by NIAMS Resource-based Center Grant # 1P30AR069625. This tissue was collected under IRB number 2020P003757. Abdominal adipose tissue was used to generate fat conditioned media.

### Lipid sources

Lipid standards were purchased from the commercial sources. Arachidonic acid (AA, Cat# U-71-A) was from Nu-Chek Prep. Hydrocortisone-d4 (cortisol-d4, Cat# 26500) was from Cayman Chemical. Triacylglycerol (TAG, Cat# T7140) and monoacylglycerol (MAG, Cat# M2015) were from Sigma. β-Glucosylceramide (β-GlcCer, Cat# 860539) and ceramide (Cer, Cat# 860518) were from Avanti Polar Lipids.

### HPLC-MS analysis of lipids from solid phase extraction fractions

Lipid fractions of chloroform, acetone, and methanol from SPE columns (triplicate) were collected in the pre-weighed vials and dried under nitrogen. The lipid quantities were then measured and redissolved in chloroform/methanol (1:1, v/v) and normalized to 0.5 mg/ml. 50 µl of normalized samples were dried under nitrogen, redissolved in 50 µl of the reverse phase starting mobile phase A with HPLC-MS run on an Agilent Poroshell 120 A, EC-C18, 3 x 50 mm, 1.9 µm reversed phase column coupled with a 3 x 5 mm, 2.7 µm guard column and analyzed by an Agilent 6546 Accurate-Mass Q-ToF/1260 series HPLC instrument. The flow rate and the binary solvent systems were used as published methods [32]. Data were analyzed by Agilent Mass Hunter. For lipidomics, data were analyzed using automated peak-picking algorithms by R package XCMS [33] and in house designed software methods [13].

#### ELISA

ELISA detection of cortisol was performed using the Cortisol ELISA Assay Kit from Eagle biosciences (COR31-K01) according to manufacturer instructions. ELISA detection of aldosterone was performed using the Aldosterone ELISA Assay Kit from Eagle biosciences (ALD31-K01) according to manufacturer instructions.

### Gene expression analysis

The human specific primers used were the following (listed 5’ to 3’): *B2M forward:* GAGGCTATCCAGCGTACTCCA, *B2M reverse:* CGGCAGGCATACTCATCTTTT, *ACTB forward:* GCTCCTCCTGAGCGCAAGTAC, *ACTB*

*reverse:* GGACTCGTCATACTCCTGCTTGC, *APOD forward:* CCACCCCAGTTAACCTCACA, *APOD reverse:* GTGCCGATGGCATAAACC, *NNMT forward:* TTGAGGTGATCTCGCAAAGTTATT, *NNMT reverse:* CTCGCCACCAGGGAGAAA, *CEBPD forward:* GGTGCCCGCTGCAGTTT, *CEBPD reverse:* CTCGCAGTTTAGTGGTGGTAAGTC, *MMP3 forward:* CTCCAACCGTGAGGAAAATC, *MMP3 reverse:* CATGGAATTTCTCTTCTCATCAAA, *IL6 forward*: AGACAGCCACTCACCTCTTCAG, *IL6 reverse*: TTCTGCCAGTGCCTCTTTGCTG, *IL1*β *forward*: GGACAAGCTGAGGAAGATGC, *IL1*β *reverse*: TCGTTATCCCATGTGTCGAA. *NR3C1α/*β forward: ACT TAC ACC TGG ATG ACC AAA T, *NR3C1α* reverse: TTC AAT ACT CAT GGT CTT ATC C, *NR3C1*β *reverse:* TCC TAT AGT TGT CGA TGA GCA T; *FABP4* forward: GCA AAG CCC ACT CCT ACA GTT, *FABP4* reverse: CTC TCT GTG CCT TTT TCC TCC T; *PPARG* forward: AGG CGA GGG CGA TCT TGA CAG, *PPARG* reverse: GAT GCG GAT GGC CAC CTC TTT; *ADIPOQ* forward: CAA CAT GCC CAT TCG CTT T, *ADIPOQ* reverse: GGA GGC CTG GTC CAC ATT AT; *LEPTIN* forward: CGG TAA GGA GAG TAT GCG GG, *LEPTIN* reverse: CAG TAG GTG CCT GGC ATT CA.

To assess adipocyte depletion in mouse joints, whole knee joints were taken, flash frozen in liquid nitrogen, crushed with a mortar and pestle and RNA isolated using TRIzol. **Mouse specific primers**: *AdipoQ* forward: AAGGAGATGCAGGTCTTCTTGGT, *AdipoQ* reverse: CACTGAACGCTGAGCGATACAT; *Hsd1* β*1* forward: CGA CAT CCA CTC TGT GCG AA, *Hsd11*β*1* reverse: TGC TGC CAT TGC TCT GCT; *Pparg* forward: CCA CAG TTG ATT TCT CCA GCA TTT C, *Pparg* reverse: CAG GTT CTA CTT TGA TCG CAC TTT G; *Cebpd* forward: TGCGAGCGACAGGAAGCT, *Cebpd* reverse: GCAATGGTAATAAGACGTAGAAAATGC; *Plin2* forward: CAG CCA ACG TCC GAG ATT G, *Plin2* reverse: CAC ATC CTT CGC CCC AGT T; *Cidec* forward: GGC GTA GTC CAT CCT TGT CA, *Cidec* reverse: GTC AGA TGA GAG ACT GGG GC.

### Tissue disaggregation

Tissue was disaggregated as previously described [34]. Briefly, cryo- preserved synovial tissue was thawed and rinsed in RPMI with 5% FBS. Samples were transferred to a gentleMACS tube containing digest media (0.2mg/mL Liberase TL, 0.1mg/mL DNase1 in RPMI). Tissue was macerated with gentleMACS followed by shaking at 37 degrees for 30 minutes. The tissue was then filtered through a 70µm cell strainer and spun down.

### Flow cytometry

Disaggregated tissues were stained with the surface markers CD45 (2D1), CD31 (WM59), CD90 (5E10), podoplanin (NC-08), CD146 (P1H12), CD14 (63D3), and CD3 on ice for 40 minutes. Fixable viability dye (UV455, eBioscience) was applied and stained on ice for 20 minutes. Cells were fixed in BD cytofix fixation buffer for 20 minutes. After fixation, HCS LipidTOX (ThermoFisher, H34475) was applied at a 1:200 dilution in PBS and stained at room temp for 30 minutes. Cells were filtered through 70-micron mesh filter and run on an LSRII (BD Biosciences).

### AMP CyTOF data

The CD45, CD3, and CD14 flow cytometry quantifications for OA and RA samples in figure 1d were calculated based on Cytof data from AMP phase 1 [2]. Leukocyte rich RA biopsies (n=8) were used for the RA group and n=12 OA samples were used.

### Micromass experiments

Fibroblasts were resuspended in Matrigel (BD Biosciences, 3354234) on ice at 200,000 cells per 35uL of Matrigel. Cold pipette tips were used to pipette 35uL of Matrigel into each well of a 12 well tissue culture plate pre-coated with poly-HEMA (Aldrich Chem Co.). The plate was placed in a 37°C incubator for 1 hour to allow the Matrigel/cell suspension to gel. Then pre-warmed media to the well. IFNγ (25ng/mL) and TNFα (2ng/mL) were added immediately and at every media change. Fat conditioned media was added at day 7 and then replenished at each subsequent media change. At day 17, micromasses were harvested by fixing in 4% paraformaldehyde and then embedded in paraffin. Sections were stained with H&E or immunofluorescently labeled with PDPN (eBioscience, 16-9381-81), Prg4 (Millipore Sigma, MABT401), Plin2 (Thermo Fisher, 15294-1-AP).

### Immunofluorescence

5μm thick sections of paraffin embedded tissue were taken and deparaffinized prior to staining with histoclear. After rehydration, antigen retrieval was performed using Tris buffer pH 9 heated to near-boiling and placed in an 85 degree Celsius oven for 30 minutes. Slides were blocked in 5% normal horse serum for 30 minutes before application of primary antibodies. The following primary antibodies were applied at 1:100 dilution for 1.5 hours at room temperature: PDPN (eBioscience, 16-9381-81), Prg4 (Millipore Sigma, MABT401), Plin2 (Thermo Fisher, 15294-1-AP), CD45 (Thermo Fisher, A304-376A-T), or COL1A1 (R&D Systems, AF6220). Secondary antibodies were applied at room temperature for 1.5 hours (donkey anti–rat AF488, 712-545-153; donkey anti–rabbit AF647, 711-605-152; donkey anti-mouse Cy3, 715-165-150; all from Jackson Immunoresearch). Slides were then treated with Sudan Black B for 15 minutes to diminish autofluorescence. This was followed by DAPI staining for 5 minutes and mounting with FluorSave™ Reagent from Sigma.

### Quantification

COL1A1 and PLIN2 staining was quantified in ImageJ by selecting the whole micromass area of each image and measuring the mean intensity.

### Oil red O staining

OCT embedded synovium was cryosectioned and sections were air dried for 5 minutes at room temperature. Working solution oil red O was prepared by making a stock solution of 300mg oil red O in 100mL isopropanol, mixing 30mL of the stock with 20mL of distilled water, mixing, and filtering prior to use. Staining procedure: Slides were fixed in 4% PFA for 10 minutes, dipped in 60% isopropanol once quickly, stained in oil red O working solution for 15 minutes, dipped in 60% isopropanol once quickly, dipped in DI water once quickly, counterstained with Mayer’s Hematoxylin for 3 minutes, dipped in DI water 10 times, and then mounted with DAKKO mounting media.

### Adipocyte quantification

Knee joints were fixed in 4% paraformaldehyde, decalcified for 2 weeks in 10% EDTA, and then dehydrated and embedded in paraffin. Sections (7 μm thick) were taken and stained with Safranin-O and Fast Green (Applied Biosciences) according to the manufacturer’s instructions. Adipocyte quantification was performed on 2 slides from the medial plateau of the tibia of each mouse. Images were opened in ImageJ and Adiposoft was used to quantify adipocyte number.

### Statistical analysis

For comparison of two groups, a two-tailed student’s T test was used to calculate significance. For three or more comparisons, an ordinary one-way ANOVA followed by Dunnett’s or Tukey’s multiple comparisons post-hoc test were used, depending on the comparisons tested, to calculate significance unless otherwise noted in figure legend. p < 0.05 was considered significant. *=p < 0.05, **= p< 0.01, ***= p< 0.001, ****=p<0.0001. Error bars represent standard deviation in every figure. Statistical testing for flow cytometry and functional assays was performed using GraphPad Prism. F values, t values, p-values, n, and df for each figure are in Data File S8. All measurements were taken from distinct samples. Data is assumed to be normally distributed.

### Data anonymization

For five donors (Table S1, cohort 2 donors 12-16), genetic variations were removed from raw fastq files using BAMboozle (v0.5.0) [35]. Sanitized fastq files will be deposited to the database of Genotypes and Phenotypes (dbGaP) within 6 months of publication.

### Data availability

Source data are provided with this paper.

## List of Supplementary Materials

Materials and Methods Fig. S1 to S12

Tables S1 to S2

Data file S1 to S8 (Excel files)

## Acknowledgments

We thank the Human Skin Disease Resource Center of Brigham and Women’s Hospital which collected anonymous, discarded adult skin plus fat samples and provided them to our lab. We thank the DF/HCC Specialized Histopathology Services Core in MGH for FFPE sections of OA and RA synovium used for immunofluorescence staining. We thank the Gift of Hope Organ & Tissue Donor Network, IL, as well as the donors’ families, for the precious tissue samples.

## Funding

National Institutes of Arthritis and Musculoskeletal and Skin Diseases, T32AR007530- 37, Immunologic Mechanisms and Rheumatic Disease (MBB)

National Institutes of Arthritis and Musculoskeletal and Skin Diseases, P30 AR070253, Microgrant (HJF)

## Author contributions

Conceptualization: HJF, MBB

Methodology: HJF, TC, GFMW, IK, KK, SG

Funding acquisition: HJF, MBB

Sample acquisition: DPS, WT, LL, RP, SC, KW, AHJ, SR, FZ

Supervision: MBB, DBM

Writing – original draft: HJF, MBB

Writing – review & editing: HJF, MBB, TC, DBM, AHJ, SR

