## Supplemental tables and figures for "Adipocytes regulate fibroblast function, and their loss contributes to fibroblast dysfunction in inflammatory diseases"

**Table S1.** Healthy Synovial Donor Information.

| **Cohort 1:** | | | | | | |
|---|---|---|---|---|---|---|
| **Donor** | **Sex** | **Age** | **BMI** | **Weight category** | **Race** | **Collins Grade of cartilage** |
| 1 | Male | 51 | 25 | Overweight | Caucasian | N/A |
| 2 | Male | 28 | 22.8 | Lean | Caucasian | N/A |
| 3 | Male | 29 | 25.85 | Overweight | Caucasian | N/A |
| 4 | Female | 41 | 27 | Overweight | Caucasian | N/A |
| 5 | Male | 54 | 24.1 | Lean | Other | N/A |
| 6 | Male | 59 | 35.82 | Obese | Caucasian | N/A |
| 7 | Female | 56 | 33.3 | Obese | Caucasian | 1 |
| 8 | Female | 58 | 21 | Lean | Caucasian | 2 |
| 9 | Female | 50 | 30.3 | Obese | Hispanic | 0 |
| 10 | Female | 65 | 24.5 | Lean | Caucasian | 1 |
| **Cohort 2:** | | | | | | |
| 11 | Female | 60 | 42.5 | Obese | | N/A |
| 12 | Female | 69 | 24.4 | Lean | White/non-hispanic | N/A |
| 13 | Male | 72 | 21.7 | Lean | White/non-hispanic | N/A |
| 14 | Male | 67 | 33.8 | Obese | White/Non-Hispanic | N/A |
| 15 | Female | 60 | 29.5 | Overweight | White/Non-Hispanic | N/A |
| 16 | Male | 41 | 19 | Lean | Black/Hispanic | N/A |

**Continued:**

| **Cohort 1:** | | | |
|---|---|---|---|
| **Donor** | **COD** | **Time to tissue collection** | **Glucocorticoid usage** |
| 1 | intracerebral hemorrage | 15.8hrs | Home: None reported, Hospital: None reported |
| 2 | Suicide (hanging) | 14.8hrs | Home: None reported, Hospital: None reported |
| 3 | drug intoxication (cocaine, alcohol, cannabinoids) | 3hrs | Home: None reported, Hospital: None reported |
| 4 | cardiac | 7.7hrs | Home: None reported, Hospital: None reported |
| 5 | cardiac | 12.5hrs | Home: None reported, Hospital: None reported |
| 6 | Acute COPD/CPA | N/A | Home: None reported, Hospital: None reported |
| 7 | Anoxia | N/A | Home: None reported, Hospital: None reported |
| 8 | Cardiac arrest | N/A | Home: None reported, Hospital: None reported |

| | | | |
|---|---|---|---|
| 9 | Suicide (burn toxic ingestion) | N/A | Home: None reported, Hospital: None reported |
| 10 | Myocardial infarction | N/A | Home: None reported, Hospital: None reported |
| **Cohort 2:** | | | |
| 11 | cardiopulmonary arrest | 12 hrs | Home: prednisone, hospital: not reported |
| 12 | Acute hypoxemic respiratory failure with right heart failure | 17hrs | Home: No steroids, Hospital: Prednisone PO 15 mg Day-7-0 |
| 13 | Respiratory failure due to hypercapnic respiratory failure, GVHD, CMML | 21hrs | Home: budesonide 3mg q 24h (inhaled), prednisolone acetate 1% opthalmic suspension for left eye, prednisone 15 mg daily. Hospital: Prednisone PO 15 mg D-3-0 |
| 14 | Enterobacter bacteremia | 24hrs | Home: none, Hospital: Hydrocortisone injection 100 mg q8h D-1 and D0 |
| 15 | Shock and respiratory failure | N/A | Home: prednisone 80 mg daily, Hospital: Hydrocortisone injection 50 mg q6h D0 |
| 16 | Intraventricular hemorrhage | 6hrs | Home: none, Hospital: Methylprednisolone 2g IV D-1; 1g IV D0 |

**Table S2.** Adipose donor Information. ****Note that MUO = metabolically unhealthy. When less than 3 MUO events are logged the patient is considered metabolically healthy obese "MHO" based on bloodwork and clinical manifestations (red highlight).*

| Patient | | Sex | Age | Weight (kg) | BMI | Group | MUO* Events ≥3 = MUO | Blood Glucose (fasted) | **Fasted Triglycerides** | HDL Chol. |
|---|---|---|---|---|---|---|---|---|---|---|
| P11 | Healthy | M | 40 | 222 | 66 | MHO | 0 | 5.9 | 1.2 | 1.01 |
| P10 | Healthy | M | 50 | 132 | 45 | MHO | 1 | 4.8 | 1.5 | 1.03 |
| P49 | Healthy | F | 57 | 147 | 56 | MHO | 0 | 4.8 | 1.5 | 1.66 |
| P08 | Healthy | F | 32 | 189 | 65.8 | MHO | 0 | 4.7 | 1.3 | 1.33 |
| P50 | Healthy | F | 37 | 134 | 50 | MHO | 1 | 5 | 0.9 | 1.14 |

**Continued:**

| Patient | **Co-morbidities: (Red Highlight = MUO Event)** | |
|---|---|---|
| P11 | depression | pulmonary embolism |
| P10 | hypertension | sleep apnea |
| P49 | PCOS | |

| | | | | |
|---|---|---|---|---|
| P08 | sleep apnea | GORD | | |
| P50 | OSA | IHD | asthma | paraoxismal Afib |

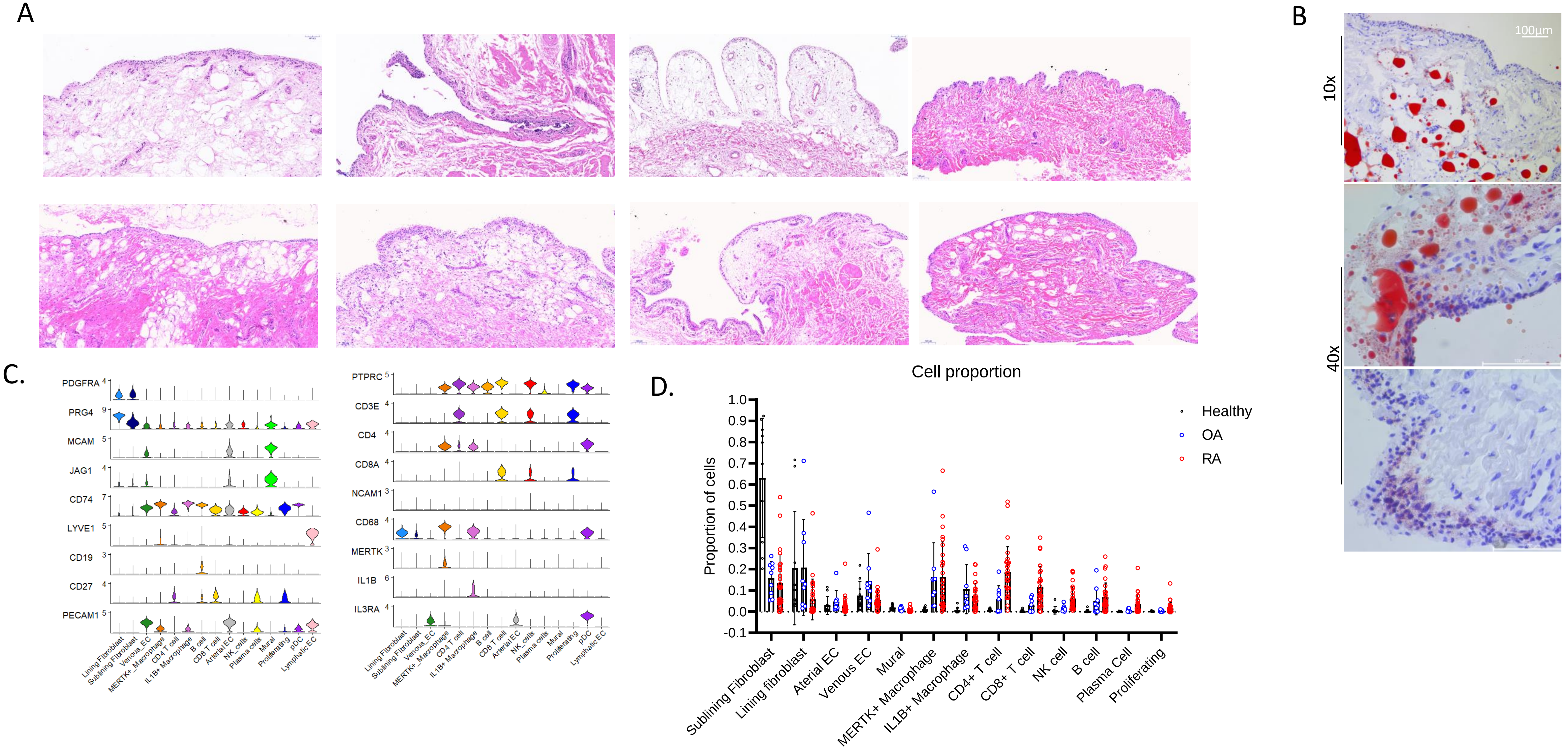

Supplementary Figure 1. A. H&E staining on paraffin embedded healthy synovial tissue sections. B. Oil red O staining of OCT embedded healthy synovium. C. Major marker expression defining each cluster. D. Cell cluster proportions among healthy, OA, and naïve RA synovial cells and UMAP separated by disease state.

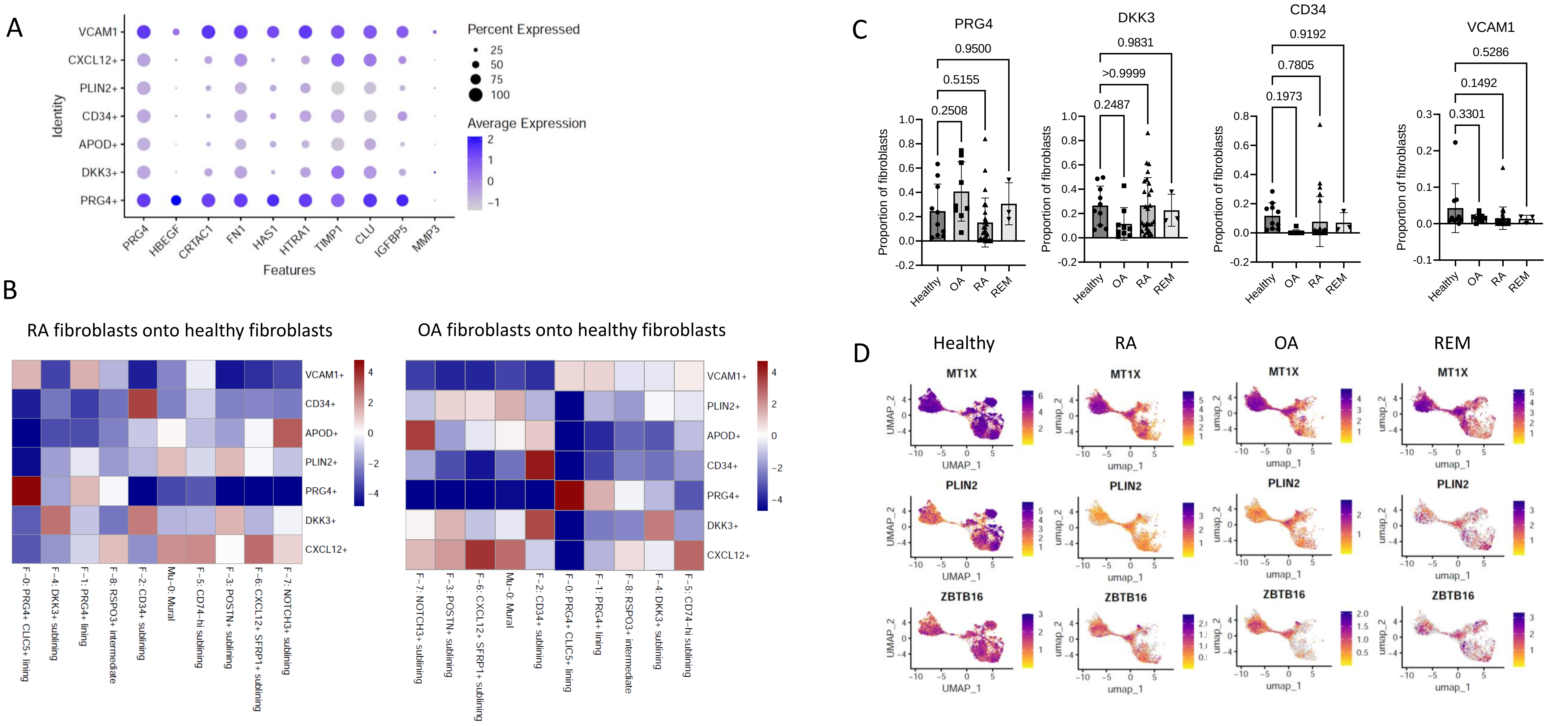

Supplementary Figure 2 S. Fig. 2. A. PRG4+ and VCAM1+ fibroblasts express high levels of lining fibroblast markers. B. Heatmaps show odds ratios for the fibroblast clusters, with rows corresponding to fibroblast clusters from healthy synovial fibroblasts and columns corresponding to fibroblast clusters from Zhang, *et al*, 2022. Blue-red color scale indicates the log(OR) for a given pair of states (OR is the ratio of odds of mapping a cluster in Zhang, *et al*, 2022 to a given healthy fibroblast cluster compared to odds of mapping other fibroblasts in Zhang, *et al*, 2022 onto the same cluster of this study), with higher values indicating greater correspondence. C. Quantification of fibroblast proportions mapping to each cluster. Df=44. PRG4: F=3.383, DKK3: F=1.410, CD34: F=1.099, VCAM1: F=1.351. D. Healthy fibroblasts globally upregulate genes which are involved in metabolism, including *PLIN2*, *MT1X*, and *ZBTB16*.

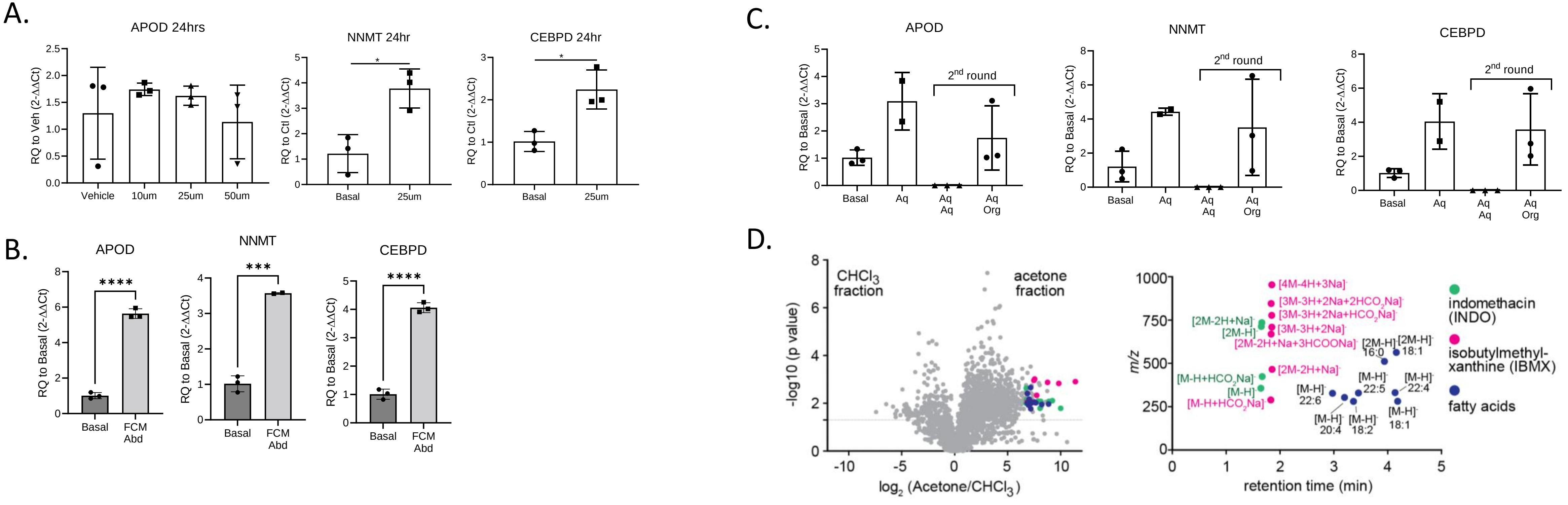

Supplementary Figure 3 A. Fibroblasts cultured with 10um oleate+10um palmitate for 24hrs do not induce APOD expression. B. Abdominally derived FCM induces *APOD*, *NNMT*, and *CEBPD* expression. C. Bligh and dyer separation of adipocyte conditioned media (ACM) shows that the active molecule is in organic and aqueous fractions of ACM (Shown in Fig. 3c). Due to incomplete separation; a second round of bligh and dyer on the aqueous fraction was performed and results in all activity going to the organic fraction ("aq org"). D. Three independently separated fractions of chloroform, acetone and methanol were normalized to the lipid weight and subject to HPLC-QToF-MS negative mode analysis. The non-stimulatory chloroform fractions were compared to the stimulatory acetone fractions by lipidomics analysis. The ions with highest fold change and intensity in the acetone fraction were selected and plot against the retention time. The fatty acid class (blue dots of seven different fatty acids), indomethacin (green dots of four alternate and multimer ion adducts), and isobutylmethylxanthine (pink dots of four alternate and multimer ion adducts) were identified.

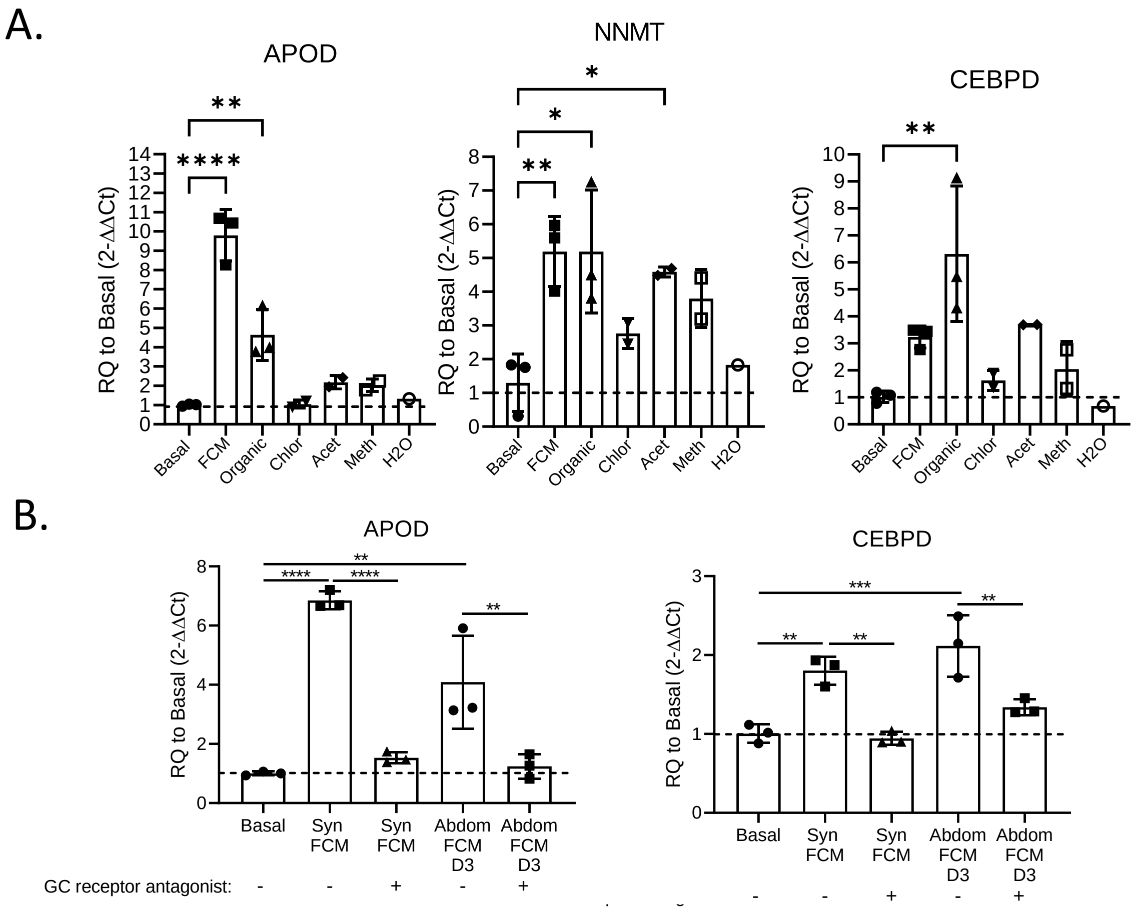

Supplementary Figure 4. A. FCM was separated using the Bligh and Dyer method into aqueous and organic phases. Then, the organic phase was taken for solid phase separation and eluted based on polarity using chloroform, acetone, methanol, and water. Testing activity of each fraction; activity was primarily in the acetone and methanol fractions of the organic phase. . B. APOD and CEBPD upregulation are suppressed by the GCR antagonist mifepristone. D3 stands for donor 3.

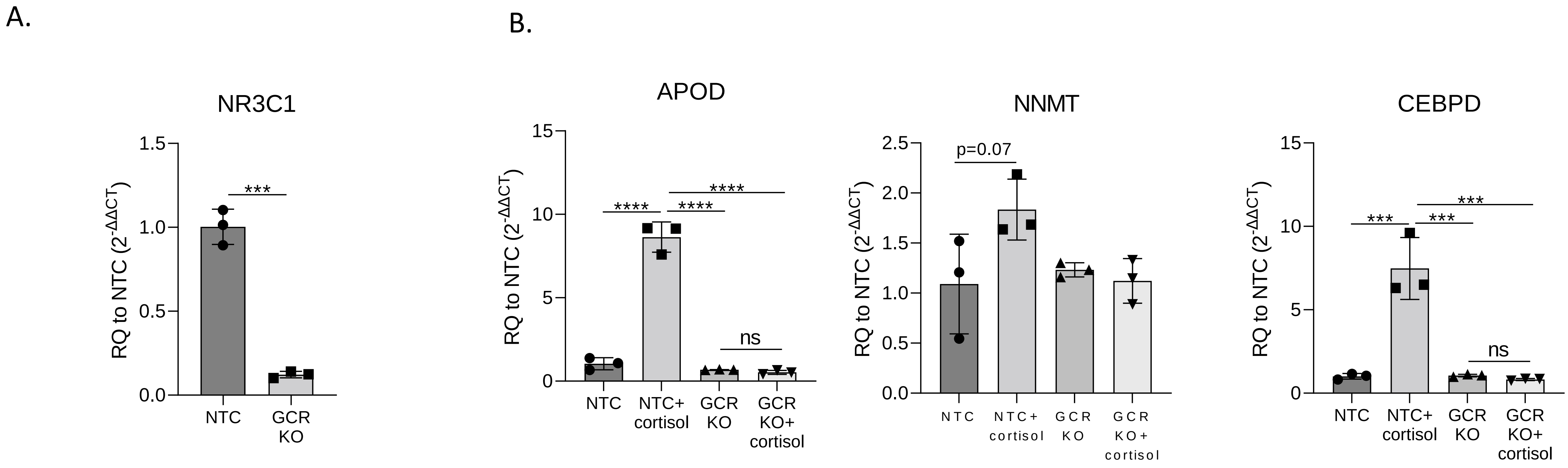

Supplementary Figure 5. A. CRISPR-Cas9 of the glucocorticoid receptor gene, *NR3C1*, results in significant reduction of *NR3C1* gene expression compared to the non-targeting control (NTC). B. *NR3C1* knockout renders cells unresponsive to cortisol.

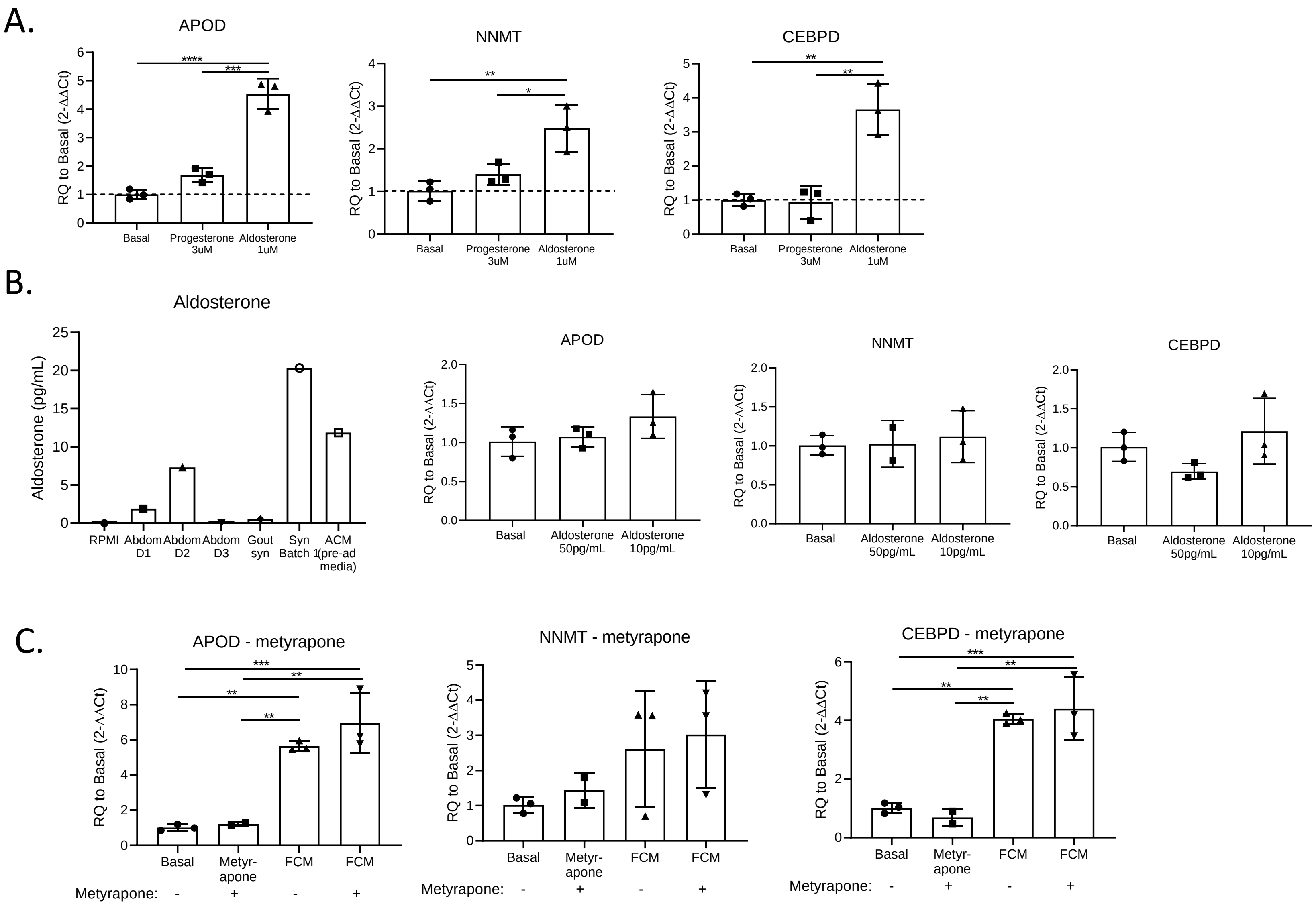

Supplementary Figure 6. A. Progesterone has no activity; aldosterone contains activity at 3µM. B. Aldosterone levels in FCM as measured by ELISA; physiologically relevant levels contain no activity. C. Blocking Hydroxysteroid 11-Beta Dehydrogenase 1 conversion of cortisone to cortisol with metyrapone does not block FCM activity.

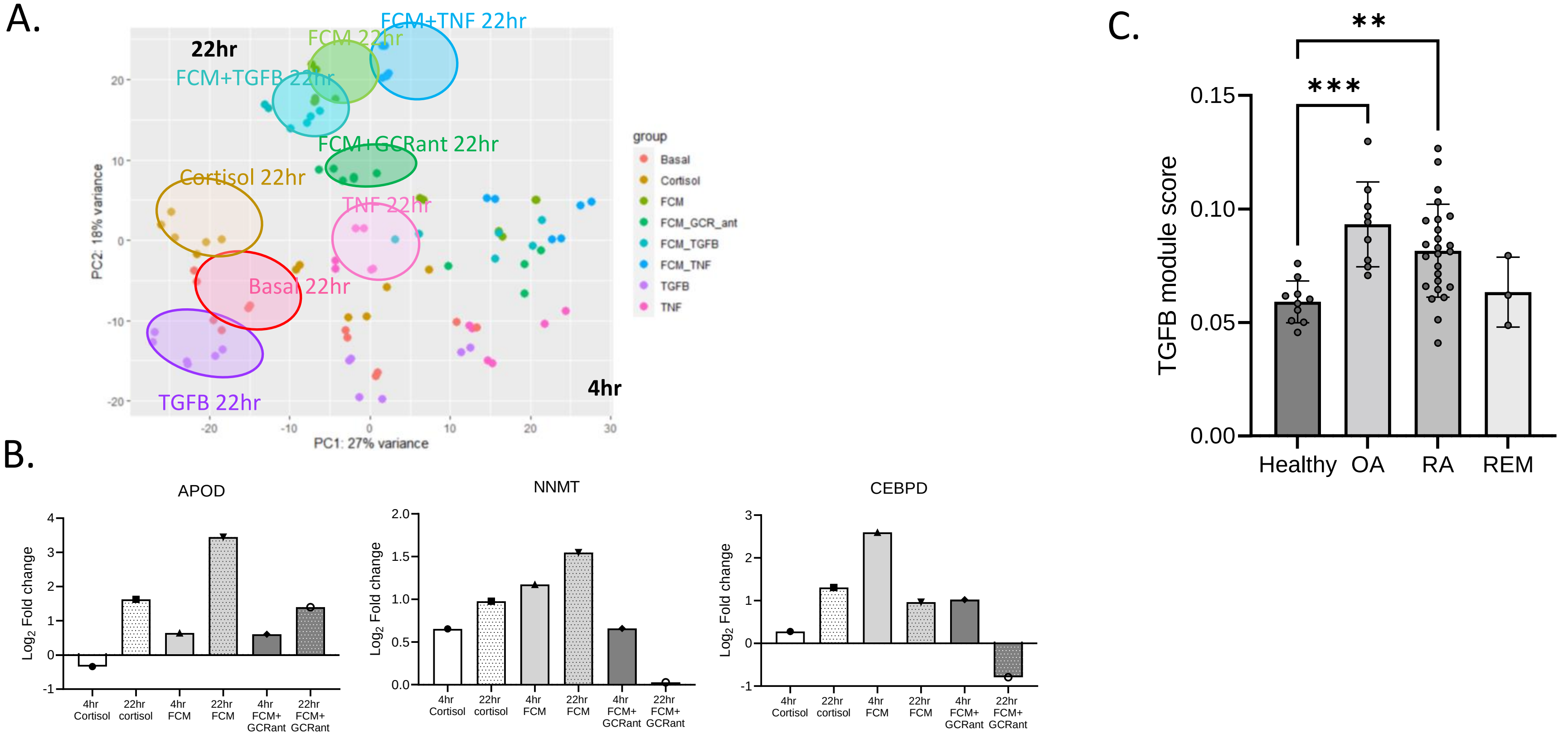

Supplementary Figure 7. A. PCA of bulk RNA sequencing samples. B. Verification that healthy fibroblast signature was induced by FCM and cortisol and blocked by the addition of a GCR antagonist. C. Bulk-RNA sequencing defined TGFB activation score was applied to a pseudobulk analysis of single cell RNA sequenced synovial fibroblasts.

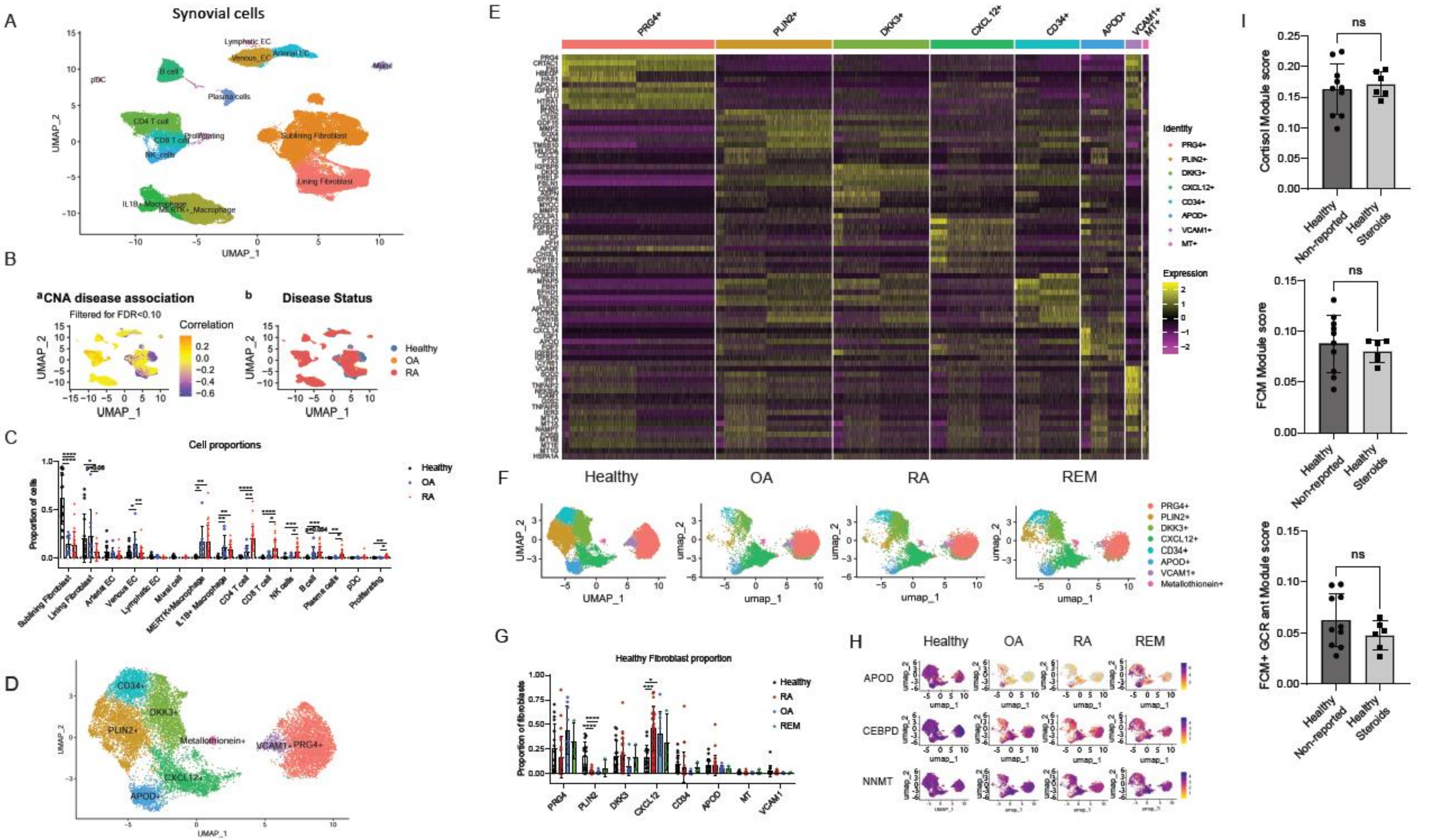

Supplementary Figure 8. Analysis of healthy steroid users and non-users. A. Synovial cells from all healthy steroid users and non-users, OA, and naïve RA donors were harmonized and clustered into a single UMAP. B. UMAP projection from (A) colored by correlation with arthritis (orange) or health (purple) using covarying neighborhood analysis (CNA). C. Cell cluster proportions among healthy, OA, and naïve RA synovial cells and UMAP separated by disease state. D. Fine clustering analysis on healthy synovial fibroblasts defines 8 distinct clusters. E. Heatmap of the top 10 DEGs per cluster. F. Symphony mapping of OA, naïve RA, and remission fibroblasts to healthy synovial fibroblast reference. G. Quantification of fibroblast proportions mapping to each cluster. H. Healthy synovium is enriched in APOD, CEBPD, and NNMT expression compared to OA and RA fibroblasts, and is partially restored in remission fibroblasts. I. Application of bulk-RNA sequencing defined module scoring to single cell pseudobulk data (reads collapsed over patient) reveals that FCM and cortisol activation is similar in fibroblasts from healthy steroid users and non-users.

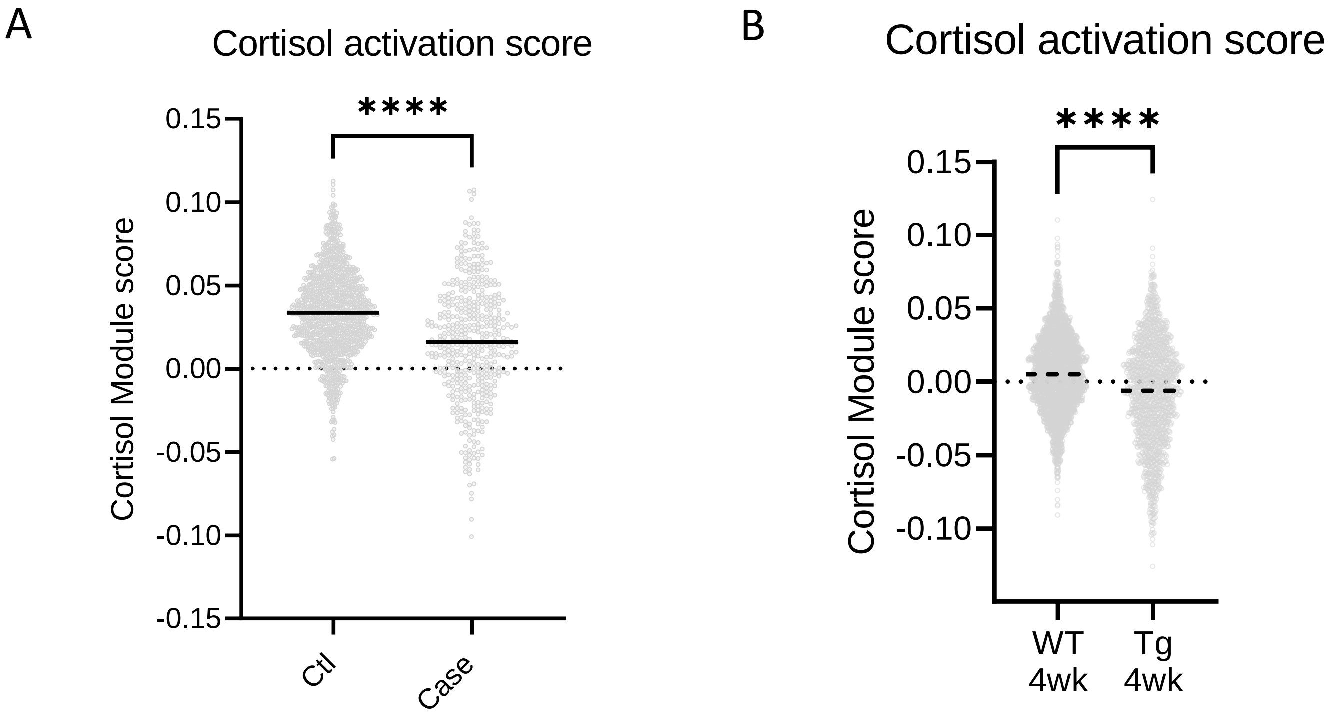

Supplementary Figure 9. A. Cortisol activation score, as defined by bulk RNA sequencing, was applied to wildtype mouse synovial sublining fibroblasts from Wei et. al.. Ctl= healthy, case=serum transfer arthritis, day 10. B. The cortisol activation score was applied to synovial fibroblasts from the Armaka et. al. WT 4wk= healthy, Tg 4wk= hTNFtg spontaneous arthritis mice

A.

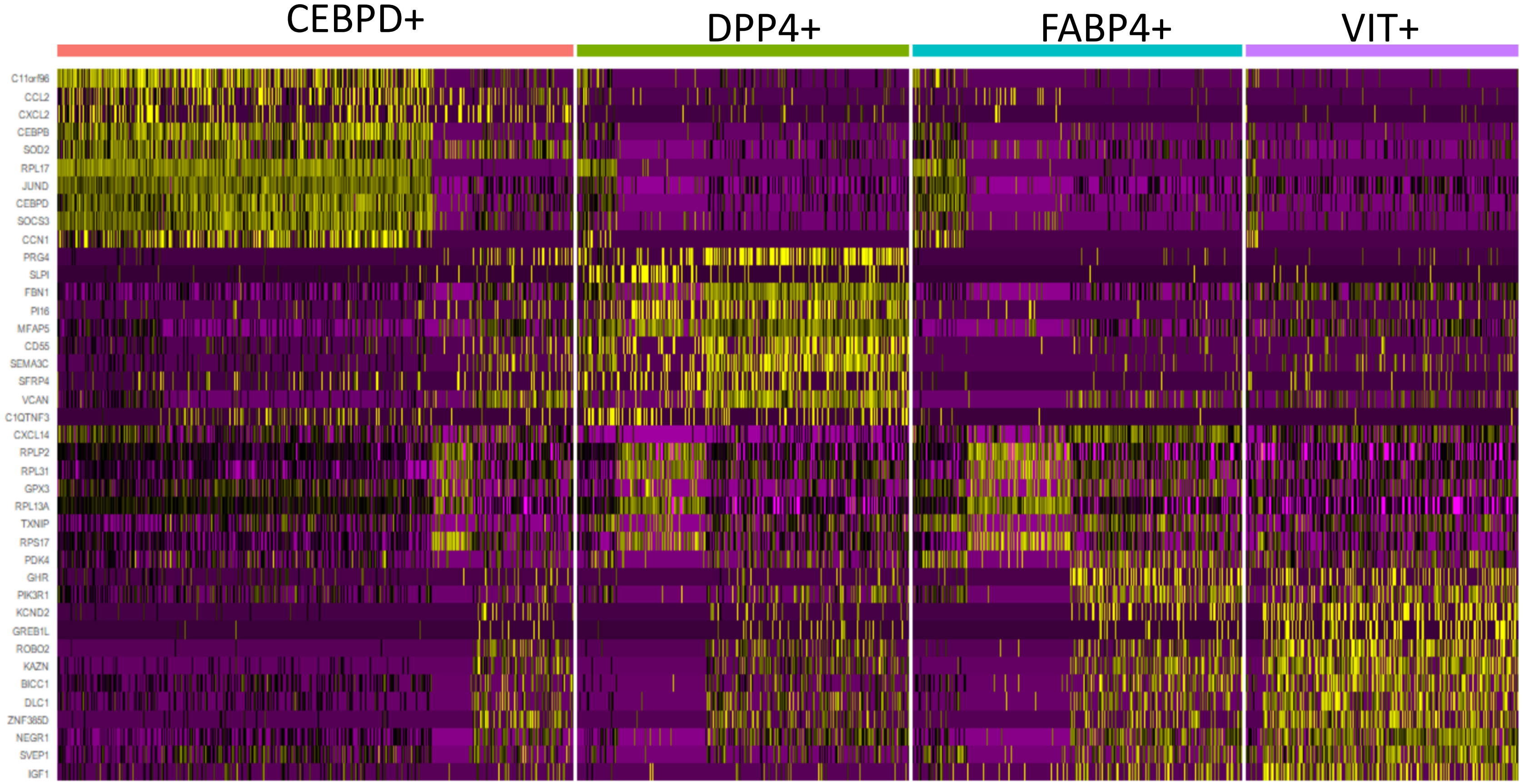

Supplementary Figure 10. A. Top ten markers of PDGFRα+ non mesothelial cells from VAT (visceral) and SAT (subcutaneous) adipose tissue.

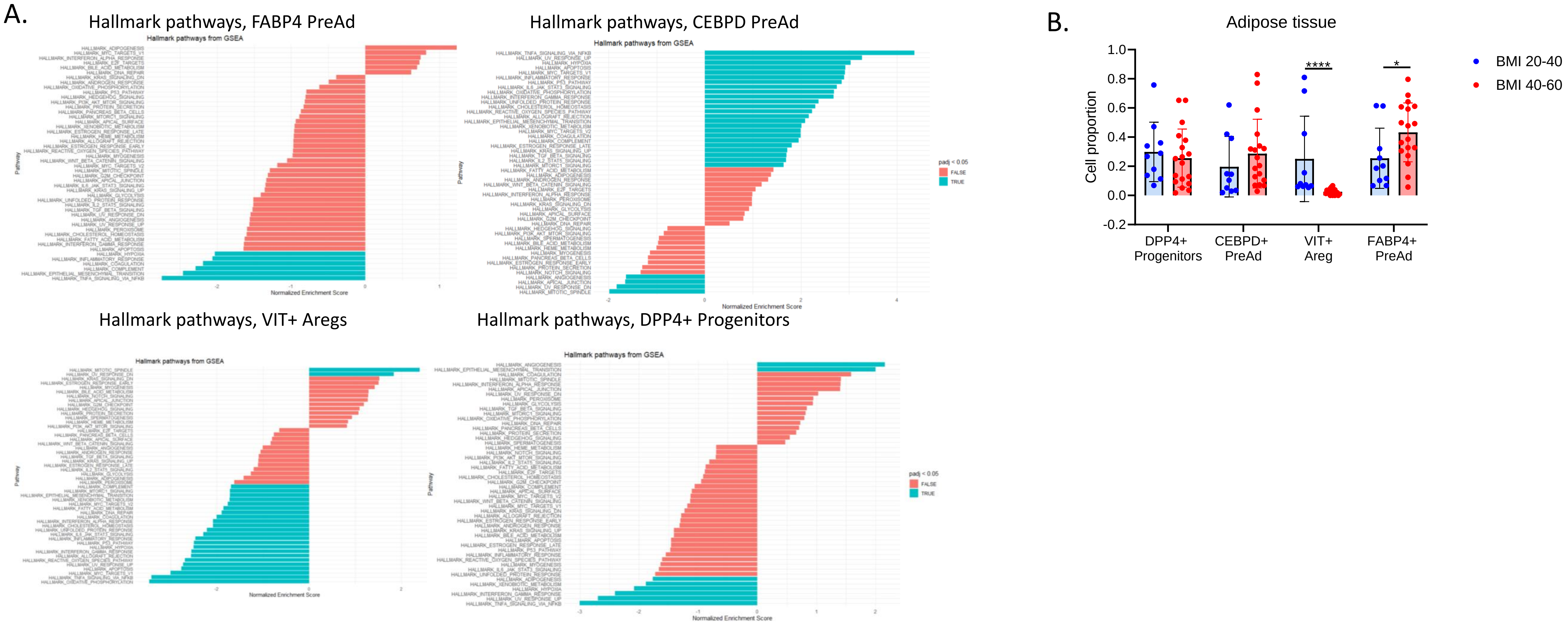

Supplementary Figure 11. A. FGSEA on adipose populations using Hallmark Pathways. B. Adipose stromal population proportions by BMI.

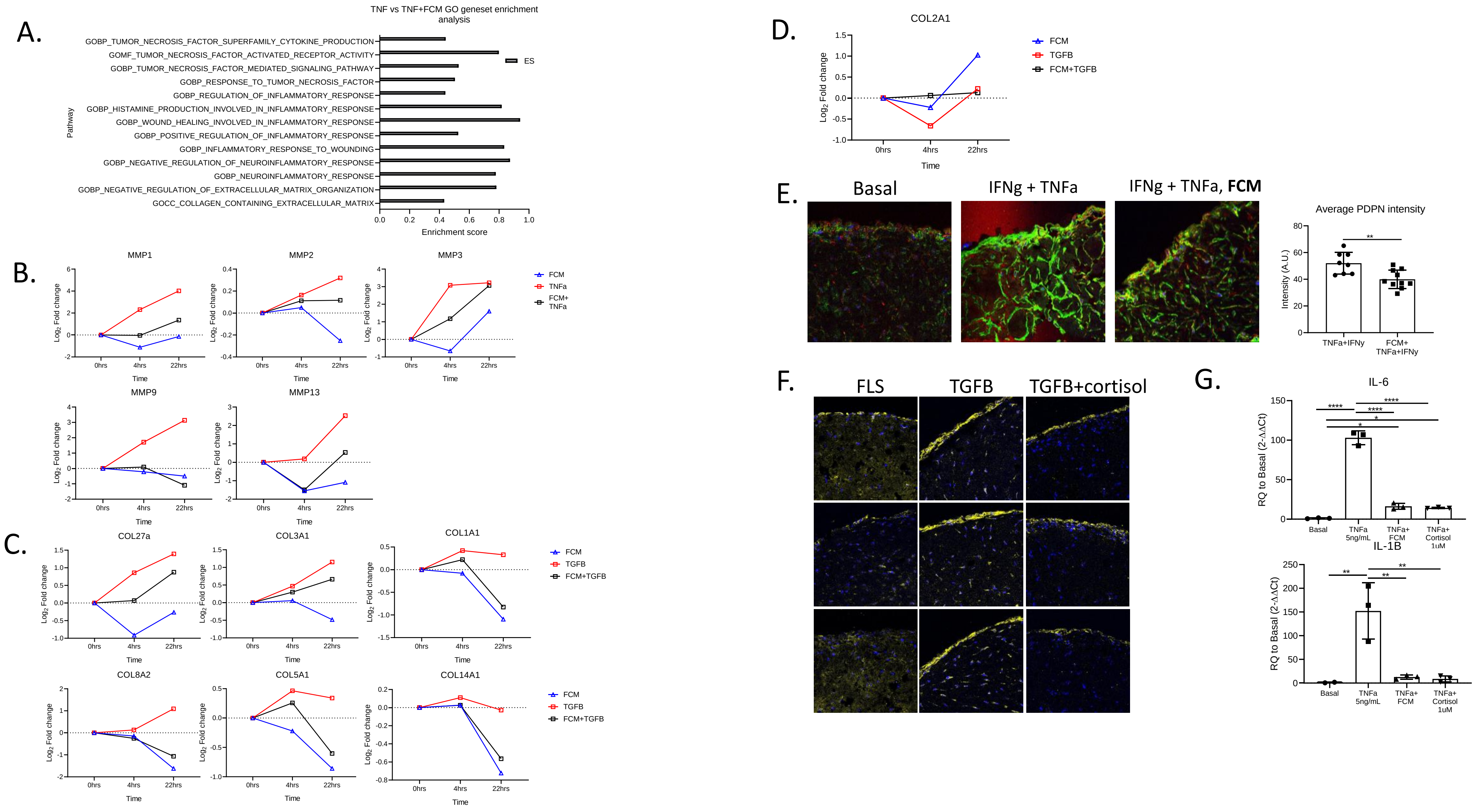

Supplementary Figure 12. A. GSEA analysis using GO terms on bulk RNA sequencing data. Enriched pathways are "rescued" by FCM to basal levels. B. FCM reduces MMP upregulation by TNFa. C. FCM rescues fibrotic collagens upregulated by TGFB. D. FCM does not impact non-fibrotic collagens. E. PDPN staining of micromass sections and intensity quantification performed in ImageJ. F. Cortisol (1uM) prevents TGFB (10ng/mL) induced fibrosis as measured by collagen 1a1 immunostaining (Day 21). G. FCM and cortisol inhibit TNFa induced pro-inflammatory changes.
