## Supplementary material for "Adipocytes regulate fibroblast function, and their loss contributes to fibroblast dysfunction in inflammatory diseases": Data file S1

Data File S1 Sublining Healthy vs RA Gene expression

| | p_val | avg_log2F( | pct.1 | pct.2 | p_val_adj |
|---|---|---|---|---|---|
| APOD | 0 | 4.953941 | 0.787 | 0.076 | 0 |
| PLIN2 | 0 | 4.368146 | 0.834 | 0.369 | 0 |
| MT1X | 0 | 4.111982 | 0.971 | 0.753 | 0 |
| GPX3 | 0 | 3.73811 | 0.977 | 0.496 | 0 |
| DEPP1 | 0 | 3.574812 | 0.822 | 0.204 | 0 |
| PTX3 | 0 | 3.566984 | 0.442 | 0.143 | 0 |
| MT1E | 0 | 3.367762 | 0.949 | 0.805 | 0 |
| MT1M | 0 | 3.144319 | 0.882 | 0.681 | 0 |
| MT2A | 0 | 3.110782 | 0.987 | 0.936 | 0 |
| GLUL | 0 | 3.066753 | 0.933 | 0.645 | 0 |
| DDIT4 | 0 | 2.927992 | 0.702 | 0.399 | 0 |
| ADH1B | 0 | 2.827494 | 0.499 | 0.006 | 0 |
| ADM | 0 | 2.641874 | 0.79 | 0.297 | 0 |
| HILPDA | 0 | 2.61784 | 0.524 | 0.305 | 0 |
| ANGPTL7 | 9.11E-292 | 2.601365 | 0.167 | 0.016 | 3.46E-287 |
| CEBPD | 0 | 2.443947 | 0.97 | 0.806 | 0 |
| C11orf96 | 0 | 2.422059 | 0.857 | 0.491 | 0 |
| CYP4B1 | 0 | 2.350877 | 0.642 | 0.013 | 0 |
| GADD45B | 0 | 2.285589 | 0.886 | 0.635 | 0 |
| MT1A | 0 | 2.215165 | 0.441 | 0.169 | 0 |
| NFKBIA | 0 | 2.047526 | 0.887 | 0.7 | 0 |
| MGST1 | 0 | 1.9633 | 0.913 | 0.507 | 0 |
| MYOC | 0 | 1.884969 | 0.313 | 0.045 | 0 |
| GALNT15 | 0 | 1.881702 | 0.864 | 0.385 | 0 |
| IGFBP6 | 0 | 1.736083 | 0.92 | 0.749 | 0 |
| ACKR3 | 0 | 1.707784 | 0.918 | 0.679 | 0 |
| TSC22D3 | 0 | 1.692977 | 0.885 | 0.742 | 0 |
| RHOB | 0 | 1.661533 | 0.926 | 0.747 | 0 |
| SLPI | 0 | 1.607485 | 0.563 | 0.142 | 0 |
| RARRES1 | 2.76E-111 | 1.606953 | 0.492 | 0.364 | 1.05E-106 |
| DUSP1 | 0 | 1.577839 | 0.962 | 0.89 | 0 |
| SOD2 | 0 | 1.56371 | 0.914 | 0.778 | 0 |
| GSN | 0 | 1.554975 | 0.999 | 0.996 | 0 |
| GADD45A | 1.94E-101 | 1.51505 | 0.498 | 0.404 | 7.37E-97 |
| TXNIP | 0 | 1.472446 | 0.965 | 0.912 | 0 |
| PNRC1 | 0 | 1.415987 | 0.944 | 0.875 | 0 |
| ZBTB16 | 0 | 1.392171 | 0.715 | 0.142 | 0 |
| PDK4 | 0 | 1.385903 | 0.577 | 0.189 | 0 |
| CXCL2 | 1.57E-163 | 1.377125 | 0.245 | 0.095 | 5.96E-159 |
| RAMP2 | 0 | 1.375645 | 0.741 | 0.479 | 0 |
| GDF15 | 0 | 1.368144 | 0.277 | 0.062 | 0 |
| MID1IP1 | 0 | 1.359122 | 0.613 | 0.284 | 0 |
| CFD | 0 | 1.325145 | 0.997 | 0.865 | 0 |
| MTRNR2L8 | 0 | 1.31798 | 0.867 | 0.753 | 0 |
| MT1G | 1.70E-146 | 1.310858 | 0.184 | 0.058 | 6.46E-142 |

| | | | | | |
|---|---|---|---|---|---|
| ADH5 | 0 | 1.280496 | 0.881 | 0.759 | 0 |
| MAOA | 0 | 1.279415 | 0.646 | 0.099 | 0 |
| ZFAND5 | 0 | 1.279346 | 0.785 | 0.54 | 0 |
| H1FX | 0 | 1.268343 | 0.833 | 0.744 | 0 |
| TSKU | 0 | 1.267889 | 0.568 | 0.281 | 0 |
| PLPP3 | 0 | 1.263314 | 0.879 | 0.712 | 0 |
| CDO1 | 0 | 1.259183 | 0.683 | 0.499 | 0 |
| KLF9 | 0 | 1.256258 | 0.847 | 0.52 | 0 |
| PLA2G2A | 0 | 1.25239 | 0.987 | 0.908 | 0 |
| NAMPT | 1.41E-223 | 1.250092 | 0.695 | 0.555 | 5.36E-219 |
| ADAMTS5 | 0 | 1.23459 | 0.657 | 0.343 | 0 |
| FTH1 | 0 | 1.20395 | 0.999 | 1 | 0 |
| ZFP36 | 0 | 1.203497 | 0.905 | 0.76 | 0 |
| ADH1C | 0 | 1.159797 | 0.288 | 0.038 | 0 |
| TGFBR3 | 0 | 1.141457 | 0.889 | 0.696 | 0 |
| PROCR | 0 | 1.139733 | 0.777 | 0.652 | 0 |
| FBLN1 | 0 | 1.138816 | 0.929 | 0.753 | 0 |
| GLRX | 3.93E-254 | 1.134449 | 0.671 | 0.527 | 1.49E-249 |
| YBX3 | 0 | 1.129092 | 0.976 | 0.95 | 0 |
| CEBPB | 0 | 1.121722 | 0.885 | 0.819 | 0 |
| SLC39A14 | 2.36E-296 | 1.121559 | 0.658 | 0.472 | 8.95E-292 |
| FKBP5 | 0 | 1.121427 | 0.666 | 0.312 | 0 |
| BCL6 | 0 | 1.108191 | 0.677 | 0.269 | 0 |
| KLF6 | 0 | 1.087967 | 0.858 | 0.736 | 0 |
| NFIL3 | 0 | 1.081063 | 0.646 | 0.281 | 0 |
| IER2 | 3.09E-290 | 1.07991 | 0.868 | 0.752 | 1.17E-285 |
| IER3 | 0.333756 | 1.077076 | 0.477 | 0.596 | 1 |
| PPP1R15A | 0 | 1.074261 | 0.849 | 0.594 | 0 |
| ZFP36L2 | 2.04E-258 | 1.069614 | 0.913 | 0.891 | 7.75E-254 |
| NOVA1 | 0 | 1.067629 | 0.805 | 0.617 | 0 |
| LAMA2 | 0 | 1.066721 | 0.571 | 0.17 | 0 |
| KLF4 | 0 | 1.052873 | 0.831 | 0.607 | 0 |
| FBLN2 | 0 | 1.051588 | 0.85 | 0.572 | 0 |
| SLC3A2 | 0 | 1.027117 | 0.783 | 0.589 | 0 |
| PCOLCE2 | 0 | 1.015667 | 0.896 | 0.656 | 0 |
| VEGFA | 0 | 1.010268 | 0.51 | 0.26 | 0 |
| UAP1 | 0 | 1.007844 | 0.842 | 0.708 | 0 |
| JUN | 0 | 0.995075 | 0.976 | 0.914 | 0 |
| NNMT | 0 | 0.99445 | 0.987 | 0.965 | 0 |
| GABARAPL | 0 | 0.993839 | 0.786 | 0.573 | 0 |
| CXCL1 | 2.48E-45 | 0.978262 | 0.156 | 0.084 | 9.40E-41 |
| G0S2 | 0.723067 | 0.977818 | 0.276 | 0.288 | 1 |
| F3 | 1.87E-265 | 0.958606 | 0.443 | 0.201 | 7.09E-261 |
| HIST1H1C | 0 | 0.950358 | 0.463 | 0.177 | 0 |
| PER1 | 0 | 0.948063 | 0.648 | 0.257 | 0 |
| MTSS1 | 0 | 0.942852 | 0.494 | 0.13 | 0 |
| CSF1 | 0 | 0.939101 | 0.67 | 0.462 | 0 |

| | | | | | |
|---|---|---|---|---|---|
| SUN2 | 0 | 0.938835 | 0.683 | 0.4 | 0 |
| NR1D1 | 0 | 0.9224 | 0.574 | 0.204 | 0 |
| DCN | 0 | 0.914075 | 1 | 0.999 | 0 |
| CHST7 | 0 | 0.898731 | 0.44 | 0.062 | 0 |
| PHC2 | 0 | 0.895994 | 0.653 | 0.423 | 0 |
| FOXO1 | 0 | 0.892358 | 0.636 | 0.358 | 0 |
| PID1 | 0 | 0.884397 | 0.546 | 0.201 | 0 |
| NUPR1 | 0 | 0.883655 | 0.967 | 0.96 | 0 |
| RPS29 | 0 | 0.88024 | 0.991 | 0.995 | 0 |
| MFAP5 | 0 | 0.877983 | 0.801 | 0.413 | 0 |
| FIBIN | 0 | 0.876463 | 0.509 | 0.236 | 0 |
| MT-ND4L | 0 | 0.874645 | 0.97 | 0.947 | 0 |
| SASH1 | 0 | 0.872704 | 0.72 | 0.515 | 0 |
| MYC | 2.76E-306 | 0.871472 | 0.536 | 0.289 | 1.05E-301 |
| PRELP | 8.91E-224 | 0.870921 | 0.789 | 0.629 | 3.38E-219 |
| CRYAB | 3.41E-137 | 0.868435 | 0.765 | 0.756 | 1.30E-132 |
| AGTR1 | 0 | 0.857925 | 0.519 | 0.169 | 0 |
| LTBP2 | 1.71E-248 | 0.850351 | 0.597 | 0.397 | 6.49E-244 |
| MIF | 0 | 0.850302 | 0.781 | 0.664 | 0 |
| H3F3B | 0 | 0.841484 | 0.997 | 0.998 | 0 |
| HSD11B1 | 0 | 0.833892 | 0.525 | 0.15 | 0 |
| NID1 | 0 | 0.833359 | 0.544 | 0.257 | 0 |
| RPS27 | 0 | 0.832768 | 1 | 1 | 0 |
| ARID5B | 0 | 0.830652 | 0.853 | 0.787 | 0 |
| ANGPTL4 | 7.81E-11 | 0.818946 | 0.45 | 0.452 | 2.97E-06 |
| DHRS3 | 0 | 0.813736 | 0.652 | 0.371 | 0 |
| ERRFI1 | 3.33E-180 | 0.811654 | 0.556 | 0.369 | 1.26E-175 |
| RPL38 | 0 | 0.806321 | 0.987 | 0.989 | 0 |
| SMIM3 | 5.32E-265 | 0.794868 | 0.542 | 0.334 | 2.02E-260 |
| VIT | 2.95E-287 | 0.78476 | 0.594 | 0.38 | 1.12E-282 |
| NEGR1 | 0 | 0.773812 | 0.497 | 0.12 | 0 |
| ABLIM1 | 0 | 0.770741 | 0.631 | 0.354 | 0 |
| H2AFJ | 0 | 0.765263 | 0.884 | 0.815 | 0 |
| USP53 | 5.47E-142 | 0.761028 | 0.56 | 0.435 | 2.08E-137 |
| ID2 | 9.55E-124 | 0.757258 | 0.662 | 0.576 | 3.63E-119 |
| H2AFZ | 3.35E-186 | 0.755827 | 0.765 | 0.728 | 1.27E-181 |
| CBLB | 0 | 0.753835 | 0.594 | 0.308 | 0 |
| CTSL | 1.04E-300 | 0.752151 | 0.876 | 0.854 | 3.96E-296 |
| CPE | 1.66E-243 | 0.745783 | 0.475 | 0.23 | 6.30E-239 |
| ANG | 0 | 0.741192 | 0.598 | 0.349 | 0 |
| MEDAG | 1.39E-283 | 0.735801 | 0.771 | 0.625 | 5.26E-279 |
| BNIP3 | 5.18E-216 | 0.732021 | 0.578 | 0.426 | 1.97E-211 |
| TFPI | 1.91E-256 | 0.731578 | 0.751 | 0.605 | 7.24E-252 |
| JUNB | 2.25E-168 | 0.7254 | 0.923 | 0.91 | 8.54E-164 |
| TWIST2 | 0 | 0.723735 | 0.519 | 0.218 | 0 |
| CCDC71L | 8.39E-58 | 0.722 | 0.42 | 0.351 | 3.18E-53 |
| SOCS3 | 1.49E-136 | 0.717841 | 0.784 | 0.714 | 5.66E-132 |

| | | | | | |
|---|---|---|---|---|---|
| RND3 | 1.49E-109 | 0.717162 | 0.655 | 0.579 | 5.65E-105 |
| AC103591. | 0 | 0.712432 | 0.321 | 0.068 | 0 |
| SPSB1 | 0 | 0.711506 | 0.575 | 0.301 | 0 |
| SLC2A3 | 5.45E-242 | 0.698109 | 0.473 | 0.255 | 2.07E-237 |
| EIF1 | 0 | 0.69058 | 0.999 | 0.999 | 0 |
| RPL37A | 0 | 0.689892 | 0.998 | 0.999 | 0 |
| WNT11 | 0 | 0.68625 | 0.4 | 0.132 | 0 |
| SQSTM1 | 0 | 0.683037 | 0.939 | 0.909 | 0 |
| HSPA1B | 4.82E-77 | 0.682966 | 0.541 | 0.468 | 1.83E-72 |
| HMGB2 | 1.16E-282 | 0.679806 | 0.645 | 0.437 | 4.41E-278 |
| ABCA9 | 0 | 0.673876 | 0.345 | 0.04 | 0 |
| NDRG1 | 5.71E-169 | 0.672727 | 0.759 | 0.679 | 2.17E-164 |
| BTG3 | 0 | 0.667727 | 0.596 | 0.374 | 0 |
| DEFB1 | 0 | 0.66638 | 0.352 | 0.103 | 0 |
| LARP6 | 0 | 0.666349 | 0.667 | 0.469 | 0 |
| MTRNR2L1 | 0 | 0.665763 | 0.982 | 0.96 | 0 |
| FOS | 1.81E-153 | 0.664609 | 0.952 | 0.92 | 6.88E-149 |
| MT-ND2 | 0 | 0.657146 | 0.998 | 0.999 | 0 |
| ADH4 | 0 | 0.657054 | 0.16 | 0.003 | 0 |
| RASD1 | 5.46E-82 | 0.65631 | 0.357 | 0.226 | 2.07E-77 |
| REV3L | 4.67E-189 | 0.652857 | 0.746 | 0.659 | 1.77E-184 |
| GPRC5A | 6.33E-226 | 0.652506 | 0.439 | 0.229 | 2.40E-221 |
| MFGE8 | 2.54E-203 | 0.651108 | 0.783 | 0.682 | 9.64E-199 |
| PTGIS | 2.26E-236 | 0.637356 | 0.443 | 0.214 | 8.57E-232 |
| DDX21 | 2.35E-153 | 0.63097 | 0.656 | 0.583 | 8.92E-149 |
| RPL37 | 0 | 0.628142 | 0.998 | 0.999 | 0 |
| SPOCK1 | 3.01E-211 | 0.628003 | 0.563 | 0.371 | 1.14E-206 |
| ABCA8 | 2.94E-205 | 0.626633 | 0.544 | 0.325 | 1.12E-200 |
| DCXR | 1.56E-140 | 0.62078 | 0.503 | 0.376 | 5.91E-136 |
| ZFAS1 | 0 | 0.619769 | 0.935 | 0.928 | 0 |
| RPS4Y1 | 5.20E-272 | 0.618627 | 0.231 | 0.046 | 1.97E-267 |
| PIM3 | 4.68E-193 | 0.618038 | 0.462 | 0.279 | 1.78E-188 |
| VASN | 1.52E-103 | 0.615874 | 0.67 | 0.631 | 5.78E-99 |
| C3 | 1.60E-249 | 0.614234 | 0.741 | 0.518 | 6.08E-245 |
| PMP22 | 1.35E-224 | 0.606735 | 0.95 | 0.967 | 5.12E-220 |
| MAP3K8 | 5.05E-205 | 0.605169 | 0.463 | 0.28 | 1.92E-200 |
| METTL7A | 9.78E-176 | 0.604749 | 0.689 | 0.596 | 3.71E-171 |
| RPL36 | 0 | 0.60325 | 0.994 | 0.996 | 0 |
| RETREG1 | 9.64E-143 | 0.602557 | 0.736 | 0.666 | 3.66E-138 |
| PQLC2L | 0 | 0.600601 | 0.336 | 0.071 | 0 |
| SNHG8 | 2.52E-226 | 0.600026 | 0.771 | 0.698 | 9.55E-222 |
| MAP1LC3E | 3.56E-261 | 0.592109 | 0.775 | 0.688 | 1.35E-256 |
| TNFRSF11I | 2.88E-59 | 0.589773 | 0.432 | 0.344 | 1.09E-54 |
| ITGA5 | 3.32E-218 | 0.580617 | 0.555 | 0.362 | 1.26E-213 |
| ST3GAL5 | 2.08E-278 | 0.579454 | 0.393 | 0.156 | 7.91E-274 |
| ANGPTL5 | 3.23E-112 | 0.579207 | 0.552 | 0.444 | 1.23E-107 |
| LTBP4 | 1.17E-150 | 0.57867 | 0.844 | 0.799 | 4.45E-146 |

| | | | | | |
|---|---|---|---|---|---|
| WTAP | 4.20E-140 | 0.570885 | 0.695 | 0.619 | 1.60E-135 |
| CILP | 2.36E-113 | 0.568526 | 0.384 | 0.232 | 8.98E-109 |
| ABCA6 | 1.01E-278 | 0.56846 | 0.414 | 0.17 | 3.83E-274 |
| RPS25 | 0 | 0.567015 | 0.995 | 0.997 | 0 |
| RHEB | 1.87E-282 | 0.558365 | 0.829 | 0.761 | 7.09E-278 |
| HMOX1 | 5.47E-128 | 0.555283 | 0.304 | 0.157 | 2.08E-123 |
| CYGB | 2.18E-171 | 0.551757 | 0.539 | 0.358 | 8.27E-167 |
| IFITM2 | 1.20E-135 | 0.551112 | 0.897 | 0.922 | 4.56E-131 |
| PIM1 | 7.66E-164 | 0.548285 | 0.341 | 0.171 | 2.91E-159 |
| ADGRD1 | 0 | 0.546801 | 0.395 | 0.131 | 0 |
| WBP2 | 9.06E-252 | 0.545868 | 0.58 | 0.383 | 3.44E-247 |
| C15orf61 | 1.58E-171 | 0.54518 | 0.57 | 0.439 | 6.00E-167 |
| STOM | 1.76E-195 | 0.543278 | 0.742 | 0.653 | 6.68E-191 |
| BTG2 | 2.92E-130 | 0.537504 | 0.581 | 0.419 | 1.11E-125 |
| UBC | 2.45E-147 | 0.536844 | 0.994 | 0.998 | 9.30E-143 |
| RPS20 | 0 | 0.535767 | 0.99 | 0.994 | 0 |
| PIK3IP1 | 3.61E-162 | 0.531829 | 0.395 | 0.23 | 1.37E-157 |
| ELL2 | 1.81E-145 | 0.528442 | 0.444 | 0.279 | 6.86E-141 |
| FBXO32 | 1.43E-77 | 0.526304 | 0.4 | 0.289 | 5.42E-73 |
| CDKN1A | 4.71E-175 | 0.524507 | 0.474 | 0.274 | 1.79E-170 |
| RSL24D1 | 2.77E-256 | 0.523856 | 0.812 | 0.763 | 1.05E-251 |
| COL12A1 | 1.29E-146 | 0.523672 | 0.679 | 0.561 | 4.91E-142 |
| BHLHE40 | 4.09E-42 | 0.52216 | 0.491 | 0.449 | 1.55E-37 |
| CITED2 | 1.91E-29 | 0.518151 | 0.452 | 0.427 | 7.24E-25 |
| ACYP1 | 2.09E-200 | 0.517997 | 0.432 | 0.239 | 7.95E-196 |
| RPL39 | 0 | 0.513983 | 0.996 | 0.998 | 0 |
| DSE | 3.20E-58 | 0.513046 | 0.537 | 0.502 | 1.21E-53 |
| PNPLA2 | 1.05E-173 | 0.511061 | 0.489 | 0.322 | 3.99E-169 |
| IL1R1 | 8.10E-141 | 0.508943 | 0.691 | 0.6 | 3.08E-136 |
| CORO6 | 5.14E-296 | 0.508705 | 0.282 | 0.074 | 1.95E-291 |
| SESTD1 | 1.14E-119 | 0.507357 | 0.634 | 0.554 | 4.32E-115 |
| PXDC1 | 7.25E-177 | 0.506888 | 0.674 | 0.562 | 2.75E-172 |
| GYPC | 5.31E-160 | 0.50279 | 0.787 | 0.754 | 2.02E-155 |
| MTHFD2 | 1.58E-249 | 0.499575 | 0.369 | 0.156 | 5.99E-245 |
| RTN4 | 1.12E-294 | 0.495727 | 0.955 | 0.949 | 4.25E-290 |
| CRISPLD2 | 2.85E-112 | 0.492696 | 0.441 | 0.292 | 1.08E-107 |
| RGS16 | 9.39E-51 | 0.492078 | 0.228 | 0.142 | 3.57E-46 |
| HAS1 | 0.000109 | 0.492026 | 0.354 | 0.356 | 1 |
| IFRD1 | 6.65E-146 | 0.490222 | 0.545 | 0.401 | 2.53E-141 |
| DDIT3 | 9.68E-78 | 0.490165 | 0.496 | 0.413 | 3.68E-73 |
| SLC43A3 | 1.89E-162 | 0.489101 | 0.519 | 0.357 | 7.17E-158 |
| HSD17B11 | 1.74E-183 | 0.488586 | 0.619 | 0.472 | 6.61E-179 |
| SLC16A7 | 5.45E-147 | 0.487189 | 0.494 | 0.336 | 2.07E-142 |
| CCNL1 | 7.17E-160 | 0.487095 | 0.807 | 0.712 | 2.72E-155 |
| ATF3 | 5.51E-162 | 0.483515 | 0.552 | 0.353 | 2.09E-157 |
| PRG4 | 3.50E-158 | 0.481384 | 0.979 | 0.851 | 1.33E-153 |
| TAF1D | 1.12E-123 | 0.47979 | 0.728 | 0.693 | 4.26E-119 |

| | | | | | |
|---|---|---|---|---|---|
| ECE1 | 6.35E-187 | 0.479445 | 0.493 | 0.302 | 2.41E-182 |
| 3-Mar | 1.10E-135 | 0.475782 | 0.356 | 0.201 | 4.17E-131 |
| EIF1B | 3.07E-180 | 0.475764 | 0.773 | 0.743 | 1.17E-175 |
| THBS1 | 1.22E-34 | 0.475434 | 0.311 | 0.24 | 4.62E-30 |
| PIK3R1 | 8.07E-67 | 0.474653 | 0.624 | 0.585 | 3.07E-62 |
| MMP14 | 4.87E-114 | 0.470271 | 0.733 | 0.678 | 1.85E-109 |
| EIF4A1 | 2.88E-147 | 0.469415 | 0.792 | 0.756 | 1.09E-142 |
| RHOQ | 1.65E-151 | 0.469104 | 0.683 | 0.574 | 6.26E-147 |
| RPS28 | 0 | 0.468272 | 0.999 | 0.999 | 0 |
| CYB5A | 2.78E-110 | 0.467644 | 0.609 | 0.552 | 1.06E-105 |
| RPS16 | 0 | 0.461735 | 0.995 | 0.998 | 0 |
| KLF15 | 0 | 0.461525 | 0.223 | 0.007 | 0 |
| TAF7 | 3.45E-167 | 0.461453 | 0.734 | 0.634 | 1.31E-162 |
| EFHD1 | 1.87E-287 | 0.460407 | 0.245 | 0.052 | 7.09E-283 |
| HSPA5 | 5.18E-41 | 0.459975 | 0.831 | 0.949 | 1.97E-36 |
| ERICH1 | 2.06E-127 | 0.459564 | 0.552 | 0.434 | 7.81E-123 |
| EPB41L4A- | 9.37E-150 | 0.459299 | 0.729 | 0.645 | 3.56E-145 |
| SDC4 | 3.39E-40 | 0.457552 | 0.37 | 0.31 | 1.29E-35 |
| CLIP1 | 2.98E-69 | 0.45668 | 0.629 | 0.604 | 1.13E-64 |
| PHGDH | 3.39E-148 | 0.456044 | 0.392 | 0.229 | 1.29E-143 |
| DPP4 | 9.42E-175 | 0.456038 | 0.439 | 0.226 | 3.58E-170 |
| STXBP6 | 1.95E-255 | 0.454009 | 0.399 | 0.169 | 7.40E-251 |
| H1F0 | 7.98E-16 | 0.453792 | 0.456 | 0.462 | 3.03E-11 |
| RPL31 | 6.45E-266 | 0.453025 | 0.948 | 0.956 | 2.45E-261 |
| TMBIM1 | 4.70E-152 | 0.452869 | 0.636 | 0.516 | 1.79E-147 |
| PERP | 1.17E-207 | 0.451761 | 0.32 | 0.13 | 4.43E-203 |
| ALDH1A1 | 1.92E-267 | 0.451543 | 0.238 | 0.052 | 7.29E-263 |
| HIST1H4C | 1.63E-78 | 0.450183 | 0.576 | 0.508 | 6.21E-74 |
| UBXN1 | 4.47E-175 | 0.4488 | 0.795 | 0.777 | 1.70E-170 |
| UBALD2 | 4.00E-186 | 0.447324 | 0.431 | 0.246 | 1.52E-181 |
| UBE2D3 | 3.72E-210 | 0.447174 | 0.84 | 0.802 | 1.41E-205 |
| ALDH2 | 1.45E-73 | 0.446894 | 0.617 | 0.587 | 5.52E-69 |
| SPRY1 | 1.06E-28 | 0.445705 | 0.455 | 0.412 | 4.02E-24 |
| MT-ATP8 | 6.11E-152 | 0.444068 | 0.753 | 0.665 | 2.32E-147 |
| MAFF | 5.46E-218 | 0.443005 | 0.463 | 0.236 | 2.07E-213 |
| HIGD1A | 2.53E-113 | 0.442975 | 0.494 | 0.388 | 9.61E-109 |
| RPLP2 | 0 | 0.442758 | 0.995 | 0.999 | 0 |
| HSPB8 | 2.72E-161 | 0.442542 | 0.381 | 0.209 | 1.03E-156 |
| MFAP4 | 1.59E-06 | 0.442026 | 0.693 | 0.761 | 0.060445 |
| NCOA7 | 1.06E-118 | 0.441777 | 0.608 | 0.505 | 4.02E-114 |
| CNOT8 | 3.91E-171 | 0.437404 | 0.382 | 0.211 | 1.48E-166 |
| PDPN | 1.78E-22 | 0.436775 | 0.623 | 0.666 | 6.76E-18 |
| KDSR | 5.63E-125 | 0.436184 | 0.655 | 0.577 | 2.14E-120 |
| FOSL2 | 4.35E-94 | 0.435222 | 0.486 | 0.375 | 1.65E-89 |
| BTF3 | 0 | 0.435017 | 0.98 | 0.99 | 0 |
| TRA2B | 7.90E-126 | 0.430499 | 0.676 | 0.607 | 3.00E-121 |
| CSRNP1 | 3.67E-199 | 0.429965 | 0.326 | 0.14 | 1.39E-194 |

| | | | | | |
|---|---|---|---|---|---|
| RPL7 | 8.24E-301 | 0.429776 | 0.991 | 0.996 | 3.13E-296 |
| GSTO1 | 2.31E-127 | 0.429751 | 0.805 | 0.801 | 8.77E-123 |
| RPL14 | 0 | 0.424773 | 0.997 | 0.998 | 0 |
| ATOH8 | 0 | 0.422282 | 0.248 | 0.036 | 0 |
| RPL23 | 7.89E-232 | 0.422012 | 0.977 | 0.986 | 3.00E-227 |
| RPS13 | 0 | 0.420511 | 0.995 | 0.998 | 0 |
| CRLF1 | 4.46E-15 | 0.418333 | 0.517 | 0.494 | 1.69E-10 |
| PNRC2 | 2.52E-119 | 0.417965 | 0.661 | 0.587 | 9.57E-115 |
| HIST1H1E | 1.32E-194 | 0.417919 | 0.312 | 0.131 | 5.02E-190 |
| MMD | 3.53E-221 | 0.411614 | 0.262 | 0.085 | 1.34E-216 |
| GBE1 | 2.53E-105 | 0.411039 | 0.415 | 0.29 | 9.62E-101 |
| BNIP3L | 7.39E-91 | 0.409905 | 0.755 | 0.745 | 2.81E-86 |
| LPCAT2 | 2.32E-238 | 0.408891 | 0.346 | 0.134 | 8.81E-234 |
| A4GALT | 6.75E-74 | 0.406588 | 0.643 | 0.582 | 2.56E-69 |
| RPL35 | 0 | 0.406471 | 0.994 | 0.998 | 0 |
| TIPARP | 5.38E-68 | 0.404616 | 0.321 | 0.22 | 2.04E-63 |
| TNFSF9 | 2.62E-91 | 0.404479 | 0.304 | 0.181 | 9.96E-87 |
| XG | 5.76E-83 | 0.402556 | 0.448 | 0.346 | 2.19E-78 |
| AXL | 1.81E-73 | 0.402515 | 0.676 | 0.623 | 6.85E-69 |
| KDM7A | 9.44E-126 | 0.402486 | 0.363 | 0.216 | 3.59E-121 |
| CNBP | 4.74E-193 | 0.40088 | 0.918 | 0.927 | 1.80E-188 |
| HNRNPF | 6.24E-148 | 0.40049 | 0.793 | 0.763 | 2.37E-143 |
| SLC19A2 | 0 | 0.399849 | 0.287 | 0.067 | 0 |
| TGIF1 | 5.21E-142 | 0.397418 | 0.453 | 0.293 | 1.98E-137 |
| SDCBP | 1.16E-101 | 0.394743 | 0.846 | 0.848 | 4.41E-97 |
| RAMP2-AS | 4.67E-223 | 0.394213 | 0.286 | 0.096 | 1.77E-218 |
| UBE2B | 2.49E-140 | 0.393108 | 0.858 | 0.856 | 9.45E-136 |
| FOXO3 | 1.69E-51 | 0.393004 | 0.53 | 0.483 | 6.41E-47 |
| C1orf21 | 6.22E-105 | 0.392523 | 0.605 | 0.499 | 2.36E-100 |
| OSBPL8 | 8.64E-56 | 0.392296 | 0.654 | 0.647 | 3.28E-51 |
| PRDX6 | 1.62E-140 | 0.391772 | 0.884 | 0.904 | 6.15E-136 |
| AAED1 | 1.00E-120 | 0.391719 | 0.508 | 0.379 | 3.80E-116 |
| AL118516. | 2.97E-208 | 0.383192 | 0.326 | 0.135 | 1.13E-203 |
| CXCL14 | 0.003508 | 0.382867 | 0.121 | 0.106 | 1 |
| ZFHX3 | 1.10E-72 | 0.382433 | 0.584 | 0.519 | 4.17E-68 |
| TGFBR2 | 1.76E-56 | 0.38203 | 0.719 | 0.714 | 6.69E-52 |
| STK24 | 1.62E-87 | 0.381948 | 0.485 | 0.389 | 6.15E-83 |
| C17orf58 | 6.68E-29 | 0.381847 | 0.364 | 0.318 | 2.54E-24 |
| PPL | 2.18E-93 | 0.380724 | 0.351 | 0.219 | 8.29E-89 |
| IFITM3 | 8.62E-57 | 0.380617 | 0.983 | 0.991 | 3.27E-52 |
| S100A6 | 2.56E-129 | 0.375963 | 1 | 1 | 9.71E-125 |
| PA2G4 | 2.63E-89 | 0.375822 | 0.712 | 0.703 | 1.00E-84 |
| TREM1 | 5.83E-53 | 0.375768 | 0.307 | 0.213 | 2.22E-48 |
| HRCT1 | 4.28E-104 | 0.375184 | 0.236 | 0.115 | 1.62E-99 |
| IFITM1 | 9.02E-84 | 0.374896 | 0.481 | 0.374 | 3.42E-79 |
| CALCRL | 4.72E-126 | 0.372811 | 0.221 | 0.09 | 1.79E-121 |
| RPL30 | 0 | 0.37152 | 0.999 | 0.999 | 0 |

| | | | | | |
|---|---|---|---|---|---|
| RPL18A | 0 | 0.37144 | 0.998 | 0.999 | 0 |
| MT-ND5 | 2.90E-189 | 0.371328 | 0.997 | 0.999 | 1.10E-184 |
| TBC1D15 | 1.41E-83 | 0.371031 | 0.477 | 0.383 | 5.35E-79 |
| FGFBP2 | 0.445228 | 0.368515 | 0.305 | 0.332 | 1 |
| SOCS1 | 6.69E-17 | 0.368312 | 0.265 | 0.228 | 2.54E-12 |
| ARHGAP6 | 2.64E-186 | 0.368135 | 0.301 | 0.122 | 1.00E-181 |
| MXI1 | 3.45E-102 | 0.366907 | 0.448 | 0.321 | 1.31E-97 |
| RAB32 | 9.09E-83 | 0.366423 | 0.648 | 0.594 | 3.45E-78 |
| POLR1D | 1.52E-96 | 0.365951 | 0.68 | 0.648 | 5.79E-92 |
| SAT1 | 5.13E-24 | 0.365328 | 0.878 | 0.927 | 1.95E-19 |
| TM4SF1 | 8.60E-12 | 0.365242 | 0.287 | 0.26 | 3.27E-07 |
| RPL8 | 5.02E-290 | 0.364174 | 0.997 | 0.999 | 1.91E-285 |
| TNFAIP2 | 0.021883 | 0.363639 | 0.623 | 0.696 | 1 |
| CREBRF | 1.65E-79 | 0.363469 | 0.54 | 0.451 | 6.27E-75 |
| MRPL33 | 1.16E-93 | 0.362945 | 0.724 | 0.71 | 4.42E-89 |
| OSER1 | 1.18E-118 | 0.362641 | 0.407 | 0.27 | 4.48E-114 |
| RPL22L1 | 2.97E-59 | 0.360641 | 0.704 | 0.703 | 1.13E-54 |
| SLC25A6 | 4.47E-118 | 0.360157 | 0.917 | 0.936 | 1.70E-113 |
| EIF4H | 8.68E-109 | 0.359058 | 0.724 | 0.688 | 3.30E-104 |
| MT-ND3 | 1.29E-221 | 0.358924 | 0.999 | 0.999 | 4.89E-217 |
| MED30 | 3.23E-123 | 0.357417 | 0.466 | 0.328 | 1.23E-118 |
| KLF2 | 4.28E-16 | 0.357297 | 0.716 | 0.713 | 1.62E-11 |
| RPS21 | 1.85E-245 | 0.357284 | 0.984 | 0.994 | 7.02E-241 |
| RPL34 | 0 | 0.357228 | 0.998 | 0.999 | 0 |
| BTG1 | 2.88E-87 | 0.354227 | 0.928 | 0.901 | 1.09E-82 |
| FSTL3 | 3.61E-91 | 0.353195 | 0.33 | 0.205 | 1.37E-86 |
| TRIB1 | 6.53E-191 | 0.35291 | 0.254 | 0.088 | 2.48E-186 |
| IGFBP3 | 0.003352 | 0.35051 | 0.138 | 0.128 | 1 |
| HERPUD1 | 3.21E-23 | 0.349134 | 0.782 | 0.838 | 1.22E-18 |
| CELF2 | 3.68E-61 | 0.348829 | 0.606 | 0.537 | 1.40E-56 |
| RPLP0 | 2.34E-115 | 0.346924 | 0.99 | 0.997 | 8.90E-111 |
| FAXDC2 | 7.18E-68 | 0.346034 | 0.356 | 0.254 | 2.73E-63 |
| TRIP10 | 2.33E-92 | 0.345631 | 0.433 | 0.312 | 8.84E-88 |
| GRINA | 5.42E-77 | 0.344803 | 0.573 | 0.515 | 2.06E-72 |
| JUND | 2.46E-91 | 0.344277 | 0.988 | 0.995 | 9.34E-87 |
| PPP2R2A | 7.23E-54 | 0.343672 | 0.474 | 0.419 | 2.74E-49 |
| DHRS7 | 7.88E-77 | 0.343329 | 0.669 | 0.631 | 2.99E-72 |
| PRKAG2 | 1.10E-103 | 0.34316 | 0.368 | 0.232 | 4.17E-99 |
| MPST | 2.62E-61 | 0.342563 | 0.467 | 0.399 | 9.95E-57 |
| TMOD1 | 1.23E-159 | 0.34216 | 0.233 | 0.084 | 4.65E-155 |
| RPL35A | 4.04E-305 | 0.341215 | 0.998 | 0.999 | 1.53E-300 |
| YBX1 | 7.19E-121 | 0.340663 | 0.955 | 0.976 | 2.73E-116 |
| IRX3 | 1.26E-31 | 0.339831 | 0.31 | 0.242 | 4.79E-27 |
| OTUD1 | 4.41E-32 | 0.339789 | 0.295 | 0.238 | 1.67E-27 |
| BLVRB | 2.64E-67 | 0.339467 | 0.663 | 0.648 | 1.00E-62 |
| CCNDBP1 | 2.32E-89 | 0.338887 | 0.514 | 0.413 | 8.81E-85 |
| LEPR | 7.48E-117 | 0.337404 | 0.194 | 0.075 | 2.84E-112 |

| | | | | | |
|---|---|---|---|---|---|
| ALDH6A1 | 1.43E-70 | 0.335877 | 0.329 | 0.229 | 5.44E-66 |
| FTL | 1.22E-90 | 0.33523 | 0.999 | 1 | 4.64E-86 |
| UBA52 | 1.04E-235 | 0.334189 | 0.993 | 0.997 | 3.96E-231 |
| VEGFD | 7.18E-224 | 0.3327 | 0.166 | 0.027 | 2.73E-219 |
| TIMP4 | 1.95E-37 | 0.331931 | 0.278 | 0.21 | 7.42E-33 |
| IRS2 | 1.81E-128 | 0.331541 | 0.28 | 0.136 | 6.87E-124 |
| FADS3 | 2.78E-92 | 0.331253 | 0.349 | 0.224 | 1.06E-87 |
| MGLL | 6.63E-133 | 0.330265 | 0.279 | 0.128 | 2.52E-128 |
| SBDS | 2.54E-79 | 0.329853 | 0.819 | 0.822 | 9.64E-75 |
| AC015912 | 6.03E-292 | 0.329734 | 0.203 | 0.031 | 2.29E-287 |
| MKNK2 | 2.06E-81 | 0.329208 | 0.382 | 0.268 | 7.84E-77 |
| CCDC59 | 2.49E-64 | 0.329052 | 0.54 | 0.49 | 9.44E-60 |
| METRNL | 2.84E-67 | 0.328735 | 0.735 | 0.692 | 1.08E-62 |
| CLK1 | 8.77E-50 | 0.328025 | 0.532 | 0.486 | 3.33E-45 |
| APCDD1 | 2.01E-130 | 0.327915 | 0.15 | 0.042 | 7.64E-126 |
| DYNLT3 | 1.27E-75 | 0.325889 | 0.44 | 0.337 | 4.84E-71 |
| LDHA | 1.16E-52 | 0.324536 | 0.963 | 0.974 | 4.40E-48 |
| CHPT1 | 7.42E-63 | 0.324359 | 0.548 | 0.497 | 2.82E-58 |
| IFNGR1 | 9.52E-62 | 0.323944 | 0.62 | 0.598 | 3.62E-57 |
| ANKRD37 | 8.98E-75 | 0.323454 | 0.321 | 0.215 | 3.41E-70 |
| SCARA5 | 2.19E-46 | 0.323133 | 0.755 | 0.708 | 8.33E-42 |
| ILF3-DT | 3.19E-46 | 0.322405 | 0.45 | 0.391 | 1.21E-41 |
| ODC1 | 1.25E-63 | 0.321083 | 0.362 | 0.272 | 4.74E-59 |
| ENO1 | 5.23E-98 | 0.320721 | 0.931 | 0.946 | 1.99E-93 |
| AC007952 | 2.45E-43 | 0.318076 | 0.355 | 0.272 | 9.31E-39 |
| RNMT | 7.28E-38 | 0.317845 | 0.598 | 0.6 | 2.77E-33 |
| FBXL3 | 1.51E-51 | 0.316391 | 0.475 | 0.421 | 5.73E-47 |
| RPL12 | 1.33E-212 | 0.315127 | 0.998 | 0.999 | 5.04E-208 |
| UXT | 3.30E-76 | 0.315103 | 0.729 | 0.725 | 1.25E-71 |
| TNFAIP3 | 1.33E-29 | 0.314986 | 0.183 | 0.128 | 5.06E-25 |
| FMO2 | 3.76E-19 | 0.313801 | 0.155 | 0.109 | 1.43E-14 |
| SH3BP5 | 1.19E-26 | 0.313226 | 0.761 | 0.807 | 4.52E-22 |
| CDC42EP4 | 3.58E-120 | 0.312564 | 0.314 | 0.167 | 1.36E-115 |
| TANK | 2.05E-65 | 0.312433 | 0.508 | 0.438 | 7.80E-61 |
| PLA2G16 | 2.89E-52 | 0.311209 | 0.474 | 0.4 | 1.10E-47 |
| ADGRG2 | 1.57E-68 | 0.311198 | 0.216 | 0.12 | 5.94E-64 |
| CYBRD1 | 1.38E-62 | 0.310648 | 0.839 | 0.862 | 5.26E-58 |
| NDUFAF2 | 1.34E-48 | 0.310222 | 0.525 | 0.497 | 5.07E-44 |
| RPAIN | 4.52E-63 | 0.309844 | 0.457 | 0.376 | 1.72E-58 |
| SSH2 | 1.85E-69 | 0.30979 | 0.274 | 0.175 | 7.04E-65 |
| FAU | 5.08E-270 | 0.307846 | 0.995 | 0.998 | 1.93E-265 |
| NOX4 | 3.28E-94 | 0.307784 | 0.347 | 0.212 | 1.25E-89 |
| SMIM14 | 5.61E-48 | 0.307366 | 0.772 | 0.797 | 2.13E-43 |
| CCNB1IP1 | 7.73E-58 | 0.306204 | 0.413 | 0.331 | 2.94E-53 |
| TKT | 1.84E-65 | 0.30617 | 0.595 | 0.553 | 6.99E-61 |
| AC245014 | 1.85E-186 | 0.306024 | 0.151 | 0.028 | 7.03E-182 |
| ATP13A3 | 5.29E-64 | 0.303062 | 0.327 | 0.23 | 2.01E-59 |

| | | | | | |
|---|---|---|---|---|---|
| RAB21 | 2.18E-58 | 0.30262 | 0.534 | 0.483 | 8.29E-54 |
| DIO3OS | 3.48E-90 | 0.301709 | 0.254 | 0.132 | 1.32E-85 |
| RRAGC | 5.48E-66 | 0.301308 | 0.41 | 0.32 | 2.08E-61 |
| EGFR | 1.55E-49 | 0.300747 | 0.574 | 0.529 | 5.87E-45 |
| LIF | 4.09E-231 | 0.300655 | 0.114 | 0.006 | 1.55E-226 |
| TMA7 | 3.45E-135 | 0.300363 | 0.926 | 0.951 | 1.31E-130 |
| GMNN | 6.46E-70 | 0.298974 | 0.234 | 0.136 | 2.45E-65 |
| TPI1 | 1.19E-55 | 0.296993 | 0.885 | 0.913 | 4.52E-51 |
| TTC32 | 6.34E-120 | 0.296683 | 0.305 | 0.16 | 2.41E-115 |
| GPC3 | 2.24E-157 | 0.29609 | 0.111 | 0.016 | 8.50E-153 |
| NTRK2 | 7.10E-49 | 0.296062 | 0.385 | 0.286 | 2.70E-44 |
| RPL22 | 2.16E-198 | 0.294893 | 0.991 | 0.995 | 8.21E-194 |
| PCSK5 | 2.21E-18 | 0.29472 | 0.33 | 0.293 | 8.39E-14 |
| LMOD1 | 7.66E-118 | 0.293369 | 0.156 | 0.05 | 2.91E-113 |
| RPL32 | 1.85E-217 | 0.292697 | 0.999 | 0.999 | 7.04E-213 |
| ARMCX1 | 1.51E-45 | 0.291943 | 0.462 | 0.41 | 5.73E-41 |
| CCDC69 | 6.92E-150 | 0.291405 | 0.233 | 0.088 | 2.63E-145 |
| CFL2 | 3.11E-70 | 0.291263 | 0.392 | 0.291 | 1.18E-65 |
| NTM | 3.29E-92 | 0.29119 | 0.246 | 0.125 | 1.25E-87 |
| MAP1B | 2.18E-19 | 0.29068 | 0.355 | 0.3 | 8.26E-15 |
| CD9 | 2.72E-85 | 0.290344 | 0.881 | 0.811 | 1.03E-80 |
| MRPS36 | 2.36E-44 | 0.290117 | 0.543 | 0.516 | 8.95E-40 |
| IMPA2 | 5.74E-117 | 0.289731 | 0.305 | 0.164 | 2.18E-112 |
| AOX1 | 2.24E-77 | 0.28759 | 0.381 | 0.259 | 8.53E-73 |
| CLDND1 | 7.72E-63 | 0.287353 | 0.498 | 0.428 | 2.93E-58 |
| RAD23A | 4.76E-60 | 0.287295 | 0.683 | 0.676 | 1.81E-55 |
| SYF2 | 5.34E-40 | 0.287198 | 0.682 | 0.701 | 2.03E-35 |
| SPRY2 | 5.84E-75 | 0.286107 | 0.274 | 0.166 | 2.22E-70 |
| CERS2 | 5.01E-55 | 0.286044 | 0.584 | 0.546 | 1.90E-50 |
| SLC1A5 | 2.31E-81 | 0.28563 | 0.334 | 0.219 | 8.78E-77 |
| ROMO1 | 2.07E-62 | 0.285624 | 0.723 | 0.718 | 7.86E-58 |
| PLCG2 | 2.83E-111 | 0.284322 | 0.44 | 0.287 | 1.07E-106 |
| GYG1 | 3.42E-56 | 0.283988 | 0.4 | 0.318 | 1.30E-51 |
| PALM | 2.85E-56 | 0.283872 | 0.423 | 0.33 | 1.08E-51 |
| SAMHD1 | 4.99E-11 | 0.283034 | 0.446 | 0.444 | 1.90E-06 |
| MT-CYB | 1.48E-126 | 0.282358 | 1 | 1 | 5.61E-122 |
| UBE2I | 2.55E-71 | 0.281648 | 0.782 | 0.786 | 9.70E-67 |
| CMSS1 | 4.03E-48 | 0.279093 | 0.345 | 0.27 | 1.53E-43 |
| CA12 | 1.43E-18 | 0.278487 | 0.318 | 0.279 | 5.43E-14 |
| HBP1 | 1.66E-41 | 0.278455 | 0.584 | 0.572 | 6.30E-37 |
| PLP2 | 1.55E-49 | 0.278224 | 0.674 | 0.616 | 5.89E-45 |
| SVEP1 | 9.63E-21 | 0.278131 | 0.37 | 0.328 | 3.66E-16 |
| RPS7 | 1.55E-174 | 0.277792 | 0.989 | 0.997 | 5.89E-170 |
| SOX4 | 2.80E-18 | 0.276799 | 0.564 | 0.728 | 1.06E-13 |
| SNRPB | 2.45E-61 | 0.276649 | 0.734 | 0.738 | 9.30E-57 |
| Z93241.1 | 2.95E-126 | 0.276414 | 0.156 | 0.047 | 1.12E-121 |
| PSAP | 1.26E-42 | 0.276045 | 0.973 | 0.994 | 4.78E-38 |

| | | | | | |
|---|---|---|---|---|---|
| SERPINE2 | 3.28E-05 | 0.27513 | 0.219 | 0.266 | 1 |
| MAN1A1 | 0.009543 | 0.274991 | 0.664 | 0.753 | 1 |
| HSPA1A | 0.062479 | 0.274979 | 0.68 | 0.792 | 1 |
| YPEL3 | 5.86E-25 | 0.274074 | 0.677 | 0.729 | 2.22E-20 |
| COQ10B | 5.88E-57 | 0.273628 | 0.477 | 0.406 | 2.23E-52 |
| KRTCAP2 | 7.71E-69 | 0.273201 | 0.745 | 0.736 | 2.93E-64 |
| IER5L | 0.805122 | 0.273195 | 0.53 | 0.634 | 1 |
| AKIRIN2 | 1.95E-63 | 0.273142 | 0.392 | 0.3 | 7.41E-59 |
| C9orf72 | 3.87E-143 | 0.273061 | 0.197 | 0.067 | 1.47E-138 |
| SRSF10 | 1.50E-41 | 0.272972 | 0.717 | 0.734 | 5.71E-37 |
| DLST | 5.20E-58 | 0.272516 | 0.37 | 0.283 | 1.98E-53 |
| TPRG1 | 1.77E-99 | 0.271917 | 0.228 | 0.109 | 6.72E-95 |
| RBM39 | 8.03E-66 | 0.271775 | 0.901 | 0.929 | 3.05E-61 |
| PPM1L | 8.13E-28 | 0.271526 | 0.32 | 0.266 | 3.09E-23 |
| SNW1 | 5.49E-43 | 0.270545 | 0.624 | 0.606 | 2.09E-38 |
| TXLNG | 1.34E-59 | 0.270313 | 0.328 | 0.236 | 5.08E-55 |
| ATP6V1F | 9.54E-33 | 0.269064 | 0.724 | 0.777 | 3.62E-28 |
| HNRNPDL | 3.00E-46 | 0.268239 | 0.835 | 0.872 | 1.14E-41 |
| ITPKC | 4.09E-149 | 0.267669 | 0.235 | 0.09 | 1.55E-144 |
| YPEL2 | 2.01E-50 | 0.26744 | 0.357 | 0.272 | 7.64E-46 |
| SLC29A1 | 1.46E-26 | 0.266903 | 0.372 | 0.319 | 5.54E-22 |
| CYP4X1 | 7.38E-264 | 0.266539 | 0.17 | 0.022 | 2.80E-259 |
| LAMP1 | 6.41E-48 | 0.266424 | 0.849 | 0.9 | 2.43E-43 |
| HES1 | 0.000548 | 0.265449 | 0.22 | 0.205 | 1 |
| TMED5 | 2.97E-40 | 0.265131 | 0.429 | 0.378 | 1.13E-35 |
| TMEM38B | 5.47E-140 | 0.265102 | 0.239 | 0.097 | 2.08E-135 |
| ETS2 | 3.78E-53 | 0.264431 | 0.279 | 0.184 | 1.44E-48 |
| ITM2A | 1.39E-64 | 0.263582 | 0.506 | 0.371 | 5.29E-60 |
| CCT7 | 5.50E-49 | 0.263066 | 0.663 | 0.674 | 2.09E-44 |
| TNS2 | 7.64E-28 | 0.262843 | 0.508 | 0.493 | 2.90E-23 |
| ATP5ME | 2.26E-50 | 0.262533 | 0.796 | 0.821 | 8.60E-46 |
| PRR13 | 2.83E-39 | 0.262306 | 0.539 | 0.52 | 1.08E-34 |
| RBMS3 | 3.92E-21 | 0.261637 | 0.577 | 0.579 | 1.49E-16 |
| COPS2 | 5.58E-35 | 0.260505 | 0.544 | 0.526 | 2.12E-30 |
| TENT5A | 0.000124 | 0.259895 | 0.609 | 0.655 | 1 |
| BRI3 | 8.16E-58 | 0.25985 | 0.897 | 0.921 | 3.10E-53 |
| RNH1 | 8.34E-65 | 0.259176 | 0.893 | 0.914 | 3.17E-60 |
| FBLN5 | 4.91E-26 | 0.259105 | 0.517 | 0.482 | 1.86E-21 |
| MMP24OS | 4.72E-35 | 0.258721 | 0.501 | 0.47 | 1.79E-30 |
| SLC25A5 | 3.61E-26 | 0.258587 | 0.643 | 0.671 | 1.37E-21 |
| TAF9 | 1.74E-31 | 0.258505 | 0.459 | 0.428 | 6.60E-27 |
| LINC01088 | 2.85E-47 | 0.258444 | 0.21 | 0.13 | 1.08E-42 |
| SAV1 | 1.45E-36 | 0.257788 | 0.482 | 0.442 | 5.51E-32 |
| ARL4C | 7.81E-12 | 0.257 | 0.479 | 0.466 | 2.96E-07 |
| PGK1 | 7.01E-11 | 0.25698 | 0.716 | 0.763 | 2.66E-06 |
| CYC1 | 1.21E-49 | 0.256873 | 0.621 | 0.604 | 4.61E-45 |
| USP10 | 5.87E-32 | 0.256813 | 0.31 | 0.254 | 2.23E-27 |

| | | | | | |
|---|---|---|---|---|---|
| SLC2A1 | 1.48E-114 | 0.25664 | 0.188 | 0.072 | 5.62E-110 |
| FST | 5.95E-46 | 0.256576 | 0.13 | 0.065 | 2.26E-41 |
| ARF1 | 4.92E-73 | 0.255919 | 0.865 | 0.873 | 1.87E-68 |
| TMEM109 | 5.16E-39 | 0.255861 | 0.625 | 0.613 | 1.96E-34 |
| VDAC2 | 1.91E-55 | 0.255607 | 0.748 | 0.762 | 7.24E-51 |
| ZC3H12A | 4.03E-88 | 0.255313 | 0.188 | 0.086 | 1.53E-83 |
| SMARCD2 | 2.02E-44 | 0.255013 | 0.406 | 0.338 | 7.66E-40 |
| RPL9 | 1.83E-150 | 0.254959 | 0.997 | 0.999 | 6.94E-146 |
| RBM8A | 1.31E-38 | 0.254812 | 0.697 | 0.744 | 4.99E-34 |
| SMAD3 | 1.81E-28 | 0.254649 | 0.458 | 0.421 | 6.86E-24 |
| CNIH4 | 2.50E-34 | 0.254 | 0.609 | 0.61 | 9.48E-30 |
| SRSF3 | 1.48E-58 | 0.253817 | 0.876 | 0.893 | 5.60E-54 |
| RPL29 | 6.08E-172 | 0.253721 | 0.996 | 0.999 | 2.31E-167 |
| RAB9A | 2.74E-42 | 0.253483 | 0.466 | 0.419 | 1.04E-37 |
| KLHL24 | 7.83E-31 | 0.252804 | 0.382 | 0.332 | 2.97E-26 |
| CLEC2B | 2.55E-30 | 0.252301 | 0.187 | 0.128 | 9.67E-26 |
| SNHG25 | 9.04E-70 | 0.252118 | 0.285 | 0.179 | 3.43E-65 |
| ERO1A | 4.74E-24 | 0.252097 | 0.318 | 0.271 | 1.80E-19 |
| TNFAIP6 | 2.02E-118 | 0.251527 | 0.515 | 0.756 | 7.68E-114 |
| FMNL2 | 2.60E-85 | 0.251399 | 0.266 | 0.15 | 9.88E-81 |
| LRRN4CL | 3.66E-31 | 0.251045 | 0.683 | 0.673 | 1.39E-26 |
| ATF4 | 1.12E-39 | 0.250472 | 0.769 | 0.78 | 4.26E-35 |
| PAMR1 | 1.23E-84 | 0.250141 | 0.23 | 0.115 | 4.69E-80 |
| CTSK | 2.88E-216 | -0.25001 | 0.672 | 0.922 | 1.10E-211 |
| EVA1B | 2.84E-103 | -0.25072 | 0.429 | 0.637 | 1.08E-98 |
| ATP5MC3 | 3.36E-120 | -0.25102 | 0.773 | 0.908 | 1.28E-115 |
| HCG18 | 3.09E-185 | -0.25111 | 0.108 | 0.34 | 1.17E-180 |
| GAMT | 4.11E-223 | -0.25117 | 0.061 | 0.297 | 1.56E-218 |
| SCAF11 | 6.99E-121 | -0.25117 | 0.58 | 0.786 | 2.65E-116 |
| PRELID1 | 1.13E-103 | -0.2516 | 0.66 | 0.825 | 4.28E-99 |
| PRDX1 | 5.76E-104 | -0.25162 | 0.905 | 0.978 | 2.19E-99 |
| SRGAP1 | 8.52E-107 | -0.25165 | 0.299 | 0.51 | 3.24E-102 |
| NRN1 | 3.71E-193 | -0.25181 | 0.06 | 0.275 | 1.41E-188 |
| PDCD6 | 5.81E-135 | -0.25192 | 0.593 | 0.805 | 2.21E-130 |
| RPL3 | 1.91E-99 | -0.25205 | 0.997 | 0.999 | 7.25E-95 |
| NEAT1 | 1.62E-87 | -0.25236 | 0.985 | 0.998 | 6.15E-83 |
| PTK7 | 1.16E-243 | -0.25256 | 0.04 | 0.278 | 4.41E-239 |
| ANKRD11 | 9.34E-107 | -0.25288 | 0.523 | 0.733 | 3.55E-102 |
| DYNC1LI2 | 3.59E-138 | -0.25317 | 0.423 | 0.664 | 1.36E-133 |
| EMC10 | 1.05E-159 | -0.25327 | 0.197 | 0.437 | 3.98E-155 |
| SELENOF | 3.18E-128 | -0.25353 | 0.726 | 0.883 | 1.21E-123 |
| PHKB | 3.22E-183 | -0.25393 | 0.194 | 0.459 | 1.22E-178 |
| BCLAF1 | 3.40E-99 | -0.25402 | 0.564 | 0.76 | 1.29E-94 |
| ERGIC1 | 5.51E-142 | -0.25472 | 0.37 | 0.622 | 2.09E-137 |
| REST | 2.37E-155 | -0.25525 | 0.25 | 0.501 | 9.01E-151 |
| ZDHHC20 | 1.34E-193 | -0.25538 | 0.105 | 0.342 | 5.08E-189 |
| SULT1A1 | 6.93E-190 | -0.25564 | 0.02 | 0.2 | 2.63E-185 |

| | | | | | |
|---|---|---|---|---|---|
| CCDC14 | 4.47E-165 | -0.25575 | 0.212 | 0.467 | 1.70E-160 |
| TGFB3 | 6.12E-215 | -0.25602 | 0.035 | 0.248 | 2.32E-210 |
| CYP27C1 | 7.91E-199 | -0.25612 | 0.042 | 0.246 | 3.00E-194 |
| GALNT11 | 3.59E-160 | -0.2563 | 0.28 | 0.538 | 1.36E-155 |
| EXTL2 | 6.50E-207 | -0.25664 | 0.127 | 0.388 | 2.47E-202 |
| AKAP11 | 1.25E-189 | -0.2568 | 0.161 | 0.419 | 4.75E-185 |
| BARX1 | 3.87E-195 | -0.25695 | 0.038 | 0.24 | 1.47E-190 |
| ICAM2 | 2.26E-213 | -0.25704 | 0.023 | 0.226 | 8.57E-209 |
| FHL3 | 6.88E-181 | -0.25708 | 0.124 | 0.354 | 2.61E-176 |
| TMEM260 | 3.15E-171 | -0.25723 | 0.092 | 0.304 | 1.20E-166 |
| RNASEH2C | 8.97E-127 | -0.25727 | 0.506 | 0.733 | 3.41E-122 |
| SLC2A12 | 3.70E-172 | -0.25746 | 0.018 | 0.183 | 1.40E-167 |
| SH3YL1 | 1.79E-205 | -0.25781 | 0.108 | 0.357 | 6.79E-201 |
| ATP5F1A | 2.44E-122 | -0.25784 | 0.569 | 0.787 | 9.27E-118 |
| BBX | 8.09E-146 | -0.258 | 0.363 | 0.626 | 3.07E-141 |
| SEC61B | 2.44E-124 | -0.25837 | 0.837 | 0.947 | 9.28E-120 |
| ADAMTSL3 | 1.01E-128 | -0.25855 | 0.117 | 0.299 | 3.83E-124 |
| TXLNA | 2.96E-193 | -0.25873 | 0.165 | 0.425 | 1.12E-188 |
| TFG | 1.31E-116 | -0.25897 | 0.557 | 0.751 | 4.98E-112 |
| GALNT5 | 2.42E-251 | -0.25897 | 0.02 | 0.246 | 9.19E-247 |
| CTNNA1 | 2.63E-119 | -0.2591 | 0.565 | 0.764 | 9.99E-115 |
| KDM5A | 6.79E-159 | -0.25922 | 0.253 | 0.51 | 2.58E-154 |
| GUCY1B1 | 7.56E-225 | -0.25932 | 0.011 | 0.209 | 2.87E-220 |
| EVL | 2.97E-147 | -0.25939 | 0.296 | 0.55 | 1.13E-142 |
| PBRM1 | 2.13E-163 | -0.2597 | 0.241 | 0.497 | 8.10E-159 |
| ASH1L | 1.68E-128 | -0.25996 | 0.479 | 0.713 | 6.38E-124 |
| GCSH | 1.60E-132 | -0.26025 | 0.32 | 0.551 | 6.06E-128 |
| ARL2BP | 6.72E-149 | -0.261 | 0.349 | 0.598 | 2.55E-144 |
| CETN2 | 2.09E-186 | -0.26104 | 0.184 | 0.443 | 7.92E-182 |
| PROS1 | 1.91E-115 | -0.26104 | 0.408 | 0.636 | 7.25E-111 |
| TMEM167 | 6.99E-117 | -0.26108 | 0.467 | 0.678 | 2.66E-112 |
| SPCS1 | 1.34E-127 | -0.26111 | 0.779 | 0.92 | 5.10E-123 |
| TGOLN2 | 5.00E-142 | -0.26119 | 0.473 | 0.719 | 1.90E-137 |
| IFI44 | 1.92E-202 | -0.26221 | 0.08 | 0.312 | 7.29E-198 |
| CCDC167 | 3.84E-228 | -0.26232 | 0.094 | 0.35 | 1.46E-223 |
| MORC4 | 5.16E-144 | -0.26248 | 0.278 | 0.522 | 1.96E-139 |
| ERP29 | 6.49E-123 | -0.26275 | 0.639 | 0.841 | 2.46E-118 |
| ITPRIPL2 | 6.39E-152 | -0.26282 | 0.26 | 0.509 | 2.43E-147 |
| SEC24D | 8.66E-163 | -0.26284 | 0.258 | 0.514 | 3.29E-158 |
| SDC1 | 3.43E-202 | -0.26303 | 0.006 | 0.183 | 1.30E-197 |
| SLC2A4RG | 7.55E-139 | -0.2633 | 0.417 | 0.662 | 2.87E-134 |
| THUMPD3 | 1.06E-163 | -0.26346 | 0.203 | 0.451 | 4.04E-159 |
| CYS1 | 5.86E-207 | -0.26402 | 0.043 | 0.256 | 2.23E-202 |
| IKBIP | 1.95E-142 | -0.2642 | 0.39 | 0.639 | 7.42E-138 |
| ZNF608 | 9.58E-197 | -0.26457 | 0.071 | 0.294 | 3.64E-192 |
| HSPA4 | 7.09E-182 | -0.26467 | 0.206 | 0.465 | 2.69E-177 |
| TM9SF2 | 1.08E-154 | -0.26503 | 0.382 | 0.646 | 4.12E-150 |

| | | | | | |
|---|---|---|---|---|---|
| RILP | 6.20E-229 | -0.26504 | 0.075 | 0.324 | 2.36E-224 |
| BAX | 2.46E-111 | -0.26523 | 0.386 | 0.599 | 9.34E-107 |
| SCX | 2.83E-114 | -0.26523 | 0.026 | 0.151 | 1.07E-109 |
| C6orf89 | 1.40E-163 | -0.26626 | 0.294 | 0.554 | 5.31E-159 |
| AP1S1 | 1.91E-189 | -0.26651 | 0.184 | 0.445 | 7.25E-185 |
| REEP3 | 1.31E-129 | -0.26656 | 0.332 | 0.56 | 4.97E-125 |
| TAP1 | 7.64E-151 | -0.26709 | 0.151 | 0.366 | 2.90E-146 |
| MSRB3 | 6.07E-173 | -0.26724 | 0.163 | 0.402 | 2.31E-168 |
| GRN | 6.02E-138 | -0.26747 | 0.648 | 0.866 | 2.29E-133 |
| RPAP2 | 5.70E-164 | -0.26753 | 0.156 | 0.389 | 2.16E-159 |
| H2AFV | 2.88E-149 | -0.26779 | 0.424 | 0.675 | 1.09E-144 |
| SHC1 | 7.38E-187 | -0.26783 | 0.193 | 0.458 | 2.80E-182 |
| FBLN7 | 9.81E-210 | -0.26825 | 0.024 | 0.221 | 3.73E-205 |
| TM9SF3 | 1.02E-128 | -0.26832 | 0.562 | 0.772 | 3.87E-124 |
| IWS1 | 1.34E-176 | -0.26889 | 0.199 | 0.454 | 5.08E-172 |
| PRKCA | 3.51E-243 | -0.26922 | 0.05 | 0.294 | 1.33E-238 |
| SRSF9 | 2.63E-138 | -0.26942 | 0.675 | 0.855 | 9.99E-134 |
| LUZP1 | 1.28E-181 | -0.26952 | 0.234 | 0.506 | 4.87E-177 |
| TP53I3 | 9.87E-168 | -0.26985 | 0.176 | 0.415 | 3.75E-163 |
| CEP290 | 1.26E-171 | -0.27015 | 0.139 | 0.374 | 4.80E-167 |
| CD82 | 5.63E-143 | -0.27017 | 0.16 | 0.375 | 2.14E-138 |
| SMAD5 | 1.10E-179 | -0.27029 | 0.25 | 0.522 | 4.19E-175 |
| USP9X | 6.07E-184 | -0.27044 | 0.185 | 0.442 | 2.30E-179 |
| SEC13 | 2.67E-157 | -0.27099 | 0.291 | 0.536 | 1.02E-152 |
| DNAJC15 | 1.51E-136 | -0.27108 | 0.442 | 0.675 | 5.72E-132 |
| PRKDC | 9.88E-134 | -0.27111 | 0.443 | 0.679 | 3.75E-129 |
| DNM1 | 4.61E-102 | -0.27116 | 0.372 | 0.579 | 1.75E-97 |
| CPXM2 | 2.19E-212 | -0.27121 | 0.023 | 0.225 | 8.32E-208 |
| ACTG1 | 5.49E-130 | -0.27122 | 0.992 | 0.999 | 2.09E-125 |
| HTR2A | 3.96E-207 | -0.27125 | 0.041 | 0.252 | 1.50E-202 |
| SDF4 | 2.64E-137 | -0.27144 | 0.622 | 0.827 | 1.00E-132 |
| FAM20C | 4.89E-106 | -0.27147 | 0.366 | 0.574 | 1.86E-101 |
| BLOC1S1 | 5.20E-138 | -0.27151 | 0.607 | 0.843 | 1.97E-133 |
| ANK3 | 9.70E-168 | -0.27157 | 0.011 | 0.167 | 3.68E-163 |
| ATP6V0D1 | 5.50E-118 | -0.27186 | 0.324 | 0.542 | 2.09E-113 |
| NEDD8 | 1.73E-144 | -0.27218 | 0.772 | 0.921 | 6.57E-140 |
| CAPRIN1 | 2.76E-165 | -0.27221 | 0.311 | 0.57 | 1.05E-160 |
| PDIA5 | 5.63E-231 | -0.27239 | 0.096 | 0.357 | 2.14E-226 |
| BTN3A1 | 2.02E-249 | -0.27263 | 0.035 | 0.271 | 7.69E-245 |
| BEND6 | 1.69E-243 | -0.27265 | 0.045 | 0.285 | 6.42E-239 |
| BRK1 | 1.30E-130 | -0.27273 | 0.746 | 0.899 | 4.94E-126 |
| HNRNPU | 1.45E-128 | -0.2729 | 0.745 | 0.901 | 5.51E-124 |
| SFPQ | 1.34E-119 | -0.27312 | 0.723 | 0.88 | 5.10E-115 |
| KPNB1 | 9.77E-145 | -0.27369 | 0.423 | 0.671 | 3.71E-140 |
| GPR153 | 2.13E-221 | -0.27373 | 0.055 | 0.286 | 8.09E-217 |
| COL5A3 | 1.05E-140 | -0.27402 | 0.083 | 0.267 | 4.01E-136 |
| C11orf24 | 5.69E-213 | -0.27414 | 0.089 | 0.329 | 2.16E-208 |

| | | | | | |
|---|---|---|---|---|---|
| ANAPC5 | 2.41E-137 | -0.27422 | 0.482 | 0.711 | 9.14E-133 |
| LAPTM4A | 4.58E-139 | -0.27461 | 0.978 | 0.996 | 1.74E-134 |
| RPS4X | 9.29E-98 | -0.27496 | 0.998 | 0.999 | 3.53E-93 |
| ACTR2 | 2.16E-132 | -0.27497 | 0.504 | 0.72 | 8.20E-128 |
| DPYD | 9.57E-191 | -0.27544 | 0.164 | 0.419 | 3.63E-186 |
| ATXN1 | 1.28E-225 | -0.27604 | 0.121 | 0.393 | 4.85E-221 |
| ACO1 | 1.63E-143 | -0.27645 | 0.252 | 0.485 | 6.20E-139 |
| FAM89B | 4.71E-139 | -0.27649 | 0.399 | 0.641 | 1.79E-134 |
| FXYD6 | 2.19E-211 | -0.27653 | 0.025 | 0.227 | 8.31E-207 |
| TMEM108 | 5.15E-175 | -0.27655 | 0.065 | 0.265 | 1.95E-170 |
| HGSNAT | 2.75E-183 | -0.27663 | 0.27 | 0.553 | 1.05E-178 |
| ABHD14A | 1.85E-185 | -0.27691 | 0.237 | 0.513 | 7.04E-181 |
| MAPRE2 | 1.73E-219 | -0.27694 | 0.112 | 0.372 | 6.56E-215 |
| LIFR | 1.41E-159 | -0.27703 | 0.153 | 0.383 | 5.34E-155 |
| TP53I13 | 3.06E-177 | -0.27728 | 0.301 | 0.58 | 1.16E-172 |
| MFAP2 | 8.51E-122 | -0.27745 | 0.22 | 0.426 | 3.23E-117 |
| GAP43 | 3.87E-145 | -0.27796 | 0.064 | 0.238 | 1.47E-140 |
| PCSK7 | 8.45E-162 | -0.27801 | 0.221 | 0.466 | 3.21E-157 |
| TXNL4A | 2.49E-140 | -0.27807 | 0.411 | 0.65 | 9.44E-136 |
| ARHGAP18 | 2.96E-186 | -0.27817 | 0.09 | 0.314 | 1.12E-181 |
| CASP1 | 5.11E-254 | -0.2787 | 0.046 | 0.293 | 1.94E-249 |
| PAXX | 9.12E-203 | -0.2788 | 0.182 | 0.453 | 3.46E-198 |
| SGCB | 7.61E-156 | -0.27897 | 0.361 | 0.62 | 2.89E-151 |
| ATP1B1 | 4.53E-180 | -0.27899 | 0.055 | 0.255 | 1.72E-175 |
| MGMT | 4.52E-186 | -0.27902 | 0.34 | 0.635 | 1.72E-181 |
| ADAMTS1 | 5.96E-38 | -0.27924 | 0.238 | 0.346 | 2.26E-33 |
| CREB5 | 8.58E-79 | -0.27926 | 0.655 | 0.807 | 3.26E-74 |
| ANO1 | 1.96E-223 | -0.27948 | 0.005 | 0.196 | 7.45E-219 |
| ARPC5 | 7.35E-125 | -0.27949 | 0.629 | 0.812 | 2.79E-120 |
| PIGT | 1.20E-136 | -0.27971 | 0.553 | 0.772 | 4.55E-132 |
| BAZ1B | 1.62E-182 | -0.28014 | 0.267 | 0.542 | 6.14E-178 |
| EPHA3 | 4.88E-222 | -0.2803 | 0.036 | 0.256 | 1.85E-217 |
| RARG | 2.24E-202 | -0.28046 | 0.127 | 0.383 | 8.51E-198 |
| PAPLN | 1.41E-234 | -0.28066 | 0.047 | 0.282 | 5.34E-230 |
| NBN | 5.87E-209 | -0.2807 | 0.152 | 0.418 | 2.23E-204 |
| CSAD | 2.78E-174 | -0.28081 | 0.178 | 0.427 | 1.06E-169 |
| ATP2A2 | 3.38E-174 | -0.28096 | 0.23 | 0.489 | 1.28E-169 |
| ALDH9A1 | 7.47E-155 | -0.28103 | 0.307 | 0.56 | 2.84E-150 |
| PTPRF | 1.72E-232 | -0.28109 | 0.063 | 0.31 | 6.53E-228 |
| ABRACL | 1.20E-194 | -0.28138 | 0.156 | 0.41 | 4.56E-190 |
| CMBL | 8.19E-169 | -0.28243 | 0.186 | 0.433 | 3.11E-164 |
| PIEZO2 | 1.72E-179 | -0.28292 | 0.039 | 0.227 | 6.55E-175 |
| LMAN1 | 3.55E-135 | -0.28299 | 0.612 | 0.809 | 1.35E-130 |
| TMOD3 | 4.01E-164 | -0.28349 | 0.395 | 0.656 | 1.52E-159 |
| GTF2H5 | 1.41E-160 | -0.28361 | 0.481 | 0.735 | 5.36E-156 |
| PJA2 | 5.99E-139 | -0.2845 | 0.553 | 0.775 | 2.27E-134 |
| SKIL | 1.51E-154 | -0.28514 | 0.272 | 0.522 | 5.75E-150 |

| | | | | | |
|---|---|---|---|---|---|
| ATP5MC1 | 1.74E-141 | -0.28514 | 0.606 | 0.824 | 6.59E-137 |
| ARHGEF12 | 2.74E-180 | -0.28548 | 0.256 | 0.526 | 1.04E-175 |
| ZNF652 | 1.59E-222 | -0.28558 | 0.15 | 0.43 | 6.03E-218 |
| EIF3A | 8.83E-129 | -0.28594 | 0.539 | 0.748 | 3.35E-124 |
| LINC00632 | 1.22E-186 | -0.28619 | 0.088 | 0.309 | 4.64E-182 |
| GBP3 | 3.81E-220 | -0.28633 | 0.071 | 0.31 | 1.45E-215 |
| COPA | 1.16E-163 | -0.28645 | 0.401 | 0.66 | 4.42E-159 |
| PSMD1 | 4.77E-172 | -0.2865 | 0.301 | 0.566 | 1.81E-167 |
| ADIRF | 1.84E-15 | -0.28681 | 0.833 | 0.826 | 6.99E-11 |
| ADD1 | 1.45E-163 | -0.28696 | 0.397 | 0.664 | 5.52E-159 |
| TIMM8B | 3.00E-167 | -0.2875 | 0.452 | 0.714 | 1.14E-162 |
| SEC61A1 | 1.24E-165 | -0.28755 | 0.31 | 0.568 | 4.69E-161 |
| LRRFIP2 | 2.54E-130 | -0.28761 | 0.32 | 0.548 | 9.64E-126 |
| PLEKHA5 | 7.15E-145 | -0.28797 | 0.29 | 0.535 | 2.72E-140 |
| VGLL4 | 5.71E-132 | -0.28828 | 0.452 | 0.69 | 2.17E-127 |
| NCOA4 | 1.96E-195 | -0.28831 | 0.238 | 0.511 | 7.44E-191 |
| CD99 | 2.96E-84 | -0.28842 | 0.937 | 0.981 | 1.12E-79 |
| ARPC1A | 1.46E-180 | -0.28853 | 0.322 | 0.598 | 5.54E-176 |
| IFIT2 | 2.28E-142 | -0.2894 | 0.053 | 0.221 | 8.66E-138 |
| EPRS | 1.25E-162 | -0.28954 | 0.37 | 0.625 | 4.74E-158 |
| NCL | 2.58E-137 | -0.28958 | 0.787 | 0.932 | 9.79E-133 |
| OLFM2 | 6.01E-172 | -0.28982 | 0.008 | 0.163 | 2.28E-167 |
| ROCK1 | 3.54E-139 | -0.28992 | 0.453 | 0.688 | 1.34E-134 |
| BCL7C | 4.68E-163 | -0.29003 | 0.441 | 0.702 | 1.78E-158 |
| MYDGF | 5.86E-140 | -0.2902 | 0.764 | 0.903 | 2.23E-135 |
| ERGIC3 | 1.16E-155 | -0.2905 | 0.452 | 0.698 | 4.39E-151 |
| HACD3 | 3.32E-202 | -0.29053 | 0.23 | 0.511 | 1.26E-197 |
| SHPRH | 1.37E-170 | -0.2913 | 0.203 | 0.453 | 5.20E-166 |
| SARAF | 3.53E-137 | -0.2924 | 0.699 | 0.873 | 1.34E-132 |
| CTNNB1 | 9.77E-133 | -0.29242 | 0.479 | 0.701 | 3.71E-128 |
| NDUFB3 | 1.96E-189 | -0.29246 | 0.418 | 0.693 | 7.44E-185 |
| OST4 | 6.82E-147 | -0.29312 | 0.874 | 0.96 | 2.59E-142 |
| ATM | 1.09E-201 | -0.29315 | 0.222 | 0.505 | 4.13E-197 |
| HES4 | 5.24E-169 | -0.29319 | 0.026 | 0.194 | 1.99E-164 |
| UQCC2 | 3.67E-191 | -0.29331 | 0.262 | 0.538 | 1.39E-186 |
| STK17B | 2.02E-175 | -0.29342 | 0.119 | 0.349 | 7.66E-171 |
| IQSEC1 | 5.12E-228 | -0.29423 | 0.118 | 0.388 | 1.94E-223 |
| WARS | 6.58E-99 | -0.29428 | 0.137 | 0.297 | 2.50E-94 |
| ROBO1 | 1.46E-219 | -0.29428 | 0.151 | 0.428 | 5.53E-215 |
| GBP4 | 2.46E-189 | -0.2943 | 0.02 | 0.2 | 9.34E-185 |
| LHFPL2 | 3.17E-177 | -0.29452 | 0.17 | 0.412 | 1.21E-172 |
| TNS1 | 3.41E-189 | -0.29468 | 0.196 | 0.46 | 1.29E-184 |
| HSPA2 | 3.26E-210 | -0.29494 | 0.072 | 0.305 | 1.24E-205 |
| PARK7 | 2.77E-163 | -0.29662 | 0.862 | 0.962 | 1.05E-158 |
| SEC23A | 2.63E-174 | -0.29671 | 0.309 | 0.573 | 9.98E-170 |
| YWHAZ | 6.50E-126 | -0.297 | 0.59 | 0.784 | 2.47E-121 |
| SPON1 | 2.19E-199 | -0.29717 | 0.021 | 0.209 | 8.33E-195 |

| | | | | | |
|---|---|---|---|---|---|
| SNRNP200 | 1.57E-191 | -0.29765 | 0.271 | 0.556 | 5.96E-187 |
| MAGEH1 | 6.59E-214 | -0.2978 | 0.224 | 0.513 | 2.50E-209 |
| NHLRC3 | 1.45E-194 | -0.29907 | 0.228 | 0.506 | 5.51E-190 |
| PDGFA | 1.99E-187 | -0.29914 | 0.066 | 0.276 | 7.55E-183 |
| B4GALT2 | 5.06E-232 | -0.29916 | 0.142 | 0.418 | 1.92E-227 |
| TTC14 | 7.45E-178 | -0.29939 | 0.366 | 0.646 | 2.83E-173 |
| NDN | 1.46E-167 | -0.29965 | 0.518 | 0.781 | 5.55E-163 |
| JTB | 2.65E-142 | -0.29985 | 0.678 | 0.85 | 1.00E-137 |
| TOP2B | 4.06E-209 | -0.29993 | 0.232 | 0.521 | 1.54E-204 |
| PYCARD | 4.53E-225 | -0.30012 | 0.146 | 0.428 | 1.72E-220 |
| KYNU | 5.03E-168 | -0.3002 | 0.056 | 0.245 | 1.91E-163 |
| SUB1 | 1.87E-145 | -0.30026 | 0.736 | 0.888 | 7.09E-141 |
| EIF4G3 | 7.96E-214 | -0.30143 | 0.21 | 0.495 | 3.02E-209 |
| ASAP2 | 7.20E-164 | -0.30148 | 0.263 | 0.518 | 2.73E-159 |
| GLS | 2.36E-165 | -0.30161 | 0.34 | 0.599 | 8.96E-161 |
| COX6C | 1.91E-156 | -0.30194 | 0.897 | 0.979 | 7.24E-152 |
| CAMK1D | 2.35E-130 | -0.30199 | 0.274 | 0.492 | 8.93E-126 |
| NDUFS8 | 1.10E-176 | -0.30201 | 0.423 | 0.689 | 4.20E-172 |
| MLF2 | 5.74E-172 | -0.30263 | 0.469 | 0.727 | 2.18E-167 |
| CDKN2A | 1.83E-171 | -0.30298 | 0.034 | 0.211 | 6.95E-167 |
| CCDC186 | 7.28E-187 | -0.30361 | 0.259 | 0.54 | 2.77E-182 |
| PRUNE2 | 2.32E-170 | -0.30452 | 0.124 | 0.348 | 8.81E-166 |
| S1PR3 | 7.28E-155 | -0.305 | 0.133 | 0.345 | 2.76E-150 |
| NDFIP1 | 3.23E-86 | -0.30503 | 0.737 | 0.869 | 1.23E-81 |
| JCHAIN | 7.45E-245 | -0.30521 | 0 | 0.201 | 2.83E-240 |
| TRIM22 | 2.63E-160 | -0.30533 | 0.216 | 0.451 | 9.98E-156 |
| VWA1 | 1.27E-241 | -0.30539 | 0.104 | 0.384 | 4.84E-237 |
| JPX | 6.01E-181 | -0.30557 | 0.315 | 0.591 | 2.28E-176 |
| SEMA3A | 4.79E-210 | -0.30561 | 0.016 | 0.208 | 1.82E-205 |
| STMP1 | 6.51E-153 | -0.30562 | 0.539 | 0.761 | 2.47E-148 |
| RAPH1 | 1.60E-213 | -0.30583 | 0.096 | 0.341 | 6.09E-209 |
| DIO2 | 3.27E-75 | -0.30726 | 0.125 | 0.259 | 1.24E-70 |
| SNED1 | 2.74E-113 | -0.30732 | 0.451 | 0.662 | 1.04E-108 |
| ADAM12 | 1.21E-209 | -0.30751 | 0.014 | 0.203 | 4.58E-205 |
| TCEAL3 | 2.09E-201 | -0.30756 | 0.256 | 0.544 | 7.93E-197 |
| SCARB2 | 4.01E-162 | -0.30759 | 0.543 | 0.778 | 1.52E-157 |
| SMC4 | 7.89E-233 | -0.30852 | 0.083 | 0.336 | 3.00E-228 |
| GOLGA4 | 2.22E-144 | -0.30876 | 0.588 | 0.802 | 8.42E-140 |
| CXCL16 | 3.21E-247 | -0.30893 | 0.104 | 0.377 | 1.22E-242 |
| LDLRAD4 | 2.30E-263 | -0.30898 | 0.04 | 0.29 | 8.73E-259 |
| CREB3L2 | 5.81E-196 | -0.30965 | 0.28 | 0.569 | 2.21E-191 |
| DDX46 | 3.48E-180 | -0.31029 | 0.474 | 0.745 | 1.32E-175 |
| TMEM230 | 1.75E-163 | -0.31048 | 0.629 | 0.835 | 6.64E-159 |
| S100A16 | 3.88E-148 | -0.31068 | 0.502 | 0.735 | 1.47E-143 |
| PHF14 | 1.35E-188 | -0.31086 | 0.384 | 0.67 | 5.13E-184 |
| GSTM3 | 1.59E-185 | -0.31197 | 0.38 | 0.661 | 6.05E-181 |
| DLX3 | 1.69E-230 | -0.31213 | 0.035 | 0.259 | 6.43E-226 |

| | | | | | |
|---|---|---|---|---|---|
| TSHZ2 | 1.89E-103 | -0.31215 | 0.556 | 0.722 | 7.19E-99 |
| FTO | 8.40E-229 | -0.31231 | 0.135 | 0.407 | 3.19E-224 |
| GCC2 | 6.17E-138 | -0.31312 | 0.519 | 0.751 | 2.34E-133 |
| TMED10 | 7.56E-158 | -0.31337 | 0.712 | 0.879 | 2.87E-153 |
| SPON2 | 2.45E-63 | -0.31377 | 0.466 | 0.633 | 9.30E-59 |
| PNISR | 7.74E-145 | -0.31379 | 0.852 | 0.965 | 2.94E-140 |
| 11-Sep | 2.70E-169 | -0.31383 | 0.404 | 0.664 | 1.02E-164 |
| SNRPN | 2.57E-194 | -0.31402 | 0.319 | 0.606 | 9.75E-190 |
| PLOD1 | 9.31E-272 | -0.31412 | 0.111 | 0.407 | 3.54E-267 |
| TCEAL4 | 3.86E-174 | -0.31475 | 0.581 | 0.827 | 1.46E-169 |
| HDDC2 | 4.54E-172 | -0.31557 | 0.409 | 0.671 | 1.72E-167 |
| PRRC2C | 3.39E-169 | -0.3159 | 0.716 | 0.891 | 1.29E-164 |
| GNG12 | 3.40E-153 | -0.31705 | 0.486 | 0.731 | 1.29E-148 |
| EXOC7 | 1.41E-227 | -0.31707 | 0.2 | 0.493 | 5.35E-223 |
| C14orf132 | 8.16E-222 | -0.31735 | 0.167 | 0.449 | 3.10E-217 |
| SLC40A1 | 1.13E-170 | -0.31741 | 0.072 | 0.273 | 4.29E-166 |
| P3H4 | 5.55E-254 | -0.31755 | 0.122 | 0.408 | 2.11E-249 |
| SEC63 | 7.45E-182 | -0.31772 | 0.445 | 0.714 | 2.83E-177 |
| DLC1 | 4.13E-166 | -0.31845 | 0.441 | 0.709 | 1.57E-161 |
| PARP9 | 1.59E-245 | -0.3186 | 0.104 | 0.376 | 6.03E-241 |
| ARL1 | 4.96E-187 | -0.31899 | 0.48 | 0.734 | 1.88E-182 |
| TUBB4B | 3.96E-133 | -0.31936 | 0.519 | 0.757 | 1.50E-128 |
| FABP5 | 1.46E-179 | -0.32026 | 0.02 | 0.192 | 5.54E-175 |
| CKAP4 | 2.55E-172 | -0.32046 | 0.333 | 0.587 | 9.68E-168 |
| GPSM2 | 3.63E-153 | -0.32083 | 0.273 | 0.516 | 1.38E-148 |
| PCM1 | 6.61E-170 | -0.32091 | 0.492 | 0.748 | 2.51E-165 |
| KAZALD1 | 2.14E-248 | -0.32106 | 0.076 | 0.339 | 8.12E-244 |
| VCL | 6.97E-145 | -0.32116 | 0.32 | 0.55 | 2.65E-140 |
| APOL2 | 2.69E-219 | -0.32131 | 0.121 | 0.383 | 1.02E-214 |
| FNDC3A | 3.00E-208 | -0.32135 | 0.275 | 0.565 | 1.14E-203 |
| SYNJ2 | 6.38E-264 | -0.32185 | 0.053 | 0.312 | 2.42E-259 |
| CUX1 | 3.34E-184 | -0.32186 | 0.315 | 0.59 | 1.27E-179 |
| COPB1 | 2.15E-189 | -0.32278 | 0.397 | 0.668 | 8.15E-185 |
| ATP5IF1 | 1.94E-176 | -0.32292 | 0.595 | 0.842 | 7.35E-172 |
| OAF | 3.73E-115 | -0.32308 | 0.234 | 0.43 | 1.42E-110 |
| RPN2 | 3.74E-176 | -0.32396 | 0.627 | 0.847 | 1.42E-171 |
| GAA | 6.50E-199 | -0.32418 | 0.329 | 0.621 | 2.47E-194 |
| HAPLN3 | 8.41E-203 | -0.32439 | 0.068 | 0.288 | 3.19E-198 |
| MT-CO1 | 3.92E-185 | -0.32479 | 0.999 | 1 | 1.49E-180 |
| RSRP1 | 3.75E-165 | -0.32485 | 0.587 | 0.829 | 1.42E-160 |
| MAF | 4.09E-180 | -0.32487 | 0.238 | 0.502 | 1.55E-175 |
| SPATS2L | 2.71E-153 | -0.32507 | 0.411 | 0.653 | 1.03E-148 |
| ARHGEF40 | 1.78E-281 | -0.32507 | 0.083 | 0.372 | 6.77E-277 |
| PLXNC1 | 4.81E-272 | -0.32519 | 0.036 | 0.288 | 1.83E-267 |
| ABI2 | 7.68E-231 | -0.32569 | 0.209 | 0.51 | 2.92E-226 |
| ARMCX3 | 9.38E-188 | -0.32659 | 0.384 | 0.664 | 3.56E-183 |
| RCN1 | 4.96E-135 | -0.3267 | 0.571 | 0.766 | 1.88E-130 |

| | | | | | |
|---|---|---|---|---|---|
| PLXDC2 | 9.15E-134 | -0.32682 | 0.48 | 0.721 | 3.47E-129 |
| NDUFA11 | 4.58E-198 | -0.32687 | 0.717 | 0.908 | 1.74E-193 |
| SLC16A4 | 3.05E-255 | -0.327 | 0.131 | 0.425 | 1.16E-250 |
| MARVELD: | 4.41E-191 | -0.32739 | 0.259 | 0.533 | 1.68E-186 |
| MAP1A | 5.91E-152 | -0.3277 | 0.32 | 0.57 | 2.25E-147 |
| DRAP1 | 4.22E-180 | -0.32814 | 0.62 | 0.834 | 1.60E-175 |
| WASF2 | 1.81E-176 | -0.32825 | 0.64 | 0.87 | 6.88E-172 |
| ACSL3 | 3.21E-196 | -0.32832 | 0.302 | 0.586 | 1.22E-191 |
| CRTAP | 2.98E-170 | -0.32855 | 0.493 | 0.735 | 1.13E-165 |
| ARF4 | 6.87E-156 | -0.32868 | 0.682 | 0.847 | 2.61E-151 |
| ZCCHC24 | 1.43E-224 | -0.33072 | 0.218 | 0.51 | 5.44E-220 |
| FKBP1A | 5.01E-172 | -0.33091 | 0.681 | 0.867 | 1.90E-167 |
| MAPK10 | 5.25E-215 | -0.33188 | 0.056 | 0.284 | 1.99E-210 |
| CTSH | 5.65E-107 | -0.33195 | 0.221 | 0.412 | 2.15E-102 |
| GOLGA3 | 5.10E-254 | -0.33221 | 0.173 | 0.478 | 1.94E-249 |
| ANXA11 | 7.45E-195 | -0.33307 | 0.54 | 0.806 | 2.83E-190 |
| LDB2 | 2.38E-243 | -0.33339 | 0.065 | 0.316 | 9.03E-239 |
| RPN1 | 1.41E-201 | -0.33371 | 0.393 | 0.674 | 5.36E-197 |
| REX1BD | 1.34E-188 | -0.33391 | 0.521 | 0.79 | 5.09E-184 |
| IFI35 | 4.02E-204 | -0.33402 | 0.255 | 0.531 | 1.53E-199 |
| PXDN | 8.10E-162 | -0.33536 | 0.236 | 0.473 | 3.08E-157 |
| PSMA2 | 3.05E-205 | -0.33578 | 0.399 | 0.676 | 1.16E-200 |
| CHD9 | 1.14E-174 | -0.33594 | 0.589 | 0.835 | 4.34E-170 |
| KIAA1324L | 1.51E-297 | -0.33638 | 0.045 | 0.323 | 5.73E-293 |
| CRELD2 | 1.20E-224 | -0.33737 | 0.284 | 0.583 | 4.56E-220 |
| DNAJB1 | 1.51E-57 | -0.33753 | 0.391 | 0.535 | 5.72E-53 |
| PALMD | 1.84E-195 | -0.33847 | 0.286 | 0.574 | 6.99E-191 |
| SUMO2 | 3.51E-239 | -0.34105 | 0.918 | 0.981 | 1.33E-234 |
| NFIA | 2.78E-144 | -0.34106 | 0.583 | 0.807 | 1.06E-139 |
| EPSTI1 | 4.03E-278 | -0.3412 | 0.016 | 0.258 | 1.53E-273 |
| SCPEP1 | 7.20E-164 | -0.34148 | 0.553 | 0.789 | 2.73E-159 |
| NUCB1 | 1.46E-185 | -0.34197 | 0.652 | 0.863 | 5.55E-181 |
| RUNX1 | 3.79E-142 | -0.34279 | 0.385 | 0.62 | 1.44E-137 |
| EDF1 | 7.75E-210 | -0.34312 | 0.885 | 0.976 | 2.94E-205 |
| ANXA2 | 4.51E-238 | -0.3432 | 0.963 | 0.996 | 1.71E-233 |
| SETX | 1.00E-240 | -0.3436 | 0.247 | 0.561 | 3.80E-236 |
| TCAF1 | 9.45E-256 | -0.34362 | 0.145 | 0.443 | 3.59E-251 |
| ENPP1 | 6.80E-107 | -0.34389 | 0.241 | 0.435 | 2.58E-102 |
| STAT3 | 1.57E-142 | -0.34395 | 0.67 | 0.843 | 5.95E-138 |
| GABPB1-A | 5.46E-158 | -0.34481 | 0.326 | 0.577 | 2.07E-153 |
| HTATSF1 | 7.99E-242 | -0.34501 | 0.228 | 0.537 | 3.03E-237 |
| MAN1A2 | 7.18E-241 | -0.34559 | 0.247 | 0.555 | 2.73E-236 |
| NDUFA4 | 5.85E-218 | -0.34563 | 0.868 | 0.968 | 2.22E-213 |
| WISP1 | 7.53E-247 | -0.3465 | 0.024 | 0.248 | 2.86E-242 |
| ZNF37A | 3.42E-244 | -0.34657 | 0.163 | 0.457 | 1.30E-239 |
| CXXC5 | 5.12E-130 | -0.34666 | 0.335 | 0.552 | 1.95E-125 |
| ADAR | 1.18E-215 | -0.34729 | 0.322 | 0.617 | 4.50E-211 |

| | | | | | |
|---|---|---|---|---|---|
| PCNX4 | 5.13E-243 | -0.34841 | 0.118 | 0.394 | 1.95E-238 |
| EID1 | 3.77E-255 | -0.34867 | 0.919 | 0.985 | 1.43E-250 |
| GTF3C6 | 1.65E-241 | -0.3492 | 0.322 | 0.628 | 6.28E-237 |
| MYH10 | 2.41E-280 | -0.34924 | 0.065 | 0.343 | 9.16E-276 |
| AKR1C1 | 2.36E-123 | -0.3496 | 0.485 | 0.703 | 8.95E-119 |
| MSRB2 | 1.92E-237 | -0.3497 | 0.352 | 0.659 | 7.30E-233 |
| SELENBP1 | 9.65E-236 | -0.35007 | 0.246 | 0.568 | 3.66E-231 |
| DAAM1 | 9.97E-221 | -0.35012 | 0.191 | 0.477 | 3.79E-216 |
| COL4A2 | 1.33E-170 | -0.35031 | 0.319 | 0.582 | 5.07E-166 |
| BANF1 | 1.85E-207 | -0.35102 | 0.55 | 0.803 | 7.02E-203 |
| NMI | 3.37E-286 | -0.35109 | 0.117 | 0.42 | 1.28E-281 |
| DKK3 | 2.48E-12 | -0.35138 | 0.348 | 0.403 | 9.43E-08 |
| PNN | 5.98E-193 | -0.35162 | 0.585 | 0.824 | 2.27E-188 |
| CHD3 | 7.27E-249 | -0.35198 | 0.169 | 0.464 | 2.76E-244 |
| KCNMA1 | 2.11E-90 | -0.35199 | 0.152 | 0.308 | 8.01E-86 |
| KIAA1217 | 1.00E-299 | -0.35204 | 0.057 | 0.341 | 3.80E-295 |
| SCP2 | 1.29E-201 | -0.35209 | 0.635 | 0.856 | 4.91E-197 |
| NT5DC2 | 3.07E-225 | -0.35216 | 0.095 | 0.342 | 1.17E-220 |
| SGK1 | 1.37E-199 | -0.35234 | 0.142 | 0.396 | 5.20E-195 |
| ABI3BP | 3.55E-177 | -0.35248 | 0.794 | 0.953 | 1.35E-172 |
| ANXA6 | 4.36E-233 | -0.35284 | 0.399 | 0.71 | 1.66E-228 |
| TUBB6 | 1.23E-151 | -0.3532 | 0.381 | 0.609 | 4.66E-147 |
| CBX5 | 4.31E-220 | -0.35348 | 0.335 | 0.632 | 1.64E-215 |
| MBTPS1 | 2.12E-238 | -0.35411 | 0.357 | 0.668 | 8.04E-234 |
| EIF4G1 | 6.01E-222 | -0.35515 | 0.331 | 0.62 | 2.28E-217 |
| CHD4 | 2.01E-205 | -0.35543 | 0.32 | 0.608 | 7.64E-201 |
| PER3 | 4.32E-220 | -0.35551 | 0.177 | 0.452 | 1.64E-215 |
| FKBP10 | 1.84E-190 | -0.35562 | 0.551 | 0.783 | 6.98E-186 |
| GOLGA2 | 8.96E-224 | -0.35567 | 0.373 | 0.674 | 3.40E-219 |
| FGF10 | 1.04E-244 | -0.35569 | 0.149 | 0.447 | 3.96E-240 |
| TRIM56 | 1.26E-214 | -0.35598 | 0.234 | 0.523 | 4.79E-210 |
| TMSB10 | 3.62E-211 | -0.35607 | 0.977 | 0.999 | 1.37E-206 |
| TRIP11 | 1.36E-198 | -0.35638 | 0.389 | 0.67 | 5.18E-194 |
| TMED9 | 6.26E-197 | -0.35646 | 0.724 | 0.886 | 2.38E-192 |
| KDELR1 | 1.90E-205 | -0.35779 | 0.713 | 0.897 | 7.23E-201 |
| RBMS1 | 7.70E-152 | -0.35779 | 0.6 | 0.808 | 2.92E-147 |
| AC020916 | 7.61E-128 | -0.35963 | 0.17 | 0.367 | 2.89E-123 |
| TCEA3 | 6.91E-265 | -0.35967 | 0.137 | 0.436 | 2.62E-260 |
| ZC3HAV1 | 1.41E-288 | -0.35985 | 0.102 | 0.405 | 5.35E-284 |
| YIF1A | 1.07E-193 | -0.36001 | 0.452 | 0.704 | 4.07E-189 |
| DNAJA1 | 5.77E-175 | -0.36012 | 0.652 | 0.859 | 2.19E-170 |
| CFL1 | 1.26E-163 | -0.36091 | 0.874 | 0.961 | 4.78E-159 |
| CHSY1 | 8.20E-265 | -0.36103 | 0.132 | 0.425 | 3.11E-260 |
| ADD3 | 2.66E-163 | -0.36116 | 0.576 | 0.816 | 1.01E-158 |
| SIL1 | 4.69E-232 | -0.36148 | 0.324 | 0.627 | 1.78E-227 |
| COL16A1 | 3.07E-198 | -0.36233 | 0.299 | 0.579 | 1.17E-193 |
| GSTK1 | 1.35E-229 | -0.36236 | 0.391 | 0.693 | 5.12E-225 |

| | | | | | |
|---|---|---|---|---|---|
| PLEKHA4 | 3.61E-247 | -0.36265 | 0.195 | 0.495 | 1.37E-242 |
| DNAJC10 | 7.46E-256 | -0.36438 | 0.259 | 0.575 | 2.83E-251 |
| AP2S1 | 2.46E-199 | -0.36452 | 0.593 | 0.814 | 9.36E-195 |
| GUCY1A1 | 6.84E-283 | -0.36533 | 0.005 | 0.238 | 2.60E-278 |
| DBP | 1.53E-215 | -0.36553 | 0.12 | 0.374 | 5.82E-211 |
| ARHGAP28 | 0 | -0.3661 | 0.048 | 0.333 | 0 |
| PLXNB2 | 4.79E-303 | -0.36626 | 0.138 | 0.467 | 1.82E-298 |
| TLN1 | 1.65E-201 | -0.36658 | 0.538 | 0.78 | 6.25E-197 |
| LRP1 | 5.57E-206 | -0.36717 | 0.923 | 0.99 | 2.11E-201 |
| FKBP7 | 1.86E-246 | -0.36734 | 0.318 | 0.627 | 7.08E-242 |
| CDC42BPA | 4.08E-254 | -0.36815 | 0.221 | 0.533 | 1.55E-249 |
| GLIS3 | 2.63E-230 | -0.36875 | 0.134 | 0.401 | 9.99E-226 |
| CBX6 | 9.70E-278 | -0.36971 | 0.209 | 0.54 | 3.68E-273 |
| PSME1 | 1.11E-168 | -0.3705 | 0.679 | 0.881 | 4.21E-164 |
| CTSF | 4.70E-193 | -0.37064 | 0.624 | 0.879 | 1.79E-188 |
| NKTR | 1.26E-197 | -0.37079 | 0.537 | 0.795 | 4.79E-193 |
| LXN | 8.71E-153 | -0.37109 | 0.212 | 0.438 | 3.31E-148 |
| RBFOX2 | 4.24E-247 | -0.37163 | 0.334 | 0.655 | 1.61E-242 |
| RSPO3 | 2.27E-115 | -0.37164 | 0.283 | 0.488 | 8.64E-111 |
| LIMS1 | 5.22E-179 | -0.37246 | 0.58 | 0.798 | 1.98E-174 |
| HLA-DQB1 | 2.30E-245 | -0.37267 | 0.011 | 0.225 | 8.73E-241 |
| 7-Sep | 1.66E-219 | -0.37278 | 0.827 | 0.947 | 6.32E-215 |
| CHST2 | 1.20E-227 | -0.37288 | 0.073 | 0.312 | 4.57E-223 |
| GNAS | 2.07E-290 | -0.37322 | 0.957 | 0.994 | 7.86E-286 |
| CNN2 | 3.98E-269 | -0.37352 | 0.159 | 0.466 | 1.51E-264 |
| GMDS | 1.07E-220 | -0.37425 | 0.176 | 0.455 | 4.06E-216 |
| ARHGAP21 | 1.91E-201 | -0.37466 | 0.372 | 0.649 | 7.27E-197 |
| TPR | 1.47E-205 | -0.37537 | 0.579 | 0.829 | 5.58E-201 |
| BICC1 | 3.97E-192 | -0.37554 | 0.367 | 0.639 | 1.51E-187 |
| BNC2 | 1.44E-246 | -0.37574 | 0.147 | 0.43 | 5.49E-242 |
| TMCO3 | 4.77E-289 | -0.37596 | 0.213 | 0.546 | 1.81E-284 |
| BPTF | 3.63E-216 | -0.377 | 0.438 | 0.724 | 1.38E-211 |
| ZNF428 | 5.91E-254 | -0.37741 | 0.415 | 0.718 | 2.24E-249 |
| ACTB | 3.39E-148 | -0.37787 | 0.992 | 0.999 | 1.29E-143 |
| PDE1A | 1.08E-306 | -0.37831 | 0.097 | 0.416 | 4.12E-302 |
| PHLDA3 | 1.24E-210 | -0.37883 | 0.291 | 0.578 | 4.72E-206 |
| SAMD9 | 1.66E-303 | -0.3795 | 0.071 | 0.368 | 6.30E-299 |
| UBE2E3 | 2.50E-235 | -0.37955 | 0.476 | 0.76 | 9.48E-231 |
| NR4A2 | 2.13E-53 | -0.38189 | 0.134 | 0.239 | 8.07E-49 |
| MYOF | 1.44E-227 | -0.38231 | 0.326 | 0.629 | 5.46E-223 |
| SSR1 | 6.72E-245 | -0.3833 | 0.339 | 0.642 | 2.55E-240 |
| PDLIM4 | 5.57E-215 | -0.38373 | 0.421 | 0.7 | 2.12E-210 |
| METRN | 1.86E-278 | -0.38398 | 0.167 | 0.474 | 7.05E-274 |
| ZNF703 | 3.44E-272 | -0.38434 | 0.172 | 0.483 | 1.31E-267 |
| SEMA5A | 9.57E-299 | -0.38453 | 0.018 | 0.276 | 3.63E-294 |
| TSPAN3 | 8.65E-249 | -0.38476 | 0.411 | 0.715 | 3.28E-244 |
| NSD1 | 0 | -0.38515 | 0.18 | 0.522 | 0 |

| | | | | | |
|---|---|---|---|---|---|
| SYTL2 | 0 | -0.38528 | 0.093 | 0.412 | 0 |
| VEGFC | 5.60E-271 | -0.38545 | 0.04 | 0.292 | 2.13E-266 |
| RNF213 | 1.02E-233 | -0.38622 | 0.298 | 0.599 | 3.87E-229 |
| EEA1 | 8.60E-192 | -0.38655 | 0.604 | 0.835 | 3.27E-187 |
| LYZ | 1.37E-301 | -0.38707 | 0.005 | 0.25 | 5.20E-297 |
| TRIM44 | 8.01E-275 | -0.38738 | 0.253 | 0.582 | 3.04E-270 |
| PRMT2 | 7.54E-256 | -0.38741 | 0.301 | 0.619 | 2.86E-251 |
| LGALS1 | 9.80E-111 | -0.38806 | 0.96 | 0.985 | 3.72E-106 |
| WLS | 2.88E-264 | -0.38893 | 0.223 | 0.544 | 1.09E-259 |
| SMARCA1 | 0 | -0.38911 | 0.155 | 0.49 | 0 |
| CHPF | 1.91E-230 | -0.39107 | 0.16 | 0.433 | 7.24E-226 |
| RHOC | 2.91E-162 | -0.39155 | 0.602 | 0.81 | 1.10E-157 |
| 2-Sep | 1.53E-245 | -0.3916 | 0.652 | 0.872 | 5.80E-241 |
| LINC01423 | 4.22E-187 | -0.39229 | 0.005 | 0.169 | 1.60E-182 |
| CD109 | 1.98E-219 | -0.39282 | 0.206 | 0.489 | 7.50E-215 |
| DAD1 | 2.75E-266 | -0.39284 | 0.869 | 0.971 | 1.04E-261 |
| DNM3OS | 0 | -0.39301 | 0.088 | 0.404 | 0 |
| FAM13C | 1.63E-301 | -0.39313 | 0.039 | 0.315 | 6.21E-297 |
| BMP1 | 5.17E-263 | -0.39553 | 0.198 | 0.501 | 1.96E-258 |
| PKM | 1.75E-123 | -0.39623 | 0.775 | 0.896 | 6.65E-119 |
| TES | 1.00E-243 | -0.39658 | 0.228 | 0.526 | 3.80E-239 |
| ITIH5 | 9.39E-182 | -0.39768 | 0.114 | 0.339 | 3.56E-177 |
| LUC7L3 | 7.16E-238 | -0.39811 | 0.601 | 0.877 | 2.72E-233 |
| TPM3 | 1.90E-208 | -0.39906 | 0.514 | 0.763 | 7.22E-204 |
| ZC3H13 | 7.45E-260 | -0.39957 | 0.34 | 0.667 | 2.83E-255 |
| RAMP3 | 6.37E-210 | -0.39969 | 0.006 | 0.188 | 2.42E-205 |
| FHL1 | 1.83E-217 | -0.40009 | 0.729 | 0.932 | 6.94E-213 |
| PLD3 | 1.57E-247 | -0.4024 | 0.641 | 0.885 | 5.98E-243 |
| COPB2 | 1.96E-258 | -0.40242 | 0.388 | 0.686 | 7.43E-254 |
| TRAPPC1 | 2.10E-286 | -0.40248 | 0.371 | 0.704 | 7.98E-282 |
| EMP2 | 3.64E-239 | -0.40362 | 0.478 | 0.789 | 1.38E-234 |
| THBS4 | 2.54E-188 | -0.4039 | 0.425 | 0.698 | 9.66E-184 |
| RNASET2 | 0 | -0.40405 | 0.102 | 0.45 | 0 |
| PDIA4 | 2.01E-256 | -0.40416 | 0.389 | 0.694 | 7.65E-252 |
| ALDH3A2 | 3.39E-241 | -0.40426 | 0.3 | 0.612 | 1.29E-236 |
| CMKLR1 | 2.98E-295 | -0.40673 | 0.046 | 0.323 | 1.13E-290 |
| MRPL51 | 1.19E-261 | -0.40791 | 0.642 | 0.882 | 4.51E-257 |
| GPX7 | 0 | -0.40801 | 0.102 | 0.442 | 0 |
| ZBTB20 | 3.25E-208 | -0.40812 | 0.725 | 0.924 | 1.23E-203 |
| PPP1CC | 6.94E-227 | -0.4083 | 0.465 | 0.741 | 2.63E-222 |
| ITGB5 | 1.00E-163 | -0.40948 | 0.611 | 0.823 | 3.81E-159 |
| NDUFC2 | 4.75E-279 | -0.40961 | 0.708 | 0.922 | 1.80E-274 |
| AKR1A1 | 7.15E-266 | -0.40985 | 0.426 | 0.735 | 2.72E-261 |
| FLRT2 | 6.55E-308 | -0.41043 | 0.088 | 0.392 | 2.49E-303 |
| ISOC2 | 1.25E-292 | -0.41064 | 0.258 | 0.587 | 4.76E-288 |
| LOXL2 | 6.32E-209 | -0.41306 | 0.125 | 0.37 | 2.40E-204 |
| SMARCA2 | 9.44E-299 | -0.41356 | 0.286 | 0.635 | 3.59E-294 |

| | | | | | |
|---|---|---|---|---|---|
| DLX4 | 2.46E-303 | -0.41374 | 0.075 | 0.379 | 9.35E-299 |
| NME3 | 6.25E-273 | -0.41419 | 0.537 | 0.83 | 2.37E-268 |
| HMGN3 | 3.93E-280 | -0.415 | 0.445 | 0.778 | 1.49E-275 |
| SDK1 | 1.23E-264 | -0.41588 | 0.193 | 0.495 | 4.67E-260 |
| SERTAD4 | 1.44E-281 | -0.41739 | 0.033 | 0.29 | 5.45E-277 |
| AL078639. | 6.84E-282 | -0.41862 | 0.177 | 0.492 | 2.60E-277 |
| TCEAL8 | 1.97E-296 | -0.4187 | 0.371 | 0.706 | 7.47E-292 |
| PTGFRN | 1.55E-280 | -0.41943 | 0.136 | 0.436 | 5.87E-276 |
| SLC39A7 | 2.88E-263 | -0.41958 | 0.402 | 0.701 | 1.09E-258 |
| FSCN1 | 0 | -0.4201 | 0.076 | 0.403 | 0 |
| ELK3 | 0 | -0.42063 | 0.184 | 0.526 | 0 |
| PTPRD | 0 | -0.42139 | 0.119 | 0.446 | 0 |
| CCPG1 | 1.10E-229 | -0.42139 | 0.627 | 0.841 | 4.17E-225 |
| NDUFB7 | 1.78E-294 | -0.42155 | 0.693 | 0.908 | 6.75E-290 |
| HSBP1 | 5.53E-283 | -0.42186 | 0.637 | 0.871 | 2.10E-278 |
| TPM1 | 3.86E-148 | -0.42384 | 0.503 | 0.73 | 1.46E-143 |
| MX1 | 8.85E-210 | -0.42474 | 0.051 | 0.269 | 3.36E-205 |
| CSRP1 | 8.98E-257 | -0.42615 | 0.359 | 0.686 | 3.41E-252 |
| TNFSF10 | 1.01E-186 | -0.42727 | 0.362 | 0.65 | 3.84E-182 |
| P3H1 | 0 | -0.42755 | 0.177 | 0.515 | 0 |
| CASC4 | 2.20E-290 | -0.42767 | 0.308 | 0.64 | 8.35E-286 |
| NAV1 | 3.46E-224 | -0.42861 | 0.326 | 0.613 | 1.32E-219 |
| FKBP11 | 0 | -0.42929 | 0.162 | 0.506 | 0 |
| SET | 8.31E-246 | -0.42952 | 0.663 | 0.882 | 3.15E-241 |
| SULF1 | 3.16E-229 | -0.43346 | 0.238 | 0.533 | 1.20E-224 |
| UBTF | 0 | -0.43354 | 0.249 | 0.601 | 0 |
| SSR2 | 5.78E-299 | -0.43408 | 0.838 | 0.963 | 2.19E-294 |
| AC245595 | 8.56E-230 | -0.43521 | 0.288 | 0.579 | 3.25E-225 |
| DYNC1H1 | 4.76E-250 | -0.4355 | 0.462 | 0.749 | 1.81E-245 |
| ACTN1 | 2.36E-243 | -0.43569 | 0.098 | 0.363 | 8.96E-239 |
| UBE2L6 | 1.42E-283 | -0.43704 | 0.249 | 0.565 | 5.37E-279 |
| SH3BGRL | 7.43E-304 | -0.43725 | 0.574 | 0.853 | 2.82E-299 |
| CDK14 | 0 | -0.4389 | 0.096 | 0.467 | 0 |
| ZMAT3 | 0 | -0.43891 | 0.078 | 0.384 | 0 |
| FAM107B | 1.83E-237 | -0.43913 | 0.247 | 0.539 | 6.93E-233 |
| CNPY4 | 0 | -0.43914 | 0.166 | 0.549 | 0 |
| C1S | 4.35E-288 | -0.44103 | 0.975 | 0.997 | 1.65E-283 |
| C12orf75 | 1.13E-275 | -0.44296 | 0.118 | 0.409 | 4.29E-271 |
| SH3PXD2A | 1.51E-306 | -0.44301 | 0.161 | 0.488 | 5.72E-302 |
| CHCHD10 | 7.75E-254 | -0.44367 | 0.234 | 0.54 | 2.94E-249 |
| MXD4 | 0 | -0.44507 | 0.284 | 0.636 | 0 |
| ST5 | 5.87E-296 | -0.44522 | 0.209 | 0.543 | 2.23E-291 |
| EIF2AK2 | 1.66E-303 | -0.44575 | 0.274 | 0.609 | 6.31E-299 |
| MMP23B | 0 | -0.44604 | 0.023 | 0.319 | 0 |
| CALM2 | 3.72E-267 | -0.44645 | 0.883 | 0.977 | 1.41E-262 |
| NFIB | 4.32E-239 | -0.44675 | 0.565 | 0.839 | 1.64E-234 |
| NDUFB2 | 4.49E-303 | -0.44924 | 0.676 | 0.91 | 1.70E-298 |

| | | | | | |
|---|---|---|---|---|---|
| ITM2B | 0 | -0.44937 | 0.996 | 0.999 | 0 |
| SKI | 0 | -0.4497 | 0.223 | 0.575 | 0 |
| ERAP2 | 0 | -0.4498 | 0.066 | 0.399 | 0 |
| LINC0111€ | 0 | -0.44989 | 0.173 | 0.534 | 0 |
| ZFP36L1 | 1.26E-157 | -0.45154 | 0.846 | 0.955 | 4.78E-153 |
| FAM20A | 1.50E-257 | -0.4522 | 0.183 | 0.473 | 5.71E-253 |
| CTSS | 0 | -0.45224 | 0.108 | 0.42 | 0 |
| EXT1 | 0 | -0.45241 | 0.164 | 0.515 | 0 |
| LAP3 | 6.83E-189 | -0.45265 | 0.29 | 0.545 | 2.59E-184 |
| GUK1 | 1.85E-302 | -0.45475 | 0.809 | 0.948 | 7.02E-298 |
| FKBP14 | 0 | -0.455 | 0.121 | 0.485 | 0 |
| EZR | 5.77E-150 | -0.45503 | 0.224 | 0.443 | 2.19E-145 |
| CCDC88A | 0 | -0.45523 | 0.209 | 0.568 | 0 |
| PDGFRA | 2.73E-217 | -0.45583 | 0.665 | 0.876 | 1.04E-212 |
| GJA1 | 7.02E-207 | -0.45698 | 0.305 | 0.571 | 2.66E-202 |
| MARCKSL1 | 3.66E-266 | -0.45833 | 0.081 | 0.351 | 1.39E-261 |
| GNAI2 | 2.22E-295 | -0.45858 | 0.634 | 0.884 | 8.44E-291 |
| TMEM176 | 1.20E-97 | -0.45916 | 0.217 | 0.379 | 4.56E-93 |
| PHLDA1 | 1.15E-152 | -0.45983 | 0.217 | 0.444 | 4.37E-148 |
| FGL2 | 4.47E-211 | -0.46068 | 0.234 | 0.519 | 1.70E-206 |
| CTSO | 0 | -0.46161 | 0.229 | 0.58 | 0 |
| ITM2C | 2.00E-207 | -0.4618 | 0.281 | 0.555 | 7.61E-203 |
| MIR99AHC | 3.51E-306 | -0.46282 | 0.264 | 0.62 | 1.33E-301 |
| P3H3 | 0 | -0.46453 | 0.172 | 0.556 | 0 |
| SCARA3 | 1.07E-155 | -0.46643 | 0.283 | 0.522 | 4.06E-151 |
| AP2M1 | 0 | -0.46671 | 0.59 | 0.854 | 0 |
| MGAT4C | 5.42E-282 | -0.46736 | 0.013 | 0.252 | 2.06E-277 |
| HLA-DQA1 | 1.79E-230 | -0.46869 | 0.005 | 0.2 | 6.78E-226 |
| OLFML1 | 0 | -0.46915 | 0.147 | 0.527 | 0 |
| NME4 | 0 | -0.47307 | 0.209 | 0.588 | 0 |
| C1orf122 | 0 | -0.47362 | 0.477 | 0.785 | 0 |
| GXYLT2 | 0 | -0.47456 | 0.079 | 0.415 | 0 |
| SVIL | 4.48E-234 | -0.47466 | 0.317 | 0.615 | 1.70E-229 |
| APLP2 | 2.49E-161 | -0.47485 | 0.874 | 0.963 | 9.47E-157 |
| CD47 | 0 | -0.47523 | 0.605 | 0.876 | 0 |
| CCL2 | 2.66E-83 | -0.47579 | 0.251 | 0.423 | 1.01E-78 |
| AKR7A2 | 0 | -0.47642 | 0.321 | 0.671 | 0 |
| PALLD | 4.07E-222 | -0.47647 | 0.298 | 0.575 | 1.55E-217 |
| IFI44L | 9.25E-290 | -0.47648 | 0.059 | 0.338 | 3.51E-285 |
| EML1 | 0 | -0.47708 | 0.145 | 0.52 | 0 |
| CDC42EP5 | 0 | -0.47923 | 0.245 | 0.631 | 0 |
| KIF5B | 0 | -0.47967 | 0.399 | 0.713 | 0 |
| MEST | 4.21E-263 | -0.48143 | 0.022 | 0.255 | 1.60E-258 |
| ANK2 | 1.26E-234 | -0.48156 | 0.302 | 0.592 | 4.79E-230 |
| APOL6 | 5.62E-293 | -0.48223 | 0.2 | 0.523 | 2.13E-288 |
| PDLIM2 | 0 | -0.48286 | 0.428 | 0.751 | 0 |
| WWTR1 | 0 | -0.48291 | 0.191 | 0.551 | 0 |

| | | | | | |
|---|---|---|---|---|---|
| TBL1XR1 | 0 | -0.48353 | 0.32 | 0.681 | 0 |
| TPD52L1 | 8.82E-265 | -0.48375 | 0.104 | 0.39 | 3.35E-260 |
| CANX | 0 | -0.48599 | 0.658 | 0.894 | 0 |
| TMEM98 | 0 | -0.48602 | 0.293 | 0.633 | 0 |
| SULF2 | 0 | -0.48633 | 0.275 | 0.639 | 0 |
| TMEM219 | 0 | -0.48696 | 0.441 | 0.782 | 0 |
| CNPY2 | 0 | -0.48793 | 0.462 | 0.8 | 0 |
| ATOX1 | 0 | -0.48808 | 0.514 | 0.804 | 0 |
| TMEM176 | 1.72E-135 | -0.48918 | 0.146 | 0.336 | 6.52E-131 |
| PDIA3 | 0 | -0.48943 | 0.865 | 0.972 | 0 |
| ENG | 0 | -0.4898 | 0.22 | 0.599 | 0 |
| GLI3 | 0 | -0.48991 | 0.182 | 0.53 | 0 |
| CLMP | 0 | -0.49002 | 0.393 | 0.72 | 0 |
| TUBB2B | 2.99E-245 | -0.49101 | 0.063 | 0.308 | 1.14E-240 |
| C1orf54 | 0 | -0.4916 | 0.097 | 0.464 | 0 |
| LRRC15 | 0 | -0.49181 | 0.001 | 0.292 | 0 |
| CALM3 | 0 | -0.49246 | 0.397 | 0.741 | 0 |
| CEMIP | 4.96E-267 | -0.49291 | 0.011 | 0.24 | 1.88E-262 |
| H3F3A | 0 | -0.49293 | 0.966 | 0.997 | 0 |
| ZKSCAN1 | 0 | -0.49496 | 0.232 | 0.621 | 0 |
| SPTAN1 | 0 | -0.49545 | 0.361 | 0.697 | 0 |
| ATP2B4 | 0 | -0.497 | 0.218 | 0.599 | 0 |
| CDON | 0 | -0.49713 | 0.145 | 0.511 | 0 |
| C4orf48 | 0 | -0.49823 | 0.079 | 0.437 | 0 |
| SMOC1 | 8.91E-268 | -0.49886 | 0.069 | 0.337 | 3.39E-263 |
| HIF1A | 3.37E-244 | -0.50051 | 0.308 | 0.603 | 1.28E-239 |
| ARPC1B | 1.08E-268 | -0.50097 | 0.484 | 0.741 | 4.09E-264 |
| AKR1C3 | 2.91E-275 | -0.50117 | 0.261 | 0.583 | 1.11E-270 |
| CCND2 | 8.81E-308 | -0.50123 | 0.24 | 0.57 | 3.35E-303 |
| FKBP2 | 0 | -0.50184 | 0.63 | 0.902 | 0 |
| CRIP2 | 1.28E-217 | -0.50323 | 0.275 | 0.553 | 4.87E-213 |
| FTX | 0 | -0.50545 | 0.282 | 0.662 | 0 |
| ABHD2 | 5.76E-282 | -0.50586 | 0.207 | 0.518 | 2.19E-277 |
| NOV | 6.56E-106 | -0.50753 | 0.138 | 0.305 | 2.49E-101 |
| TSPAN15 | 0 | -0.50819 | 0.039 | 0.346 | 0 |
| NUCKS1 | 0 | -0.51308 | 0.884 | 0.982 | 0 |
| CADM1 | 3.85E-165 | -0.51391 | 0.121 | 0.328 | 1.46E-160 |
| RUNX1T1 | 0 | -0.51516 | 0.247 | 0.61 | 0 |
| KDELR2 | 4.16E-289 | -0.51707 | 0.702 | 0.884 | 1.58E-284 |
| SGCD | 0 | -0.51714 | 0.079 | 0.474 | 0 |
| FAT1 | 0 | -0.51714 | 0.096 | 0.467 | 0 |
| TYMP | 1.21E-192 | -0.51814 | 0.605 | 0.822 | 4.58E-188 |
| HEG1 | 0 | -0.51857 | 0.144 | 0.499 | 0 |
| NOTCH2 | 0 | -0.51862 | 0.304 | 0.675 | 0 |
| MAP4K4 | 3.29E-279 | -0.51925 | 0.304 | 0.617 | 1.25E-274 |
| TMED3 | 0 | -0.52251 | 0.345 | 0.705 | 0 |
| FAM198B | 0 | -0.52256 | 0.054 | 0.438 | 0 |

| | | | | | |
|---|---|---|---|---|---|
| PLEC | 0 | -0.52289 | 0.245 | 0.616 | 0 |
| DYNLT1 | 0 | -0.52435 | 0.556 | 0.834 | 0 |
| SAMD9L | 0 | -0.52457 | 0.066 | 0.432 | 0 |
| SOX5 | 1.03E-292 | -0.52493 | 0.219 | 0.545 | 3.91E-288 |
| SESN3 | 0 | -0.52606 | 0.07 | 0.441 | 0 |
| MAP4 | 0 | -0.52666 | 0.468 | 0.78 | 0 |
| TAPBP | 0 | -0.52682 | 0.378 | 0.724 | 0 |
| PLOD2 | 0 | -0.52781 | 0.294 | 0.636 | 0 |
| ATRX | 0 | -0.52791 | 0.512 | 0.82 | 0 |
| GLG1 | 0 | -0.52825 | 0.454 | 0.804 | 0 |
| C9orf3 | 0 | -0.52843 | 0.215 | 0.601 | 0 |
| EIF5 | 0 | -0.5289 | 0.648 | 0.899 | 0 |
| P4HB | 0 | -0.53062 | 0.785 | 0.942 | 0 |
| CTSC | 8.55E-154 | -0.53109 | 0.103 | 0.295 | 3.25E-149 |
| MYL6B | 0 | -0.53162 | 0.248 | 0.657 | 0 |
| TMEM119 | 0 | -0.53317 | 0.033 | 0.392 | 0 |
| PHACTR2 | 0 | -0.53707 | 0.273 | 0.634 | 0 |
| FHL2 | 0 | -0.53824 | 0.064 | 0.393 | 0 |
| FBN1 | 2.96E-238 | -0.5387 | 0.79 | 0.956 | 1.12E-233 |
| SFRP1 | 7.63E-51 | -0.53946 | 0.294 | 0.429 | 2.90E-46 |
| ENPP2 | 0 | -0.53962 | 0.113 | 0.469 | 0 |
| NDUFA13 | 0 | -0.54008 | 0.579 | 0.856 | 0 |
| XAF1 | 0 | -0.54052 | 0.123 | 0.471 | 0 |
| POLR2J3.1 | 0 | -0.54239 | 0.25 | 0.629 | 0 |
| HLA-F | 1.94E-288 | -0.54429 | 0.235 | 0.564 | 7.38E-284 |
| GPX8 | 0 | -0.5459 | 0.382 | 0.752 | 0 |
| HSPG2 | 0 | -0.54601 | 0.47 | 0.835 | 0 |
| RAB31 | 0 | -0.54845 | 0.16 | 0.514 | 0 |
| PSMB8 | 0 | -0.54888 | 0.443 | 0.768 | 0 |
| MYL9 | 0 | -0.54971 | 0.576 | 0.875 | 0 |
| PLPP1 | 6.52E-180 | -0.54972 | 0.652 | 0.821 | 2.48E-175 |
| PLAGL1 | 1.51E-301 | -0.55102 | 0.407 | 0.729 | 5.74E-297 |
| CHI3L1 | 1.18E-107 | -0.55254 | 0.2 | 0.388 | 4.48E-103 |
| MACF1 | 0 | -0.55343 | 0.394 | 0.741 | 0 |
| IL11RA | 0 | -0.55374 | 0.157 | 0.541 | 0 |
| AIG1 | 0 | -0.55427 | 0.27 | 0.655 | 0 |
| MGST3 | 0 | -0.55429 | 0.823 | 0.979 | 0 |
| ANXA5 | 5.11E-298 | -0.55497 | 0.894 | 0.977 | 1.94E-293 |
| LAMB1 | 0 | -0.55843 | 0.426 | 0.759 | 0 |
| MYO1B | 0 | -0.55921 | 0.156 | 0.571 | 0 |
| PPP3CA | 0 | -0.55951 | 0.399 | 0.762 | 0 |
| EIF4G2 | 0 | -0.55986 | 0.646 | 0.873 | 0 |
| AHR | 1.77E-296 | -0.56165 | 0.262 | 0.584 | 6.72E-292 |
| PAM | 0 | -0.56271 | 0.518 | 0.846 | 0 |
| TAGLN2 | 2.32E-251 | -0.56469 | 0.793 | 0.916 | 8.80E-247 |
| MMP2 | 1.68E-220 | -0.56553 | 0.951 | 0.99 | 6.39E-216 |
| ZNF385D | 0 | -0.56715 | 0.1 | 0.491 | 0 |

| | | | | | |
|---|---|---|---|---|---|
| CLEC11A | 0 | -0.56869 | 0.079 | 0.404 | 0 |
| APP | 0 | -0.57034 | 0.704 | 0.942 | 0 |
| AKAP12 | 4.58E-95 | -0.57186 | 0.341 | 0.5 | 1.74E-90 |
| GOLGB1 | 0 | -0.57294 | 0.466 | 0.798 | 0 |
| TMEM196 | 1.20E-297 | -0.57357 | 0.12 | 0.433 | 4.55E-293 |
| UACA | 0 | -0.5751 | 0.352 | 0.693 | 0 |
| NORAD | 0 | -0.57645 | 0.533 | 0.837 | 0 |
| GTF2I | 0 | -0.57941 | 0.4 | 0.784 | 0 |
| S100A4 | 1.06E-245 | -0.57954 | 0.949 | 0.995 | 4.03E-241 |
| SELENOW | 0 | -0.58019 | 0.579 | 0.898 | 0 |
| IFI27L2 | 0 | -0.58101 | 0.563 | 0.869 | 0 |
| CAVIN3 | 0 | -0.58229 | 0.517 | 0.857 | 0 |
| ODF2L | 0 | -0.58379 | 0.296 | 0.685 | 0 |
| HSP90AA1 | 0 | -0.5841 | 0.929 | 0.99 | 0 |
| ZNF106 | 0 | -0.58456 | 0.321 | 0.712 | 0 |
| DDX17 | 0 | -0.5857 | 0.741 | 0.953 | 0 |
| CAPZB | 0 | -0.58644 | 0.675 | 0.895 | 0 |
| PRDX2 | 0 | -0.58651 | 0.629 | 0.914 | 0 |
| RAB13 | 0 | -0.58819 | 0.477 | 0.784 | 0 |
| TNFRSF12/ | 2.74E-203 | -0.59115 | 0.158 | 0.4 | 1.04E-198 |
| RPS10 | 0 | -0.59148 | 0.912 | 0.987 | 0 |
| FUS | 0 | -0.59223 | 0.737 | 0.951 | 0 |
| TSPO | 0 | -0.59274 | 0.768 | 0.961 | 0 |
| CALM1 | 0 | -0.59467 | 0.813 | 0.957 | 0 |
| SELENOP | 8.40E-183 | -0.59491 | 0.767 | 0.894 | 3.19E-178 |
| PRRX1 | 7.48E-288 | -0.59516 | 0.657 | 0.885 | 2.84E-283 |
| PDGFRB | 5.10E-243 | -0.59565 | 0.567 | 0.82 | 1.94E-238 |
| PDLIM3 | 0 | -0.59604 | 0.191 | 0.551 | 0 |
| VAMP5 | 0 | -0.59942 | 0.517 | 0.838 | 0 |
| MYH9 | 0 | -0.60094 | 0.287 | 0.684 | 0 |
| TPM2 | 0 | -0.60167 | 0.315 | 0.641 | 0 |
| ITGAV | 0 | -0.60561 | 0.299 | 0.703 | 0 |
| SSC5D | 0 | -0.60708 | 0.32 | 0.763 | 0 |
| TNFSF13B | 0 | -0.60898 | 0.035 | 0.366 | 0 |
| FLNA | 0 | -0.60957 | 0.502 | 0.817 | 0 |
| COL18A1 | 0 | -0.60982 | 0.257 | 0.65 | 0 |
| CLTC | 0 | -0.61017 | 0.339 | 0.739 | 0 |
| PPIB | 0 | -0.61122 | 0.929 | 0.987 | 0 |
| COPZ2 | 0 | -0.61377 | 0.56 | 0.865 | 0 |
| APOL1 | 0 | -0.61463 | 0.118 | 0.508 | 0 |
| TRPS1 | 0 | -0.61473 | 0.347 | 0.728 | 0 |
| PDIA6 | 0 | -0.6188 | 0.696 | 0.934 | 0 |
| COL8A1 | 5.12E-290 | -0.62103 | 0.156 | 0.459 | 1.94E-285 |
| MSN | 0 | -0.62229 | 0.429 | 0.782 | 0 |
| BTN3A2 | 0 | -0.6228 | 0.078 | 0.525 | 0 |
| ANTXR1 | 0 | -0.62753 | 0.205 | 0.608 | 0 |
| GPC6 | 0 | -0.63471 | 0.116 | 0.549 | 0 |

| | | | | | |
|---|---|---|---|---|---|
| ARPC2 | 0 | -0.63602 | 0.694 | 0.917 | 0 |
| TCF4 | 0 | -0.63837 | 0.538 | 0.891 | 0 |
| UBB | 0 | -0.64129 | 0.887 | 0.986 | 0 |
| ADAMTS2 | 0 | -0.64173 | 0.272 | 0.599 | 0 |
| FAM114A1 | 0 | -0.64215 | 0.502 | 0.831 | 0 |
| HLA-DMA | 0 | -0.64393 | 0.062 | 0.391 | 0 |
| C1GALT1 | 3.09E-275 | -0.64423 | 0.444 | 0.725 | 1.17E-270 |
| S100A10 | 2.23E-263 | -0.64516 | 0.981 | 0.997 | 8.49E-259 |
| PDLIM7 | 0 | -0.64549 | 0.15 | 0.582 | 0 |
| MLEC | 0 | -0.64811 | 0.269 | 0.716 | 0 |
| CALU | 0 | -0.65062 | 0.585 | 0.857 | 0 |
| MAGED1 | 0 | -0.65134 | 0.118 | 0.582 | 0 |
| CPXM1 | 0 | -0.65721 | 0.013 | 0.349 | 0 |
| PLAC9 | 0 | -0.66012 | 0.927 | 0.993 | 0 |
| CHID1 | 0 | -0.66182 | 0.301 | 0.724 | 0 |
| SSR4 | 0 | -0.662 | 0.84 | 0.971 | 0 |
| GOLIM4 | 0 | -0.66515 | 0.443 | 0.816 | 0 |
| NRP1 | 0 | -0.66556 | 0.127 | 0.621 | 0 |
| RCAN1 | 0 | -0.66574 | 0.098 | 0.411 | 0 |
| CERCAM | 0 | -0.66611 | 0.284 | 0.658 | 0 |
| ITGB8 | 0 | -0.66887 | 0.122 | 0.497 | 0 |
| DYNLL1 | 0 | -0.67085 | 0.832 | 0.972 | 0 |
| LTBP3 | 0 | -0.67411 | 0.639 | 0.945 | 0 |
| HCFC1R1 | 0 | -0.67419 | 0.35 | 0.747 | 0 |
| MAGED2 | 0 | -0.67421 | 0.442 | 0.81 | 0 |
| CRISPLD1 | 0 | -0.68059 | 0.121 | 0.584 | 0 |
| FSTL1 | 0 | -0.685 | 0.906 | 0.983 | 0 |
| SEC31A | 0 | -0.69149 | 0.493 | 0.846 | 0 |
| SRGN | 0 | -0.6936 | 0.017 | 0.376 | 0 |
| NREP | 0 | -0.69436 | 0.019 | 0.477 | 0 |
| PARP14 | 0 | -0.69469 | 0.179 | 0.6 | 0 |
| FRMD6 | 0 | -0.69787 | 0.125 | 0.548 | 0 |
| NFIX | 0 | -0.69992 | 0.5 | 0.857 | 0 |
| OSTC | 0 | -0.7003 | 0.655 | 0.912 | 0 |
| EDIL3 | 1.30E-260 | -0.70122 | 0.109 | 0.368 | 4.95E-256 |
| HLA-C | 0 | -0.70455 | 0.96 | 0.993 | 0 |
| SSPN | 0 | -0.70841 | 0.491 | 0.86 | 0 |
| MMP3 | 1.73E-100 | -0.7128 | 0.071 | 0.213 | 6.55E-96 |
| DAB2 | 0 | -0.71511 | 0.695 | 0.931 | 0 |
| LBH | 0 | -0.71685 | 0.149 | 0.5 | 0 |
| SEMA3C | 5.65E-274 | -0.7178 | 0.422 | 0.704 | 2.15E-269 |
| CTSB | 0 | -0.71837 | 0.602 | 0.887 | 0 |
| KDELR3 | 0 | -0.72268 | 0.141 | 0.594 | 0 |
| RCN3 | 0 | -0.72482 | 0.49 | 0.836 | 0 |
| CD81 | 0 | -0.72528 | 0.979 | 0.999 | 0 |
| KCTD12 | 0 | -0.72802 | 0.148 | 0.57 | 0 |
| PSD3 | 0 | -0.73103 | 0.252 | 0.677 | 0 |

| | | | | | |
|---|---|---|---|---|---|
| TPM4 | 0 | -0.73196 | 0.627 | 0.835 | 0 |
| GLIPR1 | 0 | -0.73277 | 0.259 | 0.697 | 0 |
| MYL6 | 0 | -0.73297 | 0.954 | 0.996 | 0 |
| HDLBP | 0 | -0.73358 | 0.554 | 0.865 | 0 |
| PLXDC1 | 0 | -0.74092 | 0.32 | 0.71 | 0 |
| TMSB4X | 0 | -0.75081 | 0.982 | 0.999 | 0 |
| TUBB | 0 | -0.75102 | 0.69 | 0.925 | 0 |
| ITGB1 | 0 | -0.75261 | 0.768 | 0.96 | 0 |
| BEX3 | 0 | -0.75388 | 0.459 | 0.883 | 0 |
| P4HA2 | 0 | -0.75827 | 0.272 | 0.712 | 0 |
| EFEMP2 | 0 | -0.76317 | 0.505 | 0.862 | 0 |
| HMCN1 | 0 | -0.76317 | 0.059 | 0.484 | 0 |
| MEG3 | 0 | -0.76341 | 0.623 | 0.852 | 0 |
| EMILIN1 | 0 | -0.76372 | 0.411 | 0.779 | 0 |
| EMP3 | 0 | -0.76424 | 0.783 | 0.952 | 0 |
| CSGALNAC | 0 | -0.7666 | 0.075 | 0.559 | 0 |
| PSMB9 | 0 | -0.76819 | 0.238 | 0.657 | 0 |
| AEBP1 | 1.31E-274 | -0.76905 | 0.743 | 0.898 | 4.97E-270 |
| COL6A2 | 0 | -0.76929 | 0.98 | 0.999 | 0 |
| TUBB2A | 0 | -0.77015 | 0.28 | 0.637 | 0 |
| MINOS1 | 0 | -0.77143 | 0.639 | 0.912 | 0 |
| LIMA1 | 0 | -0.77654 | 0.7 | 0.946 | 0 |
| HSPA8 | 0 | -0.7843 | 0.649 | 0.923 | 0 |
| GALNT1 | 0 | -0.78533 | 0.341 | 0.727 | 0 |
| CD248 | 0 | -0.78758 | 0.432 | 0.764 | 0 |
| CTSZ | 0 | -0.78877 | 0.494 | 0.857 | 0 |
| ZEB2 | 0 | -0.79073 | 0.393 | 0.81 | 0 |
| CALR | 0 | -0.79339 | 0.79 | 0.976 | 0 |
| PRDX4 | 0 | -0.79357 | 0.611 | 0.899 | 0 |
| MYADM | 0 | -0.79423 | 0.579 | 0.844 | 0 |
| PPFIBP1 | 0 | -0.79434 | 0.338 | 0.748 | 0 |
| NID2 | 0 | -0.79592 | 0.042 | 0.536 | 0 |
| HMGN1 | 0 | -0.79908 | 0.37 | 0.816 | 0 |
| PTMS | 0 | -0.79981 | 0.764 | 0.961 | 0 |
| AKR1C2 | 0 | -0.80145 | 0.348 | 0.671 | 0 |
| NDUFA4L2 | 2.13E-257 | -0.8016 | 0.201 | 0.506 | 8.10E-253 |
| PMEPA1 | 0 | -0.80378 | 0.351 | 0.781 | 0 |
| LTBP1 | 0 | -0.80672 | 0.374 | 0.794 | 0 |
| PSME2 | 0 | -0.81004 | 0.473 | 0.817 | 0 |
| PHPT1 | 0 | -0.81251 | 0.543 | 0.893 | 0 |
| PLAU | 0 | -0.81842 | 0.179 | 0.594 | 0 |
| C2orf40 | 0 | -0.82106 | 0.339 | 0.699 | 0 |
| CCND1 | 0 | -0.82173 | 0.114 | 0.579 | 0 |
| CD63 | 0 | -0.83133 | 0.997 | 0.999 | 0 |
| GBP1 | 0 | -0.83684 | 0.156 | 0.509 | 0 |
| HSP90B1 | 0 | -0.83855 | 0.798 | 0.982 | 0 |
| THBS3 | 0 | -0.84334 | 0.227 | 0.696 | 0 |

| | | | | | |
|---|---|---|---|---|---|
| IGHM | 0 | -0.84431 | 0 | 0.364 | 0 |
| CPQ | 0 | -0.84578 | 0.56 | 0.917 | 0 |
| VKORC1 | 0 | -0.84836 | 0.695 | 0.938 | 0 |
| TUBA1A | 0 | -0.8539 | 0.572 | 0.893 | 0 |
| STEAP4 | 5.94E-237 | -0.86142 | 0.291 | 0.564 | 2.25E-232 |
| GLT8D2 | 0 | -0.86218 | 0.212 | 0.747 | 0 |
| DST | 0 | -0.86383 | 0.705 | 0.956 | 0 |
| NUCB2 | 0 | -0.86497 | 0.434 | 0.848 | 0 |
| IGF1 | 1.06E-131 | -0.86608 | 0.341 | 0.549 | 4.02E-127 |
| AQP1 | 9.53E-204 | -0.86854 | 0.277 | 0.536 | 3.62E-199 |
| LRRC17 | 0 | -0.86924 | 0.059 | 0.5 | 0 |
| TUBA1B | 0 | -0.87286 | 0.682 | 0.946 | 0 |
| LMNA | 0 | -0.87669 | 0.886 | 0.992 | 0 |
| CALD1 | 0 | -0.88387 | 0.921 | 0.997 | 0 |
| SMOC2 | 0 | -0.88917 | 0.223 | 0.632 | 0 |
| STAT1 | 0 | -0.89967 | 0.2 | 0.599 | 0 |
| PLTP | 0 | -0.90118 | 0.77 | 0.943 | 0 |
| RRBP1 | 0 | -0.90197 | 0.725 | 0.951 | 0 |
| ANGPTL1 | 0 | -0.90674 | 0.433 | 0.81 | 0 |
| SPARCL1 | 0 | -0.90931 | 0.471 | 0.858 | 0 |
| ITGBL1 | 1.96E-254 | -0.91588 | 0.514 | 0.759 | 7.46E-250 |
| XIST | 0 | -0.9162 | 0.643 | 0.897 | 0 |
| LSP1 | 0 | -0.92106 | 0.375 | 0.874 | 0 |
| ECM2 | 0 | -0.92592 | 0.412 | 0.856 | 0 |
| ID3 | 1.16E-301 | -0.92745 | 0.451 | 0.752 | 4.39E-297 |
| MRC2 | 0 | -0.931 | 0.483 | 0.918 | 0 |
| ISG15 | 0 | -0.93161 | 0.142 | 0.554 | 0 |
| SDC2 | 0 | -0.93339 | 0.687 | 0.936 | 0 |
| IGHG1 | 0 | -0.93391 | 0 | 0.459 | 0 |
| SCG2 | 5.62E-294 | -0.93886 | 0.049 | 0.319 | 2.13E-289 |
| SERF2 | 0 | -0.93988 | 0.971 | 0.998 | 0 |
| IGLC3 | 0 | -0.94013 | 0 | 0.436 | 0 |
| FNDC1 | 0 | -0.94097 | 0.23 | 0.559 | 0 |
| TGFBI | 0 | -0.94412 | 0.188 | 0.548 | 0 |
| FAP | 0 | -0.95193 | 0.25 | 0.743 | 0 |
| VMP1 | 0 | -0.9544 | 0.581 | 0.872 | 0 |
| DPYSL3 | 0 | -0.95889 | 0.265 | 0.749 | 0 |
| HLA-A | 0 | -0.96138 | 0.983 | 0.998 | 0 |
| CYBA | 0 | -0.96894 | 0.597 | 0.945 | 0 |
| IGFBP7 | 0 | -0.97395 | 0.64 | 0.891 | 0 |
| RGS3 | 0 | -0.9859 | 0.083 | 0.529 | 0 |
| CDH11 | 0 | -0.98837 | 0.173 | 0.688 | 0 |
| C2 | 0 | -0.9951 | 0.196 | 0.715 | 0 |
| SERPINH1 | 0 | -0.99693 | 0.303 | 0.79 | 0 |
| SFRP4 | 2.62E-46 | -0.99776 | 0.276 | 0.383 | 9.94E-42 |
| GOLM1 | 0 | -0.99866 | 0.202 | 0.741 | 0 |
| LMO4 | 0 | -1.00068 | 0.415 | 0.838 | 0 |

| | | | | | |
|---|---|---|---|---|---|
| ELN | 0 | -1.00402 | 0.184 | 0.563 | 0 |
| CLEC3B | 0 | -1.00912 | 0.288 | 0.703 | 0 |
| HLA-DPA1 | 0 | -1.01988 | 0.065 | 0.48 | 0 |
| EPB41L2 | 0 | -1.0385 | 0.515 | 0.935 | 0 |
| AHNAK | 0 | -1.03916 | 0.713 | 0.976 | 0 |
| ANKH | 0 | -1.0434 | 0.232 | 0.744 | 0 |
| EFEMP1 | 0 | -1.05962 | 0.803 | 0.909 | 0 |
| PTN | 0 | -1.06176 | 0.104 | 0.492 | 0 |
| OLFML2B | 0 | -1.0643 | 0.091 | 0.664 | 0 |
| LY6E | 0 | -1.06587 | 0.552 | 0.902 | 0 |
| CRABP2 | 0 | -1.06907 | 0.096 | 0.468 | 0 |
| LGALS3BP | 0 | -1.08624 | 0.399 | 0.834 | 0 |
| VCAM1 | 0 | -1.09421 | 0.198 | 0.723 | 0 |
| RARRES3 | 0 | -1.11409 | 0.201 | 0.723 | 0 |
| HLA-DRB5 | 0 | -1.11816 | 0.017 | 0.29 | 0 |
| C1QTNF3 | 0 | -1.11816 | 0.15 | 0.58 | 0 |
| COLEC12 | 0 | -1.11827 | 0.336 | 0.843 | 0 |
| C1QTNF4 | 0 | -1.12175 | 0.088 | 0.426 | 0 |
| LOXL1 | 0 | -1.12856 | 0.25 | 0.757 | 0 |
| CYP1B1 | 0 | -1.13441 | 0.535 | 0.877 | 0 |
| IGFBP5 | 5.41E-201 | -1.14028 | 0.493 | 0.721 | 2.06E-196 |
| SCRG1 | 0 | -1.18497 | 0.034 | 0.393 | 0 |
| NBL1 | 0 | -1.21441 | 0.65 | 0.926 | 0 |
| CRTAC1 | 0 | -1.22765 | 0.455 | 0.749 | 0 |
| RARRES2 | 0 | -1.22802 | 0.133 | 0.516 | 0 |
| VCAN | 0 | -1.22968 | 0.859 | 0.947 | 0 |
| COL6A3 | 0 | -1.23201 | 0.876 | 0.986 | 0 |
| TPPP3 | 0 | -1.24468 | 0.258 | 0.681 | 0 |
| TTC3 | 0 | -1.25829 | 0.382 | 0.907 | 0 |
| IGHG4 | 0 | -1.27517 | 0 | 0.564 | 0 |
| IGHA1 | 0 | -1.28321 | 0 | 0.437 | 0 |
| APOE | 1.34E-296 | -1.28952 | 0.33 | 0.633 | 5.09E-292 |
| CHI3L2 | 6.54E-268 | -1.29124 | 0.303 | 0.588 | 2.49E-263 |
| TIMP1 | 0 | -1.29243 | 0.968 | 0.996 | 0 |
| HLA-DPB1 | 0 | -1.31648 | 0.174 | 0.645 | 0 |
| CFI | 0 | -1.33797 | 0.1 | 0.727 | 0 |
| ENAH | 0 | -1.34603 | 0.242 | 0.84 | 0 |
| ANGPTL2 | 0 | -1.34808 | 0.573 | 0.865 | 0 |
| BST2 | 0 | -1.3498 | 0.091 | 0.568 | 0 |
| IFI6 | 0 | -1.35173 | 0.306 | 0.838 | 0 |
| ISLR | 0 | -1.35454 | 0.575 | 0.945 | 0 |
| LAMA4 | 0 | -1.36262 | 0.278 | 0.897 | 0 |
| KCNQ1OT1 | 0 | -1.36289 | 0.252 | 0.742 | 0 |
| COL6A1 | 0 | -1.38229 | 0.893 | 0.996 | 0 |
| FILIP1L | 0 | -1.39317 | 0.306 | 0.783 | 0 |
| CLU | 1.15E-135 | -1.41929 | 0.909 | 0.934 | 4.38E-131 |
| MARCKS | 0 | -1.42948 | 0.548 | 0.895 | 0 |

| | | | | | |
|---|---|---|---|---|---|
| B2M | 0 | -1.46281 | 0.999 | 1 | 0 |
| IGHG3 | 0 | -1.47848 | 0.001 | 0.569 | 0 |
| OGN | 0 | -1.49253 | 0.231 | 0.704 | 0 |
| COL5A1 | 0 | -1.52263 | 0.177 | 0.789 | 0 |
| MDK | 0 | -1.55004 | 0.09 | 0.693 | 0 |
| PPIC | 0 | -1.5815 | 0.427 | 0.937 | 0 |
| HLA-B | 0 | -1.58907 | 0.929 | 0.996 | 0 |
| PRSS23 | 0 | -1.66229 | 0.331 | 0.865 | 0 |
| IGFBP4 | 0 | -1.66992 | 0.729 | 0.965 | 0 |
| LUM | 0 | -1.69689 | 0.892 | 0.998 | 0 |
| HTRA1 | 0 | -1.71004 | 0.716 | 0.986 | 0 |
| CTGF | 0 | -1.74256 | 0.625 | 0.93 | 0 |
| IGLC2 | 0 | -1.85009 | 0.001 | 0.681 | 0 |
| POSTN | 1.16E-172 | -1.8572 | 0.019 | 0.184 | 4.40E-168 |
| COL14A1 | 0 | -1.9118 | 0.566 | 0.969 | 0 |
| MXRA5 | 0 | -2.00633 | 0.053 | 0.834 | 0 |
| COL5A2 | 0 | -2.00706 | 0.333 | 0.924 | 0 |
| CXCL12 | 0 | -2.01101 | 0.588 | 0.904 | 0 |
| HLA-DRB1 | 0 | -2.0169 | 0.072 | 0.602 | 0 |
| PCOLCE | 0 | -2.03355 | 0.683 | 0.992 | 0 |
| THY1 | 0 | -2.05006 | 0.137 | 0.736 | 0 |
| ASPN | 0 | -2.06557 | 0.442 | 0.682 | 0 |
| CCDC80 | 0 | -2.1117 | 0.624 | 0.992 | 0 |
| HLA-DRA | 0 | -2.14372 | 0.046 | 0.707 | 0 |
| CD74 | 0 | -2.22309 | 0.118 | 0.753 | 0 |
| IFI27 | 0 | -2.25585 | 0.421 | 0.927 | 0 |
| TNC | 0 | -2.33965 | 0.088 | 0.744 | 0 |
| BGN | 0 | -2.42203 | 0.336 | 0.957 | 0 |
| FN1 | 0 | -2.42991 | 0.894 | 0.999 | 0 |
| DPT | 0 | -2.49044 | 0.492 | 0.963 | 0 |
| SPARC | 0 | -2.91693 | 0.573 | 0.986 | 0 |
| PTGDS | 0 | -3.30313 | 0.082 | 0.63 | 0 |
| COL1A2 | 0 | -3.4453 | 0.716 | 1 | 0 |
| IGKC | 0 | -3.62023 | 0.001 | 0.83 | 0 |
| COL1A1 | 0 | -4.1341 | 0.527 | 0.992 | 0 |
| COL3A1 | 0 | -4.14205 | 0.562 | 0.997 | 0 |
