## Supplementary material for "Adipocytes regulate fibroblast function, and their loss contributes to fibroblast dysfunction in inflammatory diseases": Data file S2

Data File S2 Sublining healthy vs OA gene expression

| | p_val | avg_log2F( | pct.1 | pct.2 | p_val_adj |
|---|---|---|---|---|---|
| PLIN2 | 0 | 4.388092 | 0.834 | 0.359 | 0 |
| MT1X | 0 | 4.089059 | 0.971 | 0.793 | 0 |
| APOD | 0 | 3.915054 | 0.787 | 0.091 | 0 |
| PTX3 | 0 | 3.718036 | 0.442 | 0.094 | 0 |
| GPX3 | 0 | 3.639022 | 0.977 | 0.534 | 0 |
| DEPP1 | 0 | 3.551096 | 0.822 | 0.179 | 0 |
| GLUL | 0 | 3.24244 | 0.933 | 0.565 | 0 |
| MT1E | 0 | 3.005107 | 0.949 | 0.82 | 0 |
| ADM | 0 | 2.993827 | 0.79 | 0.229 | 0 |
| MT1M | 0 | 2.830105 | 0.882 | 0.688 | 0 |
| ADH1B | 0 | 2.687802 | 0.499 | 0.02 | 0 |
| ANGPTL7 | 3.04E-265 | 2.666724 | 0.167 | 0.007 | 1.15E-260 |
| HILPDA | 3.74E-278 | 2.615418 | 0.524 | 0.321 | 1.42E-273 |
| MT2A | 0 | 2.604619 | 0.987 | 0.963 | 0 |
| CYP4B1 | 0 | 2.355568 | 0.642 | 0.017 | 0 |
| DDIT4 | 1.03E-219 | 2.240536 | 0.702 | 0.573 | 3.91E-215 |
| CEBPD | 0 | 2.212776 | 0.97 | 0.821 | 0 |
| GALNT15 | 0 | 2.181536 | 0.864 | 0.298 | 0 |
| MT1A | 4.13E-228 | 2.079239 | 0.441 | 0.232 | 1.57E-223 |
| C11orf96 | 0 | 2.057733 | 0.857 | 0.514 | 0 |
| MYOC | 0 | 1.943228 | 0.313 | 0.031 | 0 |
| IGFBP6 | 0 | 1.905797 | 0.92 | 0.69 | 0 |
| ACKR3 | 0 | 1.884024 | 0.918 | 0.721 | 0 |
| GSN | 0 | 1.602668 | 0.999 | 0.991 | 0 |
| TGFBR3 | 0 | 1.568855 | 0.889 | 0.588 | 0 |
| IER2 | 0 | 1.555718 | 0.868 | 0.648 | 0 |
| RAMP2 | 0 | 1.537445 | 0.741 | 0.44 | 0 |
| PRELP | 0 | 1.521258 | 0.789 | 0.538 | 0 |
| H1FX | 0 | 1.48987 | 0.833 | 0.691 | 0 |
| TSC22D3 | 0 | 1.486786 | 0.885 | 0.853 | 0 |
| MFAP5 | 0 | 1.480995 | 0.801 | 0.26 | 0 |
| GADD45B | 0 | 1.478883 | 0.886 | 0.547 | 0 |
| ZBTB16 | 0 | 1.472155 | 0.715 | 0.119 | 0 |
| SOD2 | 0 | 1.462166 | 0.914 | 0.754 | 0 |
| PCOLCE2 | 0 | 1.454015 | 0.896 | 0.619 | 0 |
| ADAMTS5 | 0 | 1.405011 | 0.657 | 0.283 | 0 |
| RHOB | 0 | 1.401133 | 0.926 | 0.76 | 0 |
| FBLN2 | 0 | 1.380223 | 0.85 | 0.474 | 0 |
| TXNIP | 0 | 1.368538 | 0.965 | 0.948 | 0 |
| MGST1 | 0 | 1.36297 | 0.913 | 0.686 | 0 |
| NFKBIA | 0 | 1.362721 | 0.887 | 0.776 | 0 |
| NAMPT | 1.15E-197 | 1.360702 | 0.695 | 0.593 | 4.36E-193 |
| SLC39A14 | 0 | 1.353008 | 0.658 | 0.412 | 0 |
| MID1IP1 | 0 | 1.351633 | 0.613 | 0.283 | 0 |
| ADH5 | 0 | 1.351583 | 0.881 | 0.752 | 0 |

| | | | | | |
|---|---|---|---|---|---|
| TSKU | 0 | 1.33828 | 0.568 | 0.244 | 0 |
| UAP1 | 0 | 1.314491 | 0.842 | 0.666 | 0 |
| FGFBP2 | 3.35E-72 | 1.296264 | 0.305 | 0.185 | 1.27E-67 |
| IER3 | 9.56E-42 | 1.279265 | 0.477 | 0.437 | 3.63E-37 |
| GADD45A | 2.60E-44 | 1.269292 | 0.498 | 0.462 | 9.86E-40 |
| FTH1 | 0 | 1.24706 | 0.999 | 1 | 0 |
| MAOA | 0 | 1.245385 | 0.646 | 0.143 | 0 |
| MT1G | 9.97E-82 | 1.22975 | 0.184 | 0.079 | 3.79E-77 |
| CDO1 | 3.52E-286 | 1.227114 | 0.683 | 0.533 | 1.34E-281 |
| CRYAB | 2.48E-258 | 1.211484 | 0.765 | 0.714 | 9.42E-254 |
| GDF15 | 8.82E-190 | 1.209732 | 0.277 | 0.089 | 3.35E-185 |
| LAMA2 | 0 | 1.130974 | 0.571 | 0.131 | 0 |
| FKBP5 | 0 | 1.117594 | 0.666 | 0.371 | 0 |
| PNRC1 | 0 | 1.112788 | 0.944 | 0.907 | 0 |
| VEGFA | 0 | 1.086834 | 0.51 | 0.216 | 0 |
| NNMT | 0 | 1.086078 | 0.987 | 0.986 | 0 |
| ADH1C | 1.99E-299 | 1.082531 | 0.288 | 0.056 | 7.56E-295 |
| MFGE8 | 0 | 1.082275 | 0.783 | 0.542 | 0 |
| YBX3 | 0 | 1.071322 | 0.976 | 0.962 | 0 |
| LTBP4 | 0 | 1.068446 | 0.844 | 0.661 | 0 |
| CXCL1 | 1.51E-29 | 1.068384 | 0.156 | 0.095 | 5.74E-25 |
| NR1D1 | 0 | 1.05433 | 0.574 | 0.136 | 0 |
| DCN | 0 | 1.04447 | 1 | 0.997 | 0 |
| KLF9 | 0 | 1.038478 | 0.847 | 0.558 | 0 |
| CXCL2 | 1.40E-73 | 1.025443 | 0.245 | 0.132 | 5.33E-69 |
| PLPP3 | 0 | 1.021734 | 0.879 | 0.749 | 0 |
| ZFP36 | 0 | 1.017653 | 0.905 | 0.701 | 0 |
| SUN2 | 0 | 1.001218 | 0.683 | 0.391 | 0 |
| SLC3A2 | 0 | 0.999339 | 0.783 | 0.627 | 0 |
| BCL6 | 0 | 0.998102 | 0.677 | 0.35 | 0 |
| PHC2 | 0 | 0.996964 | 0.653 | 0.39 | 0 |
| CFD | 0 | 0.991044 | 0.997 | 0.803 | 0 |
| MTSS1 | 0 | 0.990049 | 0.494 | 0.104 | 0 |
| LTBP2 | 0 | 0.985244 | 0.597 | 0.344 | 0 |
| GLRX | 1.53E-132 | 0.97774 | 0.671 | 0.609 | 5.80E-128 |
| KLF4 | 0 | 0.974762 | 0.831 | 0.635 | 0 |
| ABLIM1 | 0 | 0.962368 | 0.631 | 0.269 | 0 |
| G0S2 | 1.31E-19 | 0.94798 | 0.276 | 0.38 | 4.97E-15 |
| RPS29 | 0 | 0.942351 | 0.991 | 0.992 | 0 |
| PROCR | 1.41E-230 | 0.940681 | 0.777 | 0.718 | 5.35E-226 |
| FIBIN | 0 | 0.938713 | 0.509 | 0.209 | 0 |
| JUN | 0 | 0.937994 | 0.976 | 0.874 | 0 |
| ZFAND5 | 0 | 0.936862 | 0.785 | 0.639 | 0 |
| PID1 | 0 | 0.93684 | 0.546 | 0.163 | 0 |
| H3F3B | 0 | 0.93183 | 0.997 | 0.997 | 0 |
| HIST1H1C | 2.89E-278 | 0.924942 | 0.463 | 0.218 | 1.10E-273 |
| ZFP36L2 | 8.15E-194 | 0.921014 | 0.913 | 0.877 | 3.10E-189 |

| | | | | | |
|---|---|---|---|---|---|
| CHST7 | 0 | 0.912568 | 0.44 | 0.063 | 0 |
| PPP1R15A | 0 | 0.911687 | 0.849 | 0.59 | 0 |
| DHRS3 | 0 | 0.902655 | 0.652 | 0.376 | 0 |
| PMP22 | 0 | 0.894081 | 0.95 | 0.951 | 0 |
| RPS27 | 0 | 0.892488 | 1 | 0.999 | 0 |
| NUPR1 | 0 | 0.888614 | 0.967 | 0.97 | 0 |
| NID1 | 0 | 0.869904 | 0.544 | 0.229 | 0 |
| SPSB1 | 0 | 0.863973 | 0.575 | 0.212 | 0 |
| PER1 | 0 | 0.861844 | 0.648 | 0.34 | 0 |
| CSF1 | 9.30E-230 | 0.85723 | 0.67 | 0.535 | 3.53E-225 |
| RPL38 | 0 | 0.852318 | 0.987 | 0.985 | 0 |
| PTGIS | 0 | 0.841179 | 0.443 | 0.126 | 0 |
| EIF1 | 0 | 0.837326 | 0.999 | 0.998 | 0 |
| AGTR1 | 0 | 0.834522 | 0.519 | 0.21 | 0 |
| USP53 | 4.88E-146 | 0.830233 | 0.56 | 0.441 | 1.85E-141 |
| CBLB | 0 | 0.828507 | 0.594 | 0.289 | 0 |
| VIT | 4.04E-282 | 0.828306 | 0.594 | 0.374 | 1.54E-277 |
| FOXO1 | 0 | 0.819838 | 0.636 | 0.406 | 0 |
| SMIM3 | 1.49E-249 | 0.819824 | 0.542 | 0.33 | 5.66E-245 |
| JUNB | 3.82E-251 | 0.814826 | 0.923 | 0.848 | 1.45E-246 |
| CYR61 | 3.11E-181 | 0.811149 | 0.694 | 0.471 | 1.18E-176 |
| HSD11B1 | 0 | 0.80026 | 0.525 | 0.176 | 0 |
| MIF | 2.44E-298 | 0.795446 | 0.781 | 0.7 | 9.26E-294 |
| SASH1 | 2.30E-263 | 0.794655 | 0.72 | 0.58 | 8.74E-259 |
| FBLN1 | 1.71E-276 | 0.794266 | 0.929 | 0.774 | 6.50E-272 |
| TWIST2 | 0 | 0.793761 | 0.519 | 0.184 | 0 |
| CCDC71L | 1.83E-99 | 0.792612 | 0.42 | 0.299 | 6.94E-95 |
| BTG2 | 1.10E-189 | 0.792223 | 0.581 | 0.375 | 4.17E-185 |
| KLF6 | 7.49E-275 | 0.792023 | 0.858 | 0.747 | 2.85E-270 |
| BNIP3 | 1.61E-234 | 0.786295 | 0.578 | 0.402 | 6.10E-230 |
| ANG | 0 | 0.778138 | 0.598 | 0.343 | 0 |
| ZFAS1 | 0 | 0.773814 | 0.935 | 0.921 | 0 |
| MEDAG | 1.38E-299 | 0.772557 | 0.771 | 0.598 | 5.24E-295 |
| H2AFZ | 1.85E-173 | 0.759146 | 0.765 | 0.741 | 7.03E-169 |
| THBS1 | 1.64E-179 | 0.757535 | 0.311 | 0.119 | 6.22E-175 |
| CTSL | 1.53E-268 | 0.751187 | 0.876 | 0.879 | 5.82E-264 |
| COL12A1 | 2.87E-233 | 0.730163 | 0.679 | 0.51 | 1.09E-228 |
| CEBPB | 3.15E-140 | 0.724606 | 0.885 | 0.836 | 1.20E-135 |
| H2AFJ | 0 | 0.720769 | 0.884 | 0.837 | 0 |
| LARP6 | 0 | 0.719347 | 0.667 | 0.473 | 0 |
| DPP4 | 0 | 0.717253 | 0.439 | 0.11 | 0 |
| NOVA1 | 1.44E-150 | 0.71537 | 0.805 | 0.749 | 5.45E-146 |
| ITGA5 | 0 | 0.712189 | 0.555 | 0.297 | 0 |
| MT-ND4L | 0 | 0.707826 | 0.97 | 0.964 | 0 |
| NEGR1 | 0 | 0.704224 | 0.497 | 0.157 | 0 |
| GABARAPL | 6.19E-243 | 0.698589 | 0.786 | 0.696 | 2.35E-238 |
| VASN | 6.56E-121 | 0.693418 | 0.67 | 0.622 | 2.49E-116 |

| | | | | | |
|---|---|---|---|---|---|
| RARRES1 | 0.001164 | 0.693121 | 0.492 | 0.591 | 1 |
| AC103591 | 2.72E-288 | 0.691881 | 0.321 | 0.083 | 1.03E-283 |
| PDK4 | 2.58E-113 | 0.687812 | 0.577 | 0.435 | 9.80E-109 |
| METRNL | 3.06E-205 | 0.683498 | 0.735 | 0.659 | 1.16E-200 |
| RPL37A | 0 | 0.682608 | 0.998 | 0.999 | 0 |
| ABCA9 | 0 | 0.674568 | 0.345 | 0.041 | 0 |
| HMGB2 | 1.92E-220 | 0.674426 | 0.645 | 0.496 | 7.27E-216 |
| DCXR | 2.42E-149 | 0.674192 | 0.503 | 0.37 | 9.18E-145 |
| NFIL3 | 7.00E-232 | 0.673605 | 0.646 | 0.401 | 2.66E-227 |
| AXL | 3.14E-189 | 0.673207 | 0.676 | 0.562 | 1.19E-184 |
| NDRG1 | 3.78E-143 | 0.669701 | 0.759 | 0.718 | 1.43E-138 |
| HSPA1B | 3.02E-86 | 0.669284 | 0.541 | 0.455 | 1.15E-81 |
| H1F0 | 2.51E-76 | 0.668825 | 0.456 | 0.372 | 9.54E-72 |
| DDX21 | 1.26E-150 | 0.668696 | 0.656 | 0.596 | 4.78E-146 |
| UBC | 1.78E-245 | 0.666529 | 0.994 | 0.996 | 6.76E-241 |
| RTN4 | 0 | 0.66373 | 0.955 | 0.95 | 0 |
| ARID5B | 2.93E-184 | 0.657743 | 0.853 | 0.828 | 1.11E-179 |
| WNT11 | 1.73E-259 | 0.65591 | 0.4 | 0.155 | 6.56E-255 |
| CPE | 1.73E-123 | 0.651668 | 0.475 | 0.301 | 6.55E-119 |
| ADH4 | 3.78E-255 | 0.649371 | 0.16 | 0.006 | 1.44E-250 |
| SESTD1 | 8.14E-199 | 0.646047 | 0.634 | 0.491 | 3.09E-194 |
| JUND | 8.25E-306 | 0.645701 | 0.988 | 0.972 | 3.13E-301 |
| CILP | 5.68E-163 | 0.637895 | 0.384 | 0.182 | 2.16E-158 |
| BTG3 | 1.07E-231 | 0.63335 | 0.596 | 0.417 | 4.04E-227 |
| SLC43A3 | 1.31E-263 | 0.632583 | 0.519 | 0.283 | 4.97E-259 |
| TNFAIP6 | 2.45E-25 | 0.630364 | 0.515 | 0.678 | 9.31E-21 |
| PQLC2L | 0 | 0.629146 | 0.336 | 0.054 | 0 |
| PDPN | 1.54E-56 | 0.625843 | 0.623 | 0.639 | 5.86E-52 |
| MAP3K8 | 5.37E-208 | 0.625003 | 0.463 | 0.258 | 2.04E-203 |
| C3 | 2.43E-294 | 0.623201 | 0.741 | 0.46 | 9.21E-290 |
| HMOX1 | 1.78E-133 | 0.619212 | 0.304 | 0.147 | 6.77E-129 |
| HSD17B11 | 1.63E-261 | 0.61757 | 0.619 | 0.429 | 6.19E-257 |
| PDGFRL | 3.05E-158 | 0.615293 | 0.814 | 0.757 | 1.16E-153 |
| SQSTM1 | 3.32E-286 | 0.614904 | 0.939 | 0.928 | 1.26E-281 |
| C15orf61 | 4.68E-197 | 0.614749 | 0.57 | 0.423 | 1.78E-192 |
| CYGB | 2.56E-186 | 0.61464 | 0.539 | 0.348 | 9.71E-182 |
| MMP14 | 1.03E-184 | 0.61389 | 0.733 | 0.63 | 3.92E-180 |
| RPL37 | 0 | 0.610602 | 0.998 | 0.998 | 0 |
| EIF4A1 | 4.96E-219 | 0.606564 | 0.792 | 0.749 | 1.88E-214 |
| RHOQ | 8.53E-229 | 0.605466 | 0.683 | 0.532 | 3.24E-224 |
| PIM3 | 1.16E-136 | 0.602424 | 0.462 | 0.317 | 4.41E-132 |
| CELF2 | 1.83E-177 | 0.60184 | 0.606 | 0.449 | 6.94E-173 |
| SNHG8 | 7.66E-195 | 0.601317 | 0.771 | 0.735 | 2.91E-190 |
| S100A6 | 6.51E-281 | 0.600226 | 1 | 1 | 2.47E-276 |
| A4GALT | 6.41E-168 | 0.599437 | 0.643 | 0.524 | 2.43E-163 |
| 3-Mar | 3.46E-175 | 0.594058 | 0.356 | 0.172 | 1.31E-170 |
| GYPC | 1.99E-201 | 0.593557 | 0.787 | 0.748 | 7.55E-197 |

| | | | | | |
|---|---|---|---|---|---|
| MAP1LC3E | 3.59E-237 | 0.591886 | 0.775 | 0.709 | 1.36E-232 |
| KLF2 | 8.96E-62 | 0.586419 | 0.716 | 0.673 | 3.40E-57 |
| RPL36 | 0 | 0.586344 | 0.994 | 0.995 | 0 |
| PIK3IP1 | 5.73E-184 | 0.585256 | 0.395 | 0.204 | 2.18E-179 |
| ADGRD1 | 0 | 0.585244 | 0.395 | 0.105 | 0 |
| CKB | 1.60E-124 | 0.581346 | 0.356 | 0.197 | 6.06E-120 |
| SDCBP | 1.28E-192 | 0.574723 | 0.846 | 0.844 | 4.85E-188 |
| MAP1B | 1.42E-105 | 0.57366 | 0.355 | 0.203 | 5.39E-101 |
| PXDC1 | 2.88E-203 | 0.573618 | 0.674 | 0.546 | 1.09E-198 |
| RSL24D1 | 1.04E-265 | 0.566353 | 0.812 | 0.77 | 3.95E-261 |
| TNFAIP2 | 5.49E-23 | 0.564839 | 0.623 | 0.654 | 2.09E-18 |
| WTAP | 1.90E-131 | 0.563173 | 0.695 | 0.626 | 7.23E-127 |
| CORO6 | 0 | 0.561278 | 0.282 | 0.037 | 0 |
| REV3L | 3.80E-115 | 0.56099 | 0.746 | 0.714 | 1.44E-110 |
| PNPLA2 | 1.27E-184 | 0.560979 | 0.489 | 0.31 | 4.82E-180 |
| SLPI | 3.75E-194 | 0.560698 | 0.563 | 0.297 | 1.42E-189 |
| WBP2 | 1.37E-225 | 0.558265 | 0.58 | 0.395 | 5.22E-221 |
| SLC2A3 | 3.62E-153 | 0.556518 | 0.473 | 0.279 | 1.38E-148 |
| TFPI | 9.48E-117 | 0.554776 | 0.751 | 0.74 | 3.60E-112 |
| CRISPLD2 | 4.30E-146 | 0.553909 | 0.441 | 0.247 | 1.63E-141 |
| TMBIM1 | 9.00E-200 | 0.553191 | 0.636 | 0.499 | 3.42E-195 |
| HERPUD1 | 5.61E-95 | 0.552676 | 0.782 | 0.817 | 2.13E-90 |
| C1orf21 | 4.65E-191 | 0.552608 | 0.605 | 0.441 | 1.77E-186 |
| DEFB1 | 2.45E-167 | 0.550748 | 0.352 | 0.154 | 9.29E-163 |
| DSE | 9.90E-58 | 0.550422 | 0.537 | 0.515 | 3.76E-53 |
| RPS20 | 0 | 0.547804 | 0.99 | 0.995 | 0 |
| CLIP1 | 5.70E-95 | 0.546583 | 0.629 | 0.604 | 2.16E-90 |
| FOSL2 | 1.05E-155 | 0.546421 | 0.486 | 0.318 | 3.98E-151 |
| PERP | 0 | 0.537724 | 0.32 | 0.067 | 0 |
| STOM | 6.72E-173 | 0.53727 | 0.742 | 0.679 | 2.55E-168 |
| MTRNR2L1 | 0 | 0.536746 | 0.982 | 0.991 | 0 |
| ANGPTL5 | 2.19E-82 | 0.535746 | 0.552 | 0.457 | 8.31E-78 |
| MTHFD2 | 1.27E-259 | 0.535129 | 0.369 | 0.134 | 4.83E-255 |
| RAB32 | 3.93E-169 | 0.534778 | 0.648 | 0.546 | 1.49E-164 |
| ST3GAL5 | 3.23E-164 | 0.530479 | 0.393 | 0.202 | 1.23E-159 |
| ABCA6 | 1.58E-203 | 0.530057 | 0.414 | 0.188 | 6.00E-199 |
| RHEB | 6.59E-225 | 0.525697 | 0.829 | 0.804 | 2.50E-220 |
| TM4SF1 | 3.23E-75 | 0.525033 | 0.287 | 0.17 | 1.23E-70 |
| STXBP6 | 0 | 0.524002 | 0.399 | 0.119 | 0 |
| RPL14 | 0 | 0.523978 | 0.997 | 0.998 | 0 |
| ID2 | 2.39E-46 | 0.522071 | 0.662 | 0.643 | 9.08E-42 |
| PHGDH | 2.66E-202 | 0.521055 | 0.392 | 0.18 | 1.01E-197 |
| EFHD1 | 0 | 0.520549 | 0.245 | 0.016 | 0 |
| EIF1B | 6.14E-191 | 0.518189 | 0.773 | 0.75 | 2.33E-186 |
| FBXO32 | 9.42E-48 | 0.517038 | 0.4 | 0.328 | 3.58E-43 |
| RETREG1 | 5.78E-78 | 0.516256 | 0.736 | 0.747 | 2.19E-73 |
| SDC4 | 1.02E-51 | 0.514624 | 0.37 | 0.294 | 3.88E-47 |

| | | | | | |
|---|---|---|---|---|---|
| SERPINE2 | 5.74E-11 | 0.514131 | 0.219 | 0.186 | 2.18E-06 |
| STK24 | 5.89E-159 | 0.514089 | 0.485 | 0.328 | 2.24E-154 |
| RPS25 | 0 | 0.513386 | 0.995 | 0.998 | 0 |
| RND3 | 3.68E-62 | 0.511029 | 0.655 | 0.584 | 1.40E-57 |
| SOCS1 | 1.50E-86 | 0.509112 | 0.265 | 0.143 | 5.70E-82 |
| SLC16A7 | 6.98E-134 | 0.507574 | 0.494 | 0.344 | 2.65E-129 |
| XG | 1.97E-130 | 0.506285 | 0.448 | 0.302 | 7.50E-126 |
| BLVRB | 1.96E-154 | 0.504832 | 0.663 | 0.572 | 7.43E-150 |
| IFRD1 | 1.29E-150 | 0.502895 | 0.545 | 0.387 | 4.92E-146 |
| UBE2B | 1.77E-205 | 0.502172 | 0.858 | 0.847 | 6.74E-201 |
| ACYP1 | 5.66E-140 | 0.501402 | 0.432 | 0.276 | 2.15E-135 |
| UBE2D3 | 9.70E-233 | 0.501272 | 0.84 | 0.811 | 3.68E-228 |
| EPB41L4A- | 1.80E-145 | 0.500352 | 0.729 | 0.672 | 6.85E-141 |
| CNBP | 7.46E-264 | 0.500257 | 0.918 | 0.925 | 2.83E-259 |
| AHNAK2 | 1.50E-92 | 0.499556 | 0.443 | 0.313 | 5.69E-88 |
| CD55 | 1.66E-100 | 0.497164 | 0.687 | 0.607 | 6.32E-96 |
| ZFHX3 | 5.79E-114 | 0.494627 | 0.584 | 0.498 | 2.20E-109 |
| DDIT3 | 2.92E-59 | 0.487708 | 0.496 | 0.441 | 1.11E-54 |
| ECE1 | 6.58E-157 | 0.485994 | 0.493 | 0.32 | 2.50E-152 |
| MT-ND2 | 1.20E-279 | 0.483177 | 0.998 | 1 | 4.56E-275 |
| KDSR | 2.22E-135 | 0.481832 | 0.655 | 0.583 | 8.42E-131 |
| PALM | 1.86E-177 | 0.480995 | 0.423 | 0.227 | 7.06E-173 |
| ERRFI1 | 3.07E-38 | 0.479641 | 0.556 | 0.492 | 1.17E-33 |
| RPS13 | 0 | 0.479188 | 0.995 | 0.997 | 0 |
| UBXN1 | 2.24E-176 | 0.475737 | 0.795 | 0.783 | 8.50E-172 |
| TAF1D | 7.44E-107 | 0.474286 | 0.728 | 0.715 | 2.83E-102 |
| ERICH1 | 1.30E-110 | 0.472663 | 0.552 | 0.455 | 4.95E-106 |
| RPS28 | 0 | 0.470458 | 0.999 | 0.998 | 0 |
| CD44 | 9.73E-119 | 0.466462 | 0.882 | 0.892 | 3.70E-114 |
| CNOT8 | 2.34E-169 | 0.464807 | 0.382 | 0.2 | 8.87E-165 |
| KLF15 | 0 | 0.460929 | 0.223 | 0.008 | 0 |
| MAFF | 6.48E-201 | 0.45982 | 0.463 | 0.227 | 2.46E-196 |
| ALDH2 | 2.67E-70 | 0.457279 | 0.617 | 0.587 | 1.02E-65 |
| BHLHE40 | 5.42E-46 | 0.456804 | 0.491 | 0.429 | 2.06E-41 |
| GBE1 | 3.85E-114 | 0.45254 | 0.415 | 0.279 | 1.46E-109 |
| BTF3 | 0 | 0.451428 | 0.98 | 0.989 | 0 |
| RPLP2 | 0 | 0.451249 | 0.995 | 0.998 | 0 |
| PIM1 | 2.57E-119 | 0.450975 | 0.341 | 0.18 | 9.75E-115 |
| GPRC5A | 3.86E-93 | 0.44975 | 0.439 | 0.297 | 1.46E-88 |
| GSTO1 | 4.83E-122 | 0.445754 | 0.805 | 0.82 | 1.84E-117 |
| HSPB8 | 1.37E-131 | 0.445587 | 0.381 | 0.22 | 5.21E-127 |
| HNRNPF | 2.13E-164 | 0.44505 | 0.793 | 0.781 | 8.07E-160 |
| SPOCK1 | 2.87E-100 | 0.44476 | 0.563 | 0.429 | 1.09E-95 |
| UBALD2 | 9.47E-152 | 0.442924 | 0.431 | 0.26 | 3.60E-147 |
| HIST1H4C | 7.84E-51 | 0.438061 | 0.576 | 0.558 | 2.98E-46 |
| ZNF385A | 1.58E-92 | 0.437833 | 0.383 | 0.254 | 5.99E-88 |
| FOXO3 | 5.93E-57 | 0.436672 | 0.53 | 0.484 | 2.25E-52 |

| | | | | | |
|---|---|---|---|---|---|
| ALDH1A1 | 4.94E-151 | 0.435497 | 0.238 | 0.079 | 1.88E-146 |
| MXI1 | 1.53E-128 | 0.434722 | 0.448 | 0.299 | 5.81E-124 |
| HIGD1A | 6.33E-93 | 0.434661 | 0.494 | 0.402 | 2.40E-88 |
| MPST | 1.55E-103 | 0.432641 | 0.467 | 0.354 | 5.88E-99 |
| RPS16 | 0 | 0.432303 | 0.995 | 0.999 | 0 |
| RPL30 | 0 | 0.432048 | 0.999 | 0.999 | 0 |
| SPRY1 | 6.00E-23 | 0.430618 | 0.455 | 0.421 | 2.28E-18 |
| ATOH8 | 5.23E-293 | 0.42943 | 0.248 | 0.037 | 1.98E-288 |
| CYB5A | 4.28E-79 | 0.427386 | 0.609 | 0.586 | 1.62E-74 |
| HLA-E | 6.16E-196 | 0.426705 | 0.956 | 0.959 | 2.34E-191 |
| FAM180B | 1.21E-106 | 0.424205 | 0.304 | 0.156 | 4.59E-102 |
| POLR1D | 1.44E-118 | 0.423375 | 0.68 | 0.638 | 5.46E-114 |
| CAMK2N1 | 1.06E-158 | 0.422642 | 0.446 | 0.239 | 4.01E-154 |
| TNFSF9 | 5.09E-97 | 0.421842 | 0.304 | 0.167 | 1.93E-92 |
| DUSP1 | 8.74E-145 | 0.421362 | 0.962 | 0.899 | 3.32E-140 |
| IER5L | 1.35E-09 | 0.420343 | 0.53 | 0.598 | 5.12E-05 |
| RAMP2-AS | 3.69E-209 | 0.420239 | 0.286 | 0.089 | 1.40E-204 |
| RPL31 | 1.07E-200 | 0.418816 | 0.948 | 0.961 | 4.05E-196 |
| AAED1 | 1.95E-115 | 0.418 | 0.508 | 0.387 | 7.40E-111 |
| TGIF1 | 4.20E-143 | 0.416702 | 0.453 | 0.284 | 1.59E-138 |
| SOD3 | 1.21E-107 | 0.415617 | 0.772 | 0.725 | 4.58E-103 |
| BNIP3L | 8.83E-82 | 0.414811 | 0.755 | 0.769 | 3.35E-77 |
| ARHGAP6 | 4.40E-217 | 0.414806 | 0.301 | 0.096 | 1.67E-212 |
| SMS | 3.85E-137 | 0.414243 | 0.432 | 0.265 | 1.46E-132 |
| CTHRC1 | 2.49E-30 | 0.413079 | 0.364 | 0.29 | 9.46E-26 |
| IFITM1 | 3.21E-70 | 0.412466 | 0.481 | 0.405 | 1.22E-65 |
| MYC | 8.72E-126 | 0.412015 | 0.536 | 0.354 | 3.31E-121 |
| RPL18A | 0 | 0.406851 | 0.998 | 0.999 | 0 |
| ATF4 | 2.59E-107 | 0.405114 | 0.769 | 0.758 | 9.83E-103 |
| CD9 | 4.85E-90 | 0.404672 | 0.881 | 0.828 | 1.84E-85 |
| FSTL3 | 2.62E-109 | 0.403055 | 0.33 | 0.183 | 9.96E-105 |
| CD34 | 1.07E-60 | 0.402906 | 0.36 | 0.25 | 4.08E-56 |
| BRI3 | 1.34E-132 | 0.399651 | 0.897 | 0.915 | 5.11E-128 |
| HRCT1 | 1.39E-103 | 0.398573 | 0.236 | 0.106 | 5.26E-99 |
| C17orf58 | 1.07E-24 | 0.397711 | 0.364 | 0.33 | 4.08E-20 |
| CYBRD1 | 4.99E-88 | 0.396898 | 0.839 | 0.872 | 1.89E-83 |
| PRKAG2 | 5.90E-124 | 0.396474 | 0.368 | 0.211 | 2.24E-119 |
| TMEM109 | 2.09E-89 | 0.395427 | 0.625 | 0.578 | 7.94E-85 |
| GRINA | 9.23E-93 | 0.395194 | 0.573 | 0.506 | 3.51E-88 |
| MMD | 4.24E-142 | 0.394403 | 0.262 | 0.106 | 1.61E-137 |
| MGLL | 1.16E-209 | 0.394397 | 0.279 | 0.08 | 4.42E-205 |
| METTL7A | 3.11E-45 | 0.393737 | 0.689 | 0.73 | 1.18E-40 |
| EGFR | 1.50E-75 | 0.392569 | 0.574 | 0.514 | 5.71E-71 |
| AL118516. | 1.86E-177 | 0.392416 | 0.326 | 0.136 | 7.07E-173 |
| DHRS7 | 1.07E-84 | 0.390623 | 0.669 | 0.656 | 4.05E-80 |
| IL1R1 | 1.65E-67 | 0.390295 | 0.691 | 0.663 | 6.26E-63 |
| YPEL3 | 2.95E-53 | 0.387816 | 0.677 | 0.716 | 1.12E-48 |

| | | | | | |
|---|---|---|---|---|---|
| ARL4C | 2.22E-38 | 0.387453 | 0.479 | 0.416 | 8.42E-34 |
| OSBPL8 | 4.32E-46 | 0.386536 | 0.654 | 0.682 | 1.64E-41 |
| CITED2 | 3.96E-21 | 0.386398 | 0.452 | 0.438 | 1.50E-16 |
| SVEP1 | 3.05E-49 | 0.386209 | 0.37 | 0.287 | 1.16E-44 |
| CCNDBP1 | 4.93E-105 | 0.385628 | 0.514 | 0.402 | 1.87E-100 |
| FBXL3 | 1.25E-72 | 0.384054 | 0.475 | 0.402 | 4.77E-68 |
| TIMP4 | 1.12E-54 | 0.383237 | 0.278 | 0.187 | 4.26E-50 |
| TAF7 | 2.04E-95 | 0.381302 | 0.734 | 0.699 | 7.75E-91 |
| KDM7A | 3.09E-85 | 0.379552 | 0.363 | 0.24 | 1.18E-80 |
| UBA52 | 3.08E-263 | 0.378443 | 0.993 | 0.996 | 1.17E-258 |
| MFAP4 | 0.042865 | 0.377514 | 0.693 | 0.779 | 1 |
| HIST1H1E | 1.68E-104 | 0.376748 | 0.312 | 0.173 | 6.38E-100 |
| DYNLT3 | 1.19E-86 | 0.373361 | 0.44 | 0.329 | 4.50E-82 |
| MRPL33 | 6.40E-86 | 0.37332 | 0.724 | 0.74 | 2.43E-81 |
| NOX4 | 1.04E-118 | 0.37274 | 0.347 | 0.191 | 3.94E-114 |
| RBMS3 | 1.01E-49 | 0.371259 | 0.577 | 0.549 | 3.82E-45 |
| TRIP10 | 3.30E-83 | 0.368461 | 0.433 | 0.323 | 1.25E-78 |
| RPL7 | 6.50E-204 | 0.367076 | 0.991 | 0.996 | 2.47E-199 |
| SLC19A2 | 1.49E-184 | 0.366368 | 0.287 | 0.101 | 5.65E-180 |
| OSER1 | 8.86E-96 | 0.366154 | 0.407 | 0.288 | 3.36E-91 |
| FAXDC2 | 7.89E-55 | 0.365755 | 0.356 | 0.272 | 3.00E-50 |
| PPL | 2.98E-68 | 0.365035 | 0.351 | 0.235 | 1.13E-63 |
| ADAMTSL4 | 2.70E-101 | 0.363942 | 0.415 | 0.272 | 1.03E-96 |
| PRDX6 | 1.55E-104 | 0.363466 | 0.884 | 0.908 | 5.90E-100 |
| IRS2 | 5.14E-135 | 0.360702 | 0.28 | 0.123 | 1.95E-130 |
| MKNK2 | 7.24E-76 | 0.359978 | 0.382 | 0.277 | 2.75E-71 |
| APCDD1 | 3.92E-166 | 0.359485 | 0.15 | 0.023 | 1.49E-161 |
| PLA2G2A | 7.31E-65 | 0.358481 | 0.987 | 0.967 | 2.77E-60 |
| CA12 | 7.01E-35 | 0.358419 | 0.318 | 0.252 | 2.66E-30 |
| GYG1 | 1.63E-94 | 0.356699 | 0.4 | 0.274 | 6.21E-90 |
| TBC1D15 | 7.10E-59 | 0.356025 | 0.477 | 0.417 | 2.70E-54 |
| RPL35 | 3.18E-247 | 0.35567 | 0.994 | 0.998 | 1.21E-242 |
| YBX1 | 1.30E-125 | 0.353372 | 0.955 | 0.982 | 4.92E-121 |
| PPP2R2A | 3.04E-44 | 0.353266 | 0.474 | 0.441 | 1.15E-39 |
| CDKN1A | 1.64E-81 | 0.353227 | 0.474 | 0.326 | 6.24E-77 |
| FOS | 2.68E-119 | 0.35304 | 0.952 | 0.864 | 1.02E-114 |
| TTC32 | 2.76E-158 | 0.35282 | 0.305 | 0.13 | 1.05E-153 |
| DKK3 | 6.88E-14 | 0.351674 | 0.348 | 0.322 | 2.61E-09 |
| CREBRF | 9.75E-54 | 0.350432 | 0.54 | 0.495 | 3.70E-49 |
| RPL35A | 1.79E-293 | 0.349413 | 0.998 | 0.998 | 6.79E-289 |
| HSPA1A | 1.29E-07 | 0.34788 | 0.68 | 0.776 | 0.004884 |
| HTRA3 | 2.71E-15 | 0.346753 | 0.421 | 0.405 | 1.03E-10 |
| VDAC2 | 3.90E-97 | 0.346138 | 0.748 | 0.755 | 1.48E-92 |
| SERPING1 | 5.18E-149 | 0.345798 | 0.983 | 0.987 | 1.97E-144 |
| RAD23A | 4.77E-80 | 0.345694 | 0.683 | 0.682 | 1.81E-75 |
| FMNL2 | 1.09E-206 | 0.344494 | 0.266 | 0.076 | 4.16E-202 |
| RPS21 | 3.86E-195 | 0.343362 | 0.984 | 0.991 | 1.46E-190 |

| | | | | | |
|---|---|---|---|---|---|
| RNH1 | 1.29E-109 | 0.342494 | 0.893 | 0.916 | 4.91E-105 |
| PA2G4 | 2.51E-61 | 0.342387 | 0.712 | 0.744 | 9.53E-57 |
| MED30 | 5.06E-87 | 0.341523 | 0.466 | 0.364 | 1.92E-82 |
| ATP13A3 | 1.19E-76 | 0.340428 | 0.327 | 0.213 | 4.51E-72 |
| F3 | 2.29E-12 | 0.338433 | 0.443 | 0.403 | 8.71E-08 |
| CCNB1IP1 | 5.53E-58 | 0.33545 | 0.413 | 0.336 | 2.10E-53 |
| FAU | 5.36E-280 | 0.335017 | 0.995 | 0.998 | 2.03E-275 |
| FADS3 | 7.09E-83 | 0.334037 | 0.349 | 0.224 | 2.69E-78 |
| SWAP70 | 7.83E-87 | 0.334032 | 0.351 | 0.23 | 2.97E-82 |
| CERS2 | 3.60E-68 | 0.332962 | 0.584 | 0.547 | 1.37E-63 |
| GPNMB | 8.03E-20 | 0.33274 | 0.751 | 0.803 | 3.05E-15 |
| SNRPB | 7.39E-81 | 0.331637 | 0.734 | 0.751 | 2.81E-76 |
| ADGRG2 | 5.55E-79 | 0.331074 | 0.216 | 0.105 | 2.11E-74 |
| DBN1 | 3.79E-24 | 0.33093 | 0.409 | 0.386 | 1.44E-19 |
| RNMT | 7.89E-34 | 0.330823 | 0.598 | 0.623 | 3.00E-29 |
| LINC0113 | 3.95E-128 | 0.330734 | 0.26 | 0.102 | 1.50E-123 |
| HAS1 | 0.002984 | 0.330726 | 0.354 | 0.357 | 1 |
| ALDH6A1 | 6.51E-48 | 0.330236 | 0.329 | 0.252 | 2.47E-43 |
| RPL32 | 3.35E-249 | 0.329925 | 0.999 | 1 | 1.27E-244 |
| SAMHD1 | 2.73E-18 | 0.329059 | 0.446 | 0.423 | 1.04E-13 |
| SRSF3 | 2.47E-91 | 0.328668 | 0.876 | 0.907 | 9.37E-87 |
| ILF3-DT | 2.17E-36 | 0.328289 | 0.45 | 0.411 | 8.22E-32 |
| SYPL1 | 9.97E-64 | 0.327911 | 0.678 | 0.675 | 3.79E-59 |
| CXCL14 | 0.021941 | 0.32752 | 0.121 | 0.109 | 1 |
| VEGFD | 4.60E-148 | 0.32501 | 0.166 | 0.036 | 1.75E-143 |
| SMIM14 | 1.31E-45 | 0.324997 | 0.772 | 0.813 | 4.99E-41 |
| EIF4H | 3.78E-78 | 0.324483 | 0.724 | 0.714 | 1.44E-73 |
| RRAGC | 3.50E-65 | 0.324116 | 0.41 | 0.322 | 1.33E-60 |
| RAB3IL1 | 5.36E-82 | 0.322215 | 0.346 | 0.226 | 2.03E-77 |
| AKIRIN2 | 2.70E-82 | 0.319952 | 0.392 | 0.279 | 1.03E-77 |
| CHPT1 | 3.35E-50 | 0.318696 | 0.548 | 0.525 | 1.27E-45 |
| GLIPR2 | 8.85E-41 | 0.318295 | 0.523 | 0.486 | 3.36E-36 |
| PAMR1 | 6.93E-162 | 0.318133 | 0.23 | 0.066 | 2.63E-157 |
| LMCD1 | 4.41E-22 | 0.31797 | 0.319 | 0.273 | 1.68E-17 |
| LAMP1 | 1.11E-62 | 0.317814 | 0.849 | 0.909 | 4.23E-58 |
| UBE2I | 3.00E-85 | 0.317211 | 0.782 | 0.802 | 1.14E-80 |
| SRSF5 | 3.71E-67 | 0.316922 | 0.907 | 0.948 | 1.41E-62 |
| HNRNPH1 | 1.10E-37 | 0.316878 | 0.731 | 0.764 | 4.19E-33 |
| RPL8 | 7.23E-202 | 0.316042 | 0.997 | 0.999 | 2.75E-197 |
| NTM | 3.40E-100 | 0.315296 | 0.246 | 0.11 | 1.29E-95 |
| CNIH4 | 2.35E-51 | 0.314473 | 0.609 | 0.608 | 8.93E-47 |
| DLST | 3.41E-73 | 0.313004 | 0.37 | 0.264 | 1.29E-68 |
| CCDC59 | 3.44E-45 | 0.312681 | 0.54 | 0.522 | 1.31E-40 |
| SREBF1 | 6.74E-28 | 0.312263 | 0.409 | 0.369 | 2.56E-23 |
| SH3GL1 | 5.18E-47 | 0.311483 | 0.45 | 0.4 | 1.97E-42 |
| UGDH | 6.27E-32 | 0.311115 | 0.613 | 0.617 | 2.38E-27 |
| RPL22 | 9.85E-195 | 0.31086 | 0.991 | 0.997 | 3.74E-190 |

| | | | | | |
|---|---|---|---|---|---|
| ARHGDIA | 1.82E-65 | 0.310496 | 0.675 | 0.679 | 6.93E-61 |
| RBP4 | 2.91E-63 | 0.309504 | 0.268 | 0.152 | 1.10E-58 |
| ARF1 | 1.65E-96 | 0.309497 | 0.865 | 0.891 | 6.26E-92 |
| MGAT1 | 3.32E-41 | 0.309286 | 0.605 | 0.617 | 1.26E-36 |
| HSPA5 | 3.57E-83 | 0.309119 | 0.831 | 0.968 | 1.36E-78 |
| GMNN | 3.64E-62 | 0.309013 | 0.234 | 0.136 | 1.38E-57 |
| PLP2 | 1.82E-50 | 0.308282 | 0.674 | 0.639 | 6.93E-46 |
| PGK1 | 8.78E-24 | 0.308113 | 0.716 | 0.77 | 3.33E-19 |
| CCNI | 4.32E-99 | 0.308059 | 0.936 | 0.958 | 1.64E-94 |
| CDC5L | 7.74E-37 | 0.307722 | 0.543 | 0.54 | 2.94E-32 |
| AC245297. | 2.29E-41 | 0.307207 | 0.527 | 0.506 | 8.71E-37 |
| CAST | 8.15E-59 | 0.306643 | 0.894 | 0.94 | 3.09E-54 |
| RRAGA | 6.65E-66 | 0.306048 | 0.688 | 0.698 | 2.52E-61 |
| SLC25A6 | 8.07E-73 | 0.305148 | 0.917 | 0.948 | 3.06E-68 |
| ADAM33 | 1.48E-172 | 0.303117 | 0.215 | 0.055 | 5.60E-168 |
| CALCRL | 6.28E-32 | 0.302472 | 0.221 | 0.154 | 2.39E-27 |
| PRR13 | 9.24E-49 | 0.301918 | 0.539 | 0.515 | 3.51E-44 |
| SLC29A1 | 3.67E-27 | 0.301418 | 0.372 | 0.324 | 1.39E-22 |
| OTUD1 | 1.75E-27 | 0.301276 | 0.295 | 0.239 | 6.65E-23 |
| PLEKHM2 | 1.96E-42 | 0.30121 | 0.447 | 0.395 | 7.45E-38 |
| CFL2 | 1.96E-59 | 0.300995 | 0.392 | 0.305 | 7.45E-55 |
| FOXN3 | 6.23E-36 | 0.300085 | 0.577 | 0.566 | 2.37E-31 |
| SPON2 | 2.65E-06 | 0.299791 | 0.466 | 0.496 | 0.100508 |
| MT-ND3 | 1.53E-144 | 0.299632 | 0.999 | 1 | 5.81E-140 |
| RPAIN | 2.96E-42 | 0.299524 | 0.457 | 0.414 | 1.12E-37 |
| NFKBIZ | 1.31E-27 | 0.299423 | 0.615 | 0.589 | 4.96E-23 |
| NCOA7 | 7.27E-40 | 0.298978 | 0.608 | 0.594 | 2.76E-35 |
| VSIR | 1.17E-09 | 0.298476 | 0.569 | 0.631 | 4.44E-05 |
| RPL23 | 3.20E-103 | 0.297934 | 0.977 | 0.991 | 1.22E-98 |
| AC015912. | 9.33E-140 | 0.297358 | 0.203 | 0.062 | 3.54E-135 |
| TRA2B | 6.30E-48 | 0.296652 | 0.676 | 0.676 | 2.39E-43 |
| MTRNR2L8 | 5.53E-29 | 0.295827 | 0.867 | 0.855 | 2.10E-24 |
| TUBA1C | 3.11E-27 | 0.295319 | 0.414 | 0.385 | 1.18E-22 |
| WIPI1 | 1.03E-51 | 0.29522 | 0.392 | 0.314 | 3.92E-47 |
| ABTB1 | 2.26E-81 | 0.295136 | 0.277 | 0.158 | 8.58E-77 |
| SSH2 | 2.23E-44 | 0.29398 | 0.274 | 0.196 | 8.46E-40 |
| MGP | 6.90E-43 | 0.293537 | 0.981 | 0.966 | 2.62E-38 |
| FTL | 1.20E-54 | 0.293334 | 0.999 | 1 | 4.56E-50 |
| LINC01088 | 1.23E-61 | 0.293329 | 0.21 | 0.113 | 4.67E-57 |
| CLTB | 9.14E-45 | 0.29107 | 0.684 | 0.725 | 3.47E-40 |
| ENO1 | 6.45E-76 | 0.290533 | 0.931 | 0.959 | 2.45E-71 |
| CLEC2B | 3.72E-48 | 0.290337 | 0.187 | 0.105 | 1.41E-43 |
| NAGLU | 2.02E-27 | 0.290276 | 0.447 | 0.432 | 7.66E-23 |
| ANKRD37 | 7.85E-55 | 0.289253 | 0.321 | 0.231 | 2.98E-50 |
| EIF3G | 9.97E-79 | 0.289167 | 0.844 | 0.885 | 3.79E-74 |
| HDAC7 | 1.43E-47 | 0.289085 | 0.397 | 0.325 | 5.42E-43 |
| MAN1A1 | 0.036337 | 0.288988 | 0.664 | 0.783 | 1 |

| | | | | | |
|---|---|---|---|---|---|
| CLDND1 | 2.67E-54 | 0.288536 | 0.498 | 0.441 | 1.02E-49 |
| ROMO1 | 1.01E-53 | 0.288371 | 0.723 | 0.742 | 3.82E-49 |
| RPS7 | 9.32E-167 | 0.287347 | 0.989 | 0.997 | 3.54E-162 |
| HBP1 | 2.54E-37 | 0.287196 | 0.584 | 0.591 | 9.65E-33 |
| FST | 4.65E-75 | 0.286725 | 0.13 | 0.044 | 1.77E-70 |
| SYF2 | 1.00E-33 | 0.286725 | 0.682 | 0.723 | 3.80E-29 |
| CFLAR | 4.34E-27 | 0.285786 | 0.593 | 0.606 | 1.65E-22 |
| FMO2 | 2.84E-22 | 0.285526 | 0.155 | 0.101 | 1.08E-17 |
| UXT | 1.19E-51 | 0.283514 | 0.729 | 0.758 | 4.52E-47 |
| ERO1A | 2.36E-30 | 0.282015 | 0.318 | 0.26 | 8.96E-26 |
| KRTCAP2 | 4.55E-66 | 0.280919 | 0.745 | 0.733 | 1.73E-61 |
| ATP5ME | 9.12E-52 | 0.28091 | 0.796 | 0.827 | 3.46E-47 |
| SNW1 | 1.35E-39 | 0.280437 | 0.624 | 0.633 | 5.12E-35 |
| YWHAH | 2.79E-17 | 0.280328 | 0.55 | 0.563 | 1.06E-12 |
| C9orf72 | 3.46E-115 | 0.280276 | 0.197 | 0.07 | 1.31E-110 |
| TOM1 | 1.53E-50 | 0.278828 | 0.361 | 0.28 | 5.82E-46 |
| ACKR4 | 1.38E-113 | 0.278802 | 0.117 | 0.021 | 5.23E-109 |
| PPIA | 5.63E-80 | 0.278521 | 0.944 | 0.974 | 2.14E-75 |
| PTMA | 1.08E-23 | 0.278227 | 0.989 | 0.998 | 4.11E-19 |
| MRPS36 | 5.08E-30 | 0.278051 | 0.543 | 0.556 | 1.93E-25 |
| ZC3H12A | 2.79E-101 | 0.277797 | 0.188 | 0.071 | 1.06E-96 |
| ACAT1 | 1.48E-33 | 0.277341 | 0.453 | 0.421 | 5.62E-29 |
| ATP6V1F | 2.46E-32 | 0.276991 | 0.724 | 0.792 | 9.33E-28 |
| TAF9 | 3.56E-30 | 0.276451 | 0.459 | 0.437 | 1.35E-25 |
| RPL34 | 7.00E-178 | 0.276084 | 0.998 | 0.999 | 2.66E-173 |
| CDC42EP4 | 1.07E-69 | 0.27541 | 0.314 | 0.197 | 4.08E-65 |
| MAF1 | 3.04E-37 | 0.275192 | 0.598 | 0.608 | 1.16E-32 |
| LIF | 3.49E-121 | 0.274482 | 0.114 | 0.018 | 1.33E-116 |
| NDUFAF2 | 1.25E-27 | 0.274042 | 0.525 | 0.542 | 4.74E-23 |
| SMAD3 | 1.45E-30 | 0.27341 | 0.458 | 0.418 | 5.51E-26 |
| TNS2 | 3.83E-24 | 0.27339 | 0.508 | 0.511 | 1.45E-19 |
| TMA7 | 1.57E-101 | 0.273183 | 0.926 | 0.959 | 5.98E-97 |
| TGFBR2 | 4.50E-20 | 0.273164 | 0.719 | 0.769 | 1.71E-15 |
| CD59 | 2.53E-33 | 0.272906 | 0.74 | 0.789 | 9.60E-29 |
| GSTP1 | 1.12E-94 | 0.272734 | 0.95 | 0.97 | 4.24E-90 |
| TXLNG | 1.37E-52 | 0.272273 | 0.328 | 0.238 | 5.18E-48 |
| EIF3M | 2.27E-48 | 0.271948 | 0.669 | 0.691 | 8.61E-44 |
| COQ10B | 3.79E-44 | 0.271405 | 0.477 | 0.434 | 1.44E-39 |
| AC245014 | 1.36E-66 | 0.271234 | 0.151 | 0.063 | 5.16E-62 |
| LMOD1 | 1.52E-67 | 0.271232 | 0.156 | 0.065 | 5.76E-63 |
| USP10 | 4.06E-30 | 0.271045 | 0.31 | 0.256 | 1.54E-25 |
| TPRG1 | 3.23E-72 | 0.270187 | 0.228 | 0.121 | 1.23E-67 |
| ABCA1 | 6.22E-14 | 0.270013 | 0.378 | 0.366 | 2.36E-09 |
| COX7A1 | 9.75E-32 | 0.269891 | 0.625 | 0.631 | 3.70E-27 |
| ITPKC | 2.12E-121 | 0.269693 | 0.235 | 0.093 | 8.06E-117 |
| ELL2 | 4.90E-19 | 0.269599 | 0.444 | 0.41 | 1.86E-14 |
| PXN | 2.58E-87 | 0.269322 | 0.259 | 0.134 | 9.78E-83 |

| | | | | | |
|---|---|---|---|---|---|
| ARL4D | 7.69E-13 | 0.269256 | 0.385 | 0.374 | 2.92E-08 |
| IL33 | 1.31E-120 | 0.269141 | 0.207 | 0.071 | 4.99E-116 |
| ABCA8 | 1.91E-32 | 0.26898 | 0.544 | 0.492 | 7.24E-28 |
| ODC1 | 6.56E-27 | 0.267944 | 0.362 | 0.32 | 2.49E-22 |
| KLHL24 | 4.17E-24 | 0.26747 | 0.382 | 0.353 | 1.58E-19 |
| ICAM1 | 1.63E-28 | 0.267358 | 0.215 | 0.152 | 6.18E-24 |
| TRIB1 | 5.50E-103 | 0.266737 | 0.254 | 0.115 | 2.09E-98 |
| UQCRFS1 | 3.33E-42 | 0.266678 | 0.636 | 0.664 | 1.26E-37 |
| CRIM1 | 9.94E-30 | 0.266586 | 0.322 | 0.264 | 3.78E-25 |
| TANK | 3.41E-35 | 0.266105 | 0.508 | 0.486 | 1.30E-30 |
| RPL9 | 3.85E-150 | 0.266101 | 0.997 | 0.999 | 1.46E-145 |
| F10 | 1.63E-175 | 0.265773 | 0.158 | 0.023 | 6.18E-171 |
| RBM8A | 7.43E-37 | 0.265334 | 0.697 | 0.756 | 2.82E-32 |
| SPSB3 | 3.57E-24 | 0.26492 | 0.496 | 0.498 | 1.36E-19 |
| RAB7A | 7.29E-45 | 0.264853 | 0.708 | 0.746 | 2.77E-40 |
| RAB21 | 4.20E-33 | 0.264343 | 0.534 | 0.529 | 1.59E-28 |
| RPL29 | 7.62E-163 | 0.264065 | 0.996 | 0.998 | 2.90E-158 |
| PIK3R1 | 1.19E-08 | 0.263542 | 0.624 | 0.686 | 0.000451 |
| GCHFR | 4.13E-45 | 0.263137 | 0.211 | 0.131 | 1.57E-40 |
| ARHGAP1( | 2.29E-37 | 0.26309 | 0.304 | 0.235 | 8.68E-33 |
| TNFRSF11I | 0.103307 | 0.262773 | 0.432 | 0.463 | 1 |
| TMEM159 | 2.27E-39 | 0.26269 | 0.299 | 0.226 | 8.63E-35 |
| RBM3 | 5.88E-51 | 0.262664 | 0.85 | 0.886 | 2.23E-46 |
| IMPA2 | 1.16E-62 | 0.261809 | 0.305 | 0.202 | 4.39E-58 |
| CCNG2 | 1.02E-24 | 0.261705 | 0.397 | 0.365 | 3.86E-20 |
| SLC2A1 | 6.45E-96 | 0.261655 | 0.188 | 0.073 | 2.45E-91 |
| ELMO1 | 6.33E-48 | 0.261552 | 0.327 | 0.246 | 2.41E-43 |
| SNHG7 | 3.60E-22 | 0.261237 | 0.585 | 0.623 | 1.37E-17 |
| AMOTL2 | 1.84E-24 | 0.260515 | 0.331 | 0.284 | 6.99E-20 |
| PNRC2 | 6.15E-40 | 0.259423 | 0.661 | 0.662 | 2.34E-35 |
| CYP4X1 | 3.96E-174 | 0.259281 | 0.17 | 0.03 | 1.50E-169 |
| TLN2 | 4.72E-48 | 0.25909 | 0.29 | 0.205 | 1.79E-43 |
| NDRG2 | 0.269401 | 0.258758 | 0.246 | 0.271 | 1 |
| CYC1 | 2.92E-44 | 0.258735 | 0.621 | 0.624 | 1.11E-39 |
| CCDC85B | 3.65E-28 | 0.258577 | 0.691 | 0.742 | 1.38E-23 |
| MAGOH | 3.63E-45 | 0.258489 | 0.605 | 0.61 | 1.38E-40 |
| EIF4EBP1 | 1.00E-11 | 0.258111 | 0.333 | 0.327 | 3.80E-07 |
| CRLF1 | 3.42E-08 | 0.257562 | 0.517 | 0.504 | 0.001297 |
| INSR | 4.87E-43 | 0.257449 | 0.282 | 0.204 | 1.85E-38 |
| GNB2 | 2.20E-42 | 0.257443 | 0.719 | 0.752 | 8.35E-38 |
| RPL22L1 | 2.50E-16 | 0.255967 | 0.704 | 0.772 | 9.49E-12 |
| CCNL1 | 2.86E-135 | 0.255366 | 0.807 | 0.686 | 1.09E-130 |
| RPS15 | 1.38E-159 | 0.255199 | 0.998 | 0.999 | 5.24E-155 |
| SNX21 | 3.37E-15 | 0.255064 | 0.426 | 0.436 | 1.28E-10 |
| EMILIN2 | 3.92E-59 | 0.254651 | 0.36 | 0.247 | 1.49E-54 |
| RAB9A | 2.55E-34 | 0.253942 | 0.466 | 0.44 | 9.67E-30 |
| SGCE | 9.19E-48 | 0.253552 | 0.445 | 0.342 | 3.49E-43 |

| | | | | | |
|---|---|---|---|---|---|
| IFITM2 | 2.73E-16 | 0.253477 | 0.897 | 0.963 | 1.04E-11 |
| ZNF706 | 5.78E-36 | 0.253384 | 0.707 | 0.749 | 2.19E-31 |
| ARHGAP29 | 5.77E-27 | 0.253008 | 0.385 | 0.326 | 2.19E-22 |
| ORAI3 | 1.48E-32 | 0.252792 | 0.401 | 0.357 | 5.60E-28 |
| ARMCX1 | 5.21E-21 | 0.252692 | 0.462 | 0.46 | 1.98E-16 |
| ATP6V1B2 | 1.51E-32 | 0.252125 | 0.392 | 0.348 | 5.74E-28 |
| TMOD1 | 5.52E-31 | 0.251622 | 0.233 | 0.167 | 2.10E-26 |
| CSNK1A1 | 3.16E-105 | -0.25005 | 0.648 | 0.844 | 1.20E-100 |
| ZDHHC14 | 2.69E-177 | -0.25011 | 0.071 | 0.285 | 1.02E-172 |
| ITM2C | 1.93E-88 | -0.25044 | 0.281 | 0.482 | 7.33E-84 |
| SSBP2 | 7.48E-131 | -0.25052 | 0.376 | 0.645 | 2.84E-126 |
| SNRNP200 | 1.48E-152 | -0.25055 | 0.271 | 0.548 | 5.61E-148 |
| TM2D2 | 1.85E-138 | -0.25085 | 0.345 | 0.611 | 7.02E-134 |
| HNRNPA3 | 8.19E-117 | -0.25108 | 0.738 | 0.916 | 3.11E-112 |
| SVIL | 3.53E-99 | -0.25117 | 0.317 | 0.539 | 1.34E-94 |
| FAM98A | 8.20E-203 | -0.25136 | 0.124 | 0.389 | 3.12E-198 |
| LAP3 | 2.99E-108 | -0.25178 | 0.29 | 0.517 | 1.14E-103 |
| DGCR6L | 1.61E-170 | -0.25189 | 0.22 | 0.496 | 6.10E-166 |
| MGMT | 9.29E-164 | -0.25194 | 0.34 | 0.648 | 3.53E-159 |
| HAS2 | 2.12E-95 | -0.25204 | 0.129 | 0.297 | 8.07E-91 |
| EXTL2 | 6.71E-207 | -0.2525 | 0.127 | 0.404 | 2.55E-202 |
| HGF | 3.36E-192 | -0.25255 | 0.031 | 0.229 | 1.27E-187 |
| ERGIC2 | 3.60E-138 | -0.25262 | 0.405 | 0.682 | 1.37E-133 |
| FAM107B | 8.37E-124 | -0.25275 | 0.247 | 0.482 | 3.18E-119 |
| HNRNPA2I | 8.42E-109 | -0.25301 | 0.94 | 0.988 | 3.20E-104 |
| GABARAPI | 4.15E-106 | -0.25337 | 0.801 | 0.941 | 1.58E-101 |
| KITLG | 2.74E-157 | -0.25353 | 0.112 | 0.335 | 1.04E-152 |
| ADD3 | 1.81E-101 | -0.25377 | 0.576 | 0.809 | 6.86E-97 |
| JPX | 1.74E-142 | -0.25415 | 0.315 | 0.586 | 6.62E-138 |
| LSM3 | 1.13E-128 | -0.25452 | 0.555 | 0.799 | 4.28E-124 |
| RNF213 | 2.52E-147 | -0.25483 | 0.298 | 0.574 | 9.56E-143 |
| GUCY1A1 | 1.76E-178 | -0.25491 | 0.005 | 0.164 | 6.68E-174 |
| ZNF652 | 9.06E-200 | -0.2551 | 0.15 | 0.431 | 3.44E-195 |
| MAGEH1 | 8.94E-177 | -0.25521 | 0.224 | 0.51 | 3.39E-172 |
| TGOLN2 | 1.21E-134 | -0.25533 | 0.473 | 0.744 | 4.60E-130 |
| SDF4 | 3.17E-120 | -0.25579 | 0.622 | 0.847 | 1.20E-115 |
| PYCARD | 8.51E-179 | -0.25595 | 0.146 | 0.408 | 3.23E-174 |
| ZNF280D | 2.30E-181 | -0.2565 | 0.213 | 0.505 | 8.74E-177 |
| COL15A1 | 7.20E-53 | -0.25669 | 0.101 | 0.209 | 2.74E-48 |
| TMOD3 | 1.76E-139 | -0.2568 | 0.395 | 0.667 | 6.68E-135 |
| COPB1 | 5.24E-145 | -0.25705 | 0.397 | 0.679 | 1.99E-140 |
| COX6C | 4.52E-89 | -0.25775 | 0.897 | 0.973 | 1.71E-84 |
| EIF2AK4 | 4.37E-165 | -0.25799 | 0.287 | 0.577 | 1.66E-160 |
| TECR | 1.54E-145 | -0.25833 | 0.427 | 0.71 | 5.85E-141 |
| RPS18 | 4.27E-140 | -0.25835 | 0.999 | 1 | 1.62E-135 |
| CSAD | 2.35E-182 | -0.25853 | 0.178 | 0.455 | 8.93E-178 |
| FNDC4 | 3.35E-201 | -0.259 | 0.151 | 0.44 | 1.27E-196 |

| | | | | | |
|---|---|---|---|---|---|
| TMEM106 | 1.09E-146 | -0.25946 | 0.341 | 0.617 | 4.12E-142 |
| PAXX | 6.26E-194 | -0.25962 | 0.182 | 0.467 | 2.38E-189 |
| PRRC2C | 2.42E-119 | -0.25996 | 0.716 | 0.905 | 9.20E-115 |
| AL391807. | 5.94E-206 | -0.26015 | 0.045 | 0.268 | 2.26E-201 |
| CTTNBP2 | 2.90E-176 | -0.2606 | 0.16 | 0.427 | 1.10E-171 |
| NFIA | 1.70E-103 | -0.26069 | 0.583 | 0.814 | 6.46E-99 |
| NHLRC3 | 9.18E-170 | -0.26105 | 0.228 | 0.508 | 3.48E-165 |
| SLIT3 | 1.61E-49 | -0.2611 | 0.221 | 0.346 | 6.10E-45 |
| GLIS3 | 3.10E-143 | -0.26124 | 0.134 | 0.349 | 1.18E-138 |
| DPYD | 3.00E-202 | -0.26131 | 0.164 | 0.449 | 1.14E-197 |
| PIGT | 1.68E-123 | -0.26147 | 0.553 | 0.802 | 6.37E-119 |
| TXNDC15 | 2.74E-131 | -0.26162 | 0.456 | 0.721 | 1.04E-126 |
| ATXN7L3B | 6.40E-170 | -0.26164 | 0.294 | 0.593 | 2.43E-165 |
| EPRS | 2.93E-141 | -0.26205 | 0.37 | 0.635 | 1.11E-136 |
| SRSF9 | 7.30E-121 | -0.26265 | 0.675 | 0.872 | 2.77E-116 |
| S100A16 | 2.93E-105 | -0.26288 | 0.502 | 0.739 | 1.11E-100 |
| PRUNE2 | 1.24E-174 | -0.26339 | 0.124 | 0.369 | 4.71E-170 |
| SEC61B | 2.31E-117 | -0.26339 | 0.837 | 0.961 | 8.76E-113 |
| TMEM141 | 1.01E-190 | -0.26351 | 0.215 | 0.508 | 3.83E-186 |
| GAA | 2.03E-151 | -0.26365 | 0.329 | 0.616 | 7.69E-147 |
| EXOC7 | 9.19E-186 | -0.26414 | 0.2 | 0.485 | 3.49E-181 |
| IL6ST | 2.77E-86 | -0.26473 | 0.635 | 0.832 | 1.05E-81 |
| CCDC90B | 1.41E-168 | -0.26484 | 0.344 | 0.641 | 5.36E-164 |
| CETN2 | 3.93E-195 | -0.26484 | 0.184 | 0.469 | 1.49E-190 |
| NCOA4 | 2.85E-169 | -0.26524 | 0.238 | 0.512 | 1.08E-164 |
| AP1S1 | 4.18E-188 | -0.26606 | 0.184 | 0.462 | 1.59E-183 |
| AK1 | 2.20E-185 | -0.26673 | 0.154 | 0.421 | 8.35E-181 |
| ALDH7A1 | 2.52E-165 | -0.26686 | 0.185 | 0.446 | 9.56E-161 |
| RBMS1 | 4.32E-112 | -0.26706 | 0.6 | 0.823 | 1.64E-107 |
| RCN1 | 6.50E-110 | -0.26714 | 0.571 | 0.807 | 2.47E-105 |
| CTSF | 1.82E-128 | -0.26733 | 0.624 | 0.884 | 6.92E-124 |
| C12orf75 | 4.57E-124 | -0.26738 | 0.118 | 0.308 | 1.74E-119 |
| IGHG1 | 4.12E-233 | -0.26749 | 0 | 0.196 | 1.56E-228 |
| IFI44 | 2.26E-221 | -0.26768 | 0.08 | 0.338 | 8.59E-217 |
| PNISR | 7.89E-109 | -0.2677 | 0.852 | 0.96 | 3.00E-104 |
| PLOD1 | 4.28E-218 | -0.26771 | 0.111 | 0.384 | 1.62E-213 |
| RPS4Y1 | 0.000141 | -0.26792 | 0.231 | 0.242 | 1 |
| TMEM167 | 1.07E-126 | -0.268 | 0.467 | 0.717 | 4.06E-122 |
| CASP1 | 1.21E-255 | -0.26858 | 0.046 | 0.304 | 4.60E-251 |
| RCN2 | 2.98E-132 | -0.26911 | 0.508 | 0.762 | 1.13E-127 |
| ISCU | 2.58E-120 | -0.26921 | 0.735 | 0.91 | 9.80E-116 |
| EIF4G1 | 1.21E-160 | -0.26942 | 0.331 | 0.615 | 4.59E-156 |
| ZNF703 | 2.24E-177 | -0.26946 | 0.172 | 0.438 | 8.49E-173 |
| P4HA3 | 2.63E-189 | -0.26946 | 0.035 | 0.233 | 9.99E-185 |
| PARP1 | 5.96E-184 | -0.27014 | 0.228 | 0.518 | 2.26E-179 |
| EFNB1 | 4.86E-234 | -0.27049 | 0.057 | 0.309 | 1.84E-229 |
| PTPRF | 4.50E-226 | -0.2706 | 0.063 | 0.316 | 1.71E-221 |

| | | | | | |
|---|---|---|---|---|---|
| SULF1 | 2.53E-145 | -0.2706 | 0.238 | 0.499 | 9.62E-141 |
| ECHS1 | 2.50E-152 | -0.27082 | 0.494 | 0.771 | 9.51E-148 |
| EDEM3 | 1.80E-220 | -0.2709 | 0.131 | 0.415 | 6.84E-216 |
| SEC61A1 | 1.62E-160 | -0.27227 | 0.31 | 0.594 | 6.15E-156 |
| SGCA | 4.83E-133 | -0.27235 | 0.134 | 0.337 | 1.83E-128 |
| ARMCX2 | 1.58E-213 | -0.27245 | 0.144 | 0.429 | 6.01E-209 |
| ROBO1 | 6.02E-191 | -0.27258 | 0.151 | 0.423 | 2.28E-186 |
| ARHGEF12 | 3.38E-175 | -0.27266 | 0.256 | 0.547 | 1.28E-170 |
| IGHG4 | 1.50E-217 | -0.27293 | 0 | 0.184 | 5.70E-213 |
| LUZP1 | 6.60E-190 | -0.27353 | 0.234 | 0.534 | 2.51E-185 |
| C7orf50 | 4.57E-155 | -0.27357 | 0.305 | 0.579 | 1.74E-150 |
| LINC00632 | 2.83E-192 | -0.27368 | 0.088 | 0.326 | 1.07E-187 |
| YIF1A | 1.38E-125 | -0.27374 | 0.452 | 0.702 | 5.25E-121 |
| PSIP1 | 3.14E-146 | -0.27405 | 0.444 | 0.736 | 1.19E-141 |
| NDFIP1 | 2.62E-70 | -0.2741 | 0.737 | 0.881 | 9.97E-66 |
| PER3 | 3.15E-142 | -0.27448 | 0.177 | 0.405 | 1.20E-137 |
| DSEL | 4.66E-153 | -0.27465 | 0.238 | 0.496 | 1.77E-148 |
| GNAS | 1.31E-148 | -0.27515 | 0.957 | 0.995 | 4.97E-144 |
| ANXA6 | 1.04E-160 | -0.27588 | 0.399 | 0.696 | 3.95E-156 |
| C9orf47 | 6.82E-148 | -0.27638 | 0.031 | 0.191 | 2.59E-143 |
| CP | 2.96E-19 | -0.2765 | 0.541 | 0.635 | 1.12E-14 |
| SLC22A17 | 2.80E-251 | -0.27706 | 0.084 | 0.37 | 1.06E-246 |
| GALNT5 | 4.14E-277 | -0.27709 | 0.02 | 0.27 | 1.57E-272 |
| FHL2 | 4.88E-166 | -0.2771 | 0.064 | 0.264 | 1.85E-161 |
| ATRAID | 1.51E-128 | -0.27768 | 0.798 | 0.945 | 5.74E-124 |
| SETBP1 | 6.97E-160 | -0.27838 | 0.175 | 0.427 | 2.65E-155 |
| AKAP9 | 4.66E-119 | -0.2784 | 0.646 | 0.869 | 1.77E-114 |
| CRTAP | 8.78E-134 | -0.27937 | 0.493 | 0.75 | 3.34E-129 |
| CRIP2 | 1.85E-135 | -0.27986 | 0.275 | 0.528 | 7.03E-131 |
| ANP32A | 9.31E-184 | -0.27986 | 0.27 | 0.568 | 3.54E-179 |
| ERGIC3 | 1.51E-143 | -0.28006 | 0.452 | 0.716 | 5.72E-139 |
| CSRP1 | 5.33E-151 | -0.28048 | 0.359 | 0.652 | 2.02E-146 |
| HSPA4 | 1.56E-196 | -0.28186 | 0.206 | 0.496 | 5.93E-192 |
| ATP5F1A | 8.70E-130 | -0.28206 | 0.569 | 0.811 | 3.30E-125 |
| TCEAL9 | 8.54E-127 | -0.28255 | 0.598 | 0.833 | 3.24E-122 |
| ASAH1 | 1.37E-160 | -0.28273 | 0.426 | 0.726 | 5.20E-156 |
| SAMD9 | 1.97E-233 | -0.28302 | 0.071 | 0.333 | 7.49E-229 |
| ATP1B1 | 5.02E-198 | -0.28316 | 0.055 | 0.277 | 1.91E-193 |
| TIMM8B | 1.07E-155 | -0.28368 | 0.452 | 0.738 | 4.06E-151 |
| RSPO3 | 3.50E-140 | -0.28372 | 0.283 | 0.552 | 1.33E-135 |
| HLA-DQA1 | 2.47E-186 | -0.28455 | 0.005 | 0.17 | 9.37E-182 |
| BAZ1B | 8.73E-181 | -0.28478 | 0.267 | 0.563 | 3.32E-176 |
| NDN | 5.37E-137 | -0.28585 | 0.518 | 0.777 | 2.04E-132 |
| TGM2 | 4.13E-113 | -0.28619 | 0.003 | 0.107 | 1.57E-108 |
| CES1 | 9.79E-19 | -0.28648 | 0.329 | 0.406 | 3.72E-14 |
| MIA3 | 2.44E-131 | -0.28667 | 0.466 | 0.733 | 9.25E-127 |
| CBX3 | 4.28E-143 | -0.28673 | 0.524 | 0.781 | 1.62E-138 |

| | | | | | |
|---|---|---|---|---|---|
| TFDP2 | 1.00E-183 | -0.28705 | 0.318 | 0.633 | 3.80E-179 |
| SIX3 | 5.75E-212 | -0.28713 | 0.054 | 0.288 | 2.18E-207 |
| LDLRAD4 | 1.05E-285 | -0.28732 | 0.04 | 0.317 | 3.97E-281 |
| NEDD8 | 1.77E-147 | -0.28733 | 0.772 | 0.938 | 6.71E-143 |
| ACTN1 | 1.45E-172 | -0.28743 | 0.098 | 0.327 | 5.49E-168 |
| TMED1 | 1.57E-154 | -0.28751 | 0.401 | 0.685 | 5.98E-150 |
| NDUFS8 | 8.75E-161 | -0.28863 | 0.423 | 0.712 | 3.32E-156 |
| CD164 | 9.83E-113 | -0.28876 | 0.729 | 0.903 | 3.73E-108 |
| FKBP1A | 2.12E-121 | -0.28905 | 0.681 | 0.875 | 8.05E-117 |
| MMP19 | 9.22E-184 | -0.28906 | 0.089 | 0.319 | 3.50E-179 |
| ABI2 | 3.80E-203 | -0.28916 | 0.209 | 0.51 | 1.44E-198 |
| ARHGAP18 | 2.17E-209 | -0.28924 | 0.09 | 0.344 | 8.24E-205 |
| PCDH9 | 1.90E-128 | -0.28932 | 0.168 | 0.384 | 7.22E-124 |
| PRDX1 | 3.03E-127 | -0.28933 | 0.905 | 0.984 | 1.15E-122 |
| DRAP1 | 4.35E-142 | -0.28962 | 0.62 | 0.854 | 1.65E-137 |
| EPB41L1 | 1.93E-235 | -0.28977 | 0.057 | 0.31 | 7.34E-231 |
| TDO2 | 4.84E-168 | -0.29004 | 0.001 | 0.147 | 1.84E-163 |
| EIF4G3 | 4.98E-194 | -0.29014 | 0.21 | 0.499 | 1.89E-189 |
| SEMA6D | 1.70E-284 | -0.29074 | 0.009 | 0.253 | 6.44E-280 |
| PSMD1 | 2.70E-177 | -0.29111 | 0.301 | 0.596 | 1.03E-172 |
| IGHM | 9.93E-204 | -0.29126 | 0 | 0.173 | 3.77E-199 |
| LINC01503 | 8.74E-237 | -0.29197 | 0.041 | 0.28 | 3.32E-232 |
| ZCRB1 | 5.02E-169 | -0.292 | 0.453 | 0.75 | 1.91E-164 |
| PGRMC1 | 4.20E-134 | -0.29222 | 0.651 | 0.87 | 1.60E-129 |
| PRDM6 | 9.33E-243 | -0.29236 | 0.037 | 0.281 | 3.54E-238 |
| ALDH9A1 | 3.46E-168 | -0.29256 | 0.307 | 0.598 | 1.31E-163 |
| SH3KBP1 | 5.22E-202 | -0.29271 | 0.23 | 0.536 | 1.98E-197 |
| GOLGA3 | 1.31E-223 | -0.29308 | 0.173 | 0.478 | 4.96E-219 |
| CTNNAL1 | 7.69E-185 | -0.29313 | 0.219 | 0.504 | 2.92E-180 |
| BANF1 | 4.48E-148 | -0.2937 | 0.55 | 0.802 | 1.70E-143 |
| PNN | 4.97E-147 | -0.29379 | 0.585 | 0.837 | 1.89E-142 |
| TES | 6.13E-157 | -0.29401 | 0.228 | 0.485 | 2.33E-152 |
| PTGFR | 6.75E-60 | -0.29405 | 0.381 | 0.546 | 2.56E-55 |
| DDX46 | 1.44E-164 | -0.2948 | 0.474 | 0.765 | 5.46E-160 |
| WISP1 | 6.68E-221 | -0.295 | 0.024 | 0.236 | 2.54E-216 |
| SPCS3 | 2.35E-171 | -0.29564 | 0.387 | 0.683 | 8.93E-167 |
| SEC62 | 4.55E-145 | -0.29566 | 0.843 | 0.968 | 1.73E-140 |
| SUB1 | 1.23E-138 | -0.296 | 0.736 | 0.91 | 4.67E-134 |
| ITPR2 | 7.44E-218 | -0.29652 | 0.147 | 0.44 | 2.82E-213 |
| CCDC25 | 1.43E-166 | -0.29654 | 0.379 | 0.676 | 5.42E-162 |
| EMP2 | 3.11E-148 | -0.2966 | 0.478 | 0.77 | 1.18E-143 |
| FKBP7 | 7.78E-188 | -0.29713 | 0.318 | 0.623 | 2.95E-183 |
| NDUFB11 | 3.19E-149 | -0.29756 | 0.658 | 0.886 | 1.21E-144 |
| AKAP12 | 8.85E-17 | -0.29765 | 0.341 | 0.413 | 3.36E-12 |
| ADAMTS2 | 8.16E-126 | -0.29769 | 0.272 | 0.498 | 3.10E-121 |
| MESD | 4.87E-157 | -0.2984 | 0.571 | 0.837 | 1.85E-152 |
| NDUFA11 | 1.80E-159 | -0.2984 | 0.717 | 0.917 | 6.83E-155 |

| | | | | | |
|---|---|---|---|---|---|
| TPM3 | 2.96E-138 | -0.29849 | 0.514 | 0.764 | 1.12E-133 |
| PDIA5 | 1.08E-270 | -0.29855 | 0.096 | 0.399 | 4.10E-266 |
| PAPLN | 1.19E-243 | -0.29897 | 0.047 | 0.297 | 4.54E-239 |
| VGLL4 | 8.27E-117 | -0.29964 | 0.452 | 0.689 | 3.14E-112 |
| GCC2 | 5.98E-138 | -0.29972 | 0.519 | 0.781 | 2.27E-133 |
| LYZ | 6.97E-305 | -0.30076 | 0.005 | 0.257 | 2.65E-300 |
| ATP5MC3 | 1.39E-148 | -0.30126 | 0.773 | 0.934 | 5.29E-144 |
| ATM | 1.03E-211 | -0.30149 | 0.222 | 0.534 | 3.90E-207 |
| WASF2 | 2.12E-159 | -0.30161 | 0.64 | 0.898 | 8.07E-155 |
| RAMP3 | 1.56E-175 | -0.30174 | 0.006 | 0.165 | 5.92E-171 |
| SH3YL1 | 3.39E-259 | -0.30233 | 0.108 | 0.411 | 1.29E-254 |
| PCM1 | 1.38E-149 | -0.30312 | 0.492 | 0.765 | 5.22E-145 |
| SEC63 | 1.46E-168 | -0.3034 | 0.445 | 0.742 | 5.53E-164 |
| TRIM56 | 1.09E-174 | -0.30341 | 0.234 | 0.516 | 4.13E-170 |
| FBN1 | 5.99E-133 | -0.30353 | 0.79 | 0.953 | 2.28E-128 |
| TOP2B | 3.48E-214 | -0.30375 | 0.232 | 0.547 | 1.32E-209 |
| NKTR | 3.42E-159 | -0.30526 | 0.537 | 0.809 | 1.30E-154 |
| ACADVL | 5.24E-157 | -0.30569 | 0.594 | 0.852 | 1.99E-152 |
| ZNF385B | 6.89E-142 | -0.30576 | 0.262 | 0.527 | 2.62E-137 |
| FNDC1 | 3.67E-48 | -0.30578 | 0.23 | 0.357 | 1.39E-43 |
| NTNG1 | 2.63E-252 | -0.3058 | 0.008 | 0.227 | 1.00E-247 |
| PJA2 | 6.06E-149 | -0.30626 | 0.553 | 0.803 | 2.30E-144 |
| CNIH1 | 1.03E-169 | -0.30644 | 0.426 | 0.72 | 3.91E-165 |
| MYOF | 6.30E-176 | -0.30715 | 0.326 | 0.63 | 2.39E-171 |
| SETX | 8.79E-200 | -0.30865 | 0.247 | 0.553 | 3.34E-195 |
| ZCCHC24 | 1.94E-199 | -0.30869 | 0.218 | 0.511 | 7.36E-195 |
| WNT5B | 2.47E-251 | -0.30884 | 0.044 | 0.298 | 9.37E-247 |
| SKI | 2.99E-212 | -0.30886 | 0.223 | 0.536 | 1.14E-207 |
| UBE2L6 | 2.00E-190 | -0.30903 | 0.249 | 0.538 | 7.58E-186 |
| ATP5MF | 5.44E-163 | -0.30914 | 0.683 | 0.882 | 2.07E-158 |
| GBP1 | 1.72E-101 | -0.30953 | 0.156 | 0.337 | 6.53E-97 |
| RPN1 | 1.06E-175 | -0.31036 | 0.393 | 0.686 | 4.01E-171 |
| BEND6 | 1.53E-288 | -0.31045 | 0.045 | 0.326 | 5.80E-284 |
| ABHD14A | 1.85E-217 | -0.31082 | 0.237 | 0.559 | 7.03E-213 |
| CHD9 | 1.09E-158 | -0.31096 | 0.589 | 0.856 | 4.13E-154 |
| CKAP4 | 4.55E-175 | -0.31103 | 0.333 | 0.624 | 1.73E-170 |
| MAP4 | 1.09E-165 | -0.31105 | 0.468 | 0.749 | 4.14E-161 |
| GOLGA4 | 4.23E-144 | -0.31117 | 0.588 | 0.833 | 1.61E-139 |
| GNG12 | 3.18E-145 | -0.31127 | 0.486 | 0.76 | 1.21E-140 |
| SPCS1 | 1.17E-160 | -0.31144 | 0.779 | 0.944 | 4.45E-156 |
| CBX6 | 7.58E-218 | -0.312 | 0.209 | 0.521 | 2.88E-213 |
| MPG | 4.27E-160 | -0.3121 | 0.531 | 0.794 | 1.62E-155 |
| OSR2 | 2.32E-111 | -0.31253 | 0.462 | 0.698 | 8.83E-107 |
| DYNLT1 | 3.15E-157 | -0.31258 | 0.556 | 0.806 | 1.20E-152 |
| SELENOF | 4.54E-158 | -0.31298 | 0.726 | 0.912 | 1.72E-153 |
| MARCKSL1 | 2.46E-169 | -0.31309 | 0.081 | 0.292 | 9.35E-165 |
| PON2 | 1.62E-221 | -0.3139 | 0.146 | 0.428 | 6.15E-217 |

| | | | | | |
|---|---|---|---|---|---|
| DLX3 | 4.80E-283 | -0.31402 | 0.035 | 0.307 | 1.82E-278 |
| FTO | 3.53E-231 | -0.31436 | 0.135 | 0.424 | 1.34E-226 |
| PHF14 | 2.81E-185 | -0.31451 | 0.384 | 0.697 | 1.07E-180 |
| PLPP5 | 1.13E-215 | -0.31452 | 0.222 | 0.534 | 4.28E-211 |
| IKBIP | 4.66E-170 | -0.31461 | 0.39 | 0.685 | 1.77E-165 |
| HLTF | 3.43E-253 | -0.31467 | 0.125 | 0.434 | 1.30E-248 |
| HSPG2 | 2.06E-185 | -0.3148 | 0.47 | 0.798 | 7.84E-181 |
| PLIN3 | 3.07E-160 | -0.31492 | 0.524 | 0.788 | 1.16E-155 |
| KIAA1324L | 1.44E-245 | -0.31519 | 0.045 | 0.296 | 5.48E-241 |
| DPP7 | 1.80E-152 | -0.31582 | 0.704 | 0.913 | 6.82E-148 |
| HLA-F | 2.48E-171 | -0.31648 | 0.235 | 0.517 | 9.43E-167 |
| ZNF608 | 7.80E-259 | -0.3173 | 0.071 | 0.351 | 2.96E-254 |
| HNRNPU | 5.42E-130 | -0.31745 | 0.745 | 0.917 | 2.06E-125 |
| REX1BD | 2.23E-176 | -0.31765 | 0.521 | 0.808 | 8.47E-172 |
| TAGLN2 | 8.73E-86 | -0.31766 | 0.793 | 0.911 | 3.31E-81 |
| PDE4B | 1.91E-226 | -0.31853 | 0.093 | 0.365 | 7.27E-222 |
| STMP1 | 2.04E-157 | -0.31891 | 0.539 | 0.794 | 7.75E-153 |
| TIMP3 | 8.33E-14 | -0.31981 | 0.904 | 0.93 | 3.16E-09 |
| CREB3L2 | 5.27E-212 | -0.32006 | 0.28 | 0.607 | 2.00E-207 |
| SUMO2 | 8.99E-190 | -0.32076 | 0.918 | 0.98 | 3.41E-185 |
| TSHZ2 | 1.71E-119 | -0.32207 | 0.556 | 0.769 | 6.51E-115 |
| PRMT2 | 1.31E-211 | -0.32212 | 0.301 | 0.628 | 4.98E-207 |
| DBP | 1.92E-168 | -0.32241 | 0.12 | 0.352 | 7.29E-164 |
| GNAL | 6.39E-249 | -0.32356 | 0.059 | 0.32 | 2.43E-244 |
| CAPRIN1 | 5.02E-207 | -0.32392 | 0.311 | 0.624 | 1.91E-202 |
| PRRX1 | 2.52E-122 | -0.3243 | 0.657 | 0.871 | 9.58E-118 |
| FCGRT | 2.22E-163 | -0.32451 | 0.836 | 0.977 | 8.45E-159 |
| FSTL1 | 6.94E-139 | -0.32573 | 0.906 | 0.985 | 2.64E-134 |
| RSRP1 | 1.42E-152 | -0.32575 | 0.587 | 0.844 | 5.38E-148 |
| SEMA3E | 3.61E-179 | -0.32603 | 0.158 | 0.42 | 1.37E-174 |
| SHPRH | 1.95E-199 | -0.32605 | 0.203 | 0.495 | 7.40E-195 |
| ALDH3A2 | 1.18E-195 | -0.32625 | 0.3 | 0.614 | 4.46E-191 |
| PCNX4 | 2.07E-248 | -0.32687 | 0.118 | 0.416 | 7.85E-244 |
| FAM20C | 1.19E-167 | -0.32694 | 0.366 | 0.658 | 4.50E-163 |
| GALNT11 | 1.19E-218 | -0.32735 | 0.28 | 0.604 | 4.51E-214 |
| DNPH1 | 2.34E-238 | -0.32762 | 0.218 | 0.538 | 8.89E-234 |
| PSMA2 | 1.60E-196 | -0.32804 | 0.399 | 0.701 | 6.07E-192 |
| CXCL16 | 3.98E-277 | -0.32829 | 0.104 | 0.414 | 1.51E-272 |
| FSCN1 | 3.77E-279 | -0.32945 | 0.076 | 0.372 | 1.43E-274 |
| ZDBF2 | 1.54E-253 | -0.3299 | 0.049 | 0.307 | 5.86E-249 |
| TMEM260 | 6.35E-242 | -0.33005 | 0.092 | 0.369 | 2.41E-237 |
| CEP290 | 2.39E-214 | -0.33058 | 0.139 | 0.42 | 9.06E-210 |
| ID1 | 1.83E-73 | -0.33068 | 0.312 | 0.491 | 6.94E-69 |
| DNAJC15 | 3.31E-180 | -0.33075 | 0.442 | 0.731 | 1.26E-175 |
| SPTAN1 | 2.66E-182 | -0.33098 | 0.361 | 0.664 | 1.01E-177 |
| CBX5 | 1.17E-204 | -0.33118 | 0.335 | 0.648 | 4.43E-200 |
| BMP1 | 3.26E-234 | -0.33172 | 0.198 | 0.511 | 1.24E-229 |

| | | | | | |
|---|---|---|---|---|---|
| P3H4 | 1.47E-273 | -0.3319 | 0.122 | 0.438 | 5.59E-269 |
| MEST | 2.13E-204 | -0.33234 | 0.022 | 0.219 | 8.08E-200 |
| ETFB | 8.37E-190 | -0.33326 | 0.561 | 0.834 | 3.18E-185 |
| ASPH | 5.41E-145 | -0.33352 | 0.642 | 0.88 | 2.05E-140 |
| CHD3 | 1.42E-238 | -0.33376 | 0.169 | 0.477 | 5.40E-234 |
| SCP2 | 2.63E-170 | -0.33498 | 0.635 | 0.878 | 9.97E-166 |
| KDELR1 | 2.90E-175 | -0.33531 | 0.713 | 0.913 | 1.10E-170 |
| ABHD2 | 1.80E-192 | -0.3365 | 0.207 | 0.49 | 6.83E-188 |
| TRIM44 | 3.48E-230 | -0.33741 | 0.253 | 0.581 | 1.32E-225 |
| DNASE1L3 | 1.81E-185 | -0.33748 | 0.007 | 0.175 | 6.88E-181 |
| MYL9 | 4.80E-153 | -0.33761 | 0.576 | 0.847 | 1.82E-148 |
| HEG1 | 2.95E-230 | -0.33867 | 0.144 | 0.44 | 1.12E-225 |
| ZNF428 | 1.47E-205 | -0.34007 | 0.415 | 0.729 | 5.57E-201 |
| SMARCA2 | 1.23E-226 | -0.34214 | 0.286 | 0.623 | 4.68E-222 |
| DLC1 | 1.46E-163 | -0.34237 | 0.441 | 0.724 | 5.53E-159 |
| RNASEH2C | 1.00E-193 | -0.34253 | 0.506 | 0.796 | 3.81E-189 |
| HTATSF1 | 2.53E-244 | -0.34273 | 0.228 | 0.56 | 9.60E-240 |
| LUC7L3 | 1.61E-182 | -0.34292 | 0.601 | 0.87 | 6.12E-178 |
| SCARB2 | 2.41E-183 | -0.34307 | 0.543 | 0.818 | 9.14E-179 |
| MT-CO3 | 1.28E-198 | -0.3441 | 1 | 1 | 4.86E-194 |
| SORBS2 | 2.52E-246 | -0.34436 | 0.189 | 0.518 | 9.58E-242 |
| EIF2AK2 | 3.72E-227 | -0.34443 | 0.274 | 0.599 | 1.41E-222 |
| TUBA1A | 7.74E-159 | -0.34447 | 0.572 | 0.849 | 2.94E-154 |
| ARHGAP2( | 3.74E-160 | -0.34507 | 0.248 | 0.512 | 1.42E-155 |
| CDC42BPA | 1.36E-233 | -0.34561 | 0.221 | 0.543 | 5.16E-229 |
| SERTAD4-/ | 6.67E-247 | -0.3466 | 0.095 | 0.381 | 2.53E-242 |
| NSD1 | 2.82E-270 | -0.34768 | 0.18 | 0.52 | 1.07E-265 |
| AKR1C1 | 1.37E-152 | -0.34783 | 0.485 | 0.761 | 5.21E-148 |
| LNX1 | 1.63E-222 | -0.34823 | 0.165 | 0.459 | 6.19E-218 |
| GABPB1-A | 2.84E-147 | -0.34832 | 0.326 | 0.583 | 1.08E-142 |
| NPR3 | 4.11E-126 | -0.34898 | 0.025 | 0.161 | 1.56E-121 |
| AC020916. | 1.03E-93 | -0.34911 | 0.17 | 0.341 | 3.92E-89 |
| TM9SF3 | 3.05E-186 | -0.34917 | 0.562 | 0.838 | 1.16E-181 |
| C1QTNF4 | 1.00E-98 | -0.34921 | 0.088 | 0.24 | 3.81E-94 |
| CLMP | 7.12E-182 | -0.34926 | 0.393 | 0.68 | 2.70E-177 |
| TMEM108 | 6.24E-234 | -0.34938 | 0.065 | 0.318 | 2.37E-229 |
| SMARCA1 | 5.78E-290 | -0.34993 | 0.155 | 0.499 | 2.19E-285 |
| ENPP1 | 3.76E-92 | -0.35021 | 0.241 | 0.435 | 1.43E-87 |
| HTR2A | 1.22E-294 | -0.35032 | 0.041 | 0.323 | 4.62E-290 |
| TPR | 5.21E-181 | -0.35114 | 0.579 | 0.848 | 1.98E-176 |
| 2-Sep | 4.89E-190 | -0.3515 | 0.652 | 0.884 | 1.86E-185 |
| MYH9 | 2.64E-201 | -0.35167 | 0.287 | 0.595 | 1.00E-196 |
| PLXNB2 | 1.19E-280 | -0.35178 | 0.138 | 0.471 | 4.53E-276 |
| GOLGA2 | 1.81E-224 | -0.35179 | 0.373 | 0.703 | 6.87E-220 |
| CMBL | 1.49E-256 | -0.3521 | 0.186 | 0.52 | 5.68E-252 |
| ATP5MC1 | 5.18E-182 | -0.35226 | 0.606 | 0.858 | 1.97E-177 |
| GCSH | 6.14E-215 | -0.35296 | 0.32 | 0.63 | 2.33E-210 |

| | | | | | |
|---|---|---|---|---|---|
| EDF1 | 6.53E-224 | -0.35347 | 0.885 | 0.98 | 2.48E-219 |
| RUNX1 | 5.21E-165 | -0.35374 | 0.385 | 0.67 | 1.98E-160 |
| P3H1 | 1.54E-262 | -0.35393 | 0.177 | 0.499 | 5.84E-258 |
| TLN1 | 2.32E-189 | -0.35435 | 0.538 | 0.812 | 8.80E-185 |
| IFI44L | 4.68E-239 | -0.35462 | 0.059 | 0.314 | 1.78E-234 |
| ERP29 | 6.59E-189 | -0.35547 | 0.639 | 0.881 | 2.50E-184 |
| DYNC1H1 | 6.23E-198 | -0.35602 | 0.462 | 0.77 | 2.37E-193 |
| KCND3 | 0 | -0.35611 | 0.03 | 0.318 | 0 |
| PCSK1 | 2.77E-218 | -0.35633 | 0.012 | 0.212 | 1.05E-213 |
| PARK7 | 4.02E-203 | -0.35738 | 0.862 | 0.974 | 1.53E-198 |
| GNG11 | 2.85E-187 | -0.35819 | 0.47 | 0.78 | 1.08E-182 |
| RPL3 | 7.66E-255 | -0.35951 | 0.997 | 0.999 | 2.91E-250 |
| NCL | 2.03E-173 | -0.3604 | 0.787 | 0.949 | 7.69E-169 |
| ARMCX3 | 7.72E-217 | -0.36114 | 0.384 | 0.708 | 2.93E-212 |
| BICC1 | 3.06E-161 | -0.3617 | 0.367 | 0.637 | 1.16E-156 |
| BPTF | 6.94E-195 | -0.36193 | 0.438 | 0.74 | 2.64E-190 |
| UBE2E3 | 5.26E-213 | -0.362 | 0.476 | 0.782 | 2.00E-208 |
| EID1 | 2.09E-251 | -0.36219 | 0.919 | 0.989 | 7.93E-247 |
| GSTK1 | 1.87E-227 | -0.36264 | 0.391 | 0.726 | 7.10E-223 |
| ERLEC1 | 5.91E-202 | -0.36277 | 0.489 | 0.786 | 2.24E-197 |
| SYTL2 | 2.88E-273 | -0.3633 | 0.093 | 0.393 | 1.09E-268 |
| MBTPS1 | 8.03E-242 | -0.36373 | 0.357 | 0.698 | 3.05E-237 |
| FKBP10 | 1.84E-189 | -0.3654 | 0.551 | 0.812 | 7.00E-185 |
| PLAGL1 | 3.66E-186 | -0.36567 | 0.407 | 0.705 | 1.39E-181 |
| S100B | 1.33E-136 | -0.36648 | 0.036 | 0.191 | 5.05E-132 |
| SELENOP | 6.98E-129 | -0.3668 | 0.767 | 0.92 | 2.65E-124 |
| LAMB1 | 1.10E-192 | -0.36738 | 0.426 | 0.728 | 4.19E-188 |
| SFRP4 | 1.86E-05 | -0.36828 | 0.276 | 0.245 | 0.708131 |
| COL16A1 | 2.80E-194 | -0.36867 | 0.299 | 0.597 | 1.06E-189 |
| TCEAL4 | 5.96E-205 | -0.36934 | 0.581 | 0.856 | 2.26E-200 |
| KAZALD1 | 0 | -0.36935 | 0.076 | 0.399 | 0 |
| GTF3C6 | 8.53E-256 | -0.3712 | 0.322 | 0.667 | 3.24E-251 |
| SGCB | 6.26E-219 | -0.37131 | 0.361 | 0.684 | 2.38E-214 |
| FAT1 | 4.87E-295 | -0.37135 | 0.096 | 0.414 | 1.85E-290 |
| RBFOX2 | 1.17E-244 | -0.37199 | 0.334 | 0.683 | 4.45E-240 |
| CAVIN3 | 7.55E-203 | -0.37208 | 0.517 | 0.847 | 2.87E-198 |
| SPCS2 | 6.80E-209 | -0.37311 | 0.835 | 0.962 | 2.58E-204 |
| TAPBP | 3.55E-204 | -0.37349 | 0.378 | 0.687 | 1.35E-199 |
| HLA-DQB1 | 7.21E-243 | -0.37359 | 0.011 | 0.228 | 2.74E-238 |
| SCPEP1 | 1.56E-183 | -0.37383 | 0.553 | 0.824 | 5.92E-179 |
| NOTCH2 | 2.00E-229 | -0.37435 | 0.304 | 0.633 | 7.60E-225 |
| XAF1 | 7.48E-259 | -0.3759 | 0.123 | 0.43 | 2.84E-254 |
| EDIL3 | 6.07E-118 | -0.37648 | 0.109 | 0.28 | 2.31E-113 |
| MAPK10 | 5.73E-296 | -0.37659 | 0.056 | 0.354 | 2.18E-291 |
| ZFHX4 | 8.43E-227 | -0.37705 | 0.231 | 0.55 | 3.20E-222 |
| C9orf3 | 2.18E-259 | -0.37732 | 0.215 | 0.556 | 8.29E-255 |
| SH3PXD2A | 3.39E-243 | -0.3775 | 0.161 | 0.466 | 1.29E-238 |

| | | | | | |
|---|---|---|---|---|---|
| AKR1A1 | 1.77E-240 | -0.37754 | 0.426 | 0.757 | 6.74E-236 |
| BNC2 | 6.86E-257 | -0.37781 | 0.147 | 0.457 | 2.60E-252 |
| CRNDE | 9.04E-223 | -0.37792 | 0.374 | 0.704 | 3.43E-218 |
| NAV1 | 2.05E-216 | -0.37864 | 0.326 | 0.644 | 7.77E-212 |
| TRIP11 | 1.12E-204 | -0.37871 | 0.389 | 0.701 | 4.26E-200 |
| CFH | 3.16E-182 | -0.37974 | 0.777 | 0.969 | 1.20E-177 |
| RGS16 | 8.49E-13 | -0.37993 | 0.228 | 0.179 | 3.22E-08 |
| FLNA | 2.46E-203 | -0.38033 | 0.502 | 0.807 | 9.34E-199 |
| UBTF | 1.54E-276 | -0.3805 | 0.249 | 0.607 | 5.85E-272 |
| AHI1 | 2.54E-233 | -0.38149 | 0.428 | 0.764 | 9.63E-229 |
| CHSY1 | 1.43E-276 | -0.38172 | 0.132 | 0.448 | 5.41E-272 |
| 7-Sep | 1.46E-210 | -0.38207 | 0.827 | 0.963 | 5.53E-206 |
| GNAI2 | 5.99E-217 | -0.38331 | 0.634 | 0.888 | 2.28E-212 |
| APOL6 | 5.34E-250 | -0.38336 | 0.2 | 0.526 | 2.03E-245 |
| TMED9 | 5.61E-205 | -0.38346 | 0.724 | 0.914 | 2.13E-200 |
| CMKLR1 | 0 | -0.38424 | 0.046 | 0.358 | 0 |
| SGCD | 0 | -0.38426 | 0.079 | 0.419 | 0 |
| CALM2 | 6.24E-207 | -0.38426 | 0.883 | 0.977 | 2.37E-202 |
| FNDC3A | 1.33E-269 | -0.38638 | 0.275 | 0.629 | 5.04E-265 |
| SHC1 | 3.47E-282 | -0.38643 | 0.193 | 0.536 | 1.32E-277 |
| KYNU | 1.02E-265 | -0.38685 | 0.056 | 0.328 | 3.88E-261 |
| ARL1 | 2.20E-237 | -0.38782 | 0.48 | 0.786 | 8.34E-233 |
| CH25H | 1.68E-118 | -0.38837 | 0.015 | 0.137 | 6.38E-114 |
| NDUFB3 | 8.86E-272 | -0.38865 | 0.418 | 0.759 | 3.36E-267 |
| UQCC2 | 1.13E-281 | -0.38955 | 0.262 | 0.62 | 4.27E-277 |
| CTNNB1 | 3.96E-163 | -0.39032 | 0.479 | 0.738 | 1.50E-158 |
| GXYLT2 | 2.88E-295 | -0.39081 | 0.079 | 0.382 | 1.09E-290 |
| NDUFB7 | 5.72E-246 | -0.39122 | 0.693 | 0.923 | 2.17E-241 |
| ZBTB20 | 9.88E-181 | -0.39154 | 0.725 | 0.929 | 3.75E-176 |
| OST4 | 4.84E-221 | -0.39222 | 0.874 | 0.974 | 1.84E-216 |
| SERPINF1 | 6.18E-76 | -0.3923 | 0.91 | 0.927 | 2.35E-71 |
| SNRPN | 5.26E-263 | -0.39251 | 0.319 | 0.666 | 2.00E-258 |
| SARAF | 4.12E-217 | -0.39303 | 0.699 | 0.918 | 1.57E-212 |
| AKR1C2 | 2.58E-165 | -0.3934 | 0.348 | 0.636 | 9.78E-161 |
| HACD3 | 4.13E-305 | -0.39443 | 0.23 | 0.598 | 1.57E-300 |
| CUX1 | 3.44E-235 | -0.39462 | 0.315 | 0.639 | 1.31E-230 |
| PLEKHA4 | 5.36E-284 | -0.39493 | 0.195 | 0.54 | 2.03E-279 |
| TMEM98 | 3.60E-230 | -0.3956 | 0.293 | 0.615 | 1.37E-225 |
| LRRC15 | 4.02E-302 | -0.39584 | 0.001 | 0.248 | 1.53E-297 |
| KIF5B | 3.24E-238 | -0.39618 | 0.399 | 0.72 | 1.23E-233 |
| PDLIM4 | 4.37E-227 | -0.39643 | 0.421 | 0.729 | 1.66E-222 |
| ISOC2 | 1.95E-282 | -0.39668 | 0.258 | 0.612 | 7.39E-278 |
| HSBP1 | 6.61E-232 | -0.39706 | 0.637 | 0.888 | 2.51E-227 |
| CPXM1 | 1.82E-237 | -0.39718 | 0.013 | 0.225 | 6.92E-233 |
| MAN1A2 | 7.64E-293 | -0.39722 | 0.247 | 0.609 | 2.90E-288 |
| SET | 4.36E-218 | -0.39762 | 0.663 | 0.903 | 1.66E-213 |
| FLRT2 | 1.63E-307 | -0.39801 | 0.088 | 0.408 | 6.19E-303 |

| | | | | | |
|---|---|---|---|---|---|
| SSR1 | 3.39E-264 | -0.39813 | 0.339 | 0.682 | 1.29E-259 |
| PLEC | 2.42E-270 | -0.39831 | 0.245 | 0.598 | 9.19E-266 |
| CASC4 | 1.22E-264 | -0.39842 | 0.308 | 0.658 | 4.62E-260 |
| ADGRB3 | 0 | -0.39871 | 0.018 | 0.332 | 0 |
| OLFML1 | 3.77E-303 | -0.39924 | 0.147 | 0.498 | 1.43E-298 |
| SMC4 | 0 | -0.39944 | 0.083 | 0.4 | 0 |
| EXT1 | 3.83E-294 | -0.39949 | 0.164 | 0.504 | 1.45E-289 |
| SIL1 | 1.22E-265 | -0.40033 | 0.324 | 0.674 | 4.62E-261 |
| MAF | 5.49E-245 | -0.40082 | 0.238 | 0.568 | 2.09E-240 |
| TMEM119 | 0 | -0.40171 | 0.033 | 0.319 | 0 |
| CERCAM | 2.29E-251 | -0.40258 | 0.284 | 0.624 | 8.72E-247 |
| LMAN1 | 2.44E-220 | -0.40298 | 0.612 | 0.876 | 9.26E-216 |
| PPP1CC | 7.30E-178 | -0.40327 | 0.465 | 0.731 | 2.77E-173 |
| MACF1 | 7.57E-230 | -0.40334 | 0.394 | 0.716 | 2.88E-225 |
| BBX | 1.68E-258 | -0.40456 | 0.363 | 0.723 | 6.40E-254 |
| PSMB8 | 3.84E-229 | -0.40461 | 0.443 | 0.763 | 1.46E-224 |
| CCDC88A | 6.35E-281 | -0.40466 | 0.209 | 0.559 | 2.41E-276 |
| ARHGAP28 | 0 | -0.40581 | 0.048 | 0.382 | 0 |
| DNM3OS | 0 | -0.40591 | 0.088 | 0.437 | 0 |
| PTPRD | 0 | -0.40595 | 0.119 | 0.459 | 0 |
| TMEM230 | 8.02E-240 | -0.40666 | 0.629 | 0.886 | 3.04E-235 |
| NREP | 0 | -0.40695 | 0.019 | 0.349 | 0 |
| TCAF1 | 0 | -0.40712 | 0.145 | 0.507 | 0 |
| TMCO3 | 0 | -0.40733 | 0.213 | 0.583 | 0 |
| PRKDC | 1.25E-236 | -0.40823 | 0.443 | 0.764 | 4.77E-232 |
| TNFRSF12A | 8.72E-148 | -0.40879 | 0.158 | 0.379 | 3.31E-143 |
| TCEAL3 | 4.11E-282 | -0.40953 | 0.256 | 0.614 | 1.56E-277 |
| MYDGF | 2.74E-213 | -0.40972 | 0.764 | 0.94 | 1.04E-208 |
| GLCCI1 | 0 | -0.41019 | 0.052 | 0.37 | 0 |
| MRPL51 | 3.00E-245 | -0.41033 | 0.642 | 0.898 | 1.14E-240 |
| NUCB1 | 1.75E-230 | -0.4114 | 0.652 | 0.897 | 6.64E-226 |
| CCPG1 | 1.64E-226 | -0.41164 | 0.627 | 0.882 | 6.24E-222 |
| ST5 | 2.73E-280 | -0.41219 | 0.209 | 0.561 | 1.04E-275 |
| TRAPPC1 | 1.35E-287 | -0.41281 | 0.371 | 0.735 | 5.14E-283 |
| SSR2 | 6.18E-253 | -0.41284 | 0.838 | 0.968 | 2.35E-248 |
| NFIX | 2.00E-247 | -0.41365 | 0.5 | 0.828 | 7.58E-243 |
| ANK3 | 7.96E-300 | -0.41375 | 0.011 | 0.268 | 3.02E-295 |
| ZC3H13 | 9.08E-262 | -0.41476 | 0.34 | 0.691 | 3.45E-257 |
| AP2S1 | 1.46E-225 | -0.41491 | 0.593 | 0.852 | 5.53E-221 |
| FAM20A | 8.79E-264 | -0.41573 | 0.183 | 0.504 | 3.34E-259 |
| GUK1 | 4.37E-234 | -0.41702 | 0.809 | 0.956 | 1.66E-229 |
| ODF2L | 2.94E-261 | -0.41714 | 0.296 | 0.648 | 1.12E-256 |
| DNAJC10 | 1.14E-297 | -0.41826 | 0.259 | 0.625 | 4.32E-293 |
| SH3BGRL | 3.40E-262 | -0.42003 | 0.574 | 0.864 | 1.29E-257 |
| PDLIM2 | 1.48E-238 | -0.42005 | 0.428 | 0.755 | 5.62E-234 |
| SAMD11 | 0 | -0.42037 | 0.017 | 0.311 | 0 |
| GPX7 | 0 | -0.42052 | 0.102 | 0.469 | 0 |

| | | | | | |
|---|---|---|---|---|---|
| RAB31 | 0 | -0.4207 | 0.16 | 0.514 | 0 |
| PSD3 | 9.29E-268 | -0.4208 | 0.252 | 0.6 | 3.53E-263 |
| SDC2 | 1.68E-154 | -0.42315 | 0.687 | 0.886 | 6.37E-150 |
| SCARA3 | 7.66E-168 | -0.42516 | 0.283 | 0.549 | 2.91E-163 |
| CALM3 | 3.49E-270 | -0.42609 | 0.397 | 0.745 | 1.32E-265 |
| PLD3 | 2.30E-248 | -0.42624 | 0.641 | 0.902 | 8.73E-244 |
| STAT3 | 2.99E-218 | -0.42665 | 0.67 | 0.896 | 1.14E-213 |
| AP2M1 | 9.51E-261 | -0.42678 | 0.59 | 0.869 | 3.61E-256 |
| NDUFC2 | 3.34E-268 | -0.42736 | 0.708 | 0.94 | 1.27E-263 |
| C4orf48 | 0 | -0.42761 | 0.079 | 0.41 | 0 |
| AEBP1 | 1.08E-90 | -0.42945 | 0.743 | 0.842 | 4.11E-86 |
| EIF4G2 | 1.08E-252 | -0.42952 | 0.646 | 0.89 | 4.11E-248 |
| SLC16A4 | 0 | -0.42971 | 0.131 | 0.499 | 0 |
| VWA1 | 0 | -0.43127 | 0.104 | 0.473 | 0 |
| CRELD2 | 4.33E-297 | -0.43166 | 0.284 | 0.65 | 1.64E-292 |
| RPN2 | 7.81E-254 | -0.43222 | 0.627 | 0.889 | 2.97E-249 |
| TUBB | 5.69E-211 | -0.43288 | 0.69 | 0.912 | 2.16E-206 |
| FUS | 2.61E-236 | -0.43305 | 0.737 | 0.942 | 9.91E-232 |
| FGF7 | 5.85E-101 | -0.43376 | 0.415 | 0.632 | 2.22E-96 |
| CDC42EP5 | 0 | -0.43382 | 0.245 | 0.633 | 0 |
| RNASET2 | 0 | -0.43463 | 0.102 | 0.51 | 0 |
| S100A4 | 2.17E-144 | -0.43675 | 0.949 | 0.993 | 8.26E-140 |
| VEGFC | 4.94E-302 | -0.43675 | 0.04 | 0.322 | 1.88E-297 |
| ABI3BP | 8.82E-186 | -0.4374 | 0.794 | 0.957 | 3.35E-181 |
| FAM13C | 0 | -0.43751 | 0.039 | 0.376 | 0 |
| COPB2 | 9.86E-283 | -0.43764 | 0.388 | 0.733 | 3.75E-278 |
| SEMA3A | 0 | -0.43867 | 0.016 | 0.295 | 0 |
| FTX | 1.32E-285 | -0.4388 | 0.282 | 0.65 | 5.00E-281 |
| PHACTR2 | 2.71E-276 | -0.43899 | 0.273 | 0.625 | 1.03E-271 |
| UACA | 8.01E-219 | -0.43926 | 0.352 | 0.68 | 3.04E-214 |
| ANXA2 | 1.93E-285 | -0.43993 | 0.963 | 0.995 | 7.32E-281 |
| ELK3 | 0 | -0.44091 | 0.184 | 0.553 | 0 |
| CAPZB | 1.75E-237 | -0.44121 | 0.675 | 0.9 | 6.63E-233 |
| SLC39A7 | 1.28E-289 | -0.44199 | 0.402 | 0.751 | 4.86E-285 |
| CD47 | 2.46E-282 | -0.44213 | 0.605 | 0.892 | 9.32E-278 |
| TSPAN3 | 2.41E-282 | -0.44388 | 0.411 | 0.756 | 9.14E-278 |
| LIMA1 | 3.70E-239 | -0.4447 | 0.7 | 0.938 | 1.40E-234 |
| DAD1 | 2.40E-301 | -0.44476 | 0.869 | 0.979 | 9.13E-297 |
| WLS | 1.50E-293 | -0.44527 | 0.223 | 0.581 | 5.70E-289 |
| ATP5IF1 | 1.53E-262 | -0.44562 | 0.595 | 0.885 | 5.81E-258 |
| CXXC5 | 2.17E-200 | -0.44584 | 0.335 | 0.629 | 8.24E-196 |
| MXD4 | 0 | -0.44618 | 0.284 | 0.664 | 0 |
| POLR2J3.1 | 0 | -0.44633 | 0.25 | 0.625 | 0 |
| SLC2A12 | 1.40E-299 | -0.4487 | 0.018 | 0.28 | 5.30E-295 |
| MSRB2 | 0 | -0.4488 | 0.352 | 0.726 | 0 |
| CD82 | 4.89E-287 | -0.44965 | 0.16 | 0.498 | 1.86E-282 |
| TCEAL8 | 0 | -0.4498 | 0.371 | 0.748 | 0 |

| | | | | | |
|---|---|---|---|---|---|
| AIG1 | 0 | -0.45016 | 0.27 | 0.657 | 0 |
| IGLC3 | 1.63E-233 | -0.45114 | 0 | 0.196 | 6.19E-229 |
| CHD4 | 4.73E-287 | -0.45136 | 0.32 | 0.68 | 1.80E-282 |
| P3H3 | 0 | -0.45215 | 0.172 | 0.58 | 0 |
| GLI3 | 0 | -0.45528 | 0.182 | 0.529 | 0 |
| ATP2B4 | 0 | -0.45593 | 0.218 | 0.61 | 0 |
| ATOX1 | 1.64E-285 | -0.45642 | 0.514 | 0.819 | 6.25E-281 |
| MYADM | 4.51E-157 | -0.45658 | 0.579 | 0.793 | 1.71E-152 |
| C1QTNF3 | 3.34E-134 | -0.4573 | 0.15 | 0.36 | 1.27E-129 |
| MT-CO1 | 0 | -0.45791 | 0.999 | 1 | 0 |
| PALMD | 2.53E-307 | -0.45854 | 0.286 | 0.672 | 9.60E-303 |
| ZMAT3 | 0 | -0.46031 | 0.078 | 0.399 | 0 |
| TMEM219 | 0 | -0.46239 | 0.441 | 0.803 | 0 |
| SPON1 | 3.68E-301 | -0.46254 | 0.021 | 0.286 | 1.40E-296 |
| CHST2 | 0 | -0.46299 | 0.073 | 0.389 | 0 |
| CTSS | 0 | -0.46351 | 0.108 | 0.45 | 0 |
| CALM1 | 9.22E-218 | -0.46427 | 0.813 | 0.957 | 3.50E-213 |
| HCFC1R1 | 1.48E-274 | -0.46452 | 0.35 | 0.706 | 5.60E-270 |
| HLA-A | 6.40E-196 | -0.46487 | 0.983 | 0.997 | 2.43E-191 |
| AL078639. | 0 | -0.46545 | 0.177 | 0.529 | 0 |
| SAMD9L | 0 | -0.46587 | 0.066 | 0.428 | 0 |
| FKBP14 | 0 | -0.46626 | 0.121 | 0.519 | 0 |
| DDX17 | 4.60E-272 | -0.46683 | 0.741 | 0.945 | 1.75E-267 |
| IGHG3 | 0 | -0.46871 | 0.001 | 0.298 | 0 |
| TMED10 | 5.36E-287 | -0.4688 | 0.712 | 0.928 | 2.04E-282 |
| SNAI2 | 7.69E-227 | -0.46921 | 0.246 | 0.557 | 2.92E-222 |
| TSC22D1 | 1.09E-119 | -0.4695 | 0.345 | 0.587 | 4.14E-115 |
| PLAC9 | 1.66E-209 | -0.47132 | 0.927 | 0.99 | 6.31E-205 |
| CEMIP | 6.00E-285 | -0.47141 | 0.011 | 0.258 | 2.28E-280 |
| AKR7A2 | 0 | -0.47165 | 0.321 | 0.703 | 0 |
| ADAMTS1 | 4.12E-58 | -0.47208 | 0.238 | 0.376 | 1.57E-53 |
| SESN3 | 0 | -0.4729 | 0.07 | 0.443 | 0 |
| HDDC2 | 5.37E-298 | -0.47377 | 0.409 | 0.756 | 2.04E-293 |
| SERTAD4 | 0 | -0.47391 | 0.033 | 0.331 | 0 |
| ARPC2 | 1.81E-285 | -0.47405 | 0.694 | 0.92 | 6.86E-281 |
| TNFSF13B | 0 | -0.47498 | 0.035 | 0.323 | 0 |
| PKM | 1.85E-239 | -0.47503 | 0.775 | 0.952 | 7.02E-235 |
| COL6A2 | 1.54E-297 | -0.4755 | 0.98 | 0.999 | 5.85E-293 |
| MT-ND1 | 4.69E-260 | -0.47746 | 0.998 | 1 | 1.78E-255 |
| EPHA3 | 0 | -0.478 | 0.036 | 0.4 | 0 |
| EML1 | 0 | -0.47838 | 0.145 | 0.544 | 0 |
| ITM2B | 0 | -0.4784 | 0.996 | 0.999 | 0 |
| S1PR3 | 7.17E-257 | -0.48278 | 0.133 | 0.431 | 2.72E-252 |
| CTSB | 4.26E-265 | -0.48364 | 0.602 | 0.898 | 1.62E-260 |
| NFIB | 2.00E-226 | -0.48374 | 0.565 | 0.851 | 7.60E-222 |
| CRABP2 | 5.61E-167 | -0.48494 | 0.096 | 0.309 | 2.13E-162 |
| MGST3 | 2.64E-307 | -0.48543 | 0.823 | 0.977 | 1.00E-302 |

| | | | | | |
|---|---|---|---|---|---|
| ZFP36L1 | 3.40E-113 | -0.48587 | 0.846 | 0.954 | 1.29E-108 |
| TMED3 | 0 | -0.48621 | 0.345 | 0.733 | 0 |
| PDLIM7 | 0 | -0.48669 | 0.15 | 0.548 | 0 |
| EPHX1 | 5.56E-207 | -0.48797 | 0.618 | 0.883 | 2.11E-202 |
| H3F3A | 0 | -0.48813 | 0.966 | 0.996 | 0 |
| NME3 | 0 | -0.48857 | 0.537 | 0.87 | 0 |
| NDUFA4 | 0 | -0.48926 | 0.868 | 0.978 | 0 |
| WWTR1 | 0 | -0.48973 | 0.191 | 0.572 | 0 |
| SEMA5A | 0 | -0.48978 | 0.018 | 0.347 | 0 |
| IL11RA | 0 | -0.48981 | 0.157 | 0.534 | 0 |
| SULF2 | 0 | -0.49381 | 0.275 | 0.651 | 0 |
| GLG1 | 0 | -0.49462 | 0.454 | 0.82 | 0 |
| CDON | 0 | -0.49602 | 0.145 | 0.561 | 0 |
| ZKSCAN1 | 0 | -0.4962 | 0.232 | 0.654 | 0 |
| METRN | 0 | -0.49677 | 0.167 | 0.537 | 0 |
| RCAN1 | 9.21E-269 | -0.49732 | 0.098 | 0.393 | 3.50E-264 |
| EPYC | 1.98E-125 | -0.49736 | 0.004 | 0.119 | 7.51E-121 |
| SIX1 | 2.05E-203 | -0.49806 | 0.294 | 0.592 | 7.79E-199 |
| CDK14 | 0 | -0.49912 | 0.096 | 0.529 | 0 |
| ZNF106 | 0 | -0.49918 | 0.321 | 0.698 | 0 |
| PARP14 | 0 | -0.50011 | 0.179 | 0.557 | 0 |
| GOLGB1 | 1.61E-287 | -0.50052 | 0.466 | 0.799 | 6.11E-283 |
| ATRX | 0 | -0.50123 | 0.512 | 0.84 | 0 |
| ASPN | 9.03E-19 | -0.50249 | 0.442 | 0.511 | 3.43E-14 |
| VAMP5 | 3.31E-305 | -0.50261 | 0.517 | 0.839 | 1.26E-300 |
| KCNE4 | 4.10E-194 | -0.50356 | 0.211 | 0.493 | 1.56E-189 |
| TBL1XR1 | 0 | -0.50413 | 0.32 | 0.706 | 0 |
| TSPO | 0 | -0.50795 | 0.768 | 0.966 | 0 |
| TPM4 | 5.94E-198 | -0.50827 | 0.627 | 0.854 | 2.26E-193 |
| PDE1A | 0 | -0.51149 | 0.097 | 0.532 | 0 |
| FKBP11 | 0 | -0.51186 | 0.162 | 0.573 | 0 |
| ANTXR1 | 0 | -0.51441 | 0.205 | 0.59 | 0 |
| CD248 | 1.02E-233 | -0.51476 | 0.432 | 0.76 | 3.88E-229 |
| UBB | 0 | -0.51538 | 0.887 | 0.986 | 0 |
| PLXDC1 | 1.36E-229 | -0.51575 | 0.32 | 0.624 | 5.17E-225 |
| APOL1 | 0 | -0.51772 | 0.118 | 0.514 | 0 |
| NME4 | 0 | -0.51807 | 0.209 | 0.635 | 0 |
| CRISPLD1 | 0 | -0.5181 | 0.121 | 0.541 | 0 |
| CLEC11A | 8.81E-296 | -0.51839 | 0.079 | 0.379 | 3.35E-291 |
| RUNX1T1 | 0 | -0.51848 | 0.247 | 0.636 | 0 |
| SELENBP1 | 0 | -0.51872 | 0.246 | 0.691 | 0 |
| ANGPTL1 | 1.72E-236 | -0.52058 | 0.433 | 0.765 | 6.54E-232 |
| HSP90AA1 | 2.27E-268 | -0.52352 | 0.929 | 0.991 | 8.64E-264 |
| FAM114A1 | 0 | -0.52465 | 0.502 | 0.833 | 0 |
| MAP4K4 | 3.26E-243 | -0.52588 | 0.304 | 0.621 | 1.24E-238 |
| MYL6 | 0 | -0.52876 | 0.954 | 0.997 | 0 |
| PDIA4 | 0 | -0.52904 | 0.389 | 0.77 | 0 |

| | | | | | |
|---|---|---|---|---|---|
| GPX8 | 0 | -0.52911 | 0.382 | 0.789 | 0 |
| KDELR2 | 0 | -0.53073 | 0.702 | 0.923 | 0 |
| HMGN3 | 0 | -0.53103 | 0.445 | 0.829 | 0 |
| ISG15 | 0 | -0.53155 | 0.142 | 0.506 | 0 |
| COL8A1 | 1.90E-196 | -0.53186 | 0.156 | 0.413 | 7.21E-192 |
| LINC0111€ | 0 | -0.53268 | 0.173 | 0.58 | 0 |
| IGHA1 | 0 | -0.53342 | 0 | 0.322 | 0 |
| MYL6B | 0 | -0.53653 | 0.248 | 0.686 | 0 |
| CAV1 | 1.06E-108 | -0.53706 | 0.732 | 0.899 | 4.01E-104 |
| STAT1 | 2.95E-264 | -0.53723 | 0.2 | 0.527 | 1.12E-259 |
| PSMB9 | 0 | -0.53767 | 0.238 | 0.605 | 0 |
| IFI27L2 | 0 | -0.53855 | 0.563 | 0.876 | 0 |
| CNPY4 | 0 | -0.53993 | 0.166 | 0.621 | 0 |
| DLX4 | 0 | -0.54149 | 0.075 | 0.507 | 0 |
| CCND1 | 0 | -0.54184 | 0.114 | 0.466 | 0 |
| FAM198B | 0 | -0.54389 | 0.054 | 0.426 | 0 |
| CCND2 | 0 | -0.54636 | 0.24 | 0.613 | 0 |
| GGT5 | 6.59E-80 | -0.54789 | 0.288 | 0.452 | 2.50E-75 |
| BTN3A2 | 0 | -0.54991 | 0.078 | 0.496 | 0 |
| PDLIM3 | 1.36E-301 | -0.55134 | 0.191 | 0.543 | 5.16E-297 |
| SELENOW | 0 | -0.55343 | 0.579 | 0.904 | 0 |
| PROS1 | 0 | -0.55349 | 0.408 | 0.765 | 0 |
| ANGPT1 | 4.36E-237 | -0.55637 | 0.109 | 0.385 | 1.66E-232 |
| TUBB2B | 1.35E-294 | -0.5598 | 0.063 | 0.35 | 5.15E-290 |
| ZIC1 | 0 | -0.56224 | 0.178 | 0.549 | 0 |
| HLA-DMA | 0 | -0.56292 | 0.062 | 0.391 | 0 |
| RGS2 | 4.12E-179 | -0.56294 | 0.097 | 0.321 | 1.56E-174 |
| ERAP2 | 0 | -0.56448 | 0.066 | 0.483 | 0 |
| ARPC1B | 0 | -0.56485 | 0.484 | 0.806 | 0 |
| C1S | 0 | -0.56637 | 0.975 | 0.996 | 0 |
| SEMA3C | 2.23E-165 | -0.56668 | 0.422 | 0.675 | 8.45E-161 |
| PLOD2 | 0 | -0.56816 | 0.294 | 0.686 | 0 |
| NDUFB2 | 0 | -0.56833 | 0.676 | 0.936 | 0 |
| CLTC | 0 | -0.56857 | 0.339 | 0.745 | 0 |
| ITGAV | 0 | -0.56877 | 0.299 | 0.704 | 0 |
| SGK1 | 4.03E-253 | -0.56928 | 0.142 | 0.442 | 1.53E-248 |
| CHCHD10 | 0 | -0.56955 | 0.234 | 0.629 | 0 |
| P4HB | 0 | -0.5725 | 0.785 | 0.965 | 0 |
| MYO1B | 0 | -0.57504 | 0.156 | 0.604 | 0 |
| GLIPR1 | 0 | -0.57541 | 0.259 | 0.666 | 0 |
| AQP1 | 6.53E-215 | -0.57576 | 0.277 | 0.588 | 2.48E-210 |
| EIF5 | 0 | -0.57715 | 0.648 | 0.917 | 0 |
| SPARCL1 | 3.24E-222 | -0.57978 | 0.471 | 0.79 | 1.23E-217 |
| CTSO | 0 | -0.57988 | 0.229 | 0.666 | 0 |
| SNED1 | 0 | -0.58058 | 0.451 | 0.798 | 0 |
| ZNF385D | 0 | -0.58103 | 0.1 | 0.538 | 0 |
| CNPY2 | 0 | -0.58128 | 0.462 | 0.847 | 0 |

| | | | | | |
|---|---|---|---|---|---|
| COPZ2 | 0 | -0.58263 | 0.56 | 0.885 | 0 |
| EMP3 | 4.49E-267 | -0.5835 | 0.783 | 0.966 | 1.71E-262 |
| C1orf122 | 0 | -0.58461 | 0.477 | 0.847 | 0 |
| GJA1 | 0 | -0.58588 | 0.305 | 0.681 | 0 |
| RPS10 | 0 | -0.58747 | 0.912 | 0.985 | 0 |
| PAM | 0 | -0.58939 | 0.518 | 0.87 | 0 |
| MIR99AHG | 0 | -0.58981 | 0.264 | 0.718 | 0 |
| SSC5D | 0 | -0.59166 | 0.32 | 0.801 | 0 |
| TPM2 | 1.03E-221 | -0.59318 | 0.315 | 0.609 | 3.92E-217 |
| PDIA3 | 0 | -0.59516 | 0.865 | 0.984 | 0 |
| PLTP | 2.27E-215 | -0.59573 | 0.77 | 0.94 | 8.62E-211 |
| NDUFA13 | 0 | -0.59625 | 0.579 | 0.891 | 0 |
| FGF10 | 0 | -0.59872 | 0.149 | 0.592 | 0 |
| PLAU | 0 | -0.60143 | 0.179 | 0.631 | 0 |
| S100A10 | 2.47E-213 | -0.60295 | 0.981 | 0.996 | 9.37E-209 |
| THBS3 | 0 | -0.60356 | 0.227 | 0.649 | 0 |
| SEC31A | 0 | -0.60455 | 0.493 | 0.862 | 0 |
| MSN | 0 | -0.60485 | 0.429 | 0.816 | 0 |
| GEM | 1.47E-67 | -0.60506 | 0.382 | 0.518 | 5.58E-63 |
| LRP1 | 0 | -0.60523 | 0.923 | 0.993 | 0 |
| HDLBP | 0 | -0.60675 | 0.554 | 0.871 | 0 |
| LINC01423 | 5.69E-292 | -0.60723 | 0.005 | 0.249 | 2.16E-287 |
| CANX | 0 | -0.60782 | 0.658 | 0.929 | 0 |
| GTF2I | 0 | -0.6083 | 0.4 | 0.81 | 0 |
| HSPA8 | 0 | -0.60962 | 0.649 | 0.936 | 0 |
| IGFBP7 | 2.56E-154 | -0.61039 | 0.64 | 0.849 | 9.72E-150 |
| KDELR3 | 0 | -0.61067 | 0.141 | 0.581 | 0 |
| EEA1 | 0 | -0.6119 | 0.604 | 0.894 | 0 |
| LXN | 7.35E-258 | -0.61381 | 0.212 | 0.523 | 2.79E-253 |
| C1orf54 | 0 | -0.61825 | 0.097 | 0.593 | 0 |
| TSPAN15 | 0 | -0.61842 | 0.039 | 0.465 | 0 |
| GPC6 | 0 | -0.62009 | 0.116 | 0.569 | 0 |
| LBH | 6.89E-277 | -0.62286 | 0.149 | 0.463 | 2.62E-272 |
| XIST | 1.53E-134 | -0.62383 | 0.643 | 0.68 | 5.82E-130 |
| NRP1 | 0 | -0.62531 | 0.127 | 0.63 | 0 |
| ENG | 0 | -0.62956 | 0.22 | 0.694 | 0 |
| AKR1C3 | 0 | -0.63107 | 0.261 | 0.629 | 0 |
| LSP1 | 0 | -0.63278 | 0.375 | 0.799 | 0 |
| GOLIM4 | 0 | -0.6341 | 0.443 | 0.855 | 0 |
| TUBA1B | 8.38E-298 | -0.63528 | 0.682 | 0.932 | 3.18E-293 |
| ELN | 0 | -0.63547 | 0.184 | 0.538 | 0 |
| MAGED1 | 0 | -0.63581 | 0.118 | 0.609 | 0 |
| KCTD12 | 0 | -0.63616 | 0.148 | 0.596 | 0 |
| TCF4 | 0 | -0.63678 | 0.538 | 0.895 | 0 |
| NORAD | 0 | -0.6391 | 0.533 | 0.869 | 0 |
| TRPS1 | 0 | -0.64052 | 0.347 | 0.769 | 0 |
| PMEPA1 | 0 | -0.64061 | 0.351 | 0.758 | 0 |

| | | | | | |
|---|---|---|---|---|---|
| SOX5 | 0 | -0.64529 | 0.219 | 0.614 | 0 |
| HIF1A | 0 | -0.64621 | 0.308 | 0.698 | 0 |
| ANK2 | 5.91E-187 | -0.64807 | 0.302 | 0.571 | 2.24E-182 |
| NR4A2 | 3.17E-103 | -0.64914 | 0.134 | 0.295 | 1.20E-98 |
| PRDX2 | 0 | -0.65055 | 0.629 | 0.931 | 0 |
| CPQ | 0 | -0.65306 | 0.56 | 0.905 | 0 |
| C7 | 4.95E-141 | -0.65482 | 0.075 | 0.261 | 1.88E-136 |
| EFEMP2 | 0 | -0.65718 | 0.505 | 0.871 | 0 |
| RCN3 | 0 | -0.6602 | 0.49 | 0.857 | 0 |
| TWISTNB | 3.41E-69 | -0.66202 | 0.348 | 0.521 | 1.30E-64 |
| FRMD6 | 0 | -0.66257 | 0.125 | 0.566 | 0 |
| HLA-DRB5 | 4.77E-278 | -0.66364 | 0.017 | 0.262 | 1.81E-273 |
| COL18A1 | 0 | -0.66485 | 0.257 | 0.705 | 0 |
| CALU | 0 | -0.6728 | 0.585 | 0.906 | 0 |
| PSME2 | 0 | -0.6743 | 0.473 | 0.85 | 0 |
| RAB13 | 0 | -0.67503 | 0.477 | 0.816 | 0 |
| DST | 0 | -0.68271 | 0.705 | 0.962 | 0 |
| CHID1 | 0 | -0.68651 | 0.301 | 0.775 | 0 |
| CLEC3B | 5.33E-269 | -0.68686 | 0.288 | 0.603 | 2.03E-264 |
| APP | 0 | -0.68811 | 0.704 | 0.952 | 0 |
| TUBB2A | 0 | -0.6883 | 0.28 | 0.634 | 0 |
| NPTX2 | 0 | -0.69316 | 0.007 | 0.3 | 0 |
| FKBP2 | 0 | -0.69961 | 0.63 | 0.946 | 0 |
| HMCN1 | 0 | -0.70089 | 0.059 | 0.457 | 0 |
| MINOS1 | 0 | -0.70111 | 0.639 | 0.922 | 0 |
| DYNLL1 | 0 | -0.70441 | 0.832 | 0.983 | 0 |
| PDGFRA | 0 | -0.70507 | 0.665 | 0.931 | 0 |
| MGAT4C | 0 | -0.70634 | 0.013 | 0.398 | 0 |
| ANGPTL2 | 0 | -0.70749 | 0.573 | 0.881 | 0 |
| NUCKS1 | 0 | -0.71122 | 0.884 | 0.99 | 0 |
| LTBP1 | 0 | -0.7124 | 0.374 | 0.793 | 0 |
| P4HA2 | 0 | -0.71659 | 0.272 | 0.747 | 0 |
| AHR | 0 | -0.71901 | 0.262 | 0.652 | 0 |
| FAP | 0 | -0.71966 | 0.25 | 0.662 | 0 |
| IGFBP5 | 2.35E-102 | -0.72016 | 0.493 | 0.686 | 8.92E-98 |
| MAGED2 | 0 | -0.72237 | 0.442 | 0.848 | 0 |
| LRRC17 | 0 | -0.7239 | 0.059 | 0.453 | 0 |
| PPFIBP1 | 0 | -0.73116 | 0.338 | 0.764 | 0 |
| CD81 | 0 | -0.73453 | 0.979 | 0.999 | 0 |
| PTMS | 0 | -0.73898 | 0.764 | 0.965 | 0 |
| THBS4 | 0 | -0.74304 | 0.425 | 0.795 | 0 |
| LOXL1 | 0 | -0.74735 | 0.25 | 0.652 | 0 |
| PHPT1 | 0 | -0.75714 | 0.543 | 0.912 | 0 |
| CTSZ | 0 | -0.76518 | 0.494 | 0.883 | 0 |
| NBL1 | 0 | -0.77261 | 0.65 | 0.896 | 0 |
| DAB2 | 0 | -0.78177 | 0.695 | 0.93 | 0 |
| PRDX4 | 0 | -0.78246 | 0.611 | 0.926 | 0 |

| | | | | | |
|---|---|---|---|---|---|
| TGFBI | 9.38E-240 | -0.7858 | 0.188 | 0.483 | 3.56E-235 |
| EMILIN1 | 0 | -0.78682 | 0.411 | 0.822 | 0 |
| TPD52L1 | 0 | -0.78832 | 0.104 | 0.523 | 0 |
| PPIB | 0 | -0.7918 | 0.929 | 0.992 | 0 |
| CHI3L1 | 3.79E-284 | -0.79526 | 0.2 | 0.549 | 1.44E-279 |
| MLEC | 0 | -0.80389 | 0.269 | 0.802 | 0 |
| PDIA6 | 0 | -0.8107 | 0.696 | 0.964 | 0 |
| SMOC2 | 0 | -0.8126 | 0.223 | 0.632 | 0 |
| OSTC | 0 | -0.81872 | 0.655 | 0.946 | 0 |
| DPYSL3 | 0 | -0.8248 | 0.265 | 0.729 | 0 |
| LTBP3 | 0 | -0.8251 | 0.639 | 0.961 | 0 |
| PLPP1 | 0 | -0.82878 | 0.652 | 0.871 | 0 |
| ITGB1 | 0 | -0.83332 | 0.768 | 0.978 | 0 |
| BEX3 | 0 | -0.83465 | 0.459 | 0.902 | 0 |
| ECM2 | 0 | -0.83675 | 0.412 | 0.835 | 0 |
| APLP2 | 0 | -0.84636 | 0.874 | 0.978 | 0 |
| CHMP1B | 1.74E-63 | -0.85342 | 0.444 | 0.581 | 6.61E-59 |
| GALNT1 | 0 | -0.85417 | 0.341 | 0.774 | 0 |
| AHNAK | 0 | -0.85763 | 0.713 | 0.979 | 0 |
| RRBP1 | 0 | -0.86117 | 0.725 | 0.972 | 0 |
| CSGALNAC | 0 | -0.86601 | 0.075 | 0.612 | 0 |
| CALD1 | 0 | -0.86643 | 0.921 | 0.996 | 0 |
| HMGN1 | 0 | -0.86768 | 0.37 | 0.863 | 0 |
| SERPINH1 | 0 | -0.86932 | 0.303 | 0.79 | 0 |
| SSR4 | 0 | -0.87149 | 0.84 | 0.985 | 0 |
| ZEB2 | 0 | -0.87413 | 0.393 | 0.851 | 0 |
| PPP3CA | 0 | -0.88252 | 0.399 | 0.851 | 0 |
| C1GALT1 | 0 | -0.89207 | 0.444 | 0.822 | 0 |
| B2M | 0 | -0.89234 | 0.999 | 1 | 0 |
| IFI6 | 0 | -0.89432 | 0.306 | 0.799 | 0 |
| TCIM | 7.25E-183 | -0.89975 | 0.06 | 0.268 | 2.75E-178 |
| NDUFA4L2 | 3.99E-287 | -0.90601 | 0.201 | 0.543 | 1.52E-282 |
| SSPN | 0 | -0.9067 | 0.491 | 0.904 | 0 |
| LMNA | 0 | -0.90733 | 0.886 | 0.991 | 0 |
| IGLC2 | 0 | -0.90758 | 0.001 | 0.483 | 0 |
| CALR | 0 | -0.90994 | 0.79 | 0.984 | 0 |
| GLT8D2 | 0 | -0.91285 | 0.212 | 0.796 | 0 |
| CD63 | 0 | -0.93615 | 0.997 | 0.998 | 0 |
| SERF2 | 0 | -0.93638 | 0.971 | 0.999 | 0 |
| MRPS6 | 2.17E-178 | -0.94512 | 0.347 | 0.615 | 8.26E-174 |
| TMEM196 | 0 | -0.94721 | 0.12 | 0.62 | 0 |
| ITGB8 | 0 | -0.95027 | 0.122 | 0.593 | 0 |
| NID2 | 0 | -0.95313 | 0.042 | 0.61 | 0 |
| LY6E | 0 | -0.95612 | 0.552 | 0.928 | 0 |
| EPB41L2 | 0 | -0.95673 | 0.515 | 0.942 | 0 |
| LMO4 | 0 | -0.97058 | 0.415 | 0.838 | 0 |
| NUCB2 | 0 | -0.97693 | 0.434 | 0.89 | 0 |

| | | | | | |
|---|---|---|---|---|---|
| ENPP2 | 0 | -0.98528 | 0.113 | 0.603 | 0 |
| RGS3 | 0 | -0.98553 | 0.083 | 0.566 | 0 |
| MMP1 | 1.48E-109 | -0.98878 | 0.008 | 0.115 | 5.63E-105 |
| SRGN | 0 | -0.99043 | 0.017 | 0.457 | 0 |
| C2orf40 | 0 | -0.99164 | 0.339 | 0.775 | 0 |
| MEG3 | 0 | -0.99781 | 0.623 | 0.909 | 0 |
| ITGBL1 | 1.88E-206 | -0.99868 | 0.514 | 0.752 | 7.15E-202 |
| VKORC1 | 0 | -1.00815 | 0.695 | 0.951 | 0 |
| SMOC1 | 0 | -1.01346 | 0.069 | 0.499 | 0 |
| COL5A1 | 0 | -1.01486 | 0.177 | 0.721 | 0 |
| MRC2 | 0 | -1.01541 | 0.483 | 0.947 | 0 |
| SFRP2 | 6.21E-32 | -1.03586 | 0.366 | 0.24 | 2.36E-27 |
| RARRES3 | 0 | -1.04155 | 0.201 | 0.755 | 0 |
| OLFML2B | 0 | -1.05746 | 0.091 | 0.69 | 0 |
| COL6A1 | 0 | -1.06259 | 0.893 | 0.996 | 0 |
| CDH11 | 0 | -1.07365 | 0.173 | 0.729 | 0 |
| C2 | 0 | -1.07421 | 0.196 | 0.788 | 0 |
| CYBA | 0 | -1.07464 | 0.597 | 0.969 | 0 |
| BST2 | 0 | -1.07663 | 0.091 | 0.578 | 0 |
| ISLR | 0 | -1.09098 | 0.575 | 0.938 | 0 |
| PTN | 0 | -1.09452 | 0.104 | 0.478 | 0 |
| CYP1B1 | 0 | -1.09873 | 0.535 | 0.913 | 0 |
| SLC5A3 | 1.25E-195 | -1.1161 | 0.111 | 0.347 | 4.74E-191 |
| GOLM1 | 0 | -1.12135 | 0.202 | 0.769 | 0 |
| KCNQ1OT1 | 0 | -1.12529 | 0.252 | 0.712 | 0 |
| HLA-DPA1 | 0 | -1.12542 | 0.065 | 0.544 | 0 |
| TPPP3 | 0 | -1.12978 | 0.258 | 0.692 | 0 |
| HLA-B | 0 | -1.13421 | 0.929 | 0.99 | 0 |
| LGALS3BP | 0 | -1.14263 | 0.399 | 0.919 | 0 |
| HSP90B1 | 0 | -1.14933 | 0.798 | 0.993 | 0 |
| VMP1 | 0 | -1.16478 | 0.581 | 0.888 | 0 |
| COLEC12 | 0 | -1.18425 | 0.336 | 0.898 | 0 |
| HLA-DPB1 | 0 | -1.18974 | 0.174 | 0.612 | 0 |
| STEAP4 | 0 | -1.2138 | 0.291 | 0.74 | 0 |
| ID3 | 0 | -1.21861 | 0.451 | 0.8 | 0 |
| MARCKS | 0 | -1.23237 | 0.548 | 0.861 | 0 |
| OGN | 0 | -1.25677 | 0.231 | 0.681 | 0 |
| COL6A3 | 0 | -1.26343 | 0.876 | 0.974 | 0 |
| PHLDA1 | 0 | -1.28198 | 0.217 | 0.537 | 0 |
| MDK | 0 | -1.28213 | 0.09 | 0.609 | 0 |
| TTC3 | 0 | -1.2846 | 0.382 | 0.928 | 0 |
| MMP3 | 0 | -1.28831 | 0.071 | 0.463 | 0 |
| LAMA4 | 0 | -1.30853 | 0.278 | 0.904 | 0 |
| VCAM1 | 0 | -1.33429 | 0.198 | 0.854 | 0 |
| RARRES2 | 0 | -1.37376 | 0.133 | 0.495 | 0 |
| TMEM176 | 0 | -1.37825 | 0.217 | 0.723 | 0 |
| EFEMP1 | 0 | -1.38124 | 0.803 | 0.962 | 0 |

| | | | | | |
|---|---|---|---|---|---|
| CTGF | 0 | -1.39754 | 0.625 | 0.886 | 0 |
| SFRP1 | 1.23E-287 | -1.40237 | 0.294 | 0.604 | 4.66E-283 |
| TMEM176 | 0 | -1.41855 | 0.146 | 0.682 | 0 |
| FILIP1L | 0 | -1.43589 | 0.306 | 0.753 | 0 |
| APOE | 0 | -1.46335 | 0.33 | 0.839 | 0 |
| ANKH | 0 | -1.47638 | 0.232 | 0.823 | 0 |
| LUM | 0 | -1.49592 | 0.892 | 0.998 | 0 |
| PPIC | 0 | -1.50577 | 0.427 | 0.954 | 0 |
| IFI27 | 0 | -1.51323 | 0.421 | 0.838 | 0 |
| CHI3L2 | 0 | -1.53467 | 0.303 | 0.765 | 0 |
| SCG2 | 0 | -1.56809 | 0.049 | 0.432 | 0 |
| IGFBP2 | 1.08E-177 | -1.60389 | 0.033 | 0.218 | 4.09E-173 |
| ENAH | 0 | -1.61147 | 0.242 | 0.858 | 0 |
| VCAN | 0 | -1.64501 | 0.859 | 0.934 | 0 |
| CRTAC1 | 0 | -1.64676 | 0.455 | 0.839 | 0 |
| CFI | 0 | -1.66859 | 0.1 | 0.875 | 0 |
| MXRA5 | 0 | -1.66902 | 0.053 | 0.804 | 0 |
| IGF1 | 2.52E-290 | -1.73724 | 0.341 | 0.635 | 9.57E-286 |
| PRSS23 | 0 | -1.74742 | 0.331 | 0.88 | 0 |
| HLA-DRB1 | 0 | -1.74896 | 0.072 | 0.618 | 0 |
| SCRG1 | 0 | -1.75044 | 0.034 | 0.582 | 0 |
| COL5A2 | 0 | -1.82769 | 0.333 | 0.921 | 0 |
| CD74 | 0 | -1.85515 | 0.118 | 0.727 | 0 |
| TIMP1 | 0 | -1.89829 | 0.968 | 0.998 | 0 |
| HLA-DRA | 0 | -1.94166 | 0.046 | 0.667 | 0 |
| CXCL12 | 0 | -1.97724 | 0.588 | 0.935 | 0 |
| THY1 | 0 | -2.01303 | 0.137 | 0.83 | 0 |
| HTRA1 | 0 | -2.08557 | 0.716 | 0.982 | 0 |
| CCDC80 | 0 | -2.10185 | 0.624 | 0.994 | 0 |
| BGN | 0 | -2.14356 | 0.336 | 0.95 | 0 |
| DPT | 0 | -2.14671 | 0.492 | 0.919 | 0 |
| PCOLCE | 0 | -2.1759 | 0.683 | 0.995 | 0 |
| SPARC | 0 | -2.19079 | 0.573 | 0.983 | 0 |
| COL14A1 | 0 | -2.21281 | 0.566 | 0.952 | 0 |
| IGKC | 0 | -2.23025 | 0.001 | 0.662 | 0 |
| CLU | 0 | -2.23577 | 0.909 | 0.967 | 0 |
| IGFBP4 | 0 | -2.27285 | 0.729 | 0.986 | 0 |
| TNC | 0 | -2.58459 | 0.088 | 0.824 | 0 |
| FN1 | 0 | -2.81523 | 0.894 | 0.999 | 0 |
| COL1A2 | 0 | -2.92938 | 0.716 | 1 | 0 |
| COL1A1 | 0 | -3.01181 | 0.527 | 0.981 | 0 |
| COL3A1 | 0 | -3.64806 | 0.562 | 0.998 | 0 |
| PTGDS | 0 | -3.71089 | 0.082 | 0.808 | 0 |
