## Supplementary material for "Adipocytes regulate fibroblast function, and their loss contributes to fibroblast dysfunction in inflammatory diseases": Data file S3

**Data file S3 Bulk RNAseq pathway activation modules**

| FCM activation genes | Cortisol activation genes | FCM+GCR ant activation genes |
|---|---|---|
| CSF3 | ZBTB16 | CSF3 |
| SAA1 | CIDEC | EDNRB |
| RHCG | MMP7 | AREG |
| EDNRB | SAA1 | RHCG |
| STEAP4 | RGCC | CXCL6 |
| ZBTB16 | ANGPTL7 | CXCL8 |
| CXCL5 | LEP | CXCL3 |
| SAA2 | ALOX15B | CXCL5 |
| AREG | FKBP5 | CXCL1 |
| PDK4 | SAA2 | CCL20 |
| CCL20 | FLVCR2 | PDK4 |
| FLVCR2 | RAB4B-EGLN2 | CA2 |
| MT3 | MCTP1 | STC1 |
| OLAH | MAOA | NR4A2 |
| GALNT15 | GLDN | LYVE1 |
| CXCL3 | EDNRB | FLVCR2 |
| TLR2 | PLCE1-AS1 | TFPI2 |
| CXCL1 | NKD2 | CXCL2 |
| ALOX15B | TMEM145 | MUC13 |
| CIDEC | SORBS1 | IL33 |
| CXCL6 | CPM | CXCR4 |
| MCTP1 | STEAP4 | IBSP |
| NKD2 | MT1X | RASD1 |
| SLC7A2 | CRISPLD2 | PITPNC1 |
| MAOA | CACNB2 | GALNT15 |
| LYVE1 | GLUL | NKD2 |
| CXCL8 | GALNT15 | DAW1 |
| AKR1C1 | TSC22D3 | SCARA5 |
| CA2 | COL11A1 | AKR1C1 |
| DAW1 | OMD | PTGES |
| MT1G | FOXO1 | VMO1 |
| AKR1C2 | GPM6B | CCR7 |
| MT1JP | TLR2 | PILRA |
| MT1X | ADH1B | RAB4B-EGLN2 |
| CXCL2 | PKD1P3 | PECAM1 |
| MT1E | RASL11A | C15orf48 |
| HSD11B1 | CORIN | IL1RN |
| FKBP5 | AOC2 | SMOX |
| STC1 | PTK2B | GPRC5A |
| HAS1 | METTL7A | BCL2A1 |
| RASD1 | FAM107A | NR4A1 |
| LEP | ANKRD1 | PTHLH |
| MMP7 | DNAJC6 | HAS1 |
| MUC13 | NAV2-AS5 | FGFBP2 |
| IL1RL1 | TRABD2B | RAB27B |

| | | |
|---|---|---|
| CPM | IPO5P1 | FAM87B |
| MT1L | DELEC1 | OAS1 |
| RGCC | IMPA2 | STEAP4 |
| CCR7 | ALOX5AP | AKR1C2 |
| PITPNC1 | CRYAB | LIF |
| NR4A1 | MT1JP | PREX1 |
| CRISPLD2 | MT1M | MT1F |
| MT1M | GRIA1 | ETV4 |
| MT2A | ACKR2 | PID1 |
| ANGPTL4 | NKD1 | NTRK1 |
| RIPOR2 | TRIM29 | HSD11B1 |
| MAOB | LINC01554 | TREM1 |
| MT1A | MAOB | TMEM145 |
| PILRA | PDGFD | GK |
| APOD | SORT1 | TMEM233 |
| PID1 | TIMP4 | MT1JP |
| NR4A2 | HIF3A | PTGS2 |
| SMOX | CYP19A1 | GPAT3 |
| PTGES | AOX1 | PAQR5 |
| MMD | PMS2P6 | DUSP4 |
| TBX2 | ITGA10 | MT1G |
| GPM6B | ACTC1 | TMEM158 |
| SERPINA3 | SLAMF8 | MT1X |
| SORBS1 | POM121L9P | PTPN22 |
| FGFBP2 | PRELP | CHMP1B |
| PTPN22 | MT1E | MT2A |
| TFPI2 | LMOD1 | EREG |
| GLDN | MT1L | MMP10 |
| TRABD2B | RASL11B | MT1E |
| DELEC1 | FPR1 | IL1B |
| FPR1 | IRAG1 | MT3 |
| PF4 | ADAMTS9 | SLC6A15 |
| PF4V1 | MT1G | GNG11 |
| VMO1 | MT1F | MMP3 |
| RAB4B-EGLN2 | INMT | IGFBP1 |
| CD38 | ADARB1 | PF4V1 |
| INSC | KIF6 | PDE4B |
| RGS2 | PDK4 | MT1L |
| NRCAM | HSPB3 | SERPINB2 |
| PREX1 | GLRX | TLR2 |
| HPDL | MAP2 | OLAH |
| B3GNT5 | SAMHD1 | GPR183 |
| LNCOG | MT2A | DNASE1L3 |
| AKAP12 | DUSP4 | LUCAT1 |
| SIRPB1 | SPINT2 | RASD2 |
| SLC44A1 | NCAM1 | FAM167A |
| RAB38 | FIBIN | MFSD2A |

| | | |
|---|---|---|
| STEAP1 | OLAH | ANGPTL4 |
| SLC6A15 | PILRA | CUL4B |
| IFI44L | SLC38A4 | EGR3 |
| ZC3H12A | SCAMP5 | IL11 |
| SOD2 | TEX2 | KYNU |
| FAM167A | RNF144B | SLCO4A1 |
| GLUL | LYVE1 | MMP1 |
| OAS1 | TG | SAT1 |
| AMPH | DUSP23 | WNT5A |
| MT1F | IRS2 | TBX2 |
| SAT1 | HSPB1 | ARHGAP6 |
| DNER | P2RY11 | CHST2 |
| CCL7 | CCR7 | RGS17 |
| PKD1P3 | NEXN | FAM124A |
| MFSD2A | NEBL | NAMPTP1 |
| NR4A3 | PCDH9 | MTHFD2L |
| PDGFRL | PDGFRL | ENTPD3 |
| CYP19A1 | AOC3 | IL6 |
| MAP2 | GGT5 | SERPIND1 |
| TRIM29 | NPIPB13 | DNER |
| CCNE2 | NRCAM | PF4 |
| TMEM145 | RIPOR2 | KLHL13 |
| CUL4B | MYOSLID | SLC19A3 |
| ETV4 | APOD | MT1A |
| GLRX | BANK1 | MMD |
| LINC02432 | SLC26A6 | PTGDR |
| RAB27B | C11orf52 | CA9 |
| SLAMF8 | ST6GAL1 | DOCK4 |
| CHST2 | ITGA9 | FCER1G |
| TIMP4 | CALCRL | ABCA6 |
| CHMP1B | CNKSR3 | PTGER4 |
| GAS1 | GLCCI1 | RIPOR2 |
| FCER1G | MAMDC2 | NAMPT |
| ADAMTS4 | SERPINA3 | ISG20 |
| NAMPTP1 | DEPP1 | RWDD4P2 |
| SLC39A8 | FAT4 | MCTP1 |
| NFKBIZ | PCA3 | SAA2 |
| TREM1 | MOSMO | DDX10 |
| DUSP1 | ALCAM | RAB38 |
| CCN4 | CORO6 | GFPT2 |
| SLCO4A1 | SNRPEP2 | HHIPL2 |
| TG | TNC | MT1M |
| GNG11 | RAMP1 | NECTIN1 |
| GMNN | HCK | BTBD11 |
| PDE7B | JAG1 | BMP6 |
| DUSP5 | IL16 | C10orf90 |
| NECTIN1 | ANGPTL1 | ACKR3 |

| | | |
|---|---|---|
| GK | PAG1 | PDE7B |
| IBSP | SMARCD2 | EPB41L3 |
| MEDAG | SUSD2 | ITPRIP |
| EGR3 | PARD3B | UCN2 |
| FADS1 | MTSS1 | CDCP1 |
| FAM111B | HSPB2 | MEDAG |
| NAMPT | IGF2 | SIRPB1 |
| AOC2 | AMPH | ZP1 |
| C15orf48 | HOXC11 | ZC3H12A |
| PRELP | KLF13 | TRIM29 |
| SCARA5 | NT5DC3 | SPRY4 |
| PTHLH | PDE4DIPP2 | INSC |
| BCL2A1 | PRKD1 | PDE4D |
| CHI3L2 | ABCC2 | B3GNT5 |
| SEMA4D | MYPN | STEAP1 |
| MYPN | AGFG2 | TSHZ2 |
| ABCA6 | SLC44A1 | CCL7 |
| CFAP69 | FOS | RIPK4 |
| CCL13 | GSTT2 | PLA2G4A |
| SLC19A3 | SERPINA5 | NR4A3 |
| TMEM132B | LIPE | TFPI |
| HPD | HIP1 | BDKRB1 |
| GRIA1 | ITGA5 | CSGALNACT1 |
| LINC01554 | CCN4 | PPP1R14C |
| STEAP1B | OLFML3 | PCSK1 |
| GOLGA6L4 | LRRN3 | KRT81 |
| SLC4A11 | XPNPEP2 | SLC2A13 |
| TFAP4 | CEBPD | BDKRB2 |
| GPR3 | TCEAL4 | HPDL |
| MGAM | ARRDC2 | ESM1 |
| TBC1D8 | PPARG | TNFAIP6 |
| MTHFD2L | ADM | PALMD |
| CHST7 | DDR1 | DUSP6 |
| SLC19A2 | CARMIL1 | IFI44L |
| GPRC5A | IKZF2 | NDP |
| CTSC | CPNE7 | ABCA1 |
| STON1-GTF2A1L | HSD11B1 | SOX5 |
| DUSP4 | FTH1P16 | S1PR1 |
| CDCA7 | LINC00598 | BDH1 |
| TFPI | PRKAG2 | ADAMTS4 |
| LUCAT1 | CLDN7 | MLKL |
| IL33 | FAM167A | GMNN |
| FOXO1 | CERS6 | AKAP12 |
| PHC2 | TXNIP | PGF |
| KRT8P47 | DYNC1I1 | ZNF385D |
| LDHAP3 | ADGRV1 | KIF6 |
| CAMK2N1 | PRUNE2 | SEMA4D |

| | | |
|---|---|---|
| RASSF8-AS1 | SLF1 | PKD1P3 |
| SLC22A4 | MCAM | GLDC |
| NAV2-AS5 | FLVCR1 | CFAP69 |
| IRAK3 | DUSP10 | GAS1 |
| EPB41L3 | PER1 | THBD |
| SPON1 | ABCA6 | PHLDA1 |
| LDHAP4 | NPR3 | PLAT |
| EGR1 | NRP2 | ZNF850 |
| RIBC2 | ZNF485 | AMPH |
| AOX1 | AFAP1L1 | FAM47E |
| RSPO3 | KCNB1 | ADAMTS9 |
| MIOS | LNCOG | TG |
| ITGA10 | STON1 | SFRP1 |
| TBXAS1 | SLC8A1-AS1 | LINC01060 |
| MAN1A1 | NXPH3 | IRAK2 |
| CFAP58 | ABLIM3 | IL21R |
| LDHA | TMEM265 | MLPH |
| IL1RN | MIR155HG | PTGER2 |
| RGS17 | HPD | PORCN |
| LDHAP7 | ANGPT1 | CD38 |
| SLC2A13 | MYCBP2 | PLXNA4 |
| BDH1 | LPAR6 | CRISPLD2 |
| IL18R1 | FADS1 | ANKH |
| MYSM1 | FOXO3 | LINC01119 |
| ELK3 | MB21D2 | HOMER1 |
| METTL9 | VIT | ARRDC2 |
| RRM2 | MYSM1 | CHST7 |
| TWIST2 | SORBS2 | C1GALT1C1 |
| SERPIND1 | NR2F1 | CD55 |
| KLF17 | SLC46A3 | SPON1 |
| PTGDR | GPX3 | SEPSECS-AS1 |
| CDC6 | IRAK3 | DUSP5 |
| CAMK2B | MME | NPTX1 |
| METTL7A | DHCR24 | ST3GAL5 |
| SPRY4 | CCDC88B | ZC2HC1C |
| MND1 | SLPI | SLC5A3 |
| MYOSLID | TWIST1 | ITGB1-DT |
| SQOR | KLHL29 | CREM |
| STEAP2 | GPR89A | LINC02432 |
| PECAM1 | B3GALT4 | SRPX2 |
| PHKA1 | RTN4IP1 | LIMS3 |
| L3MBTL3 | HACD4 | EDNRA |
| SLC26A6 | REEP6 | SMIM3 |
| TWIST1 | MIOS | VASH1-AS1 |
| MAP3K5 | DGCR6 | ANKRD29 |
| MLKL | FGF14 | SQOR |
| RNF144A | FAM43A | BTNL9 |

| | | |
|---|---|---|
| POM121L9P | LINC00968 | IL16 |
| AMPD3 | CMPK1 | DHRS13 |
| NEBL | FMO5 | STEAP1B |
| PDE4B | LPIN3 | CFAP58 |
| TNC | PMP22 | GAPDHP1 |
| CDC45 | SSB | ABHD6 |
| PPARG | SLC66A1L | INHBA |
| WNT5A | ANKRD13B | PTPRN |
| HIPK2 | IFI44L | TARID |
| DIRAS3 | ACVRL1 | TBX3 |
| OGFRL1 | RIMS1 | LAMA1 |
| SEPSECS-AS1 | VCL | KCNK1 |
| CD83 | COPS8 | ITGA2 |
| SINHCAF | KCNE4 | MYSM1 |
| ORC1 | KLF9 | TBC1D8 |
| ZDHHC15 | ANGPTL4 | PMEPA1 |
| RPS26P19 | PDE11A | SIPA1L2 |
| MMP3 | ZBTB6 | PHC2 |
| CHAC2 | TCEAL1 | DHRS9 |
| HS3ST3B1 | SOS2 | FBRSL1 |
| CSGALNACT1 | EOGT | C2CD2 |
| RNF152 | COMP | TLE3 |
| ALOX5AP | RGMA | TFR2 |
| EREG | LAMB1 | RNF152 |
| GFPT2 | TLR4 | PSD4 |
| GPX3 | FGD4 | ELOVL3 |
| SERF1A | TTPAL | CSF1R |
| C9orf72 | LINC00607 | CTSS |
| LINC00607 | DPT | TMEM132B |
| ITPRIP | BDH1 | SLC7A8 |
| BICRA | CAV1 | CPNE4 |
| NNMT | SH3RF3 | RPLP0P2 |
| PTGS1 | SLX4IP | CTSL |
| BMP6 | ZFP36L2 | PFKP |
| LINC01348 | SOX13 | CAMK2B |
| SRPX | ANKEF1 | AGPAT4 |
| CCDC88B | CACHD1 | KCTD14 |
| NCR3LG1 | MPP6 | SLC4A11 |
| PTGER2 | MSANTD2 | HSD17B2 |
| TBX3 | LAMA2 | RNF145 |
| E2F1 | WASF3 | POU2F2 |
| C1QTNF1 | ARHGEF19 | LDHAP3 |
| MCM10 | WTIP | SLC26A6 |
| PAQR5 | TBCE | CCL13 |
| LINC00545 | DANCR | FGL2 |
| NKRF | CHST2 | RPSAP52 |
| ATAD5 | CDH4 | APOD |

| | | |
|---|---|---|
| IL6 | RHOBTB3 | CLGN |
| ZSWIM9 | SPRED1 | CACNA2D3 |
| LDLRAD3 | GAS6-DT | UHRF2 |
| LYSMD2 | CILP2 | SLC44A1 |
| ACKR3 | NABP1 | LRRK2 |
| ENTPD1 | SEPSECS-AS1 | RNF182 |
| CENPE | ATP1B1 | TSC22D1 |
| UGP2 | MTRNR2L10 | DYSF |
| KYNU | LMCD1 | NDUFV2P1 |
| C10orf90 | KCNK6 | PTPRU |
| PDE4D | ELOVL3 | CRPPA |
| ABLIM3 | HOTAIRM1 | RIPOR3 |
| LAMA1 | KATNAL2 | PPP1R3C |
| DSCC1 | NNMT | LINC01588 |
| GABBR2 | PTCH1 | IRS1 |
| DCN | PABPC1P4 | HMOX1 |
| S1PR3 | DUSP5 | ADGRG1 |
| SLC35F2 | MXI1 | TEX26-AS1 |
| ARNTL | APOL2 | SEMA3F |
| E2F8 | EEPD1 | ST6GALNAC4 |
| ITGA5 | HPS5 | PTGFR |
| IMPA2 | LDHA | PAX8 |
| LIMS3 | C12orf60 | BCORL1 |
| TSHZ2 | COL8A1 | PAPPA2 |
| PCSK1 | CALHM5 | SYNDIG1 |
| PLXNA4 | ENAH | TTN |
| CMPK1 | FBLN5 | OTUD3 |
| FTH1P16 | OSER1-DT | ANO7 |
| TMEM164 | THAP11 | CASP9 |
| MPHOSPH6 | CCDC69 | LIPE |
| CLSPN | KLF6 | SLC22A4 |
| KCTD14 | ROCK2 | SHOX |
| DRAXIN | SMIM43 | BDNF |
| HSD17B2 | USP18 | ESYT3 |
| TMEM201 | ZNF175 | SOD2 |
| HIP1 | SLC26A2 | HPD |
| FAM124A | C9orf72 | SLC38A5 |
| FEN1 | ZFP36 | IGF2BP3 |
| EPAS1 | TMOD2 | IER3 |
| RPS2P32 | SELENOP | GRB14 |
| C1GALT1C1 | ZNF232-AS1 | ADGRE2 |
| PAQR4 | TMEM14A | CD82 |
| EDNRA | CASTOR1 | ENOSF1 |
| HELLS | DLX5 | ADGRL4 |
| RPGR | LINC02202 | ABCA8 |
| ADH1B | ADH5 | ST3GAL1 |
| PLK4 | ELF1 | C11orf87 |

| | | |
|---|---|---|
| STK39 | CTNNB1 | PHLDA2 |
| ADM | SLC7A7 | LAMA3 |
| ARRDC2 | SMIM10 | LINC00545 |
| LIF | ADHFE1 | HYAL1 |
| AMD1 | LDHAP4 | FSBP |
| TFR2 | CYB5A | GPAA1P2 |
| MCM3 | HAUS3 | CPM |
| CKAP2L | WHAMMP3 | OAS2 |
| UHRF1 | ZNF215 | BCL2L2-PABPN1 |
| GPAT3 | PHC2 | AMPD3 |
| POU2F2 | LGR5 | MTFP1 |
| ERCC6L | SKP2 | ISM1 |
| SOX13 | HNMT | PYGB |
| CLUH | THBS1 | AHI1 |
| ST3GAL5 | NBPF20 | ARNTL |
| HOMER1 | PTPDC1 | GLIS3 |
| STAMBPL1 | DNAJB4 | SLC35G2 |
| CEP55 | PPM1B | TWIST1 |
| DUSP23 | STK39 | EGFR-AS1 |
| PKN3 | LRRC37A | RASSF8 |
| OTUD3 | PLXNA2 | SLF1 |
| LINC01119 | GADD45B | CP |
| MLPH | MFGE8 | GPR3 |
| TMEM132A | RUNX2 | TMEM132A |
| RASD2 | HSPA2 | SRGN |
| BEND3 | MINCR | RETREG1 |
| TNFAIP3 | LINC01089 | PFKFB4 |
| PIGV | DDIT4 | TGIF1 |
| PKMYT1 | TMEM150A | PPARG |
| DEPDC1B | GON7 | ADGRV1 |
| CTNNB1 | THBS1-AS1 | FPR1 |
| MGST1 | MMP19 | EGR1 |
| ITGB1-DT | REV3L | CCNE2 |
| KCNK1 | PLXNB1 | LPXN |
| ESCO2 | CCDC51 | CDON |
| CYP7B1 | FBXO32 | LRRN3 |
| KAT2A | LINC00565 | EPOR |
| MRM1 | RGS2 | GLRX |
| IL16 | RASD1 | GLA |
| ELOVL3 | PGM2L1 | MYPN |
| EXO1 | ATP10A | MGAT5 |
| CSF1R | FABP5P7 | BZW1 |
| ID1 | IGF2BP2 | LPAR6 |
| WDR76 | GPC4 | SERPINE2 |
| ZNF215 | FNBP1L | DUSP1 |
| RPIA | ZNF443 | ANKEF1 |
| THADA | THBS1-IT1 | KRT8P47 |

| | | |
|---|---|---|
| CMTM7 | PLA2G5 | QPCTL |
| APOL3 | PLCB4 | HBEGF |
| LIPE | ELN | PTP4A1 |
| H2BC11 | KCTD12 | METTL9 |
| NDC1 | PRXL2C | P2RY11 |
| FTH1P8 | DDAH1 | MAML3 |
| LRRK2 | TRAK2 | ADGRL2 |
| C5AR2 | GASK1B | LRP8 |
| SLC19A1 | PDXK | STX1A |
| FAM107A | DHRS12 | GNG2 |
| CRYAB | NRF1 | HNRNPUL2 |
| PSMC3IP | ZNF506 | NINJ2 |
| UPP1 | CCND3 | EMP2 |
| FSBP | ITPKC | PPT2 |
| AGPAT5 | AVIL | SYTL3 |
| GLDC | OFD1 | SYT12 |
| EMP2 | MPC1 | LDLRAD3 |
| FAM87B | FSTL3 | CD83 |
| NDUFV2P1 | ARNTL | ACYP1 |
| MCM2 | ASPN | SLC39A8 |
| SHMT1 | ARSK | VEGFA |
| GGT5 | LDHAP3 | LRRC8C |
| SERPINB2 | TMTC1 | RFX8 |
| ATP1B3 | CPPED1 | LDHA |
| KCTD12 | FZD4 | ANPEP |
| POLA2 | SNAI2 | HERC2P8 |
| MARS2 | CSF1 | TNFRSF1B |
| PRIM1 | GCNT1 | GPR161 |
| RAD54B | RCBTB2 | PRDX6-AS1 |
| GON7 | DUSP1 | SOX13 |
| PLXNA2 | GCAT | MAN1A1 |
| SRGN | DNAJC21 | FUT2 |
| STON1 | HSD17B11 | IRF2BPL |
| E2F2 | LBH | MILR1 |
| TNFAIP8L3 | CNN2 | MPHOSPH6 |
| PRKCH | DCN | LINC00888 |
| ZFP36L2 | JCAD | MCL1 |
| MYCBP | ENDOD1 | TCEAL9 |
| PTK2B | GPR89B | NQO2 |
| TMEM251 | YPEL2 | MCF2L2 |
| L2HGDH | CMIP | RGL3 |
| BRCA1 | UGP2 | ZNF175 |
| ARHGEF19 | PIK3CD | SLC20A1 |
| RASSF8 | BCORL1 | CNIH3 |
| MME | INCA1 | RALGAPA1 |
| PAX8 | AASS | TDP2 |
| CDCA5 | HDX | RIPK2 |

| | | |
|---|---|---|
| ERGIC1 | DAAM2 | STON1 |
| FAM111A | CKB | SPRY3 |
| SPHK1 | CLIC3 | NCALD |
| RPP25 | JAGN1 | SPATA5L1 |
| P2RY11 | SSH2 | PHKA1 |
| MTFP1 | PRDM2 | GSAP |
| NBPF20 | MOCS1 | ADAT2 |
| XPNPEP2 | FAHD2B | ANXA10 |
| NID1 | SLC38A6 | TNFRSF10D |
| DOCK4 | CTSC | RAB3D |
| OAS3 | TNFAIP8L3 | FOXC1 |
| TTLL12 | HIP1R | ID1 |
| PDXP | SAT1 | PITPNM1 |
| PDP2 | SETMAR | IRAK3 |
| PMEPA1 | TJP2 | ZNF697 |
| SERPINA5 | TNS2 | HERC2P5 |
| NFKBIA | FAIM | KIF13B |
| POLN | SLC45A1 | SYTL4 |
| HMGA2 | HIGD1A | FAH |
| MYC | RNF115 | TMTC4 |
| HAGHL | MINDY4 | ITGB3 |
| ANPEP | RBM14-RBM4 | AKR1C3 |
| GK5 | TACC1 | PCDHGB9P |
| USP31 | DSTN | SRRD |
| TMTC1 | ACTA2 | SPRY2 |
| RAB20 | NR2F1-AS1 | SLC1A2 |
| SUV39H2 | ZKSCAN4 | FOLR3 |
| DANCR | AAMDC | PLEK2 |
| BCL2L2-PABPN1 | MBLAC2 | PNP |
| SLC45A1 | PEAK1 | BMERB1 |
| SYNJ1 | SSBP2 | ATP1B3 |
| CHCHD4 | ANKRD36BP1 | MRPS6 |
| QPCTL | ABHD18 | ENO2 |
| PRDM10 | OSBPL5 | PART1 |
| PPP1R14C | CUTC | PLIN2 |
| RBL1 | NDOR1 | MIR155HG |
| DTL | TRAM1L1 | SMN1 |
| BTBD1 | SERPINB7 | SLC16A4 |
| NETO2 | OXCT1 | RAB20 |
| CNKSR3 | ITSN1 | COL7A1 |
| CERS6 | C15orf61 | FGF13 |
| CENPQ | KLHL25 | PLEKHN1 |
| GAPDHP1 | NUDT4 | FAM225B |
| PLEK2 | TNFRSF19 | PTGS1 |
| SLC5A6 | SLX4 | ZNF462 |
| C18orf54 | NEDD9 | BMI1 |
| S1PR1 | ZNF589 | ACSL4 |

| | | |
|---|---|---|
| PCLAF | PCGF5 | NDUFV2 |
| MCL1 | MEGF9 | RNF215 |
| H6PD | DCUN1D3 | CXCL16 |
| HSP90AB2P | BTG2 | KAT2A |
| CCNA2 | RNF141 | PPP3R1 |
| KLHL2 | PGGHG | NR1H3 |
| PCCA-DT | ELL2 | ZMIZ1-AS1 |
| BZW1 | NSUN6 | AKR1B1 |
| FIBIN | SKAP2 | FANCM |
| SENP3-EIF4A1 | CALCOCO2 | TSKU |
| ABALON | ZNF30 | MEG9 |
| EEF1AKMT4 | DKK1 | EEF1AKMT4 |
| EIF2AK3 | ZNF823 | STAMBPL1 |
| ASB13 | ABCC3 | AP1S2 |
| NUDCD1 | DSCC1 | ERGIC1 |
| CHAF1A | TMEM30B | BTN3A2 |
| SNTB1 | DDTL | PGM2L1 |
| RTN4IP1 | C7orf25 | COMMD3 |
| SLC38A5 | UST | DNAJC12 |
| NASP | KNSTRN | ITGA10 |
| SLC26A1 | ZNF768 | HIPK2 |
| KLHL13 | MPC2 | CLCF1 |
| EMP1 | H2AJ | GDPD3 |
| MMS22L | HMGN5 | RPGR |
| ADGRV1 | NUDT16L2P | FNDC4 |
| DEPDC1 | NUDCD1 | LDHAP4 |
| TNFAIP6 | MYC | MEIS1 |
| LPAR6 | GCLC | NID2 |
| PTPDC1 | SCAND2P | TMEM53 |
| CCDC13 | ITGA1 | TBX15 |
| ASF1B | ICAM3 | SH3BP5L |
| LINC01140 | ZBED5-AS1 | ZNF639 |
| FABP5P7 | BAG5 | SERPINA3 |
| UBE2T | DSE | POLG2 |
| CELF2 | BCL2L1 | ELK3 |
| SRPX2 | PDLIM1 | ETV1 |
| SMIM13 | SLC8A1 | SLC35F2 |
| PORCN | LMO7 | PDXP |
| FAH | DPYD | MRM1 |
| KRT81 | ARMC8 | SMIM29 |
| CHAF1B | ZNF28 | FAM225A |
| FANCB | ZHX3 | VAMP4 |
| HOXC11 | PLEKHG2 | LAPTM5 |
| LRRC20 | ANG | GYPC |
| AGFG1 | DEPTOR | NTNG1 |
| URB2 | ID1 | LIMS4 |
| GINS1 | ANXA6 | SLC22A23 |

| | | |
|---|---|---|
| ACSL4 | DRAM2 | GLDN |
| SLC43A3 | EYA2 | MINPP1 |
| RNF115 | SLC7A6 | TRH |
| RAB3D | | RAMP1 |
| NECTIN3 | | MCAM |
| TOP1 | | FAM131A |
| LRP8 | | HDAC8 |
| C1QTNF2 | | ALAS1 |
| PTGFR | | PPFIBP1 |
| NIPA1 | | L3HYPDH |
| U2AF1 | | GON7 |
| KLF9 | | ZNF808 |
| FANCM | | RPL17P50 |
| ARHGEF39 | | RNF113A |
| ARSK | | KLHL29 |
| NPL | | NAF1 |
| ARHGAP6 | | SH2B3 |
| CEP72 | | NINL |
| HAUS3 | | ADK |
| SLC25A15 | | CD58 |
| NCAPD3 | | CMTM7 |
| PMAIP1 | | TMEM251 |
| FZD4 | | INSYN2A |
| ZBTB33 | | HDX |
| GSTO1 | | NAGS |
| UCN2 | | ARL6 |
| LRRC45 | | PIP5KL1 |
| TMEM171 | | TFIP11-DT |
| ARHGAP11B | | IRF3 |
| NIPAL2 | | CDK5RAP2 |
| PPM1B | | SERPINA5 |
| ZFP36L1 | | SLC18B1 |
| NCAPH | | MLLT11 |
| XRCC2 | | SLC9B2 |
| HS3ST3A1 | | SMPDL3A |
| PFKP | | FYN |
| PLA2G2A | | ADSS2 |
| NOLC1 | | SCN1B |
| SRRD | | ICAM2 |
| AFAP1L1 | | ALDOC |
| ZNF485 | | ICAM3 |
| KIF14 | | ZNF215 |
| EBPL | | ZNF284 |
| MCM7 | | ING2 |
| MYZAP | | TOP1 |
| CDK1 | | NOTCH4 |
| RANBP1 | | BLID |

KIF4A
CERS6-AS1
ANKRD1
SERINC2
MCM8
SMIM3
ADAT2
PTPRG
FOXO3
C3
FOS
OAS2
GNG2
SLC25A10
NPAS2
SEC14L2
WDR12
LAMB1
GPATCH4
MYBL2
NXT2
PCA3
MACIR
MAGI3
ELL2
MRPL32
SPR
SORD
ITSN1
MGLL
MCAM
ZNF731P
PGM2L1
DHFR
CEBPD
MAML3
DUSP6
H3C12
TUT7
INCENP
HBEGF
SESTD1
SMIM10
ARL6
ESPL1
NT5DC3
RFC2
PIK3R3
CRLS1
SPTLC3
SH3PXD2B
RPS5P2
ZBTB33
AGFG1
CTNNB1
TRAPPC14
KIF3C
DEPDC7
BOC
LDHAP7
SYNJ1
ESF1
ZSCAN12P1
IL4R
IMPA2
FAM111B
IFI30
ACKR4
DCBLD2
ABL1
DNAJC1
SLX1A-SULT1A3
C1QTNF1
INSYN2B
FAM180A
SEMA3A
RASSF8-AS1
SLC35B4
ERFE
SLC7A7
CHUK
MARCKSL1
LAMB3
C9orf72
PDE3A
DOCK5
PLAU
PLXNA2
TRIM44
ADGRA3
MBLAC2
TNFAIP3
IL17RC
TRAF3IP2

| | |
|---|---|
| RFC4 | GLCE |
| TCF19 | GPR39 |
| BTG3 | SLK |
| HMGN3 | STK39 |
| CPNE4 | DNAJC25 |
| SH3RF3 | PLAUR |
| ICAM3 | GJA1 |
| ZNF443 | KCTD12 |
| GJC1 | APIP |
| KIF18B | C17orf100 |
| FANCI | APOBEC3D |
| TIPIN | RGS12 |
| IGF2 | WLS |
| PYGB | MGST1 |
| ABHD3 | LINC02454 |
| NR1H3 | BACE1-AS |
| RAD54L | LXN |
| KLHL8 | CCDC57 |
| ANKRD9 | PLEKHG5 |
| MIR155HG | GSTO1 |
| IRS2 | PRELP |
| BRI3BP | SMAP2 |
| LMNB2 | TBXAS1 |
| SCML1 | SORD |
| ZDHHC9 | THAP11 |
| CNOT11 | RARG |
| SIK2 | TRO |
| PHLDA1 | BET1 |
| CCND3 | MEG8 |
| JCAD | H6PD |
| CTSS | SESTD1 |
| PGP | NBPF20 |
| TGIF1 | WDR12 |
| CENPX | P4HA3 |
| SNRPD1 | KLHL2 |
| PRDM1 | DNAJC3 |
| PTRH2 | PRRG1 |
| SLC20A1 | HMGN3 |
| NAGS | ARSK |
| RAD51 | LAMA5 |
| OSR2 | NFKBIZ |
| CCDC138 | RORA |
| PRPF19 | ALG6 |
| BUB1B | DTNA |
| TFB1M | TMED5 |
| UTP20 | DLX3 |
| ATAD2 | TNFRSF25 |

ZNF433-AS1
ARHGAP18
STRN
ANKEF1
RIPOR3
IL7R
SYT12
USB1
DDX11
ZNF395
WDHD1
NDUFV2
HSPA14
SPRED1
GCAT
CCDC69
E2F4
PRR5L
MELK
SFXN2
RECQL4
PANX1
RASA2
FGD4
WBP1LP2
SLC7A7
HASPIN
NUDT8
ALG6
KLHL29
GNL2
AADAT
SMC2
ABCC3
FRMD4A
PNP
DNASE1L3
PHLDA2
H3C8
IRF3
DSN1
SS18L2
NOP16
OMD
CDK2
AURKB
MCM5
MRTFB
HOMEZ
CLMP
PCCA
HMGA2-AS1
PPP3CA
ZNF433-AS1
SHISA9
CFB
TMEM164
DLX4
ENTPD7
LPCAT4
PABPC1P4
ZNF771
POGLUT1
MYCBP
CISD2
CYGB
EEF1A1P11
PYCARD
SLC30A1
ZNF589
HSPA8P1
EPAS1
GPR155
DSCC1
MRPL19
SPHK1
SOCS6
IFNAR2
AKAP11
SRPX
DANCR
APOL3
B4GALT3
ABHD2
STRIP2
TIMP4
CYP27C1
MKKS
ESPL1
CTSC
DNAJC24
ZNF248
FADS1
MORC2-AS1

FYN
KIF15
ALG5
GLA
ANKRD34A
GLIS3
POLR3K
ABHD18
DNAAF2
DCP2
CENPJ
SYTL4
SPRY2
MAP2K4
ZNF749
ALMS1
HOMEZ
G0S2
ERRFI1
NKX3-1
ZNF331
FANCA
ALG3
PINX1
GLCE
CDCA4
CENPU
CD82
PDCD5
TRIP13
PISD
LSM6
CCNF
MCM4
NAB1
IRS1
GEMIN4
STIL
SYTL3
SRD5A3
STIM2
POP1
UBE2N
DHX37
HSPE1
THAP11
AUNIP
CPEB4
PCCA-DT
ID2
PLCD4
CCT6P3
GNB5
APEX2
CHST6
TOP1MT
MMAA
NMRK1
PPTC7
BOLA2B
EXTL2
KDM7A-DT
FABP5P7
SEMA4B
NXT2
ANKRD9
G0S2
SLC9A5
ARMCX2
HS3ST3B1
B4GAT1
MAGED4
ADAMTS6
INPP4B
ACOX2
PROCR
ECH1
EEF1E1
ARMCX6
NBPF9
TFB1M
CLDND1
ORAI1
TRMT61B
HGF
MAFK
NRXN2
NAB1
ADPGK
ITGA1
IGSF8
ZBED8
GTF3C1
MRPL32

GSAP
PASK
RRS1
EPOR
LTBP1
FAM20C
NAV2
FTH1
POLD1
CCDC86
MBLAC2
KATNAL2
ZWINT
MB21D2
PSRC1
VRK1
IRAK2
LAMA3
SLC18B1
GCSHP5
TEKT4P2
CIP2A
H4C9
LMNB1
JAGN1
MED11
FURIN
FADS2
SDHAF3
ANKRD35
IFNAR2
NOP58
PFAS
HMBS
FABP5
ENTPD7
LPCAT1
CSTF2
ETS2
LINC01588
PLEKHG2
CARD6
GPR89A
ZBTB24
INAFM1
TOMM40
H1-5
FMNL2
ATG2A
MGLL
EFL1
SPAG4
A4GALT
HAUS3
ZBTB24
SEPTIN6
RCAN1
SMIM8
P4HA1
GOLGA8A
MPG
URB2
AMD1
PRRX2
MTRF1
TAF13
KYAT1
ALDH3A2
HMBS
TBPL1
ARHGEF19
CLK1
NASP
JAGN1
NBN
CDK2AP2
ADAMTS5
GLRX2
DPY19L3
GRK5
PC
ARHGAP12
PACC1

AGPAT4
GRWD1
MMP19
EEF1E1
CACNB3
PBK
NCAPG
FAM225A
KIF20B
SMN1
MRPS12
CFLAR
COL18A1
TMPO
SLC35B4
CMTM4
PPP1R21
TRMT10C
SLC39A14
PLCXD1
MKI67
EDEM3
SLC35D1
MRPS17
SKA3
ACYP1
GGH
SLC35G2
CDKN3
DTNA
PCNA
NUF2
PHC1
PPP3R1
GGCT
GEMIN6
ANGPT1
COA7
TXNRD3
CPSF3
SLC9B2
SH2D3A
THOP1
MTERF3
LIN9
BOLA2B
NR2C2AP

TFRC
GLRX2
PPP1R3C
HIP1R
ZNF589
ATP13A3
DEPDC7
SPC24
SLC38A6
STAT4
MRPS28
SPOUT1
ATIC
RFC3
TEX2
SLC7A8
BRIX1
GINS2
MGAT5
MGME1
MAD2L1
RPUSD1
TICRR
PTP4A1
C22orf46
EXOSC9
BIRC5
RRM1
PER1
HMMR
GYPC
LYAR
PTDSS1
RCC1
GNPNAT1
NFIA
MTF1
ATAD3A
TMTC4
NOP56
PAPOLG
SLC36A4
MTF2
SAMHD1
BCL2L1
IL4R
ST3GAL1

SMIM43
SKP2
PPIF
DCLRE1B
IKZF3
NBPF9
CEBPZ
CGAS
C2CD2
CACHD1
ACOT7
CMAHP
TOP2A
TPM3P9
PARP4
PC
C9orf116
FAM225B
PRRG1
MCM6
MRPL50
SHISA9
PARP1
CHEK1
PPAN
RUVBL1
RHOBTB3
WBP1L
PRRX2
SLX4IP
TDP2
GMPS
FIGNL1
APIP
HYAL3
DHX35
KPNA4
SNRPE
C7orf25
C1orf112
NUDT6
ENOSF1
UST
HROB
MAPK14
CHN1
PRORP

PYCARD
FASTKD5
AKAP1
UHMK1
EEF1AKNMT
SH2B3
CMTM6
SRPK1
COQ2
TIMELESS
RAB31
C19orf48
