## Supplementary material for "Adipocytes regulate fibroblast function, and their loss contributes to fibroblast dysfunction in inflammatory diseases": Data file S4

**Data file S4 AMP symphony clusters**

| AMP Population | Number of healthy cells mapping to this cluster |
|---|---|
| B-0: CD24+CD27+CD11b+ switched memory | 1 |
| B-1: CD24++CD27+IgM+ unswitched memory | 1 |
| B-2: IgM+IgD+TCL1A+ naive | 4 |
| B-3: IgM+IgD+CD1c+ MZ-like | 1 |
| B-4: AICDA+BCL6+ GC-like | 0 |
| B-5: CD11c+LAMP1+ ABC | 0 |
| B-6: IgM+ plasma | 0 |
| B-7: HLA-DR+IgG+ plasmablast | 1 |
| B-8: IgG1+IgG3+ plasma | 1 |
| M-0: MERTK+ SELENOP+ LYVE1+ | 62 |
| M-1: MERTK+ SELENOP+ LYVE1- | 21 |
| M-2: MERTK+ S100A8+ | 34 |
| M-3: MERTK+ HBEGF+ | 2 |
| M-4: SPP1+ | 20 |
| M-5: C1QA+ | 24 |
| M-6: STAT1+ CXCL10+ | 0 |
| M-7: IL1B+ FCN1+ | 32 |
| M-8: PLCG2+ | 4 |
| M-9: DC3 | 3 |
| M-10: DC2 | 11 |
| M-11: DC4 | 8 |
| M-12: DC1 | 1 |
| M-13: pDC | 0 |
| M-14: LAMP3+ | 0 |
| CD4+ IL7R+CCR5+ memory | 16 |
| CD4+ IL7R+ memory | 35 |
| CD4+ CD161+ memory | 37 |
| CD4+ GZMK+ memory | 5 |
| CD4+ naive | 7 |
| CD4+ CD25-low Treg | 2 |
| CD4+ CD146+ memory | 1 |
| CD4+ GNLY+ | 0 |
| CD4+ Tfh/Tph | 0 |
| CD4+ OX40+NR3C1+ | 0 |
| CD4+ Tph | 0 |
| CD8+ GZMB+/TEMRA | 29 |
| CD8+ CD45ROlow/naive | 3 |
| CD8+ GZMK/B+ memory | 8 |
| CD8+ activated/NK-like | 1 |
| Innate-like | 1 |
| Vdelta1 | 2 |
| Vdelta2 | 1 |
| CD38+ | 2 |
| Proliferating | 2 |
| MT-high (low quality) | 9 |

| | |
|---|---|
| E-0: SPARC+ capillary | 490 |
| E-1: LIFR+ venular | 429 |
| E-2: ICAM1+ venular | 376 |
| E-3: NOTCH4+ arteriolar | 475 |
| E-4: Lymphatic | 128 |
| Mu-0: Mural | 275 |
| F-0: PRG4+ CLIC5+ lining | 7535 |
| F-1: PRG4+ lining | 276 |
| F-2: CD34+ sublining | 2567 |
| F-3: POSTN+ sublining | 1757 |
| F-4: DKK3+ sublining | 412 |
| F-5: CD74-hi sublining | 659 |
| F-6: CXCL12+ SFRP1+ sublining | 2898 |
| F-7: NOTCH3+ sublining | 513 |
| F-8: RSPO3+ intermediate | 148 |
| NK-0: CD56dim CD16+ IFNG- | 14 |
| NK-1: CD56dim CD16+ IFNG+CD160+ | 8 |
| NK-2: CD56dim CD16+ IFNG+CD160- | 6 |
| NK-3: CD56dim CD16+ GZMB- | 0 |
| NK-4: CD56bright CD16- GZMA+CD160+ | 0 |
| NK-5: CD56bright CD16- GZMA+CD69+ | 1 |
| NK-6: CD56bright CD16- GNLY+ | 1 |
| NK-7: CD56bright CD16- GNLY+CD69+ | 0 |
| NK-8: CD56bright CD16- IFN response | 0 |
| NK-9: MT-high | 0 |
| NK-10: PCNA+ Proliferating | 2 |
| NK-11: MKI67+ Proliferating | 1 |
| NK-12: IL7R+ ILC | 1 |
| NK-13: IL7R+CD161+ ILC | 0 |
