## Supplementary material for "Adipocytes regulate fibroblast function, and their loss contributes to fibroblast dysfunction in inflammatory diseases": Data file S5

**Data File S5 Adipose cluster differentially expressed genes**

| | p_val | avg_log2F( | pct.1 | pct.2 | p_val_adj |
|---|---|---|---|---|---|
| KAZN | 0 | 1.874265 | 0.721 | 0.29 | 0 |
| RPL11 | 0 | -1.08048 | 0.732 | 0.898 | 0 |
| RPS8 | 0 | -0.74501 | 0.797 | 0.913 | 0 |
| NEGR1 | 0 | 1.580913 | 0.825 | 0.526 | 0 |
| RPL5 | 0 | -1.17293 | 0.524 | 0.769 | 0 |
| DPYD | 0 | 1.319686 | 0.499 | 0.217 | 0 |
| S100A10 | 0 | -1.07914 | 0.757 | 0.888 | 0 |
| S100A11 | 0 | -1.35996 | 0.359 | 0.672 | 0 |
| S100A6 | 0 | -0.76596 | 0.9 | 0.958 | 0 |
| RPS27 | 0 | -1.71735 | 0.468 | 0.812 | 0 |
| H3F3A | 0 | -1.78544 | 0.137 | 0.514 | 0 |
| RPS7 | 0 | -1.63859 | 0.329 | 0.676 | 0 |
| RPS27A | 0 | -1.29461 | 0.618 | 0.839 | 0 |
| COL3A1 | 0 | 0.821229 | 0.882 | 0.678 | 0 |
| PARD3B | 0 | 1.347604 | 0.56 | 0.235 | 0 |
| PTMA | 0 | -1.29269 | 0.479 | 0.765 | 0 |
| COL6A3 | 0 | 0.99055 | 0.85 | 0.66 | 0 |
| RPL32 | 0 | -1.27225 | 0.679 | 0.881 | 0 |
| ZNF385D | 0 | 1.598941 | 0.562 | 0.202 | 0 |
| RPL15 | 0 | -1.62191 | 0.433 | 0.746 | 0 |
| RPSA | 0 | -1.56044 | 0.224 | 0.582 | 0 |
| RPL14 | 0 | -1.2958 | 0.5 | 0.778 | 0 |
| RPL29 | 0 | -2.23124 | 0.264 | 0.68 | 0 |
| PTPRG | 0 | 1.52172 | 0.677 | 0.332 | 0 |
| ROBO2 | 0 | 1.829785 | 0.619 | 0.208 | 0 |
| RPL24 | 0 | -1.20047 | 0.53 | 0.778 | 0 |
| ZBTB20 | 0 | 1.283443 | 0.798 | 0.532 | 0 |
| FNDC3B | 0 | 1.241403 | 0.584 | 0.335 | 0 |
| NAALADL2 | 0 | 1.266998 | 0.505 | 0.232 | 0 |
| RPL35A | 0 | -1.00428 | 0.626 | 0.852 | 0 |
| RPL9 | 0 | -1.86888 | 0.388 | 0.721 | 0 |
| ANK2 | 0 | 1.177749 | 0.73 | 0.504 | 0 |
| RPS3A | 0 | -2.00883 | 0.331 | 0.714 | 0 |
| RPL37 | 0 | -0.97703 | 0.635 | 0.836 | 0 |
| RPS23 | 0 | -1.47528 | 0.565 | 0.832 | 0 |
| RPS14 | 0 | -1.16352 | 0.674 | 0.881 | 0 |
| EBF1 | 0 | 1.361355 | 0.716 | 0.453 | 0 |
| RPS18 | 0 | -1.57335 | 0.627 | 0.87 | 0 |
| RPL10A | 0 | -1.52133 | 0.424 | 0.739 | 0 |
| FKBP5 | 0 | 1.41549 | 0.65 | 0.425 | 0 |
| EEF1A1 | 0 | -2.15445 | 0.493 | 0.804 | 0 |
| LAMA2 | 0 | 1.527639 | 0.861 | 0.54 | 0 |
| RPS12 | 0 | -1.25068 | 0.695 | 0.89 | 0 |
| UST | 0 | 1.264528 | 0.447 | 0.183 | 0 |
| COL1A2 | 0 | 0.762719 | 0.964 | 0.842 | 0 |

| | | | | | |
|---|---|---|---|---|---|
| KCND2 | 0 | 1.987893 | 0.469 | 0.115 | 0 |
| DLC1 | 0 | 1.611703 | 0.682 | 0.358 | 0 |
| RPL7 | 0 | -2.41011 | 0.107 | 0.566 | 0 |
| RPL30 | 0 | -1.36969 | 0.575 | 0.826 | 0 |
| RPL8 | 0 | -1.55395 | 0.462 | 0.758 | 0 |
| SVEP1 | 0 | 1.581437 | 0.727 | 0.436 | 0 |
| RPL12 | 0 | -1.08607 | 0.634 | 0.85 | 0 |
| RPL7A | 0 | -1.79611 | 0.428 | 0.75 | 0 |
| BICC1 | 0 | 1.822627 | 0.663 | 0.368 | 0 |
| RPS24 | 0 | -1.22599 | 0.621 | 0.842 | 0 |
| RPS13 | 0 | -1.32723 | 0.52 | 0.804 | 0 |
| FTH1 | 0 | -1.41235 | 0.679 | 0.828 | 0 |
| EEF1G | 0 | -1.73364 | 0.02 | 0.35 | 0 |
| FAU | 0 | -1.38036 | 0.425 | 0.737 | 0 |
| RPS3 | 0 | -1.51369 | 0.471 | 0.759 | 0 |
| RPS25 | 0 | -1.17685 | 0.512 | 0.792 | 0 |
| IGFBP6 | 0 | -1.65929 | 0.48 | 0.768 | 0 |
| CD63 | 0 | -0.99621 | 0.668 | 0.798 | 0 |
| RPS26 | 0 | -2.02766 | 0.04 | 0.414 | 0 |
| RPL41 | 0 | -1.50708 | 0.587 | 0.844 | 0 |
| NACA | 0 | -1.56824 | 0.351 | 0.686 | 0 |
| NAV3 | 0 | 1.474805 | 0.524 | 0.22 | 0 |
| IGF1 | 0 | 1.597392 | 0.55 | 0.245 | 0 |
| RPL6 | 0 | -1.40958 | 0.431 | 0.726 | 0 |
| RPL21 | 0 | -2.5047 | 0.116 | 0.593 | 0 |
| DCLK1 | 0 | 1.288615 | 0.744 | 0.459 | 0 |
| TPT1 | 0 | -1.26351 | 0.84 | 0.939 | 0 |
| B2M | 0 | -0.91707 | 0.735 | 0.883 | 0 |
| RORA | 0 | 1.496941 | 0.68 | 0.384 | 0 |
| RPLP1 | 0 | -1.2591 | 0.77 | 0.934 | 0 |
| RPS2 | 0 | -1.91335 | 0.392 | 0.687 | 0 |
| RPS15A | 0 | -1.4303 | 0.509 | 0.819 | 0 |
| MT2A | 0 | -1.81835 | 0.59 | 0.8 | 0 |
| RPL13 | 0 | -1.12426 | 0.807 | 0.886 | 0 |
| RPL26 | 0 | -1.59417 | 0.486 | 0.79 | 0 |
| SNHG29 | 0 | -1.45216 | 0.014 | 0.332 | 0 |
| RPL23A | 0 | -1.30685 | 0.505 | 0.768 | 0 |
| RPL19 | 0 | -1.35305 | 0.61 | 0.844 | 0 |
| EIF1 | 0 | -1.50749 | 0.599 | 0.837 | 0 |
| ABCA9 | 0 | 1.385652 | 0.708 | 0.459 | 0 |
| ABCA10 | 0 | 1.47583 | 0.548 | 0.274 | 0 |
| H3F3B | 0 | -1.1431 | 0.574 | 0.767 | 0 |
| GREB1L | 0 | 1.946179 | 0.282 | 0.065 | 0 |
| RPL17 | 0 | -2.50605 | 0.06 | 0.416 | 0 |
| RPS15 | 0 | -1.50395 | 0.482 | 0.802 | 0 |
| RPS28 | 0 | -1.24273 | 0.485 | 0.756 | 0 |
| RPL18A | 0 | -2.34795 | 0.216 | 0.676 | 0 |

| | | | | | |
|---|---|---|---|---|---|
| RPS16 | 0 | -1.23457 | 0.577 | 0.826 | 0 |
| RPS19 | 0 | -1.25276 | 0.538 | 0.786 | 0 |
| RPL18 | 0 | -1.5279 | 0.44 | 0.736 | 0 |
| FTL | 0 | -1.41957 | 0.698 | 0.88 | 0 |
| RPL13A | 0 | -1.08712 | 0.708 | 0.901 | 0 |
| RPS9 | 0 | -1.57019 | 0.463 | 0.752 | 0 |
| RPL28 | 0 | -1.63377 | 0.433 | 0.757 | 0 |
| RPS5 | 0 | -1.51537 | 0.365 | 0.701 | 0 |
| SAMHD1 | 0 | 1.326421 | 0.653 | 0.411 | 0 |
| LGALS1 | 0 | -0.99581 | 0.652 | 0.834 | 0 |
| RPL3 | 0 | -1.72531 | 0.415 | 0.751 | 0 |
| SLC25A6 | 0 | -1.65209 | 0.023 | 0.362 | 0 |
| CD99 | 0 | -1.99282 | 0.029 | 0.42 | 0 |
| TMSB4X | 0 | -0.93697 | 0.866 | 0.958 | 0 |
| RPS4X | 0 | -1.47783 | 0.58 | 0.833 | 0 |
| RPL36A | 0 | -1.40632 | 0.087 | 0.409 | 0 |
| RPL39 | 0 | -1.33156 | 0.479 | 0.765 | 0 |
| RPL10 | 0 | -1.97436 | 0.604 | 0.874 | 0 |
| LINC00486 | 0 | 0.942155 | 0.894 | 0.383 | 0 |
| MKL1 | 0 | 1.246506 | 0.334 | 0.097 | 0 |
| NFIA | 3.32E-307 | 1.037598 | 0.74 | 0.572 | 1.21E-302 |
| NME2 | 6.17E-307 | -1.34793 | 0.042 | 0.348 | 2.26E-302 |
| RPLP0 | 2.64E-306 | -1.11947 | 0.483 | 0.73 | 9.64E-302 |
| RPL22 | 3.18E-305 | -1.19585 | 0.396 | 0.687 | 1.16E-300 |
| RPS6 | 1.63E-304 | -0.73509 | 0.781 | 0.906 | 5.96E-300 |
| SEPTIN7 | 8.32E-302 | -1.08785 | 0.025 | 0.322 | 3.04E-297 |
| RPL37A | 1.59E-301 | -0.86904 | 0.668 | 0.847 | 5.80E-297 |
| CELF2 | 1.65E-301 | 1.089029 | 0.672 | 0.449 | 6.03E-297 |
| GAS5 | 1.76E-301 | -1.20206 | 0.013 | 0.299 | 6.44E-297 |
| MAML2 | 1.56E-299 | 1.436955 | 0.487 | 0.243 | 5.69E-295 |
| S100A4 | 1.03E-297 | -0.88704 | 0.743 | 0.869 | 3.78E-293 |
| MT-ATP8 | 2.30E-297 | 0.954953 | 0.737 | 0.501 | 8.40E-293 |
| UBA52 | 2.85E-297 | -1.09123 | 0.484 | 0.732 | 1.04E-292 |
| EEF1D | 8.24E-292 | -1.36811 | 0.25 | 0.569 | 3.01E-287 |
| ATP5F1E | 8.41E-291 | -1.04839 | 0.013 | 0.292 | 3.08E-286 |
| ACTG1 | 1.28E-290 | -1.16391 | 0.521 | 0.724 | 4.67E-286 |
| GADD45B | 5.72E-290 | -1.85895 | 0.225 | 0.529 | 2.09E-285 |
| VIM | 2.59E-288 | -0.8534 | 0.938 | 0.916 | 9.47E-284 |
| PARD3 | 2.80E-286 | 1.22723 | 0.431 | 0.189 | 1.02E-281 |
| EMP3 | 2.91E-280 | -1.4012 | 0.149 | 0.474 | 1.06E-275 |
| SLIT3 | 8.10E-280 | 0.990378 | 0.697 | 0.513 | 2.96E-275 |
| SNED1 | 5.20E-276 | 1.155609 | 0.543 | 0.292 | 1.90E-271 |
| EEF1B2 | 1.10E-273 | -1.28025 | 0.17 | 0.489 | 4.02E-269 |
| MGP | 2.53E-273 | -1.27708 | 0.891 | 0.92 | 9.26E-269 |
| RPL36 | 1.49E-272 | -1.09893 | 0.467 | 0.731 | 5.43E-268 |
| TACC1 | 2.39E-272 | 1.000817 | 0.735 | 0.552 | 8.72E-268 |
| BTF3 | 1.69E-268 | -1.19204 | 0.284 | 0.584 | 6.17E-264 |

| | | | | | |
|---|---|---|---|---|---|
| UBC | 4.23E-267 | -1.45115 | 0.601 | 0.737 | 1.54E-262 |
| MAN1A1 | 1.75E-266 | 1.279584 | 0.591 | 0.394 | 6.39E-262 |
| ATP5MG | 1.99E-266 | -0.86267 | 0.01 | 0.269 | 7.29E-262 |
| SBF2 | 1.59E-263 | 1.165521 | 0.475 | 0.241 | 5.82E-259 |
| CLU | 3.88E-263 | -1.3147 | 0.302 | 0.604 | 1.42E-258 |
| LAMC1 | 2.34E-262 | 1.029904 | 0.664 | 0.475 | 8.56E-258 |
| REV3L | 6.21E-261 | 1.072785 | 0.633 | 0.418 | 2.27E-256 |
| TCF4 | 9.56E-261 | 0.995853 | 0.671 | 0.5 | 3.49E-256 |
| DLG2 | 5.63E-258 | 1.174381 | 0.369 | 0.145 | 2.06E-253 |
| LDHA | 4.02E-256 | -1.29923 | 0.213 | 0.516 | 1.47E-251 |
| GAPDH | 4.66E-256 | -1.25331 | 0.269 | 0.569 | 1.70E-251 |
| RUNX1T1 | 1.32E-252 | 1.157491 | 0.426 | 0.2 | 4.82E-248 |
| RPL34 | 1.46E-252 | -0.60592 | 0.818 | 0.924 | 5.35E-248 |
| COL5A2 | 1.96E-252 | 1.105179 | 0.602 | 0.393 | 7.16E-248 |
| PTEN | 3.72E-252 | 1.481192 | 0.55 | 0.356 | 1.36E-247 |
| ATP5PF | 1.59E-251 | -0.77209 | 0.013 | 0.263 | 5.80E-247 |
| MKLN1 | 2.93E-251 | 1.094509 | 0.488 | 0.249 | 1.07E-246 |
| GPX4 | 6.21E-251 | -1.18086 | 0.338 | 0.625 | 2.27E-246 |
| RNASEK | 9.50E-250 | -0.79478 | 0.02 | 0.274 | 3.47E-245 |
| MIF | 1.66E-249 | -1.03494 | 0.104 | 0.398 | 6.07E-245 |
| CCN5 | 3.49E-248 | -1.76668 | 0.018 | 0.267 | 1.27E-243 |
| WWOX | 3.59E-248 | 1.114018 | 0.404 | 0.174 | 1.31E-243 |
| MALAT1 | 5.27E-248 | 0.416436 | 1 | 0.997 | 1.92E-243 |
| ACTB | 8.55E-248 | -0.96695 | 0.577 | 0.759 | 3.13E-243 |
| GPC3 | 2.21E-247 | 0.846199 | 0.743 | 0.526 | 8.08E-243 |
| RPL35 | 9.21E-247 | -0.9215 | 0.564 | 0.788 | 3.37E-242 |
| ATP6V0C | 1.08E-246 | -0.78587 | 0.011 | 0.256 | 3.93E-242 |
| CST3 | 1.18E-246 | -1.09943 | 0.662 | 0.755 | 4.32E-242 |
| CEBPB | 2.95E-246 | -1.6156 | 0.149 | 0.424 | 1.08E-241 |
| ATP5MC2 | 2.49E-245 | -0.79789 | 0.01 | 0.253 | 9.09E-241 |
| CCN1 | 8.79E-244 | -1.62208 | 0.026 | 0.278 | 3.21E-239 |
| AUTS2 | 3.37E-243 | 1.153977 | 0.464 | 0.234 | 1.23E-238 |
| EIF4A1 | 1.90E-242 | -1.31404 | 0.19 | 0.466 | 6.94E-238 |
| MT-ND1 | 3.60E-242 | -0.7219 | 0.848 | 0.896 | 1.31E-237 |
| ZFPM2 | 1.24E-241 | 1.059052 | 0.48 | 0.248 | 4.53E-237 |
| ATP5MC3 | 5.41E-241 | -0.75877 | 0.014 | 0.256 | 1.98E-236 |
| SERTAD1 | 3.44E-240 | -1.36971 | 0.048 | 0.31 | 1.26E-235 |
| ARHGAP6 | 1.38E-239 | 1.222491 | 0.333 | 0.132 | 5.05E-235 |
| SNHG5 | 6.73E-239 | -0.82012 | 0.011 | 0.249 | 2.46E-234 |
| FAM46A | 1.05E-237 | 1.032139 | 0.32 | 0.114 | 3.83E-233 |
| OAZ1 | 1.27E-236 | -1.12929 | 0.286 | 0.578 | 4.65E-232 |
| NPM1 | 1.28E-236 | -1.18779 | 0.173 | 0.469 | 4.67E-232 |
| C11orf96 | 9.38E-236 | -2.33337 | 0.121 | 0.384 | 3.43E-231 |
| PLAGL1 | 3.35E-235 | 1.253585 | 0.51 | 0.308 | 1.22E-230 |
| HSPA8 | 5.60E-235 | -1.33608 | 0.129 | 0.413 | 2.05E-230 |
| DMD | 6.25E-234 | 1.084835 | 0.351 | 0.14 | 2.28E-229 |
| DYNLL1 | 7.49E-234 | -1.20175 | 0.182 | 0.478 | 2.74E-229 |

| | | | | | |
|---|---|---|---|---|---|
| TIMP1 | 8.29E-234 | -1.21617 | 0.416 | 0.669 | 3.03E-229 |
| SUMO2 | 1.67E-233 | -1.12323 | 0.176 | 0.475 | 6.10E-229 |
| PHACTR1 | 1.75E-233 | 1.115518 | 0.201 | 0.05 | 6.41E-229 |
| SOCS3 | 3.88E-229 | -1.69571 | 0.24 | 0.486 | 1.42E-224 |
| ADIRF | 6.09E-229 | -1.30229 | 0.216 | 0.501 | 2.23E-224 |
| RPL36AL | 4.26E-227 | -1.12024 | 0.226 | 0.528 | 1.56E-222 |
| LAMB1 | 2.68E-226 | 1.209548 | 0.527 | 0.32 | 9.79E-222 |
| MYC | 1.33E-225 | -1.36046 | 0.176 | 0.449 | 4.87E-221 |
| USP53 | 2.73E-225 | 0.97968 | 0.622 | 0.413 | 9.99E-221 |
| PDGFD | 5.31E-225 | 1.314775 | 0.368 | 0.165 | 1.94E-220 |
| ATP5PO | 1.49E-224 | -0.64609 | 0.005 | 0.227 | 5.43E-220 |
| ATP5F1D | 6.31E-224 | -0.67895 | 0.01 | 0.236 | 2.31E-219 |
| FTX | 7.51E-221 | 1.14059 | 0.518 | 0.315 | 2.74E-216 |
| CCL2 | 1.81E-218 | -2.42924 | 0.061 | 0.304 | 6.62E-214 |
| ABLIM1 | 2.29E-218 | 0.893834 | 0.596 | 0.366 | 8.36E-214 |
| TOMM7 | 5.04E-216 | -1.15335 | 0.266 | 0.561 | 1.84E-211 |
| SEMA3A | 6.54E-216 | 1.298316 | 0.265 | 0.092 | 2.39E-211 |
| ATP5PD | 6.91E-216 | -0.63227 | 0.009 | 0.228 | 2.53E-211 |
| CDK14 | 3.73E-215 | 1.195273 | 0.366 | 0.169 | 1.36E-210 |
| EGFR | 1.58E-214 | 1.074633 | 0.544 | 0.353 | 5.78E-210 |
| CYR61 | 1.88E-214 | 0.882026 | 0.345 | 0.135 | 6.88E-210 |
| CCN2 | 1.42E-213 | -1.45154 | 0.021 | 0.247 | 5.19E-209 |
| TLE5 | 1.97E-213 | -0.66139 | 0.014 | 0.235 | 7.20E-209 |
| KCNT2 | 3.10E-213 | 1.049355 | 0.26 | 0.086 | 1.13E-208 |
| ATP5F1B | 5.26E-213 | -0.63008 | 0.009 | 0.224 | 1.92E-208 |
| SH3BGRL3 | 2.17E-212 | -1.37643 | 0.214 | 0.491 | 7.94E-208 |
| MT-ATP6 | 2.50E-212 | -0.93493 | 0.93 | 0.908 | 9.15E-208 |
| MT1X | 7.65E-212 | -1.93553 | 0.249 | 0.499 | 2.80E-207 |
| LINC0245[illegible] | 1.45E-211 | 0.826562 | 0.182 | 0.043 | 5.31E-207 |
| MT1A | 1.24E-210 | -1.99979 | 0.124 | 0.374 | 4.53E-206 |
| CHCHD2 | 1.69E-210 | -1.04607 | 0.225 | 0.516 | 6.17E-206 |
| REX1BD | 1.72E-210 | -0.614 | 0.012 | 0.228 | 6.28E-206 |
| PFDN5 | 8.31E-210 | -1.09675 | 0.286 | 0.566 | 3.04E-205 |
| TTC28 | 1.47E-209 | 1.075157 | 0.366 | 0.165 | 5.38E-205 |
| LINC0096[illegible] | 4.40E-207 | 0.806284 | 0.246 | 0.078 | 1.61E-202 |
| LPP | 9.16E-207 | 0.90519 | 0.591 | 0.392 | 3.35E-202 |
| FBXL7 | 1.78E-206 | 1.106007 | 0.418 | 0.211 | 6.52E-202 |
| PRKN | 5.51E-206 | 0.95013 | 0.287 | 0.105 | 2.01E-201 |
| SDK1 | 8.82E-206 | 0.941359 | 0.407 | 0.188 | 3.22E-201 |
| NOVA1 | 1.57E-205 | 0.821396 | 0.726 | 0.577 | 5.73E-201 |
| ATP5F1C | 2.17E-205 | -0.58574 | 0.009 | 0.218 | 7.91E-201 |
| ARL17B | 3.31E-205 | 0.947786 | 0.156 | 0.034 | 1.21E-200 |
| TUBB4B | 6.49E-205 | -1.13552 | 0.128 | 0.393 | 2.37E-200 |
| RFX2 | 7.48E-205 | 1.095861 | 0.434 | 0.23 | 2.73E-200 |
| SNHG6 | 2.64E-204 | -0.62856 | 0.009 | 0.218 | 9.67E-200 |
| IFITM3 | 8.71E-204 | -0.91914 | 0.562 | 0.729 | 3.18E-199 |
| PPIA | 5.46E-203 | -1.02326 | 0.279 | 0.569 | 2.00E-198 |

| | | | | | |
|---|---|---|---|---|---|
| H2AFZ | 4.23E-202 | -1.06768 | 0.056 | 0.297 | 1.55E-197 |
| YBX1 | 9.42E-202 | -0.86543 | 0.279 | 0.556 | 3.44E-197 |
| CDKN1A | 1.17E-201 | -1.17162 | 0.107 | 0.363 | 4.28E-197 |
| MT1M | 2.47E-201 | -1.69213 | 0.202 | 0.454 | 9.03E-197 |
| DNAJB1 | 3.78E-201 | -1.33002 | 0.108 | 0.367 | 1.38E-196 |
| JUNB | 3.30E-200 | -1.45772 | 0.457 | 0.624 | 1.20E-195 |
| LINC01578 | 3.65E-200 | -0.63818 | 0.014 | 0.224 | 1.34E-195 |
| PRKG1 | 1.53E-198 | 1.21237 | 0.44 | 0.247 | 5.60E-194 |
| CD9 | 2.76E-196 | -1.14321 | 0.116 | 0.37 | 1.01E-191 |
| SEPTIN2 | 1.27E-195 | -0.60179 | 0.015 | 0.222 | 4.63E-191 |
| RPL27A | 1.86E-195 | -0.81856 | 0.63 | 0.825 | 6.81E-191 |
| ATP5MD | 5.78E-195 | -0.5731 | 0.008 | 0.207 | 2.11E-190 |
| RBMS3 | 1.12E-194 | 1.049481 | 0.515 | 0.329 | 4.10E-190 |
| CEBPD | 2.47E-194 | -1.6384 | 0.335 | 0.538 | 9.04E-190 |
| CLIC1 | 6.54E-194 | -0.99343 | 0.167 | 0.44 | 2.39E-189 |
| PAM | 7.69E-194 | 0.93716 | 0.569 | 0.377 | 2.81E-189 |
| PTBP2 | 2.83E-193 | 1.050975 | 0.312 | 0.132 | 1.03E-188 |
| PTN | 3.36E-193 | -1.63637 | 0.035 | 0.249 | 1.23E-188 |
| LDB2 | 5.76E-193 | 1.268432 | 0.338 | 0.161 | 2.10E-188 |
| SOD2 | 1.62E-191 | -1.61904 | 0.368 | 0.578 | 5.92E-187 |
| RTRAF | 1.75E-191 | -0.54611 | 0.007 | 0.203 | 6.40E-187 |
| PTPRS | 6.93E-191 | 0.789191 | 0.332 | 0.143 | 2.53E-186 |
| SH3D19 | 8.73E-191 | 0.971903 | 0.584 | 0.414 | 3.19E-186 |
| RBFOX1 | 9.67E-190 | 0.765616 | 0.396 | 0.18 | 3.54E-185 |
| CFH | 7.27E-189 | 0.739797 | 0.733 | 0.586 | 2.66E-184 |
| RPS6KA5 | 1.15E-188 | 1.118685 | 0.304 | 0.128 | 4.21E-184 |
| LINC02511 | 1.46E-188 | 1.156367 | 0.385 | 0.189 | 5.33E-184 |
| ARL15 | 2.91E-188 | 1.047826 | 0.246 | 0.086 | 1.06E-183 |
| KRTCAP2 | 1.73E-187 | -0.75155 | 0.076 | 0.318 | 6.34E-183 |
| LINC01091 | 4.62E-187 | 1.091012 | 0.177 | 0.048 | 1.69E-182 |
| TOMM6 | 7.45E-186 | -0.51539 | 0.007 | 0.198 | 2.72E-181 |
| BST2 | 1.26E-185 | -1.16207 | 0.063 | 0.288 | 4.59E-181 |
| ABCA9-AS1 | 1.69E-185 | 0.818507 | 0.214 | 0.066 | 6.19E-181 |
| RAD51B | 3.05E-185 | 0.997356 | 0.321 | 0.136 | 1.11E-180 |
| HSP90AB1 | 1.05E-184 | -0.97205 | 0.404 | 0.647 | 3.83E-180 |
| SAT1 | 2.62E-184 | -1.29917 | 0.274 | 0.522 | 9.58E-180 |
| TMEM35B | 1.48E-183 | -0.53432 | 0.019 | 0.22 | 5.40E-179 |
| C3 | 1.58E-183 | 0.601078 | 0.834 | 0.682 | 5.77E-179 |
| PLXDC2 | 4.74E-183 | 0.922676 | 0.558 | 0.367 | 1.73E-178 |
| AP001528. | 8.57E-183 | 0.822893 | 0.187 | 0.052 | 3.13E-178 |
| CIAO2B | 9.53E-183 | -0.49833 | 0.007 | 0.195 | 3.48E-178 |
| MICOS10 | 3.29E-182 | -0.52476 | 0.006 | 0.194 | 1.20E-177 |
| AKAP9 | 5.96E-182 | 0.872718 | 0.609 | 0.443 | 2.18E-177 |
| PEAK1 | 8.42E-182 | 0.959492 | 0.366 | 0.176 | 3.08E-177 |
| HSPE1 | 1.65E-181 | -0.9578 | 0.077 | 0.313 | 6.02E-177 |
| PRDX1 | 2.99E-181 | -0.94001 | 0.19 | 0.457 | 1.09E-176 |
| EIF1B | 3.17E-181 | -0.86284 | 0.089 | 0.328 | 1.16E-176 |

| | | | | | |
|---|---|---|---|---|---|
| BNC2 | 1.47E-179 | 0.991251 | 0.348 | 0.167 | 5.36E-175 |
| MT-CO1 | 2.89E-179 | -0.67549 | 0.908 | 0.917 | 1.06E-174 |
| ABCA6 | 4.20E-179 | 0.990846 | 0.554 | 0.388 | 1.54E-174 |
| ATP5IF1 | 6.85E-179 | -0.50761 | 0.009 | 0.197 | 2.50E-174 |
| MFAP4 | 9.18E-179 | -0.99253 | 0.482 | 0.67 | 3.35E-174 |
| XIST | 1.04E-178 | 0.739364 | 0.621 | 0.432 | 3.81E-174 |
| S100A13 | 1.73E-178 | -0.97347 | 0.392 | 0.646 | 6.31E-174 |
| ZFAS1 | 8.82E-178 | -0.93708 | 0.198 | 0.465 | 3.22E-173 |
| MME | 1.27E-177 | 1.072329 | 0.26 | 0.101 | 4.64E-173 |
| PRDX2 | 1.29E-177 | -0.83894 | 0.074 | 0.307 | 4.70E-173 |
| MGEA5 | 2.33E-177 | 0.765468 | 0.307 | 0.125 | 8.53E-173 |
| AL031599. | 6.21E-177 | 1.010956 | 0.161 | 0.041 | 2.27E-172 |
| PIM1 | 1.64E-175 | -0.93811 | 0.064 | 0.283 | 6.01E-171 |
| KIAA1217 | 1.69E-175 | 1.139542 | 0.284 | 0.117 | 6.19E-171 |
| DENND2A | 2.09E-175 | 1.020739 | 0.296 | 0.125 | 7.63E-171 |
| CXCL12 | 3.26E-175 | 0.731674 | 0.749 | 0.596 | 1.19E-170 |
| MGST3 | 2.31E-174 | -0.90334 | 0.207 | 0.477 | 8.46E-170 |
| THBS1 | 5.21E-174 | -1.45556 | 0.041 | 0.243 | 1.91E-169 |
| SNHG32 | 5.45E-174 | -0.50006 | 0.007 | 0.188 | 1.99E-169 |
| MYO1D | 5.67E-174 | 1.076804 | 0.293 | 0.126 | 2.07E-169 |
| RPS10 | 6.26E-174 | -1.01584 | 0.139 | 0.391 | 2.29E-169 |
| RBIS | 6.59E-174 | -0.48351 | 0.007 | 0.188 | 2.41E-169 |
| ATP5ME | 8.98E-174 | -0.5014 | 0.011 | 0.196 | 3.28E-169 |
| CALD1 | 1.08E-172 | 0.604681 | 0.83 | 0.749 | 3.96E-168 |
| AKAP13 | 7.25E-172 | 0.920047 | 0.575 | 0.422 | 2.65E-167 |
| RNASE4 | 3.72E-171 | -0.75734 | 0.134 | 0.377 | 1.36E-166 |
| GABARAP | 8.40E-171 | -0.65485 | 0.142 | 0.393 | 3.07E-166 |
| GULP1 | 2.03E-170 | 1.128535 | 0.297 | 0.136 | 7.43E-166 |
| COL1A1 | 2.20E-169 | 0.541382 | 0.827 | 0.661 | 8.04E-165 |
| PTPN9 | 6.32E-169 | 1.082059 | 0.21 | 0.073 | 2.31E-164 |
| CCDC39 | 1.47E-168 | 0.908844 | 0.266 | 0.103 | 5.36E-164 |
| EIF3G | 1.58E-168 | -0.75967 | 0.09 | 0.324 | 5.79E-164 |
| CYCS | 2.33E-168 | -0.89988 | 0.069 | 0.287 | 8.52E-164 |
| NFIB | 1.13E-167 | 0.768331 | 0.657 | 0.537 | 4.12E-163 |
| ATP5MPL | 1.37E-167 | -0.48096 | 0.009 | 0.185 | 5.01E-163 |
| MT-CYB | 1.91E-167 | -0.73302 | 0.926 | 0.9 | 6.98E-163 |
| SMYD3 | 2.24E-167 | 1.08259 | 0.28 | 0.118 | 8.18E-163 |
| GRIA1 | 7.81E-167 | 0.923893 | 0.117 | 0.023 | 2.86E-162 |
| AC105402 | 9.53E-167 | 0.673851 | 0.103 | 0.017 | 3.48E-162 |
| RPLP2 | 3.64E-166 | -0.64698 | 0.743 | 0.881 | 1.33E-161 |
| RAN | 3.80E-166 | -0.79705 | 0.137 | 0.385 | 1.39E-161 |
| COL6A6 | 7.48E-166 | 1.214742 | 0.224 | 0.083 | 2.73E-161 |
| SSR2 | 1.26E-165 | -0.8355 | 0.159 | 0.413 | 4.61E-161 |
| SDHD | 3.71E-165 | -0.60605 | 0.007 | 0.18 | 1.36E-160 |
| SOD1 | 8.76E-165 | -0.88483 | 0.217 | 0.483 | 3.20E-160 |
| TPI1 | 9.57E-165 | -0.79699 | 0.116 | 0.358 | 3.50E-160 |
| TAGLN2 | 1.25E-164 | -0.93389 | 0.226 | 0.484 | 4.58E-160 |

| | | | | | |
|---|---|---|---|---|---|
| TCF12 | 1.33E-164 | 0.968026 | 0.366 | 0.186 | 4.85E-160 |
| CCDC85B | 1.56E-164 | -0.60935 | 0.039 | 0.237 | 5.71E-160 |
| MZT2B | 1.88E-164 | -0.6124 | 0.102 | 0.338 | 6.88E-160 |
| GSTP1 | 2.52E-164 | -0.91105 | 0.309 | 0.561 | 9.22E-160 |
| RACK1 | 7.63E-164 | -0.85946 | 0.487 | 0.634 | 2.79E-159 |
| ATP5MF | 7.80E-164 | -0.44992 | 0.008 | 0.181 | 2.85E-159 |
| GMDS-AS1 | 1.21E-163 | 0.931729 | 0.189 | 0.059 | 4.41E-159 |
| MBD5 | 2.26E-162 | 1.013147 | 0.291 | 0.13 | 8.27E-158 |
| PLP2 | 4.29E-161 | -0.83167 | 0.075 | 0.291 | 1.57E-156 |
| FAM129A | 8.12E-160 | 0.778881 | 0.3 | 0.127 | 2.97E-155 |
| ATP5MC1 | 2.14E-159 | -0.42374 | 0.006 | 0.172 | 7.83E-155 |
| HSPA1A | 3.72E-159 | -1.6947 | 0.129 | 0.355 | 1.36E-154 |
| TRAPPC5 | 3.77E-159 | -0.44386 | 0.014 | 0.189 | 1.38E-154 |
| KCNQ1 | 6.28E-159 | 0.678778 | 0.124 | 0.027 | 2.29E-154 |
| RPL27 | 2.16E-158 | -0.76595 | 0.533 | 0.745 | 7.89E-154 |
| RNF115 | 3.40E-158 | 1.264281 | 0.395 | 0.237 | 1.24E-153 |
| VDAC2 | 9.42E-158 | -0.68376 | 0.065 | 0.277 | 3.44E-153 |
| BCAS3 | 2.92E-157 | 0.848485 | 0.312 | 0.14 | 1.07E-152 |
| DDAH2 | 2.92E-157 | -0.82128 | 0.155 | 0.4 | 1.07E-152 |
| VPS13B | 4.24E-157 | 0.904842 | 0.33 | 0.159 | 1.55E-152 |
| CYBA | 4.92E-157 | -0.56968 | 0.081 | 0.3 | 1.80E-152 |
| FAM13A | 8.31E-157 | 1.057715 | 0.396 | 0.229 | 3.04E-152 |
| 11-Sep | 9.21E-157 | 0.666563 | 0.288 | 0.119 | 3.37E-152 |
| STAG1 | 1.65E-155 | 0.916654 | 0.429 | 0.25 | 6.04E-151 |
| APBB2 | 3.29E-155 | 1.015552 | 0.328 | 0.163 | 1.20E-150 |
| ANKS1B | 5.70E-155 | 0.924475 | 0.239 | 0.092 | 2.08E-150 |
| ZFP36 | 2.08E-154 | -1.00625 | 0.669 | 0.742 | 7.59E-150 |
| PRDX5 | 2.33E-154 | -0.84508 | 0.152 | 0.396 | 8.51E-150 |
| LY6E | 2.38E-154 | -0.85428 | 0.11 | 0.336 | 8.72E-150 |
| GYPC | 5.25E-154 | -0.81016 | 0.109 | 0.336 | 1.92E-149 |
| TNFRSF12/ | 1.20E-153 | -0.87038 | 0.03 | 0.209 | 4.39E-149 |
| HLA-A | 1.37E-153 | -0.88253 | 0.31 | 0.554 | 5.00E-149 |
| COX4I1 | 1.81E-153 | -0.87213 | 0.246 | 0.507 | 6.62E-149 |
| MIR99AHG | 2.27E-153 | 1.117008 | 0.468 | 0.319 | 8.30E-149 |
| H19 | 2.75E-153 | -0.78257 | 0.025 | 0.203 | 1.00E-148 |
| POLR2F | 3.40E-153 | -0.74696 | 0.013 | 0.182 | 1.24E-148 |
| COX7C | 3.79E-153 | -0.95056 | 0.23 | 0.489 | 1.38E-148 |
| TOMM5 | 6.26E-153 | -0.46627 | 0.031 | 0.216 | 2.29E-148 |
| VIT | 7.19E-153 | 0.966866 | 0.394 | 0.221 | 2.63E-148 |
| MRPS24 | 1.23E-152 | -0.4269 | 0.008 | 0.17 | 4.51E-148 |
| MED13L | 1.83E-152 | 0.966561 | 0.472 | 0.31 | 6.71E-148 |
| C20orf194 | 3.65E-152 | 0.917326 | 0.256 | 0.106 | 1.34E-147 |
| MAGI2 | 5.86E-152 | 1.024623 | 0.327 | 0.162 | 2.14E-147 |
| ZEB1 | 4.90E-149 | 0.923031 | 0.448 | 0.274 | 1.79E-144 |
| RBM25 | 1.91E-148 | 0.777161 | 0.621 | 0.497 | 6.99E-144 |
| CZIB | 2.70E-148 | -0.41424 | 0.007 | 0.164 | 9.86E-144 |
| RFWD2 | 2.88E-148 | 0.789639 | 0.211 | 0.075 | 1.05E-143 |

| | | | | | |
|---|---|---|---|---|---|
| COL15A1 | 1.49E-147 | 1.086402 | 0.369 | 0.211 | 5.43E-143 |
| CYTH3 | 2.05E-147 | 1.016416 | 0.299 | 0.145 | 7.48E-143 |
| PDLIM2 | 6.30E-147 | -0.72128 | 0.07 | 0.274 | 2.30E-142 |
| HLA-C | 4.41E-146 | -0.83651 | 0.234 | 0.478 | 1.61E-141 |
| FXYD5 | 1.08E-145 | -0.82869 | 0.045 | 0.23 | 3.96E-141 |
| SEPTIN11 | 1.55E-145 | -0.44723 | 0.016 | 0.181 | 5.66E-141 |
| NLGN1 | 3.79E-145 | 0.857938 | 0.168 | 0.052 | 1.39E-140 |
| ELF1 | 7.00E-145 | 0.971174 | 0.482 | 0.325 | 2.56E-140 |
| AC062004 | 1.10E-144 | 0.869233 | 0.154 | 0.045 | 4.01E-140 |
| CDKAL1 | 1.10E-144 | 0.806454 | 0.274 | 0.12 | 4.01E-140 |
| FXYD1 | 1.64E-144 | -0.82356 | 0.156 | 0.389 | 5.98E-140 |
| DIAPH2 | 2.29E-144 | 0.905504 | 0.398 | 0.228 | 8.37E-140 |
| MYO9A | 4.32E-144 | 0.900156 | 0.389 | 0.219 | 1.58E-139 |
| SLC25A5 | 7.89E-144 | -0.67911 | 0.064 | 0.26 | 2.88E-139 |
| RPS20 | 1.41E-143 | -0.64975 | 0.626 | 0.801 | 5.14E-139 |
| RPS11 | 1.84E-143 | -0.59873 | 0.656 | 0.793 | 6.71E-139 |
| PARK7 | 2.75E-143 | -0.7489 | 0.179 | 0.423 | 1.00E-138 |
| C7 | 1.19E-142 | -1.25793 | 0.13 | 0.331 | 4.33E-138 |
| ATP6V1F | 2.02E-142 | -0.74715 | 0.054 | 0.244 | 7.38E-138 |
| NFATC2 | 2.35E-142 | 1.137088 | 0.369 | 0.209 | 8.58E-138 |
| TRMT112 | 2.92E-142 | -0.75236 | 0.106 | 0.324 | 1.07E-137 |
| MAGI2-AS | 4.49E-142 | -0.42265 | 0.011 | 0.168 | 1.64E-137 |
| CSTB | 6.04E-142 | -0.84906 | 0.144 | 0.375 | 2.21E-137 |
| MICOS13 | 9.57E-142 | -0.40092 | 0.008 | 0.16 | 3.50E-137 |
| PTPN13 | 1.37E-141 | 1.000282 | 0.348 | 0.186 | 5.01E-137 |
| PPP1R14B | 6.51E-141 | -0.57605 | 0.027 | 0.194 | 2.38E-136 |
| LUC7L3 | 1.59E-140 | 0.876428 | 0.523 | 0.381 | 5.79E-136 |
| GHR | 1.80E-140 | 0.97057 | 0.341 | 0.181 | 6.57E-136 |
| IER3 | 5.81E-140 | -1.65675 | 0.044 | 0.216 | 2.12E-135 |
| ATP6V0E1 | 1.01E-139 | -0.7406 | 0.185 | 0.428 | 3.68E-135 |
| SEC11A | 1.18E-139 | -0.75739 | 0.153 | 0.387 | 4.31E-135 |
| ARHGAP26 | 2.09E-139 | 0.918212 | 0.277 | 0.127 | 7.64E-135 |
| CUTA | 5.54E-139 | -0.69605 | 0.074 | 0.273 | 2.03E-134 |
| GNG10 | 2.53E-138 | -0.38303 | 0.014 | 0.171 | 9.24E-134 |
| TENT5A | 4.12E-138 | -0.64872 | 0.011 | 0.163 | 1.51E-133 |
| UAP1 | 4.80E-138 | -0.85015 | 0.189 | 0.416 | 1.75E-133 |
| UBE2V1 | 7.89E-138 | -0.39406 | 0.027 | 0.195 | 2.89E-133 |
| SNRPB | 8.11E-138 | -0.61353 | 0.091 | 0.298 | 2.96E-133 |
| PRELID1 | 1.59E-137 | -0.44953 | 0.042 | 0.22 | 5.81E-133 |
| MYL12A | 1.77E-137 | -0.68868 | 0.184 | 0.424 | 6.48E-133 |
| ZNF706 | 2.41E-137 | -0.63719 | 0.085 | 0.287 | 8.81E-133 |
| LRMDA | 2.85E-137 | 0.904656 | 0.304 | 0.15 | 1.04E-132 |
| SNHG16 | 4.44E-137 | -0.38693 | 0.005 | 0.149 | 1.62E-132 |
| IGFBP5 | 6.47E-137 | -1.04108 | 0.428 | 0.628 | 2.36E-132 |
| GABARAPL | 6.80E-137 | -0.7218 | 0.129 | 0.353 | 2.48E-132 |
| HINT1 | 1.02E-136 | -0.83789 | 0.171 | 0.407 | 3.73E-132 |
| SEC61B | 1.08E-136 | -0.71444 | 0.164 | 0.398 | 3.95E-132 |

| | | | | | |
|---|---|---|---|---|---|
| GPHN | 2.00E-136 | 0.823176 | 0.273 | 0.122 | 7.32E-132 |
| SSH2 | 2.29E-136 | 0.905557 | 0.312 | 0.157 | 8.36E-132 |
| RABAC1 | 2.94E-136 | -0.79817 | 0.205 | 0.444 | 1.07E-131 |
| UQCRB | 4.28E-136 | -0.77546 | 0.203 | 0.446 | 1.57E-131 |
| BRK1 | 4.45E-136 | -0.76779 | 0.101 | 0.311 | 1.63E-131 |
| FAM35A | 4.50E-136 | 0.749382 | 0.199 | 0.072 | 1.65E-131 |
| EPB41L4A- | 4.92E-136 | -0.67171 | 0.05 | 0.23 | 1.80E-131 |
| PLEKHA5 | 7.08E-136 | 0.939928 | 0.446 | 0.294 | 2.59E-131 |
| SPCS2 | 8.86E-136 | -0.62682 | 0.138 | 0.367 | 3.24E-131 |
| S100A16 | 1.51E-135 | -0.80338 | 0.056 | 0.239 | 5.53E-131 |
| DAD1 | 1.54E-135 | -0.69244 | 0.175 | 0.411 | 5.63E-131 |
| UQCRH | 1.55E-135 | -0.74874 | 0.119 | 0.336 | 5.66E-131 |
| PTPRD | 2.11E-135 | 0.832112 | 0.27 | 0.123 | 7.71E-131 |
| RNH1 | 2.73E-135 | -0.6983 | 0.162 | 0.39 | 9.99E-131 |
| HERC1 | 3.14E-135 | 0.85418 | 0.281 | 0.133 | 1.15E-130 |
| EIF3F | 3.52E-135 | -0.63842 | 0.114 | 0.329 | 1.29E-130 |
| VKORC1 | 4.14E-135 | -0.68706 | 0.136 | 0.362 | 1.51E-130 |
| PPDPF | 6.07E-135 | -0.68554 | 0.18 | 0.415 | 2.22E-130 |
| VEGFB | 9.84E-135 | -0.5462 | 0.088 | 0.29 | 3.60E-130 |
| EIF3K | 1.28E-134 | -0.77595 | 0.156 | 0.384 | 4.68E-130 |
| LUM | 1.33E-134 | -0.83584 | 0.795 | 0.812 | 4.88E-130 |
| EDF1 | 2.48E-134 | -0.75049 | 0.22 | 0.464 | 9.07E-130 |
| RRAS | 3.73E-134 | -0.51291 | 0.069 | 0.262 | 1.36E-129 |
| SIK3 | 3.76E-134 | 1.010643 | 0.359 | 0.203 | 1.37E-129 |
| COMMD6 | 4.42E-134 | -0.84217 | 0.182 | 0.42 | 1.61E-129 |
| CDK2AP1 | 4.55E-134 | -0.39079 | 0.029 | 0.195 | 1.66E-129 |
| SAP18 | 6.43E-134 | -0.72262 | 0.161 | 0.393 | 2.35E-129 |
| EXT1 | 1.59E-133 | 0.769406 | 0.388 | 0.218 | 5.83E-129 |
| ATP5PB | 1.76E-133 | -0.35768 | 0.005 | 0.146 | 6.42E-129 |
| LGALS3 | 2.42E-133 | -0.72364 | 0.492 | 0.666 | 8.85E-129 |
| NPC2 | 2.48E-133 | -0.8333 | 0.191 | 0.427 | 9.07E-129 |
| C4orf3 | 3.68E-133 | -0.71029 | 0.13 | 0.35 | 1.35E-128 |
| BRI3 | 8.27E-133 | -0.4956 | 0.117 | 0.328 | 3.02E-128 |
| MRPL51 | 8.73E-133 | -0.70311 | 0.051 | 0.23 | 3.19E-128 |
| CTSL | 1.42E-132 | -0.84386 | 0.199 | 0.425 | 5.21E-128 |
| TSC22D1 | 2.12E-132 | -0.94342 | 0.121 | 0.325 | 7.73E-128 |
| AC013394 | 2.40E-132 | 0.682148 | 0.183 | 0.064 | 8.79E-128 |
| MAFF | 4.43E-132 | -0.8708 | 0.12 | 0.323 | 1.62E-127 |
| CKB | 8.22E-132 | -0.64914 | 0.049 | 0.224 | 3.01E-127 |
| FCGRT | 1.37E-131 | -0.76612 | 0.199 | 0.429 | 5.02E-127 |
| TXN | 4.57E-131 | -0.91718 | 0.279 | 0.509 | 1.67E-126 |
| COPE | 1.07E-130 | -0.58916 | 0.073 | 0.265 | 3.93E-126 |
| AL445250. | 1.26E-130 | 0.808453 | 0.101 | 0.022 | 4.62E-126 |
| NDUFB10 | 3.44E-130 | -0.71281 | 0.097 | 0.301 | 1.26E-125 |
| BSG | 4.08E-130 | -0.71117 | 0.118 | 0.33 | 1.49E-125 |
| LARGE1 | 4.36E-130 | 0.817838 | 0.259 | 0.116 | 1.59E-125 |
| ARL2 | 4.87E-130 | -0.68336 | 0.066 | 0.252 | 1.78E-125 |

| | | | | | |
|---|---|---|---|---|---|
| RAMP2 | 1.06E-129 | -0.87224 | 0.094 | 0.286 | 3.87E-125 |
| EVA1B | 3.93E-129 | -0.39796 | 0.044 | 0.215 | 1.44E-124 |
| ATP5F1A | 7.83E-129 | -0.35421 | 0.006 | 0.144 | 2.86E-124 |
| MT1E | 3.64E-128 | -1.19115 | 0.338 | 0.544 | 1.33E-123 |
| FKBP8 | 5.49E-128 | -0.61927 | 0.079 | 0.271 | 2.01E-123 |
| KRT10 | 8.67E-128 | -0.38318 | 0.1 | 0.305 | 3.17E-123 |
| MT-CO2 | 3.27E-127 | -0.58913 | 0.923 | 0.896 | 1.19E-122 |
| EXOC6B | 3.68E-127 | 0.77643 | 0.255 | 0.114 | 1.34E-122 |
| FIS1 | 3.86E-127 | -0.65557 | 0.096 | 0.296 | 1.41E-122 |
| C5orf42 | 6.78E-127 | 0.697607 | 0.19 | 0.07 | 2.48E-122 |
| TMSB10 | 8.05E-127 | -0.4093 | 0.887 | 0.926 | 2.94E-122 |
| CLASP2 | 8.98E-127 | 0.818323 | 0.24 | 0.105 | 3.28E-122 |
| NR2F1 | 1.22E-126 | -0.69038 | 0.016 | 0.161 | 4.45E-122 |
| GNG5 | 2.60E-126 | -0.64115 | 0.107 | 0.311 | 9.51E-122 |
| PRXL2C | 2.62E-126 | -0.36762 | 0.006 | 0.142 | 9.59E-122 |
| EHBP1 | 3.44E-126 | 1.043994 | 0.38 | 0.233 | 1.26E-121 |
| BPTF | 6.22E-126 | 0.828771 | 0.489 | 0.347 | 2.27E-121 |
| IFI27 | 3.70E-125 | -0.94428 | 0.136 | 0.339 | 1.35E-120 |
| CFL1 | 5.16E-125 | -0.80325 | 0.298 | 0.528 | 1.88E-120 |
| AC005237. | 9.23E-125 | 0.682178 | 0.154 | 0.049 | 3.37E-120 |
| AGAP1 | 9.51E-125 | 0.756375 | 0.341 | 0.18 | 3.47E-120 |
| SRSF11 | 1.11E-124 | 0.699358 | 0.584 | 0.475 | 4.06E-120 |
| CLTB | 1.17E-124 | -0.46371 | 0.095 | 0.293 | 4.27E-120 |
| CAMK2N1 | 1.77E-124 | -0.56829 | 0.065 | 0.242 | 6.48E-120 |
| NEDD8 | 2.06E-124 | -0.71522 | 0.143 | 0.361 | 7.53E-120 |
| PTMS | 3.18E-124 | -0.59131 | 0.16 | 0.377 | 1.16E-119 |
| FOXP2 | 6.05E-124 | 0.80339 | 0.219 | 0.091 | 2.21E-119 |
| FBLN2 | 8.38E-124 | 0.566161 | 0.714 | 0.603 | 3.06E-119 |
| NTRK2 | 1.23E-123 | 0.698221 | 0.506 | 0.35 | 4.49E-119 |
| ATRAID | 1.31E-123 | -0.59577 | 0.12 | 0.33 | 4.78E-119 |
| EIF3E | 1.52E-123 | -0.74514 | 0.196 | 0.428 | 5.57E-119 |
| SNX3 | 1.67E-123 | -0.5817 | 0.164 | 0.385 | 6.09E-119 |
| HLA-B | 3.00E-123 | -0.80127 | 0.296 | 0.528 | 1.10E-118 |
| NDUFA4 | 4.45E-123 | -0.72579 | 0.238 | 0.478 | 1.63E-118 |
| GUK1 | 5.13E-123 | -0.58336 | 0.162 | 0.385 | 1.88E-118 |
| C19orf53 | 1.05E-122 | -0.6537 | 0.09 | 0.281 | 3.83E-118 |
| AL359915. | 1.14E-122 | 0.5743 | 0.109 | 0.026 | 4.17E-118 |
| C1QTNF3 | 1.64E-122 | -1.12129 | 0.016 | 0.156 | 6.00E-118 |
| H2AFJ | 1.68E-122 | -0.71837 | 0.169 | 0.39 | 6.12E-118 |
| VPS28 | 2.01E-122 | -0.63142 | 0.111 | 0.314 | 7.33E-118 |
| SKP1 | 2.36E-122 | -0.74566 | 0.248 | 0.482 | 8.64E-118 |
| IRF1 | 2.64E-122 | -0.90472 | 0.119 | 0.312 | 9.64E-118 |
| METRNL | 2.79E-122 | -0.54677 | 0.069 | 0.248 | 1.02E-117 |
| EIF5A | 6.51E-122 | -0.57098 | 0.1 | 0.294 | 2.38E-117 |
| SPARCL1 | 8.76E-122 | -1.17739 | 0.208 | 0.393 | 3.20E-117 |
| KCNQ1OT1 | 1.03E-121 | 1.145268 | 0.365 | 0.22 | 3.75E-117 |
| PSMD8 | 1.04E-121 | -0.50658 | 0.094 | 0.289 | 3.81E-117 |

| | | | | | |
|---|---|---|---|---|---|
| NNMT | 2.28E-121 | -1.00153 | 0.398 | 0.586 | 8.32E-117 |
| GPRC5A | 2.47E-121 | -0.69766 | 0.126 | 0.324 | 9.02E-117 |
| PPP1R12B | 3.96E-121 | 0.855875 | 0.26 | 0.124 | 1.45E-116 |
| LAPTM4A | 4.12E-121 | -0.67089 | 0.523 | 0.679 | 1.51E-116 |
| RTF2 | 5.16E-121 | -0.31377 | 0.005 | 0.135 | 1.89E-116 |
| DCTN3 | 9.71E-121 | -0.50055 | 0.066 | 0.245 | 3.55E-116 |
| VAMP2 | 1.38E-120 | -0.64819 | 0.081 | 0.265 | 5.04E-116 |
| TBX15 | 1.42E-120 | 0.785978 | 0.166 | 0.059 | 5.21E-116 |
| CSMD1 | 1.58E-120 | 0.630959 | 0.216 | 0.088 | 5.76E-116 |
| DRAP1 | 2.09E-120 | -0.48689 | 0.076 | 0.26 | 7.64E-116 |
| APRT | 2.15E-120 | -0.55316 | 0.105 | 0.303 | 7.84E-116 |
| PNRC1 | 3.37E-120 | -0.95859 | 0.406 | 0.568 | 1.23E-115 |
| HIGD2A | 4.39E-120 | -0.65563 | 0.074 | 0.254 | 1.60E-115 |
| TMA7 | 5.73E-120 | -0.73596 | 0.247 | 0.487 | 2.10E-115 |
| CBLB | 6.16E-120 | 0.811342 | 0.461 | 0.316 | 2.25E-115 |
| CAMK2D | 7.70E-120 | 1.005749 | 0.44 | 0.308 | 2.81E-115 |
| EEF2 | 9.90E-120 | -0.68954 | 0.328 | 0.562 | 3.62E-115 |
| EEF2K | 1.81E-119 | 0.915693 | 0.247 | 0.115 | 6.60E-115 |
| STMP1 | 3.43E-119 | -0.31343 | 0.006 | 0.135 | 1.26E-114 |
| SRP14 | 8.75E-119 | -0.71783 | 0.406 | 0.616 | 3.20E-114 |
| TNXB | 9.68E-119 | -0.71086 | 0.246 | 0.467 | 3.54E-114 |
| NBL1 | 1.06E-118 | -0.56188 | 0.082 | 0.263 | 3.86E-114 |
| RSPO3 | 1.48E-118 | -0.7132 | 0.057 | 0.22 | 5.42E-114 |
| MSR1 | 2.34E-118 | 0.813303 | 0.139 | 0.043 | 8.55E-114 |
| CD151 | 1.52E-117 | -0.66125 | 0.126 | 0.328 | 5.55E-113 |
| AKT3 | 1.69E-117 | 0.891335 | 0.364 | 0.217 | 6.18E-113 |
| WDR83OS | 3.62E-117 | -0.65067 | 0.129 | 0.334 | 1.32E-112 |
| FOXN3 | 3.73E-117 | 0.887744 | 0.481 | 0.357 | 1.36E-112 |
| CTSF | 6.14E-117 | -0.42909 | 0.093 | 0.281 | 2.24E-112 |
| ALDH1A3 | 6.17E-117 | -0.73783 | 0.059 | 0.224 | 2.26E-112 |
| LRPAP1 | 7.39E-117 | -0.52791 | 0.106 | 0.303 | 2.70E-112 |
| MON2 | 1.07E-116 | 0.847227 | 0.317 | 0.173 | 3.89E-112 |
| TRIR | 1.12E-116 | -0.35576 | 0.117 | 0.318 | 4.10E-112 |
| SH3RF1 | 2.10E-116 | 0.883152 | 0.219 | 0.097 | 7.68E-112 |
| PLAT | 5.06E-116 | -0.74202 | 0.062 | 0.226 | 1.85E-111 |
| SLIT2 | 1.04E-115 | 0.911101 | 0.3 | 0.161 | 3.79E-111 |
| SOS2 | 1.65E-115 | 0.923525 | 0.283 | 0.147 | 6.02E-111 |
| NXT1 | 3.78E-115 | -0.3828 | 0.02 | 0.16 | 1.38E-110 |
| CRYAB | 3.92E-115 | -0.72366 | 0.097 | 0.282 | 1.43E-110 |
| SPIDR | 9.24E-115 | 0.847462 | 0.275 | 0.137 | 3.38E-110 |
| PRMT1 | 7.10E-114 | -0.39694 | 0.045 | 0.202 | 2.59E-109 |
| EIF3I | 1.31E-113 | -0.54572 | 0.105 | 0.295 | 4.79E-109 |
| AL592183. | 1.97E-113 | 0.699026 | 0.169 | 0.062 | 7.21E-109 |
| ANKRD36 | 3.53E-113 | 0.800863 | 0.301 | 0.159 | 1.29E-108 |
| PFDN2 | 4.24E-113 | -0.55427 | 0.093 | 0.276 | 1.55E-108 |
| RPL22L1 | 5.90E-113 | -0.71034 | 0.054 | 0.215 | 2.16E-108 |
| NDUFA11 | 6.85E-113 | -0.62109 | 0.145 | 0.354 | 2.50E-108 |

| | | | | | |
|---|---|---|---|---|---|
| IER2 | 1.33E-112 | -0.96499 | 0.158 | 0.35 | 4.87E-108 |
| MARCKSL1 | 2.22E-112 | -0.47158 | 0.011 | 0.138 | 8.13E-108 |
| COX6B1 | 3.59E-112 | -0.71756 | 0.175 | 0.391 | 1.31E-107 |
| MACROD2 | 9.45E-112 | 0.751929 | 0.19 | 0.076 | 3.45E-107 |
| SAT2 | 1.01E-111 | -0.51435 | 0.075 | 0.249 | 3.68E-107 |
| ARHGAP24 | 1.28E-111 | 0.933517 | 0.353 | 0.215 | 4.67E-107 |
| ANKRD36C | 1.55E-111 | 0.757108 | 0.424 | 0.275 | 5.66E-107 |
| ADD3 | 2.89E-111 | 0.661353 | 0.58 | 0.459 | 1.06E-106 |
| TMEM132 | 3.62E-111 | 0.897657 | 0.213 | 0.094 | 1.32E-106 |
| BLVRB | 4.23E-111 | -0.56883 | 0.115 | 0.309 | 1.55E-106 |
| ATRX | 4.40E-111 | 0.755061 | 0.502 | 0.374 | 1.61E-106 |
| COX6A1 | 5.08E-111 | -0.67428 | 0.166 | 0.379 | 1.86E-106 |
| TSPO | 1.19E-110 | -0.63502 | 0.17 | 0.382 | 4.34E-106 |
| NENF | 3.19E-110 | -0.39237 | 0.136 | 0.34 | 1.17E-105 |
| SNU13 | 7.89E-110 | -0.63797 | 0.117 | 0.311 | 2.88E-105 |
| RPS21 | 9.63E-110 | -0.65633 | 0.524 | 0.699 | 3.52E-105 |
| ZCCHC11 | 1.29E-109 | 0.643448 | 0.203 | 0.084 | 4.70E-105 |
| TMEM59 | 1.65E-109 | -0.65271 | 0.278 | 0.499 | 6.02E-105 |
| RPL4 | 3.80E-109 | -0.73658 | 0.397 | 0.623 | 1.39E-104 |
| SMDT1 | 4.74E-109 | -0.58223 | 0.057 | 0.216 | 1.73E-104 |
| TNKS | 4.90E-109 | 0.78379 | 0.227 | 0.104 | 1.79E-104 |
| 2-Sep | 5.47E-109 | 0.510026 | 0.313 | 0.158 | 2.00E-104 |
| TMEM14C | 7.53E-109 | -0.51478 | 0.125 | 0.323 | 2.75E-104 |
| PAMR1 | 9.33E-109 | -0.56555 | 0.085 | 0.257 | 3.41E-104 |
| RARRES1 | 1.68E-108 | -1.10932 | 0.038 | 0.18 | 6.13E-104 |
| SSR4 | 1.70E-108 | -0.6281 | 0.177 | 0.388 | 6.21E-104 |
| OSTC | 2.57E-108 | -0.48905 | 0.115 | 0.307 | 9.40E-104 |
| INVS | 6.11E-108 | 0.782642 | 0.165 | 0.062 | 2.23E-103 |
| MYL12B | 6.66E-108 | -0.65487 | 0.372 | 0.579 | 2.44E-103 |
| RPS4Y1 | 9.09E-108 | -0.65082 | 0.012 | 0.136 | 3.32E-103 |
| COL4A2 | 1.17E-107 | 0.886977 | 0.492 | 0.376 | 4.28E-103 |
| RAB5IF | 1.80E-107 | -0.26693 | 0.005 | 0.123 | 6.57E-103 |
| HNRNPA1 | 1.88E-107 | -0.69397 | 0.217 | 0.436 | 6.86E-103 |
| RBM8A | 2.19E-107 | -0.54038 | 0.12 | 0.311 | 8.00E-103 |
| TCF7L2 | 2.23E-107 | 0.844351 | 0.383 | 0.248 | 8.16E-103 |
| MYDGF | 2.51E-107 | -0.41966 | 0.128 | 0.324 | 9.17E-103 |
| CARMIL1 | 2.85E-107 | 0.932228 | 0.296 | 0.162 | 1.04E-102 |
| ABCA8 | 3.77E-107 | 0.805601 | 0.53 | 0.415 | 1.38E-102 |
| CCDC71L | 3.84E-107 | -0.89719 | 0.063 | 0.218 | 1.40E-102 |
| SPCS1 | 4.19E-107 | -0.48114 | 0.115 | 0.306 | 1.53E-102 |
| UBE2S | 4.32E-107 | -0.35707 | 0.012 | 0.137 | 1.58E-102 |
| NDUFB11 | 6.36E-107 | -0.56141 | 0.102 | 0.285 | 2.32E-102 |
| BZW1 | 7.62E-107 | -0.49238 | 0.15 | 0.352 | 2.78E-102 |
| PSMA6 | 8.90E-107 | -0.27339 | 0.032 | 0.174 | 3.25E-102 |
| CTGF | 1.25E-106 | 0.541873 | 0.299 | 0.149 | 4.56E-102 |
| BTG1 | 1.90E-106 | -0.6124 | 0.179 | 0.38 | 6.94E-102 |
| C16orf45 | 2.52E-106 | 0.552831 | 0.248 | 0.115 | 9.22E-102 |

| | | | | | |
|---|---|---|---|---|---|
| PLAAT3 | 2.94E-106 | -0.3149 | 0.004 | 0.119 | 1.07E-101 |
| MT-ND4 | 4.32E-106 | -0.43037 | 0.95 | 0.931 | 1.58E-101 |
| BANF1 | 4.49E-106 | -0.52448 | 0.064 | 0.226 | 1.64E-101 |
| PHB2 | 5.56E-106 | -0.49473 | 0.06 | 0.219 | 2.03E-101 |
| IMMP2L | 8.11E-106 | 0.903806 | 0.27 | 0.142 | 2.96E-101 |
| SRSF9 | 8.32E-106 | -0.28038 | 0.069 | 0.236 | 3.04E-101 |
| HSPA1B | 1.16E-105 | -1.10628 | 0.085 | 0.248 | 4.25E-101 |
| EXOC4 | 1.45E-105 | 0.788696 | 0.258 | 0.129 | 5.31E-101 |
| FGF10-AS1 | 2.54E-105 | 0.67328 | 0.114 | 0.033 | 9.29E-101 |
| RSL24D1 | 2.70E-105 | -0.56996 | 0.109 | 0.293 | 9.87E-101 |
| AC002480 | 3.00E-105 | 0.608835 | 0.2 | 0.083 | 1.10E-100 |
| MAF1 | 5.18E-105 | -0.31131 | 0.028 | 0.164 | 1.89E-100 |
| UXT | 7.79E-105 | -0.60162 | 0.072 | 0.236 | 2.85E-100 |
| MXRA7 | 9.59E-105 | -0.38251 | 0.121 | 0.311 | 3.51E-100 |
| RPS29 | 1.13E-104 | -0.52718 | 0.668 | 0.802 | 4.14E-100 |
| SLC9A9 | 1.65E-104 | 0.944837 | 0.248 | 0.126 | 6.03E-100 |
| KLF2 | 2.12E-104 | -1.02609 | 0.197 | 0.375 | 7.76E-100 |
| CYB5A | 3.25E-104 | -0.5655 | 0.13 | 0.32 | 1.19E-99 |
| CD55 | 4.50E-104 | -1.1032 | 0.139 | 0.311 | 1.64E-99 |
| EIF3L | 4.92E-104 | -0.57276 | 0.06 | 0.217 | 1.80E-99 |
| RAB34 | 5.26E-104 | -0.49178 | 0.068 | 0.23 | 1.92E-99 |
| ANAPC11 | 6.62E-104 | -0.62104 | 0.108 | 0.289 | 2.42E-99 |
| SCAPER | 8.83E-104 | 0.784475 | 0.258 | 0.13 | 3.23E-99 |
| PFN1 | 1.33E-103 | -0.54086 | 0.183 | 0.391 | 4.87E-99 |
| SEC61G | 2.47E-103 | -0.58777 | 0.122 | 0.31 | 9.02E-99 |
| DEPP1 | 5.01E-103 | -1.2003 | 0.007 | 0.121 | 1.83E-98 |
| SF3B5 | 5.23E-103 | -0.55908 | 0.095 | 0.27 | 1.91E-98 |
| BRAF | 7.13E-103 | 0.877515 | 0.302 | 0.171 | 2.61E-98 |
| NF1 | 7.32E-103 | 0.816118 | 0.311 | 0.178 | 2.67E-98 |
| FMO2 | 7.78E-103 | -0.90048 | 0.09 | 0.25 | 2.85E-98 |
| RGS16 | 8.74E-103 | -1.06573 | 0.028 | 0.158 | 3.19E-98 |
| MRFAP1 | 1.19E-102 | -0.60071 | 0.175 | 0.383 | 4.36E-98 |
| OST4 | 1.19E-102 | -0.66425 | 0.214 | 0.432 | 4.36E-98 |
| RFX3 | 1.48E-102 | 0.710916 | 0.173 | 0.069 | 5.42E-98 |
| ANTXR1 | 2.32E-102 | 0.80436 | 0.293 | 0.161 | 8.46E-98 |
| AC069208 | 7.24E-102 | 0.633809 | 0.182 | 0.072 | 2.65E-97 |
| NID1 | 8.07E-102 | 0.778527 | 0.429 | 0.299 | 2.95E-97 |
| KIF1B | 1.50E-101 | 0.806041 | 0.347 | 0.212 | 5.47E-97 |
| COX5B | 2.43E-101 | -0.68485 | 0.176 | 0.382 | 8.89E-97 |
| CDC26 | 3.86E-101 | -0.41468 | 0.02 | 0.146 | 1.41E-96 |
| ADAMTSL1 | 4.26E-101 | 0.815091 | 0.233 | 0.112 | 1.56E-96 |
| GET3 | 4.48E-101 | -0.2503 | 0.004 | 0.113 | 1.64E-96 |
| USP33 | 4.88E-101 | 0.862524 | 0.295 | 0.168 | 1.78E-96 |
| VAMP5 | 5.15E-101 | -0.64352 | 0.092 | 0.262 | 1.88E-96 |
| SLC3A2 | 5.57E-101 | -0.41316 | 0.048 | 0.193 | 2.04E-96 |
| PHPT1 | 5.57E-101 | -0.6834 | 0.134 | 0.324 | 2.04E-96 |
| TSHZ2 | 8.87E-101 | 0.753581 | 0.599 | 0.527 | 3.24E-96 |

| | | | | | |
|---|---|---|---|---|---|
| CHD9 | 1.08E-100 | 0.726025 | 0.513 | 0.404 | 3.95E-96 |
| MAP3K5 | 1.53E-100 | 0.780293 | 0.21 | 0.095 | 5.60E-96 |
| NFKBIA | 1.65E-100 | -0.90634 | 0.418 | 0.581 | 6.02E-96 |
| NDUFAF3 | 2.16E-100 | -0.47379 | 0.063 | 0.218 | 7.90E-96 |
| CAV1 | 4.33E-100 | -0.69937 | 0.249 | 0.449 | 1.58E-95 |
| IFITM2 | 4.61E-100 | -0.69126 | 0.246 | 0.457 | 1.69E-95 |
| ERP29 | 4.67E-100 | -0.53273 | 0.081 | 0.246 | 1.71E-95 |
| LAMTOR4 | 4.68E-100 | -0.6005 | 0.144 | 0.337 | 1.71E-95 |
| SIK2 | 5.15E-100 | 0.824192 | 0.276 | 0.149 | 1.88E-95 |
| SCMH1 | 5.56E-100 | 0.772134 | 0.213 | 0.098 | 2.03E-95 |
| MTRNR2L8 | 5.94E-100 | 1.240249 | 0.49 | 0.391 | 2.17E-95 |
| GCSH | 6.35E-100 | -0.31729 | 0.01 | 0.126 | 2.32E-95 |
| ERH | 8.22E-100 | -0.52964 | 0.12 | 0.304 | 3.01E-95 |
| NCOA1 | 9.48E-100 | 0.848968 | 0.329 | 0.199 | 3.46E-95 |
| PLAAT4 | 9.72E-100 | -0.2766 | 0.003 | 0.11 | 3.55E-95 |
| MAP2K2 | 1.09E-99 | -0.30715 | 0.061 | 0.215 | 4.00E-95 |
| DDT | 1.21E-99 | -0.52681 | 0.053 | 0.201 | 4.41E-95 |
| PAN3 | 1.48E-99 | 0.764281 | 0.293 | 0.159 | 5.40E-95 |
| CD81 | 1.80E-99 | -0.70153 | 0.52 | 0.614 | 6.57E-95 |
| CXCL2 | 1.87E-99 | -1.65195 | 0.055 | 0.192 | 6.83E-95 |
| ESR1 | 2.03E-99 | 0.711852 | 0.139 | 0.049 | 7.43E-95 |
| SIVA1 | 3.29E-99 | -0.54836 | 0.095 | 0.267 | 1.20E-94 |
| EIF4A2 | 3.58E-99 | -0.60761 | 0.191 | 0.396 | 1.31E-94 |
| TASP1 | 5.79E-99 | 0.803302 | 0.205 | 0.094 | 2.12E-94 |
| MIDN | 6.55E-99 | -0.57171 | 0.101 | 0.269 | 2.40E-94 |
| ACACB | 8.87E-99 | 0.648509 | 0.268 | 0.137 | 3.24E-94 |
| IL16 | 9.68E-99 | 0.682035 | 0.194 | 0.085 | 3.54E-94 |
| KDM6B | 9.91E-99 | -0.58333 | 0.046 | 0.186 | 3.62E-94 |
| NDUFC2 | 1.61E-98 | -0.4574 | 0.146 | 0.341 | 5.87E-94 |
| HTRA1 | 1.93E-98 | -0.5464 | 0.23 | 0.435 | 7.06E-94 |
| PGLS | 2.10E-98 | -0.29379 | 0.06 | 0.212 | 7.69E-94 |
| SFRP2 | 2.42E-98 | -1.24161 | 0.167 | 0.328 | 8.86E-94 |
| NUPR1 | 2.69E-98 | -0.73555 | 0.294 | 0.504 | 9.85E-94 |
| PDZRN3 | 2.76E-98 | 0.893463 | 0.215 | 0.103 | 1.01E-93 |
| JUND | 2.86E-98 | -1.37357 | 0.584 | 0.613 | 1.04E-93 |
| PLAUR | 2.97E-98 | -0.61974 | 0.076 | 0.23 | 1.08E-93 |
| CACNA2D1 | 2.99E-98 | 0.778473 | 0.261 | 0.136 | 1.09E-93 |
| CSNK2B | 4.78E-98 | -0.55035 | 0.083 | 0.247 | 1.75E-93 |
| 7-Sep | 8.68E-98 | 0.314406 | 0.394 | 0.219 | 3.17E-93 |
| PCBD1 | 1.11E-97 | -0.45163 | 0.039 | 0.176 | 4.06E-93 |
| PSMA2 | 1.68E-97 | -0.32315 | 0.081 | 0.245 | 6.13E-93 |
| COX7A2L | 2.01E-97 | -0.44617 | 0.064 | 0.218 | 7.36E-93 |
| RHOC | 2.89E-97 | -0.53549 | 0.092 | 0.259 | 1.05E-92 |
| COL4A1 | 3.26E-97 | 0.776186 | 0.503 | 0.383 | 1.19E-92 |
| BTBD9 | 4.23E-97 | 0.640179 | 0.169 | 0.067 | 1.55E-92 |
| PHB | 4.52E-97 | -0.43938 | 0.052 | 0.197 | 1.65E-92 |
| LAMTOR1 | 4.66E-97 | -0.47314 | 0.077 | 0.237 | 1.70E-92 |

| | | | | | |
|---|---|---|---|---|---|
| TRIB1 | 4.80E-97 | -0.51844 | 0.019 | 0.139 | 1.76E-92 |
| MACF1 | 7.53E-97 | 0.800484 | 0.485 | 0.375 | 2.75E-92 |
| DAAM1 | 7.59E-97 | 0.878876 | 0.302 | 0.179 | 2.77E-92 |
| AC007388. | 7.84E-97 | 0.542438 | 0.104 | 0.03 | 2.87E-92 |
| BAZ2B | 1.51E-96 | 0.756002 | 0.337 | 0.203 | 5.52E-92 |
| SEPTIN10 | 1.65E-96 | -0.26478 | 0.008 | 0.118 | 6.03E-92 |
| TOMM20 | 1.95E-96 | -0.41772 | 0.133 | 0.321 | 7.11E-92 |
| CD59 | 2.12E-96 | -0.62594 | 0.264 | 0.47 | 7.74E-92 |
| MAP1LC3A | 2.92E-96 | -0.38008 | 0.074 | 0.232 | 1.07E-91 |
| SLC66A3 | 3.55E-96 | -0.25725 | 0.004 | 0.109 | 1.30E-91 |
| GNB2L1 | 3.94E-96 | -1.83834 | 0.014 | 0.127 | 1.44E-91 |
| MIR100HG | 4.31E-96 | -0.31343 | 0.007 | 0.116 | 1.58E-91 |
| ECH1 | 4.33E-96 | -0.46751 | 0.074 | 0.231 | 1.58E-91 |
| FBXO11 | 4.75E-96 | 0.844261 | 0.32 | 0.19 | 1.73E-91 |
| ETFB | 4.81E-96 | -0.49385 | 0.078 | 0.237 | 1.76E-91 |
| SRI | 5.05E-96 | -0.46145 | 0.14 | 0.33 | 1.85E-91 |
| TBC1D5 | 5.13E-96 | 0.748242 | 0.292 | 0.164 | 1.87E-91 |
| SERF2 | 6.24E-96 | -0.55184 | 0.618 | 0.76 | 2.28E-91 |
| FMNL2 | 7.08E-96 | 0.785039 | 0.298 | 0.168 | 2.59E-91 |
| TMBIM4 | 8.05E-96 | -0.3783 | 0.162 | 0.36 | 2.94E-91 |
| STUB1 | 9.10E-96 | -0.26238 | 0.062 | 0.214 | 3.33E-91 |
| MIR4527H | 1.04E-95 | 0.474002 | 0.118 | 0.037 | 3.82E-91 |
| AIP | 1.16E-95 | -0.47356 | 0.037 | 0.169 | 4.25E-91 |
| SUB1 | 1.23E-95 | -0.56576 | 0.204 | 0.411 | 4.50E-91 |
| PPP3CA | 1.39E-95 | 0.867669 | 0.393 | 0.27 | 5.07E-91 |
| PIM3 | 2.47E-95 | -0.49225 | 0.048 | 0.185 | 9.04E-91 |
| CDV3 | 2.88E-95 | -0.40986 | 0.104 | 0.274 | 1.05E-90 |
| SENP6 | 3.62E-95 | 0.840814 | 0.404 | 0.28 | 1.32E-90 |
| PSMB1 | 3.86E-95 | -0.562 | 0.205 | 0.413 | 1.41E-90 |
| RBM6 | 3.90E-95 | 0.802148 | 0.344 | 0.217 | 1.43E-90 |
| RPS19BP1 | 5.14E-95 | -0.5152 | 0.052 | 0.194 | 1.88E-90 |
| PIK3R1 | 5.21E-95 | 0.71369 | 0.525 | 0.418 | 1.90E-90 |
| BIRC6 | 6.48E-95 | 0.720815 | 0.304 | 0.174 | 2.37E-90 |
| HEBP1 | 7.08E-95 | -0.33433 | 0.052 | 0.196 | 2.59E-90 |
| MEG3 | 8.74E-95 | 0.516695 | 0.604 | 0.491 | 3.19E-90 |
| NOP53 | 1.43E-94 | -0.40543 | 0.205 | 0.408 | 5.22E-90 |
| TUBA1B | 2.49E-94 | -0.66738 | 0.308 | 0.506 | 9.09E-90 |
| FER | 2.65E-94 | 0.71164 | 0.341 | 0.203 | 9.68E-90 |
| C18orf32 | 2.85E-94 | -0.27222 | 0.046 | 0.184 | 1.04E-89 |
| FAM89B | 3.46E-94 | -0.3109 | 0.017 | 0.133 | 1.26E-89 |
| SYNE1 | 3.86E-94 | 0.812141 | 0.482 | 0.378 | 1.41E-89 |
| AMD1 | 4.82E-94 | -0.4881 | 0.076 | 0.229 | 1.76E-89 |
| UBXN1 | 5.17E-94 | -0.54638 | 0.104 | 0.273 | 1.89E-89 |
| CTSZ | 6.60E-94 | -0.27024 | 0.057 | 0.203 | 2.41E-89 |
| AP2M1 | 1.04E-93 | -0.4921 | 0.101 | 0.269 | 3.79E-89 |
| NDUFS5 | 1.29E-93 | -0.60603 | 0.251 | 0.465 | 4.72E-89 |
| PSMB3 | 1.40E-93 | -0.47634 | 0.119 | 0.296 | 5.10E-89 |

| | | | | | |
|---|---|---|---|---|---|
| MAT2A | 1.42E-93 | -0.52392 | 0.068 | 0.215 | 5.18E-89 |
| CFAP69 | 2.37E-93 | 0.734121 | 0.188 | 0.084 | 8.66E-89 |
| FAF1 | 2.53E-93 | 0.777786 | 0.247 | 0.128 | 9.23E-89 |
| MPC2 | 5.42E-93 | -0.4154 | 0.054 | 0.195 | 1.98E-88 |
| PAPD4 | 5.75E-93 | 0.569271 | 0.154 | 0.059 | 2.10E-88 |
| TNRC6A | 1.18E-92 | 0.770674 | 0.352 | 0.225 | 4.31E-88 |
| COX7A2 | 1.24E-92 | -0.66838 | 0.16 | 0.351 | 4.52E-88 |
| KIAA0368 | 1.37E-92 | 0.578297 | 0.226 | 0.106 | 5.00E-88 |
| COPRS | 1.60E-92 | -0.28388 | 0.05 | 0.189 | 5.83E-88 |
| ZNF638 | 2.41E-92 | 0.747075 | 0.403 | 0.277 | 8.82E-88 |
| SNHG15 | 2.81E-92 | -0.43074 | 0.027 | 0.149 | 1.03E-87 |
| PKD2 | 2.92E-92 | 0.918933 | 0.366 | 0.244 | 1.07E-87 |
| GOLGA4 | 3.45E-92 | 0.657937 | 0.528 | 0.415 | 1.26E-87 |
| SELENOW | 5.96E-92 | -0.28509 | 0.157 | 0.35 | 2.18E-87 |
| LAMTOR5 | 6.13E-92 | -0.56539 | 0.109 | 0.279 | 2.24E-87 |
| SNHG7 | 7.80E-92 | -0.30638 | 0.04 | 0.171 | 2.85E-87 |
| CKS1B | 9.42E-92 | -0.49511 | 0.042 | 0.174 | 3.44E-87 |
| COPZ2 | 9.55E-92 | -0.28767 | 0.072 | 0.224 | 3.49E-87 |
| PTGDS | 1.24E-91 | -1.68549 | 0.132 | 0.278 | 4.53E-87 |
| MKL2 | 2.29E-91 | 0.573744 | 0.102 | 0.03 | 8.38E-87 |
| SERPINE1 | 2.53E-91 | -0.99529 | 0.077 | 0.22 | 9.25E-87 |
| AC034206 | 2.65E-91 | 0.419227 | 0.221 | 0.099 | 9.68E-87 |
| ATG7 | 3.70E-91 | 0.679285 | 0.207 | 0.097 | 1.35E-86 |
| PLD1 | 3.73E-91 | 0.752097 | 0.152 | 0.06 | 1.36E-86 |
| PCSK5 | 6.67E-91 | 0.816058 | 0.266 | 0.146 | 2.44E-86 |
| TECR | 7.17E-91 | -0.39006 | 0.032 | 0.157 | 2.62E-86 |
| LARP6 | 8.12E-91 | -0.41233 | 0.084 | 0.24 | 2.97E-86 |
| COX16 | 9.01E-91 | -0.27761 | 0.022 | 0.139 | 3.29E-86 |
| BLOC1S1 | 1.08E-90 | -0.6062 | 0.09 | 0.247 | 3.93E-86 |
| C9orf16 | 1.09E-90 | -0.51175 | 0.036 | 0.163 | 3.97E-86 |
| SFRP4 | 2.49E-90 | -1.49408 | 0.072 | 0.206 | 9.09E-86 |
| CEP112 | 2.51E-90 | 0.736086 | 0.223 | 0.111 | 9.19E-86 |
| SGCD | 3.14E-90 | 0.769598 | 0.24 | 0.126 | 1.15E-85 |
| ARF5 | 4.02E-90 | -0.25125 | 0.031 | 0.155 | 1.47E-85 |
| TMEM230 | 4.61E-90 | -0.45315 | 0.098 | 0.263 | 1.68E-85 |
| PLK3 | 5.24E-90 | -0.40819 | 0.063 | 0.205 | 1.92E-85 |
| RPL38 | 5.42E-90 | -0.65932 | 0.546 | 0.734 | 1.98E-85 |
| TMEM98 | 5.84E-90 | -0.39206 | 0.064 | 0.209 | 2.13E-85 |
| TUFM | 6.03E-90 | -0.37013 | 0.07 | 0.22 | 2.20E-85 |
| FHIT | 6.59E-90 | 0.678067 | 0.188 | 0.084 | 2.41E-85 |
| CDC42BPA | 6.91E-90 | 0.784856 | 0.336 | 0.211 | 2.52E-85 |
| SF3B6 | 8.57E-90 | -0.46977 | 0.106 | 0.274 | 3.13E-85 |
| PLTP | 9.30E-90 | -0.59293 | 0.208 | 0.4 | 3.40E-85 |
| AURKAIP1 | 1.02E-89 | -0.44357 | 0.062 | 0.205 | 3.74E-85 |
| PGAM1 | 1.23E-89 | -0.49237 | 0.035 | 0.161 | 4.50E-85 |
| MAGI3 | 1.28E-89 | 0.76119 | 0.238 | 0.123 | 4.67E-85 |
| PCCA | 2.48E-89 | 0.719319 | 0.212 | 0.103 | 9.07E-85 |

| | | | | | |
|---|---|---|---|---|---|
| EPHX1 | 2.76E-89 | -0.56373 | 0.194 | 0.381 | 1.01E-84 |
| LAMA4 | 2.76E-89 | 0.657288 | 0.559 | 0.472 | 1.01E-84 |
| ATG101 | 5.58E-89 | -0.3312 | 0.028 | 0.148 | 2.04E-84 |
| HNRNPF | 6.86E-89 | -0.4414 | 0.127 | 0.301 | 2.51E-84 |
| ZFAND3 | 7.69E-89 | 0.713935 | 0.275 | 0.154 | 2.81E-84 |
| WISP2 | 1.05E-88 | 0.254405 | 0.391 | 0.221 | 3.84E-84 |
| LINC02802 | 1.18E-88 | -0.26568 | 0.007 | 0.109 | 4.30E-84 |
| ECM1 | 2.38E-88 | -0.64444 | 0.151 | 0.322 | 8.69E-84 |
| FGF14 | 2.81E-88 | 0.775188 | 0.202 | 0.095 | 1.03E-83 |
| COX6C | 3.23E-88 | -0.65908 | 0.276 | 0.49 | 1.18E-83 |
| POMP | 3.40E-88 | -0.49875 | 0.155 | 0.34 | 1.24E-83 |
| SBDS | 3.43E-88 | -0.452 | 0.144 | 0.324 | 1.25E-83 |
| FUNDC2 | 3.51E-88 | -0.44462 | 0.074 | 0.223 | 1.28E-83 |
| AP2S1 | 5.06E-88 | -0.44152 | 0.082 | 0.235 | 1.85E-83 |
| REXO2 | 5.95E-88 | -0.50778 | 0.16 | 0.347 | 2.17E-83 |
| EFCAB2 | 6.21E-88 | 0.794104 | 0.198 | 0.094 | 2.27E-83 |
| RNF7 | 7.54E-88 | -0.43819 | 0.082 | 0.236 | 2.76E-83 |
| CFI | 7.79E-88 | -0.41662 | 0.023 | 0.137 | 2.85E-83 |
| MAP4 | 8.61E-88 | 0.742792 | 0.462 | 0.353 | 3.15E-83 |
| CACNA2D3 | 3.39E-87 | 0.674036 | 0.198 | 0.091 | 1.24E-82 |
| NDUFB8 | 3.63E-87 | -0.41176 | 0.123 | 0.296 | 1.33E-82 |
| MZT2A | 4.96E-87 | -0.29645 | 0.063 | 0.205 | 1.81E-82 |
| PPARG | 5.28E-87 | 0.849859 | 0.287 | 0.169 | 1.93E-82 |
| SOD3 | 7.92E-87 | -0.66161 | 0.279 | 0.45 | 2.89E-82 |
| CYGB | 8.30E-87 | -0.36935 | 0.093 | 0.247 | 3.03E-82 |
| NDUFB7 | 8.81E-87 | -0.59669 | 0.138 | 0.315 | 3.22E-82 |
| PTGES3 | 9.26E-87 | -0.45034 | 0.18 | 0.372 | 3.38E-82 |
| PDLIM1 | 9.85E-87 | -0.5237 | 0.156 | 0.336 | 3.60E-82 |
| TNFSF12 | 1.14E-86 | -0.28461 | 0.042 | 0.169 | 4.16E-82 |
| TGFBR2 | 1.28E-86 | 0.753704 | 0.44 | 0.331 | 4.67E-82 |
| CTNND2 | 1.30E-86 | 0.561723 | 0.116 | 0.039 | 4.76E-82 |
| C16orf62 | 1.78E-86 | 0.466389 | 0.124 | 0.043 | 6.50E-82 |
| ASH1L | 1.78E-86 | 0.771895 | 0.43 | 0.317 | 6.52E-82 |
| CSRNP1 | 2.95E-86 | -0.40472 | 0.072 | 0.216 | 1.08E-81 |
| HSPD1 | 9.46E-86 | -0.58292 | 0.168 | 0.35 | 3.46E-81 |
| FTO | 9.48E-86 | 0.731378 | 0.216 | 0.108 | 3.46E-81 |
| DNAJA1 | 1.48E-85 | -0.60101 | 0.269 | 0.462 | 5.39E-81 |
| PDLIM4 | 1.56E-85 | -0.34792 | 0.034 | 0.155 | 5.70E-81 |
| NDUFB4 | 1.62E-85 | -0.59486 | 0.123 | 0.291 | 5.93E-81 |
| MPG | 3.29E-85 | -0.36025 | 0.05 | 0.181 | 1.20E-80 |
| ADRM1 | 5.03E-85 | -0.3898 | 0.028 | 0.144 | 1.84E-80 |
| DYNLT1 | 5.33E-85 | -0.39997 | 0.08 | 0.229 | 1.95E-80 |
| BAG1 | 6.03E-85 | -0.27481 | 0.053 | 0.186 | 2.20E-80 |
| MID1 | 7.13E-85 | 0.767675 | 0.166 | 0.072 | 2.61E-80 |
| ERC1 | 9.12E-85 | 0.770831 | 0.279 | 0.162 | 3.33E-80 |
| DPP7 | 1.24E-84 | -0.28502 | 0.076 | 0.223 | 4.52E-80 |
| APOD | 1.51E-84 | 0.307046 | 0.919 | 0.752 | 5.50E-80 |

| | | | | | |
|---|---|---|---|---|---|
| MED12L | 1.60E-84 | 0.655798 | 0.199 | 0.092 | 5.85E-80 |
| CCT7 | 1.82E-84 | -0.34914 | 0.071 | 0.214 | 6.64E-80 |
| TLN2 | 2.28E-84 | 0.77659 | 0.306 | 0.189 | 8.35E-80 |
| KDELR1 | 2.83E-84 | -0.45609 | 0.17 | 0.356 | 1.04E-79 |
| ATP6V0B | 3.35E-84 | -0.38236 | 0.06 | 0.196 | 1.23E-79 |
| SELENOM | 3.53E-84 | -0.40885 | 0.27 | 0.462 | 1.29E-79 |
| COMT | 3.70E-84 | -0.33001 | 0.13 | 0.303 | 1.35E-79 |
| GTF2IRD2E | 3.90E-84 | 0.590001 | 0.154 | 0.063 | 1.43E-79 |
| CHMP2A | 4.60E-84 | -0.45421 | 0.113 | 0.277 | 1.68E-79 |
| CHMP5 | 5.02E-84 | -0.39748 | 0.068 | 0.209 | 1.83E-79 |
| FKBP1A | 5.78E-84 | -0.52578 | 0.178 | 0.363 | 2.11E-79 |
| EIF4H | 6.26E-84 | -0.39453 | 0.087 | 0.237 | 2.29E-79 |
| CAB39L | 6.36E-84 | 0.798888 | 0.382 | 0.267 | 2.32E-79 |
| ATP9B | 6.53E-84 | 0.687144 | 0.196 | 0.094 | 2.39E-79 |
| MTRNR2L1 | 7.35E-84 | -0.48986 | 0.401 | 0.603 | 2.69E-79 |
| ANKRD17 | 1.31E-83 | 0.710511 | 0.304 | 0.184 | 4.80E-79 |
| GTF2A2 | 1.46E-83 | -0.34402 | 0.078 | 0.225 | 5.34E-79 |
| BAD | 1.53E-83 | -0.40363 | 0.052 | 0.183 | 5.59E-79 |
| COPS6 | 1.99E-83 | -0.35617 | 0.079 | 0.226 | 7.26E-79 |
| SRP9 | 2.76E-83 | -0.3688 | 0.089 | 0.241 | 1.01E-78 |
| ARC | 2.81E-83 | -0.96921 | 0.01 | 0.107 | 1.03E-78 |
| GSTM5 | 2.97E-83 | -0.49487 | 0.045 | 0.17 | 1.08E-78 |
| UFC1 | 4.62E-83 | -0.44554 | 0.065 | 0.202 | 1.69E-78 |
| PSMD9 | 6.03E-83 | -0.25163 | 0.036 | 0.155 | 2.20E-78 |
| RHEB | 6.73E-83 | -0.28915 | 0.128 | 0.299 | 2.46E-78 |
| ADI1 | 8.18E-83 | -0.46896 | 0.087 | 0.234 | 2.99E-78 |
| ZNF428 | 1.05E-82 | -0.32883 | 0.042 | 0.166 | 3.85E-78 |
| CAVIN3 | 1.11E-82 | -0.26429 | 0.138 | 0.311 | 4.06E-78 |
| TCF21 | 1.40E-82 | -0.47183 | 0.009 | 0.106 | 5.13E-78 |
| LPAR1 | 1.67E-82 | 0.678962 | 0.319 | 0.198 | 6.10E-78 |
| SNHG8 | 1.95E-82 | -0.49661 | 0.119 | 0.282 | 7.13E-78 |
| CDC42 | 1.97E-82 | -0.47304 | 0.234 | 0.431 | 7.21E-78 |
| HIF3A | 2.11E-82 | 0.639308 | 0.134 | 0.052 | 7.71E-78 |
| EIF3H | 2.51E-82 | -0.50625 | 0.155 | 0.333 | 9.19E-78 |
| DLEU2 | 2.63E-82 | 0.701792 | 0.176 | 0.079 | 9.61E-78 |
| PID1 | 2.96E-82 | 0.82202 | 0.441 | 0.34 | 1.08E-77 |
| RARRES2 | 3.53E-82 | -0.73789 | 0.253 | 0.424 | 1.29E-77 |
| DOK6 | 3.66E-82 | 0.616844 | 0.113 | 0.039 | 1.34E-77 |
| NEU1 | 3.82E-82 | -0.52691 | 0.047 | 0.171 | 1.40E-77 |
| OPHN1 | 6.72E-82 | 0.673491 | 0.27 | 0.153 | 2.46E-77 |
| RERE | 7.55E-82 | 0.781651 | 0.307 | 0.19 | 2.76E-77 |
| MRPL43 | 7.79E-82 | -0.447 | 0.05 | 0.178 | 2.85E-77 |
| NDUFA13 | 7.89E-82 | -0.38997 | 0.204 | 0.396 | 2.88E-77 |
| ITFG1 | 9.59E-82 | 0.848762 | 0.29 | 0.177 | 3.50E-77 |
| RER1 | 1.06E-81 | -0.30923 | 0.079 | 0.224 | 3.87E-77 |
| PRRX1 | 1.18E-81 | 0.609856 | 0.563 | 0.484 | 4.32E-77 |
| RABGAP1L | 1.23E-81 | 0.694378 | 0.281 | 0.162 | 4.49E-77 |

| | | | | | |
|---|---|---|---|---|---|
| POLR2J | 1.58E-81 | -0.49177 | 0.059 | 0.191 | 5.76E-77 |
| MT-ND3 | 1.63E-81 | -0.51179 | 0.898 | 0.836 | 5.95E-77 |
| NDUFB5 | 2.34E-81 | -0.36195 | 0.046 | 0.17 | 8.56E-77 |
| NME3 | 2.95E-81 | -0.3819 | 0.081 | 0.227 | 1.08E-76 |
| SRSF2 | 2.97E-81 | -0.37468 | 0.15 | 0.325 | 1.09E-76 |
| MRPL18 | 3.26E-81 | -0.39684 | 0.037 | 0.155 | 1.19E-76 |
| SLC39A1 | 4.07E-81 | -0.38298 | 0.055 | 0.184 | 1.49E-76 |
| TMEM109 | 4.17E-81 | -0.39617 | 0.087 | 0.235 | 1.52E-76 |
| SDF2 | 5.51E-81 | -0.34727 | 0.047 | 0.172 | 2.02E-76 |
| SUMO1 | 7.62E-81 | -0.38412 | 0.1 | 0.255 | 2.79E-76 |
| HNRNPAB | 7.75E-81 | -0.29714 | 0.103 | 0.257 | 2.83E-76 |
| PCED1B | 9.17E-81 | 0.583314 | 0.116 | 0.041 | 3.35E-76 |
| SELENOK | 1.21E-80 | -0.25047 | 0.122 | 0.286 | 4.44E-76 |
| KLF12 | 1.42E-80 | 0.721342 | 0.185 | 0.086 | 5.20E-76 |
| GSTO1 | 1.88E-80 | -0.43968 | 0.102 | 0.256 | 6.89E-76 |
| TTC17 | 2.01E-80 | 0.76172 | 0.328 | 0.209 | 7.34E-76 |
| NCOR1 | 2.01E-80 | 0.788978 | 0.361 | 0.251 | 7.36E-76 |
| CBR4 | 2.14E-80 | 0.633796 | 0.2 | 0.098 | 7.84E-76 |
| USP47 | 2.25E-80 | 0.780964 | 0.329 | 0.212 | 8.22E-76 |
| CENPP | 2.27E-80 | 0.573889 | 0.217 | 0.107 | 8.28E-76 |
| MDH1 | 2.28E-80 | -0.39156 | 0.08 | 0.224 | 8.33E-76 |
| CCDC7 | 2.58E-80 | 0.671542 | 0.14 | 0.056 | 9.43E-76 |
| DYNLRB1 | 4.06E-80 | -0.53611 | 0.15 | 0.323 | 1.48E-75 |
| PSMB6 | 6.61E-80 | -0.42168 | 0.116 | 0.277 | 2.42E-75 |
| UBL5 | 1.20E-79 | -0.55237 | 0.191 | 0.38 | 4.39E-75 |
| PRICKLE2 | 2.03E-79 | 0.710024 | 0.243 | 0.135 | 7.41E-75 |
| ITCH | 2.20E-79 | 0.686451 | 0.233 | 0.125 | 8.04E-75 |
| CAPG | 2.51E-79 | -0.46201 | 0.038 | 0.154 | 9.18E-75 |
| TRAK1 | 2.70E-79 | 0.63461 | 0.172 | 0.079 | 9.86E-75 |
| CAMLG | 2.72E-79 | -0.45542 | 0.123 | 0.283 | 9.94E-75 |
| EIF3M | 2.87E-79 | -0.35595 | 0.091 | 0.24 | 1.05E-74 |
| COL5A1 | 2.94E-79 | 0.720127 | 0.34 | 0.226 | 1.08E-74 |
| MYL9 | 4.51E-79 | -0.64618 | 0.193 | 0.371 | 1.65E-74 |
| TSPAN4 | 4.56E-79 | -0.47548 | 0.179 | 0.361 | 1.67E-74 |
| ISCU | 4.77E-79 | -0.35398 | 0.138 | 0.31 | 1.74E-74 |
| ANG | 7.07E-79 | -0.36398 | 0.06 | 0.189 | 2.58E-74 |
| COBLL1 | 8.19E-79 | 0.758571 | 0.279 | 0.166 | 2.99E-74 |
| CACNA1C | 8.71E-79 | 0.637457 | 0.143 | 0.059 | 3.18E-74 |
| JMJD1C | 8.99E-79 | 0.658974 | 0.54 | 0.455 | 3.29E-74 |
| DUSP1 | 1.03E-78 | -0.98187 | 0.448 | 0.56 | 3.76E-74 |
| MDH2 | 1.19E-78 | -0.35415 | 0.068 | 0.203 | 4.35E-74 |
| PGRMC1 | 1.80E-78 | -0.28264 | 0.146 | 0.319 | 6.60E-74 |
| BUD31 | 1.82E-78 | -0.38935 | 0.085 | 0.229 | 6.65E-74 |
| IGSF10 | 2.22E-78 | 0.719146 | 0.17 | 0.079 | 8.13E-74 |
| SQSTM1 | 2.71E-78 | -0.51051 | 0.242 | 0.429 | 9.89E-74 |
| ANXA2 | 2.82E-78 | -0.48789 | 0.708 | 0.79 | 1.03E-73 |
| G0S2 | 3.96E-78 | -0.84385 | 0.027 | 0.132 | 1.45E-73 |

| | | | | | |
|---|---|---|---|---|---|
| AP000311. | 4.21E-78 | 0.518647 | 0.189 | 0.087 | 1.54E-73 |
| UGDH | 5.68E-78 | -0.65782 | 0.156 | 0.315 | 2.08E-73 |
| PNISR | 7.44E-78 | 0.587831 | 0.593 | 0.53 | 2.72E-73 |
| ST13 | 8.68E-78 | -0.49656 | 0.155 | 0.326 | 3.17E-73 |
| NR4A1 | 1.02E-77 | -0.54246 | 0.223 | 0.395 | 3.72E-73 |
| EFNA5 | 1.25E-77 | 0.673396 | 0.178 | 0.082 | 4.58E-73 |
| HSPB1 | 1.27E-77 | -0.6305 | 0.257 | 0.444 | 4.64E-73 |
| MAGED2 | 1.39E-77 | -0.34604 | 0.096 | 0.245 | 5.09E-73 |
| PTX3 | 1.55E-77 | -1.5658 | 0.022 | 0.121 | 5.65E-73 |
| GEM | 1.79E-77 | -0.65353 | 0.087 | 0.221 | 6.56E-73 |
| NDUFAB1 | 2.08E-77 | -0.39589 | 0.087 | 0.231 | 7.59E-73 |
| PRDX6 | 2.30E-77 | -0.56285 | 0.277 | 0.468 | 8.41E-73 |
| SENP5 | 2.35E-77 | 0.714795 | 0.246 | 0.137 | 8.58E-73 |
| HSPB2 | 2.47E-77 | -0.37698 | 0.038 | 0.152 | 9.03E-73 |
| TKT | 2.87E-77 | -0.32723 | 0.06 | 0.189 | 1.05E-72 |
| ANKRD12 | 3.09E-77 | 0.620283 | 0.552 | 0.467 | 1.13E-72 |
| UBR3 | 3.69E-77 | 0.6782 | 0.239 | 0.132 | 1.35E-72 |
| TRAPPC2L | 3.85E-77 | -0.39719 | 0.052 | 0.175 | 1.41E-72 |
| SET | 4.44E-77 | -0.41242 | 0.125 | 0.283 | 1.62E-72 |
| LINC-PINT | 4.86E-77 | 0.878581 | 0.396 | 0.294 | 1.77E-72 |
| ADAMTS9- | 5.81E-77 | 0.725288 | 0.152 | 0.066 | 2.12E-72 |
| MAP4K3 | 5.89E-77 | 0.739558 | 0.204 | 0.105 | 2.15E-72 |
| ALKBH7 | 6.08E-77 | -0.41345 | 0.034 | 0.146 | 2.22E-72 |
| MBNL1 | 7.13E-77 | 0.684033 | 0.484 | 0.391 | 2.61E-72 |
| TIMM13 | 1.43E-76 | -0.52083 | 0.069 | 0.2 | 5.24E-72 |
| RAC1 | 1.44E-76 | -0.36606 | 0.276 | 0.474 | 5.25E-72 |
| CRIP1 | 1.53E-76 | -0.59723 | 0.296 | 0.469 | 5.57E-72 |
| DYNC2H1 | 1.68E-76 | 0.724996 | 0.25 | 0.142 | 6.16E-72 |
| RPS17 | 1.75E-76 | -0.81721 | 0.505 | 0.698 | 6.38E-72 |
| VAPA | 2.51E-76 | -0.25174 | 0.16 | 0.337 | 9.17E-72 |
| ADK | 2.97E-76 | 0.588888 | 0.29 | 0.171 | 1.08E-71 |
| PLPP3 | 3.70E-76 | 0.523194 | 0.657 | 0.571 | 1.35E-71 |
| SLC25A3 | 3.85E-76 | -0.50085 | 0.26 | 0.454 | 1.41E-71 |
| DEGS1 | 4.81E-76 | -0.25283 | 0.08 | 0.22 | 1.76E-71 |
| TAX1BP3 | 5.68E-76 | -0.41192 | 0.07 | 0.201 | 2.08E-71 |
| YWHAH | 5.87E-76 | -0.35927 | 0.074 | 0.207 | 2.15E-71 |
| ISM1 | 7.59E-76 | 0.766936 | 0.186 | 0.092 | 2.77E-71 |
| GPM6A | 8.90E-76 | 0.486781 | 0.171 | 0.076 | 3.25E-71 |
| ARL4D | 9.56E-76 | -0.52314 | 0.015 | 0.11 | 3.50E-71 |
| LHFPL6 | 1.07E-75 | 0.625862 | 0.637 | 0.57 | 3.90E-71 |
| RANBP1 | 1.27E-75 | -0.25858 | 0.078 | 0.216 | 4.66E-71 |
| GSTM3 | 1.63E-75 | -0.41191 | 0.084 | 0.224 | 5.97E-71 |
| COL4A3BP | 1.93E-75 | 0.50843 | 0.156 | 0.067 | 7.06E-71 |
| PPA1 | 1.98E-75 | -0.41511 | 0.135 | 0.297 | 7.24E-71 |
| RNASEH2C | 2.17E-75 | -0.361 | 0.085 | 0.226 | 7.92E-71 |
| CCDC88A | 2.22E-75 | 0.769089 | 0.296 | 0.185 | 8.13E-71 |
| LMAN2 | 2.41E-75 | -0.31032 | 0.094 | 0.24 | 8.81E-71 |

| | | | | | |
|---|---|---|---|---|---|
| PLCB1 | 2.46E-75 | 0.658999 | 0.246 | 0.134 | 8.98E-71 |
| HSP90B1 | 2.85E-75 | 0.481011 | 0.656 | 0.603 | 1.04E-70 |
| PTK2 | 2.86E-75 | 0.684978 | 0.282 | 0.171 | 1.04E-70 |
| MAP3K20 | 3.08E-75 | 0.824651 | 0.374 | 0.271 | 1.12E-70 |
| TMEM147 | 3.25E-75 | -0.3578 | 0.045 | 0.161 | 1.19E-70 |
| TMEM14B | 5.47E-75 | -0.33374 | 0.077 | 0.214 | 2.00E-70 |
| ENOSF1 | 5.60E-75 | 0.650924 | 0.195 | 0.098 | 2.05E-70 |
| NBEAL1 | 6.12E-75 | 0.552089 | 0.362 | 0.238 | 2.24E-70 |
| DIP2B | 7.12E-75 | 0.59465 | 0.143 | 0.06 | 2.60E-70 |
| ZFY | 7.44E-75 | 0.682315 | 0.172 | 0.08 | 2.72E-70 |
| POLR1D | 7.50E-75 | -0.33494 | 0.096 | 0.241 | 2.74E-70 |
| PSMG2 | 9.68E-75 | -0.37938 | 0.112 | 0.265 | 3.54E-70 |
| SRSF7 | 1.03E-74 | -0.48739 | 0.203 | 0.382 | 3.76E-70 |
| TMEM258 | 1.20E-74 | -0.50698 | 0.172 | 0.348 | 4.39E-70 |
| RGCC | 1.29E-74 | -0.79239 | 0.099 | 0.232 | 4.71E-70 |
| UBB | 1.34E-74 | -0.56878 | 0.446 | 0.625 | 4.88E-70 |
| TNFRSF1A | 1.73E-74 | -0.26643 | 0.133 | 0.294 | 6.32E-70 |
| NDUFA12 | 2.15E-74 | -0.37434 | 0.077 | 0.213 | 7.86E-70 |
| ACSS3 | 2.83E-74 | 0.656595 | 0.136 | 0.056 | 1.03E-69 |
| HSBP1 | 3.17E-74 | -0.39728 | 0.148 | 0.316 | 1.16E-69 |
| NKTR | 4.37E-74 | 0.764968 | 0.44 | 0.351 | 1.60E-69 |
| LCORL | 5.57E-74 | 0.65643 | 0.156 | 0.069 | 2.04E-69 |
| CTSD | 5.58E-74 | -0.4639 | 0.162 | 0.329 | 2.04E-69 |
| ALDH2 | 5.78E-74 | -0.32738 | 0.123 | 0.277 | 2.11E-69 |
| TMEM179 | 5.89E-74 | -0.36318 | 0.057 | 0.179 | 2.15E-69 |
| SERP1 | 6.66E-74 | -0.2779 | 0.127 | 0.286 | 2.43E-69 |
| MDK | 7.17E-74 | -0.39423 | 0.025 | 0.127 | 2.62E-69 |
| MYCBP2 | 9.71E-74 | 0.669916 | 0.379 | 0.267 | 3.55E-69 |
| LDLRAD4 | 1.13E-73 | 0.586167 | 0.152 | 0.066 | 4.14E-69 |
| SKIV2L2 | 1.22E-73 | 0.463553 | 0.157 | 0.068 | 4.44E-69 |
| ULK4 | 1.67E-73 | 0.547969 | 0.15 | 0.064 | 6.11E-69 |
| TMEM50A | 2.13E-73 | -0.40195 | 0.202 | 0.385 | 7.77E-69 |
| ZFHX4 | 2.27E-73 | 0.750624 | 0.294 | 0.187 | 8.31E-69 |
| XBP1 | 2.39E-73 | -0.33337 | 0.128 | 0.284 | 8.75E-69 |
| HSPB6 | 2.56E-73 | -0.3966 | 0.131 | 0.285 | 9.35E-69 |
| SASH1 | 2.80E-73 | 0.742751 | 0.378 | 0.274 | 1.02E-68 |
| POLE4 | 5.27E-73 | -0.35429 | 0.034 | 0.141 | 1.93E-68 |
| PSMB5 | 7.57E-73 | -0.35729 | 0.093 | 0.234 | 2.77E-68 |
| POLR2K | 8.15E-73 | -0.30609 | 0.057 | 0.179 | 2.98E-68 |
| IAH1 | 9.55E-73 | -0.31439 | 0.071 | 0.201 | 3.49E-68 |
| LRBA | 1.35E-72 | 0.597797 | 0.176 | 0.083 | 4.94E-68 |
| PCNX2 | 1.93E-72 | 0.654842 | 0.19 | 0.095 | 7.05E-68 |
| AC023469 | 2.55E-72 | 0.458446 | 0.175 | 0.081 | 9.32E-68 |
| IER5L | 2.61E-72 | -0.37868 | 0.012 | 0.1 | 9.54E-68 |
| CARHSP1 | 2.75E-72 | -0.48095 | 0.095 | 0.234 | 1.00E-67 |
| HCFC1R1 | 2.94E-72 | -0.52771 | 0.072 | 0.2 | 1.07E-67 |
| SF1 | 3.12E-72 | -0.35216 | 0.152 | 0.318 | 1.14E-67 |

| | | | | | |
|---|---|---|---|---|---|
| NIPBL | 3.52E-72 | 0.679691 | 0.373 | 0.262 | 1.29E-67 |
| HDAC8 | 3.95E-72 | 0.756706 | 0.187 | 0.094 | 1.44E-67 |
| EDNRB | 4.39E-72 | -0.36639 | 0.052 | 0.168 | 1.60E-67 |
| GADD45A | 4.67E-72 | -0.47913 | 0.077 | 0.205 | 1.71E-67 |
| SPTSSA | 4.88E-72 | -0.36904 | 0.042 | 0.154 | 1.78E-67 |
| VDAC1 | 4.92E-72 | -0.35282 | 0.082 | 0.215 | 1.80E-67 |
| COX8A | 5.41E-72 | -0.49871 | 0.15 | 0.313 | 1.98E-67 |
| NOP10 | 5.89E-72 | -0.45636 | 0.082 | 0.217 | 2.15E-67 |
| ARID5A | 6.06E-72 | -0.36387 | 0.02 | 0.114 | 2.21E-67 |
| AC007319. | 1.04E-71 | 0.65376 | 0.181 | 0.086 | 3.81E-67 |
| HNRNPA0 | 1.05E-71 | -0.34339 | 0.164 | 0.335 | 3.83E-67 |
| C12orf57 | 1.27E-71 | -0.55646 | 0.267 | 0.458 | 4.64E-67 |
| PIN1 | 1.45E-71 | -0.36875 | 0.046 | 0.16 | 5.31E-67 |
| ITIH5 | 2.03E-71 | 0.66099 | 0.475 | 0.395 | 7.43E-67 |
| IER5 | 3.59E-71 | -0.36065 | 0.062 | 0.183 | 1.31E-66 |
| FAM120B | 3.74E-71 | 0.531974 | 0.157 | 0.071 | 1.37E-66 |
| MAP1LC3E | 4.13E-71 | -0.29506 | 0.116 | 0.267 | 1.51E-66 |
| PRR16 | 4.21E-71 | 0.662235 | 0.141 | 0.059 | 1.54E-66 |
| HECTD4 | 4.29E-71 | 0.652307 | 0.2 | 0.103 | 1.57E-66 |
| ZHX3 | 4.33E-71 | 0.742515 | 0.233 | 0.13 | 1.58E-66 |
| YIPF3 | 4.74E-71 | -0.34914 | 0.062 | 0.184 | 1.73E-66 |
| KCTD3 | 5.87E-71 | 0.733771 | 0.217 | 0.12 | 2.15E-66 |
| RAB5C | 6.71E-71 | -0.3669 | 0.066 | 0.19 | 2.45E-66 |
| WDR60 | 7.33E-71 | 0.738421 | 0.253 | 0.151 | 2.68E-66 |
| GRASP | 1.00E-70 | -0.32177 | 0.026 | 0.125 | 3.66E-66 |
| HMGN2 | 1.78E-70 | -0.42909 | 0.192 | 0.37 | 6.52E-66 |
| SPTBN1 | 1.86E-70 | 0.518109 | 0.599 | 0.545 | 6.81E-66 |
| SON | 2.06E-70 | 0.509936 | 0.592 | 0.545 | 7.53E-66 |
| CIB1 | 2.27E-70 | -0.42518 | 0.078 | 0.207 | 8.29E-66 |
| SCAND1 | 2.33E-70 | -0.28253 | 0.082 | 0.214 | 8.53E-66 |
| DNAJC19 | 4.06E-70 | -0.34969 | 0.059 | 0.178 | 1.49E-65 |
| BTG3 | 4.07E-70 | -0.27862 | 0.102 | 0.244 | 1.49E-65 |
| UACA | 4.28E-70 | 0.747111 | 0.348 | 0.243 | 1.56E-65 |
| MRPL40 | 5.14E-70 | -0.37259 | 0.04 | 0.148 | 1.88E-65 |
| MAPK8 | 5.20E-70 | 0.76586 | 0.209 | 0.113 | 1.90E-65 |
| CSRP1 | 5.46E-70 | -0.32776 | 0.066 | 0.189 | 2.00E-65 |
| BORCS7 | 5.77E-70 | -0.41263 | 0.069 | 0.194 | 2.11E-65 |
| NR2F2-AS1 | 7.44E-70 | 0.798756 | 0.145 | 0.065 | 2.72E-65 |
| COX7B | 7.57E-70 | -0.48179 | 0.125 | 0.275 | 2.77E-65 |
| CAMKMT | 8.07E-70 | 0.599787 | 0.135 | 0.057 | 2.95E-65 |
| ATF4 | 8.07E-70 | -0.53098 | 0.229 | 0.41 | 2.95E-65 |
| KANTR | 8.38E-70 | 0.50545 | 0.118 | 0.046 | 3.06E-65 |
| VEGFA | 8.67E-70 | -0.4009 | 0.055 | 0.169 | 3.17E-65 |
| ARF1 | 1.19E-69 | -0.36647 | 0.203 | 0.384 | 4.36E-65 |
| PITPNC1 | 1.58E-69 | 0.70391 | 0.224 | 0.124 | 5.76E-65 |
| CRABP2 | 1.59E-69 | -0.52292 | 0.121 | 0.263 | 5.81E-65 |
| LAMTOR2 | 1.87E-69 | -0.40413 | 0.046 | 0.157 | 6.83E-65 |

| | | | | | |
|---|---|---|---|---|---|
| ARL6IP1 | 2.05E-69 | -0.35098 | 0.098 | 0.237 | 7.51E-65 |
| GLUL | 2.27E-69 | -0.71769 | 0.397 | 0.554 | 8.30E-65 |
| VASN | 2.83E-69 | -0.36328 | 0.08 | 0.207 | 1.03E-64 |
| UGGT2 | 2.83E-69 | 0.710326 | 0.308 | 0.199 | 1.04E-64 |
| NDUFS7 | 3.26E-69 | -0.407 | 0.067 | 0.191 | 1.19E-64 |
| STXBP5 | 3.84E-69 | 0.672642 | 0.168 | 0.08 | 1.40E-64 |
| C12orf10 | 4.36E-69 | -0.32377 | 0.017 | 0.107 | 1.59E-64 |
| CCT5 | 5.01E-69 | -0.28065 | 0.063 | 0.183 | 1.83E-64 |
| F10 | 6.11E-69 | -0.37396 | 0.146 | 0.303 | 2.23E-64 |
| AC006453. | 6.33E-69 | 0.524267 | 0.153 | 0.069 | 2.32E-64 |
| HAX1 | 6.36E-69 | -0.31985 | 0.036 | 0.14 | 2.32E-64 |
| MRPL34 | 6.38E-69 | -0.38961 | 0.034 | 0.136 | 2.33E-64 |
| PRELP | 7.29E-69 | -0.4736 | 0.223 | 0.397 | 2.66E-64 |
| FBXL17 | 8.47E-69 | 0.689831 | 0.186 | 0.094 | 3.10E-64 |
| CCDC146 | 8.98E-69 | 0.683477 | 0.213 | 0.116 | 3.28E-64 |
| MANF | 1.81E-68 | -0.26096 | 0.05 | 0.163 | 6.63E-64 |
| LDHB | 2.58E-68 | -0.35552 | 0.113 | 0.258 | 9.45E-64 |
| COX14 | 2.80E-68 | -0.40408 | 0.053 | 0.167 | 1.02E-63 |
| AC021351. | 2.89E-68 | 0.36973 | 0.105 | 0.038 | 1.06E-63 |
| BTG2 | 3.22E-68 | -0.61841 | 0.236 | 0.4 | 1.18E-63 |
| DOPEY1 | 5.42E-68 | 0.427625 | 0.111 | 0.041 | 1.98E-63 |
| ZNF593 | 5.87E-68 | -0.33316 | 0.033 | 0.135 | 2.15E-63 |
| ACVR2A | 6.65E-68 | 0.665662 | 0.145 | 0.066 | 2.43E-63 |
| PHACTR2 | 7.28E-68 | 0.669263 | 0.297 | 0.19 | 2.66E-63 |
| VPS54 | 7.32E-68 | 0.620707 | 0.176 | 0.086 | 2.67E-63 |
| NDUFS6 | 7.42E-68 | -0.4285 | 0.097 | 0.233 | 2.71E-63 |
| MLF2 | 8.10E-68 | -0.38009 | 0.059 | 0.176 | 2.96E-63 |
| ARPC3 | 1.15E-67 | -0.41602 | 0.175 | 0.344 | 4.21E-63 |
| NCALD | 1.48E-67 | 0.680286 | 0.256 | 0.155 | 5.42E-63 |
| EIF4A3 | 1.88E-67 | -0.382 | 0.093 | 0.224 | 6.86E-63 |
| DOCK9 | 1.94E-67 | 0.586511 | 0.242 | 0.137 | 7.08E-63 |
| CDON | 2.06E-67 | 0.693415 | 0.223 | 0.127 | 7.52E-63 |
| C12orf60 | 2.29E-67 | 0.443212 | 0.154 | 0.069 | 8.38E-63 |
| MAFB | 2.29E-67 | -0.66561 | 0.111 | 0.245 | 8.39E-63 |
| CAMK1D | 2.80E-67 | 0.621758 | 0.308 | 0.193 | 1.02E-62 |
| GABARAPL | 2.85E-67 | -0.38369 | 0.112 | 0.251 | 1.04E-62 |
| EIF6 | 3.03E-67 | -0.3446 | 0.055 | 0.17 | 1.11E-62 |
| ARL6IP4 | 3.36E-67 | -0.34174 | 0.174 | 0.342 | 1.23E-62 |
| MYOF | 4.01E-67 | 0.727538 | 0.343 | 0.242 | 1.46E-62 |
| ARF4 | 7.39E-67 | -0.39488 | 0.174 | 0.34 | 2.70E-62 |
| AFF3 | 7.88E-67 | 0.5803 | 0.128 | 0.054 | 2.88E-62 |
| STK38 | 8.70E-67 | 0.661348 | 0.22 | 0.123 | 3.18E-62 |
| PTPRM | 9.24E-67 | 0.582392 | 0.236 | 0.134 | 3.38E-62 |
| PRKCE | 9.56E-67 | 0.548971 | 0.148 | 0.067 | 3.49E-62 |
| MFGE8 | 1.17E-66 | -0.35776 | 0.183 | 0.35 | 4.28E-62 |
| ATP6V1G1 | 1.90E-66 | -0.33398 | 0.146 | 0.305 | 6.93E-62 |
| SNRPG | 1.92E-66 | -0.4523 | 0.077 | 0.201 | 7.02E-62 |

| | | | | | |
|---|---|---|---|---|---|
| LINC00558 | 2.30E-66 | 0.464002 | 0.176 | 0.084 | 8.42E-62 |
| PSMA1 | 2.71E-66 | -0.3568 | 0.126 | 0.274 | 9.91E-62 |
| LSM7 | 2.82E-66 | -0.40112 | 0.084 | 0.212 | 1.03E-61 |
| POLR2E | 3.65E-66 | -0.35305 | 0.079 | 0.205 | 1.34E-61 |
| RWDD1 | 3.93E-66 | -0.46145 | 0.153 | 0.311 | 1.44E-61 |
| MAP2K5 | 5.20E-66 | 0.595367 | 0.16 | 0.076 | 1.90E-61 |
| AHSA1 | 5.20E-66 | -0.25135 | 0.048 | 0.157 | 1.90E-61 |
| CAST | 5.49E-66 | 0.483802 | 0.605 | 0.552 | 2.01E-61 |
| MGST1 | 6.27E-66 | -0.54899 | 0.457 | 0.604 | 2.29E-61 |
| KANSL1 | 6.54E-66 | 0.710552 | 0.229 | 0.135 | 2.39E-61 |
| HERPUD1 | 6.55E-66 | -0.40719 | 0.108 | 0.247 | 2.39E-61 |
| PKM | 7.95E-66 | -0.44082 | 0.122 | 0.264 | 2.90E-61 |
| EXTL3 | 9.39E-66 | 0.461171 | 0.177 | 0.086 | 3.43E-61 |
| ARPC1A | 9.92E-66 | -0.33385 | 0.042 | 0.147 | 3.63E-61 |
| FOXO1 | 1.00E-65 | 0.696868 | 0.339 | 0.236 | 3.67E-61 |
| PRDX4 | 1.07E-65 | -0.30305 | 0.149 | 0.307 | 3.93E-61 |
| WDFY3 | 1.24E-65 | 0.648754 | 0.238 | 0.137 | 4.55E-61 |
| PSMB8 | 1.39E-65 | -0.30651 | 0.043 | 0.147 | 5.09E-61 |
| SOCS1 | 1.51E-65 | -0.27709 | 0.016 | 0.101 | 5.51E-61 |
| ESD | 1.90E-65 | -0.40202 | 0.169 | 0.334 | 6.95E-61 |
| ENY2 | 1.92E-65 | -0.33814 | 0.156 | 0.317 | 7.04E-61 |
| GRHPR | 1.95E-65 | -0.29151 | 0.062 | 0.179 | 7.12E-61 |
| CCDC107 | 2.19E-65 | -0.32738 | 0.064 | 0.181 | 8.00E-61 |
| SNRPN | 2.38E-65 | -0.26519 | 0.049 | 0.159 | 8.69E-61 |
| ADH5 | 3.12E-65 | -0.34453 | 0.135 | 0.284 | 1.14E-60 |
| COX5A | 3.48E-65 | -0.27705 | 0.06 | 0.175 | 1.27E-60 |
| TPM2 | 3.63E-65 | -0.42845 | 0.124 | 0.265 | 1.33E-60 |
| CNTN3 | 4.37E-65 | 0.59285 | 0.113 | 0.045 | 1.60E-60 |
| 10-Sep | 9.64E-65 | 0.408198 | 0.177 | 0.086 | 3.53E-60 |
| ARPC1B | 1.06E-64 | -0.41995 | 0.103 | 0.237 | 3.86E-60 |
| TIAM2 | 1.06E-64 | 0.458588 | 0.119 | 0.048 | 3.88E-60 |
| CIRBP | 1.18E-64 | -0.56316 | 0.359 | 0.539 | 4.30E-60 |
| C16orf52 | 1.47E-64 | 0.420774 | 0.122 | 0.049 | 5.39E-60 |
| UBE2L6 | 1.54E-64 | -0.39404 | 0.043 | 0.146 | 5.61E-60 |
| CNIH4 | 1.55E-64 | -0.30756 | 0.077 | 0.201 | 5.68E-60 |
| TRIO | 2.41E-64 | 0.365327 | 0.409 | 0.275 | 8.82E-60 |
| TMEM219 | 3.31E-64 | -0.2796 | 0.096 | 0.229 | 1.21E-59 |
| EMC7 | 3.33E-64 | -0.30854 | 0.096 | 0.23 | 1.22E-59 |
| EIF4G3 | 3.40E-64 | 0.682244 | 0.282 | 0.181 | 1.24E-59 |
| NDUFV2 | 3.93E-64 | -0.28345 | 0.115 | 0.258 | 1.44E-59 |
| ENO1 | 4.20E-64 | -0.40957 | 0.209 | 0.38 | 1.53E-59 |
| MIR222HG | 5.78E-64 | -0.42493 | 0.031 | 0.124 | 2.11E-59 |
| TMEM70 | 8.38E-64 | -0.31295 | 0.039 | 0.139 | 3.06E-59 |
| CNTNAP2 | 1.02E-63 | 0.440608 | 0.154 | 0.071 | 3.71E-59 |
| SAMD4A | 1.52E-63 | 0.700419 | 0.321 | 0.216 | 5.56E-59 |
| SGK1 | 1.56E-63 | -0.48712 | 0.073 | 0.187 | 5.71E-59 |
| AC006059.1 | 1.57E-63 | 0.453791 | 0.141 | 0.062 | 5.72E-59 |

| | | | | | |
|---|---|---|---|---|---|
| MARK1 | 1.64E-63 | 0.653784 | 0.125 | 0.054 | 6.01E-59 |
| PSMB2 | 1.78E-63 | -0.26842 | 0.079 | 0.202 | 6.49E-59 |
| FAM102B | 1.78E-63 | 0.694597 | 0.158 | 0.078 | 6.52E-59 |
| PICALM | 2.11E-63 | 0.624633 | 0.27 | 0.17 | 7.72E-59 |
| FBXO42 | 2.42E-63 | 0.635541 | 0.156 | 0.076 | 8.85E-59 |
| ID3 | 2.61E-63 | -0.8334 | 0.087 | 0.204 | 9.55E-59 |
| COTL1 | 2.69E-63 | -0.25574 | 0.075 | 0.195 | 9.83E-59 |
| ICAM1 | 2.95E-63 | -0.75065 | 0.067 | 0.176 | 1.08E-58 |
| PDCD5 | 3.11E-63 | -0.39029 | 0.131 | 0.277 | 1.14E-58 |
| RGS2 | 3.20E-63 | -0.63958 | 0.065 | 0.175 | 1.17E-58 |
| FILIP1 | 3.65E-63 | 0.786219 | 0.32 | 0.221 | 1.33E-58 |
| CHST11 | 3.88E-63 | 0.488672 | 0.176 | 0.088 | 1.42E-58 |
| REEP5 | 4.46E-63 | -0.29709 | 0.209 | 0.384 | 1.63E-58 |
| NTAN1 | 4.83E-63 | -0.25535 | 0.069 | 0.187 | 1.77E-58 |
| NDUFB2 | 4.95E-63 | -0.46169 | 0.179 | 0.342 | 1.81E-58 |
| MRPL54 | 5.04E-63 | -0.30837 | 0.047 | 0.151 | 1.84E-58 |
| STK3 | 5.12E-63 | 0.671539 | 0.334 | 0.231 | 1.87E-58 |
| TRAF3IP2-/ | 8.18E-63 | 0.678119 | 0.176 | 0.09 | 2.99E-58 |
| HSPG2 | 8.87E-63 | 0.576042 | 0.47 | 0.396 | 3.24E-58 |
| WIPF1 | 9.01E-63 | 0.702562 | 0.256 | 0.159 | 3.29E-58 |
| ATAD2B | 1.22E-62 | 0.620972 | 0.184 | 0.096 | 4.46E-58 |
| HSPH1 | 1.38E-62 | -0.74509 | 0.073 | 0.186 | 5.04E-58 |
| PRKD1 | 1.48E-62 | 0.576831 | 0.144 | 0.066 | 5.40E-58 |
| EMP2 | 1.77E-62 | -0.38665 | 0.241 | 0.414 | 6.47E-58 |
| DENND1A | 2.43E-62 | 0.548137 | 0.142 | 0.065 | 8.88E-58 |
| PCDH7 | 3.32E-62 | 0.63681 | 0.144 | 0.067 | 1.21E-57 |
| RAB13 | 3.48E-62 | -0.55214 | 0.053 | 0.157 | 1.27E-57 |
| PLIN2 | 3.74E-62 | -0.62146 | 0.21 | 0.362 | 1.37E-57 |
| NDUFS8 | 4.29E-62 | -0.38104 | 0.05 | 0.154 | 1.57E-57 |
| AGTR1 | 4.40E-62 | 0.941482 | 0.313 | 0.227 | 1.61E-57 |
| JAGN1 | 7.60E-62 | -0.30848 | 0.029 | 0.121 | 2.78E-57 |
| ATP10A | 1.02E-61 | 0.548351 | 0.152 | 0.072 | 3.73E-57 |
| JOSD2 | 1.08E-61 | -0.37362 | 0.031 | 0.123 | 3.95E-57 |
| ADAMTS1: | 1.18E-61 | 0.634584 | 0.145 | 0.069 | 4.32E-57 |
| HLA-E | 1.22E-61 | -0.5228 | 0.422 | 0.585 | 4.47E-57 |
| PSMB9 | 2.09E-61 | -0.35092 | 0.033 | 0.127 | 7.64E-57 |
| TIMM23B | 2.15E-61 | 0.638354 | 0.167 | 0.083 | 7.85E-57 |
| ATF3 | 2.51E-61 | -0.62626 | 0.233 | 0.385 | 9.18E-57 |
| GADD45GI | 3.90E-61 | -0.25218 | 0.111 | 0.247 | 1.43E-56 |
| CDC14B | 3.93E-61 | 0.614119 | 0.171 | 0.088 | 1.44E-56 |
| UBE2I | 4.39E-61 | -0.29977 | 0.122 | 0.263 | 1.60E-56 |
| ZNHIT1 | 5.04E-61 | -0.38818 | 0.133 | 0.276 | 1.84E-56 |
| NIFK | 6.52E-61 | -0.33091 | 0.048 | 0.15 | 2.38E-56 |
| CYP7B1 | 7.00E-61 | 0.842647 | 0.248 | 0.157 | 2.56E-56 |
| POLR2L | 7.97E-61 | -0.59469 | 0.26 | 0.434 | 2.91E-56 |
| PPIC | 8.37E-61 | -0.46846 | 0.143 | 0.285 | 3.06E-56 |
| MYL6B | 1.02E-60 | -0.32086 | 0.047 | 0.149 | 3.74E-56 |

| | | | | | |
|---|---|---|---|---|---|
| NOP16 | 1.06E-60 | -0.27466 | 0.027 | 0.115 | 3.86E-56 |
| TUBA1A | 1.06E-60 | -0.6611 | 0.335 | 0.49 | 3.88E-56 |
| TMED2 | 1.17E-60 | -0.30912 | 0.136 | 0.282 | 4.28E-56 |
| PDPN | 1.36E-60 | -0.50522 | 0.051 | 0.151 | 4.97E-56 |
| TALDO1 | 1.69E-60 | -0.32044 | 0.086 | 0.208 | 6.17E-56 |
| ADAMTS4 | 2.20E-60 | -0.55646 | 0.022 | 0.105 | 8.05E-56 |
| NBPF15 | 3.32E-60 | 0.477246 | 0.125 | 0.054 | 1.21E-55 |
| XRN1 | 4.30E-60 | 0.709228 | 0.282 | 0.184 | 1.57E-55 |
| ARHGDIA | 4.52E-60 | -0.25801 | 0.113 | 0.25 | 1.65E-55 |
| FIBP | 5.32E-60 | -0.30968 | 0.025 | 0.111 | 1.95E-55 |
| XPR1 | 5.53E-60 | 0.575078 | 0.147 | 0.07 | 2.02E-55 |
| WDPCP | 5.54E-60 | 0.602337 | 0.191 | 0.102 | 2.03E-55 |
| HIPK3 | 6.31E-60 | 0.771594 | 0.371 | 0.29 | 2.31E-55 |
| RRBP1 | 1.14E-59 | 0.571746 | 0.543 | 0.49 | 4.18E-55 |
| SEC24B | 1.17E-59 | 0.561727 | 0.166 | 0.084 | 4.26E-55 |
| ATG14 | 1.51E-59 | 0.568062 | 0.141 | 0.066 | 5.54E-55 |
| XYLT1 | 1.72E-59 | 0.577388 | 0.217 | 0.123 | 6.27E-55 |
| IL1RAPL1 | 2.19E-59 | 0.373654 | 0.101 | 0.039 | 8.02E-55 |
| MRPL11 | 2.73E-59 | -0.27653 | 0.04 | 0.136 | 9.98E-55 |
| PCBP1 | 2.99E-59 | -0.32046 | 0.15 | 0.299 | 1.09E-54 |
| CNN3 | 3.06E-59 | -0.35619 | 0.171 | 0.326 | 1.12E-54 |
| PIGP | 3.30E-59 | -0.30628 | 0.026 | 0.112 | 1.21E-54 |
| GRN | 3.56E-59 | -0.29417 | 0.136 | 0.278 | 1.30E-54 |
| IDH3G | 3.85E-59 | -0.25507 | 0.046 | 0.146 | 1.41E-54 |
| VPS29 | 4.46E-59 | -0.29695 | 0.118 | 0.254 | 1.63E-54 |
| TPRG1 | 4.50E-59 | 0.634479 | 0.115 | 0.048 | 1.64E-54 |
| TRERF1 | 4.85E-59 | 0.548358 | 0.154 | 0.076 | 1.77E-54 |
| FOCAD | 6.12E-59 | 0.494954 | 0.112 | 0.047 | 2.24E-54 |
| POU2F1 | 7.01E-59 | 0.659658 | 0.176 | 0.093 | 2.56E-54 |
| SCFD2 | 7.60E-59 | 0.540846 | 0.151 | 0.073 | 2.78E-54 |
| DIP2C | 8.25E-59 | 0.515282 | 0.168 | 0.086 | 3.01E-54 |
| IMPDH2 | 8.86E-59 | -0.36652 | 0.078 | 0.194 | 3.24E-54 |
| UQCR10 | 9.11E-59 | -0.44447 | 0.108 | 0.237 | 3.33E-54 |
| POLR2I | 9.32E-59 | -0.38193 | 0.052 | 0.154 | 3.41E-54 |
| AP001347. | 1.01E-58 | 0.598288 | 0.123 | 0.054 | 3.68E-54 |
| SMOC2 | 1.05E-58 | 0.858903 | 0.333 | 0.248 | 3.85E-54 |
| MRPL14 | 1.09E-58 | -0.25905 | 0.041 | 0.136 | 3.98E-54 |
| SETD5 | 1.16E-58 | 0.647574 | 0.289 | 0.193 | 4.23E-54 |
| DPH6 | 1.39E-58 | 0.739575 | 0.153 | 0.076 | 5.07E-54 |
| RAD50 | 1.55E-58 | -0.26749 | 0.026 | 0.111 | 5.68E-54 |
| TNRC6B | 1.57E-58 | 0.632816 | 0.373 | 0.28 | 5.74E-54 |
| PEBP1 | 1.61E-58 | -0.49997 | 0.334 | 0.511 | 5.89E-54 |
| DDX3X | 1.92E-58 | -0.30956 | 0.221 | 0.389 | 7.02E-54 |
| MYL6 | 1.96E-58 | -0.47544 | 0.612 | 0.73 | 7.17E-54 |
| AES | 2.12E-58 | 0.316679 | 0.161 | 0.078 | 7.73E-54 |
| DYM | 2.12E-58 | 0.581832 | 0.195 | 0.108 | 7.74E-54 |
| EPS15 | 2.54E-58 | 0.713842 | 0.285 | 0.195 | 9.28E-54 |

| | | | | | |
|---|---|---|---|---|---|
| ATXN3 | 2.66E-58 | 0.700707 | 0.253 | 0.162 | 9.71E-54 |
| UQCR11 | 3.34E-58 | -0.51814 | 0.158 | 0.305 | 1.22E-53 |
| ST5 | 3.44E-58 | 0.713398 | 0.2 | 0.117 | 1.26E-53 |
| FGL2 | 3.62E-58 | 0.688559 | 0.323 | 0.229 | 1.32E-53 |
| UBE4B | 4.43E-58 | 0.575871 | 0.166 | 0.085 | 1.62E-53 |
| AGO3 | 5.33E-58 | 0.712159 | 0.278 | 0.184 | 1.95E-53 |
| EBF2 | 5.54E-58 | 0.699192 | 0.285 | 0.19 | 2.02E-53 |
| TANC1 | 5.93E-58 | 0.483143 | 0.1 | 0.04 | 2.17E-53 |
| GPATCH8 | 5.96E-58 | 0.691872 | 0.291 | 0.196 | 2.18E-53 |
| MVB12A | 7.92E-58 | -0.2721 | 0.039 | 0.132 | 2.89E-53 |
| FREM1 | 9.42E-58 | 0.506885 | 0.115 | 0.049 | 3.44E-53 |
| TMEM57 | 1.20E-57 | 0.413859 | 0.122 | 0.053 | 4.40E-53 |
| PNRC2 | 1.28E-57 | -0.34101 | 0.144 | 0.286 | 4.69E-53 |
| ROBO1 | 1.65E-57 | 0.644492 | 0.285 | 0.187 | 6.02E-53 |
| HSD3B7 | 1.73E-57 | -0.42691 | 0.027 | 0.111 | 6.34E-53 |
| CCT3 | 2.58E-57 | -0.30828 | 0.093 | 0.214 | 9.42E-53 |
| ZCCHC6 | 3.21E-57 | 0.402814 | 0.128 | 0.057 | 1.17E-52 |
| AF241726. | 4.48E-57 | 0.372951 | 0.115 | 0.048 | 1.64E-52 |
| SMIM19 | 4.59E-57 | -0.27142 | 0.086 | 0.204 | 1.68E-52 |
| HSP90AA1 | 5.18E-57 | -0.64624 | 0.601 | 0.717 | 1.89E-52 |
| TMEM256 | 5.40E-57 | -0.40586 | 0.047 | 0.144 | 1.97E-52 |
| NUDC | 5.46E-57 | -0.27608 | 0.126 | 0.262 | 1.99E-52 |
| LRCH3 | 6.03E-57 | 0.58874 | 0.202 | 0.115 | 2.20E-52 |
| ZFYVE21 | 6.10E-57 | -0.27485 | 0.045 | 0.142 | 2.23E-52 |
| MAGI1 | 8.28E-57 | 0.419272 | 0.101 | 0.04 | 3.03E-52 |
| UBR2 | 9.24E-57 | 0.618282 | 0.219 | 0.13 | 3.38E-52 |
| SPON2 | 9.55E-57 | -0.35081 | 0.174 | 0.321 | 3.49E-52 |
| MRPL27 | 2.08E-56 | -0.28523 | 0.051 | 0.151 | 7.60E-52 |
| EYS | 2.19E-56 | 0.332113 | 0.103 | 0.041 | 8.01E-52 |
| KIF13A | 2.26E-56 | 0.631434 | 0.216 | 0.127 | 8.26E-52 |
| SWI5 | 2.32E-56 | -0.27699 | 0.028 | 0.113 | 8.47E-52 |
| KLF10 | 2.88E-56 | -0.36736 | 0.095 | 0.211 | 1.05E-51 |
| YWHAQ | 5.00E-56 | -0.25198 | 0.223 | 0.395 | 1.83E-51 |
| TPM4 | 5.40E-56 | -0.449 | 0.194 | 0.34 | 1.97E-51 |
| MRPS15 | 5.80E-56 | -0.28844 | 0.048 | 0.146 | 2.12E-51 |
| HTRA3 | 6.12E-56 | -0.57731 | 0.164 | 0.298 | 2.24E-51 |
| CTSC | 6.25E-56 | -0.4184 | 0.031 | 0.116 | 2.29E-51 |
| TUBA1C | 7.85E-56 | -0.3234 | 0.079 | 0.189 | 2.87E-51 |
| CTSH | 1.03E-55 | -0.42436 | 0.045 | 0.138 | 3.76E-51 |
| IL6 | 1.23E-55 | -0.8062 | 0.057 | 0.152 | 4.51E-51 |
| FAM172A | 1.33E-55 | 0.612015 | 0.301 | 0.204 | 4.85E-51 |
| BAG3 | 1.45E-55 | -0.28502 | 0.086 | 0.2 | 5.30E-51 |
| OSBPL9 | 1.52E-55 | 0.706289 | 0.243 | 0.155 | 5.55E-51 |
| TMEM208 | 1.55E-55 | -0.25005 | 0.052 | 0.151 | 5.68E-51 |
| GRID2 | 1.67E-55 | 0.362118 | 0.127 | 0.057 | 6.10E-51 |
| ITSN1 | 1.95E-55 | 0.576915 | 0.26 | 0.166 | 7.13E-51 |
| DENND4C | 2.04E-55 | 0.606523 | 0.24 | 0.146 | 7.44E-51 |

| | | | | | |
|---|---|---|---|---|---|
| BHLHE40 | 2.68E-55 | -0.30539 | 0.055 | 0.152 | 9.81E-51 |
| CLEC2B | 2.79E-55 | -0.35603 | 0.15 | 0.289 | 1.02E-50 |
| ARMCX4 | 2.86E-55 | 0.558131 | 0.127 | 0.058 | 1.04E-50 |
| TEX264 | 3.13E-55 | -0.2768 | 0.031 | 0.116 | 1.15E-50 |
| SEC24D | 3.42E-55 | 0.727685 | 0.22 | 0.134 | 1.25E-50 |
| AVL9 | 3.55E-55 | 0.53861 | 0.128 | 0.059 | 1.30E-50 |
| TRAPPC1 | 3.72E-55 | -0.26719 | 0.052 | 0.15 | 1.36E-50 |
| IGF2 | 4.76E-55 | 0.497753 | 0.229 | 0.137 | 1.74E-50 |
| SLC20A1 | 5.07E-55 | -0.29359 | 0.06 | 0.16 | 1.85E-50 |
| PRG4 | 5.36E-55 | -2.50995 | 0.102 | 0.196 | 1.96E-50 |
| GRPEL1 | 8.54E-55 | -0.30252 | 0.026 | 0.107 | 3.12E-50 |
| NSMCE2 | 8.74E-55 | 0.682851 | 0.213 | 0.126 | 3.20E-50 |
| COA3 | 1.15E-54 | -0.30254 | 0.045 | 0.139 | 4.19E-50 |
| ARHGEF3 | 1.16E-54 | 0.453041 | 0.147 | 0.072 | 4.23E-50 |
| HIGD1A | 1.38E-54 | -0.29427 | 0.033 | 0.119 | 5.05E-50 |
| NHP2 | 1.40E-54 | -0.31291 | 0.085 | 0.199 | 5.11E-50 |
| AIDA | 1.73E-54 | -0.25855 | 0.04 | 0.129 | 6.32E-50 |
| IL32 | 1.81E-54 | -0.61187 | 0.042 | 0.13 | 6.61E-50 |
| SRSF3 | 1.83E-54 | -0.38793 | 0.235 | 0.399 | 6.68E-50 |
| PNKD | 1.85E-54 | -0.38854 | 0.06 | 0.16 | 6.75E-50 |
| TOMM22 | 2.28E-54 | -0.27674 | 0.044 | 0.136 | 8.33E-50 |
| RHOBTB3 | 2.64E-54 | 0.747148 | 0.448 | 0.384 | 9.66E-50 |
| ISOC2 | 3.12E-54 | -0.31216 | 0.023 | 0.102 | 1.14E-49 |
| BAX | 3.31E-54 | -0.2527 | 0.026 | 0.107 | 1.21E-49 |
| KMT2C | 3.42E-54 | 0.605656 | 0.297 | 0.203 | 1.25E-49 |
| TNRC6C | 3.48E-54 | 0.58678 | 0.205 | 0.12 | 1.27E-49 |
| PI16 | 3.96E-54 | -1.00135 | 0.153 | 0.265 | 1.45E-49 |
| CNBP | 6.78E-54 | -0.33586 | 0.267 | 0.444 | 2.48E-49 |
| WDR70 | 7.06E-54 | 0.520446 | 0.198 | 0.113 | 2.58E-49 |
| HPSE2 | 7.66E-54 | 0.527369 | 0.105 | 0.044 | 2.80E-49 |
| TBCB | 7.68E-54 | -0.32643 | 0.086 | 0.199 | 2.81E-49 |
| YPEL5 | 8.98E-54 | -0.25037 | 0.081 | 0.192 | 3.28E-49 |
| SUPT4H1 | 9.27E-54 | -0.35346 | 0.045 | 0.137 | 3.39E-49 |
| INSR | 9.29E-54 | 0.627492 | 0.248 | 0.159 | 3.40E-49 |
| PMF1 | 1.15E-53 | -0.26909 | 0.026 | 0.107 | 4.21E-49 |
| LSM2 | 1.32E-53 | -0.28466 | 0.038 | 0.126 | 4.82E-49 |
| FLYWCH2 | 1.56E-53 | -0.27544 | 0.033 | 0.119 | 5.71E-49 |
| ILK | 1.74E-53 | -0.26193 | 0.092 | 0.21 | 6.35E-49 |
| AC016708 | 2.35E-53 | 0.341223 | 0.289 | 0.176 | 8.59E-49 |
| APH1A | 2.40E-53 | -0.26111 | 0.058 | 0.158 | 8.76E-49 |
| SNRPE | 2.61E-53 | -0.42299 | 0.102 | 0.22 | 9.55E-49 |
| SCLT1 | 3.05E-53 | 0.666142 | 0.188 | 0.106 | 1.11E-48 |
| MRPS36 | 3.06E-53 | -0.2721 | 0.056 | 0.154 | 1.12E-48 |
| MRPL36 | 3.33E-53 | -0.30042 | 0.029 | 0.11 | 1.22E-48 |
| ARIH1 | 3.42E-53 | 0.66989 | 0.272 | 0.185 | 1.25E-48 |
| PDE8A | 3.63E-53 | 0.579429 | 0.204 | 0.119 | 1.33E-48 |
| LRP1B | 3.86E-53 | 0.522761 | 0.199 | 0.114 | 1.41E-48 |

| | | | | | |
|---|---|---|---|---|---|
| C16orf89 | 4.43E-53 | -0.29639 | 0.025 | 0.103 | 1.62E-48 |
| RBMS3-AS | 6.34E-53 | 0.441495 | 0.121 | 0.054 | 2.32E-48 |
| ARHGAP32 | 6.51E-53 | 0.671509 | 0.172 | 0.094 | 2.38E-48 |
| STOML2 | 6.62E-53 | -0.26734 | 0.052 | 0.147 | 2.42E-48 |
| LSM5 | 7.84E-53 | -0.31162 | 0.056 | 0.153 | 2.87E-48 |
| RPS27L | 8.10E-53 | -0.5048 | 0.238 | 0.398 | 2.96E-48 |
| GBF1 | 9.69E-53 | 0.593839 | 0.161 | 0.085 | 3.54E-48 |
| AKR1A1 | 1.11E-52 | -0.25872 | 0.086 | 0.2 | 4.05E-48 |
| PRKRIP1 | 1.16E-52 | 0.579642 | 0.176 | 0.097 | 4.24E-48 |
| SLC2A13 | 1.20E-52 | 0.570907 | 0.15 | 0.077 | 4.40E-48 |
| PMP22 | 1.71E-52 | -0.46484 | 0.393 | 0.557 | 6.25E-48 |
| NDUFA8 | 1.79E-52 | -0.26989 | 0.037 | 0.123 | 6.55E-48 |
| CTDSPL | 1.97E-52 | 0.785588 | 0.236 | 0.153 | 7.21E-48 |
| COL6A1 | 2.04E-52 | 0.31489 | 0.78 | 0.715 | 7.44E-48 |
| CCDC124 | 2.75E-52 | -0.29421 | 0.042 | 0.131 | 1.00E-47 |
| GNG11 | 3.59E-52 | -0.39923 | 0.242 | 0.405 | 1.31E-47 |
| FGD4 | 4.33E-52 | 0.594857 | 0.157 | 0.083 | 1.58E-47 |
| SIGIRR | 4.78E-52 | -0.25627 | 0.05 | 0.143 | 1.75E-47 |
| PIAS1 | 5.72E-52 | 0.617036 | 0.225 | 0.139 | 2.09E-47 |
| MARCKS | 7.07E-52 | 0.511263 | 0.531 | 0.47 | 2.58E-47 |
| OSBPL10 | 8.09E-52 | 0.556926 | 0.145 | 0.073 | 2.96E-47 |
| MAP7D3 | 9.27E-52 | 0.622251 | 0.292 | 0.205 | 3.39E-47 |
| NFX1 | 9.35E-52 | 0.506903 | 0.171 | 0.092 | 3.42E-47 |
| TAF15 | 1.33E-51 | 0.647384 | 0.233 | 0.147 | 4.88E-47 |
| TXN2 | 1.34E-51 | -0.32439 | 0.064 | 0.164 | 4.88E-47 |
| PDGFRL | 1.35E-51 | -0.46128 | 0.279 | 0.429 | 4.94E-47 |
| USP2-AS1 | 1.70E-51 | 0.407037 | 0.125 | 0.058 | 6.23E-47 |
| FAM162A | 1.79E-51 | -0.2873 | 0.061 | 0.16 | 6.54E-47 |
| AK1 | 2.12E-51 | -0.33074 | 0.031 | 0.112 | 7.75E-47 |
| TMEM135 | 2.37E-51 | 0.599485 | 0.153 | 0.08 | 8.68E-47 |
| SSBP1 | 2.76E-51 | -0.30777 | 0.144 | 0.279 | 1.01E-46 |
| REPS1 | 2.90E-51 | 0.534636 | 0.131 | 0.064 | 1.06E-46 |
| HMCN1 | 3.31E-51 | 0.652982 | 0.205 | 0.122 | 1.21E-46 |
| KIAA1328 | 3.49E-51 | 0.641741 | 0.164 | 0.088 | 1.28E-46 |
| PSMC5 | 4.33E-51 | -0.27377 | 0.131 | 0.261 | 1.58E-46 |
| ANXA1 | 5.45E-51 | -0.50315 | 0.733 | 0.758 | 1.99E-46 |
| ITM2B | 5.45E-51 | -0.36492 | 0.685 | 0.712 | 1.99E-46 |
| NDUFA6 | 5.62E-51 | -0.29295 | 0.083 | 0.191 | 2.06E-46 |
| NANS | 7.01E-51 | -0.28687 | 0.044 | 0.132 | 2.56E-46 |
| C10orf76 | 8.46E-51 | 0.394761 | 0.102 | 0.043 | 3.09E-46 |
| SLPI | 9.02E-51 | -1.31005 | 0.034 | 0.111 | 3.30E-46 |
| TMEM106 | 1.02E-50 | -0.31955 | 0.027 | 0.104 | 3.73E-46 |
| TRAPPC9 | 1.40E-50 | 0.465676 | 0.109 | 0.048 | 5.12E-46 |
| ABHD18 | 1.63E-50 | 0.442037 | 0.112 | 0.05 | 5.96E-46 |
| PHLDA1 | 1.70E-50 | -0.45974 | 0.095 | 0.203 | 6.22E-46 |
| PELI2 | 1.79E-50 | 0.70744 | 0.164 | 0.09 | 6.55E-46 |
| TEAD1 | 1.84E-50 | 0.607638 | 0.245 | 0.16 | 6.71E-46 |

| | | | | | |
|---|---|---|---|---|---|
| AKR1B1 | 1.85E-50 | -0.28408 | 0.047 | 0.136 | 6.76E-46 |
| ABHD14B | 1.98E-50 | -0.29927 | 0.049 | 0.14 | 7.24E-46 |
| MIEN1 | 2.06E-50 | -0.29323 | 0.026 | 0.102 | 7.52E-46 |
| AFF1 | 2.44E-50 | 0.610725 | 0.277 | 0.188 | 8.92E-46 |
| LRRC4C | 3.42E-50 | 0.515372 | 0.153 | 0.08 | 1.25E-45 |
| NRG1 | 3.99E-50 | 0.305235 | 0.1 | 0.042 | 1.46E-45 |
| NDUFB3 | 4.70E-50 | -0.26303 | 0.045 | 0.133 | 1.72E-45 |
| DPY30 | 5.03E-50 | -0.29455 | 0.061 | 0.157 | 1.84E-45 |
| DENND5B | 7.47E-50 | 0.507011 | 0.156 | 0.082 | 2.73E-45 |
| C17orf58 | 7.91E-50 | -0.3645 | 0.079 | 0.18 | 2.89E-45 |
| RNF181 | 9.22E-50 | -0.32533 | 0.073 | 0.175 | 3.37E-45 |
| DPT | 1.03E-49 | -0.61638 | 0.569 | 0.647 | 3.76E-45 |
| SOX6 | 1.24E-49 | 0.353334 | 0.107 | 0.047 | 4.54E-45 |
| PQBP1 | 1.29E-49 | -0.25025 | 0.037 | 0.119 | 4.71E-45 |
| CALN1 | 1.38E-49 | 0.350287 | 0.101 | 0.043 | 5.05E-45 |
| FAM69A | 1.43E-49 | 0.397215 | 0.153 | 0.078 | 5.22E-45 |
| ROCK2 | 1.44E-49 | 0.641544 | 0.378 | 0.296 | 5.27E-45 |
| ANAPC16 | 1.61E-49 | -0.31675 | 0.13 | 0.257 | 5.88E-45 |
| PPM1L | 1.79E-49 | 0.582984 | 0.121 | 0.057 | 6.55E-45 |
| PSME1 | 1.80E-49 | -0.37634 | 0.185 | 0.332 | 6.56E-45 |
| SH3KBP1 | 1.92E-49 | 0.562667 | 0.227 | 0.143 | 7.01E-45 |
| AK6 | 1.92E-49 | -0.25682 | 0.049 | 0.138 | 7.02E-45 |
| SMIM20 | 1.97E-49 | -0.30696 | 0.033 | 0.113 | 7.21E-45 |
| SVIL-AS1 | 2.00E-49 | 0.368093 | 0.103 | 0.044 | 7.31E-45 |
| MT-CO3 | 2.00E-49 | -0.40431 | 0.958 | 0.914 | 7.31E-45 |
| BICRAL | 2.47E-49 | 0.486925 | 0.137 | 0.068 | 9.02E-45 |
| IFI27L2 | 4.05E-49 | -0.44049 | 0.106 | 0.22 | 1.48E-44 |
| GAS6 | 4.14E-49 | -0.34722 | 0.187 | 0.32 | 1.51E-44 |
| ID4 | 4.24E-49 | -0.34576 | 0.042 | 0.126 | 1.55E-44 |
| GTF2H5 | 5.33E-49 | -0.31408 | 0.086 | 0.193 | 1.95E-44 |
| FAM208B | 5.90E-49 | 0.399959 | 0.159 | 0.082 | 2.16E-44 |
| GSTK1 | 7.17E-49 | -0.31093 | 0.11 | 0.227 | 2.62E-44 |
| DCTN6 | 8.21E-49 | -0.25063 | 0.059 | 0.152 | 3.00E-44 |
| SORCS2 | 9.39E-49 | 0.42316 | 0.123 | 0.058 | 3.43E-44 |
| PPIB | 1.17E-48 | -0.39426 | 0.383 | 0.539 | 4.29E-44 |
| DHFR | 1.25E-48 | 0.26207 | 0.327 | 0.21 | 4.57E-44 |
| DOCK1 | 1.31E-48 | 0.545034 | 0.213 | 0.131 | 4.79E-44 |
| SNRPF | 1.45E-48 | -0.30087 | 0.101 | 0.215 | 5.29E-44 |
| EIF3D | 1.98E-48 | -0.2682 | 0.101 | 0.214 | 7.24E-44 |
| TYMP | 2.04E-48 | -0.25115 | 0.106 | 0.219 | 7.45E-44 |
| PHF21A | 2.05E-48 | 0.595589 | 0.182 | 0.105 | 7.51E-44 |
| TIAM1 | 2.88E-48 | 0.411702 | 0.181 | 0.1 | 1.05E-43 |
| VTI1A | 3.88E-48 | 0.56296 | 0.171 | 0.096 | 1.42E-43 |
| NDUFA2 | 3.97E-48 | -0.37677 | 0.103 | 0.217 | 1.45E-43 |
| KDELC2 | 4.10E-48 | 0.350103 | 0.127 | 0.06 | 1.50E-43 |
| COG5 | 5.35E-48 | 0.508791 | 0.164 | 0.09 | 1.95E-43 |
| RASAL2 | 5.35E-48 | 0.454623 | 0.295 | 0.196 | 1.95E-43 |

| | | | | | |
|---|---|---|---|---|---|
| LITAF | 6.16E-48 | -0.34529 | 0.112 | 0.226 | 2.25E-43 |
| TBC1D4 | 6.49E-48 | 0.59878 | 0.161 | 0.089 | 2.37E-43 |
| UBE2A | 9.87E-48 | -0.25199 | 0.074 | 0.174 | 3.61E-43 |
| NSA2 | 1.21E-47 | -0.29801 | 0.149 | 0.278 | 4.42E-43 |
| LGALS3BP | 1.25E-47 | -0.38758 | 0.307 | 0.467 | 4.58E-43 |
| RNF38 | 1.53E-47 | 0.513074 | 0.135 | 0.069 | 5.59E-43 |
| LSM8 | 2.49E-47 | -0.27456 | 0.08 | 0.182 | 9.09E-43 |
| ATP11B | 2.57E-47 | 0.602035 | 0.171 | 0.098 | 9.40E-43 |
| SLC44A1 | 2.61E-47 | 0.644718 | 0.277 | 0.194 | 9.55E-43 |
| SSBP2 | 3.47E-47 | 0.767689 | 0.273 | 0.198 | 1.27E-42 |
| ANKAR | 3.69E-47 | 0.529373 | 0.133 | 0.067 | 1.35E-42 |
| ATXN7L1 | 3.73E-47 | 0.373076 | 0.101 | 0.044 | 1.36E-42 |
| NCOA2 | 3.95E-47 | 0.568824 | 0.189 | 0.112 | 1.44E-42 |
| SETBP1 | 4.48E-47 | 0.65569 | 0.27 | 0.187 | 1.64E-42 |
| ID2 | 5.35E-47 | -0.4348 | 0.116 | 0.227 | 1.96E-42 |
| ACAA2 | 5.38E-47 | -0.31974 | 0.072 | 0.168 | 1.97E-42 |
| EIF4B | 5.68E-47 | -0.29268 | 0.132 | 0.255 | 2.08E-42 |
| HILPDA | 6.31E-47 | -0.32214 | 0.037 | 0.114 | 2.31E-42 |
| ECM2 | 6.73E-47 | 0.609645 | 0.425 | 0.367 | 2.46E-42 |
| MTSS1 | 7.59E-47 | 0.665672 | 0.246 | 0.163 | 2.77E-42 |
| PPP6R3 | 2.11E-46 | 0.617545 | 0.215 | 0.136 | 7.70E-42 |
| PPP1R15A | 2.49E-46 | -0.46416 | 0.33 | 0.475 | 9.10E-42 |
| RBX1 | 2.52E-46 | -0.35452 | 0.153 | 0.283 | 9.23E-42 |
| TMEM176 | 3.16E-46 | -0.35176 | 0.167 | 0.294 | 1.16E-41 |
| HAS1 | 3.98E-46 | -0.60808 | 0.037 | 0.111 | 1.46E-41 |
| AC093249. | 4.18E-46 | 0.369628 | 0.107 | 0.048 | 1.53E-41 |
| CYTOR | 4.32E-46 | -0.28978 | 0.092 | 0.195 | 1.58E-41 |
| SREBF2 | 4.96E-46 | 0.496687 | 0.135 | 0.07 | 1.81E-41 |
| HTT | 5.37E-46 | 0.521015 | 0.146 | 0.078 | 1.96E-41 |
| CUL5 | 5.67E-46 | 0.565605 | 0.213 | 0.133 | 2.07E-41 |
| AL591368. | 5.98E-46 | 0.372331 | 0.118 | 0.056 | 2.18E-41 |
| ZHX2 | 6.13E-46 | 0.452597 | 0.143 | 0.075 | 2.24E-41 |
| ACYP2 | 6.42E-46 | 0.61637 | 0.177 | 0.103 | 2.35E-41 |
| TBCK | 6.63E-46 | 0.540786 | 0.136 | 0.07 | 2.42E-41 |
| PSME2 | 7.07E-46 | -0.30378 | 0.085 | 0.188 | 2.58E-41 |
| GSE1 | 7.64E-46 | 0.427699 | 0.133 | 0.068 | 2.79E-41 |
| SOS1 | 9.96E-46 | 0.661902 | 0.223 | 0.144 | 3.64E-41 |
| FBXL20 | 1.02E-45 | 0.534129 | 0.166 | 0.093 | 3.73E-41 |
| SNRPD2 | 1.02E-45 | -0.42918 | 0.197 | 0.338 | 3.73E-41 |
| DCAF10 | 1.05E-45 | 0.612013 | 0.194 | 0.117 | 3.84E-41 |
| ARL6IP5 | 1.12E-45 | -0.34459 | 0.332 | 0.501 | 4.08E-41 |
| PLXNA4 | 1.22E-45 | 0.518252 | 0.14 | 0.074 | 4.46E-41 |
| ANKRD28 | 1.52E-45 | 0.709171 | 0.336 | 0.26 | 5.54E-41 |
| TNFAIP6 | 1.69E-45 | -0.88099 | 0.168 | 0.277 | 6.17E-41 |
| ZNF292 | 1.76E-45 | 0.642752 | 0.239 | 0.158 | 6.43E-41 |
| SLC38A9 | 1.80E-45 | 0.471381 | 0.102 | 0.046 | 6.57E-41 |
| SCFD1 | 1.87E-45 | 0.660205 | 0.277 | 0.199 | 6.82E-41 |

| | | | | | |
|---|---|---|---|---|---|
| AGO4 | 2.23E-45 | 0.536108 | 0.113 | 0.054 | 8.16E-41 |
| CCNDBP1 | 2.45E-45 | -0.29185 | 0.071 | 0.166 | 8.97E-41 |
| ZNF609 | 2.48E-45 | 0.538441 | 0.179 | 0.104 | 9.05E-41 |
| TRPC1 | 2.75E-45 | 0.538333 | 0.141 | 0.074 | 1.01E-40 |
| MRPS21 | 3.32E-45 | -0.38951 | 0.13 | 0.25 | 1.21E-40 |
| DNPH1 | 3.43E-45 | -0.25454 | 0.035 | 0.111 | 1.25E-40 |
| GTF2IRD2 | 3.79E-45 | 0.456983 | 0.127 | 0.064 | 1.38E-40 |
| MRPL57 | 4.10E-45 | -0.34242 | 0.05 | 0.133 | 1.50E-40 |
| BABAM2 | 4.31E-45 | 0.653987 | 0.165 | 0.094 | 1.58E-40 |
| TROVE2 | 5.03E-45 | 0.318129 | 0.138 | 0.069 | 1.84E-40 |
| HIP1 | 5.38E-45 | 0.570115 | 0.152 | 0.083 | 1.97E-40 |
| ANKRD36E | 7.47E-45 | 0.4944 | 0.169 | 0.096 | 2.73E-40 |
| PPP1R10 | 7.62E-45 | -0.27036 | 0.037 | 0.113 | 2.79E-40 |
| AL138752. | 8.06E-45 | 0.365639 | 0.102 | 0.046 | 2.95E-40 |
| NDUFS3 | 9.04E-45 | -0.28377 | 0.051 | 0.135 | 3.30E-40 |
| STARD9 | 1.14E-44 | 0.50441 | 0.179 | 0.104 | 4.16E-40 |
| MBNL2 | 1.24E-44 | 0.742848 | 0.322 | 0.254 | 4.52E-40 |
| FOSL2 | 1.51E-44 | -0.25355 | 0.084 | 0.183 | 5.53E-40 |
| PSMB7 | 1.56E-44 | -0.29202 | 0.134 | 0.256 | 5.72E-40 |
| ZNF521 | 1.57E-44 | 0.482579 | 0.165 | 0.094 | 5.75E-40 |
| THADA | 1.59E-44 | 0.464451 | 0.145 | 0.077 | 5.82E-40 |
| AC010894 | 2.80E-44 | 0.370151 | 0.106 | 0.049 | 1.02E-39 |
| ATXN1 | 3.84E-44 | 0.559259 | 0.206 | 0.129 | 1.40E-39 |
| NDUFB9 | 4.35E-44 | -0.33369 | 0.158 | 0.287 | 1.59E-39 |
| ACSBG2 | 4.76E-44 | 0.302814 | 0.109 | 0.05 | 1.74E-39 |
| CTHRC1 | 5.06E-44 | -0.62455 | 0.115 | 0.212 | 1.85E-39 |
| ROCK1 | 8.40E-44 | 0.62827 | 0.389 | 0.324 | 3.07E-39 |
| HMGN3 | 1.07E-43 | -0.29653 | 0.145 | 0.268 | 3.92E-39 |
| VWA8 | 1.18E-43 | 0.437275 | 0.11 | 0.052 | 4.31E-39 |
| PPP6R2 | 1.19E-43 | 0.571851 | 0.143 | 0.078 | 4.35E-39 |
| CCBE1 | 1.31E-43 | 0.696635 | 0.189 | 0.118 | 4.78E-39 |
| TOX2 | 1.33E-43 | 0.432199 | 0.132 | 0.068 | 4.85E-39 |
| RGS10 | 1.39E-43 | -0.28068 | 0.05 | 0.132 | 5.09E-39 |
| PA2G4 | 1.44E-43 | -0.30335 | 0.162 | 0.291 | 5.27E-39 |
| RAD23A | 1.45E-43 | -0.2598 | 0.13 | 0.248 | 5.28E-39 |
| TSIX | 1.45E-43 | 0.294905 | 0.116 | 0.055 | 5.29E-39 |
| AHI1 | 1.56E-43 | 0.718727 | 0.366 | 0.303 | 5.71E-39 |
| ANGPTL1 | 1.60E-43 | 0.8174 | 0.282 | 0.211 | 5.85E-39 |
| SUMF1 | 2.12E-43 | 0.657067 | 0.202 | 0.127 | 7.76E-39 |
| CDK19 | 2.13E-43 | 0.396043 | 0.111 | 0.053 | 7.79E-39 |
| FCHSD2 | 3.52E-43 | 0.448197 | 0.115 | 0.056 | 1.29E-38 |
| GOLGB1 | 3.57E-43 | 0.599656 | 0.403 | 0.332 | 1.31E-38 |
| LPCAT2 | 4.33E-43 | 0.582758 | 0.125 | 0.064 | 1.58E-38 |
| RPS6KC1 | 5.26E-43 | 0.518632 | 0.13 | 0.067 | 1.92E-38 |
| FOSL1 | 6.00E-43 | -0.29234 | 0.051 | 0.132 | 2.19E-38 |
| MRPL55 | 6.62E-43 | -0.33876 | 0.042 | 0.118 | 2.42E-38 |
| NT5C2 | 6.71E-43 | 0.561522 | 0.208 | 0.132 | 2.45E-38 |

| | | | | | |
|---|---|---|---|---|---|
| PDAP1 | 8.11E-43 | -0.25209 | 0.135 | 0.254 | 2.97E-38 |
| USP34 | 8.13E-43 | 0.597177 | 0.277 | 0.2 | 2.97E-38 |
| PSMC3 | 8.90E-43 | -0.25712 | 0.099 | 0.204 | 3.25E-38 |
| PATJ | 1.07E-42 | 0.410821 | 0.107 | 0.051 | 3.93E-38 |
| EIF5 | 1.37E-42 | -0.25931 | 0.243 | 0.392 | 5.00E-38 |
| ROR1 | 1.52E-42 | 0.474336 | 0.141 | 0.076 | 5.54E-38 |
| PMM1 | 2.20E-42 | -0.25142 | 0.033 | 0.104 | 8.02E-38 |
| SMC5 | 2.46E-42 | 0.610342 | 0.32 | 0.245 | 8.98E-38 |
| CCDC152 | 2.63E-42 | 0.346747 | 0.123 | 0.062 | 9.62E-38 |
| ZRANB3 | 2.90E-42 | 0.356172 | 0.109 | 0.052 | 1.06E-37 |
| SSR3 | 3.49E-42 | -0.27316 | 0.197 | 0.336 | 1.27E-37 |
| RHOA | 3.52E-42 | -0.4366 | 0.349 | 0.507 | 1.29E-37 |
| ARFGEF2 | 3.96E-42 | 0.423865 | 0.115 | 0.057 | 1.45E-37 |
| DBI | 4.07E-42 | -0.31048 | 0.235 | 0.386 | 1.49E-37 |
| PSMA7 | 4.51E-42 | -0.37501 | 0.311 | 0.476 | 1.65E-37 |
| PHIP | 5.00E-42 | 0.657663 | 0.305 | 0.231 | 1.83E-37 |
| ARID2 | 5.35E-42 | 0.46224 | 0.156 | 0.088 | 1.96E-37 |
| NBPF11 | 7.14E-42 | 0.363334 | 0.11 | 0.053 | 2.61E-37 |
| TTLL5 | 8.48E-42 | 0.412304 | 0.111 | 0.054 | 3.10E-37 |
| TBCA | 9.01E-42 | -0.25046 | 0.213 | 0.357 | 3.29E-37 |
| NEMF | 9.87E-42 | 0.598287 | 0.277 | 0.2 | 3.61E-37 |
| BOD1L1 | 1.58E-41 | 0.634008 | 0.384 | 0.321 | 5.79E-37 |
| ZDHHC14 | 1.68E-41 | 0.415089 | 0.108 | 0.052 | 6.15E-37 |
| JAZF1 | 2.07E-41 | 0.594006 | 0.184 | 0.114 | 7.55E-37 |
| RBPMS | 2.19E-41 | 0.800827 | 0.287 | 0.22 | 8.01E-37 |
| RASD1 | 2.25E-41 | -0.88087 | 0.193 | 0.3 | 8.24E-37 |
| CEP350 | 2.70E-41 | 0.541607 | 0.263 | 0.182 | 9.86E-37 |
| ZRANB2 | 2.83E-41 | 0.633031 | 0.352 | 0.286 | 1.04E-36 |
| TWIST1 | 3.21E-41 | -0.27828 | 0.104 | 0.204 | 1.17E-36 |
| CP | 3.72E-41 | 0.535422 | 0.151 | 0.085 | 1.36E-36 |
| WRN | 4.36E-41 | 0.554792 | 0.152 | 0.086 | 1.59E-36 |
| PDSS2 | 4.82E-41 | 0.476725 | 0.134 | 0.071 | 1.76E-36 |
| TRPC4AP | 5.48E-41 | 0.60631 | 0.173 | 0.104 | 2.00E-36 |
| UBE2E2 | 5.75E-41 | 0.646233 | 0.269 | 0.191 | 2.10E-36 |
| NBPF9 | 5.85E-41 | 0.374161 | 0.106 | 0.051 | 2.14E-36 |
| KDM3B | 6.56E-41 | 0.510366 | 0.144 | 0.08 | 2.40E-36 |
| HMGB1 | 6.65E-41 | -0.38414 | 0.47 | 0.608 | 2.43E-36 |
| CYB5R3 | 6.82E-41 | -0.29637 | 0.236 | 0.384 | 2.49E-36 |
| MYH10 | 7.39E-41 | 0.538009 | 0.175 | 0.106 | 2.70E-36 |
| AC058822 | 7.73E-41 | 0.31984 | 0.109 | 0.052 | 2.83E-36 |
| SERBP1 | 9.35E-41 | -0.29997 | 0.249 | 0.398 | 3.42E-36 |
| RAB3GAP2 | 1.78E-40 | 0.486157 | 0.159 | 0.092 | 6.51E-36 |
| C14orf37 | 1.79E-40 | 0.355282 | 0.104 | 0.049 | 6.54E-36 |
| PXN | 1.93E-40 | 0.481563 | 0.255 | 0.175 | 7.06E-36 |
| PIGL | 2.41E-40 | 0.461669 | 0.134 | 0.072 | 8.80E-36 |
| LEPROT | 2.52E-40 | -0.28053 | 0.257 | 0.403 | 9.19E-36 |
| VPS45 | 2.52E-40 | 0.472431 | 0.113 | 0.057 | 9.20E-36 |

| | | | | | |
|---|---|---|---|---|---|
| UGP2 | 3.37E-40 | -0.35608 | 0.198 | 0.322 | 1.23E-35 |
| RBPJ | 4.44E-40 | 0.509518 | 0.44 | 0.395 | 1.62E-35 |
| FN1 | 5.86E-40 | 0.446318 | 0.468 | 0.384 | 2.14E-35 |
| ACACA | 7.55E-40 | 0.399738 | 0.13 | 0.068 | 2.76E-35 |
| EGR1 | 8.24E-40 | -0.41203 | 0.556 | 0.66 | 3.01E-35 |
| TAOK3 | 8.26E-40 | 0.550648 | 0.244 | 0.168 | 3.02E-35 |
| CDK4 | 8.40E-40 | -0.27314 | 0.061 | 0.144 | 3.07E-35 |
| CDKN1C | 1.07E-39 | -0.28445 | 0.185 | 0.31 | 3.92E-35 |
| CYP1B1 | 1.26E-39 | -0.43883 | 0.044 | 0.115 | 4.61E-35 |
| ARID4B | 1.59E-39 | 0.605424 | 0.375 | 0.314 | 5.83E-35 |
| DST | 2.02E-39 | 0.314944 | 0.558 | 0.491 | 7.38E-35 |
| STEAP4 | 2.12E-39 | 0.495246 | 0.204 | 0.131 | 7.73E-35 |
| PIBF1 | 2.41E-39 | 0.622167 | 0.202 | 0.131 | 8.79E-35 |
| ESYT2 | 2.50E-39 | 0.630076 | 0.332 | 0.262 | 9.14E-35 |
| MTA3 | 2.64E-39 | 0.470886 | 0.133 | 0.073 | 9.66E-35 |
| SLC39A11 | 3.83E-39 | 0.405202 | 0.109 | 0.054 | 1.40E-34 |
| UBR1 | 4.17E-39 | 0.52173 | 0.178 | 0.109 | 1.52E-34 |
| ADM | 4.45E-39 | -0.88141 | 0.193 | 0.296 | 1.63E-34 |
| THRB | 5.25E-39 | 0.613003 | 0.176 | 0.109 | 1.92E-34 |
| TCF7L1 | 6.35E-39 | 0.633153 | 0.242 | 0.173 | 2.32E-34 |
| CDC27 | 6.64E-39 | 0.684871 | 0.244 | 0.172 | 2.43E-34 |
| ANGPTL4 | 1.24E-38 | -0.4675 | 0.144 | 0.246 | 4.53E-34 |
| ZBTB16 | 1.44E-38 | 0.600054 | 0.465 | 0.426 | 5.26E-34 |
| EBF3 | 1.65E-38 | 0.583615 | 0.183 | 0.117 | 6.03E-34 |
| EEA1 | 1.76E-38 | 0.52443 | 0.448 | 0.397 | 6.42E-34 |
| KDM2A | 1.94E-38 | 0.55165 | 0.228 | 0.155 | 7.11E-34 |
| CLASP1 | 2.11E-38 | 0.434206 | 0.146 | 0.083 | 7.71E-34 |
| PPP1R12A | 2.56E-38 | 0.581804 | 0.347 | 0.282 | 9.36E-34 |
| NLRP1 | 3.30E-38 | 0.399753 | 0.104 | 0.051 | 1.21E-33 |
| PLAU | 4.34E-38 | -0.37462 | 0.064 | 0.143 | 1.58E-33 |
| SMG6 | 4.80E-38 | 0.571473 | 0.201 | 0.131 | 1.75E-33 |
| ASAP1 | 4.86E-38 | 0.551425 | 0.248 | 0.173 | 1.78E-33 |
| TANGO6 | 5.19E-38 | 0.377619 | 0.106 | 0.052 | 1.90E-33 |
| NFAT5 | 5.50E-38 | 0.631057 | 0.288 | 0.218 | 2.01E-33 |
| ITGA11 | 5.52E-38 | 0.468028 | 0.205 | 0.131 | 2.02E-33 |
| SUPT3H | 5.87E-38 | 0.4623 | 0.117 | 0.061 | 2.15E-33 |
| SPG11 | 6.13E-38 | 0.486936 | 0.158 | 0.093 | 2.24E-33 |
| APOE | 6.18E-38 | -0.98745 | 0.134 | 0.22 | 2.26E-33 |
| PRR13 | 9.72E-38 | -0.36161 | 0.051 | 0.125 | 3.55E-33 |
| ORC2 | 1.01E-37 | 0.430414 | 0.107 | 0.053 | 3.67E-33 |
| MLLT10 | 1.37E-37 | 0.480572 | 0.16 | 0.096 | 4.99E-33 |
| NEXN | 1.39E-37 | 0.6678 | 0.324 | 0.26 | 5.09E-33 |
| ENTPD1-A! | 1.68E-37 | 0.568656 | 0.126 | 0.068 | 6.15E-33 |
| CDK13 | 1.99E-37 | 0.590003 | 0.23 | 0.159 | 7.28E-33 |
| CWF19L2 | 2.33E-37 | 0.546752 | 0.235 | 0.162 | 8.50E-33 |
| CNTRL | 2.40E-37 | 0.530662 | 0.148 | 0.086 | 8.76E-33 |
| FAM214A | 2.55E-37 | 0.564384 | 0.183 | 0.116 | 9.31E-33 |

| | | | | | |
|---|---|---|---|---|---|
| THOC2 | 2.88E-37 | 0.57097 | 0.291 | 0.221 | 1.05E-32 |
| RABGAP1 | 2.94E-37 | 0.47061 | 0.156 | 0.092 | 1.07E-32 |
| CACNA1A | 4.39E-37 | 0.341188 | 0.108 | 0.054 | 1.60E-32 |
| ATOX1 | 4.54E-37 | -0.33654 | 0.119 | 0.219 | 1.66E-32 |
| MBTD1 | 6.06E-37 | 0.408658 | 0.107 | 0.054 | 2.21E-32 |
| DENND4A | 6.87E-37 | 0.426045 | 0.195 | 0.123 | 2.51E-32 |
| DOCK4 | 7.95E-37 | 0.369198 | 0.247 | 0.166 | 2.91E-32 |
| PCNX4 | 1.35E-36 | 0.583092 | 0.173 | 0.107 | 4.92E-32 |
| ANKRD26 | 1.41E-36 | 0.537992 | 0.162 | 0.098 | 5.15E-32 |
| NPAS3 | 1.51E-36 | 0.372009 | 0.1 | 0.049 | 5.50E-32 |
| BBS9 | 3.35E-36 | 0.478552 | 0.113 | 0.059 | 1.22E-31 |
| RB1CC1 | 3.96E-36 | 0.509483 | 0.348 | 0.279 | 1.45E-31 |
| SIPA1L3 | 4.19E-36 | 0.317847 | 0.102 | 0.05 | 1.53E-31 |
| MRPL21 | 5.52E-36 | -0.27018 | 0.054 | 0.128 | 2.02E-31 |
| VPS13D | 9.27E-36 | 0.514107 | 0.173 | 0.109 | 3.39E-31 |
| ZC3H13 | 9.85E-36 | 0.56171 | 0.328 | 0.258 | 3.60E-31 |
| C11orf49 | 9.89E-36 | 0.534716 | 0.151 | 0.09 | 3.61E-31 |
| AL445426. | 1.04E-35 | 0.512972 | 0.132 | 0.075 | 3.78E-31 |
| SERPINF1 | 1.21E-35 | -0.38699 | 0.636 | 0.683 | 4.42E-31 |
| CALR | 2.05E-35 | -0.26246 | 0.274 | 0.415 | 7.50E-31 |
| ECHDC2 | 2.10E-35 | 0.606884 | 0.299 | 0.238 | 7.66E-31 |
| NBPF14 | 2.26E-35 | 0.344526 | 0.138 | 0.077 | 8.26E-31 |
| PNN | 2.67E-35 | 0.625642 | 0.391 | 0.346 | 9.75E-31 |
| SETD2 | 2.79E-35 | 0.556383 | 0.25 | 0.179 | 1.02E-30 |
| TRPM7 | 3.01E-35 | 0.526409 | 0.202 | 0.133 | 1.10E-30 |
| USP25 | 4.26E-35 | 0.522818 | 0.172 | 0.108 | 1.56E-30 |
| AP3B1 | 5.52E-35 | 0.547966 | 0.213 | 0.144 | 2.02E-30 |
| SEC22A | 5.80E-35 | 0.511805 | 0.106 | 0.055 | 2.12E-30 |
| PRICKLE1 | 8.00E-35 | 0.555873 | 0.139 | 0.081 | 2.93E-30 |
| SIPA1L1 | 9.19E-35 | 0.382328 | 0.11 | 0.057 | 3.36E-30 |
| TUBB | 1.00E-34 | -0.40362 | 0.272 | 0.409 | 3.66E-30 |
| PDS5B | 1.37E-34 | 0.525335 | 0.197 | 0.131 | 5.02E-30 |
| SRGAP1 | 1.53E-34 | 0.408005 | 0.304 | 0.231 | 5.61E-30 |
| AMBRA1 | 1.77E-34 | 0.353787 | 0.209 | 0.134 | 6.48E-30 |
| PSEN1 | 2.31E-34 | 0.493596 | 0.167 | 0.104 | 8.44E-30 |
| NFATC3 | 2.40E-34 | 0.377543 | 0.131 | 0.073 | 8.77E-30 |
| BMPR2 | 2.43E-34 | 0.565936 | 0.259 | 0.189 | 8.87E-30 |
| SYN3 | 2.57E-34 | 0.255714 | 0.226 | 0.144 | 9.38E-30 |
| STIM1 | 3.18E-34 | 0.487973 | 0.189 | 0.123 | 1.16E-29 |
| YTHDC1 | 3.68E-34 | 0.552033 | 0.275 | 0.207 | 1.34E-29 |
| ITGB1BP1 | 4.20E-34 | -0.25161 | 0.196 | 0.322 | 1.53E-29 |
| ATG10 | 4.95E-34 | 0.421353 | 0.149 | 0.088 | 1.81E-29 |
| CEP83 | 5.48E-34 | 0.452849 | 0.117 | 0.064 | 2.00E-29 |
| TTC3 | 8.95E-34 | 0.519864 | 0.388 | 0.334 | 3.27E-29 |
| CREBBP | 1.35E-33 | 0.49254 | 0.226 | 0.159 | 4.94E-29 |
| COX7A1 | 1.44E-33 | -0.28421 | 0.139 | 0.244 | 5.26E-29 |
| PUM2 | 1.62E-33 | 0.512049 | 0.162 | 0.101 | 5.92E-29 |

| | | | | | |
|---|---|---|---|---|---|
| CCAR1 | 1.76E-33 | 0.53269 | 0.222 | 0.155 | 6.43E-29 |
| SCAF11 | 2.92E-33 | 0.521991 | 0.425 | 0.385 | 1.07E-28 |
| APPBP2 | 3.11E-33 | 0.565096 | 0.181 | 0.119 | 1.14E-28 |
| CNST | 3.44E-33 | 0.503823 | 0.132 | 0.077 | 1.26E-28 |
| SLC16A1-A | 4.16E-33 | 0.482114 | 0.132 | 0.076 | 1.52E-28 |
| MITF | 4.70E-33 | 0.643999 | 0.224 | 0.162 | 1.72E-28 |
| NIPAL2 | 5.02E-33 | 0.502652 | 0.124 | 0.07 | 1.84E-28 |
| CLDN11 | 5.06E-33 | -0.2596 | 0.078 | 0.156 | 1.85E-28 |
| PLCL2 | 5.32E-33 | 0.421124 | 0.117 | 0.064 | 1.95E-28 |
| GAREM1 | 6.02E-33 | 0.452566 | 0.116 | 0.063 | 2.20E-28 |
| SLTM | 7.02E-33 | 0.553324 | 0.329 | 0.27 | 2.57E-28 |
| BACH2 | 7.99E-33 | 0.372503 | 0.128 | 0.072 | 2.92E-28 |
| CDR1-antis | 8.15E-33 | 0.28569 | 0.102 | 0.051 | 2.98E-28 |
| TAF1 | 8.67E-33 | 0.432269 | 0.144 | 0.085 | 3.17E-28 |
| ANKIB1 | 9.96E-33 | 0.558366 | 0.213 | 0.148 | 3.64E-28 |
| RAB8B | 1.05E-32 | 0.676942 | 0.263 | 0.202 | 3.85E-28 |
| PALLD | 1.31E-32 | 0.566437 | 0.359 | 0.305 | 4.78E-28 |
| MRPL33 | 1.37E-32 | -0.26253 | 0.116 | 0.21 | 5.00E-28 |
| MICU2 | 1.41E-32 | 0.642568 | 0.195 | 0.133 | 5.15E-28 |
| RNPC3 | 2.08E-32 | 0.486914 | 0.157 | 0.098 | 7.59E-28 |
| GPC6 | 3.42E-32 | 0.434693 | 0.149 | 0.09 | 1.25E-27 |
| CNTLN | 4.52E-32 | 0.590217 | 0.186 | 0.125 | 1.65E-27 |
| RPAP2 | 4.52E-32 | 0.471598 | 0.162 | 0.101 | 1.65E-27 |
| ERRFI1 | 4.95E-32 | -0.25746 | 0.296 | 0.428 | 1.81E-27 |
| SNX29 | 5.77E-32 | 0.474464 | 0.214 | 0.149 | 2.11E-27 |
| UBR5 | 8.81E-32 | 0.568656 | 0.229 | 0.165 | 3.22E-27 |
| RDH10 | 1.17E-31 | -0.29448 | 0.046 | 0.109 | 4.29E-27 |
| ZCCHC7 | 1.27E-31 | 0.507454 | 0.205 | 0.141 | 4.63E-27 |
| CEP290 | 1.54E-31 | 0.533489 | 0.17 | 0.11 | 5.62E-27 |
| FOXO3 | 1.80E-31 | 0.573914 | 0.374 | 0.32 | 6.57E-27 |
| AKAP10 | 1.98E-31 | 0.40199 | 0.101 | 0.053 | 7.23E-27 |
| SYNE2 | 2.08E-31 | 0.485249 | 0.268 | 0.198 | 7.59E-27 |
| FOXP1 | 2.10E-31 | 0.45247 | 0.379 | 0.322 | 7.69E-27 |
| SRSF4 | 2.12E-31 | 0.571463 | 0.336 | 0.282 | 7.74E-27 |
| ELF2 | 2.12E-31 | 0.532016 | 0.335 | 0.275 | 7.76E-27 |
| CASK | 2.43E-31 | 0.512578 | 0.182 | 0.12 | 8.89E-27 |
| ZSWIM6 | 2.50E-31 | 0.421401 | 0.204 | 0.136 | 9.12E-27 |
| SOX4 | 2.89E-31 | -0.30577 | 0.087 | 0.166 | 1.06E-26 |
| ADGRL2 | 3.02E-31 | 0.495867 | 0.124 | 0.072 | 1.10E-26 |
| WSB1 | 3.81E-31 | 0.432743 | 0.498 | 0.481 | 1.39E-26 |
| CDK17 | 4.16E-31 | 0.56098 | 0.151 | 0.095 | 1.52E-26 |
| YLPM1 | 4.74E-31 | 0.427741 | 0.141 | 0.085 | 1.73E-26 |
| USP4 | 4.93E-31 | 0.437187 | 0.121 | 0.069 | 1.80E-26 |
| YBX3 | 5.79E-31 | -0.30576 | 0.461 | 0.569 | 2.12E-26 |
| KIAA1109 | 6.71E-31 | 0.486655 | 0.218 | 0.155 | 2.45E-26 |
| MTRNR2L3 | 9.26E-31 | 0.349908 | 0.171 | 0.106 | 3.39E-26 |
| CAPN7 | 9.58E-31 | 0.514844 | 0.144 | 0.089 | 3.50E-26 |

| | | | | | |
|---|---|---|---|---|---|
| TGFBR3 | 1.30E-30 | 0.286802 | 0.453 | 0.383 | 4.74E-26 |
| ALDH1A1 | 1.47E-30 | -0.35236 | 0.178 | 0.278 | 5.38E-26 |
| UTRN | 1.49E-30 | 0.382245 | 0.352 | 0.282 | 5.45E-26 |
| PHC3 | 1.61E-30 | 0.477935 | 0.143 | 0.088 | 5.89E-26 |
| TPM1 | 1.63E-30 | -0.37679 | 0.267 | 0.389 | 5.95E-26 |
| CNKSR2 | 1.64E-30 | 0.398346 | 0.115 | 0.065 | 5.98E-26 |
| SH3PXD2B | 1.93E-30 | 0.478702 | 0.247 | 0.178 | 7.07E-26 |
| NCOA6 | 2.53E-30 | 0.388394 | 0.119 | 0.068 | 9.26E-26 |
| ZNF280D | 2.78E-30 | 0.522347 | 0.194 | 0.133 | 1.02E-25 |
| SPRED2 | 2.84E-30 | 0.556823 | 0.134 | 0.081 | 1.04E-25 |
| OSBPL8 | 2.92E-30 | 0.594856 | 0.357 | 0.31 | 1.07E-25 |
| TENM1 | 3.23E-30 | 0.384413 | 0.12 | 0.068 | 1.18E-25 |
| RBM5 | 3.26E-30 | 0.509033 | 0.243 | 0.182 | 1.19E-25 |
| TIMP3 | 3.34E-30 | -0.31506 | 0.696 | 0.745 | 1.22E-25 |
| ECE1 | 4.49E-30 | 0.501272 | 0.207 | 0.145 | 1.64E-25 |
| PDXDC1 | 5.29E-30 | 0.411592 | 0.173 | 0.113 | 1.93E-25 |
| ITPR2 | 5.33E-30 | 0.440755 | 0.129 | 0.076 | 1.95E-25 |
| LSP1 | 6.08E-30 | -0.31287 | 0.2 | 0.317 | 2.22E-25 |
| DBN1 | 7.52E-30 | -0.28598 | 0.1 | 0.181 | 2.75E-25 |
| COMMD1( | 8.94E-30 | 0.546876 | 0.158 | 0.102 | 3.27E-25 |
| GLG1 | 1.13E-29 | 0.476017 | 0.347 | 0.295 | 4.14E-25 |
| SMURF1 | 1.15E-29 | 0.410449 | 0.111 | 0.062 | 4.20E-25 |
| ARFGEF1 | 1.46E-29 | 0.505031 | 0.179 | 0.12 | 5.34E-25 |
| NID2 | 1.95E-29 | 0.502596 | 0.142 | 0.089 | 7.12E-25 |
| SLMAP | 2.03E-29 | 0.578056 | 0.158 | 0.101 | 7.42E-25 |
| RLF | 2.12E-29 | 0.56771 | 0.201 | 0.142 | 7.74E-25 |
| KDM5A | 2.34E-29 | 0.473342 | 0.211 | 0.15 | 8.54E-25 |
| TCERG1 | 2.82E-29 | 0.443748 | 0.224 | 0.16 | 1.03E-24 |
| CADPS2 | 3.01E-29 | 0.404292 | 0.127 | 0.075 | 1.10E-24 |
| ITGA9 | 4.42E-29 | 0.395062 | 0.121 | 0.07 | 1.61E-24 |
| ATR | 4.43E-29 | 0.421651 | 0.136 | 0.082 | 1.62E-24 |
| EFR3A | 5.45E-29 | 0.548819 | 0.165 | 0.109 | 1.99E-24 |
| LMNA | 5.67E-29 | -0.44119 | 0.601 | 0.659 | 2.07E-24 |
| RALGAPA1 | 6.94E-29 | 0.548228 | 0.221 | 0.161 | 2.54E-24 |
| MDN1 | 6.97E-29 | 0.373842 | 0.117 | 0.067 | 2.55E-24 |
| EVI5 | 7.15E-29 | 0.525635 | 0.219 | 0.158 | 2.61E-24 |
| RAPGEF2 | 7.38E-29 | 0.43614 | 0.11 | 0.062 | 2.70E-24 |
| R3HDM1 | 7.80E-29 | 0.422689 | 0.113 | 0.064 | 2.85E-24 |
| EPB41L2 | 8.84E-29 | 0.507119 | 0.45 | 0.426 | 3.23E-24 |
| MXRA5 | 9.66E-29 | 0.539356 | 0.177 | 0.121 | 3.53E-24 |
| MEF2A | 9.68E-29 | 0.482381 | 0.289 | 0.226 | 3.54E-24 |
| ZFP36L2 | 1.25E-28 | -0.49933 | 0.505 | 0.582 | 4.58E-24 |
| KANSL1L | 1.35E-28 | 0.470441 | 0.135 | 0.082 | 4.95E-24 |
| UPF2 | 1.45E-28 | 0.471493 | 0.17 | 0.113 | 5.29E-24 |
| SEC31A | 1.72E-28 | 0.502751 | 0.371 | 0.327 | 6.30E-24 |
| TYW1 | 1.89E-28 | 0.390612 | 0.11 | 0.062 | 6.92E-24 |
| FOXN2 | 1.92E-28 | 0.47196 | 0.107 | 0.06 | 7.02E-24 |

| | | | | | |
|---|---|---|---|---|---|
| SDCCAG8 | 1.98E-28 | 0.553714 | 0.204 | 0.146 | 7.23E-24 |
| POGZ | 2.12E-28 | 0.508492 | 0.193 | 0.134 | 7.74E-24 |
| NXN | 2.57E-28 | 0.432331 | 0.117 | 0.068 | 9.40E-24 |
| USP24 | 3.23E-28 | 0.388136 | 0.108 | 0.061 | 1.18E-23 |
| PRRC2C | 3.37E-28 | 0.362266 | 0.524 | 0.509 | 1.23E-23 |
| ITSN2 | 3.54E-28 | 0.519207 | 0.199 | 0.139 | 1.30E-23 |
| PER3 | 4.68E-28 | 0.630405 | 0.2 | 0.145 | 1.71E-23 |
| PARP4 | 5.16E-28 | 0.47846 | 0.159 | 0.103 | 1.89E-23 |
| PACSIN2 | 5.31E-28 | 0.434675 | 0.155 | 0.1 | 1.94E-23 |
| ARHGAP21 | 6.56E-28 | 0.611473 | 0.366 | 0.33 | 2.40E-23 |
| GNAS | 7.25E-28 | -0.25102 | 0.506 | 0.602 | 2.65E-23 |
| DDX17 | 7.62E-28 | 0.453846 | 0.441 | 0.415 | 2.79E-23 |
| FAXDC2 | 8.46E-28 | 0.48168 | 0.157 | 0.103 | 3.09E-23 |
| AC008440 | 8.61E-28 | 0.288178 | 0.154 | 0.095 | 3.15E-23 |
| NUMB | 9.76E-28 | 0.554875 | 0.216 | 0.159 | 3.57E-23 |
| CMSS1 | 1.09E-27 | 0.420556 | 0.22 | 0.159 | 4.00E-23 |
| RB1 | 1.17E-27 | 0.456794 | 0.13 | 0.079 | 4.27E-23 |
| PI4KA | 1.18E-27 | 0.413161 | 0.116 | 0.068 | 4.32E-23 |
| HMBOX1 | 1.26E-27 | 0.433824 | 0.13 | 0.079 | 4.60E-23 |
| C1orf21 | 1.45E-27 | 0.486911 | 0.433 | 0.403 | 5.28E-23 |
| GBE1 | 1.56E-27 | 0.514072 | 0.224 | 0.162 | 5.71E-23 |
| FNBP1 | 1.62E-27 | 0.460707 | 0.192 | 0.134 | 5.93E-23 |
| RBMS1 | 1.71E-27 | 0.460818 | 0.44 | 0.414 | 6.27E-23 |
| MAP2K4 | 1.73E-27 | 0.436038 | 0.113 | 0.065 | 6.32E-23 |
| CELF1 | 2.23E-27 | 0.474678 | 0.179 | 0.123 | 8.16E-23 |
| KDM6A | 2.39E-27 | 0.371335 | 0.113 | 0.066 | 8.73E-23 |
| RICTOR | 2.58E-27 | 0.508224 | 0.199 | 0.14 | 9.43E-23 |
| GNG2 | 2.98E-27 | 0.465447 | 0.128 | 0.078 | 1.09E-22 |
| PTPRT | 3.01E-27 | 0.270247 | 0.113 | 0.064 | 1.10E-22 |
| WDFY2 | 3.73E-27 | 0.451004 | 0.173 | 0.116 | 1.36E-22 |
| SOBP | 3.75E-27 | 0.521947 | 0.166 | 0.111 | 1.37E-22 |
| KLHL29 | 3.96E-27 | 0.396494 | 0.134 | 0.083 | 1.45E-22 |
| RNGTT | 5.20E-27 | 0.401924 | 0.1 | 0.056 | 1.90E-22 |
| ANKRD13C | 5.93E-27 | 0.457768 | 0.142 | 0.09 | 2.17E-22 |
| SNX9 | 6.07E-27 | 0.570948 | 0.39 | 0.358 | 2.22E-22 |
| ANGPT1 | 8.61E-27 | 0.580794 | 0.141 | 0.091 | 3.15E-22 |
| FRMD6 | 9.62E-27 | 0.588032 | 0.194 | 0.14 | 3.52E-22 |
| ZNF438 | 1.15E-26 | 0.384964 | 0.117 | 0.069 | 4.22E-22 |
| TAGLN | 1.21E-26 | -0.48957 | 0.181 | 0.271 | 4.41E-22 |
| NEK11 | 1.30E-26 | 0.389604 | 0.115 | 0.068 | 4.75E-22 |
| EPN2 | 1.33E-26 | 0.60189 | 0.205 | 0.152 | 4.87E-22 |
| TBK1 | 1.39E-26 | 0.503428 | 0.153 | 0.101 | 5.09E-22 |
| TBC1D16 | 1.48E-26 | 0.353932 | 0.127 | 0.077 | 5.40E-22 |
| MED13 | 1.94E-26 | 0.392796 | 0.207 | 0.15 | 7.09E-22 |
| BDP1 | 1.99E-26 | 0.522135 | 0.286 | 0.229 | 7.28E-22 |
| MTMR3 | 2.17E-26 | 0.425346 | 0.139 | 0.087 | 7.93E-22 |
| CALM2 | 2.18E-26 | -0.31278 | 0.459 | 0.599 | 7.98E-22 |

| | | | | | |
|---|---|---|---|---|---|
| KIAA0319L | 2.57E-26 | 0.386947 | 0.113 | 0.066 | 9.41E-22 |
| OXR1 | 2.76E-26 | 0.559408 | 0.218 | 0.162 | 1.01E-21 |
| RC3H1 | 3.48E-26 | 0.461867 | 0.148 | 0.096 | 1.27E-21 |
| PLA2R1 | 3.97E-26 | 0.496207 | 0.155 | 0.104 | 1.45E-21 |
| TAF2 | 4.05E-26 | 0.436152 | 0.122 | 0.074 | 1.48E-21 |
| ASPN | 4.61E-26 | -0.33502 | 0.237 | 0.336 | 1.69E-21 |
| CPQ | 4.94E-26 | 0.519248 | 0.427 | 0.41 | 1.80E-21 |
| CDC42SE2 | 4.96E-26 | 0.428965 | 0.111 | 0.065 | 1.81E-21 |
| STK39 | 5.20E-26 | 0.366563 | 0.102 | 0.058 | 1.90E-21 |
| ARID4A | 5.25E-26 | 0.473471 | 0.187 | 0.13 | 1.92E-21 |
| OGFRL1 | 5.34E-26 | 0.71442 | 0.249 | 0.2 | 1.95E-21 |
| GIGYF2 | 6.38E-26 | 0.487005 | 0.19 | 0.134 | 2.33E-21 |
| BOC | 8.11E-26 | 0.591266 | 0.293 | 0.247 | 2.96E-21 |
| AC242426 | 8.34E-26 | 0.285401 | 0.136 | 0.082 | 3.05E-21 |
| MPDZ | 1.09E-25 | 0.510769 | 0.184 | 0.13 | 3.98E-21 |
| USP32 | 1.18E-25 | 0.440359 | 0.141 | 0.089 | 4.32E-21 |
| KANK1 | 1.37E-25 | 0.275642 | 0.153 | 0.098 | 5.01E-21 |
| USP48 | 1.59E-25 | 0.515361 | 0.184 | 0.13 | 5.80E-21 |
| GTDC1 | 1.65E-25 | 0.39604 | 0.108 | 0.063 | 6.02E-21 |
| UBE3C | 1.87E-25 | 0.378033 | 0.188 | 0.129 | 6.83E-21 |
| ETV6 | 1.95E-25 | 0.379521 | 0.106 | 0.061 | 7.13E-21 |
| MRC2 | 2.15E-25 | 0.472883 | 0.352 | 0.311 | 7.86E-21 |
| CTSK | 2.44E-25 | -0.3264 | 0.302 | 0.422 | 8.92E-21 |
| ABI3BP | 2.87E-25 | 0.271639 | 0.519 | 0.492 | 1.05E-20 |
| FILIP1L | 3.18E-25 | 0.425933 | 0.47 | 0.439 | 1.16E-20 |
| IQGAP1 | 3.84E-25 | 0.42795 | 0.396 | 0.355 | 1.40E-20 |
| MOCS2 | 4.03E-25 | -0.27464 | 0.046 | 0.1 | 1.47E-20 |
| DMTF1 | 4.42E-25 | 0.424864 | 0.119 | 0.073 | 1.61E-20 |
| TMCC1 | 4.85E-25 | 0.352982 | 0.115 | 0.068 | 1.77E-20 |
| EID1 | 6.41E-25 | -0.29516 | 0.493 | 0.622 | 2.34E-20 |
| ST3GAL3 | 6.85E-25 | 0.394814 | 0.1 | 0.057 | 2.50E-20 |
| ZNF721 | 7.17E-25 | 0.451784 | 0.154 | 0.103 | 2.62E-20 |
| FYN | 7.24E-25 | 0.575873 | 0.359 | 0.331 | 2.64E-20 |
| ARHGAP1( | 7.31E-25 | 0.589653 | 0.295 | 0.25 | 2.67E-20 |
| KDM4C | 9.57E-25 | 0.367115 | 0.136 | 0.086 | 3.50E-20 |
| PDS5A | 1.11E-24 | 0.47405 | 0.221 | 0.165 | 4.07E-20 |
| ELN | 1.13E-24 | 0.463744 | 0.368 | 0.316 | 4.14E-20 |
| SPOCK1 | 1.27E-24 | 0.575045 | 0.27 | 0.221 | 4.63E-20 |
| DDX21 | 1.44E-24 | -0.27658 | 0.298 | 0.419 | 5.25E-20 |
| SOX5 | 1.44E-24 | 0.495774 | 0.181 | 0.127 | 5.28E-20 |
| PHLPP1 | 1.63E-24 | 0.348563 | 0.109 | 0.064 | 5.95E-20 |
| ITM2A | 1.80E-24 | 0.328534 | 0.482 | 0.447 | 6.57E-20 |
| CLMP | 1.90E-24 | 0.503426 | 0.293 | 0.243 | 6.93E-20 |
| NIN | 2.00E-24 | 0.472097 | 0.181 | 0.127 | 7.30E-20 |
| CDK5RAP2 | 2.37E-24 | 0.391131 | 0.136 | 0.088 | 8.66E-20 |
| MSH3 | 2.52E-24 | 0.389135 | 0.106 | 0.063 | 9.22E-20 |
| HDAC4 | 2.69E-24 | 0.349103 | 0.113 | 0.068 | 9.85E-20 |

| | | | | | |
|---|---|---|---|---|---|
| PIP5K1A | 3.45E-24 | 0.408328 | 0.122 | 0.076 | 1.26E-19 |
| FOXJ3 | 3.53E-24 | 0.447186 | 0.117 | 0.072 | 1.29E-19 |
| ZFHX3 | 3.82E-24 | 0.454051 | 0.247 | 0.187 | 1.40E-19 |
| MID1IP1 | 4.39E-24 | -0.25892 | 0.076 | 0.138 | 1.60E-19 |
| LYST | 5.26E-24 | 0.441469 | 0.17 | 0.118 | 1.92E-19 |
| NBAS | 5.39E-24 | 0.395181 | 0.126 | 0.079 | 1.97E-19 |
| APC | 6.93E-24 | 0.448194 | 0.153 | 0.103 | 2.53E-19 |
| ANKRD11 | 7.89E-24 | 0.501697 | 0.35 | 0.312 | 2.89E-19 |
| RASA1 | 9.42E-24 | 0.397626 | 0.133 | 0.085 | 3.44E-19 |
| BMPER | 1.06E-23 | 0.524707 | 0.121 | 0.077 | 3.88E-19 |
| TOGARAM | 1.13E-23 | 0.407217 | 0.124 | 0.077 | 4.15E-19 |
| RFTN2 | 1.22E-23 | 0.481273 | 0.106 | 0.064 | 4.44E-19 |
| TRHDE | 1.42E-23 | 0.5016 | 0.116 | 0.072 | 5.21E-19 |
| CCDC84 | 1.53E-23 | 0.397103 | 0.121 | 0.076 | 5.58E-19 |
| PBRM1 | 1.62E-23 | 0.482255 | 0.221 | 0.169 | 5.93E-19 |
| DCAF6 | 1.64E-23 | 0.499257 | 0.169 | 0.119 | 6.01E-19 |
| TPST1 | 1.78E-23 | 0.573839 | 0.251 | 0.198 | 6.52E-19 |
| SMARCC1 | 2.20E-23 | 0.381917 | 0.161 | 0.109 | 8.03E-19 |
| NPEPPS | 2.24E-23 | 0.450934 | 0.256 | 0.204 | 8.18E-19 |
| TRA2A | 2.28E-23 | 0.480774 | 0.282 | 0.234 | 8.32E-19 |
| SPAG17 | 2.36E-23 | 0.552327 | 0.114 | 0.069 | 8.62E-19 |
| ARID1B | 2.44E-23 | 0.558334 | 0.297 | 0.254 | 8.92E-19 |
| USP8 | 2.68E-23 | 0.481887 | 0.241 | 0.187 | 9.81E-19 |
| RBM26 | 3.22E-23 | 0.471426 | 0.172 | 0.122 | 1.18E-18 |
| FAM126B | 3.54E-23 | 0.377488 | 0.113 | 0.069 | 1.29E-18 |
| MAMDC2 | 3.63E-23 | 0.390219 | 0.213 | 0.154 | 1.33E-18 |
| NRP1 | 3.77E-23 | 0.608509 | 0.279 | 0.233 | 1.38E-18 |
| FBN1 | 4.22E-23 | -0.30275 | 0.694 | 0.613 | 1.54E-18 |
| RNF149 | 4.25E-23 | 0.486208 | 0.29 | 0.241 | 1.55E-18 |
| LSAMP | 5.70E-23 | 0.403235 | 0.161 | 0.11 | 2.08E-18 |
| CCDC186 | 5.96E-23 | 0.480539 | 0.268 | 0.215 | 2.18E-18 |
| CDR2 | 7.05E-23 | 0.563551 | 0.144 | 0.097 | 2.58E-18 |
| DNMBP | 8.59E-23 | 0.425952 | 0.125 | 0.079 | 3.14E-18 |
| PDZD2 | 8.96E-23 | 0.2957 | 0.131 | 0.082 | 3.27E-18 |
| ELMO1 | 9.09E-23 | 0.358657 | 0.124 | 0.078 | 3.32E-18 |
| TBC1D22A | 1.26E-22 | 0.366339 | 0.131 | 0.084 | 4.59E-18 |
| ELP4 | 1.27E-22 | 0.438851 | 0.104 | 0.062 | 4.64E-18 |
| RIC1 | 1.52E-22 | 0.392346 | 0.108 | 0.066 | 5.56E-18 |
| RPRD2 | 1.76E-22 | 0.391671 | 0.137 | 0.09 | 6.44E-18 |
| ZNF148 | 1.83E-22 | 0.527187 | 0.209 | 0.159 | 6.70E-18 |
| CFLAR | 1.94E-22 | 0.468449 | 0.335 | 0.293 | 7.11E-18 |
| MTMR2 | 2.02E-22 | 0.374321 | 0.113 | 0.07 | 7.37E-18 |
| PPM1B | 2.16E-22 | 0.511486 | 0.173 | 0.125 | 7.90E-18 |
| SFRP1 | 2.31E-22 | -0.31955 | 0.198 | 0.28 | 8.44E-18 |
| TAOK1 | 2.78E-22 | 0.568591 | 0.229 | 0.175 | 1.02E-17 |
| DLG1 | 2.81E-22 | 0.421782 | 0.155 | 0.106 | 1.03E-17 |
| HECTD1 | 3.11E-22 | 0.452606 | 0.255 | 0.204 | 1.13E-17 |

| | | | | | |
|---|---|---|---|---|---|
| IPO9 | 3.42E-22 | 0.407911 | 0.113 | 0.07 | 1.25E-17 |
| RABEP1 | 3.92E-22 | 0.522238 | 0.216 | 0.164 | 1.43E-17 |
| FARP2 | 4.64E-22 | 0.371489 | 0.133 | 0.086 | 1.69E-17 |
| GABPB2 | 6.14E-22 | 0.437489 | 0.133 | 0.088 | 2.25E-17 |
| PTPRK | 6.88E-22 | 0.398514 | 0.115 | 0.072 | 2.51E-17 |
| SEMA3D | 7.40E-22 | 0.560575 | 0.174 | 0.126 | 2.70E-17 |
| NUP153 | 8.73E-22 | 0.511018 | 0.179 | 0.131 | 3.19E-17 |
| QRICH1 | 9.50E-22 | 0.385341 | 0.12 | 0.077 | 3.47E-17 |
| RPL31 | 1.34E-21 | -0.46181 | 0.631 | 0.753 | 4.91E-17 |
| FAR1 | 1.41E-21 | 0.482814 | 0.153 | 0.108 | 5.14E-17 |
| AGPAT4 | 1.61E-21 | 0.394846 | 0.101 | 0.062 | 5.89E-17 |
| MAPK14 | 1.63E-21 | 0.446903 | 0.145 | 0.098 | 5.94E-17 |
| LATS1 | 1.77E-21 | 0.400978 | 0.125 | 0.082 | 6.48E-17 |
| ZMIZ1 | 1.85E-21 | 0.422855 | 0.155 | 0.108 | 6.77E-17 |
| SLC30A9 | 2.20E-21 | 0.434168 | 0.138 | 0.093 | 8.03E-17 |
| NOX4 | 2.35E-21 | 0.342659 | 0.1 | 0.059 | 8.58E-17 |
| CREBRF | 2.51E-21 | 0.409577 | 0.255 | 0.205 | 9.19E-17 |
| HERC2 | 2.55E-21 | 0.415094 | 0.141 | 0.096 | 9.32E-17 |
| GLYR1 | 2.68E-21 | 0.416764 | 0.139 | 0.093 | 9.80E-17 |
| CPED1 | 2.83E-21 | 0.518155 | 0.232 | 0.185 | 1.03E-16 |
| MIB1 | 3.00E-21 | 0.497545 | 0.15 | 0.104 | 1.10E-16 |
| RASA2 | 3.16E-21 | 0.414345 | 0.142 | 0.096 | 1.15E-16 |
| LARP4B | 3.35E-21 | 0.346291 | 0.113 | 0.071 | 1.22E-16 |
| SCAF8 | 4.30E-21 | 0.403305 | 0.122 | 0.079 | 1.57E-16 |
| CPB1 | 5.30E-21 | -0.33864 | 0.056 | 0.107 | 1.94E-16 |
| NUAK1 | 6.93E-21 | 0.382334 | 0.103 | 0.063 | 2.53E-16 |
| LONP2 | 7.77E-21 | 0.466762 | 0.207 | 0.159 | 2.84E-16 |
| APPL2 | 8.09E-21 | 0.477207 | 0.178 | 0.129 | 2.96E-16 |
| JUN | 8.73E-21 | -0.81285 | 0.716 | 0.691 | 3.19E-16 |
| ZFYVE9 | 9.13E-21 | 0.43072 | 0.127 | 0.084 | 3.34E-16 |
| RUSC2 | 9.97E-21 | 0.399731 | 0.1 | 0.06 | 3.64E-16 |
| PLEKHA2 | 1.01E-20 | 0.41391 | 0.115 | 0.074 | 3.67E-16 |
| CSPP1 | 1.07E-20 | 0.402008 | 0.14 | 0.094 | 3.90E-16 |
| CDK11A | 1.13E-20 | 0.375856 | 0.137 | 0.092 | 4.15E-16 |
| SETX | 1.41E-20 | 0.44497 | 0.213 | 0.163 | 5.14E-16 |
| IGF1R | 1.57E-20 | 0.394364 | 0.142 | 0.097 | 5.74E-16 |
| GAPVD1 | 1.73E-20 | 0.444238 | 0.153 | 0.107 | 6.33E-16 |
| TRIM56 | 1.77E-20 | 0.513574 | 0.191 | 0.144 | 6.47E-16 |
| NEK1 | 2.00E-20 | 0.412559 | 0.159 | 0.112 | 7.31E-16 |
| SPATA6 | 2.24E-20 | 0.56343 | 0.198 | 0.153 | 8.20E-16 |
| GAB1 | 2.36E-20 | 0.374456 | 0.113 | 0.072 | 8.63E-16 |
| VRK2 | 2.50E-20 | 0.384088 | 0.104 | 0.064 | 9.14E-16 |
| DPYSL2 | 3.00E-20 | 0.384257 | 0.382 | 0.356 | 1.10E-15 |
| CDK12 | 3.08E-20 | 0.395918 | 0.201 | 0.153 | 1.13E-15 |
| CEP126 | 3.59E-20 | 0.453822 | 0.202 | 0.153 | 1.31E-15 |
| PAFAH1B1 | 4.01E-20 | 0.513667 | 0.306 | 0.268 | 1.47E-15 |
| MCPH1 | 4.17E-20 | 0.339474 | 0.121 | 0.079 | 1.52E-15 |

| | | | | | |
|---|---|---|---|---|---|
| UQCRQ | 4.92E-20 | -0.27158 | 0.205 | 0.301 | 1.80E-15 |
| GPX3 | 5.05E-20 | -0.38331 | 0.531 | 0.622 | 1.84E-15 |
| HELZ | 5.35E-20 | 0.43487 | 0.155 | 0.11 | 1.95E-15 |
| FAM193A | 5.50E-20 | 0.370037 | 0.125 | 0.082 | 2.01E-15 |
| SREK1 | 5.53E-20 | 0.506139 | 0.315 | 0.28 | 2.02E-15 |
| NR2C1 | 5.75E-20 | 0.42311 | 0.128 | 0.086 | 2.10E-15 |
| CEP95 | 6.94E-20 | 0.414394 | 0.14 | 0.096 | 2.54E-15 |
| RNF111 | 8.35E-20 | 0.319107 | 0.153 | 0.105 | 3.05E-15 |
| PER2 | 9.50E-20 | 0.578575 | 0.253 | 0.215 | 3.47E-15 |
| KMT5B | 1.14E-19 | 0.511293 | 0.179 | 0.133 | 4.16E-15 |
| EFCAB13 | 1.14E-19 | 0.317888 | 0.104 | 0.064 | 4.17E-15 |
| RFTN1 | 1.30E-19 | 0.471913 | 0.218 | 0.172 | 4.75E-15 |
| OSR2 | 1.56E-19 | -0.29023 | 0.145 | 0.22 | 5.70E-15 |
| MARK3 | 1.77E-19 | 0.459752 | 0.226 | 0.18 | 6.48E-15 |
| COL5A3 | 2.38E-19 | 0.475533 | 0.13 | 0.089 | 8.72E-15 |
| FBXW7 | 2.71E-19 | 0.411133 | 0.161 | 0.116 | 9.91E-15 |
| FBLN1 | 2.87E-19 | -0.36491 | 0.589 | 0.638 | 1.05E-14 |
| KIAA2026 | 2.91E-19 | 0.443396 | 0.173 | 0.127 | 1.06E-14 |
| RUFY3 | 3.00E-19 | 0.472145 | 0.294 | 0.252 | 1.10E-14 |
| ERCC6L2 | 3.14E-19 | 0.368404 | 0.123 | 0.081 | 1.15E-14 |
| ALPK1 | 3.25E-19 | 0.373246 | 0.108 | 0.069 | 1.19E-14 |
| KMT2E | 3.31E-19 | 0.436597 | 0.329 | 0.293 | 1.21E-14 |
| BMPR1A | 3.94E-19 | 0.425261 | 0.145 | 0.101 | 1.44E-14 |
| LRRFIP2 | 4.63E-19 | 0.578774 | 0.222 | 0.18 | 1.69E-14 |
| ANO6 | 5.19E-19 | 0.551429 | 0.269 | 0.229 | 1.90E-14 |
| SLC25A12 | 6.49E-19 | 0.343203 | 0.117 | 0.076 | 2.37E-14 |
| ACIN1 | 6.76E-19 | 0.429027 | 0.194 | 0.148 | 2.47E-14 |
| AGL | 6.91E-19 | 0.40573 | 0.146 | 0.103 | 2.52E-14 |
| TET1 | 8.81E-19 | 0.344525 | 0.119 | 0.078 | 3.22E-14 |
| NUP107 | 9.08E-19 | 0.402379 | 0.129 | 0.088 | 3.32E-14 |
| DDIT3 | 1.10E-18 | -0.36147 | 0.06 | 0.109 | 4.02E-14 |
| SFSWAP | 1.14E-18 | 0.493849 | 0.209 | 0.165 | 4.15E-14 |
| BAZ1B | 1.19E-18 | 0.44295 | 0.237 | 0.191 | 4.37E-14 |
| ATP2B4 | 1.45E-18 | 0.42397 | 0.242 | 0.196 | 5.31E-14 |
| EVA1C | 1.47E-18 | 0.413405 | 0.119 | 0.079 | 5.39E-14 |
| MAPK10 | 1.94E-18 | 0.591654 | 0.169 | 0.127 | 7.09E-14 |
| SECISBP2L | 1.97E-18 | 0.413836 | 0.178 | 0.133 | 7.19E-14 |
| TSC1 | 2.07E-18 | 0.320841 | 0.101 | 0.064 | 7.57E-14 |
| NSMAF | 2.38E-18 | 0.401629 | 0.12 | 0.08 | 8.69E-14 |
| VPS13C | 2.42E-18 | 0.411855 | 0.254 | 0.209 | 8.86E-14 |
| ACER3 | 2.50E-18 | 0.361089 | 0.104 | 0.067 | 9.13E-14 |
| RIN2 | 2.74E-18 | 0.438717 | 0.132 | 0.092 | 1.00E-13 |
| MRPS6 | 3.40E-18 | -0.31482 | 0.166 | 0.242 | 1.24E-13 |
| DENND6A | 3.42E-18 | 0.412279 | 0.11 | 0.072 | 1.25E-13 |
| PIK3CA | 3.75E-18 | 0.436225 | 0.161 | 0.118 | 1.37E-13 |
| RNF19A | 3.89E-18 | 0.499501 | 0.317 | 0.283 | 1.42E-13 |
| PACS1 | 3.96E-18 | 0.404117 | 0.13 | 0.089 | 1.45E-13 |

| | | | | | |
|---|---|---|---|---|---|
| EMX2OS | 4.07E-18 | 0.335903 | 0.108 | 0.07 | 1.49E-13 |
| NUP160 | 4.25E-18 | 0.368216 | 0.109 | 0.071 | 1.55E-13 |
| USP3 | 4.51E-18 | 0.352217 | 0.139 | 0.097 | 1.65E-13 |
| CCNL1 | 5.17E-18 | -0.28607 | 0.418 | 0.519 | 1.89E-13 |
| KAT6B | 5.21E-18 | 0.476216 | 0.236 | 0.194 | 1.91E-13 |
| ERCC1 | 5.23E-18 | 0.47479 | 0.265 | 0.227 | 1.91E-13 |
| CPSF6 | 5.79E-18 | 0.47092 | 0.165 | 0.124 | 2.12E-13 |
| SLF2 | 7.12E-18 | 0.345056 | 0.115 | 0.076 | 2.60E-13 |
| FGD5 | 7.55E-18 | 0.292691 | 0.107 | 0.068 | 2.76E-13 |
| TRIM2 | 7.69E-18 | 0.392981 | 0.112 | 0.074 | 2.81E-13 |
| FRMD4A | 8.11E-18 | 0.2764 | 0.189 | 0.139 | 2.96E-13 |
| ARHGEF26 | 1.01E-17 | 0.378411 | 0.128 | 0.087 | 3.69E-13 |
| FAM13B | 1.04E-17 | 0.332876 | 0.128 | 0.087 | 3.80E-13 |
| TNPO3 | 1.07E-17 | 0.341963 | 0.11 | 0.072 | 3.90E-13 |
| UVRAG | 1.07E-17 | 0.464926 | 0.156 | 0.114 | 3.91E-13 |
| NAA35 | 1.29E-17 | 0.355965 | 0.109 | 0.071 | 4.71E-13 |
| WAC | 1.36E-17 | 0.47496 | 0.272 | 0.235 | 4.96E-13 |
| METTL7A | 1.77E-17 | 0.341559 | 0.42 | 0.407 | 6.47E-13 |
| BICD1 | 1.78E-17 | 0.26766 | 0.106 | 0.067 | 6.52E-13 |
| CRISPLD2 | 1.94E-17 | 0.405187 | 0.367 | 0.331 | 7.10E-13 |
| MRPL1 | 1.95E-17 | 0.368351 | 0.13 | 0.089 | 7.13E-13 |
| FGF10 | 2.08E-17 | 0.511624 | 0.153 | 0.112 | 7.62E-13 |
| CDK11B | 2.36E-17 | 0.320988 | 0.129 | 0.088 | 8.62E-13 |
| BRWD1 | 3.54E-17 | 0.434272 | 0.159 | 0.118 | 1.29E-12 |
| UPF3A | 3.60E-17 | 0.443051 | 0.248 | 0.209 | 1.32E-12 |
| MEF2C | 3.86E-17 | 0.379998 | 0.108 | 0.072 | 1.41E-12 |
| BCL2 | 3.96E-17 | 0.460944 | 0.122 | 0.084 | 1.45E-12 |
| RAB3GAP1 | 4.73E-17 | 0.384797 | 0.171 | 0.128 | 1.73E-12 |
| LINC01133 | 6.30E-17 | -0.40153 | 0.082 | 0.131 | 2.30E-12 |
| ATF7IP | 6.32E-17 | 0.381675 | 0.123 | 0.085 | 2.31E-12 |
| ZMYND8 | 7.40E-17 | 0.338689 | 0.123 | 0.084 | 2.71E-12 |
| KLF4 | 8.13E-17 | -0.38272 | 0.391 | 0.463 | 2.97E-12 |
| ZMYM2 | 8.69E-17 | 0.419004 | 0.172 | 0.131 | 3.17E-12 |
| TFDP2 | 9.27E-17 | 0.522874 | 0.241 | 0.203 | 3.39E-12 |
| DDIT4 | 9.44E-17 | -0.36336 | 0.076 | 0.126 | 3.45E-12 |
| TRIM33 | 9.82E-17 | 0.397709 | 0.142 | 0.102 | 3.59E-12 |
| GFRA1 | 9.83E-17 | 0.486766 | 0.186 | 0.146 | 3.59E-12 |
| CFD | 1.23E-16 | -0.5773 | 0.978 | 0.887 | 4.51E-12 |
| VPS53 | 1.27E-16 | 0.347272 | 0.141 | 0.1 | 4.65E-12 |
| CYP20A1 | 1.30E-16 | 0.457899 | 0.211 | 0.171 | 4.74E-12 |
| PTPN14 | 1.44E-16 | 0.373679 | 0.115 | 0.077 | 5.27E-12 |
| PCM1 | 1.45E-16 | 0.395946 | 0.317 | 0.282 | 5.32E-12 |
| TERF2 | 1.49E-16 | 0.314517 | 0.108 | 0.072 | 5.44E-12 |
| RNF19B | 1.58E-16 | -0.36188 | 0.062 | 0.105 | 5.78E-12 |
| WARS2 | 1.96E-16 | 0.407659 | 0.136 | 0.096 | 7.17E-12 |
| FRS2 | 2.03E-16 | 0.387826 | 0.116 | 0.079 | 7.42E-12 |
| ZDHHC20 | 2.35E-16 | 0.39332 | 0.106 | 0.07 | 8.60E-12 |

| | | | | | |
|---|---|---|---|---|---|
| GRB10 | 2.86E-16 | 0.333703 | 0.102 | 0.067 | 1.04E-11 |
| IWS1 | 3.07E-16 | 0.458852 | 0.216 | 0.174 | 1.12E-11 |
| LINC0123S | 3.10E-16 | 0.306003 | 0.128 | 0.087 | 1.13E-11 |
| DCTN4 | 3.26E-16 | 0.314023 | 0.104 | 0.068 | 1.19E-11 |
| TC2N | 4.18E-16 | 0.383838 | 0.119 | 0.082 | 1.53E-11 |
| PHACTR4 | 5.29E-16 | 0.344019 | 0.126 | 0.088 | 1.93E-11 |
| ENAH | 5.31E-16 | 0.49254 | 0.212 | 0.173 | 1.94E-11 |
| ZNF704 | 5.45E-16 | 0.460545 | 0.182 | 0.142 | 1.99E-11 |
| AOX1 | 5.54E-16 | 0.43002 | 0.241 | 0.201 | 2.03E-11 |
| SMCHD1 | 5.71E-16 | 0.498668 | 0.215 | 0.177 | 2.09E-11 |
| SLC20A2 | 5.93E-16 | 0.339287 | 0.113 | 0.077 | 2.17E-11 |
| MFSD14C | 6.15E-16 | 0.375612 | 0.118 | 0.081 | 2.25E-11 |
| COL12A1 | 6.47E-16 | 0.449859 | 0.383 | 0.372 | 2.36E-11 |
| ADAM33 | 6.51E-16 | 0.49569 | 0.218 | 0.182 | 2.38E-11 |
| SLC25A43 | 6.56E-16 | 0.424993 | 0.105 | 0.07 | 2.40E-11 |
| ZDHHC17 | 7.46E-16 | 0.385072 | 0.13 | 0.093 | 2.73E-11 |
| CLASRP | 7.90E-16 | 0.379982 | 0.106 | 0.071 | 2.89E-11 |
| HERC4 | 7.92E-16 | 0.393809 | 0.172 | 0.131 | 2.89E-11 |
| RIPK1 | 9.00E-16 | 0.350657 | 0.129 | 0.091 | 3.29E-11 |
| XRRA1 | 1.01E-15 | 0.317849 | 0.101 | 0.067 | 3.69E-11 |
| PTPN12 | 1.22E-15 | 0.441092 | 0.238 | 0.2 | 4.45E-11 |
| ATXN2 | 1.38E-15 | 0.461918 | 0.174 | 0.135 | 5.05E-11 |
| COLEC12 | 1.52E-15 | 0.536147 | 0.342 | 0.327 | 5.57E-11 |
| PPP2R3A | 1.85E-15 | 0.460288 | 0.125 | 0.088 | 6.75E-11 |
| FNBP4 | 2.60E-15 | 0.446141 | 0.28 | 0.248 | 9.52E-11 |
| JAK2 | 2.81E-15 | 0.348714 | 0.151 | 0.111 | 1.03E-10 |
| SLC7A6 | 2.82E-15 | 0.378362 | 0.111 | 0.076 | 1.03E-10 |
| PPFIBP1 | 2.95E-15 | 0.517793 | 0.241 | 0.207 | 1.08E-10 |
| BTAF1 | 3.11E-15 | 0.379896 | 0.131 | 0.094 | 1.14E-10 |
| ZNF532 | 3.20E-15 | 0.321785 | 0.104 | 0.07 | 1.17E-10 |
| SRPK2 | 3.52E-15 | 0.304633 | 0.229 | 0.187 | 1.29E-10 |
| NTM | 4.04E-15 | 0.281979 | 0.151 | 0.109 | 1.48E-10 |
| IP6K2 | 4.15E-15 | 0.352434 | 0.203 | 0.164 | 1.52E-10 |
| PREX2 | 4.30E-15 | 0.548813 | 0.124 | 0.089 | 1.57E-10 |
| CCDC14 | 4.41E-15 | 0.385205 | 0.147 | 0.109 | 1.61E-10 |
| DOCK11 | 4.57E-15 | 0.390204 | 0.175 | 0.134 | 1.67E-10 |
| PCNX1 | 4.77E-15 | 0.326017 | 0.132 | 0.094 | 1.74E-10 |
| MAP1B | 4.80E-15 | 0.33318 | 0.505 | 0.488 | 1.75E-10 |
| CHL1 | 5.99E-15 | 0.423332 | 0.276 | 0.243 | 2.19E-10 |
| NCOA3 | 7.45E-15 | 0.345755 | 0.148 | 0.11 | 2.72E-10 |
| FGF7 | 7.94E-15 | 0.398572 | 0.373 | 0.35 | 2.90E-10 |
| ERBIN | 8.78E-15 | 0.37942 | 0.162 | 0.123 | 3.21E-10 |
| IREB2 | 8.99E-15 | 0.36731 | 0.126 | 0.09 | 3.29E-10 |
| PRKAG2 | 1.01E-14 | 0.386009 | 0.16 | 0.122 | 3.70E-10 |
| ZNF131 | 1.19E-14 | 0.441188 | 0.183 | 0.145 | 4.34E-10 |
| DNM1 | 1.19E-14 | 0.450736 | 0.214 | 0.178 | 4.35E-10 |
| THOC1 | 1.21E-14 | 0.389969 | 0.131 | 0.094 | 4.43E-10 |

| | | | | | |
|---|---|---|---|---|---|
| HOOK3 | 1.26E-14 | 0.497308 | 0.267 | 0.236 | 4.60E-10 |
| DNM2 | 1.37E-14 | 0.358528 | 0.104 | 0.072 | 5.02E-10 |
| RIPOR3 | 1.38E-14 | 0.373314 | 0.111 | 0.077 | 5.03E-10 |
| IRAK3 | 1.42E-14 | 0.428539 | 0.192 | 0.153 | 5.18E-10 |
| DIP2A | 1.48E-14 | 0.306803 | 0.1 | 0.067 | 5.42E-10 |
| BCL6 | 1.51E-14 | 0.477372 | 0.224 | 0.189 | 5.52E-10 |
| DDHD2 | 1.53E-14 | 0.327532 | 0.104 | 0.071 | 5.58E-10 |
| RAF1 | 1.56E-14 | 0.427302 | 0.158 | 0.122 | 5.70E-10 |
| AUH | 1.58E-14 | 0.385028 | 0.117 | 0.083 | 5.79E-10 |
| ADNP | 1.63E-14 | 0.410719 | 0.134 | 0.099 | 5.98E-10 |
| ASPH | 1.78E-14 | 0.455632 | 0.388 | 0.383 | 6.50E-10 |
| GOLGA8A | 1.84E-14 | 0.362996 | 0.104 | 0.072 | 6.74E-10 |
| PRPF4B | 2.15E-14 | 0.439827 | 0.322 | 0.299 | 7.84E-10 |
| YAF2 | 2.32E-14 | 0.448717 | 0.142 | 0.106 | 8.48E-10 |
| HIBCH | 2.38E-14 | 0.45562 | 0.153 | 0.117 | 8.72E-10 |
| VGLL4 | 2.43E-14 | 0.50052 | 0.211 | 0.179 | 8.89E-10 |
| SMARCA1 | 2.46E-14 | 0.445029 | 0.244 | 0.208 | 8.99E-10 |
| PPFIA1 | 2.53E-14 | 0.409673 | 0.169 | 0.131 | 9.23E-10 |
| GOLGA2 | 2.58E-14 | 0.373588 | 0.268 | 0.235 | 9.45E-10 |
| U2SURP | 2.81E-14 | 0.437569 | 0.309 | 0.284 | 1.03E-09 |
| PROCR | 3.06E-14 | -0.35447 | 0.244 | 0.312 | 1.12E-09 |
| JPX | 3.24E-14 | 0.461334 | 0.24 | 0.206 | 1.18E-09 |
| CCNL2 | 3.46E-14 | 0.452547 | 0.205 | 0.171 | 1.26E-09 |
| PBX3 | 3.56E-14 | 0.345325 | 0.13 | 0.095 | 1.30E-09 |
| COPA | 3.70E-14 | 0.470127 | 0.244 | 0.214 | 1.35E-09 |
| THY1 | 3.82E-14 | -0.30466 | 0.384 | 0.468 | 1.40E-09 |
| WDR26 | 3.85E-14 | 0.331848 | 0.141 | 0.104 | 1.41E-09 |
| YTHDC2 | 3.86E-14 | 0.402785 | 0.116 | 0.082 | 1.41E-09 |
| CNTN4 | 4.12E-14 | 0.274238 | 0.105 | 0.071 | 1.51E-09 |
| PER1 | 4.49E-14 | 0.433227 | 0.233 | 0.2 | 1.64E-09 |
| PPIP5K2 | 5.19E-14 | 0.406823 | 0.149 | 0.113 | 1.90E-09 |
| CUL3 | 6.05E-14 | 0.513539 | 0.221 | 0.189 | 2.21E-09 |
| MICU1 | 6.24E-14 | 0.422212 | 0.18 | 0.144 | 2.28E-09 |
| AC007563. | 6.73E-14 | -0.35135 | 0.064 | 0.102 | 2.46E-09 |
| BCLAF1 | 7.64E-14 | 0.376097 | 0.376 | 0.367 | 2.79E-09 |
| ODF2L | 8.96E-14 | 0.462616 | 0.247 | 0.215 | 3.28E-09 |
| DDI2 | 9.18E-14 | 0.275835 | 0.101 | 0.069 | 3.36E-09 |
| C1RL | 9.41E-14 | 0.444143 | 0.165 | 0.13 | 3.44E-09 |
| MEG8 | 9.45E-14 | 0.472943 | 0.209 | 0.172 | 3.45E-09 |
| AHCTF1 | 9.84E-14 | 0.442507 | 0.18 | 0.144 | 3.60E-09 |
| KPNA6 | 1.00E-13 | 0.369188 | 0.141 | 0.106 | 3.65E-09 |
| PIK3C2A | 1.03E-13 | 0.373112 | 0.181 | 0.142 | 3.77E-09 |
| SF3B1 | 1.10E-13 | 0.450712 | 0.377 | 0.376 | 4.04E-09 |
| LIMA1 | 1.15E-13 | 0.251395 | 0.541 | 0.554 | 4.21E-09 |
| CHD8 | 1.29E-13 | 0.296752 | 0.118 | 0.084 | 4.71E-09 |
| RNF216 | 1.53E-13 | 0.401486 | 0.157 | 0.121 | 5.59E-09 |
| FARP1 | 1.56E-13 | 0.368906 | 0.144 | 0.108 | 5.69E-09 |

| | | | | | |
|---|---|---|---|---|---|
| NAV2 | 1.56E-13 | 0.333767 | 0.156 | 0.118 | 5.71E-09 |
| TMEM131 | 1.67E-13 | 0.344428 | 0.144 | 0.108 | 6.09E-09 |
| NAA25 | 1.79E-13 | 0.311009 | 0.1 | 0.069 | 6.54E-09 |
| RUFY2 | 2.17E-13 | 0.388914 | 0.145 | 0.11 | 7.94E-09 |
| GRK5 | 2.28E-13 | 0.357185 | 0.135 | 0.1 | 8.32E-09 |
| CUL4A | 2.41E-13 | 0.401739 | 0.145 | 0.111 | 8.81E-09 |
| PHTF2 | 2.97E-13 | 0.401564 | 0.12 | 0.087 | 1.09E-08 |
| STX12 | 3.07E-13 | 0.626554 | 0.279 | 0.26 | 1.12E-08 |
| PSMD5 | 3.07E-13 | 0.384349 | 0.101 | 0.07 | 1.12E-08 |
| WWC2 | 3.25E-13 | 0.397831 | 0.14 | 0.105 | 1.19E-08 |
| MAVS | 3.39E-13 | 0.304215 | 0.114 | 0.081 | 1.24E-08 |
| PRKAR2A | 3.91E-13 | 0.382267 | 0.181 | 0.145 | 1.43E-08 |
| WASHC4 | 4.36E-13 | 0.386005 | 0.133 | 0.099 | 1.60E-08 |
| MAP3K2 | 4.80E-13 | 0.431754 | 0.223 | 0.191 | 1.76E-08 |
| NBN | 5.75E-13 | 0.422097 | 0.15 | 0.116 | 2.10E-08 |
| HP1BP3 | 5.84E-13 | 0.372655 | 0.406 | 0.407 | 2.14E-08 |
| LAMB2 | 6.00E-13 | 0.362913 | 0.38 | 0.374 | 2.19E-08 |
| NR3C1 | 6.33E-13 | 0.420823 | 0.318 | 0.299 | 2.31E-08 |
| XIAP | 7.26E-13 | 0.3416 | 0.161 | 0.126 | 2.65E-08 |
| KAT6A | 7.44E-13 | 0.425342 | 0.178 | 0.144 | 2.72E-08 |
| TMEM245 | 8.77E-13 | 0.297617 | 0.107 | 0.075 | 3.20E-08 |
| CAB39 | 9.17E-13 | 0.394074 | 0.124 | 0.092 | 3.35E-08 |
| USF3 | 1.07E-12 | 0.369044 | 0.102 | 0.072 | 3.91E-08 |
| FRYL | 1.11E-12 | 0.305511 | 0.116 | 0.083 | 4.07E-08 |
| RSF1 | 1.16E-12 | 0.393734 | 0.324 | 0.298 | 4.23E-08 |
| ZFC3H1 | 1.20E-12 | 0.467497 | 0.223 | 0.192 | 4.40E-08 |
| LIFR | 1.32E-12 | 0.472219 | 0.213 | 0.18 | 4.82E-08 |
| PRSS23 | 1.74E-12 | -0.26027 | 0.257 | 0.328 | 6.34E-08 |
| PIP4K2A | 1.76E-12 | 0.339539 | 0.11 | 0.079 | 6.43E-08 |
| FOSB | 2.04E-12 | -0.29069 | 0.673 | 0.667 | 7.45E-08 |
| QKI | 2.34E-12 | 0.507489 | 0.321 | 0.309 | 8.55E-08 |
| GCC2 | 2.42E-12 | 0.469737 | 0.264 | 0.237 | 8.85E-08 |
| ARHGAP18 | 2.66E-12 | 0.509422 | 0.155 | 0.125 | 9.72E-08 |
| TTC37 | 2.81E-12 | 0.417547 | 0.2 | 0.167 | 1.03E-07 |
| PHKB | 2.84E-12 | 0.378643 | 0.138 | 0.106 | 1.04E-07 |
| FBXW11 | 3.73E-12 | 0.3452 | 0.117 | 0.085 | 1.36E-07 |
| RALGPS2 | 3.87E-12 | 0.414612 | 0.145 | 0.113 | 1.42E-07 |
| SRPX | 4.36E-12 | 0.290396 | 0.492 | 0.472 | 1.60E-07 |
| ACAP2 | 4.73E-12 | 0.462021 | 0.219 | 0.189 | 1.73E-07 |
| TPP2 | 4.75E-12 | 0.313866 | 0.127 | 0.094 | 1.74E-07 |
| ZNF106 | 4.83E-12 | 0.404259 | 0.285 | 0.262 | 1.76E-07 |
| KIF13B | 5.27E-12 | 0.327962 | 0.115 | 0.083 | 1.93E-07 |
| STAT5B | 6.48E-12 | 0.338751 | 0.124 | 0.092 | 2.37E-07 |
| MYO6 | 6.69E-12 | 0.312565 | 0.126 | 0.094 | 2.44E-07 |
| SLC5A3 | 7.50E-12 | 0.353009 | 0.169 | 0.135 | 2.74E-07 |
| SULF1 | 7.63E-12 | 0.454426 | 0.203 | 0.169 | 2.79E-07 |
| DCAF7 | 8.01E-12 | 0.3406 | 0.113 | 0.083 | 2.93E-07 |

| | | | | | |
|---|---|---|---|---|---|
| AFF4 | 8.49E-12 | 0.460056 | 0.27 | 0.248 | 3.10E-07 |
| TBC1D23 | 8.77E-12 | 0.383746 | 0.145 | 0.113 | 3.21E-07 |
| INO80D | 9.12E-12 | 0.388625 | 0.158 | 0.125 | 3.33E-07 |
| EHMT1 | 9.25E-12 | 0.411709 | 0.153 | 0.121 | 3.38E-07 |
| GNAQ | 9.64E-12 | 0.413389 | 0.189 | 0.157 | 3.52E-07 |
| IL13RA1 | 1.03E-11 | 0.420459 | 0.238 | 0.21 | 3.77E-07 |
| PODN | 1.11E-11 | 0.336191 | 0.408 | 0.411 | 4.07E-07 |
| WBP4 | 1.20E-11 | 0.445013 | 0.229 | 0.2 | 4.38E-07 |
| NMT2 | 1.23E-11 | 0.31544 | 0.108 | 0.078 | 4.51E-07 |
| TMEM168 | 1.28E-11 | 0.373131 | 0.104 | 0.075 | 4.68E-07 |
| DMXL1 | 1.29E-11 | 0.374127 | 0.127 | 0.096 | 4.70E-07 |
| KDM7A | 1.36E-11 | 0.293253 | 0.147 | 0.112 | 4.97E-07 |
| CEP85L | 1.57E-11 | 0.38311 | 0.127 | 0.097 | 5.74E-07 |
| TNNT3 | 1.71E-11 | 0.469941 | 0.22 | 0.194 | 6.24E-07 |
| PDE7A | 2.03E-11 | 0.29808 | 0.102 | 0.073 | 7.42E-07 |
| KPNB1 | 2.26E-11 | 0.357584 | 0.266 | 0.24 | 8.25E-07 |
| INO80 | 2.51E-11 | 0.336862 | 0.132 | 0.101 | 9.19E-07 |
| AQR | 2.60E-11 | 0.297904 | 0.1 | 0.071 | 9.50E-07 |
| NSRP1 | 2.64E-11 | 0.379247 | 0.243 | 0.214 | 9.66E-07 |
| ARHGEF12 | 3.27E-11 | 0.363543 | 0.168 | 0.136 | 1.19E-06 |
| PPP3CC | 3.51E-11 | 0.30111 | 0.115 | 0.085 | 1.28E-06 |
| ABCA5 | 3.53E-11 | 0.3654 | 0.102 | 0.074 | 1.29E-06 |
| TOP2B | 3.72E-11 | 0.383279 | 0.153 | 0.121 | 1.36E-06 |
| ADCY3 | 4.17E-11 | 0.379608 | 0.151 | 0.121 | 1.52E-06 |
| MOB3B | 4.19E-11 | 0.40635 | 0.114 | 0.086 | 1.53E-06 |
| LSM14A | 4.41E-11 | 0.407976 | 0.214 | 0.186 | 1.61E-06 |
| RBL2 | 5.19E-11 | 0.404998 | 0.157 | 0.127 | 1.90E-06 |
| XPC | 5.39E-11 | 0.300103 | 0.138 | 0.107 | 1.97E-06 |
| GATAD2B | 6.20E-11 | 0.323053 | 0.133 | 0.102 | 2.27E-06 |
| CC2D2A | 6.72E-11 | 0.341534 | 0.11 | 0.081 | 2.46E-06 |
| TLK1 | 7.38E-11 | 0.357371 | 0.166 | 0.136 | 2.70E-06 |
| CLIP1 | 7.76E-11 | 0.299458 | 0.329 | 0.304 | 2.84E-06 |
| SPATS2 | 7.97E-11 | 0.341144 | 0.125 | 0.096 | 2.91E-06 |
| LRP6 | 8.24E-11 | 0.376693 | 0.142 | 0.113 | 3.01E-06 |
| MEIS1 | 8.29E-11 | 0.393854 | 0.119 | 0.09 | 3.03E-06 |
| RANBP9 | 9.67E-11 | 0.295385 | 0.117 | 0.088 | 3.53E-06 |
| PPP3CB | 9.94E-11 | 0.466118 | 0.159 | 0.131 | 3.63E-06 |
| RCOR3 | 1.10E-10 | 0.286819 | 0.103 | 0.075 | 4.02E-06 |
| SEMA3C | 1.18E-10 | -0.5787 | 0.245 | 0.29 | 4.31E-06 |
| TNRC18 | 1.20E-10 | 0.313279 | 0.107 | 0.079 | 4.37E-06 |
| SMARCA2 | 1.25E-10 | 0.387458 | 0.26 | 0.237 | 4.55E-06 |
| SLC41A2 | 1.29E-10 | 0.284468 | 0.109 | 0.08 | 4.70E-06 |
| UBAP2 | 1.29E-10 | 0.307659 | 0.1 | 0.072 | 4.72E-06 |
| PPL | 1.46E-10 | 0.377297 | 0.224 | 0.198 | 5.32E-06 |
| HIVEP2 | 1.66E-10 | 0.270674 | 0.132 | 0.102 | 6.05E-06 |
| SMURF2 | 1.73E-10 | 0.326236 | 0.196 | 0.163 | 6.31E-06 |
| N4BP2L2 | 1.79E-10 | 0.381474 | 0.391 | 0.399 | 6.54E-06 |

| | | | | | |
|---|---|---|---|---|---|
| IGFBP7 | 1.90E-10 | -0.55932 | 0.509 | 0.513 | 6.94E-06 |
| SLCO3A1 | 2.00E-10 | 0.38196 | 0.173 | 0.143 | 7.32E-06 |
| ATM | 2.14E-10 | 0.312707 | 0.133 | 0.104 | 7.82E-06 |
| CD36 | 2.32E-10 | 0.288077 | 0.133 | 0.102 | 8.47E-06 |
| NCOR2 | 2.42E-10 | 0.287551 | 0.157 | 0.127 | 8.84E-06 |
| PHF3 | 2.80E-10 | 0.398559 | 0.265 | 0.241 | 1.02E-05 |
| PUM1 | 3.12E-10 | 0.388537 | 0.196 | 0.168 | 1.14E-05 |
| WWP1 | 3.26E-10 | 0.384323 | 0.151 | 0.123 | 1.19E-05 |
| NFE2L1 | 3.58E-10 | 0.353895 | 0.201 | 0.173 | 1.31E-05 |
| DYRK1A | 3.72E-10 | 0.328339 | 0.111 | 0.084 | 1.36E-05 |
| RPS6KA3 | 3.89E-10 | 0.411876 | 0.189 | 0.16 | 1.42E-05 |
| SUGP2 | 4.74E-10 | 0.381484 | 0.131 | 0.104 | 1.73E-05 |
| STX18 | 5.13E-10 | 0.284965 | 0.103 | 0.076 | 1.88E-05 |
| WDR11 | 6.25E-10 | 0.310082 | 0.102 | 0.076 | 2.29E-05 |
| EZH1 | 6.45E-10 | 0.373915 | 0.152 | 0.123 | 2.36E-05 |
| EIF2B3 | 6.48E-10 | 0.402365 | 0.146 | 0.117 | 2.37E-05 |
| EP300 | 6.69E-10 | 0.336873 | 0.145 | 0.117 | 2.45E-05 |
| C6orf62 | 6.81E-10 | 0.286299 | 0.174 | 0.144 | 2.49E-05 |
| PDE1A | 7.03E-10 | 0.332528 | 0.1 | 0.074 | 2.57E-05 |
| DNM1L | 7.85E-10 | 0.425041 | 0.187 | 0.16 | 2.87E-05 |
| FUT8 | 8.13E-10 | 0.388238 | 0.126 | 0.099 | 2.97E-05 |
| TRIP11 | 8.48E-10 | 0.443819 | 0.232 | 0.208 | 3.10E-05 |
| WASF3 | 9.02E-10 | 0.359638 | 0.134 | 0.106 | 3.30E-05 |
| ATP5E | 9.27E-10 | -0.35938 | 0.36 | 0.279 | 3.39E-05 |
| UHRF1BP1 | 9.40E-10 | 0.336426 | 0.156 | 0.127 | 3.44E-05 |
| KLF7 | 1.04E-09 | 0.336484 | 0.135 | 0.107 | 3.80E-05 |
| SEC63 | 1.08E-09 | 0.325526 | 0.304 | 0.283 | 3.93E-05 |
| KCTD12 | 1.11E-09 | 0.433249 | 0.154 | 0.128 | 4.05E-05 |
| CD2AP | 1.20E-09 | 0.353264 | 0.104 | 0.078 | 4.40E-05 |
| ENPP2 | 1.38E-09 | 0.400695 | 0.147 | 0.12 | 5.03E-05 |
| SHPRH | 1.57E-09 | 0.359807 | 0.13 | 0.103 | 5.75E-05 |
| SENP7 | 1.58E-09 | 0.323813 | 0.134 | 0.106 | 5.79E-05 |
| PKN2 | 2.07E-09 | 0.426231 | 0.211 | 0.187 | 7.57E-05 |
| NAV1 | 2.26E-09 | 0.352522 | 0.24 | 0.214 | 8.27E-05 |
| MIGA1 | 2.45E-09 | 0.360692 | 0.138 | 0.111 | 8.97E-05 |
| RNF168 | 2.66E-09 | 0.273769 | 0.131 | 0.103 | 9.73E-05 |
| IL6ST | 2.70E-09 | 0.293584 | 0.453 | 0.473 | 9.85E-05 |
| PTGFR | 2.85E-09 | 0.304328 | 0.122 | 0.095 | 0.000104 |
| MAP3K3 | 3.27E-09 | 0.2906 | 0.103 | 0.078 | 0.000119 |
| AGPS | 3.33E-09 | 0.402535 | 0.129 | 0.104 | 0.000122 |
| EIF2AK2 | 3.64E-09 | 0.375464 | 0.204 | 0.179 | 0.000133 |
| MAP4K5 | 3.77E-09 | 0.398107 | 0.192 | 0.167 | 0.000138 |
| PHLDB1 | 4.11E-09 | 0.39054 | 0.238 | 0.217 | 0.00015 |
| MEIS2 | 4.11E-09 | 0.562156 | 0.214 | 0.193 | 0.00015 |
| MED15 | 4.50E-09 | 0.260246 | 0.111 | 0.085 | 0.000165 |
| MIS18BP1 | 4.99E-09 | 0.398155 | 0.133 | 0.107 | 0.000182 |
| MLH3 | 5.23E-09 | 0.316698 | 0.11 | 0.085 | 0.000191 |

| | | | | | |
|---|---|---|---|---|---|
| MAP4K4 | 5.79E-09 | 0.295597 | 0.209 | 0.182 | 0.000212 |
| SERINC5 | 6.39E-09 | 0.277084 | 0.1 | 0.075 | 0.000234 |
| IBTK | 6.77E-09 | 0.348271 | 0.146 | 0.12 | 0.000247 |
| AMPH | 6.82E-09 | 0.329805 | 0.104 | 0.08 | 0.000249 |
| CUL1 | 9.29E-09 | 0.298356 | 0.133 | 0.107 | 0.00034 |
| PLA2G2A | 9.49E-09 | -0.52642 | 0.278 | 0.32 | 0.000347 |
| ASCC3 | 1.31E-08 | 0.348655 | 0.138 | 0.112 | 0.000478 |
| YAP1 | 1.32E-08 | 0.501438 | 0.167 | 0.144 | 0.000484 |
| AP1G1 | 1.45E-08 | 0.306493 | 0.109 | 0.085 | 0.00053 |
| MECP2 | 1.60E-08 | 0.392648 | 0.162 | 0.137 | 0.000585 |
| OPA1 | 1.61E-08 | 0.332312 | 0.126 | 0.1 | 0.000587 |
| ARHGAP2( | 1.68E-08 | 0.32723 | 0.101 | 0.078 | 0.000614 |
| TRIP12 | 1.69E-08 | 0.379997 | 0.191 | 0.167 | 0.000618 |
| ZNF518A | 1.85E-08 | 0.29271 | 0.109 | 0.085 | 0.000678 |
| PDGFRA | 1.95E-08 | 0.307981 | 0.411 | 0.422 | 0.000712 |
| FNDC3A | 2.02E-08 | 0.442065 | 0.212 | 0.19 | 0.000737 |
| CHD2 | 2.20E-08 | 0.420714 | 0.229 | 0.209 | 0.000805 |
| KMT2A | 2.20E-08 | 0.405087 | 0.258 | 0.242 | 0.000805 |
| SNX4 | 2.23E-08 | 0.333041 | 0.124 | 0.1 | 0.000816 |
| GLI3 | 2.47E-08 | 0.264574 | 0.133 | 0.108 | 0.000904 |
| LARP7 | 2.93E-08 | 0.346698 | 0.232 | 0.21 | 0.001072 |
| RCOR1 | 2.95E-08 | 0.370278 | 0.113 | 0.089 | 0.001079 |
| LRRK1 | 3.19E-08 | 0.283341 | 0.105 | 0.081 | 0.001165 |
| ATF6 | 3.26E-08 | 0.352207 | 0.172 | 0.147 | 0.001192 |
| ZFR | 4.27E-08 | 0.31393 | 0.23 | 0.208 | 0.001559 |
| ARNT | 4.31E-08 | 0.284701 | 0.107 | 0.084 | 0.001575 |
| MAX | 4.46E-08 | 0.295875 | 0.168 | 0.143 | 0.00163 |
| PHF20L1 | 5.06E-08 | 0.408814 | 0.171 | 0.148 | 0.001849 |
| CDADC1 | 5.17E-08 | 0.355783 | 0.112 | 0.089 | 0.001889 |
| KLHL24 | 5.53E-08 | 0.307414 | 0.128 | 0.103 | 0.002023 |
| IGF2R | 5.62E-08 | 0.280045 | 0.131 | 0.106 | 0.002054 |
| PTPN11 | 5.89E-08 | 0.331826 | 0.15 | 0.125 | 0.002153 |
| USO1 | 5.92E-08 | 0.366504 | 0.191 | 0.169 | 0.002165 |
| STAT6 | 6.07E-08 | 0.31686 | 0.159 | 0.135 | 0.002218 |
| USP15 | 6.59E-08 | 0.39526 | 0.218 | 0.199 | 0.002408 |
| ADAMTSL3 | 6.68E-08 | 0.427941 | 0.166 | 0.143 | 0.00244 |
| GPBP1L1 | 7.04E-08 | 0.368408 | 0.123 | 0.099 | 0.002574 |
| KIDINS220 | 7.75E-08 | 0.391905 | 0.255 | 0.24 | 0.002831 |
| ZKSCAN1 | 8.56E-08 | 0.304464 | 0.174 | 0.149 | 0.003129 |
| PPP2CB | 8.80E-08 | 0.483934 | 0.224 | 0.209 | 0.003215 |
| DICER1 | 8.87E-08 | 0.326263 | 0.135 | 0.111 | 0.00324 |
| TAF3 | 9.08E-08 | 0.325314 | 0.125 | 0.101 | 0.003319 |
| DSE | 9.11E-08 | 0.344532 | 0.264 | 0.245 | 0.003331 |
| ACSL1 | 1.01E-07 | 0.328117 | 0.111 | 0.089 | 0.003708 |
| CCDC102B | 1.03E-07 | 0.477305 | 0.115 | 0.093 | 0.00375 |
| UBE2K | 1.03E-07 | 0.381865 | 0.229 | 0.21 | 0.003756 |
| GSK3B | 1.11E-07 | 0.315363 | 0.111 | 0.088 | 0.004047 |

| | | | | | |
|---|---|---|---|---|---|
| AP2A2 | 1.29E-07 | 0.276446 | 0.117 | 0.094 | 0.004716 |
| PPP6C | 1.34E-07 | 0.317043 | 0.11 | 0.088 | 0.004899 |
| TLK2 | 1.36E-07 | 0.304469 | 0.121 | 0.098 | 0.004971 |
| LONRF1 | 1.40E-07 | 0.441924 | 0.16 | 0.139 | 0.005109 |
| PRKCA | 1.42E-07 | 0.368399 | 0.124 | 0.101 | 0.005207 |
| TPPP3 | 1.51E-07 | -0.278 | 0.263 | 0.317 | 0.005534 |
| RBBP6 | 1.55E-07 | 0.393371 | 0.28 | 0.272 | 0.005668 |
| CRY2 | 1.69E-07 | 0.307272 | 0.117 | 0.095 | 0.006185 |
| RIF1 | 1.82E-07 | 0.401173 | 0.235 | 0.217 | 0.006668 |
| 7-Mar | 2.01E-07 | 0.372065 | 0.18 | 0.16 | 0.007331 |
| SPG7 | 2.16E-07 | 0.327446 | 0.141 | 0.118 | 0.007877 |
| PSME4 | 2.31E-07 | 0.291724 | 0.14 | 0.116 | 0.008444 |
| SESTD1 | 2.45E-07 | 0.403244 | 0.259 | 0.248 | 0.008971 |
| TP53BP1 | 2.49E-07 | 0.28806 | 0.104 | 0.082 | 0.009089 |
| TMEM41B | 3.17E-07 | 0.316627 | 0.113 | 0.091 | 0.011578 |
| UBAP1 | 3.27E-07 | 0.374231 | 0.164 | 0.142 | 0.011963 |
| IPMK | 3.97E-07 | 0.30538 | 0.115 | 0.092 | 0.014502 |
| R3HDM2 | 3.98E-07 | 0.370572 | 0.165 | 0.144 | 0.01453 |
| TPR | 4.03E-07 | 0.291671 | 0.347 | 0.342 | 0.014733 |
| FABP4 | 4.22E-07 | -0.6231 | 0.168 | 0.202 | 0.015436 |
| SNX14 | 4.30E-07 | 0.316489 | 0.121 | 0.099 | 0.015717 |
| NSD1 | 4.32E-07 | 0.277362 | 0.157 | 0.133 | 0.015805 |
| RPL23 | 4.50E-07 | -0.25828 | 0.674 | 0.754 | 0.016432 |
| TDRD3 | 4.76E-07 | 0.267676 | 0.114 | 0.092 | 0.017385 |
| HNRNPLL | 4.80E-07 | 0.320979 | 0.1 | 0.079 | 0.017558 |
| AR | 5.00E-07 | 0.3977 | 0.138 | 0.118 | 0.018267 |
| FGFR1 | 5.03E-07 | 0.264615 | 0.451 | 0.486 | 0.018379 |
| XAF1 | 5.07E-07 | 0.276842 | 0.124 | 0.102 | 0.018531 |
| AEBP2 | 5.44E-07 | 0.274846 | 0.133 | 0.11 | 0.019893 |
| RHOB | 5.46E-07 | -0.28123 | 0.432 | 0.486 | 0.019957 |
| KPNA1 | 5.62E-07 | 0.265617 | 0.118 | 0.095 | 0.020537 |
| NFKB1 | 5.67E-07 | 0.255531 | 0.208 | 0.187 | 0.020719 |
| COL16A1 | 6.54E-07 | 0.34471 | 0.213 | 0.196 | 0.023907 |
| CHD6 | 6.83E-07 | 0.31249 | 0.156 | 0.135 | 0.024963 |
| VCL | 7.11E-07 | 0.419431 | 0.258 | 0.248 | 0.025985 |
| POLK | 7.32E-07 | 0.32509 | 0.144 | 0.122 | 0.026746 |
| PRPF40A | 7.61E-07 | 0.357483 | 0.267 | 0.256 | 0.027813 |
| OGT | 7.86E-07 | 0.374911 | 0.169 | 0.149 | 0.028738 |
| GALNT15 | 8.08E-07 | -0.30555 | 0.176 | 0.211 | 0.029515 |
| UBE3A | 8.37E-07 | 0.409633 | 0.217 | 0.202 | 0.030577 |
| NFATC2IP | 9.05E-07 | 0.298882 | 0.106 | 0.086 | 0.033063 |
| H6PD | 9.38E-07 | 0.338824 | 0.153 | 0.133 | 0.034272 |
| YME1L1 | 1.04E-06 | 0.29155 | 0.24 | 0.224 | 0.038054 |
| SECISBP2 | 1.10E-06 | 0.349298 | 0.151 | 0.131 | 0.040106 |
| SLC19A2 | 1.10E-06 | 0.389658 | 0.273 | 0.264 | 0.040375 |
| PAPSS2 | 1.23E-06 | 0.256712 | 0.165 | 0.141 | 0.044967 |
| FAM114A1 | 1.31E-06 | 0.387285 | 0.301 | 0.301 | 0.048039 |

| | | | | | |
|---|---|---|---|---|---|
| SENP2 | 1.51E-06 | 0.257636 | 0.1 | 0.079 | 0.055208 |
| EPB41L3 | 1.67E-06 | 0.276253 | 0.235 | 0.215 | 0.060904 |
| SPPL3 | 1.74E-06 | 0.322769 | 0.136 | 0.116 | 0.063755 |
| CHD3 | 1.86E-06 | 0.288945 | 0.132 | 0.111 | 0.068037 |
| NMT1 | 1.94E-06 | 0.250192 | 0.139 | 0.118 | 0.071009 |
| RABL6 | 2.02E-06 | 0.282291 | 0.116 | 0.096 | 0.073892 |
| PRKDC | 2.45E-06 | 0.392564 | 0.186 | 0.169 | 0.089585 |
| NMD3 | 2.47E-06 | 0.403523 | 0.116 | 0.097 | 0.090371 |
| CTDSPL2 | 2.57E-06 | 0.306298 | 0.129 | 0.108 | 0.093783 |
| RFC1 | 2.80E-06 | 0.252857 | 0.194 | 0.175 | 0.102359 |
| EXOC1 | 2.86E-06 | 0.290863 | 0.145 | 0.124 | 0.104688 |
| DCP1A | 3.27E-06 | 0.341011 | 0.136 | 0.118 | 0.119347 |
| OLFML2B | 3.32E-06 | 0.325164 | 0.132 | 0.114 | 0.121396 |
| CCDC91 | 3.41E-06 | 0.310197 | 0.13 | 0.11 | 0.124814 |
| TMEM87A | 3.57E-06 | 0.437339 | 0.264 | 0.261 | 0.130549 |
| HGSNAT | 3.58E-06 | 0.384108 | 0.16 | 0.143 | 0.130712 |
| CDC42BPB | 3.97E-06 | 0.25212 | 0.149 | 0.128 | 0.145112 |
| MTR | 4.29E-06 | 0.288555 | 0.13 | 0.11 | 0.156781 |
| USP9X | 4.33E-06 | 0.269582 | 0.129 | 0.109 | 0.158261 |
| BBX | 5.72E-06 | 0.319119 | 0.234 | 0.224 | 0.208917 |
| PPP4R3B | 5.73E-06 | 0.336194 | 0.158 | 0.14 | 0.209366 |
| CENPC | 5.80E-06 | 0.312948 | 0.188 | 0.171 | 0.211832 |
| LUC7L | 7.02E-06 | 0.379102 | 0.187 | 0.172 | 0.256466 |
| MTUS1 | 7.65E-06 | 0.272499 | 0.14 | 0.121 | 0.279667 |
| PPP2R5C | 7.73E-06 | 0.270976 | 0.149 | 0.131 | 0.282507 |
| CDC16 | 8.21E-06 | 0.295995 | 0.126 | 0.107 | 0.300007 |
| RSRC1 | 8.35E-06 | 0.322625 | 0.196 | 0.18 | 0.305327 |
| DCAF5 | 8.53E-06 | 0.276826 | 0.132 | 0.113 | 0.311612 |
| ST3GAL5 | 8.90E-06 | 0.282006 | 0.14 | 0.121 | 0.325135 |
| STK17B | 8.95E-06 | 0.379286 | 0.161 | 0.145 | 0.327014 |
| TMF1 | 9.52E-06 | 0.357045 | 0.223 | 0.21 | 0.348095 |
| FCHO2 | 1.01E-05 | 0.293358 | 0.126 | 0.107 | 0.368042 |
| GGNBP2 | 1.04E-05 | 0.352372 | 0.233 | 0.223 | 0.379976 |
| ARHGAP12 | 1.43E-05 | 0.260342 | 0.157 | 0.138 | 0.521684 |
| PTPRA | 1.63E-05 | 0.36539 | 0.273 | 0.27 | 0.595152 |
| NBR1 | 2.19E-05 | 0.307355 | 0.117 | 0.1 | 0.80026 |
| ABL1 | 2.39E-05 | 0.364438 | 0.202 | 0.189 | 0.874054 |
| DYNC1I2 | 2.43E-05 | 0.298466 | 0.273 | 0.27 | 0.887373 |
| PPIG | 2.55E-05 | 0.273286 | 0.322 | 0.325 | 0.93203 |
| LATS2 | 2.62E-05 | 0.251625 | 0.15 | 0.133 | 0.957602 |
| CLTC | 2.73E-05 | 0.288876 | 0.195 | 0.18 | 0.997091 |
| MAN2A1 | 2.79E-05 | 0.291421 | 0.122 | 0.104 | 1 |
| CRY1 | 3.00E-05 | 0.321206 | 0.112 | 0.094 | 1 |
| PAK3 | 3.03E-05 | 0.300476 | 0.109 | 0.091 | 1 |
| STAG2 | 3.14E-05 | 0.26345 | 0.197 | 0.182 | 1 |
| 6-Mar | 3.32E-05 | 0.352879 | 0.182 | 0.17 | 1 |
| MYO1B | 3.39E-05 | 0.314172 | 0.142 | 0.125 | 1 |

| | | | | | |
|---|---|---|---|---|---|
| MPHOSPH | 3.62E-05 | 0.322064 | 0.321 | 0.332 | 1 |
| RBFOX2 | 3.93E-05 | 0.257611 | 0.281 | 0.274 | 1 |
| KTN1 | 5.09E-05 | 0.292275 | 0.339 | 0.346 | 1 |
| REV1 | 5.50E-05 | 0.322589 | 0.158 | 0.143 | 1 |
| TTC14 | 6.32E-05 | 0.334496 | 0.17 | 0.157 | 1 |
| CTNNA1 | 7.26E-05 | 0.341367 | 0.276 | 0.276 | 1 |
| ATF2 | 7.72E-05 | 0.291917 | 0.104 | 0.088 | 1 |
| ILKAP | 7.95E-05 | 0.303684 | 0.104 | 0.089 | 1 |
| RBM28 | 9.53E-05 | 0.290418 | 0.114 | 0.098 | 1 |
| ZMYM4 | 0.000101 | 0.253674 | 0.147 | 0.131 | 1 |
| CTNNB1 | 0.000108 | 0.316004 | 0.252 | 0.248 | 1 |
| ZNHIT6 | 0.000108 | 0.351271 | 0.109 | 0.094 | 1 |
| PPP4R3A | 0.000109 | 0.254564 | 0.12 | 0.104 | 1 |
| NAA15 | 0.000113 | 0.309903 | 0.141 | 0.126 | 1 |
| FUBP1 | 0.000115 | 0.300798 | 0.143 | 0.128 | 1 |
| NEK7 | 0.000122 | 0.317815 | 0.101 | 0.086 | 1 |
| SMAD2 | 0.000126 | 0.309073 | 0.112 | 0.097 | 1 |
| TMOD3 | 0.000126 | 0.409063 | 0.252 | 0.25 | 1 |
| MAGT1 | 0.000127 | 0.285559 | 0.16 | 0.146 | 1 |
| OSMR | 0.000151 | 0.440887 | 0.251 | 0.245 | 1 |
| SRRM1 | 0.000167 | 0.260233 | 0.355 | 0.375 | 1 |
| LONRF3 | 0.000175 | 0.307267 | 0.106 | 0.091 | 1 |
| MTDH | 0.000179 | 0.2924 | 0.397 | 0.428 | 1 |
| GPX1 | 0.000196 | -0.42067 | 0.113 | 0.091 | 1 |
| PDIA3 | 0.000211 | 0.281817 | 0.451 | 0.495 | 1 |
| RAB1A | 0.000214 | 0.347491 | 0.308 | 0.325 | 1 |
| TCF25 | 0.000229 | 0.287377 | 0.311 | 0.323 | 1 |
| PCYOX1 | 0.000235 | 0.320377 | 0.274 | 0.279 | 1 |
| PLEKHA4 | 0.000241 | 0.289486 | 0.141 | 0.126 | 1 |
| ARRDC3 | 0.00025 | 0.274647 | 0.121 | 0.107 | 1 |
| NUP54 | 0.000256 | 0.361152 | 0.127 | 0.114 | 1 |
| AZI2 | 0.000265 | 0.366581 | 0.193 | 0.185 | 1 |
| SRP54 | 0.000271 | 0.339589 | 0.133 | 0.12 | 1 |
| PCSK7 | 0.000272 | 0.27372 | 0.118 | 0.104 | 1 |
| GABPB1-A | 0.000279 | 0.405861 | 0.217 | 0.212 | 1 |
| WAPL | 0.000289 | 0.284603 | 0.138 | 0.124 | 1 |
| LRRFIP1 | 0.000308 | 0.283056 | 0.301 | 0.304 | 1 |
| ERICH1 | 0.000311 | 0.288128 | 0.162 | 0.15 | 1 |
| ADGRE5 | 0.000328 | 0.28656 | 0.128 | 0.114 | 1 |
| BTBD7 | 0.00037 | 0.29752 | 0.149 | 0.135 | 1 |
| ACBD3 | 0.000379 | 0.386699 | 0.212 | 0.207 | 1 |
| CLIC4 | 0.000419 | 0.355102 | 0.259 | 0.26 | 1 |
| CDH11 | 0.000423 | 0.387816 | 0.164 | 0.155 | 1 |
| SAV1 | 0.000461 | 0.357732 | 0.202 | 0.196 | 1 |
| PRRC2B | 0.000467 | 0.295601 | 0.175 | 0.165 | 1 |
| ETFDH | 0.000579 | 0.278191 | 0.106 | 0.093 | 1 |
| LNPK | 0.000644 | 0.322155 | 0.122 | 0.11 | 1 |

| | | | | | |
|---|---|---|---|---|---|
| NIPSNAP2 | 0.000652 | 0.428001 | 0.144 | 0.134 | 1 |
| RECK | 0.000669 | 0.323688 | 0.197 | 0.19 | 1 |
| CAND1 | 0.000704 | 0.298775 | 0.116 | 0.103 | 1 |
| PHYKPL | 0.000712 | 0.369097 | 0.164 | 0.156 | 1 |
| ARMC8 | 0.000729 | 0.281864 | 0.106 | 0.093 | 1 |
| CLINT1 | 0.00073 | 0.340555 | 0.206 | 0.201 | 1 |
| OSBPL1A | 0.000732 | 0.381333 | 0.223 | 0.222 | 1 |
| CIR1 | 0.000754 | 0.298981 | 0.209 | 0.203 | 1 |
| TBL1XR1 | 0.000829 | 0.322132 | 0.197 | 0.192 | 1 |
| TNKS2 | 0.000836 | 0.285537 | 0.122 | 0.109 | 1 |
| SESN1 | 0.000844 | 0.298495 | 0.106 | 0.093 | 1 |
| GPATCH2L | 0.000845 | 0.268963 | 0.109 | 0.096 | 1 |
| CWC27 | 0.000846 | 0.251198 | 0.122 | 0.109 | 1 |
| DNAJC1 | 0.000962 | 0.440754 | 0.197 | 0.194 | 1 |
| SMC6 | 0.001058 | 0.337437 | 0.108 | 0.095 | 1 |
| CRYBG3 | 0.001181 | 0.328951 | 0.169 | 0.16 | 1 |
| EIF3A | 0.00119 | 0.257359 | 0.337 | 0.352 | 1 |
| TOP1 | 0.001206 | 0.288186 | 0.319 | 0.334 | 1 |
| CCPG1 | 0.001307 | 0.343103 | 0.277 | 0.289 | 1 |
| EPS8 | 0.001313 | 0.404312 | 0.26 | 0.269 | 1 |
| ZNF644 | 0.001316 | 0.330478 | 0.193 | 0.187 | 1 |
| SPART | 0.001446 | 0.295658 | 0.197 | 0.248 | 1 |
| YIPF4 | 0.001461 | 0.369344 | 0.179 | 0.173 | 1 |
| SERPINH1 | 0.001513 | 0.347479 | 0.265 | 0.275 | 1 |
| ZNF83 | 0.00175 | 0.35212 | 0.196 | 0.191 | 1 |
| DYNC1H1 | 0.001883 | 0.333722 | 0.247 | 0.249 | 1 |
| ZC3H7A | 0.001919 | 0.26728 | 0.118 | 0.106 | 1 |
| PXDN | 0.002031 | 0.296184 | 0.134 | 0.123 | 1 |
| FRA10AC1 | 0.002106 | 0.282044 | 0.136 | 0.126 | 1 |
| PDCD6IP | 0.002163 | 0.303325 | 0.182 | 0.175 | 1 |
| PPP2R5E | 0.002189 | 0.268421 | 0.148 | 0.138 | 1 |
| ACTR2 | 0.002222 | 0.312961 | 0.241 | 0.243 | 1 |
| SF3B2 | 0.002549 | 0.254884 | 0.275 | 0.282 | 1 |
| PHF14 | 0.002753 | 0.354571 | 0.196 | 0.195 | 1 |
| ANPEP | 0.0029 | 0.265971 | 0.148 | 0.139 | 1 |
| ATP5G2 | 0.003102 | -0.45293 | 0.154 | 0.131 | 1 |
| EWSR1 | 0.003398 | 0.356084 | 0.219 | 0.221 | 1 |
| ARCN1 | 0.00351 | 0.288116 | 0.151 | 0.144 | 1 |
| MAN1A2 | 0.003828 | 0.367485 | 0.179 | 0.173 | 1 |
| VCAN | 0.003899 | -0.45504 | 0.415 | 0.42 | 1 |
| PSMD1 | 0.004043 | 0.267461 | 0.153 | 0.145 | 1 |
| TANK | 0.004071 | 0.369206 | 0.195 | 0.192 | 1 |
| CSNK1A1 | 0.00441 | 0.282141 | 0.361 | 0.395 | 1 |
| MAN1C1 | 0.004539 | 0.260348 | 0.101 | 0.092 | 1 |
| PRPF38B | 0.004697 | 0.306923 | 0.224 | 0.224 | 1 |
| SBNO1 | 0.004701 | 0.258004 | 0.11 | 0.1 | 1 |
| ATP5L | 0.004928 | -0.43149 | 0.183 | 0.157 | 1 |

| | | | | | |
|---|---|---|---|---|---|
| MAPK1 | 0.005048 | 0.253824 | 0.142 | 0.134 | 1 |
| PSMA3-AS | 0.005718 | 0.292624 | 0.155 | 0.149 | 1 |
| SIL1 | 0.006022 | 0.278745 | 0.157 | 0.15 | 1 |
| BRD4 | 0.00653 | 0.289336 | 0.201 | 0.201 | 1 |
| STX8 | 0.00711 | 0.308678 | 0.195 | 0.194 | 1 |
| ATP5O | 0.007993 | -0.31693 | 0.109 | 0.093 | 1 |
| SP100 | 0.008397 | 0.298005 | 0.261 | 0.271 | 1 |
| CASC4 | 0.008556 | 0.297922 | 0.23 | 0.234 | 1 |
| RARS2 | 0.0088 | 0.269527 | 0.113 | 0.104 | 1 |
| CAMSAP2 | 0.009212 | 0.302962 | 0.148 | 0.14 | 1 |
| ZC3H6 | 0.009914 | 0.2672 | 0.121 | 0.112 | 1 |
| SMC3 | 0.010348 | 0.272021 | 0.21 | 0.209 | 1 |
| SNTB2 | 0.010502 | 0.324589 | 0.238 | 0.244 | 1 |
| KHDRBS3 | 0.010636 | 0.337702 | 0.139 | 0.133 | 1 |
| NRDC | 0.011293 | 0.304149 | 0.185 | 0.183 | 1 |
| ANAPC5 | 0.011365 | 0.313653 | 0.227 | 0.233 | 1 |
| YES1 | 0.011994 | 0.283161 | 0.135 | 0.128 | 1 |
| RAB6A | 0.012308 | 0.291263 | 0.161 | 0.156 | 1 |
| ZNF451 | 0.012664 | 0.28819 | 0.146 | 0.139 | 1 |
| SVIL | 0.014976 | 0.351375 | 0.236 | 0.243 | 1 |
| DHX15 | 0.015586 | 0.319094 | 0.14 | 0.135 | 1 |
| GNB1 | 0.016704 | 0.309107 | 0.237 | 0.245 | 1 |
| HSD17B12 | 0.016753 | 0.333485 | 0.177 | 0.176 | 1 |
| SNX1 | 0.017452 | 0.319547 | 0.126 | 0.12 | 1 |
| RDX | 0.019463 | 0.336525 | 0.214 | 0.219 | 1 |
| TBC1D2B | 0.019472 | 0.251834 | 0.104 | 0.097 | 1 |
| SLC25A33 | 0.019685 | 0.321343 | 0.12 | 0.144 | 1 |
| PRKCI | 0.020101 | 0.257405 | 0.103 | 0.095 | 1 |
| HSD17B4 | 0.023204 | 0.319997 | 0.134 | 0.13 | 1 |
| SRRM2 | 0.024344 | 0.255721 | 0.405 | 0.454 | 1 |
| IL15RA | 0.025481 | 0.261824 | 0.117 | 0.11 | 1 |
| MLF1 | 0.026639 | 0.517532 | 0.173 | 0.177 | 1 |
| DHX36 | 0.02686 | 0.26649 | 0.267 | 0.281 | 1 |
| RRAS2 | 0.029351 | 0.262814 | 0.09 | 0.107 | 1 |
| RAB28 | 0.036285 | 0.306812 | 0.102 | 0.096 | 1 |
| NCOA7 | 0.038282 | 0.261166 | 0.27 | 0.284 | 1 |
| ZEB2 | 0.041124 | 0.262925 | 0.296 | 0.316 | 1 |
| WIPI1 | 0.042203 | 0.294839 | 0.11 | 0.104 | 1 |
| KIAA0232 | 0.042809 | 0.308443 | 0.13 | 0.126 | 1 |
| SP110 | 0.04322 | 0.285132 | 0.121 | 0.116 | 1 |
| PRDM2 | 0.044866 | 0.312673 | 0.225 | 0.232 | 1 |
| STRN3 | 0.047389 | 0.319332 | 0.151 | 0.149 | 1 |
| SPPL2A | 0.054805 | 0.318151 | 0.21 | 0.217 | 1 |
| XPO1 | 0.063196 | 0.275226 | 0.128 | 0.126 | 1 |
| CHIC2 | 0.065128 | 0.330807 | 0.135 | 0.16 | 1 |
| SLC35F5 | 0.077791 | 0.290783 | 0.107 | 0.103 | 1 |
| TRIP10 | 0.080984 | 0.252929 | 0.249 | 0.267 | 1 |

| | | | | | |
|---|---|---|---|---|---|
| SEC23A | 0.081668 | 0.356689 | 0.15 | 0.152 | 1 |
| ADAM17 | 0.09547 | 0.35957 | 0.193 | 0.201 | 1 |
| NDUFS1 | 0.096883 | 0.25712 | 0.126 | 0.124 | 1 |
| WDR41 | 0.104299 | 0.266292 | 0.105 | 0.102 | 1 |
| PLSCR4 | 0.106997 | 0.375496 | 0.213 | 0.227 | 1 |
| EPC1 | 0.113907 | 0.268854 | 0.217 | 0.228 | 1 |
| SPECC1 | 0.120784 | 0.273889 | 0.1 | 0.097 | 1 |
| MINDY2 | 0.124652 | 0.330496 | 0.139 | 0.139 | 1 |
| MORC4 | 0.131833 | 0.290041 | 0.122 | 0.121 | 1 |
| CCSER2 | 0.137522 | 0.300049 | 0.177 | 0.183 | 1 |
| NAA50 | 0.147092 | 0.356787 | 0.178 | 0.184 | 1 |
| ANTXR2 | 0.153596 | 0.295179 | 0.22 | 0.231 | 1 |
| CD46 | 0.164499 | 0.2634 | 0.186 | 0.218 | 1 |
| ZNF326 | 0.169223 | 0.259313 | 0.119 | 0.118 | 1 |
| TMEM263 | 0.16965 | 0.338352 | 0.181 | 0.189 | 1 |
| SAMD4B | 0.180658 | 0.300215 | 0.16 | 0.165 | 1 |
| GPBP1 | 0.183176 | 0.306679 | 0.255 | 0.279 | 1 |
| RPN2 | 0.188562 | 0.26122 | 0.208 | 0.245 | 1 |
| UBE2E1 | 0.194986 | 0.316708 | 0.131 | 0.133 | 1 |
| SLK | 0.195652 | 0.262937 | 0.235 | 0.247 | 1 |
| VEGFC | 0.212303 | 0.301641 | 0.089 | 0.102 | 1 |
| ASXL1 | 0.215545 | 0.294501 | 0.163 | 0.188 | 1 |
| MFAP5 | 0.217594 | -0.31648 | 0.629 | 0.567 | 1 |
| PRMT2 | 0.226278 | 0.262878 | 0.194 | 0.203 | 1 |
| DERA | 0.234464 | 0.264298 | 0.115 | 0.131 | 1 |
| CSAD | 0.247862 | 0.287788 | 0.12 | 0.121 | 1 |
| SLC2A3 | 0.257317 | 0.261188 | 0.299 | 0.326 | 1 |
| AASS | 0.262463 | 0.297634 | 0.141 | 0.16 | 1 |
| GPATCH2 | 0.265213 | 0.267706 | 0.107 | 0.107 | 1 |
| ZC2HC1A | 0.289016 | 0.275565 | 0.138 | 0.157 | 1 |
| PARVA | 0.307892 | 0.259049 | 0.184 | 0.212 | 1 |
| ALDOA | 0.312798 | -0.38865 | 0.128 | 0.134 | 1 |
| MAF | 0.336668 | 0.273603 | 0.209 | 0.223 | 1 |
| SPAG9 | 0.357118 | 0.275609 | 0.293 | 0.325 | 1 |
| VPS36 | 0.374043 | 0.251418 | 0.14 | 0.144 | 1 |
| STX5 | 0.384455 | 0.324242 | 0.129 | 0.133 | 1 |
| ADD1 | 0.385558 | 0.258724 | 0.21 | 0.242 | 1 |
| UBE2H | 0.395805 | 0.298195 | 0.223 | 0.258 | 1 |
| ELL2 | 0.408988 | 0.263267 | 0.206 | 0.216 | 1 |
| JAK1 | 0.417823 | 0.283858 | 0.304 | 0.345 | 1 |
| EVL | 0.437752 | 0.262461 | 0.138 | 0.144 | 1 |
| LIMS1 | 0.44568 | 0.286734 | 0.259 | 0.285 | 1 |
| P4HA1 | 0.45032 | 0.28742 | 0.147 | 0.154 | 1 |
| SAFB | 0.485 | 0.254064 | 0.117 | 0.12 | 1 |
| SUCLG2 | 0.490666 | 0.279848 | 0.144 | 0.151 | 1 |
| KHDRBS1 | 0.544527 | 0.298892 | 0.258 | 0.289 | 1 |
| ENPEP | 0.565936 | 0.292757 | 0.099 | 0.101 | 1 |

| | | | | | |
|---|---|---|---|---|---|
| LDLR | 0.600449 | 0.31904 | 0.146 | 0.155 | 1 |
| NUTM2B-/ | 0.634172 | 0.284309 | 0.145 | 0.154 | 1 |
| SNRNP70 | 0.652179 | 0.342892 | 0.217 | 0.242 | 1 |
| UFL1 | 0.692284 | 0.255748 | 0.138 | 0.146 | 1 |
| TSC22D2 | 0.710721 | 0.273255 | 0.192 | 0.213 | 1 |
| SEC13 | 0.728433 | 0.255036 | 0.137 | 0.146 | 1 |
| ARFGAP3 | 0.753648 | 0.326327 | 0.202 | 0.223 | 1 |
| NRIP1 | 0.788261 | 0.274091 | 0.173 | 0.191 | 1 |
| SLC25A36 | 0.800613 | 0.255245 | 0.179 | 0.198 | 1 |
| NSD3 | 0.819496 | 0.329694 | 0.186 | 0.208 | 1 |
| ACSL3 | 0.832904 | 0.342435 | 0.138 | 0.148 | 1 |
| SMARCA5 | 0.835243 | 0.254056 | 0.169 | 0.182 | 1 |
| TBC1D15 | 0.845096 | 0.318068 | 0.133 | 0.141 | 1 |
| RNF130 | 0.89598 | 0.384893 | 0.209 | 0.237 | 1 |
| GLUD1 | 0.908154 | 0.258654 | 0.19 | 0.209 | 1 |
| AFDN | 0.91441 | 0.289854 | 0.129 | 0.139 | 1 |
| UBA2 | 0.935021 | 0.251454 | 0.161 | 0.176 | 1 |
| DYNC2LI1 | 0.936205 | 0.30955 | 0.13 | 0.141 | 1 |
| TTC39C | 0.945268 | 0.250409 | 0.098 | 0.102 | 1 |
| VGLL3 | 0.958425 | 0.311854 | 0.167 | 0.182 | 1 |
| TMEM43 | 0.974581 | 0.263457 | 0.226 | 0.254 | 1 |
