## Supplementary material for "Adipocytes regulate fibroblast function, and their loss contributes to fibroblast dysfunction in inflammatory diseases": Data file S6

**Data File S6 22hr Basal vs FCM+TGFB FGSEA**

| pathway | pval | padj | ES | NES | nMoreEvents | size | leadingEdge |
|---|---|---|---|---|---|---|---|
| GOBP_DEVELOPMENTAL_GROWTH | 0.0018622 | 0.1400576 | -0.4111521 | -1.4729489 | 0 | 231 | GDF15 GDF5 CD9 DPYSL2 CRABP2 HEG1 CXCL12 ARID5B KCNK2 GATA6 BMP4 CDKN1A ERCC2 GAP43 GAMT ERCC6 GSN EXT1 ATF5 APP BNC2 ADM CSF1 FTO CCN3 CPQ AKIRIN1 ADRB2 FKBP8 DDR2 FZD7 BMPR1A AFG3L2 [illegible] |
| GOBP_GROWTH | 0.00186916 | 0.1400576 | -0.3630934 | -1.3637647 | 0 | 376 | GDF15 FHL1 CAMK2D KAZALD1 GDF5 GAS6 JADE1 CD9 DPYSL2 CRABP2 HEG1 CXCL12 ARID5B KCNK2 GATA6 BMP4 IGFBP3 CCDC85B CDKN1A ERCC2 CREG1 GAP43 GAMT ERCC6 GSN EXT1 ATF5 EGFR APP BNC2 ADM BDKRB1 [illegible] |
| GOBP_REGULATION_OF_INTRINSIC_APOPTOTIC_SIGNALING_PATHWAY_IN_RE | 0.00186916 | 0.1400576 | -0.9726192 | -1.4660666 | 0 | 3 | ARHGEF2 BDKRB2 BAD |
| GOBP_REGULATION_OF_RELAXATION_OF_CARDIAC_MUSCLE | 0.00186916 | 0.1400576 | -0.9858081 | -1.4859468 | 0 | 3 | CAMK2D |
| GOBP_SKIN_DEVELOPMENT | 0.0018797 | 0.1400576 | -0.4188711 | -1.4383965 | 0 | 161 | KRT19 LDB2 GATA6 FST LGR5 COL1A1 FLG KRTAP2-3 ERCC2 KRT18 KRT16 EGFR KRTAP1-5 COL1A2 CTSL KAZN KRT34 ERRFI1 KRT14 FOSL2 GBA AQP3 EPHA2 COL5A3 ITGA3 CSTA KRT80 ABCB6 COL5A1 KRT33B LDB1 LAMA5 [illegible] |
| GOBP_CELL_MORPHOGENESIS | 0.00188324 | 0.1400576 | -0.3618325 | -1.3795003 | 0 | 439 | CDC42EP3 ARHGAP35 ATF4 CDH11 DPYSL2 ARHGEF2 CRABP2 HEG1 CXCL12 CDC42EP1 CDC42EP2 DOK5 FLOT1 AXL FEZ1 CAP2 GAP43 LIMS2 DAPK3 EXT1 EGFR FBLN1 APP DVL2 DLC1 CDC42EP4 FES GREM1 CCKAR HEXB CDH6 [illegible] |
| GOBP_ANIMAL_ORGAN_MORPHOGENESIS | 0.00188679 | 0.1400576 | -0.4092104 | -1.5564878 | 0 | 426 | FHL1 DAB2 FHL2 EFEMP1 CITED2 ATF4 HEG1 GREM2 ARID5B ALPK2 GATA6 BMP4 ADAMTS1 FST COL1A1 ACTA2 GLG1 CAV1 GAMT LIMS2 EXT1 ADM ACVR1 FGF5 CRIP1 DVL2 IFT57 DLC1 COL5A2 CSF1 ALDH1A3 AP2A1 AQP1 [illegible] |
| GOBP_REGULATION_OF_TRANSMEMBRANE_RECEPTOR_PROTEIN_SERINE_TH | 0.00188679 | 0.1400576 | -0.5047398 | -1.6647008 | 0 | 110 | GDF15 DAB2 GDF5 CITED2 GOT1 BMPER GREM2 BMP4 FST ACVR2A GLG1 CAV1 ACVR1 FLCN CCN3 ASPN GREM1 FKBP8 CAV2 CRIM1 BMPR1A FBN1 HSPA1A ITGA3 GDF10 INHBE BCL9L JAK2 EMILIN1 LEMD2 CTDSPL2 GPC3 [illegible] |
| GOMF_NERVE_GROWTH_FACTOR_BINDING | 0.00189036 | 0.1400576 | 0.99830053 | 1.33613606 | 0 | 1 | FURIN |
| GOBP_MUSCLE_STRUCTURE_DEVELOPMENT | 0.00189036 | 0.1400576 | -0.4082366 | -1.4745921 | 0 | 257 | GDF15 FHL1 DDIT3 FHL2 ITGA11 KRT19 CD9 HEG1 CAPN2 ARID5B ALPK2 GATA6 BMP4 IGFBP3 FLOT1 FLNC HMG20B CAV1 EHD2 GPER1 ADM ACVR1 CCN3 EEF2 HDAC9 GREM1 AKIRIN1 CAV2 ASS1 FZD7 EPAS1 GPCPD1 BMPR1A [illegible] |
| GOCC_CELL_SUBSTRATE_JUNCTION | 0.00189036 | 0.1400576 | -0.4771884 | -1.6920519 | 0 | 213 | FHL1 DAB2 FHL2 ITGA11 ITGB8 CSPG4 CD9 ARHGEF2 AJUBA CAPN2 CDC42EP1 ASAP3 LAYN FLOT1 FLNC CORO2B CAV1 LIMS2 AFAP1 ITGBL1 GSN EGFR HMGA1 ARL2 DLC1 CAPN5 FES AHNAK LIMA1 ARMC5 CAV2 ACTN4 [illegible] |
| GOCC_ANCHORING_JUNCTION | 0.00189394 | 0.1400576 | -0.4451083 | -1.6878732 | 0 | 412 | FHL1 DAB2 FHL2 CLDN11 ITGA11 ITGB8 CDH11 CSPG4 CD9 ARHGEF2 AJUBA HEG1 CAPN2 CDC42EP1 ASAP3 CCDC85B LAYN FLOT1 FLNC AMOTL2 CORO2B CAV1 LIMS2 CADM1 AFAP1 KRT18 ITGBL1 GSN EGFR APP HMGA1 ARL2 [illegible] |
| GOCC_INTERMEDIATE_FILAMENT | 0.00189394 | 0.1400576 | -0.6915422 | -2.0539732 | 0 | 63 | CLDN11 KRT19 IFFO1 FLG KRTAP2-3 LDLRAP1 KRT18 KRT16 GPER1 KRTAP1-5 KRT34 KRT14 KRT80 CLIP1 KRT33B DST INA KRT86 KRT15 KRT17 |
| GOBP_DORSAL_VENTRAL_PATTERN_FORMATION | 0.00189753 | 0.1400576 | -0.6581765 | -1.7303598 | 0 | 30 | DDIT3 BMP4 ACVR1 GREM1 FKBP8 IFT52 BMPR1A GLI1 GLI3 CHRD |
| GOBP_REGULATION_OF_CELLULAR_RESPONSE_TO_GROWTH_FACTOR_STIMU | 0.00190114 | 0.1400576 | -0.5013421 | -1.6603335 | 0 | 118 | DAB2 GDF5 CITED2 GOT1 BMPER GREM2 DOK5 BMP4 FST CREB3L1 GLG1 CAV1 ATP2B4 FLCN ASPN GREM1 FKBP8 CAV2 CRIM1 FBN1 HSPA1A ITGA3 BCL9L EPN2 EMILIN1 GPC1 CD63 DSTYK FGF2 LEMD2 CTDSPL2 GPC3 FGF1 [illegible] |
| GOBP_MONOCYTE_DIFFERENTIATION | 0.0019084 | 0.1400576 | -0.6905391 | -1.6851884 | 0 | 22 | BMP4 IFI16 JUN CSF1 IL34 CDK6 FES FOXP1 CD74 IRF7 CTNNBIP1 |
| GOBP_MYOBLAST_FUSION | 0.0019084 | 0.1400576 | -0.7406182 | -1.806913 | 0 | 22 | GDF15 CD9 CAPN2 FLOT1 EHD2 CDB1 ADAM12 DYRK1B EHD1 |
| GOBP_REGULATION_OF_RELAXATION_OF_MUSCLE | 0.00191205 | 0.1400576 | -0.9651337 | -1.6276878 | 0 | 5 | CAMK2D |
| GOBP_SYNCYTIUM_FORMATION | 0.00191571 | 0.1400576 | -0.6818714 | -1.7718166 | 0 | 29 | GDF15 CD9 CAPN2 FLOT1 EHD2 CDB1 ADAM12 CAMK1 DYRK1B EHD1 |
| GOMF_CALCIUM_ION_BINDING | 0.00191939 | 0.1400576 | -0.3872569 | -1.417863 | 0 | 286 | EFEMP1 GAS6 CD248 CDH11 ANXA11 HEG1 CAPN2 FLG HPCAL1 EHD2 GSN ANXA7 FBLN1 CRELD2 CBLB FKBP10 FBLN2 FKBP9 ASPN EPDR1 CDH6 CALM3 ACTN4 FBN1 EHD4 ANXA10 IDS ESYT1 EHD1 DCHS1 EFHC1 ACTN1 CDH13 [illegible] |
| GOCC_EXTERNAL_ENCAPSULATING_STRUCTURE | 0.00192308 | 0.1400576 | -0.4044194 | -1.4722737 | 0 | 278 | GDF15 ADAMTSL1 EFEMP1 KAZALD1 CD248 CSPG4 LGALS3 ANXA11 CCDC80 BMPER CXCL12 ADAMTS1 COL1A1 FLG ACAN GLG1 COL12A1 ANXA7 FBLN1 FGL2 COL1A2 CTSL FBLN2 HAPLN3 CCN3 LAMA4 ASPN GREM1 GPC4 [illegible] |
| GOCC_INTERMEDIATE_FILAMENT_CYTOSKELETON | 0.00192308 | 0.1400576 | -0.6032302 | -1.8579508 | 0 | 77 | CLDN11 KRT19 IFFO1 FLG KRTAP2-3 LDLRAP1 KRT18 KRT16 GPER1 KRTAP1-5 INSL6 KRT34 KRT14 KRT80 CLIP1 KRT33B DST INA KRT86 EXD2 KRT15 KRT17 DDX60 |
| GOBP_TRANSDIFFERENTIATION | 0.00193424 | 0.1400576 | -0.9903565 | -1.375868 | 0 | 2 | GOT1 |
| GOBP_CALCIUM_DEPENDENT_CELL_CELL_ADHESION_VIA_PLASMA_MEMBRA | 0.00193424 | 0.1400576 | -0.7238476 | -1.7901295 | 0 | 23 | CDH11 AJUBA CDH6 DCHS1 CDH13 ARVCF CDHR3 CDH23 |
| GOBP_INTRINSIC_APOPTOTIC_SIGNALING_PATHWAY_IN_RESPONSE_TO_END | 0.00194175 | 0.1400576 | -0.6738753 | -1.764571 | 0 | 31 | DDIT3 CHAC1 ATF4 CREB3L1 ERP29 BAX BOK HSPA1A GRINA |
| GOBP_EPITHELIUM_DEVELOPMENT | 0.00194553 | 0.1400576 | -0.3672357 | -1.4113765 | 0 | 500 | DAB2 KRT19 CITED2 ARHGAP35 ATF4 LGALS3 BMPER AJUBA HEG1 LDB2 GATA6 BMP4 FST LGR5 FLG CDKN1A KRTAP2-3 ACTA2 ERCC2 CAV1 KRT18 EXT1 KRT16 ANXA7 EGFR ADM ACVR1 DVL2 IFT57 KRTAP1-5 DLC1 CSF1 ALDH1A3 [illegible] |
| GOBP_SEX_DETERMINATION | 0.00195695 | 0.1400576 | -0.9417643 | -1.6520012 | 0 | 6 | CITED2 |
| GOBP_NEGATIVE_REGULATION_OF_SIGNALING | 0.00197239 | 0.1400576 | -0.3651245 | -1.412732 | 0 | 548 | GDF15 DDIT3 DAB2 FHL2 CHAC1 GAS6 JADE1 IGFBP6 ARHGAP35 LGALS3 DUSP14 ARHGEF2 BMPER AJUBA HEG1 CXCL12 GREM2 ALPK2 BDKRB2 BMP4 FST IGFBP3 DEPTOR CREB3L1 GLG1 CAV1 ADA EGFR FBLN1 GPER1 GLIS2 [illegible] |
| GOBP_TISSUE_REGENERATION | 0.0019802 | 0.1400576 | -0.6390224 | -1.7277684 | 0 | 35 | CD9 CDKN1A GAP43 GSN CCN3 CPQ AKIRIN1 FZD7 GPX1 CDB1 DAG1 FGF10 IGFBP1 |
| GOBP_ORGANONITROGEN_COMPOUND_BIOSYNTHETIC_PROCESS | 0.00198413 | 0.1400576 | 0.35309241 | 1.40802484 | 0 | 753 | HS3ST3B1 CTNNB1 CMPK1 GFPT2 FURIN B4GALT1 GSTO1 GLUL INHBA ACLY CTPS1 FTSJ1 GALNT15 DNAJC3 C1GALT1C1 IL33 HAS1 GAPDH B4GALT3 GOT2 ALAS1 FARSA ENO1 HARS1 ALG3 ACOT7 ALDOA DCN DBI EIF4A1 ADK [illegible] |
| GOCC_CHROMOSOME | 0.00198413 | 0.1400576 | 0.34571959 | 1.37010541 | 0 | 696 | CTNNB1 DCTN4 ARNTL HMGB1 FEN1 IL33 HNRNPA2B1 EXOSC10 FOXO3 ELOA FOXO1 EGR1 HDAC8 BMS1 DDX21 HELLS DLX5 BUB3 ELK3 EXOSC9 ASF1B CDCA5 DHX9 ADNP HAT1 HIF1A KLF7 DYNC1LI2 DSN1 BHLHE40 AURKB [illegible] |
| GOBP_NEGATIVE_REGULATION_OF_RESPONSE_TO_STIMULUS | 0.00198413 | 0.1400576 | -0.3682233 | -1.4382201 | 0 | 642 | GDF15 DDIT3 DAB2 FHL2 CHAC1 GAS6 JADE1 IGFBP6 ARHGAP35 LGALS3 CD9 DUSP14 ARHGEF2 BMPER AJUBA HEG1 CXCL12 GREM2 ALPK2 BDKRB2 BMP4 FST IGFBP3 DEPTOR CREB3L1 GLG1 CAV1 ERCC6 ADA IFI16 IL20RB [illegible] |
| GOBP_RESPONSE_TO_OXYGEN_CONTAINING_COMPOUND | 0.00198807 | 0.1400576 | 0.35887789 | 1.43215618 | 0 | 755 | [illegible] |
| GOBP_CELLULAR_RESPONSE_TO_OXYGEN_CONTAINING_COMPOUND | 0.00199601 | 0.1400576 | 0.38225138 | 1.48305151 | 0 | 520 | [illegible] |
| GOBP_PROTEIN_PHOSPHORYLATION | 0.00199601 | 0.1400576 | -0.3365045 | -1.3240903 | 0 | 730 | [illegible] |
| GOBP_BIOLOGICAL_ADHESION | 0.00199601 | 0.1400576 | -0.3406886 | -1.3391506 | 0 | 708 | [illegible] |
| GOMF_SIGNALING_RECEPTOR_BINDING | 0.002 | 0.1400576 | 0.34303363 | 1.35965431 | 0 | 711 | [illegible] |
| GOBP_PROTEIN_N_LINKED_GLYCOSYLATION | 0.00201207 | 0.1400576 | 0.77212263 | 2.08564559 | 0 | 35 | [illegible] |
| GOMF_RNA_BINDING | 0.00201207 | 0.1400576 | 0.38881634 | 1.52560357 | 0 | 632 | [illegible] |
| GOCC_CATALYTIC_COMPLEX | 0.00201207 | 0.1400576 | 0.37114231 | 1.45146511 | 0 | 605 | [illegible] |
| GOBP_REGULATION_OF_PHOSPHORUS_METABOLIC_PROCESS | 0.00201207 | 0.1400576 | -0.3308565 | -1.2973326 | 0 | 691 | [illegible] |
| GOBP_CARBOHYDRATE_DERIVATIVE_METABOLIC_PROCESS | 0.0020202 | 0.1400576 | 0.39175187 | 1.52425955 | 0 | 555 | [illegible] |
| GOBP_REGULATION_OF_CELL_DIFFERENTIATION | 0.0020202 | 0.1400576 | -0.3381672 | -1.3178162 | 0 | 640 | [illegible] |
| GOBP_DNA_REPLICATION_INDEPENDENT_NUCLEOSOME_ORGANIZATION | 0.0020284 | 0.1400576 | 0.63087686 | 1.72829475 | 0 | 39 | [illegible] |
| GOBP_CHROMOSOME_ORGANIZATION | 0.00203252 | 0.1400576 | 0.44712424 | 1.73928407 | 0 | 556 | [illegible] |
| GOBP_REGULATION_OF_CELL_CYCLE | 0.00203252 | 0.1400576 | 0.35703296 | 1.38857424 | 0 | 558 | [illegible] |
| GOBP_MITOTIC_SISTER_CHROMATID_SEGREGATION | 0.00203666 | 0.1400576 | 0.60175905 | 1.89435551 | 0 | 81 | [illegible] |
| GOBP_RIBOSOMAL_LARGE_SUBUNIT_BIOGENESIS | 0.00204082 | 0.1400576 | 0.86596245 | 1.84166626 | 0 | 13 | [illegible] |
| GOCC_CHAPERONIN_CONTAINING_T_COMPLEX | 0.00204499 | 0.1400576 | 0.93263536 | 1.76237161 | 0 | 8 | [illegible] |
| GOBP_REGULATION_OF_CHROMOSOME_SEGREGATION | 0.00204918 | 0.1400576 | 0.66167456 | 1.84905227 | 0 | 42 | [illegible] |
| GOMF_CXCR_CHEMOKINE_RECEPTOR_BINDING | 0.00204918 | 0.1400576 | 0.88394492 | 1.83866566 | 0 | 12 | [illegible] |
| GOBP_METAPHASE_ANAPHASE_TRANSITION_OF_CELL_CYCLE | 0.00204918 | 0.1400576 | 0.67165222 | 1.81389254 | 0 | 36 | [illegible] |
| GOMF_CHEMOKINE_ACTIVITY | 0.00205339 | 0.1400576 | 0.78817197 | 2.1126 | 0 | 33 | [illegible] |
| GOBP_ANTIMICROBIAL_HUMORAL_IMMUNE_RESPONSE_MEDIATED_BY_ANT | 0.00205339 | 0.1400576 | 0.79750573 | 2.09363241 | 0 | 31 | [illegible] |
| GOCC_CATALYTIC_STEP_2_SPLICEOSOME | 0.00205339 | 0.1400576 | 0.74785828 | 2.00200397 | 0 | 33 | [illegible] |
| GOBP_ANTIMICROBIAL_HUMORAL_RESPONSE | 0.00205761 | 0.1400576 | 0.68779693 | 1.9495456 | 0 | 50 | [illegible] |
| GOBP_CHROMOSOME_SEPARATION | 0.00205761 | 0.1400576 | 0.65268954 | 1.83708172 | 0 | 47 | [illegible] |
| [illegible] | [illegible] | [illegible] | [illegible] | [illegible] | [illegible] | [illegible] | [illegible] |
