## Supplementary material for "Adipocytes regulate fibroblast function, and their loss contributes to fibroblast dysfunction in inflammatory diseases": Data file S7

**Data File S7 22hr Basal vs FCM+TGFB FGSEA**

| pathway | pval | padj | ES | NES | nMoreEvents | size | leadingEdge |
|---|---|---|---|---|---|---|---|
| GOBP_DEVELOPMENTAL_GROWTH | 0.0018622 | 0.1400576 | -0.4111521 | -1.4729489 | 0 | 231 | [illegible] |
| GOBP_GROWTH | 0.00186916 | 0.1400576 | -0.3630934 | -1.3637647 | 0 | 376 | [illegible] |
| GOBP_REGULATION_OF_INTRINSIC_APOPTOTIC_SIGNALING_PATHWAY_IN_RE | 0.00186916 | 0.1400576 | -0.9726192 | -1.4660666 | 0 | 3 | ARHGEF2 BDKRB2 BAD |
| GOBP_REGULATION_OF_RELAXATION_OF_CARDIAC_MUSCLE | 0.00186916 | 0.1400576 | -0.9858081 | -1.4859468 | 0 | 3 | CAMK2D |
| GOBP_SKIN_DEVELOPMENT | 0.0018797 | 0.1400576 | -0.4188711 | -1.4383965 | 0 | 161 | [illegible] |
| GOBP_CELL_MORPHOGENESIS | 0.00188324 | 0.1400576 | -0.3618325 | -1.3795003 | 0 | 439 | [illegible] |
| GOBP_ANIMAL_ORGAN_MORPHOGENESIS | 0.00188679 | 0.1400576 | -0.4092104 | -1.5564878 | 0 | 428 | [illegible] |
| GOBP_REGULATION_OF_TRANSMEMBRANE_RECEPTOR_PROTEIN_SERINE_TH | 0.00188679 | 0.1400576 | -0.5047398 | -1.6647008 | 0 | 110 | [illegible] |
| GOMF_NERVE_GROWTH_FACTOR_BINDING | 0.00189036 | 0.1400576 | 0.99830153 | 1.33613606 | 0 | 1 | FURIN |
| GOBP_MUSCLE_STRUCTURE_DEVELOPMENT | 0.00189036 | 0.1400576 | -0.4082366 | -1.4745921 | 0 | 257 | [illegible] |
| GOCC_CELL_SUBSTRATE_JUNCTION | 0.00189036 | 0.1400576 | -0.4771884 | -1.6920519 | 0 | 213 | [illegible] |
| GOCC_ANCHORING_JUNCTION | 0.00189394 | 0.1400576 | -0.4451083 | -1.6878732 | 0 | 412 | [illegible] |
| GOCC_INTERMEDIATE_FILAMENT | 0.00189394 | 0.1400576 | -0.6915422 | -2.0539732 | 0 | 63 | CLDN11 KRT19 IFFO1 FLG KRTAP2-3 LDLRAP1 KRT18 KRT16 GPER1 KRTAP1-5 KRT34 KRT14 KRT80 CLIP1 KRT33B DST INA KRT86 KRT15 KRT17 |
| GOBP_DORSAL_VENTRAL_PATTERN_FORMATION | 0.00189753 | 0.1400576 | -0.6581765 | -1.7303598 | 0 | 30 | DDIT3 BMP4 ACVR1 GREM1 FKBP8 IFT52 BMPR1A GLI1 GLI3 CHRD |
| GOBP_REGULATION_OF_CELLULAR_RESPONSE_TO_GROWTH_FACTOR_STIMU | 0.00190114 | 0.1400576 | -0.5013421 | -1.6603335 | 0 | 118 | [illegible] |
| GOBP_MONOCYTE_DIFFERENTIATION | 0.0019084 | 0.1400576 | -0.6905391 | -1.6851884 | 0 | 22 | BMP4 IFI16 JUN CSF1 IL34 CDK6 FES FOXP1 CD74 IRF7 CTNNBIP1 |
| GOBP_MYOBLAST_FUSION | 0.0019084 | 0.1400576 | -0.7406182 | -1.806913 | 0 | 22 | GDF15 CD9 CAPN2 FLOT1 EHD2 CD81 ADAM12 DYRK1B EHD1 |
| GOBP_REGULATION_OF_RELAXATION_OF_MUSCLE | 0.00191205 | 0.1400576 | -0.9651337 | -1.6276878 | 0 | 5 | CAMK2D |
| GOBP_SYNCYTIUM_FORMATION | 0.00191571 | 0.1400576 | -0.6818714 | -1.7718166 | 0 | 29 | GDF15 CD9 CAPN2 FLOT1 EHD2 CD81 ADAM12 CAMK1 DYRK1B EHD1 |
| GOMF_CALCIUM_ION_BINDING | 0.00191939 | 0.1400576 | -0.3872569 | -1.417863 | 0 | 286 | [illegible] |
| GOCC_EXTERNAL_ENCAPSULATING_STRUCTURE | 0.00192308 | 0.1400576 | -0.4046194 | -1.4722737 | 0 | 278 | [illegible] |
| GOCC_INTERMEDIATE_FILAMENT_CYTOSKELETON | 0.00192308 | 0.1400576 | -0.6032302 | -1.8579508 | 0 | 77 | [illegible] |
| GOBP_TRANSDIFFERENTIATION | 0.00193424 | 0.1400576 | -0.9903565 | -1.375868 | 0 | 2 | GOT1 |
| GOBP_CALCIUM_DEPENDENT_CELL_CELL_ADHESION_VIA_PLASMA_MEMBRA | 0.00193424 | 0.1400576 | -0.7238476 | -1.7901295 | 0 | 23 | CDH11 AJUBA CDH6 DCHS1 CDH13 ARVCF CDHR3 CDH23 |
| GOBP_INTRINSIC_APOPTOTIC_SIGNALING_PATHWAY_IN_RESPONSE_TO_END | 0.00194175 | 0.1400576 | -0.6738753 | -1.764571 | 0 | 31 | DDIT3 CHAC1 ATF4 CREB3L1 ERP29 BAX BOK HSPA1A GRINA |
| GOBP_EPITHELIUM_DEVELOPMENT | 0.00194553 | 0.1400576 | -0.3672357 | -1.4113765 | 0 | 500 | [illegible] |
| GOBP_SEX_DETERMINATION | 0.00195695 | 0.1400576 | -0.9417643 | -1.6520012 | 0 | 6 | CITED2 |
| GOBP_NEGATIVE_REGULATION_OF_SIGNALING | 0.00197239 | 0.1400576 | -0.3651245 | -1.412732 | 0 | 548 | [illegible] |
| GOBP_TISSUE_REGENERATION | 0.0019802 | 0.1400576 | -0.6390224 | -1.7277684 | 0 | 35 | CD9 CDKN1A GAP43 GSN CCN3 CPQ AKIRIN1 FZD7 GPX1 CD81 DAG1 FGF10 IGFBP5 |
| GOBP_ORGANONITROGEN_COMPOUND_BIOSYNTHETIC_PROCESS | 0.00198413 | 0.1400576 | 0.35309241 | 1.40802484 | 0 | 753 | [illegible] |
| GOCC_CHROMOSOME | 0.00198413 | 0.1400576 | 0.34571958 | 1.37010541 | 0 | 696 | [illegible] |
| GOBP_NEGATIVE_REGULATION_OF_RESPONSE_TO_STIMULUS | 0.00198413 | 0.1400576 | -0.3682233 | -1.4382201 | 0 | 642 | [illegible] |
| GOBP_RESPONSE_TO_OXYGEN_CONTAINING_COMPOUND | 0.00198807 | 0.1400576 | 0.35887789 | 1.43215618 | 0 | 755 | [illegible] |
| GOBP_CELLULAR_RESPONSE_TO_OXYGEN_CONTAINING_COMPOUND | 0.00199601 | 0.1400576 | 0.38225138 | 1.48305151 | 0 | 520 | [illegible] |
| GOBP_PROTEIN_PHOSPHORYLATION | 0.00199601 | 0.1400576 | -0.3365045 | -1.3240903 | 0 | 730 | [illegible] |
| GOBP_BIOLOGICAL_ADHESION | 0.00199601 | 0.1400576 | -0.3406886 | -1.3391506 | 0 | 708 | [illegible] |
| GOMF_SIGNALING_RECEPTOR_BINDING | 0.002 | 0.1400576 | 0.34303363 | 1.35965431 | 0 | 711 | [illegible] |
| GOBP_PROTEIN_N_LINKED_GLYCOSYLATION | 0.00201207 | 0.1400576 | 0.77212263 | 2.08564559 | 0 | 35 | [illegible] |
| GOMF_RNA_BINDING | 0.00201207 | 0.1400576 | 0.38881634 | 1.52560357 | 0 | 632 | [illegible] |
| GOCC_CATALYTIC_COMPLEX | 0.00201207 | 0.1400576 | 0.37114231 | 1.45146511 | 0 | 605 | [illegible] |
| GOBP_REGULATION_OF_PHOSPHORUS_METABOLIC_PROCESS | 0.00201207 | 0.1400576 | -0.3308565 | -1.2973326 | 0 | 691 | [illegible] |
| GOBP_CARBOHYDRATE_DERIVATIVE_METABOLIC_PROCESS | 0.0020202 | 0.1400576 | 0.39175187 | 1.52425955 | 0 | 555 | [illegible] |
| GOBP_REGULATION_OF_CELL_DIFFERENTIATION | 0.0020202 | 0.1400576 | -0.3381672 | -1.3178162 | 0 | 640 | [illegible] |
| GOBP_DNA_REPLICATION_INDEPENDENT_NUCLEOSOME_ORGANIZATION | 0.0020284 | 0.1400576 | 0.63087686 | 1.72829475 | 0 | 39 | [illegible] |
| GOBP_CHROMOSOME_ORGANIZATION | 0.00203252 | 0.1400576 | 0.44712424 | 1.73928807 | 0 | 556 | [illegible] |
| GOBP_REGULATION_OF_CELL_CYCLE | 0.00203252 | 0.1400576 | 0.35703296 | 1.38857424 | 0 | 558 | [illegible] |
| GOBP_MITOTIC_SISTER_CHROMATID_SEGREGATION | 0.00203666 | 0.1400576 | 0.60175905 | 1.89435551 | 0 | 81 | [illegible] |
| GOBP_RIBOSOMAL_LARGE_SUBUNIT_BIOGENESIS | 0.00204082 | 0.1400576 | 0.86596245 | 1.84166626 | 0 | 13 | [illegible] |
| GOCC_CHAPERONIN_CONTAINING_T_COMPLEX | 0.00204499 | 0.1400576 | 0.93263536 | 1.76237161 | 0 | 8 | CCT2 CCT6A CCT7 CCT8 CCT3 CCT4 CCT5 |
| GOBP_REGULATION_OF_CHROMOSOME_SEGREGATION | 0.00204918 | 0.1400576 | 0.66167456 | 1.84905227 | 0 | 42 | [illegible] |
| GOMF_CXCR_CHEMOKINE_RECEPTOR_BINDING | 0.00204918 | 0.1400576 | 0.88394492 | 1.83866566 | 0 | 12 | CXCL1 CXCL8 CXCL3 CXCL6 CXCL2 CXCL5 |
| GOBP_METAPHASE_ANAPHASE_TRANSITION_OF_CELL_CYCLE | 0.00204918 | 0.1400576 | 0.67165222 | 1.81349254 | 0 | 36 | [illegible] |
| GOMF_CHEMOKINE_ACTIVITY | 0.00205339 | 0.1400576 | 0.78917197 | 2.1126 | 0 | 33 | CXCL1 CXCL8 CXCL3 CXCL6 CXCL2 CXCL5 CCL20 |
| GOBP_ANTIMICROBIAL_HUMORAL_IMMUNE_RESPONSE_MEDIATED_BY_ANT | 0.00205339 | 0.1400576 | 0.79750573 | 2.09263241 | 0 | 31 | [illegible] |
| GOCC_CATALYTIC_STEP_2_SPLICEOSOME | 0.00205339 | 0.1400576 | 0.74785828 | 2.00200397 | 0 | 33 | [illegible] |
| GOBP_ANTIMICROBIAL_HUMORAL_RESPONSE | 0.00205761 | 0.1400576 | 0.68779693 | 1.9495456 | 0 | 50 | [illegible] |
| GOBP_CHROMOSOME_SEPARATION | 0.00205761 | 0.1400576 | 0.65268954 | 1.83738172 | 0 | 47 | [illegible] |
| GOBP_CARBOHYDRATE_CATABOLIC_PROCESS | 0.00205761 | 0.1400576 | 0.52429871 | 1.6213066 | 0 | 76 | [illegible] |
| GOBP_NUCLEOTIDE_PHOSPHORYLATION | 0.00206186 | 0.1400576 | 0.64778214 | 1.83084956 | 0 | 49 | [illegible] |
| GOBP_NEUTROPHIL_CHEMOTAXIS | 0.00206186 | 0.1400576 | 0.62389278 | 1.79796969 | 0 | 54 | [illegible] |
| GOCC_NUCLEAR_PROTEIN_CONTAINING_COMPLEX | 0.00206186 | 0.1400576 | 0.43230123 | 1.66384341 | 0 | 457 | [illegible] |
| GOBP_CARBOHYDRATE_DERIVATIVE_BIOSYNTHETIC_PROCESS | 0.00206186 | 0.1400576 | 0.42679139 | 1.60036423 | 0 | 345 | [illegible] |
| GOBP_RESPONSE_TO_LIPID | 0.00206186 | 0.1400576 | 0.39651312 | 1.51076657 | 0 | 397 | [illegible] |
| GOBP_MITOTIC_CELL_CYCLE | 0.00206186 | 0.1400576 | 0.37202165 | 1.43232834 | 0 | 461 | [illegible] |
| GOCC_CARDIAC_MYOFIBRIL | 0.00206186 | 0.1400576 | 0.99891631 | 1.42612376 | 0 | 2 | CRYAB DES |
| GOBP_ENDOPLASMIC_RETICULUM_TO_GOLGI_VESICLE_MEDIATED_TRANSPO | 0.00206612 | 0.1400576 | 0.56193279 | 1.81056309 | 0 | 93 | [illegible] |
| GOBP_BINDING_OF_SPERM_TO_ZONA_PELLUCIDA | 0.00207039 | 0.1400576 | 0.86619565 | 2.01763323 | 0 | 19 | B4GALT1 CCT2 ALDOA CCT7 CCT8 CCT3 CCT4 CCT5 |
| GOBP_CELLULAR_RESPONSE_TO_BIOTIC_STIMULUS | 0.00207039 | 0.1400576 | 0.54571954 | 1.77192008 | 0 | 100 | [illegible] |
| GOBP_CHEMOKINE_PRODUCTION | 0.00207039 | 0.1400576 | 0.62447543 | 1.74265816 | 0 | 43 | [illegible] |
| GOBP_CENTROSOME_DUPLICATION | 0.00207039 | 0.1400576 | 0.62267071 | 1.73782251 | 0 | 43 | [illegible] |
| GOBP_RESPONSE_TO_INTERLEUKIN_1 | 0.00207039 | 0.1400576 | 0.55025819 | 1.67870993 | 0 | 70 | [illegible] |
| GOMF_HELICASE_ACTIVITY | 0.00207039 | 0.1400576 | 0.50443234 | 1.6378629 | 0 | 100 | [illegible] |
| GOBP_MYELOID_LEUKOCYTE_MIGRATION | 0.00207039 | 0.1400576 | 0.50388228 | 1.63607691 | 0 | 100 | [illegible] |
| GOBP_CELLULAR_RESPONSE_TO_MOLECULE_OF_BACTERIAL_ORIGIN | 0.00207469 | 0.1400576 | 0.60842999 | 1.93665184 | 0 | 88 | [illegible] |
| GOBP_NUCLEOSOME_ASSEMBLY | 0.00207469 | 0.1400576 | 0.55286047 | 1.75979185 | 0 | 88 | [illegible] |
| GOBP_DNA_PACKAGING | 0.00207469 | 0.1400576 | 0.50398847 | 1.6905684 | 0 | 120 | [illegible] |
| GOBP_RESPONSE_TO_BACTERIUM | 0.00207469 | 0.1400576 | 0.42938747 | 1.57702778 | 0 | 263 | [illegible] |
| GOBP_NUCLEOBASE_CONTAINING_SMALL_MOLECULE_METABOLIC_PROCESS | 0.00207469 | 0.1400576 | 0.42131858 | 1.55601909 | 0 | 278 | [illegible] |
| GOBP_MYELOID_LEUKOCYTE_MEDIATED_IMMUNITY | 0.00207469 | 0.1400576 | 0.39119657 | 1.44477208 | 0 | 278 | [illegible] |
| GOBP_CARBOHYDRATE_METABOLIC_PROCESS | 0.00207469 | 0.1400576 | 0.37944661 | 1.40086777 | 0 | 288 | [illegible] |
| GOBP_RNA_PROCESSING | 0.002079 | 0.1400576 | 0.47783656 | 1.77070449 | 0 | 318 | [illegible] |
| GOBP_PYRUVATE_METABOLIC_PROCESS | 0.002079 | 0.1400576 | 0.60597814 | 1.72058571 | 0 | 51 | [illegible] |
| GOBP_CELLULAR_RESPONSE_TO_DNA_DAMAGE_STIMULUS | 0.002079 | 0.1400576 | 0.38422196 | 1.44556988 | 0 | 361 | [illegible] |
| GOBP_ORGANELLE_LOCALIZATION | 0.002079 | 0.1400576 | 0.38223514 | 1.41069826 | 0 | 286 | [illegible] |
| GOBP_RNA_SPLICING_VIA_TRANSESTERIFICATION_REACTIONS | 0.00208333 | 0.1400576 | 0.61073277 | 2.01815309 | 0 | 116 | [illegible] |
| GOBP_REGULATION_OF_TELOMERE_MAINTENANCE | 0.00208333 | 0.1400576 | 0.72907873 | 1.85798359 | 0 | 29 | [illegible] |
| GOBP_PROTEIN_LOCALIZATION_TO_CHROMOSOME | 0.00208333 | 0.1400576 | 0.66637025 | 1.80724627 | 0 | 38 | [illegible] |
| GOBP_REGULATION_OF_CHROMOSOME_SEPARATION | 0.00208333 | 0.1400576 | 0.66547104 | 1.80480754 | 0 | 38 | [illegible] |
| GOBP_DNA_REPLICATION | 0.00208333 | 0.1400576 | 0.53690035 | 1.7653539 | 0 | 112 | [illegible] |
| GOBP_RNA_LOCALIZATION | 0.00208333 | 0.1400576 | 0.56179887 | 1.71602053 | 0 | 72 | [illegible] |
| GOBP_MITOTIC_CELL_CYCLE_CHECKPOINT | 0.00208333 | 0.1400576 | 0.55264115 | 1.698863 | 0 | 74 | [illegible] |
| GOBP_CHROMATIN_ASSEMBLY_OR_DISASSEMBLY | 0.00208333 | 0.1400576 | 0.51271424 | 1.69425297 | 0 | 114 | [illegible] |
| GOBP_DNA_DEPENDENT_DNA_REPLICATION | 0.00208768 | 0.1400576 | 0.60088203 | 1.76158221 | 0 | 58 | [illegible] |
| GOCC_CONDENSED_CHROMOSOME_CENTROMERIC_REGION | 0.00208768 | 0.1400576 | 0.57042022 | 1.67227853 | 0 | 58 | [illegible] |
| GOCC_ZONA_PELLUCIDA_RECEPTOR_COMPLEX | 0.00208768 | 0.1400576 | 0.96784824 | 1.66755067 | 0 | 5 | CCT2 CCT8 CCT3 CCT4 |
| GOMF_KETOSTEROID_MONOOXYGENASE_ACTIVITY | 0.00208768 | 0.1400576 | 0.96667414 | 1.66552776 | 0 | 5 | AKR1C2 AKR1C1 AKR1C3 |
| GOBP_DNA_METABOLIC_PROCESS | 0.00208768 | 0.1400576 | 0.40669286 | 1.54688623 | 0 | 392 | [illegible] |
| GOBP_MICROTUBULE_CYTOSKELETON_ORGANIZATION | 0.00208768 | 0.1400576 | 0.37872557 | 1.39815719 | 0 | 290 | [illegible] |
| GOBP_RESPONSE_TO_CHEMOKINE | 0.00209205 | 0.1400576 | 0.7331237 | 2.09028513 | 0 | 52 | [illegible] |
| GOMF_CYTOKINE_ACTIVITY | 0.00209205 | 0.1400576 | 0.54818586 | 1.82705373 | 0 | 117 | [illegible] |
| GOMF_CYTOKINE_RECEPTOR_BINDING | 0.00209205 | 0.1400576 | 0.54762844 | 1.81413782 | 0 | 115 | [illegible] |
| GOCC_CHROMOSOME_CENTROMERIC_REGION | 0.00209205 | 0.1400576 | 0.5687743 | 1.81065042 | 0 | 89 | [illegible] |
| GOBP_GRANULOCYTE_MIGRATION | 0.00209205 | 0.1400576 | 0.55030864 | 1.68389527 | 0 | 73 | [illegible] |
| GOBP_PROTEIN_FOLDING | 0.00209205 | 0.1400576 | 0.49170149 | 1.65765129 | 0 | 123 | [illegible] |
| GOBP_NUCLEOSOME_ORGANIZATION | 0.00209205 | 0.1400576 | 0.49177517 | 1.59671155 | 0 | 101 | [illegible] |
| GOMF_ATPASE_ACTIVITY | 0.00209205 | 0.1400576 | 0.40286431 | 1.49090665 | 0 | 302 | [illegible] |
| GOBP_MYELOID_LEUKOCYTE_ACTIVATION | 0.00209205 | 0.1400576 | 0.38419526 | 1.42923417 | 0 | 325 | [illegible] |
| GOBP_SISTER_CHROMATID_SEGREGATION | 0.00209644 | 0.1400576 | 0.6228814 | 2.00635515 | 0 | 96 | [illegible] |
| GOBP_TELOMERE_ORGANIZATION | 0.00209644 | 0.1400576 | 0.65341097 | 1.97892727 | 0 | 67 | [illegible] |
| GOBP_HUMORAL_IMMUNE_RESPONSE | 0.00209644 | 0.1400576 | 0.54175049 | 1.78199652 | 0 | 111 | [illegible] |
| GOCC_CONDENSED_CHROMOSOME | 0.00209644 | 0.1400576 | 0.53570512 | 1.7177422 | 0 | 92 | [illegible] |
| GOCC_KINETOCHORE | 0.00209644 | 0.1400576 | 0.55170777 | 1.67090789 | 0 | 67 | [illegible] |
| GOCC_NUCLEAR_CHROMOSOME | 0.00209644 | 0.1400576 | 0.51699588 | 1.66539561 | 0 | 96 | [illegible] |
| GOMF_OXIDOREDUCTASE_ACTIVITY_ACTING_ON_CH_OH_GROUP_OF_DONO | 0.00209644 | 0.1400576 | 0.49829168 | 1.59774369 | 0 | 92 | [illegible] |
| GOBP_SPINDLE_ORGANIZATION | 0.00209644 | 0.1400576 | 0.4908527 | 1.57392261 | 0 | 92 | [illegible] |
| GOBP_CELL_DIVISION | 0.00209644 | 0.1400576 | 0.37309809 | 1.37437946 | 0 | 290 | [illegible] |
| GOMF_CHEMOKINE_RECEPTOR_BINDING | 0.00210084 | 0.1400576 | 0.75617861 | 2.04607188 | 0 | 37 | CXCL1 CXCL8 CXCL3 CXCL6 CXCL2 CXCL5 CCL20 CNIH4 |
| GOMF_RNA_HELICASE_ACTIVITY | 0.00210084 | 0.1400576 | 0.5664452 | 1.69781822 | 0 | 64 | [illegible] |
| GOBP_GRANULOCYTE_CHEMOTAXIS | 0.00210084 | 0.1400576 | 0.55685185 | 1.66857233 | 0 | 64 | [illegible] |
| GOMF_CATALYTIC_ACTIVITY_ACTING_ON_RNA | 0.00210084 | 0.1400576 | 0.46039388 | 1.58929037 | 0 | 165 | [illegible] |
| GOBP_REGULATION_OF_CHROMOSOME_ORGANIZATION | 0.00210526 | 0.1400576 | 0.68181391 | 2.21894882 | 0 | 107 | [illegible] |
| GOBP_NEGATIVE_REGULATION_OF_HORMONE_SECRETION | 0.00210526 | 0.1400576 | 0.71335268 | 1.8304405 | 0 | 30 | INHBA IL11 LIF JAGN1 IRS1 KLF7 ENY2 ADRA2A LEP |
| GOBP_NEUTROPHIL_MIGRATION | 0.00210526 | 0.1400576 | 0.60941163 | 1.80801279 | 0 | 62 | [illegible] |
| GOBP_TELOMERE_MAINTENANCE_VIA_TELOMERE_LENGTHENING | 0.00210526 | 0.1400576 | 0.71244916 | 1.8061081 | 0 | 28 | [illegible] |
| GOMF_UNFOLDED_PROTEIN_BINDING | 0.00210526 | 0.1400576 | 0.58144493 | 1.75776119 | 0 | 66 | [illegible] |
| GOCC_RIBONUCLEOPROTEIN_COMPLEX | 0.00210526 | 0.1400576 | 0.45887716 | 1.63271606 | 0 | 197 | [illegible] |
| GOBP_GLYCOPROTEIN_BIOSYNTHETIC_PROCESS | 0.00210526 | 0.1400576 | 0.46172685 | 1.61458833 | 0 | 181 | [illegible] |
| GOBP_GLYCOSYLATION | 0.00210526 | 0.1400576 | 0.46881193 | 1.59298185 | 0 | 131 | [illegible] |
| GOBP_POSITIVE_REGULATION_OF_CHROMOSOME_ORGANIZATION | 0.0021097 | 0.1400576 | 0.69978772 | 2.05284638 | 0 | 59 | [illegible] |
| GOBP_RIBOSOME_BIOGENESIS | 0.0021097 | 0.1400576 | 0.61907677 | 1.95694883 | 0 | 86 | [illegible] |
| GOCC_SPLICEOSOMAL_COMPLEX | 0.0021097 | 0.1400576 | 0.62279613 | 1.85318033 | 0 | 63 | [illegible] |
| GOBP_NUCLEOSIDE_DIPHOSPHATE_METABOLIC_PROCESS | 0.0021097 | 0.1400576 | 0.56820216 | 1.69371256 | 0 | 63 | [illegible] |
| GOBP_GLYCOPROTEIN_METABOLIC_PROCESS | 0.0021097 | 0.1400576 | 0.42100646 | 1.51197201 | 0 | 218 | [illegible] |
| GOBP_LEUKOCYTE_MIGRATION | 0.0021097 | 0.1400576 | 0.40995095 | 1.47141271 | 0 | 217 | [illegible] |
| GOCC_CHROMOSOMAL_REGION | 0.00211416 | 0.1400576 | 0.54892129 | 1.8720832 | 0 | 140 | [illegible] |
| GOBP_MRNA_METABOLIC_PROCESS | 0.00211416 | 0.1400576 | 0.43875213 | 1.60577582 | 0 | 257 | [illegible] |
| GOBP_NCRNA_METABOLIC_PROCESS | 0.00211864 | 0.1400576 | 0.4711626 | 1.63486038 | 0 | 172 | [illegible] |
| GOBP_ORGANELLE_FISSION | 0.00211864 | 0.1400576 | 0.42546536 | 1.52518677 | 0 | 216 | [illegible] |
| GOBP_MRNA_PROCESSING | 0.00212314 | 0.1400576 | 0.49751224 | 1.73063677 | 0 | 176 | [illegible] |
| GOBP_DNA_CONFORMATION_CHANGE | 0.00212314 | 0.1400576 | 0.48529595 | 1.68451048 | 0 | 173 | [illegible] |
| GOBP_NUCLEAR_CHROMOSOME_SEGREGATION | 0.00212766 | 0.1400576 | 0.56567064 | 1.90772201 | 0 | 128 | [illegible] |
| GOBP_NCRNA_PROCESSING | 0.00212766 | 0.1400576 | 0.53382826 | 1.81102136 | 0 | 136 | [illegible] |
| GOBP_RESPONSE_TO_MOLECULE_OF_BACTERIAL_ORIGIN | 0.00212766 | 0.1400576 | 0.48329725 | 1.66479303 | 0 | 161 | [illegible] |
| GOBP_RIBONUCLEOPROTEIN_COMPLEX_BIOGENESIS | 0.0021322 | 0.1400576 | 0.49027634 | 1.67654817 | 0 | 145 | [illegible] |
| GOMF_BILE_ACID_BINDING | 0.0021322 | 0.1400576 | 0.97784228 | 1.57207327 | 0 | 4 | AKR1C2 AKR1C1 AKR1C3 |
| GOBP_CHROMOSOME_SEGREGATION | 0.00213675 | 0.1400576 | 0.54782529 | 1.87626117 | 0 | 148 | [illegible] |
| GOBP_RNA_SPLICING | 0.00213675 | 0.1400576 | 0.50466337 | 1.7259602 | 0 | 146 | [illegible] |
| GOBP_POSITIVE_REGULATION_OF_TRANSFORMING_GROWTH_FACTOR_BETA | 0.00214133 | 0.1400576 | 0.98105175 | 1.49025741 | 0 | 3 | FURIN ATP6AP2 |
| GOBP_CELLULAR_RESPONSE_TO_LIPID | 0.00215054 | 0.1400576 | 0.44817711 | 1.62461608 | 0 | 245 | [illegible] |
| GOBP_GENERATION_OF_PRECURSOR_METABOLITES_AND_ENERGY | 0.00215983 | 0.1400576 | 0.4518572 | 1.62349128 | 0 | 225 | [illegible] |
| GOBP_MUSCLE_CELL_DIFFERENTIATION | 0.00374532 | 0.18894716 | -0.4234259 | -1.4438588 | 1 | 148 | [illegible] |
| GOBP_PLASMA_MEMBRANE_RAFT_ASSEMBLY | 0.00375235 | 0.18894716 | -0.961485 | -1.5458988 | 1 | 4 | FLOT1 CAV1 CAV2 |
| GOBP_REGULATION_OF_ACTIN_FILAMENT_BASED_PROCESS | 0.0037594 | 0.18894716 | -0.4131797 | -1.4482245 | 1 | 187 | [illegible] |
| GOBP_STRIATED_MUSCLE_CELL_DIFFERENTIATION | 0.00377358 | 0.18894716 | -0.4950254 | -1.628887 | 1 | 108 | [illegible] |
| GOBP_NEGATIVE_REGULATION_OF_INTRACELLULAR_SIGNAL_TRANSDUCTION | 0.00378788 | 0.18894716 | -0.419845 | -1.4749241 | 1 | 195 | [illegible] |
| GOCC_ENDOCYTIC_VESICLE | 0.00380228 | 0.18894716 | -0.4246353 | -1.4263221 | 1 | 136 | [illegible] |
| GOBP_POSITIVE_REGULATION_OF_PHOSPHORUS_METABOLIC_PROCESS | 0.00380952 | 0.18894716 | -0.3426538 | -1.3060894 | 1 | 446 | [illegible] |
| GOBP_B_CELL_CHEMOTAXIS | 0.00382409 | 0.18894716 | -0.9387091 | -1.5848095 | 1 | 5 | GAS6 CH25H HSD3B7 |
| GOBP_RESPONSE_TO_MINERALOCORTICOID | 0.00383142 | 0.18894716 | -0.7328172 | -1.7023622 | 1 | 18 | CDKN1A FOSL1 GPER1 CALM3 CCN2 CCND1 KRAS |
| GOBP_LOCOMOTION | 0.00385356 | 0.18894716 | -0.3095741 | -1.2363649 | 1 | 847 | [illegible] |
| GOBP_CARDIAC_SEPTUM_MORPHOGENESIS | 0.00385356 | 0.18894716 | -0.7006813 | -1.651225 | 1 | 19 | CITED2 GATA6 BMP4 ACVR1 BMPR1A DHRS3 FZD2 |
| GOMF_LBD_DOMAIN_BINDING | 0.00386847 | 0.18894716 | -0.9878255 | -1.3723318 | 1 | 2 | CITED2 |
| GOMF_HYALURONIC_ACID_BINDING | 0.00389105 | 0.18894716 | -0.8284442 | -1.73522 | 1 | 12 | CEMIP ACAN LAYN HAPLN3 HAPLN1 CD44 |
| GOBP_REGULATION_OF_PROTEIN_LOCALIZATION_TO_MEMBRANE | 0.00390625 | 0.18894716 | -0.5375316 | -1.6504619 | 1 | 80 | [illegible] |
| GOBP_NEGATIVE_REGULATION_OF_DEVELOPMENTAL_PROCESS | 0.00393701 | 0.18894716 | -0.3662401 | -1.3547171 | 1 | 340 | [illegible] |
| GOCC_CORTICAL_ACTIN_CYTOSKELETON | 0.0039604 | 0.18894716 | -0.6315524 | -1.7075753 | 1 | 35 | [illegible] |
| GOMF_ADENYL_NUCLEOTIDE_BINDING | 0.00399202 | 0.18894716 | 0.32836824 | 1.30705237 | 1 | 748 | [illegible] |
| GOBP_CELL_CYCLE | 0.00401606 | 0.18894716 | 0.34266798 | 1.36953354 | 1 | 795 | [illegible] |
| GOBP_IMMUNE_EFFECTOR_PROCESS | 0.00401606 | 0.18894716 | 0.34268856 | 1.33557605 | 1 | 560 | [illegible] |
| GOBP_PROTEIN_LOCALIZATION_TO_KINETOCHORE | 0.00402414 | 0.18894716 | 0.88129952 | 1.74430232 | 1 | 10 | BUB3 AURKB CDK1 CENPQ CENPT HASPIN IK |
| GOBP_CELL_ACTIVATION | 0.0040568 | 0.18894716 | 0.33849883 | 1.33648955 | 1 | 667 | [illegible] |
| GOCC_ORGANELLE_SUBCOMPARTMENT | 0.0040568 | 0.18894716 | 0.32184473 | 1.28664722 | 1 | 808 | [illegible] |
| GOBP_MEIOTIC_CELL_CYCLE | 0.00407332 | 0.18894716 | 0.52251311 | 1.64488692 | 1 | 81 | [illegible] |
| GOBP_PROTEIN_STABILIZATION | 0.00407332 | 0.18894716 | 0.51582583 | 1.62383517 | 1 | 81 | [illegible] |
| GOBP_REGULATION_OF_TELOMERE_MAINTENANCE_VIA_TELOMERE_LENGTH | 0.00408163 | 0.18894716 | 0.72039793 | 1.78700116 | 1 | 26 | [illegible] |
| GOMF_STEROID_DEHYDROGENASE_ACTIVITY | 0.00408163 | 0.18894716 | 0.71244286 | 1.76726802 | 1 | 26 | AKR1C2 AKR1C1 HSD11B1 HSD17B10 HSD17B11 |
| GOBP_NAD_METABOLIC_PROCESS | 0.00408163 | 0.18894716 | 0.71150794 | 1.76494887 | 1 | 26 | LDHA GPI GAPDH ENO1 ALDOA ENO2 ADPGK CD38 HK1 BPGM |
| GOBP_ESTABLISHMENT_OF_PROTEIN_LOCALIZATION_TO_CHROMOSOME | 0.00408163 | 0.18894716 | 0.82095835 | 1.74595481 | 1 | 13 | [illegible] |
| GOBP_ENDOPLASMIC_RETICULUM_TO_CYTOSOL_TRANSPORT | 0.00408163 | 0.18894716 | 0.80914258 | 1.72082587 | 1 | 13 | AUP1 EDEM2 HM13 EDEM1 BCAP31 DERL1 DERL2 ERLEC1 |
| GOBP_MULTICELLULAR_ORGANISM_AGING | 0.00408163 | 0.18894716 | 0.79767375 | 1.69643481 | 1 | 13 | CRYAB COMP CISD2 LEP |
| GOBP_INFLAMMATORY_RESPONSE | 0.00408163 | 0.18894716 | 0.38171467 | 1.4294537 | 1 | 338 | [illegible] |
| GOBP_REGULATION_OF_ESTABLISHMENT_OF_PROTEIN_LOCALIZATION_TO_C | 0.00408998 | 0.18894716 | 0.90491902 | 1.70999691 | 1 | 8 | CCT2 CCT6A CCT7 CCT8 CCT3 CCT4 CCT5 DKC1 |
| GOBP_POSITIVE_REGULATION_OF_ESTABLISHMENT_OF_PROTEIN_LOCALIZAT | 0.00408998 | 0.18894716 | 0.90491902 | 1.70999691 | 1 | 8 | CCT2 CCT6A CCT7 CCT8 CCT3 CCT4 CCT5 DKC1 |
| GOBP_POSITIVE_REGULATION_OF_TELOMERASE_RNA_LOCALIZATION_TO_CA | 0.00408998 | 0.18894716 | 0.90491902 | 1.70999691 | 1 | 8 | CCT2 CCT6A CCT7 CCT8 CCT3 CCT4 CCT5 DKC1 |
| GOBP_PROTEIN_LOCALIZATION_TO_CHROMOSOME_TELOMERIC_REGION | 0.00409836 | 0.18894716 | 0.82217619 | 1.81801301 | 1 | 15 | [illegible] |
| GOBP_RIBOSOME_ASSEMBLY | 0.00409836 | 0.18894716 | 0.82774146 | 1.78169478 | 1 | 14 | [illegible] |
| GOBP_POSITIVE_REGULATION_OF_MACROPHAGE_ACTIVATION | 0.00409836 | 0.18894716 | 0.85091084 | 1.76995252 | 1 | 12 | CTSC IL33 KARS1 IL1RL1 HSPD1 |
| GOBP_RESPONSE_TO_PROSTAGLANDIN | 0.00409836 | 0.18894716 | 0.79621156 | 1.71382738 | 1 | 14 | AKR1C2 CCR7 AKAP8 AKT1 GNG2 ADCY6 |
| GOBP_RRNA_METABOLIC_PROCESS | 0.00411523 | 0.18894716 | 0.54321876 | 1.65702205 | 1 | 68 | [illegible] |
| GOBP_MEIOTIC_CELL_CYCLE_PROCESS | 0.00411523 | 0.18894716 | 0.53416186 | 1.63184826 | 1 | 69 | [illegible] |
| GOBP_MINERALOCORTICOID_METABOLIC_PROCESS | 0.00412371 | 0.18894716 | 0.89794561 | 1.73729044 | 1 | 9 | BMP6 BMP2 |
| GOBP_PROTEIN_LOCALIZATION_TO_NUCLEAR_BODY | 0.00412371 | 0.18894716 | 0.88101376 | 1.70452193 | 1 | 9 | CCT2 CCT6A CCT7 CCT8 CCT3 CCT4 CCT5 DKC1 |
| GOBP_PROTEIN_LOCALIZATION_TO_NUCLEOPLASM | 0.00412371 | 0.18894716 | 0.88101376 | 1.70452193 | 1 | 9 | CCT2 CCT6A CCT7 CCT8 CCT3 CCT4 CCT5 DKC1 |
| GOBP_REGULATION_OF_LIPOPROTEIN_LIPASE_ACTIVITY | 0.00412371 | 0.18894716 | 0.83628794 | 1.69525193 | 1 | 11 | ANGPTL4 FURIN APOA5 APOC3 |
| GOMF_ANDROSTAN_3_ALPHA_17_BETA_DIOL_DEHYDROGENASE_ACTIVITY | 0.00412371 | 0.18894716 | 0.90683827 | 1.67755086 | 1 | 7 | AKR1C2 AKR1C1 HSD17B10 |
| GOBP_POSITIVE_REGULATION_OF_NUCLEOTIDE_BIOSYNTHETIC_PROCESS | 0.00412371 | 0.18894716 | 0.98694015 | 1.40902574 | 1 | 2 | ENO1 GUCA1B |
| GOBP_PROTEIN_CONTAINING_COMPLEX_SUBUNIT_ORGANIZATION | 0.00413223 | 0.18894716 | 0.3258879 | 1.30486825 | 1 | 833 | [illegible] |
| GOBP_NADH_METABOLIC_PROCESS | 0.00414079 | 0.18894716 | 0.79554818 | 1.85307378 | 1 | 19 | GPI GAPDH ENO1 ALDOA ENO2 ADPGK HK1 BPGM |
| GOMF_RIBONUCLEOTIDE_BINDING | 0.00414938 | 0.18894716 | 0.32182547 | 1.29100217 | 1 | 869 | [illegible] |
| GOBP_ANAPHASE_PROMOTING_COMPLEX_DEPENDENT_CATABOLIC_PROCES | 0.00416667 | 0.18894716 | 0.71268577 | 1.79972555 | 1 | 27 | [illegible] |
| GOCC_PRERIBOSOME | 0.00416667 | 0.18894716 | 0.78011879 | 1.7946429 | 1 | 18 | [illegible] |
| GOBP_GLUCOSE_CATABOLIC_PROCESS | 0.00416667 | 0.18894716 | 0.74941286 | 1.72400472 | 1 | 18 | GPI GAPDH ENO1 ALDOA ENO2 ADPGK HK1 BPGM |
| GOBP_CHROMATIN_ORGANIZATION_INVOLVED_IN_REGULATION_OF_TRANSC | 0.00416667 | 0.18894716 | 0.48036476 | 1.53650283 | 1 | 90 | [illegible] |
| GOMF_G_PROTEIN_COUPLED_RECEPTOR_BINDING | 0.00416667 | 0.18894716 | 0.44862544 | 1.51762638 | 1 | 126 | [illegible] |
| GOBP_SISTER_CHROMATID_COHESION | 0.0041841 | 0.18894716 | 0.76528159 | 1.84755118 | 1 | 22 | [illegible] |
| GOBP_GLYCOLYTIC_PROCESS_THROUGH_FRUCTOSE_6_PHOSPHATE | 0.0041841 | 0.18894716 | 0.8011427 | 1.81877368 | 1 | 17 | GPI GAPDH ENO1 ALDOA ENO2 ADPGK HK1 BPGM |
| GOCC_ENDOPLASMIC_RETICULUM_GOLGI_INTERMEDIATE_COMPARTMENT | 0.0041841 | 0.18894716 | 0.60382645 | 1.72162661 | 1 | 52 | [illegible] |
| GOMF_ESTRADIOL_17_BETA_DEHYDROGENASE_ACTIVITY | 0.0041841 | 0.18894716 | 0.77074172 | 1.71286593 | 1 | 16 | AKR1C2 AKR1C1 HSD17B10 HSD17B11 HSD17B2 HSD17B7 |
| GOBP_OLIGOSACCHARIDE_LIPID_INTERMEDIATE_BIOSYNTHETIC_PROCESS | 0.0041841 | 0.18894716 | 0.76402018 | 1.69792824 | 1 | 16 | ALG3 ALG2 DPAGT1 ALG8 DOLPP1 ALG6 ALG11 ALG1 DHDDS |
| GOBP_LEUKOCYTE_CHEMOTAXIS | 0.00419287 | 0.18894716 | 0.4740953 | 1.55965623 | 1 | 111 | [illegible] |
| GOBP_REGULATION_OF_PROTEIN_MODIFICATION_BY_SMALL_PROTEIN_CON | 0.00419287 | 0.18894716 | 0.4773685 | 1.53863795 | 1 | 95 | [illegible] |
| GOBP_SPERM_EGG_RECOGNITION | 0.00421053 | 0.18894716 | 0.6791721 | 1.74273382 | 1 | 30 | B4GALT1 CCT2 ALDOA CCT7 CCT8 CCT3 CCT4 CCT5 |
| GOBP_RNA_PHOSPHODIESTER_BOND_HYDROLYSIS | 0.00421053 | 0.18894716 | 0.51644068 | 1.5612476 | 1 | 66 | [illegible] |
| GOBP_PROTEIN_DNA_COMPLEX_SUBUNIT_ORGANIZATION | 0.00422833 | 0.18894716 | 0.4750755 | 1.62024255 | 1 | 140 | [illegible] |
| GOBP_DNA_RECOMBINATION | 0.00423729 | 0.18894716 | 0.47547423 | 1.55366186 | 1 | 110 | [illegible] |
| GOBP_GOLGI_VESICLE_TRANSPORT | 0.00424628 | 0.18894716 | 0.41828181 | 1.45485915 | 1 | 176 | [illegible] |
| GOBP_CHROMATIN_ORGANIZATION | 0.00424628 | 0.18894716 | 0.38186599 | 1.44683577 | 1 | 383 | [illegible] |
| GOBP_REGULATION_OF_PROTEIN_STABILITY | 0.00425532 | 0.18894716 | 0.46986087 | 1.5863448 | 1 | 129 | [illegible] |
| GOBP_CELL_CHEMOTAXIS | 0.0042735 | 0.18894716 | 0.44404247 | 1.5208127 | 1 | 148 | [illegible] |
| GOBP_REGULATION_OF_VASCULATURE_DEVELOPMENT | 0.0042735 | 0.18894716 | 0.43699183 | 1.48996344 | 1 | 142 | [illegible] |
| GOBP_DNA_REPAIR | 0.00431034 | 0.18971368 | 0.39857349 | 1.42918249 | 1 | 222 | [illegible] |
| GOCC_CELL_CELL_JUNCTION | 0.00557621 | 0.22284759 | -0.3831268 | -1.3802176 | 2 | 248 | [illegible] |
| GOCC_COLLAGEN_CONTAINING_EXTRACELLULAR_MATRIX | 0.00559701 | 0.22284759 | -0.3935268 | -1.408377 | 2 | 229 | [illegible] |
| GOBP_HEART_DEVELOPMENT | 0.00560748 | 0.22284759 | -0.403953 | -1.4337063 | 2 | 208 | [illegible] |
| GOBP_PEPTIDYL_TYROSINE_MODIFICATION | 0.00562852 | 0.22284759 | -0.3960726 | -1.3885746 | 2 | 189 | [illegible] |
| GOBP_REGULATION_OF_ACTIN_FILAMENT_ORGANIZATION | 0.00562852 | 0.22284759 | -0.4231954 | -1.4237395 | 2 | 130 | [illegible] |
| GOBP_REGULATION_OF_GLYCOGEN_CATABOLIC_PROCESS | 0.00567108 | 0.22284759 | 0.99640324 | 1.33359537 | 2 | 1 | HMGB1 |
| GOBP_POSITIVE_REGULATION_OF_TOLL_LIKE_RECEPTOR_9_SIGNALING_PATH | 0.00567108 | 0.22284759 | 0.99640324 | 1.33359537 | 2 | 1 | HMGB1 |
| GOBP_OSSIFICATION | 0.00567108 | 0.22284759 | -0.4033082 | -1.4036992 | 2 | 179 | [illegible] |
| GOBP_MORPHOGENESIS_OF_AN_EPITHELIUM | 0.0056926 | 0.22284759 | -0.384447 | -1.3692173 | 2 | 219 | [illegible] |
| GOBP_RESPONSE_TO_EXTRACELLULAR_STIMULUS | 0.0056926 | 0.22284759 | -0.4038211 | -1.4137043 | 2 | 197 | [illegible] |
| GOBP_RESPONSE_TO_INTERFERON_GAMMA | 0.00571429 | 0.22284759 | -0.4814686 | -1.5642746 | 2 | 102 | [illegible] |
| GOCC_VACUOLE | 0.00572519 | 0.22284759 | -0.346285 | -1.2940115 | 2 | 365 | [illegible] |
| GOBP_INNATE_IMMUNE_RESPONSE | 0.00575816 | 0.22284759 | -0.3508897 | -1.3038664 | 2 | 348 | [illegible] |
| GOBP_POSITIVE_REGULATION_OF_KIDNEY_DEVELOPMENT | 0.00580271 | 0.22284759 | -0.9851494 | -1.3686141 | 2 | 2 | BMP4 |
| GOBP_RETINOIC_ACID_METABOLIC_PROCESS | 0.00580271 | 0.22284759 | -0.6553316 | -1.6206844 | 2 | 23 | [illegible] |
| GOBP_POSITIVE_REGULATION_OF_CYSTEINE_TYPE_ENDOPEPTIDASE_ACTIVIT | 0.00580271 | 0.22284759 | -0.8907935 | -1.6426555 | 2 | 7 | GSN BAX FAS JAK2 |
| GOBP_POSITIVE_REGULATION_OF_MYOBLAST_FUSION | 0.00580271 | 0.22284759 | -0.8990604 | -1.6578998 | 2 | 7 | GDF15 CAPN2 FLOT1 EHD2 |
| GOCC_COSTAMERE | 0.00580271 | 0.22284759 | -0.8063971 | -1.6655306 | 2 | 11 | AHNAK2 KRT19 FLNC AHNAK ANK2 ANK3 DAG1 |
| GOBP_PHOSPHOLIPID_DEPHOSPHORYLATION | 0.00580271 | 0.22284759 | -0.8389684 | -1.7328033 | 2 | 11 | FIG4 INPP5B EPHX2 INPP5E INPP5K |
| GOCC_SARCOLEMMA | 0.00581395 | 0.22284759 | -0.5409688 | -1.6224951 | 2 | 68 | [illegible] |
| GOBP_REGULATION_OF_PROTEIN_LOCALIZATION_TO_CELL_PERIPHERY | 0.00581395 | 0.22284759 | -0.5724119 | -1.6469675 | 2 | 50 | [illegible] |
| GOMF_TRANSMEMBRANE_RECEPTOR_PROTEIN_KINASE_ACTIVITY | 0.00581395 | 0.22284759 | -0.5771553 | -1.6606154 | 2 | 50 | [illegible] |
| GOBP_RHO_PROTEIN_SIGNAL_TRANSDUCTION | 0.00584795 | 0.22284759 | -0.5245783 | -1.6235576 | 2 | 82 | [illegible] |
| GOBP_REGULATION_OF_PROTEIN_PHOSPHORYLATION | 0.00588235 | 0.22284759 | -0.3229083 | -1.2512394 | 2 | 558 | [illegible] |
| GOBP_INOSITOL_PHOSPHATE_MEDIATED_SIGNALING | 0.00592885 | 0.22284759 | -0.6484404 | -1.6209978 | 2 | 25 | FHL2 ATP2B4 ADGRB2 HOMER3 |
| GOBP_T_CELL_ACTIVATION_VIA_T_CELL_RECEPTOR_CONTACT_WITH_ANTIGE | 0.00594059 | 0.22284759 | -0.8060319 | -1.6288907 | 2 | 10 | LGALS3 FGL2 CD81 |
| GOBP_PHOSPHATIDYLINOSITOL_DEPHOSPHORYLATION | 0.00594059 | 0.22284759 | -0.8309362 | -1.6792192 | 2 | 10 | FIG4 INPP5B INPP5E INPP5K |
| GOBP_TELOMERASE_RNA_LOCALIZATION | 0.00603622 | 0.22284759 | 0.86993993 | 1.72181899 | 2 | 10 | CCT2 EXOSC10 CCT6A CCT7 CCT8 CCT3 CCT4 CCT5 DKC1 DCP2 |
| GOBP_REGULATION_OF_PROTEIN_LOCALIZATION_TO_CHROMOSOME_TELOM | 0.00603622 | 0.22284759 | 0.86851338 | 1.71899551 | 2 | 10 | CCT2 CCT6A CCT7 CCT8 CCT3 CCT4 CCT5 GNL3 DKC1 GNL3L |
| GOBP_POSITIVE_REGULATION_OF_CHROMATIN_ORGANIZATION | 0.00603622 | 0.22284759 | 0.62533856 | 1.68507589 | 2 | 36 | [illegible] |
| GOMF_EXONUCLEASE_ACTIVITY_ACTIVE_WITH_EITHER_RIBO_OR_DEOXYRIBC | 0.00603622 | 0.22284759 | 0.6159673 | 1.65976964 | 2 | 36 | [illegible] |
| GOBP_CLEAVAGE_INVOLVED_IN_RRNA_PROCESSING | 0.00612245 | 0.22284759 | 0.78829123 | 1.67645948 | 2 | 13 | EXOSC10 BMS1 EXOSC9 ABT1 KRI1 EXOSC3 BOP1 EXOSC8 |
| GOBP_MATURATION_OF_5_8S_RRNA_FROM_TRICISTRONIC_RRNA_TRANSCRI | 0.00612245 | 0.22284759 | 0.7820068 | 1.6631046 | 2 | 13 | EXOSC10 EXOSC9 ABT1 FTSJ3 KRI1 EXOSC3 BOP1 EXOSC8 |
| GOBP_REGULATION_OF_MEMBRANE_PROTEIN_ECTODOMAIN_PROTEOLYSIS | 0.00613497 | 0.22284759 | 0.87043229 | 1.64482843 | 2 | 8 | FURIN ADRA2A APOE |
| GOBP_PROTEIN_LOCALIZATION_TO_CHROMOSOME_CENTROMERIC_REGION | 0.00614754 | 0.22284759 | 0.78506197 | 1.68982813 | 2 | 14 | BUB3 AURKB CDK1 CENPQ CENPT HASPIN IK BUB1B CTCF |
| GOCC_SPINDLE_MICROTUBULE | 0.00614754 | 0.22284759 | 0.59296841 | 1.65705544 | 2 | 42 | [illegible] |
| GOBP_ALDEHYDE_BIOSYNTHETIC_PROCESS | 0.00618557 | 0.22284759 | 0.8935199 | 1.65291285 | 2 | 7 | BMP6 BMP2 KDM3A |
| GOBP_ENERGY_DERIVATION_BY_OXIDATION_OF_ORGANIC_COMPOUNDS | 0.00618557 | 0.22284759 | 0.46461807 | 1.50308154 | 2 | 96 | [illegible] |
| GOBP_POSITIVE_REGULATION_OF_TELOMERE_MAINTENANCE | 0.00622407 | 0.22284759 | 0.73019628 | 1.74552304 | 2 | 21 | [illegible] |
| GOBP_POSITIVE_REGULATION_OF_CHEMOKINE_PRODUCTION | 0.00625 | 0.22284759 | 0.69023198 | 1.74302362 | 2 | 27 | [illegible] |
| GOBP_REGULATION_OF_ALDOSTERONE_METABOLIC_PROCESS | 0.00626305 | 0.22284759 | 0.93977091 | 1.61917494 | 2 | 5 | BMP6 BMP2 |
| GOMF_PEPTIDE_DISULFIDE_OXIDOREDUCTASE_ACTIVITY | 0.00626305 | 0.22284759 | 0.93201412 | 1.6058104 | 2 | 5 | GLRX GSTO1 GSTO2 GLRX2 |
| GOMF_GLUTATHIONE_DISULFIDE_OXIDOREDUCTASE_ACTIVITY | 0.00626305 | 0.22284759 | 0.93201412 | 1.6058104 | 2 | 5 | GLRX GSTO1 GSTO2 GLRX2 |
| GOBP_PYRIMIDINE_NUCLEOSIDE_DIPHOSPHATE_METABOLIC_PROCESS | 0.00626305 | 0.22284759 | 0.93149665 | 1.60491539 | 2 | 5 | CMPK1 ENTPD7 DTYMK |
| GOBP_LIPOXIN_METABOLIC_PROCESS | 0.00626305 | 0.22284759 | 0.92864598 | 1.60000727 | 2 | 5 | ALOX15B ALOX5AP |
| GOBP_NEGATIVE_REGULATION_OF_INSULIN_SECRETION_INVOLVED_IN_CELL | 0.00626305 | 0.22284759 | 0.92639288 | 1.59612531 | 2 | 5 | JAGN1 KLF7 ENY2 ADRA2A FKBP1B |
| GOMF_CHLORIDE_ION_BINDING | 0.00626305 | 0.22284759 | 0.92628453 | 1.59593862 | 2 | 5 | CTSC LCN1 AMY2A |
| GOBP_MATURATION_OF_5_8S_RRNA | 0.00627615 | 0.22284759 | 0.7816033 | 1.77441485 | 2 | 17 | [illegible] |
| GOBP_RIBOSE_PHOSPHATE_BIOSYNTHETIC_PROCESS | 0.00630252 | 0.22284759 | 0.45801717 | 1.49149029 | 2 | 105 | [illegible] |
| GOMF_TRANSFERASE_ACTIVITY_TRANSFERRING_GLYCOSYL_GROUPS | 0.00631579 | 0.22284759 | 0.4346948 | 1.47681714 | 2 | 135 | [illegible] |
| GOBP_ORGANOPHOSPHATE_METABOLIC_PROCESS | 0.00631579 | 0.22284759 | 0.35132297 | 1.33700283 | 2 | 406 | [illegible] |
| GOBP_RESPONSE_TO_GLYCOPROTEIN | 0.00639659 | 0.22284759 | 0.96810758 | 1.55642283 | 2 | 4 | EDEM2 EDEM1 EDEM3 |
| GOMF_C_X_C_CHEMOKINE_BINDING | 0.00639659 | 0.22284759 | 0.96803222 | 1.55630167 | 2 | 4 | HMGB1 ACKR3 A2M |
| GOBP_PROTEIN_SULFATION | 0.00639659 | 0.22284759 | 0.96672868 | 1.554206 | 2 | 4 | HS3ST3B1 CHST4 |
| GOBP_CYTOKINE_MEDIATED_SIGNALING_PATHWAY | 0.00641026 | 0.22284759 | 0.37275453 | 1.40458104 | 2 | 372 | [illegible] |
| GOBP_UBIQUITIN_DEPENDENT_GLYCOPROTEIN_ERAD_PATHWAY | 0.00642398 | 0.22284759 | 0.97244004 | 1.47717587 | 2 | 3 | EDEM2 EDEM1 EDEM3 |
| GOMF_MANNOSYL_OLIGOSACCHARIDE_1_2_ALPHA_MANNOSIDASE_ACTIVITI | 0.00642398 | 0.22284759 | 0.97244004 | 1.47717587 | 2 | 3 | EDEM2 EDEM1 EDEM3 |
| GOMF_MANNOSIDASE_ACTIVITY | 0.00642398 | 0.22284759 | 0.97244004 | 1.47717587 | 2 | 3 | EDEM2 EDEM1 EDEM3 |
| GOMF_MANNOSYL_OLIGOSACCHARIDE_MANNOSIDASE_ACTIVITY | 0.00642398 | 0.22284759 | 0.97244004 | 1.47717587 | 2 | 3 | EDEM2 EDEM1 EDEM3 |
| GOBP_SPINDLE_ASSEMBLY | 0.00643777 | 0.22284759 | 0.53018615 | 1.55881189 | 2 | 61 | [illegible] |
| GOBP_PROTEIN_LOCALIZATION_TO_MEMBRANE | 0.00736648 | 0.24502802 | -0.3784418 | -1.3577128 | 3 | 232 | [illegible] |
| GOBP_PROTEIN_LOCALIZATION_TO_CELL_PERIPHERY | 0.00749064 | 0.24502802 | -0.4076112 | -1.3868919 | 3 | 146 | [illegible] |
| GOBP_AXON_CHOICE_POINT_RECOGNITION | 0.00750469 | 0.24502802 | -0.9497305 | -1.5270005 | 3 | 4 | GAP43 APP |
| GOBP_HISTAMINE_SECRETION | 0.00750469 | 0.24502802 | -0.9586424 | -1.5413294 | 3 | 4 | ADA |
| GOBP_HISTAMINE_TRANSPORT | 0.00750469 | 0.24502802 | -0.9586424 | -1.5413294 | 3 | 4 | ADA |
| GOBP_POSITIVE_REGULATION_OF_CELL_ADHESION_MOLECULE_PRODUCTION | 0.00750469 | 0.24502802 | -0.9589023 | -1.5417472 | 3 | 6 | FLOT1 CAV1 |
| GOCC_EXTRINSIC_COMPONENT_OF_PRESYNAPTIC_MEMBRANE | 0.00756144 | 0.24502802 | 0.99620342 | 1.33332793 | 3 | 1 | HIP1 |
| GOBP_TISSUE_MORPHOGENESIS | 0.00756144 | 0.24502802 | -0.3682898 | -1.3303001 | 3 | 257 | [illegible] |
| GOBP_EMBRYO_DEVELOPMENT | 0.00757576 | 0.24502802 | -0.3342306 | -1.2676595 | 3 | 417 | [illegible] |
| GOCC_CELL_LEADING_EDGE | 0.00757576 | 0.24502802 | -0.3765303 | -1.3423969 | 3 | 220 | [illegible] |
| GOBP_REGULATION_OF_CALCIUM_MEDIATED_SIGNALING | 0.00759013 | 0.24502802 | -0.6294816 | -1.6254837 | 3 | 28 | FHL2 BST1 ADA ATP2B4 HOMER3 GBP1 CDH13 DYRK2 |
| GOBP_REGENERATION | 0.00761905 | 0.24502802 | -0.5006683 | -1.5904871 | 3 | 91 | [illegible] |
| GOBP_REGULATION_OF_MONOCYTE_DIFFERENTIATION | 0.00763359 | 0.24502802 | -0.7429419 | -1.6715642 | 3 | 16 | JUN CSF1 IL34 CDK6 FES FOXP1 CD74 IRF7 CTNNBIP1 |
| GOCC_CORTICAL_CYTOSKELETON | 0.00764818 | 0.24502802 | -0.5940662 | -1.6976293 | 3 | 46 | [illegible] |
| GOBP_TRANSFORMING_GROWTH_FACTOR_BETA_RECEPTOR_SIGNALING_PAT | 0.00766284 | 0.24502802 | -0.49934 | -1.5545473 | 3 | 86 | [illegible] |
| GOBP_POSITIVE_REGULATION_OF_PROTEIN_LOCALIZATION_TO_CELL_PERIPH | 0.00766284 | 0.24502802 | -0.613898 | -1.5951903 | 3 | 29 | [illegible] |
| GOBP_HEART_MORPHOGENESIS | 0.00767754 | 0.24502802 | -0.4865689 | -1.5316313 | 3 | 87 | [illegible] |
| GOBP_ENZYME_LINKED_RECEPTOR_PROTEIN_SIGNALING_PATHWAY | 0.00772201 | 0.24502802 | -0.3358574 | -1.2901982 | 3 | 498 | [illegible] |
