## Supplementary material for "Adipocytes regulate fibroblast function, and their loss contributes to fibroblast dysfunction in inflammatory diseases": Data file S8

| Data file S8. Statistical information | |
|---|---|
| Figure 1. | B. Healthy n= 8, RA n=3, p=0.001, t=4.813, df=9 |
| Figure 2. | E. DF=44. Healthy n= 10, OA n= 9, RA n= 26, REM n=3. |
| | **APOD**: F=10.98, Healthy vs OA p<0.0001, Healthy vs RA p<0.0001, Healthy vs REM p=0.2637. |
| | **PLIN2**: F=12.70, Healthy vs OA p<0.0001, Healthy vs RA p<0.0001, Healthy v REM p=0.0717. |
| | **CXCL12**: F=6.769, Healthy vs OA p= 0.0056, Healthy vs RA p=0.003, Healthy vs REM p=0.5072 |
| Figure 3. | A. DF=6, **APOD**: t=7.741 p=0.0002 n=4, **NNMT**: t=10.25 p<0.0001 n=4, **CEBPD**: t=4.938 p=0.0026 n=4 |
| | B. DF=4, **APOD**: t=5.740 p=0.0046 n=3, **NNMT**: t=4.410 p=0.0116 n=3, **CEBPD**: t=5.187 p=0.0066 n=3 |
| | C. DF=9, Ctl n=3, ACM n=3, Organic n=3, Inter-phase n=3, Aqueus n=2; |
| | **APOD**: F=55.56, Ctl vs. ACM P<0.0001, Ctl vs. Organic p=0.032, Ctl vs. Inter- phase p=0.7546, Ctl vs. Aqueous p=0.8512; |
| | **NNMT**: F=33.86, Ctl vs. ACM p<0.0001, Ctl vs. Organic p=0.0039, Ctl vs. Inter- phase p=0.8746, Ctl vs. Aqueous p= 0.0487; |
| | **CEBPD**: F=6.310, Ctl vs. ACM p=0.0088, Ctl vs. Organic p=0.4583, Ctl vs. Inter- phase p=0.9967, Ctl vs. Aqueous p=0.1838 |
| | D. DF=23, Ctl n=6, ACM n=3, CHCL3 n=5, Acetone n=5, Methanol n=5, H2O n=5; |
| | **APOD:** F=4.070, Ctl vs. ACM p=0.2206, Ctl vs. CHCL3 p=0.9868, Ctl vs. Acetone p=0.0085, Ctl vs. Methanol p=0.7568, Ctl vs. H2O p=0.9805; |
| | **NNMT:** F=4.665, Ctl vs. ACM p=0.7741, Ctl vs. CHCL3 p=0.9607, Ctl vs. Acetone p=0.0034, Ctl vs. Methanol p=0.9996, Ctl vs. H2O p=0.9997; |
| | **CEBPD:** F=7.261, Ctl vs. ACM p=0.0101, Ctl vs. CHCL3 p=0.9792, Ctl vs. Acetone p=0.0014, Ctl vs. Methanol p=0.9997, Ctl vs. H2O p=0.9921 |
| Figure 4. | A. DF=14, Basal n=3, Differentiation media n=3, ACM n=3, -Dexamethasone n=3, -Indomethacin n=3, -Insulin n=3, -IBMX n=3; |
| | **APOD**: F=12.03, Basal vs. Differentiation media p=0.0002, Basal vs. ACM p<0.0001, Basal vs. -Dexamethasone p=0.9778, Basal vs. -Indomethacin p=0.0204, Basal vs. -Insulin p=0.0367, Basal vs. -IBMX p=0.0283; |
| | **NNMT**: F= 6.561,Basal vs. Differentiation media p=0.1722, Basal vs. ACM p=0.0013, Basal vs. -Dexamethasone p=0.9997, Basal vs. -Indomethacin p=0.3975, Basal vs. -Insulin p=0.2896, Basal vs. -IBMX p=0.9999; |
| | **CEBPD:** F=7.247, Basal vs. Differentiation media p=0.5578, Basal vs. ACM p=0.0015, Basal vs. -Dexamethasone p=0.7883, Basal vs. -Indomethacin p=0.2181, Basal vs. -Insulin p=0.0367, Basal vs. -IBMX p=0.9975 |
| | C. DF=9, Basal n=3, FCM n=3, Organic n=3, Chlor n=2, Acet n=2, Meth n=2, H2O n=1; |
| | **APOD:** F=34.22, Basal vs. FCM p<0.0001, Basal vs. Organic p=0.0038, Basal vs. Chlor p>0.9999, Basal vs. Acet p=0.5766, Basal vs. Meth p=0.698, Basal vs. H2O p=0.9985; |
| | **NNMT**: F=5.005, Basal vs. FCM p=0.0099, Basal vs. Organic p=0.01, Basal vs. Chlor p=0.5769, Basal vs. Acet p=0.0468, Basal vs. Meth p=0.1506, Basal vs. H2O p=0.9961; |
| | **CEBPD**: F= 5.992, Basal vs. FCM p=0.23, Basal vs. Organic p=0.0029, Basal vs. Chlor p=0.9881, Basal vs. Acet p=0.1802, Basal vs. Meth p=0.8879, Basal vs. H2O p=0.9997 |
| | F. DF=4, Basal n=3, Cortisol n=3; **APOD**: t=4.760, p=0.0089; **NNMT**: t=3.182, p=0.0335; **CEBPD**: t=9.430, p=0.0007 |
| | G. DF=8,n=3 |
| | **APOD**: Row Factor: NTC vs GCR KO: F=683.7, P<0.0001; Column Factor: Control vs FCM: F=1173, P<0.0001, Interaction: F=659.4, P<0.0001 |
| | WT:Control vs. WT:FCM: p<0.0001, WT:Control vs. GCR KO:Control p=0.9998, WT:Control vs. GCR KO:FCM p=0.0026, WT:FCM vs. GCR KO:Control p<0.0001, WT:FCM vs. GCR KO:FCM p<0.0001, GCR KO:Control vs. GCR KO:FCM p=0.0018 |
| | **NNMT**: Row Factor: NTC vs GCR KO: F=380.8, P<0.0001; Column Factor: Control vs FCM: F=663.9, p<0.0001; Interaction: F=399.3, P<0.0001 |
| | WT:Control vs. WT:FCM: p<0.0001, WT:Control vs. GCR KO:Control p=0.9866, WT:Control vs. GCR KO:FCM p= 0.0096, WT:FCM vs. GCR KO:Control p<0.0001, WT:FCM vs. GCR KO:FCM p<0.0001, GCR KO:Control vs. GCR KO:FCM p=0.0148 |
| | **CEBPD**: Row Factor: NTC vs GCR KO: F=165.0, P<0.0001; Column Factor: Control vs FCM: F=797.1, p<0.0001; Interaction: F=166.7, P<0.0001 |
| | WT:Control vs. WT:FCM p<0.001, WT:Control vs. GCR KO:Control p>0.9999, WT:Control vs. GCR KO:FCM p<0.0001, WT:FCM vs. GCR KO:Control p<0.0001, WT:FCM vs. GCR KO:FCM p<0.0001, GCR KO:Control vs. GCR KO:FCM p<0.0001 |
| | I. Healthy: n=10, OA: n=9, RA: n=25, REM: n=3. Kruskil-Wallis statistic: Cortisol score: 25.70, FCM score: 19.52, FCM+GCR ant: 15.38. |
| | **Cortisol score:** Healthy vs OA p= 0.0026, Healthy vs RA p<0.0001, Healthy vs REM p>0.9999, OA vs RA p>0.9999, OA vs REM p>0.9999, RA vs REM p= 0.5001. |
| | **FCM score:** Healthy vs OA p= 0.0576, Healthy vs RA p<0.0001, Healthy vs REM p= 0.0769, OA vs RA p>0.9999, OA vs REM p>0.9999, RA vs REM p>0.9999. |
| | **FCM+ GCR ant score**: Healthy vs OA p>0.9999, Healthy vs RA p= 0.0024, Healthy vs REM p= 0.4651, OA vs RA p= 0.0740, OA vs REM p>0.9999, RA vs REM p>0.9999. |
| | J. Ctl n=2, depleted n=2. p= 0.0191. t=7.125, df=2 |
| | K. Ctl n=3, depleted n=4. DF=5, AdipoQ: p= 0.0088, t=4.160, Hsd11B1: p= 0.0220, t=3.277, Pparg: p= 0.0546, t=2.498, Cebpd: p= 0.0559, t=2.479, Plin2: p= 0.0187, t=3.823, Cidec: p= 0.0442, t=2.672 |
| Figure 5. | B. n=28 donors. Kruskil-Wallis statistic: Cortisol score: 18.20, FCM score: 22.85, FCM+GCR ant: 17.86, TGFB score: 15.91. |
| | **Cortisol score p-values:** DDP4 vs CEBPD: p= 0.0009, DDP4 vs FABP4: p= 0.0024, DDP4 vs PPARG: p= 0.3089, CEBPD vs FABP4: p= >0.9999, CEBPD vs PPARG: p= 0.5819, FABP4 vs PPARG: p= 0.9508. |
| | **FCM score p values**: DDP4 vs CEBPD: p<0.0001, DDP4 vs FABP4: p= 0.0723, DDP4 vs PPARG: p= 0.4833, CEBPD vs FABP4: p= 0.1577, CEBPD vs PPARG: p= 0.0361, FABP4 vs PPARG: p>0.9999. |
| | **FCM+ GCR antagonist score p-values:** DDP4 vs CEBPD: p= 0.0006, DDP4 vs FABP4: p> 0.9999, DDP4 vs PPARG: p> 0.9999, CEBPD vs FABP4: p= 0.0295, CEBPD vs PPARG: p= 0.0085, FABP4 vs PPARG: p>0.9999. |
| | **TGFB score p-values:** DDP4 vs CEBPD: p> 0.9999, DDP4 vs FABP4: p= 0.0031, DDP4 vs PPARG: p= 0.1774, CEBPD vs FABP4: p= 0.0111, CEBPD vs PPARG: p= 0.3978, FABP4 vs PPARG: p> 0.9999 |
| | C. Healthy n= 10, OA n= 9, RA n= 25, REM n=3. DF: 43, |
| | **DPP4+ progenitor:** F= 2.850, Healthy vs OA p-value= 0.6917, Healthy vs RA p-value= 0.3918, Healthy vs REM p-value= 0.4229, OA vs RA p-value= 0.9953, OA vs REM p-value= 0.1190, RA vs REM p-value= 0.0543, |
| | **CEBPD+ PreAd:** F= 2.944, Healthy vs OA p-value= 0.844, Healthy vs RA p-value= 0.1273, Healthy vs REM p-value= 0.8367, OA vs RA p-value= 0.6353, OA vs REM p-value= 0.5112, RA vs REM p-value= 0.1195, |
| | **FABP4+ PreAd**: F= 6.090, Healthy vs OA p-value= 0.1307, Healthy vs RA p-value= 0.2790, Healthy vs REM p-value= 0.1033, OA vs RA p-value= 0.8106, OA vs REM p-value= 0.0021, RA vs REM p-value= 0.0039, |
| | **VIT+Areg:** F= 2.710, Healthy vs OA p-value= 0.9493, Healthy vs RA p-value= 0.0621, Healthy vs REM p-value= 0.9136, OA vs RA p-value= 0.2663, OA vs REM p-value= 0.9926, RA vs REM p-value= 0.8179 |
| Figure 6. | A. n=3, DF=8, F=6.841. Basal vs TNFa p-value= 0.0208, Basal vs TNFa+FCM p-value= 0.9796, Basal vs TNFa+cortisol p-value= 0.9999, TNFa vs TNFa+FCM p-value= 0.0353, |
| | TNFa vs TNFa+cortisol p-value= 0.0228, TNFa+FCM vs TNFa+cortisol p-value= 0.9882 |
| | B. FLS n=5, TGFB n=8, TGFB+cortisol n=8. DF=18, F=8.805. FLS vs TGFB p-value= 0.0149, FLS vs TGFB+cortisol p-value= 0.9630, TGFB vs TGFB+cortisol p-value= 0.0030 |
| | C. n=3 DF= 18, |
| | **APOD**: F= 48.39, Cortisol vs Basal p< 0.0001, Cortisol vs Cortisol+TNFa p= 0.0086, Cortisol vs Cortisol +TGFB p<0.0001, Cortisol vs Cortisol+IL17 p= 0.9875, Cortisol vs Cortisol+ IL1B p= 0.9863, |
| | Cortisol vs Cortisol+IFNy p<0.0001, Cortisol vs Cortisol +TNFa+IFNy p<0.0001, Cortisol vs Cortisol+TNFa+IL17 p<0.0001, |
| | **CEBPD**: F= 47.54, Cortisol vs Basal p< 0.0001, Cortisol vs Cortisol+TNFa p>0.9999, Cortisol vs Cortisol +TGFB p< 0.0001, Cortisol vs Cortisol+IL17 p= 0.0183, Cortisol vs Cortisol+ IL1B p= 0.9834, |
| | Cortisol vs Cortisol+IFNy p= 0.9920, Cortisol vs Cortisol +TNFa+IFNy p= 0.3960, Cortisol vs Cortisol+TNFa+IL17 p= 0.9994, |
| | **PLIN2**: F= 62.38, Cortisol vs Basal p= 0.0091, Cortisol vs Cortisol+TNFa p= 0.0771, Cortisol vs Cortisol +TGFB p< 0.0001, Cortisol vs Cortisol+IL17 p= 0.0445, Cortisol vs Cortisol+ IL1B p= 0.0035, |
| | Cortisol vs Cortisol+IFNy p< 0.0001, Cortisol vs Cortisol +TNFa+IFNy p< 0.0001, Cortisol vs Cortisol+TNFa+IL17 p= 0.3853 |
| | D. n=3, DF=8, |
| | **ADIPOQ**: Row Factor: basal vs GCR ant: F=8.919, P=0.0174; Column Factor: basal vs ADM: F=8.923, P=0.0174, Interaction: F=8.92, P=0.0174 |
| | WT:Basal vs. WT:ADM: -=0.0124, WT:Basal vs. Antagonist:Basa:l p>0.9999, WT:Basal vs. Antagonist:ADM: p>0.9999, WT:ADM vs. Antagonist:Basal: p=0.0124, WT:ADM vs. Antagonist:ADM: p=0.0124, Antagonist:Basal vs. Antagonist:ADM p>0.9999 |
| | **FABP4**: Row Factor: basal vs GCR ant: F=207.1, P<0.0001; Column Factor: basal vs ADM: F=207.4, P<0.0001; Interaction: F=207.4, P<0.0001 |
| | WT:Basal vs. WT:ADM p<0.0001, WT:Basal vs. Antagonist:Basal p>0.9999, WT:Basal vs. Antagonist:ADM p>0.9999, WT:ADM vs. Antagonist:Basal p<0.0001, WT:ADM vs. Antagonist:ADM p<0.0001, Antagonist:Basal vs. Antagonist:ADM p>0.9999 |
| | **LEPTIN**: Row Factor: basal vs GCR ant: F=6.689, P=0.0323; Column Factor: basal vs ADM: F=21.59, P=0.0017; Interaction: F=7.818, P=0.0233 |
| | WT:Basal vs. WT:ADM: p=0.0034, WT:Basal vs. Antagonist:Basal: p=0.9987, WT:Basal vs. Antagonist:ADM: p=0.5026, WT:ADM vs. Antagonist:Basal: p=0.0040, WT:ADM vs. Antagonist:ADM: p=0.0216, Antagonist:Basal vs. Antagonist:ADM: p=0.5830 |
| | **PPARG**: Row Factor: basal vs GCR ant: F=30.80, P=0.0005; Column Factor: basal vs ADM: F=30.38, P=0.0006; Interaction: F=41.52, P=0.0002 |
| | WT:Basal vs. WT:ADM: p=0.0001, WT:Basal vs. Antagonist:Basal: p=0.9187, WT:Basal vs. Antagonist:ADM: p>0.9999, WT:ADM vs. Antagonist:Basal: p=0.0002, WT:ADM vs. Antagonist:ADM: p=0.0001, Antagonist:Basal vs. Antagonist:ADM: p=0.9095 |
| | E. n=6, DF=20, F= 37.05, FLS+ADM vs FLS p<0.0001, FLS+ADM vs FLS+ADM+TGFB p<0.0001, FLS+ADM vs FLS+ADM+TNFa+IFNy p= 0.0088 |
