## Supplementary material for "Adipocytes regulate fibroblast function, and their loss contributes to fibroblast dysfunction in inflammatory diseases": Source data

| | | |
|---|---|---|
| **Subfigure:** | **Figure 1B** | |
| Plot name: | Total lipidtox+ requant | |
| **Groups:** | Healthy | RA |
| **Data points:** | 73.3 | 71.5 |
| | 87.9 | 44.2 |
| | 92.5 | 30.83 |
| | 82.7 | |
| | 90.5 | |
| | 83.2 | |
| | 92.4 | |
| | 83.3 | |

| | |
|---|---|
| **Subfigure** | **Figure 2E** |
| **Plot name** | No steroid users Fibroblast symphony redo fibroblast proportions |

| Plot name: | **APOD** | | | |
|---|---|---|---|---|
| **Groups:** | Healthy | OA | RA | REM |
| **Data points:** | 0.200787 | 0.003881 | 0.017857 | 0.089367 |
| | 0.083333 | 0.005195 | 0.00201 | 0.057143 |
| | 0.059172 | 0.003367 | 0 | 0.032699 |
| | 0.03064351 | 0.000296 | 0.007092 | |
| | 0.078947 | 0 | 0.003119 | |
| | 0.111111 | 0.064391 | 0.035714 | |
| | 0.085766 | 0.003522 | 0.027778 | |
| | 0.316867 | 0.008485 | 0.00641 | |
| | 0.035828 | 0.002681 | 0.005716 | |
| | 0.06746032 | | 0.001666 | |
| | | | 0.044304 | |
| | | | 0 | |
| | | | 0.057143 | |
| | | | 0 | |
| | | | 0.025385 | |
| | | | 0.053333 | |
| | | | 0.030626 | |
| | | | 0.040816 | |
| | | | 0 | |
| | | | 0.008467 | |
| | | | 0.017391 | |
| | | | 0.000951 | |
| | | | 0.006092 | |
| | | | 0.044444 | |
| | | | 0.075758 | |
| | | | 0.00813 | |

| Plot name: | **PLIN2** | | | |
|---|---|---|---|---|
| **Groups:** | Healthy | OA | RA | REM |
| **Data points:** | 0.240157 | 0.002687 | 0 | 0.102376 |
| | 0.008681 | 0 | 0.005025 | 0.005357 |
| | 0.025148 | 0 | 0 | 0 |
| | 0.02247191 | 0.003852 | 0.003546 | |
| | 0.026316 | 0.016611 | 0 | |
| | 0.245791 | 0.003103 | 0 | |
| | 0.122263 | 0.000271 | 0 | |
| | 0.189132 | 0.006667 | 0.003205 | |
| | 0.044586 | 0.002681 | 0.008406 | |

| | | | |
|---|---|---|---|
| | 0.09656085 | | 0.006108 |
| | | | 0.012658 |
| | | | 0 |
| | | | 0 |
| | | | 0 |
| | | | 0.011786 |
| | | | 0.013333 |
| | | | 0.011984 |
| | | | 0 |
| | | | 0 |
| | | | 0.008467 |
| | | | 0.008696 |
| | | | 0.001427 |
| | | | 0.004352 |
| | | | 0.022222 |
| | | | 0 |
| | | | 0.00813 |

| Plot name: | **CXCL12** | | | |
|---|---|---|---|---|
| **Groups:** | Healthy | OA | RA | REM |
| **Data points:** | 0.232283 | 0.450746 | 0.348214 | 0.244344 |
| | 0.175347 | 0.246753 | 0.630151 | 0.558036 |
| | 0.152367 | 0.713805 | 0.53159 | 0.060367 |
| | 0.08273749 | 0.206815 | 0.585106 | |
| | 0.197368 | 0.255814 | 0.372141 | |
| | 0.148148 | 0.403413 | 0.535714 | |
| | 0.054745 | 0.2246 | 0.777778 | |
| | 0.048694 | 0.795152 | 0.775641 | |
| | 0.093949 | 0.689008 | 0.403497 | |
| | 0.01851852 | | 0.673515 | |
| | | | 0.607595 | |
| | | | 0.666667 | |
| | | | 0.028571 | |
| | | | 0.628571 | |
| | | | 0.572983 | |
| | | | 0.153333 | |
| | | | 0.054594 | |
| | | | 0.653061 | |
| | | | 0.576923 | |
| | | | 0.469941 | |
| | | | 0.356522 | |
| | | | 0.103186 | |
| | | | 0.848564 | |
| | | | 0.377778 | |
| | | | 0.174242 | |
| | | | 0.162602 | |

| | | |
|---|---|---|
| **Subfigure** | **Figure 3A** | |
| Plot name: | **APOD FCM** | |
| **Groups:** | Ctl24hr | FCM 24hrs |
| **Data points:** | 0.539979 | 33.08554 |
| | 1.851923 | 36.28757 |
| | 1.022474 | 24.84947 |
| | 0.927824 | 20.88724 |
| Plot name: | **CEBPD FCM** | |
| **Groups:** | Ctl24hr | FCM 24hrs |
| **Data points:** | 0.622427 | 5.950986 |
| | 1.606615 | 4.520253 |
| | 0.811458 | 3.108383 |
| | 1.431651 | 3.794082 |
| Plot name: | **NNMT FCM** | |
| **Groups:** | Ctl24hr | FCM 24hrs |
| **Data points:** | 0.903425 | 3.799024 |
| | 1.106899 | 3.768273 |
| | 1.355412 | 2.98176 |
| | 0.805345 | 3.246198 |

| | | | | |
|---|---|---|---|---|
| **Subfigure** | **Figure 3B** | *ACM APOD* | | |
| Plot name | **APOD 24hr** | | | |
| **Groups** | F+pre-adipoc | F+D10 ACM | Lipid(25um C | Adipocyte |
| **Data points:** | 0.712715 | 26.76263 | 1.808841 | 44.89339 |
| | 0.946219 | 47.19433 | 1.453424 | 61.96739 |
| | 1.482833 | 33.15777 | 1.604197 | 36.8182 |

| | | |
|---|---|---|
| Plot name | **APOD** | |
| **Groups** | Basal | ACM |
| **Data points:** | 0.712715 | 26.76263 |
| | 0.946219 | 47.19433 |
| | 1.482833 | 33.15777 |

| | | | | |
|---|---|---|---|---|
| **Subfigure** | **Figure 3B** | *ACM CEBPD* | | |
| Plot name | **CEBPD 24hr** | | | |
| **Groups** | F+pre-adipoc | F+D10 ACM | Lipid(25um C | Adipocyte |
| **Data points:** | 0.97585 | 33.7188 | 2.777179 | 2.898269 |
| | 0.805848 | 21.85071 | 1.997244 | 6.321196 |
| | 1.27164 | 43.07787 | 1.959392 | 2.987849 |

| | | |
|---|---|---|
| Plot name | **CEBPD** | |
| **Groups** | Basal | ACM |
| **Data points:** | 0.97585 | 33.7188 |
| | 0.805848 | 21.85071 |
| | 1.27164 | 43.07787 |

| | | | | |
|---|---|---|---|---|
| **Subfigure** | **Figure 3B** | *ACM NNMT* | | |
| Plot name | **NNMT 24hr** | | | |
| **Groups** | F+pre-adipoc | F+D10 ACM | Lipid(25um C | Adipocyte |
| **Data points:** | 0.382818 | 41.36908 | 4.023972 | 4.987896 |
| | 1.417727 | 31.99132 | 2.91402 | 12.02894 |
| | 1.842532 | 66.43516 | 4.393607 | 4.414797 |

| | | |
|---|---|---|
| Plot name | **NNMT** | |
| **Groups** | Basal | ACM |
| **Data points:** | 0.382818 | 41.36908 |
| | 1.417727 | 31.99132 |
| | 1.842532 | 66.43516 |

| | | | | | |
|---|---|---|---|---|---|
| **Subfigure** | **Figure 3C** | Bligh and dyer sep | | | |
| Plot name | APOD | | | | |
| **Groups:** | Ctl | ACM | Organic | Inter-phase | Aqueous |
| **Data points:** | 1.048717 | 7.226886 | 2.286 | 0.489909 | 0.891 |
| | 1.039106 | 5.273952 | 2.708 | 0.56008 | 1.889 |
| | 0.917659 | 6.646367 | 2.325 | 0.688368 | |

| | | | | | |
|---|---|---|---|---|---|
| **Subfigure** | **Figure 3C** | Bligh and dyer sep | | | |
| Plot name | CEBPD | | | | |
| **Groups:** | Ctl | ACM | Organic | Inter-phase | Aqueous |
| **Data points:** | 1.03907 | 5.540594 | 2.978 | 0.116005 | 11.09 |
| | 1.084485 | 17.5762 | 5.914 | 0.203115 | 6.04 |
| | 0.887425 | 19.41679 | 7.967 | 0.215823 | |

| | | | | | |
|---|---|---|---|---|---|
| **Subfigure** | **Figure 3C** | Bligh and dyer sep | | | |
| Plot name | NNMT | | | | |
| **Groups:** | Ctl | ACM | Organic | Inter-phase | Aqueous |
| **Data points:** | 1.139678 | 12.89173 | 6.138 | 2.251032 | 5.499 |
| | 1.47462 | 10.47504 | 5.054 | 1.417849 | 2.914 |
| | 0.595029 | 9.142356 | 5.25 | 1.625916 | |

**Subfigure:** **Figure 3D** *Organic sep ACM*

Plot name: **APOD organic sep ACM**

| **Groups:** | Ctl | ACM | CHCL3 | Acetone | Methanol | H2O |
|---|---|---|---|---|---|---|
| **Data points:** | 1.188302 | 4.511553 | 0.9843619 | 6.8873225 | 1.7407621 | 1.3813961 |
| | 0.907954 | 1.728345 | 0.6916277 | 3.7536616 | 1.3612762 | 1.0985119 |
| | 0.92685 | 1.67464 | 2.956695 | 4.093943 | 1.782197 | 0.299696 |
| | 1.20612 | | 1.256146 | 1.407925 | 2.111086 | 0.202303 |
| | 0.886731 | | 0.859671 | 1.303763 | 1.797065 | 0.192638 |
| | 0.935013 | | | | | |

Plot name: **CEBPD organic sep ACM**

| **Groups:** | Ctl | ACM | CHCL3 | Acetone | Methanol | H2O |
|---|---|---|---|---|---|---|
| **Data points:** | 1.243065 | 11.34217 | 0.9442631 | 7.2548371 | 1.1807627 | 0.9566924 |
| | 0.93025 | 5.203173 | 0.465892 | 3.7797704 | 0.662026 | 0.8153834 |
| | 0.864781 | 4.769543 | 4.892772 | 15.56418 | 3.3527 | 0.089203 |
| | 0.269254 | | 2.122363 | 5.46506 | 1.978836 | 0.503917 |
| | 1.645169 | | 1.52999 | 5.185017 | 0.405369 | 0.458703 |
| | 2.257498 | | | | | |

Plot name: **NNMT organic sep ACM**

| **Groups:** | Ctl | ACM | CHCL3 | Acetone | Methanol | H2O |
|---|---|---|---|---|---|---|
| **Data points:** | 0.952732 | 7.55837 | 1.1661163 | 12.281323 | 2.1349021 | 1.0285258 |
| | 1.049613 | 2.777607 | 0.8873151 | 5.1257016 | 1.0602545 | 1.0606888 |
| | 3.053308 | 3.336423 | 7.536973 | 25.87507 | 4.209523 | 0.956987 |
| | 2.276108 | | 4.955411 | 11.98843 | 2.334292 | 1.064068 |
| | 0.143892 | | 0.921372 | 3.346568 | 0.787659 | 0.442567 |

| | | | | |
|---|---|---|---|---|
| **Subfigure** | Figure 3E | | | |
| **Plot name:** | Arachidonic acid | | | |
| | **Groups:** | **Data points:** | | |
| | CHCl3 | 407 | 1598 | 1022 |
| | Acetone | 1753820 | 1270749 | 1780757 |
| | MeOH | 23633 | 21504 | 24541 |
| **Plot name:** | Indo | | | |
| | **Groups:** | **Data points:** | | |
| | CHCl3 | 417 | 1015 | 492 |
| | Acetone | 760935 | 656364 | 879211 |
| | MeOH | 25075 | 34275 | 38352 |
| **Plot name:** | IBMX | | | |
| | **Groups:** | **Data points:** | | |
| | CHCl3 | 16728 | 12004 | 15204 |
| | Acetone | 147379300 | 138565100 | 155154400 |
| | MeOH | 8051786 | 7473690 | 7850672 |
| **Plot name:** | DEX | | | |
| | **Groups:** | **Data points:** | | |
| | CHCl3 | 796 | 0 | 185 |
| | Acetone | 49951 | 34555 | 49832 |
| | MeOH | 1912 | 1349 | 1164 |

**Subfigure:** **Figure 4A** *Media minus 1*

Plot name: **APOD media minus 1**

| **Groups:** | Basal | Differentiatio | ACM | -Dexamethas | -Indomethaci | -Insulin | -IBMX |
|---|---|---|---|---|---|---|---|
| **Data points:** | 0.995447 | 3.579025 | 2.962632 | 1.507455 | 2.930434 | 2.185781 | 2.084882 |
| | 0.947527 | 3.594167 | 3.499974 | 1.161469 | 2.191092 | 2.513392 | 2.305251 |
| | 1.060207 | 4.062736 | 5.403734 | 1.134645 | 2.499039 | 2.498874 | 2.995404 |

Plot name: **CEBPD media minus 1**

| **Groups:** | Basal | Differentiatio | ACM | -Dexamethas | -Indomethaci | -Insulin | -IBMX |
|---|---|---|---|---|---|---|---|
| **Data points:** | 1.061975 | 1.477236 | 2.362334 | 0.3586 | 2.076923 | 1.94731 | 1.163361 |
| | 1.010852 | 1.295933 | 2.446429 | 0.837571 | 1.438917 | 1.929155 | 0.968119 |
| | 0.931533 | 2.107869 | 4.554491 | 0.404882 | 2.257469 | 2.361349 | 1.381355 |

Plot name: **NNMT media minus 1**

| **Groups:** | Basal | Differentiatio | ACM | -Dexamethas | -Indomethaci | -Insulin | -IBMX |
|---|---|---|---|---|---|---|---|
| **Data points:** | 1.155419 | 2.907071 | 3.882214 | 1.285309 | 3.774053 | 2.975535 | 1.37075 |
| | 0.756409 | 2.33862 | 4.618243 | 0.986883 | 2.871192 | 3.041189 | 0.910622 |
| | 1.144205 | 4.264722 | 8.657537 | 0.090386 | 1.275774 | 2.542553 | 0.46903 |

| **Subfigure** | **Figure 4C** | Organic separation FCM |
|---|---|---|

| Plot name: | **APOD** | | | | | | |
|---|---|---|---|---|---|---|---|
| **Groups:** | Basal | FCM | Organic | Chlor | Acet | Meth | H2O |
| **Data points:** | 1.040972 | 10.43423 | 6.149834 | 1.198433 | 2.423436 | 2.24639 | 1.328755 |
| | 1.025129 | 10.67942 | 3.979882 | 0.894809 | 1.939626 | 1.795963 | |
| | 0.937092 | 8.266852 | 3.772838 | | | | |

| Plot name: | **CEBPD** | | | | | | |
|---|---|---|---|---|---|---|---|
| **Groups:** | Basal | FCM | Organic | Chlor | Acet | Meth | H2O |
| **Data points:** | 0.776253 | 2.770064 | 9.140127 | 1.371875 | 3.685994 | 2.76563 | 0.680693 |
| | 1.081079 | 3.490485 | 5.480822 | 1.885114 | 3.695523 | 1.333074 | |
| | 1.191625 | 3.443808 | 4.323851 | | | | |

| Plot name: | **NNMT** | | | | | | |
|---|---|---|---|---|---|---|---|
| **Groups:** | Basal | FCM | Organic | Chlor | Acet | Meth | H2O |
| **Data points:** | 1.829266 | 5.965263 | 7.254524 | 2.452889 | 4.692599 | 4.410499 | 1.831986 |
| | 1.758248 | 5.598708 | 4.503539 | 3.080118 | 4.481904 | 3.18897 | |
| | 0.310916 | 4.015756 | 3.807868 | | | | |

**Subfigure:** **Figure 4E**

Plot name: **Cortisol Quantification (ng/mL)**

| **Groups:** | pre-adipocyte | D1 | D2 | D3 | gout | 7289L | 7289R |
|---|---|---|---|---|---|---|---|
| **Data points:** | 0.335991 | 0.375745 | 0.359181 | 0.506001 | 0.256827 | 3.344631 | 2.267108 |
| | 0.599495 | 0.548643 | 0.487051 | 0.519485 | 0.425621 | 3.538595 | 2.349119 |
| | 0.225931 | 0.301378 | 0.308581 | 0.537278 | 0.42836 | 6.046057 | 2.540101 |
| | 0.321297 | 0.450858 | 0.402784 | 0.565238 | 0.403655 | 3.376738 | 2.284757 |
| | 0.232113 | 0.41057 | | 0.733494 | 0.349566 | 3.387092 | 2.517541 |

| | | |
|---|---|---|
| **Subfigure** | **Figure 4F** | *cort 2.5ng* |
| Plot name: | **APOD cort 2.5ng** | |
| **Groups:** | Basal | Cortisol2.5ng/mL |
| **Data points:** | 0.985411 | 2.562156 |
| | 0.84972 | 3.079003 |
| | 1.194281 | 2.012513 |
| Plot name: | **CEBPD cort 2.5ng** | |
| **Groups:** | Basal | Cortisol2.5ng/mL |
| **Data points:** | 0.821954 | 2.714509 |
| | 1.034113 | 2.960409 |
| | 1.17648 | 2.449235 |
| Plot name: | **NNMT cort 2.5ng** | |
| **Groups:** | Basal | Cortisol2.5ng/mL |
| **Data points:** | 1.054918 | 2.686432 |
| | 1.22128 | 2.043152 |
| | 0.776187 | 1.593646 |

| | | | | | | | |
|---|---|---|---|---|---|---|---|
| **Subfigure** | **Figure 4G** | | | | | | |
| Plot name: | **APOD GCR KO** | | | | | | |
| **Groups:** | NTC | NTC+FCM | GCRKO | GCRKO+FCM | | | |
| **Data points:** | 1.083704 | 48.25122 | 0.700997 | 7.326604 | | | |
| | 1.384909 | 48.79603 | 0.679473 | 8.249307 | | | |
| | 0.666297 | 44.32448 | 0.674142 | 6.237942 | | | |
| Plot name: | **APOD** | | | | | | |
| **Groups:** | | Control | | | FCM | | |
| **Data points:** | WT | 1.083704 | 1.384909 | 0.666297 | 48.25122 | 48.79603 | 44.32448 |
| | GCR KO | 0.700997 | 0.679473 | 0.674142 | 7.326604 | 8.249307 | 6.237942 |
| Plot name: | **CEBPD GCR KO** | | | | | | |
| **Groups:** | NTC | NTC+FCM | GCRKO | GCRKO+FCM | | | |
| **Data points:** | 0.824145 | 31.50467 | 0.981106 | 13.81482 | | | |
| | 1.045933 | 28.41719 | 1.062518 | 10.26223 | | | |
| | 1.160091 | 29.44665 | 1.125935 | 11.2439 | | | |
| Plot name: | **NNMT GCR KO** | | | | | | |
| **Groups:** | NTC | NTC+FCM | GCRKO | GCRKO+FCM | | | |
| **Data points:** | 0.544991 | 15.35703 | 1.231839 | 2.422469 | | | |
| | 1.519002 | 15.15819 | 1.302082 | 2.903124 | | | |
| | 1.207959 | 14.05844 | 1.160267 | 3.591044 | | | |

**Subfigure** **Figure 4I** Bulk Module Scores

Plot name: **No steroid users Cortisol module score**

| **Groups:** | Healthy | OA | RA | REM |
|---|---|---|---|---|
| **Data points:** | 0.098041 | 0.02006787 | 0.04139838 | 0.08087492 |
| | 0.157596 | 0.01685633 | 0.03326525 | 0.10893103 |
| | 0.224444 | 0.08935383 | 0.06981858 | 0.0582495 |
| | 0.165339 | 0.07572551 | 0.05568622 | |
| | 0.164132 | 0.05415646 | 0.1007513 | |
| | 0.119049 | 0.05841005 | 0.05950265 | |
| | 0.220185 | 0.05906615 | 0.05091152 | |
| | 0.185089 | 0.06553337 | 0.02997531 | |
| | 0.175247 | 0.0443005 | 0.04803191 | |
| | 0.119264 | | 0.03444974 | |
| | | | 0.03042602 | |
| | | | 0.05669758 | |
| | | | 0.06538121 | |
| | | | 0.03908128 | |
| | | | 0.04248803 | |
| | | | 0.07230956 | |
| | | | 0.05563602 | |
| | | | 0.04967355 | |
| | | | 0.04325817 | |
| | | | 0.02923332 | |
| | | | 0.0606451 | |
| | | | 0.063729 | |
| | | | 0.04782548 | |
| | | | 0.02275634 | |
| | | | 0.05763688 | |

Plot name: **No steroid users FCM module score**

| **Groups:** | Healthy | OA | RA | REM |
|---|---|---|---|---|
| **Data points:** | 0.042612 | 0.013514 | 0.039306 | 0.048597 |
| | 0.079728 | 0.021433 | 0.02152 | 0.061443 |
| | 0.107215 | 0.072122 | 0.035299 | 0.008997 |
| | 0.130702 | 0.062943 | 0.037669 | 0.048597 |
| | 0.060483 | 0.051723 | 0.040454 | 0.061443 |
| | 0.097929 | 0.038336 | 0.012735 | 0.008997 |
| | 0.113681 | 0.044075 | 0.035756 | |
| | 0.088338 | 0.042037 | 0.012836 | |
| | 0.102388 | 0.034204 | 0.05034 | |
| | 0.05432 | | 0.011763 | |
| | | | 0.024919 | |
| | | | 0.067296 | |
| | | | 0.068819 | |
| | | | 0.036204 | |

| | | | |
|---|---|---|---|
| | | | 0.007126 |
| | | | 0.045732 |
| | | | 0.059743 |
| | | | 0.02155 |
| | | | 0.029938 |
| | | | 0.027385 |
| | | | 0.04056 |
| | | | 0.037882 |
| | | | 0.028345 |
| | | | 0.028608 |
| | | | 0.013144 |

| Plot name: | **No steroid users FCM+GCR ant module score** | | | |
|---|---|---|---|---|
| **Groups:** | Healthy | OA | RA | REM |
| **Data points:** | 0.03561 | 0.018263 | 0.032832 | 0.039441 |
| | 0.053041 | 0.032631 | 0.028078 | 0.044054 |
| | 0.05589 | 0.07529 | 0.027789 | 0.003798 |
| | 0.097697 | 0.060648 | 0.031405 | |
| | 0.027374 | 0.060879 | 0.024709 | |
| | 0.085124 | 0.036424 | 0.005616 | |
| | 0.083739 | 0.052403 | 0.042326 | |
| | 0.050756 | 0.0402 | 0.005688 | |
| | 0.096811 | 0.051665 | 0.051409 | |
| | 0.037771 | | 0.007594 | |
| | | | 0.026944 | |
| | | | 0.069087 | |
| | | | 0.052479 | |
| | | | 0.035255 | |
| | | | -0.01119 | |
| | | | 0.030705 | |
| | | | 0.050531 | |
| | | | 0.014575 | |
| | | | 0.017329 | |
| | | | 0.009829 | |
| | | | 0.033978 | |
| | | | 0.04727 | |
| | | | 0.026348 | |
| | | | 0.032792 | |
| | | | 0.005514 | |

| | | |
|---|---|---|
| **Subfigure** | **Figure 4J** | Adiposoft measurements |
| Plot name | **Adipocyte cell number** | |
| **Groups** | CTL | Depleted |
| **Data points:** | 163.5 | 1 |
| | 123.5 | 1 |

**Subfigure** **Figure 4K**

8wk IA 50ng diphtheria depletion qPCR

| Plot name: | **CEBPD** | | |
|---|---|---|---|
| **Groups:** | Ctl | Depleted | CL |
| **Data points:** | 0.79267 | 0.763739 | 0.472598 |
| | 0.989643 | 0.660842 | 0.478329 |
| | 1.274762 | 0.736326 | 0.507234 |
| | | 0.726303 | 0.786588 |

| Plot name: | **AdipoQ** | | |
|---|---|---|---|
| **Groups:** | Ctl | Depleted | CL |
| **Data points:** | 1.099203 | 0.54443 | 0.650829 |
| | 0.735423 | 0.307582 | 0.775254 |
| | 1.237042 | 0.501765 | 0.606 |
| | | 0.286243 | 0.63303 |

| Plot name: | **11HSD1** | | |
|---|---|---|---|
| **Groups:** | Ctl | Depleted | CL |
| **Data points:** | 0.921118 | 0.374443 | 0.465016 |
| | 0.732879 | 0.374761 | 0.520417 |
| | 1.481332 | 0.438328 | 0.571328 |
| | | 0.489816 | 0.568545 |

| Plot name: | **Plin2** | | |
|---|---|---|---|
| **Groups:** | Ctl | Depleted | CL |
| **Data points:** | 1.200849 | 0.591702 | 0.618973 |
| | 1.006427 | 0.483107 | 0.702387 |
| | 0.827426 | 0.627045 | 0.534028 |

| Plot name: | **Pparg** | |
|---|---|---|
| **Groups:** | Ctl | Depleted |
| **Data points:** | 1.158604 | 0.843083 |
| | 1.001787 | 0.646165 |
| | 0.861568 | 0.871162 |
| | | 0.715111 |

| Plot name: | **Cidec** | |
|---|---|---|
| **Groups:** | Ctl | Depleted |
| **Data points:** | 0.913233 | 0.422029 |
| | 1.734167 | 0.295584 |
| | 0.631433 | 0.324813 |
| | | 0.351819 |

**Subfigure** **Figure 5B** Module Scoring Adipose Atlas

Plot name: **Fixed cortisol score**

| **Groups:** | DPP4+ | CEBPD+ | FABP4+ | PPARG+ |
|---|---|---|---|---|
| **Data points:** | 0.133384 | 0.158012 | 0.147343 | 0.150762 |
| | 0.164576 | 0.202193 | 0.196663 | 0.206588 |
| | 0.125692 | 0.149444 | 0.153714 | 0.129794 |
| | 0.106082 | 0.145943 | 0.144834 | 0.073641 |
| | 0.085665 | 0.129602 | 0.12955 | 0.020381 |
| | 0.099837 | 0.134459 | 0.141568 | 0.089097 |
| | 0.099325 | 0.167337 | 0.161138 | 0.052334 |
| | 0.123102 | 0.143467 | 0.164191 | 0.10424 |
| | 0.066472 | 0.127291 | 0.08586 | 0.083276 |
| | 0.081434 | 0.122055 | 0.119969 | 0.087175 |
| | 0.138064 | 0.142373 | 0.119299 | 0.080441 |
| | 0.106922 | 0.16492 | 0.143881 | 0.106755 |
| | 0.043695 | 0.154775 | 0.146334 | 0.117865 |
| | 0.105008 | 0.177974 | 0.167199 | 0.180846 |
| | 0.071274 | 0.182243 | 0.177224 | 0.206944 |
| | 0.136767 | 0.179091 | 0.172013 | 0.17672 |
| | 0.076979 | 0.116079 | 0.134681 | 0.226389 |
| | 0.056734 | 0.112085 | 0.158202 | 0.190998 |
| | 0.140848 | 0.189116 | 0.174647 | 0.231153 |
| | 0.158416 | 0.221451 | 0.199583 | 0.185761 |
| | 0.158144 | 0.209607 | 0.183929 | 0.19099 |
| | 0.183299 | 0.228991 | 0.223517 | 0.207344 |
| | 0.14078 | 0.175682 | 0.19049 | 0.149222 |
| | 0.189871 | 0.240035 | 0.226269 | |
| | 0.168191 | 0.184286 | 0.195507 | |
| | 0.115842 | 0.182856 | 0.190046 | |
| | 0.164663 | 0.205428 | 0.193101 | |
| | 0.151761 | 0.172209 | 0.151048 | |

Plot name: **Fiixed FCM score**

| **Groups:** | DPP4+ | CEBPD+ | FABP4+ | PPARG+ |
|---|---|---|---|---|
| **Data points:** | 0.08536 | 0.117251 | 0.083536 | 0.104191 |
| | 0.094522 | 0.126627 | 0.09028 | 0.114543 |
| | 0.071534 | 0.103251 | 0.075017 | 0.058297 |
| | 0.049776 | 0.076615 | 0.081708 | 0.027548 |
| | 0.044564 | 0.089856 | 0.079777 | 0.009197 |
| | 0.039221 | 0.065481 | 0.054973 | 0.080445 |
| | 0.045021 | 0.10343 | 0.07997 | 0.030607 |
| | 0.055058 | 0.073677 | 0.070958 | 0.048663 |
| | 0.04016 | 0.078228 | 0.063356 | 0.060957 |
| | 0.050357 | 0.06668 | 0.072843 | 0.072335 |

| | | | |
|---|---|---|---|
| 0.048966 | 0.079372 | 0.047542 | 0.058051 |
| 0.085482 | 0.137897 | 0.112067 | 0.053295 |
| 0.031231 | 0.088747 | 0.07134 | 0.040885 |
| 0.0502 | 0.084647 | 0.067421 | 0.118145 |
| 0.059569 | 0.101629 | 0.084753 | 0.101463 |
| 0.090562 | 0.144333 | 0.129221 | 0.088062 |
| 0.05657 | 0.087378 | 0.085952 | 0.112882 |
| 0.010129 | 0.067684 | 0.07608 | 0.105418 |
| 0.098645 | 0.1402 | 0.096449 | 0.106622 |
| 0.084505 | 0.120587 | 0.100318 | 0.117291 |
| 0.091312 | 0.13363 | 0.094648 | 0.103964 |
| 0.087436 | 0.15286 | 0.116602 | 0.121699 |
| 0.073052 | 0.173575 | 0.098721 | 0.096435 |
| 0.094988 | 0.118583 | 0.108828 | |
| 0.100926 | 0.165509 | 0.115022 | |
| 0.055105 | 0.111676 | 0.104955 | |
| 0.12643 | 0.176755 | 0.120265 | |
| 0.065379 | 0.12371 | 0.091459 | |

| Plot name: | **Fixed FCM + GCR ant score** | | | |
|---|---|---|---|---|
| **Groups:** | DPP4+ | CEBPD+ | FABP4+ | PPARG+ |
| **Data points:** | 0.069479 | 0.093257 | 0.055848 | 0.079139 |
| | 0.082565 | 0.103382 | 0.063192 | 0.091991 |
| | 0.054572 | 0.085749 | 0.045054 | 0.03722 |
| | 0.039095 | 0.066822 | 0.060447 | 0.047148 |
| | 0.038901 | 0.073118 | 0.054621 | 0.0000182 |
| | 0.022257 | 0.032798 | 0.030847 | 0.053956 |
| | 0.035789 | 0.073535 | 0.060042 | 0.02734 |
| | 0.043679 | 0.050837 | 0.050536 | 0.01187 |
| | 0.044653 | 0.067922 | 0.041884 | 0.038445 |
| | 0.040237 | 0.062622 | 0.058694 | 0.059859 |
| | 0.043596 | 0.059062 | 0.027938 | 0.010318 |
| | 0.077917 | 0.128801 | 0.091918 | 0.028421 |
| | 0.027365 | 0.065027 | 0.047367 | 0.003883 |
| | 0.004435 | 0.062231 | 0.039377 | 0.108367 |
| | 0.063155 | 0.067913 | 0.059946 | 0.0689 |
| | 0.052365 | 0.107902 | 0.084933 | 0.074943 |
| | 0.038833 | 0.074021 | 0.06927 | 0.078273 |
| | 0.014027 | 0.04091 | 0.035576 | 0.076618 |
| | 0.096038 | 0.127751 | 0.07308 | 0.069426 |
| | 0.061041 | 0.087126 | 0.072475 | 0.092654 |
| | 0.086541 | 0.120073 | 0.083282 | 0.093244 |
| | 0.063587 | 0.11821 | 0.078866 | 0.08636 |
| | 0.05802 | 0.155475 | 0.071593 | 0.054413 |
| | 0.07119 | 0.079875 | 0.071454 | |
| | 0.086665 | 0.155612 | 0.082642 | |
| | 0.062155 | 0.096 | 0.090038 | |

| | | | |
|---|---|---|---|
| | 0.115833 | 0.159754 | 0.082374 |
| | 0.051031 | 0.102206 | 0.074397 |

| Plot name: | **Fixed TGFB score** | | | |
|---|---|---|---|---|
| **Groups:** | DPP4+ | CEBPD+ | FABP4+ | PPARG+ |
| **Data points:** | 0.097401 | 0.085044 | 0.058212 | 0.103494 |
| | 0.108027 | 0.10244 | 0.064321 | 0.119149 |
| | 0.099007 | 0.075387 | 0.050174 | 0.020819 |
| | 0.075281 | 0.07759 | 0.063411 | 0.032147 |
| | 0.073547 | 0.063735 | 0.045113 | 0.021924 |
| | 0.074418 | 0.054402 | 0.056767 | 0.065451 |
| | 0.097143 | 0.083805 | 0.074467 | 0.031413 |
| | 0.07916 | 0.089769 | 0.061292 | 0.059709 |
| | 0.091757 | 0.079802 | 0.063525 | 0.070182 |
| | 0.083349 | 0.072087 | 0.073061 | 0.037679 |
| | 0.077399 | 0.061404 | 0.04705 | 0.039164 |
| | 0.090869 | 0.114389 | 0.073277 | 0.038469 |
| | 0.053728 | 0.054181 | 0.049861 | 0.013009 |
| | 0.055251 | 0.095613 | 0.044946 | 0.125357 |
| | 0.075057 | 0.08655 | 0.045701 | 0.114474 |
| | 0.085704 | 0.10995 | 0.069203 | 0.10069 |
| | 0.083614 | 0.059789 | 0.069408 | 0.098862 |
| | 0.059192 | 0.069433 | 0.058297 | 0.117439 |
| | 0.157288 | 0.143911 | 0.091483 | 0.104017 |
| | 0.129872 | 0.113696 | 0.09825 | 0.137088 |
| | 0.134321 | 0.144184 | 0.088869 | 0.107051 |
| | 0.119258 | 0.126628 | 0.085373 | 0.119614 |
| | 0.135307 | 0.16051 | 0.090578 | 0.063909 |
| | 0.13438 | 0.097595 | 0.088432 | |
| | 0.160326 | 0.16949 | 0.112126 | |
| | 0.117837 | 0.118641 | 0.085917 | |
| | 0.183401 | 0.159535 | 0.106587 | |
| | 0.099009 | 0.080058 | 0.057729 | |

**Subfigure** **Figure 5C** Symphony prop adipose

Plot name: **No steroid users FABP4 + prop**

| **Groups:** | Healthy | OA | RA | REM |
|---|---|---|---|---|
| **Data points:** | 0.244094 | 0.04091 | 0.046296 | 0.036765 |
| | 0.065972 | 0.013575 | 0.033777 | 0.106696 |
| | 0.093195 | 0.009375 | 0.051173 | 0.314409 |
| | 0.019408 | 0.003219 | 0.055385 | |
| | 0.039474 | 0.022013 | 0.032226 | |
| | 0.082492 | 0.053603 | 0.146341 | |
| | 0.096715 | 0.015502 | 0.052632 | |
| | 0.025406 | 0.0408614 | 0.076923 | |
| | 0.027866 | 0.002398 | 0.031897 | |
| | 0.048942 | | 0.048725 | |
| | | | 0.017751 | |
| | | | 0 | |
| | | | 0.037037 | |
| | | | 0.02381 | |
| | | | 0.088346 | |
| | | | 0.034682 | |
| | | | 0.019004 | |
| | | | 0.02 | |
| | | | 0 | |
| | | | 0.008489 | |
| | | | 0 | |
| | | | 0.009743 | |
| | | | 0.081315 | |
| | | | 0.020408 | |
| | | | 0.06135 | |

Plot name: **No steroid users VIT + prop**

| **Groups:** | Healthy | OA | RA | REM |
|---|---|---|---|---|
| **Data points:** | 0.007874 | 0.019778 | 0.046296 | 0.003959 |
| | 0.006944 | 0.002262 | 0.071648 | 0.079464 |
| | 0.001479 | 0.015625 | 0.185501 | 0.014373 |
| | 0.001021 | 0.001341 | 0.058462 | |
| | 0 | 0 | 0.04431 | |
| | 0.003367 | 0.028399 | 0.219512 | |
| | 0.00365 | 0.005485 | 0.184211 | |
| | 0 | 0.09828824 | 0.384615 | |
| | 0.001592 | 0.007194 | 0.007805 | |
| | 0 | | 0.057224 | |
| | | | 0.04142 | |
| | | | 0 | |
| | | | 0 | |

| | | | |
|---|---|---|---|
| | | | 0.02381 |
| | | | 0.081767 |
| | | | 0 |
| | | | 0.005898 |
| | | | 0.08 |
| | | | 0.085714 |
| | | | 0.011885 |
| | | | 0.015385 |
| | | | 0.007529 |
| | | | 0.127163 |
| | | | 0 |
| | | | 0.006135 |

| Plot name: | **No steroid users CEBPD+** | | | |
|---|---|---|---|---|
| **Groups:** | Healthy | OA | RA | REM |
| **Data points:** | 0.413386 | 0.245191 | 0.314815 | 0.649887 |
| | 0.076389 | 0.165158 | 0.413511 | 0.472768 |
| | 0.720414 | 0.63125 | 0.400853 | 0.662594 |
| | 0.689479 | 0.570279 | 0.48 | |
| | 0.355263 | 0.427673 | 0.23565 | |
| | 0.774411 | 0.264821 | 0.219512 | |
| | 0.324818 | 0.190794 | 0.421053 | |
| | 0.864502 | 0.60022087 | 0.297659 | |
| | 0.561306 | 0.697842 | 0.332542 | |
| | 0.140212 | | 0.34051 | |
| | | | 0.538462 | |
| | | | 0.533333 | |
| | | | 0.037037 | |
| | | | 0.380952 | |
| | | | 0.307331 | |
| | | | 0.127168 | |
| | | | 0.193316 | |
| | | | 0.36 | |
| | | | 0.371429 | |
| | | | 0.588285 | |
| | | | 0.476923 | |
| | | | 0.124889 | |
| | | | 0.439446 | |
| | | | 0.244898 | |
| | | | 0.190184 | |

| Plot name: | **No steroid users DPP4+** | | | |
|---|---|---|---|---|
| **Groups:** | Healthy | OA | RA | REM |
| **Data points:** | 0.334646 | 0.694121 | 0.592593 | 0.309389 |
| | 0.850694 | 0.819005 | 0.481064 | 0.341071 |
| | 0.184911 | 0.34375 | 0.362473 | 0.008624 |
| | 0.290092 | 0.425161 | 0.406154 | |

| | | |
|---|---|---|
| 0.605263 | 0.550314 | 0.687815 |
| 0.139731 | 0.653177 | 0.414634 |
| 0.574818 | 0.788218 | 0.342105 |
| 0.110092 | 0.26062949 | 0.240803 |
| 0.409236 | 0.292566 | 0.627757 |
| 0.810847 | | 0.553541 |
| | | 0.402367 |
| | | 0.466667 |
| | | 0.925926 |
| | | 0.571429 |
| | | 0.522556 |
| | | 0.83815 |
| | | 0.781782 |
| | | 0.54 |
| | | 0.542857 |
| | | 0.391341 |
| | | 0.507692 |
| | | 0.857839 |
| | | 0.352076 |
| | | 0.734694 |
| | | 0.742331 |

| | | | | |
|---|---|---|---|---|
| **Subfigure** | **Figure 6A** | | | |
| Plot name | **MMP3** | | | |
| **Groups:** | Basal | TNFa | TNFa+FCM | TNFa+cortisol |
| **Data points:** | 1.361975 | 24.64536 | 1.839665 | 2.325829 |
| | 0.949269 | 10.36904 | 2.003845 | 0.540395 |
| | 0.773467 | 9.077676 | 3.334641 | 0.929604 |

| | | | |
|---|---|---|---|
| **Subfigure** | **Figure 6B** | | |
| **Plot name** | Col1a1 edge intensity | | |
| **Groups** | FLS | TGFB | TGFB+cortisol |
| **Data points:** | 4.999 | 8.296 | 8.213 |
| | 7.314 | 7.732 | 8.214 |
| | 5.399 | 8.923 | 4.333 |
| | 5.611 | 14.727 | 3.078 |
| | 5.252 | 9.653 | 4.904 |
| | | 12.059 | 4.923 |
| | | 13.753 | 4.162 |
| | | 4.931 | 5.03 |

| | |
|---|---|
| **Subfigure** | **Figure 6C** |

| | | | | | | | |
|---|---|---|---|---|---|---|---|
| Plot name | **APOD** | | | | | | |
| **Groups:** | Basal | Cortisol100nl | Cortisol+TNFa | Cortisol+TGFI | Cortisol+IL17 | Cortisol+IL1B | Cortisol+IFNy |
| **Data points:** | 0.861414 | 5.20451 | 4.561561 | 1.08095 | 6.11667 | 5.876499 | 3.174562 |
| | 1.08921 | 6.012408 | 4.725494 | 1.005819 | 6.60156 | 5.67909 | 3.480355 |
| | 1.065802 | 6.769653 | 4.261138 | 1.004148 | 4.518144 | 5.66985 | 3.164754 |

| | | | | |
|---|---|---|---|---|
| Plot name | **CEBPD** | | | |
| **Groups:** | Cortisol+IL1B | Cortisol+IFNy | Cortisol+TNFa | Cortisol+TNFa+IL17 |
| **Data points:** | 2.560851 | 2.231825 | 2.0885 | 2.236528 |
| | 1.955214 | 2.093477 | 2.110232 | 2.482398 |
| | 2.077686 | 2.311389 | 1.874029 | 2.027507 |

| | | | | |
|---|---|---|---|---|
| Plot name | **PLIN2** | | | |
| **Groups:** | Cortisol+IL1B | Cortisol+IFNy | Cortisol+TNFa | Cortisol+TNFa+IL17 |
| **Data points:** | 1.477635 | 0.826388 | 0.760033 | 1.373567 |
| | 1.589963 | 0.636207 | 0.757369 | 1.493193 |
| | 1.725057 | 0.626518 | 0.848396 | 1.381632 |

| Cortisol+TNFa | Cortisol+TNFa+IL17 |
|---|---|
| 2.622241 | 3.98504 |
| 2.909427 | 3.530978 |
| 2.86687 | 3.289893 |

**Subfigure:** **Figure 6d**

| Plot name | **AdipoQ** | | | |
|---|---|---|---|---|
| **Groups:** | Basal | GCR ant | ADM | ADM+GCR ant |
| **Data points:** | 1.155582 | 2.233045 | 9115.767 | 5.813415 |
| | 1.088201 | 1.304428 | 36976.13 | 1.59311 |
| | 0.795225 | 1.127955 | 36660.78 | 5.662678 |

| Plot name | **FABP4** | | | |
|---|---|---|---|---|
| **Groups:** | Basal | GCR ant | ADM | ADM+GCR ant |
| **Data points:** | 1.08319 | 8.031553 | 6386.523 | 1.980758 |
| | 1.02003 | 1.222711 | 6332.751 | 1.493309 |
| | 0.90507 | 2.838988 | 7782.258 | 7.003865 |

| Plot name | **Leptin** | | | |
|---|---|---|---|---|
| **Groups:** | Basal | GCR ant | ADM | ADM+GCR ant |
| **Data points:** | 0.587212 | 2.192149 | 8.18216 | 1.073795 |
| | 1.885933 | 1.099426 | 5.94897 | 1.898921 |
| | 0.902982 | 0.573173 | 6.590022 | 5.20471 |

| Plot name | **PPARG** | | | |
|---|---|---|---|---|
| **Groups:** | Basal | GCR ant | ADM | ADM+GCR ant |
| **Data points:** | 1.352179 | 2.251176 | 13.36543 | 0.865168 |
| | 0.75451 | 3.021121 | 19.04823 | 1.120011 |
| | 0.980169 | 1.759758 | 23.42651 | 0.935073 |

| | | | | |
|---|---|---|---|---|
| **Subfigure** | **Figure 6E** | | | |
| **Plot name** | PLIN2 staining | | | |
| **Groups:** | FLS | FLS+ADM | FLS+ADM+TG | FLS+ADM+TNFa+IFNγ |
| **Data points:** | 1.120439 | 1.599452 | 0.698989 | 1.312176 |
| | 1.076238 | 1.331163 | 0.651705 | 1.448187 |
| | 1.007367 | 1.477129 | 0.746274 | 1.487047 |
| | 1.065959 | 1.471989 | 0.586945 | 1.365285 |
| | 0.818229 | 1.854377 | 0.825424 | 1.079016 |

**Subfigure:** Figure 1D
**Plot Name:** Fixed Cell Proportions

| | Healthy | | | | |
|---|---|---|---|---|---|
| **Sublining Fibr** | 0.327586 | 0.857664 | 0.696855 | 0.906484 | 0.251043 |
| **Lining fibrobla** | 0.126437 | 0.10219 | 0.049057 | 0.034913 | 0.685675 |
| **Aterial EC** | 0.114943 | 0.00365 | 0.011321 | 0.004364 | 0.002782 |
| **Venous EC** | 0.218391 | 0.007299 | 0.157233 | 0.041771 | 0.031293 |
| **Mural** | 0.034483 | 0.007299 | 0.038994 | 0.006234 | 0.019471 |
| **MERTK+ Macr** | 0.028736 | 0 | 0.021384 | 0.003741 | 0.002782 |
| **IL1B+ Macrop** | 0.04023 | 0.00365 | 0.002516 | 0 | 0 |
| **CD4+ T cell** | 0.005747 | 0.018248 | 0.01761 | 0.001247 | 0.004172 |
| **CD8+ T cell** | 0.017241 | 0 | 0.001258 | 0 | 0 |
| **NK cell** | 0.057471 | 0 | 0.001258 | 0 | 0.000695 |
| **B cell** | 0.022989 | 0 | 0 | 0 | 0 |
| **Plasma Cell** | 0 | 0 | 0.002516 | 0.001247 | 0.002086 |
| **Proliferating** | 0.005747 | 0 | 0 | 0 | 0 |

| | | | | | | |
|---|---|---|---|---|---|---|
| | | | | | | **OA** |
| 0.922424 | 0.521437 | 0.202297 | 0.797688 | 0.828452 | | 0.2072 |
| 0.029091 | 0.230591 | 0.715548 | 0.056647 | 0.026499 | | 0.37024 |
| 0.009697 | 0.100811 | 0.007951 | 0.031214 | 0.006974 | | 0.03397 |
| 0.027879 | 0.079954 | 0.028269 | 0.061272 | 0.107392 | | 0.09851 |
| 0.007273 | 0.017381 | 0.007067 | 0.028902 | 0.027894 | | 0.02582 |
| 0 | 0.00927 | 0.016784 | 0.015029 | 0 | | 0.02649 |
| 0 | 0.017381 | 0.00265 | 0.002312 | 0 | | 0.07337 |
| 0.002424 | 0.011587 | 0.009717 | 0.004624 | 0.002789 | | 0.05842 |
| 0 | 0.005794 | 0.004417 | 0.001156 | 0 | | 0.03329 |
| 0 | 0.005794 | 0.001767 | 0 | 0 | | 0.00747 |
| 0 | 0 | 0 | 0 | 0 | | 0.05231 |
| 0.001212 | 0 | 0.003534 | 0.001156 | 0 | | 0.01223 |
| 0 | 0 | 0 | 0 | 0 | | 0.00068 |

| | | | | | | |
|---|---|---|---|---|---|---|
| 0.054 | 0.08777 | 0.18294 | 0.1247 | 0.26209 | 0.05631 | 0.21922 |
| 0.144 | 0.01567 | 0.71044 | 0.03357 | 0.11268 | 0.32758 | 0.02532 |
| 0.182 | 0.05643 | 0.01122 | 0.04077 | 0.01408 | 0.04932 | 0.01841 |
| 0.466 | 0.12853 | 0.02806 | 0.20384 | 0.13533 | 0.05179 | 0.06444 |
| 0.008 | 0.01254 | 0.01347 | 0.00959 | 0.00735 | 0.02137 | 0.00173 |
| 0.078 | 0.20219 | 0.02806 | 0.56595 | 0.09124 | 0.15125 | 0.15478 |
| 0.048 | 0.30721 | 0.02357 | 0.01679 | 0.04348 | 0.12043 | 0.02244 |
| 0 | 0.10031 | 0.00112 | 0.0024 | 0.08696 | 0.08015 | 0.18872 |
| 0.004 | 0.03448 | 0 | 0 | 0.07655 | 0.04973 | 0.06847 |
| 0.016 | 0.02351 | 0 | 0.0024 | 0.04593 | 0.04809 | 0.01611 |
| 0 | 0.02821 | 0.00112 | 0 | 0.10839 | 0.03617 | 0.19505 |
| 0 | 0.00157 | 0 | 0 | 0.00857 | 0 | 0.01726 |
| 0 | 0.00157 | 0 | 0 | 0.00735 | 0.00781 | 0.00806 |

| | **RA** | | | | |
|---|---|---|---|---|---|
| 0.22662 | 0.056818 | 0.275986 | 0.046181 | 0.129338 | 0.060893 |
| 0.1295 | 0.002841 | 0.172043 | 0.063943 | 0 | 0.138024 |
| 0.02158 | 0.025568 | 0.010753 | 0.010657 | 0.028391 | 0.010825 |
| 0.11151 | 0.039773 | 0.02509 | 0.005329 | 0.056782 | 0.033829 |
| 0.01799 | 0.005682 | 0 | 0 | 0.003155 | 0.014885 |
| 0.15468 | 0.664773 | 0.0681 | 0.01421 | 0.227129 | 0.116373 |
| 0.29496 | 0.065341 | 0.050179 | 0.095915 | 0.07571 | 0.177267 |
| 0.0036 | 0.065341 | 0.150538 | 0.349911 | 0.217666 | 0.177267 |
| 0 | 0.025568 | 0.146953 | 0.097691 | 0.078864 | 0.143437 |
| 0.01079 | 0.036932 | 0.021505 | 0.040853 | 0.044164 | 0.056834 |
| 0.01799 | 0.008523 | 0.028674 | 0.213144 | 0.082019 | 0.037889 |
| 0 | 0 | 0.050179 | 0.031972 | 0.041009 | 0.006766 |
| 0.01079 | 0.002841 | 0 | 0.030195 | 0.015773 | 0.02571 |

| | | | | | | |
|---|---|---|---|---|---|---|
| 0.012448 | 0.041237 | 0.038667 | 0.452442 | 0.168397 | 0.111765 | 0.055556 |
| 0.002075 | 0 | 0.008 | 0.131105 | 0.136101 | 0.011765 | 0.018519 |
| 0.006224 | 0.010309 | 0.005333 | 0.025707 | 0.021915 | 0.017647 | 0 |
| 0 | 0.051546 | 0.034667 | 0.069409 | 0.025375 | 0.035294 | 0.092593 |
| 0.008299 | 0.020619 | 0 | 0 | 0.001153 | 0.005882 | 0 |
| 0.06639 | 0.268041 | 0.4 | 0.134961 | 0.351788 | 0.182353 | 0.018519 |
| 0.060166 | 0.041237 | 0.069333 | 0.01671 | 0.10842 | 0.223529 | 0.222222 |
| 0.278008 | 0.237113 | 0.149333 | 0.028278 | 0.071511 | 0.182353 | 0.111111 |
| 0.29668 | 0.216495 | 0.118667 | 0.020566 | 0.036909 | 0.052941 | 0.074074 |
| 0.033195 | 0.020619 | 0.016 | 0.011568 | 0.014994 | 0.117647 | 0.185185 |
| 0.155602 | 0.030928 | 0.045333 | 0.014139 | 0.028835 | 0.052941 | 0.092593 |
| 0.053942 | 0.041237 | 0.096 | 0.09383 | 0.031142 | 0 | 0.074074 |
| 0.026971 | 0.020619 | 0.018667 | 0.001285 | 0.00346 | 0.005882 | 0.055556 |

| | | | | | | |
|---|---|---|---|---|---|---|
| 0.222222 | 0.153846 | 0.117834 | 0 | 0 | 0.056122 | 0.539846 |
| 0 | 0.076923 | 0.084395 | 0 | 0 | 0 | 0 |
| 0 | 0 | 0.017516 | 0 | 0.075472 | 0.035714 | 0.061697 |
| 0.055556 | 0.115385 | 0.047771 | 0 | 0.037736 | 0.05102 | 0.293059 |
| 0 | 0 | 0.006369 | 0 | 0 | 0.002551 | 0.002571 |
| 0 | 0.038462 | 0.025478 | 0.037037 | 0.037736 | 0.02551 | 0.07455 |
| 0.055556 | 0 | 0.049363 | 0 | 0.018868 | 0.09949 | 0.017995 |
| 0.5 | 0.230769 | 0.194268 | 0.518519 | 0.226415 | 0.211735 | 0.005141 |
| 0 | 0.153846 | 0.078025 | 0.074074 | 0.301887 | 0.216837 | 0 |
| 0.111111 | 0.192308 | 0.148089 | 0.037037 | 0.188679 | 0.045918 | 0.005141 |
| 0 | 0.038462 | 0.203822 | 0.259259 | 0.09434 | 0.084184 | 0 |
| 0 | 0 | 0.015924 | 0.074074 | 0.018868 | 0.038265 | 0 |
| 0.055556 | 0 | 0.011146 | 0 | 0 | 0.132653 | 0 |

| | | | | | | |
|---|---|---|---|---|---|---|
| 0.225806 | 0.027397 | 0.208279 | 0.117021 | 0.046225 | 0.196203 | 0.01046 |
| 0.096774 | 0.013699 | 0.007762 | 0.010638 | 0.46379 | 0.044304 | 0 |
| 0.225806 | 0.041096 | 0.002587 | 0.053191 | 0.003082 | 0.023734 | 0.01046 |
| 0.096774 | 0.041096 | 0.028461 | 0.042553 | 0.016949 | 0.044304 | 0.043933 |
| 0 | 0 | 0.002587 | 0.021277 | 0.004622 | 0.004747 | 0.002092 |
| 0.032258 | 0.315068 | 0.006468 | 0.340426 | 0.234206 | 0.098101 | 0.449791 |
| 0.064516 | 0.136986 | 0.021992 | 0.138298 | 0.032357 | 0.007911 | 0.064854 |
| 0.064516 | 0.164384 | 0.232859 | 0.085106 | 0.117103 | 0.143987 | 0.110879 |
| 0.064516 | 0.191781 | 0.349288 | 0.138298 | 0.029276 | 0.060127 | 0.186192 |
| 0.064516 | 0.041096 | 0.075032 | 0.010638 | 0.020031 | 0.023734 | 0.058577 |
| 0.064516 | 0.027397 | 0.049159 | 0 | 0.016949 | 0.121835 | 0.041841 |
| 0 | 0 | 0.01423 | 0.010638 | 0.015408 | 0.205696 | 0.01046 |
| 0 | 0 | 0.001294 | 0.031915 | 0 | 0.025316 | 0.01046 |

| | |
|---|---|
| 0.111675 | 0.293478 |
| 0 | 0.119565 |
| 0.030457 | 0.01087 |
| 0.096447 | 0.076087 |
| 0.035533 | 0 |
| 0.147208 | 0.217391 |
| 0.106599 | 0.01087 |
| 0.248731 | 0.108696 |
| 0.101523 | 0.021739 |
| 0.091371 | 0.01087 |
| 0.030457 | 0.076087 |
| 0 | 0.054348 |
| 0 | 0 |

**Subfigure:** Figure 2C

Plot Name: **CD34 Proportions**

| **Groups:** | Healthy | OA | RA | REM |
|---|---|---|---|---|
| **Data Points:** | 0.165354 | 0 | 0 | 0.149321 |
| | 0.057292 | 0 | 0.001005 | 0.038393 |
| | 0.02071 | 0 | 0 | 0.023715 |
| | 0.02962206 | 0 | 0.021277 | |
| | 0.065789 | 0 | 0 | |
| | 0.144781 | 0.046936 | 0 | |
| | 0.20438 | 0 | 0 | |
| | 0.1729 | 0 | 0.003205 | |
| | 0.026274 | 0 | 0.011096 | |
| | 0.28439153 | | 0.00111 | |
| | | | 0.012658 | |
| | | | 0.222222 | |
| | | | 0.742857 | |
| | | | 0 | |
| | | | 0.000907 | |
| | | | 0.313333 | |
| | | | 0.032623 | |
| | | | 0 | |
| | | | 0 | |
| | | | 0.018628 | |
| | | | 0 | |
| | | | 0 | |
| | | | 0 | |
| | | | 0.266667 | |
| | | | 0.340909 | |
| | | | 0.02439 | |

Plot Name: **DKK3 Proportions**

| **Groups:** | Healthy | OA | RA | REM |
|---|---|---|---|---|
| **Data Points:** | 0.066929 | 0.038209 | 0.616071 | 0.158937 |
| | 0.399306 | 0.083117 | 0.057286 | 0.142857 |
| | 0.276627 | 0.074074 | 0.019608 | 0.378369 |
| | 0.07252298 | 0.009481 | 0.379433 | |
| | 0.328947 | 0.428571 | 0.028067 | |
| | 0.20202 | 0.221877 | 0.357143 | |
| | 0.485401 | 0.048496 | 0.055556 | |
| | 0.221595 | 0.106061 | 0.105769 | |
| | 0.102707 | 0.018767 | 0.44889 | |
| | 0.49603175 | | 0.131038 | |
| | | | 0.240506 | |
| | | | 0.111111 | |
| | | | 0.114286 | |
| | | | 0.257143 | |

| | | | |
|---|---|---|---|
| | | | 0.079782 |
| | | | 0.466667 |
| | | | 0.863515 |
| | | | 0.122449 |
| | | | 0.115385 |
| | | | 0.480102 |
| | | | 0.608696 |
| | | | 0.019496 |
| | | | 0.043516 |
| | | | 0.266667 |
| | | | 0.393939 |
| | | | 0.544715 |

| Plot Name: | **PRG4 Proportions** | | | |
|---|---|---|---|---|
| **Groups:** | Healthy | OA | RA | REM |
| **Data Points:** | 0.07874 | 0.481493 | 0.008929 | 0.236991 |
| | 0.267361 | 0.644156 | 0.299497 | 0.179018 |
| | 0.446746 | 0.208754 | 0.422658 | 0.503054 |
| | 0.53932584 | 0.743704 | 0 | |
| | 0.236842 | 0.289037 | 0.582121 | |
| | 0.136364 | 0.252133 | 0.071429 | |
| | 0.047445 | 0.707667 | 0.138889 | |
| | 0.040226 | 0.069091 | 0.067308 | |
| | 0.633758 | 0.268097 | 0.112307 | |
| | 0.02777778 | | 0.172127 | |
| | | | 0.044304 | |
| | | | 0 | |
| | | | 0.057143 | |
| | | | 0.114286 | |
| | | | 0.296464 | |
| | | | 0 | |
| | | | 0.006658 | |
| | | | 0.183673 | |
| | | | 0.153846 | |
| | | | 0.013548 | |
| | | | 0.008696 | |
| | | | 0.840704 | |
| | | | 0.087032 | |
| | | | 0.022222 | |
| | | | 0.015152 | |
| | | | 0.235772 | |

| Plot Name: | **VCAM1 Proportions** | | | |
|---|---|---|---|---|
| **Groups:** | Healthy | OA | RA | REM |
| **Data Points:** | 0.015748 | 0.022985 | 0.008929 | 0.018665 |
| | 0.008681 | 0.020779 | 0.005025 | 0.019196 |
| | 0.019231 | 0 | 0.026144 | 0.001797 |

| | | |
|---|---|---|
| 0.2226762 | 0.035852 | 0.003546 |
| 0.065789 | 0.009967 | 0.014553 |
| 0.011785 | 0.008146 | 0 |
| 0 | 0.015443 | 0 |
| 0.010586 | 0.014545 | 0.038462 |
| 0.062898 | 0.018767 | 0.010087 |
| 0.00925926 | | 0.014436 |
| | | 0.037975 |
| | | 0 |
| | | 0 |
| | | 0 |
| | | 0.012693 |
| | | 0 |
| | | 0 |
| | | 0 |
| | | 0.153846 |
| | | 0.000847 |
| | | 0 |
| | | 0.034237 |
| | | 0.010444 |
| | | 0 |
| | | 0 |
| | | 0.01626 |

**Subfigure:** Figure 3A

| | | | | |
|---|---|---|---|---|
| Plot Name: | **APOD 24hrs** | | | |
| **Groups:** | Vehicle | 10um | 25um | 50um |
| **Data Points:** | 0.310808 | 1.866668 | 1.808841 | 1.416998 |
| | 1.782699 | 1.716304 | 1.453424 | 1.633529 |
| | 1.804805 | 1.639781 | 1.604197 | 0.356226 |

| | | |
|---|---|---|
| Plot Name: | **CEBPD 24hrs** | |
| **Groups:** | Basal | 25um |
| **Data Points:** | 0.97585 | 2.777179 |
| | 0.805848 | 1.997244 |
| | 1.27164 | 1.959392 |

| | | |
|---|---|---|
| Plot Name: | **NNMT 24hrs** | |
| **Groups:** | Basal | 25um |
| **Data Points:** | 0.382818 | 4.023972 |
| | 1.417727 | 2.91402 |
| | 1.842532 | 4.393607 |

**Subfigure:** Figure 3B

| | | |
|---|---|---|
| Plot Name: | **APOD** | |
| **Groups:** | Basal | FCMAbd |
| **Data Points:** | 0.985411 | 5.45717 |
| | 0.84972 | 5.951404 |
| | 1.194281 | 5.501386 |

| | | |
|---|---|---|
| Plot Name: | **CEBPD** | |
| **Groups:** | Basal | FCMAbd |
| **Data Points:** | 0.821954 | 4.018024 |
| | 1.034113 | 3.90836 |
| | 1.17648 | 4.251961 |

| | | |
|---|---|---|
| Plot Name: | **NNMT** | |
| **Groups:** | Basal | FCMAbd |
| **Data Points:** | 1.054918 | 3.583791 |
| | 1.22128 | 3.559226 |
| | 0.776187 | |

**Subfigure:** Figure 3C

Plot Name: **APOD**

| **Groups:** | Basal | Aq | Aq Aq | AqOrg |
| --- | --- | --- | --- | --- |
| **Data Points:** | 1.341617 | 3.840194 | 0 | 1.099872 |
| | 0.919385 | 2.348831 | 0 | 3.111517 |
| | 0.810726 | | 0 | 1.02644 |

Plot Name: **CEBPD**

| **Groups:** | Basal | Aq | Aq Aq | AqOrg |
| --- | --- | --- | --- | --- |
| **Data Points:** | 0.730678 | 2.88854 | 0 | 2.02184 |
| | 1.199201 | 5.203357 | 0 | 5.96264 |
| | 1.141254 | | 0 | 2.752959 |

Plot Name: **NNMT**

| **Groups:** | Basal | Aq | Aq Aq | AqOrg |
| --- | --- | --- | --- | --- |
| **Data Points:** | 2.220124 | 4.283171 | 0 | 0.961929 |
| | 0.917263 | 4.577128 | 0 | 6.539973 |
| | 0.491054 | | 0 | 3.029997 |

**Subfigure:** Figure 4A

| Plot Name: | **APOD** | | | | | |
|---|---|---|---|---|---|---|
| **Groups:** | Basal | FCM | Organic | Chlor | Acet | Meth | H2O |
| **Data Points:** | 1.040972 | 10.43423 | 6.149834 | 1.198433 | 2.423436 | 2.24639 | 1.328755 |
| | 1.025129 | 10.67942 | 3.979882 | 0.894809 | 1.939626 | 1.795963 | |
| | 0.937092 | 8.266852 | 3.772838 | | | | |

| Plot Name: | **CEBPD** | | | | | | |
|---|---|---|---|---|---|---|---|
| **Groups:** | Basal | FCM | Organic | Chlor | Acet | Meth | H2O |
| **Data Points:** | 0.776253 | 2.770064 | 9.140127 | 1.371875 | 3.685994 | 2.76563 | 0.680693 |
| | 1.081079 | 3.490485 | 5.480822 | 1.885114 | 3.695523 | 1.333074 | |
| | 1.191625 | 3.443808 | 4.323851 | | | | |

| Plot Name: | **NNMT** | | | | | | |
|---|---|---|---|---|---|---|---|
| **Groups:** | Basal | FCM | Organic | Chlor | Acet | Meth | H2O |
| **Data Points:** | 1.829266 | 5.965263 | 7.254524 | 2.452889 | 4.692599 | 4.410499 | 1.831986 |
| | 1.758248 | 5.598708 | 4.503539 | 3.080118 | 4.481904 | 3.18897 | |
| | 0.310916 | 4.015756 | 3.807868 | | | | |

**Subfigure:** Figure 4B

Plot Name: **APOD**

| | | | | | |
|---|---|---|---|---|---|
| **Groups:** | Basal | Syn FCM | SynFCM | AbdomFCMD3 | AbdomFCMD3 |
| **Data Points:** | 1.004168 | 6.665856 | 1.466672 | 3.135848287 | 0.820002749 |
| | 1.066575 | 6.690581 | 1.741185 | 5.909080655 | 1.648911508 |
| | 0.93369 | 7.202219 | 1.380369 | 3.22207759 | 1.263688519 |

Plot Name: **CEBPD**

| | | | | | |
|---|---|---|---|---|---|
| **Groups:** | Basal | Syn FCM | SynFCM | AbdomFCMD3 | AbdomFCMD3 |
| **Data Points:** | 1.116771 | 1.602208 | 0.901762 | 2.14680194 | 1.285236613 |
| | 0.880453 | 1.873068 | 1.037665 | 2.490209334 | 1.453818641 |
| | 1.01702 | 1.934168 | 0.893987 | 1.712761853 | 1.275421616 |

**Subfigure:** Figure 5A

| | | |
|---|---|---|
| Plot Name: | **NR3C1** | |
| **Groups:** | NTC | GCRKO |
| **Data Points:** | 0.893561 | 0.101824 |
| | 1.014984 | 0.124495 |
| | 1.102596 | 0.140905 |

**Subfigure:** Figure 5B

Plot Name: **APOD**

| **Groups:** | NTC | NTC+cortisol | GCRKO | GCRKO+cortisol |
|---|---|---|---|---|
| **Data Points:** | 1.083704 | 7.593016 | 0.700997 | 0.650027 |
| | 1.384909 | 9.168721 | 0.679473 | 0.520914 |
| | 0.666297 | 9.145486 | 0.674142 | 0.416555 |

Plot Name: **CEBPD**

| **Groups:** | NTC | NTC+cortisol | GCRKO | GCRKO+cortisol |
|---|---|---|---|---|
| **Data Points:** | 0.824145 | 9.61091 | 0.981106 | 0.86668 |
| | 1.045933 | 6.505773 | 1.062518 | 0.852114 |
| | 1.160091 | 6.311837 | 1.125935 | 0.753448 |

Plot Name: **NNMT**

| **Groups:** | NTC | NTC+cortisol | GCRKO | GCRKO+cortisol |
|---|---|---|---|---|
| **Data Points:** | 0.544991 | 2.185565 | 1.231839 | 1.146236 |
| | 1.519002 | 1.635865 | 1.302082 | 1.330945 |
| | 1.207959 | 1.683418 | 1.160267 | 0.888789 |

| | | | |
|---|---|---|---|
| **Subfigure:** | Figure 6A | *Mifepristone targets* | |
| Plot Name: | **APOD** | | |
| **Groups:** | Basal | Progesterone3 | Aldosterone1uM |
| **Data Points:** | 0.985411 | 1.926601 | 4.823711 |
| | 0.84972 | 1.712594 | 3.932405 |
| | 1.194281 | 1.422885 | 4.876468 |
| Plot Name: | **CEBPD** | | |
| **Groups:** | Basal | Progesterone3 | Aldosterone1uM |
| **Data Points:** | 0.821954 | 0.392126 | 4.433595 |
| | 1.034113 | 1.235005 | 2.925625 |
| | 1.17648 | 1.186862 | 3.627989 |
| Plot Name: | **NNMT** | | |
| **Groups:** | Basal | Progesterone3 | Aldosterone1uM |
| **Data Points:** | 1.054918 | 1.692776 | 3.007317 |
| | 1.22128 | 1.28745 | 2.503151 |
| | 0.776187 | 1.232983 | 1.930761 |

**Subfigure:** Figure 6B

| Plot Name: | **Aldosterone** | | | |
|---|---|---|---|---|
| **Groups:** | RPMI | AbdomD1 | AbdomD2 | AbdomD3 |
| **Data Points:** | 0 | 1.93289 | 7.319363 | 0 |

| Plot Name: | **APOD** | | |
|---|---|---|---|
| **Groups:** | Basal | Aldosterone50pg/mL | Aldosterone10pg/mL |
| **Data Points:** | 1.163094 | 1.179823 | 1.646188 |
| | 0.799152 | 1.107814 | 1.102291 |
| | 1.075859 | 0.928278 | 1.254332 |

| Plot Name: | **CEBPD** | | |
|---|---|---|---|
| **Groups:** | Basal | Aldosterone50pg/mL | Aldosterone10pg/mL |
| **Data Points:** | 0.82911 | 0.653037 | 1.694425 |
| | 1.002227 | 0.62174 | 1.036825 |
| | 1.203433 | 0.809047 | 0.90663 |

| Plot Name: | **NNMT** | | |
|---|---|---|---|
| **Groups:** | Basal | Aldosterone50pg/mL | Aldosterone10pg/mL |
| **Data Points:** | 1.141793 | 1.23705 | 1.476788 |
| | 0.979336 | 0.812844 | 1.052513 |
| | 0.894295 | | 0.823735 |

| Goutsyn | SynBatch 1 | ACM(pre-ad media) |
|---|---|---|
| 0.511262 | 20.3141 | 11.88481 |

**Subfigure:** Figure 6C *Metyrapone*

| | | | | |
|---|---|---|---|---|
| Plot Name: | **APOD** | | | |
| **Groups:** | Basal | Metyr-apone | FCM | FCM |
| **Data Points:** | 0.985411 | 1.145596 | 5.45717 | 8.886424 |
| | 0.84972 | 1.284209 | 5.951404 | 5.759797 |
| | 1.194281 | | 5.501386 | 6.195771 |

| | | | | |
|---|---|---|---|---|
| Plot Name: | **CEBPD** | | | |
| **Groups:** | Basal | Metyr-apone | FCM | FCM |
| **Data Points:** | 0.821954 | 0.897778 | 4.018024 | 5.559428 |
| | 1.034113 | 0.474412 | 3.90836 | 4.196979 |
| | 1.17648 | | 4.251961 | 3.469234 |

| | | | | |
|---|---|---|---|---|
| Plot Name: | **NNMT** | | | |
| **Groups:** | Basal | Metyr-apone | FCM | FCM |
| **Data Points:** | 1.054918 | 1.086307 | 3.583791 | 4.198681 |
| | 1.22128 | 1.801518 | 3.559226 | 3.553773 |
| | 0.776187 | | 0.703747 | 1.314594 |

**Subfigure:** Figure 7B

Plot Name: **APOD**

| **Groups:** | 4hr Cortisol | 22hr cortisol | 4hr FCM | 22hr FCM | 4hr FCM+GCI | 22hr FCM+GCRant |
|---|---|---|---|---|---|---|
| **Data Points:** | -0.338008 | 1.628923 | 0.64524 | 3.44615 | 0.608063 | 1.398747 |

Plot Name: **CEBPD**

| **Groups:** | 4hr Cortisol | 22hr cortisol | 4hr FCM | 22hr FCM | 4hr FCM+GCI | 22hr FCM+GCRant |
|---|---|---|---|---|---|---|
| **Data Points:** | 0.2782581 | 1.307501 | 2.597603 | 0.964418 | 1.022253 | -0.79103 |

Plot Name: **NNMT**

| **Groups:** | 4hr Cortisol | 22hr cortisol | 4hr FCM | 22hr FCM | 4hr FCM+GCI | 22hr FCM+GCRant |
|---|---|---|---|---|---|---|
| **Data Points:** | 0.6547576 | 0.977202 | 1.173594 | 1.548054 | 0.659341 | 0.02945 |

**Subfigure:** Figure 7C

Plot Name: **No steroid user's TGFB score**

| **Groups:** | Healthy | OA | RA | REM |
| --- | --- | --- | --- | --- |
| **Data Points:** | 0.061704 | 0.070671 | 0.071414 | 0.062012 |
| | 0.075939 | 0.077368 | 0.065952 | 0.079372 |
| | 0.045622 | 0.094134 | 0.086819 | 0.048696 |
| | 0.055366 | 0.096847 | 0.103122 | |
| | 0.050048 | 0.108413 | 0.083881 | |
| | 0.050727 | 0.074471 | 0.126595 | |
| | 0.060221 | 0.101025 | 0.065552 | |
| | 0.058361 | 0.129728 | 0.064485 | |
| | 0.070047 | 0.086462 | 0.074755 | |
| | 0.062425 | | 0.061014 | |
| | | | 0.060343 | |
| | | | 0.120692 | |
| | | | 0.094857 | |
| | | | 0.040933 | |
| | | | 0.095384 | |
| | | | 0.108384 | |
| | | | 0.079114 | |
| | | | 0.096831 | |
| | | | 0.080228 | |
| | | | 0.081247 | |
| | | | 0.068241 | |
| | | | 0.082827 | |
| | | | 0.084328 | |
| | | | 0.090739 | |
| | | | 0.051209 | |

**Subfigure:** **Figure 8C**

Plot Name: **AMP QC cell proportions merged with healthy**

| | **Healthy** | | | | |
|---|---|---|---|---|---|
| **Sublining Fibr** | 0.861314 | 0.434058 | 0.527231 | 0.460116 | 0.860424 |
| **Lining Fibrobl** | 0.10219 | 0.012319 | 0.224797 | 0.4 | 0.060071 |
| **Arterial EC** | 0.00365 | 0.280435 | 0.105446 | 0.045087 | 0.016784 |
| **Venous EC** | 0 | 0.176812 | 0.073001 | 0.039306 | 0.015018 |
| **Lymphatic EC** | 0.00365 | 0.012319 | 0.002317 | 0.028902 | 0.007067 |
| **Mural cell** | 0.007299 | 0.018116 | 0.017381 | 0.002312 | 0.00265 |
| **MERTK+Macr** | 0 | 0.01087 | 0.00927 | 0.015029 | 0.016784 |
| **IL1B+ Macrop** | 0.00365 | 0.008696 | 0.017381 | 0.002312 | 0.003534 |
| **CD4 T cell** | 0.014599 | 0.021739 | 0.011587 | 0.004624 | 0.013251 |
| **CD8 T cell** | 0 | 0.005797 | 0.008111 | 0.001156 | 0.000883 |
| **NK cells** | 0 | 0.00942 | 0.003476 | 0 | 0.001767 |
| **B cell** | 0.00365 | 0.005072 | 0 | 0.001156 | 0 |
| **Plasma cells** | 0 | 0.002174 | 0 | 0 | 0.001767 |
| **pDC** | | | | 0 | 0 |
| **Proliferating** | 0 | 0.002174 | 0 | 0 | 0 |

| | | | | | | |
|---|---|---|---|---|---|---|
| 0.37931 | 0.730823 | 0.735849 | 0.942643 | 0.285814 | 0.933333 | 0.21145091 |
| 0.086207 | 0.131102 | 0.01761 | 0.002494 | 0.65299 | 0.019394 | 0.71177615 |
| 0.12069 | 0.026499 | 0.057862 | 0.009975 | 0.005563 | 0.009697 | 0.01040989 |
| 0.201149 | 0.079498 | 0.101887 | 0.031796 | 0.018081 | 0.021818 | 0.02667534 |
| 0.034483 | 0.027894 | 0 | 0.001247 | 0.006954 | 0.006061 | 0.00455433 |
| 0 | 0.001395 | 0.038994 | 0.006234 | 0.019471 | 0.007273 | 0.00065062 |
| 0.028736 | 0 | 0.01761 | 0.003117 | 0.002782 | 0 | 0.01171113 |
| 0.04023 | 0 | 0.008805 | 0.000623 | 0 | 0 | 0.00390371 |
| 0.011494 | 0.002789 | 0.016352 | 0.000623 | 0.004172 | 0.002424 | 0.00845804 |
| 0.022989 | 0 | 0 | 0 | 0 | 0 | 0.00325309 |
| 0.045977 | 0 | 0.001258 | 0 | 0.000695 | 0 | 0.00455433 |
| 0.022989 | 0 | 0.001258 | 0.000623 | 0.002086 | 0 | 0.00130124 |
| 0 | 0 | 0.001258 | 0.000623 | 0.001391 | 0 | 0.00065062 |
| 0 | 0 | 0.001258 | 0 | 0 | 0 | |
| 0.005747 | 0 | 0 | 0 | 0 | 0 | 0.00065062 |

| | | | | OA | |
|---|---|---|---|---|---|
| 0.921452 | 0.919204 | 0.449225 | 0.278957 | 0.199864 | 0.045988 |
| 0.015182 | 0.005269 | 0.175559 | 0.618479 | 0.36342 | 0.144814 |
| 0.009241 | 0.006148 | 0.166954 | 0.033599 | 0.040719 | 0.179061 |
| 0.030363 | 0.017857 | 0.11704 | 0.028294 | 0.098066 | 0.458904 |
| 0.00066 | 0.008489 | 0.056799 | 0.014589 | 0.027146 | 0.009785 |
| 0 | 0.021663 | 0 | 0.001326 | | |
| 0.00066 | 0.013759 | 0.017212 | 0.004863 | 0.029861 | 0.08317 |
| 0 | 0.002635 | 0.006885 | 0.000884 | 0.072616 | 0.052838 |
| 0.019142 | 0.002342 | 0.001721 | 0.005305 | 0.066508 | 0.005871 |
| 0 | 0 | 0 | 0.008842 | 0.023074 | 0.002935 |
| 0 | 0.000585 | 0 | 0.00221 | 0.008144 | 0.016634 |
| 0.00132 | 0.002049 | 0.005164 | 0.000884 | 0.058025 | 0 |
| 0.00198 | 0 | 0.003442 | 0.001326 | 0.011198 | 0 |
| | | | | 0.001018 | 0 |
| 0 | 0 | 0 | 0.000442 | 0.000339 | 0 |

| | | | | | | |
|---|---|---|---|---|---|---|
| 0.077662 | 0.033208 | 0.115959 | 0.264941 | 0.057322 | 0.23499 | 0.214153 |
| 0.028022 | 0.866095 | 0.058553 | 0.119224 | 0.316736 | 0.007469 | 0.122905 |
| 0.066453 | 0.00857 | 0.029851 | 0.016636 | 0.052929 | 0.015513 | 0.029795 |
| 0.118495 | 0.024103 | 0.191734 | 0.127542 | 0.053138 | 0.061764 | 0.108007 |
| 0.008807 | 0.016604 | 0.008037 | 0.009858 | 0.021548 | 0.001436 | 0.024209 |

| | | | | | | |
|---|---|---|---|---|---|---|
| 0.195356 | 0.029459 | 0.570608 | 0.090265 | 0.149163 | 0.159437 | 0.130354 |
| 0.29944 | 0.020354 | 0.019518 | 0.04313 | 0.123431 | 0.018098 | 0.327747 |
| 0.108086 | 0.000536 | 0.001148 | 0.096118 | 0.098326 | 0.191037 | 0.007449 |
| 0.038431 | 0 | 0 | 0.063463 | 0.029707 | 0.063487 | 0 |
| 0.020817 | 0 | 0.004592 | 0.047135 | 0.053347 | 0.014076 | 0.007449 |
| 0.036029 | 0.001071 | 0 | 0.104128 | 0.03431 | 0.205401 | 0.022346 |
| 0.001601 | 0 | 0 | 0.007394 | 0.000209 | 0.017811 | 0 |
| 0 | 0 | 0 | 0.009242 | 0.007531 | 0.001436 | 0.005587 |
| 0.000801 | 0 | 0 | 0.000924 | 0.002301 | 0.008044 | 0 |

**RA**

| | | | | | |
|---|---|---|---|---|---|
| 0.451069 | 0.145401 | 0.118541 | 0.020408 | 0.045262 | 0.27598 |
| 0.138691 | 0.147774 | 0 | 0.030612 | 0.004243 | 0.204429 |
| 0.022683 | 0.017211 | 0.018237 | 0.010204 | 0.02546 | 0.015332 |
| 0.062865 | 0.017211 | 0.030395 | 0.081633 | 0.032532 | 0.020443 |
| 0.005185 | 0 | 0 | 0 | 0 | 0 |
| 0.000648 | 0.00178 | 0.00304 | 0 | 0.002829 | 0.001704 |
| 0.139987 | 0.358457 | 0.182371 | 0.081633 | 0.671853 | 0.080068 |
| 0.018795 | 0.107418 | 0.231003 | 0.163265 | 0.057992 | 0.039182 |
| 0.033701 | 0.074184 | 0.179331 | 0.132653 | 0.091938 | 0.151618 |
| 0.01361 | 0.032641 | 0.039514 | 0.081633 | 0.02546 | 0.085179 |
| 0.01361 | 0.018991 | 0.139818 | 0.204082 | 0.029703 | 0.032368 |
| 0.01685 | 0.033234 | 0.048632 | 0.061224 | 0.009901 | 0.030664 |
| 0.081011 | 0.04095 | 0.00304 | 0.081633 | 0.001414 | 0.057922 |
| 0.000648 | 0 | 0 | 0.010204 | 0 | 0 |
| 0.000648 | 0.004748 | 0.006079 | 0.040816 | 0.001414 | 0.005111 |

| | | | | | | |
|---|---|---|---|---|---|---|
| 0.056995 | 0.126761 | 0.053073 | 0.15384613 | 0.18000004 | 0.116369 | 0 |
| 0.060449 | 0 | 0.14176 | 0 | 0.08000002 | 0.114042 | 0 |
| 0.006908 | 0.031299 | 0.011872 | 0 | 0.02 | 0.017843 | 0.016667 |
| 0.007772 | 0.037559 | 0.027933 | 0 | 0.08000002 | 0.03879 | 0 |
| 0 | 0 | 0 | 0.02564102 | 0 | 0 | 0 |
| 0.000864 | 0.00313 | 0.011872 | 0 | 0 | 0.004655 | 0 |
| 0.013817 | 0.225352 | 0.106145 | 0 | 0.02 | 0.024825 | 0.05 |
| 0.08981 | 0.104851 | 0.180866 | 0.10256412 | 0.02 | 0.046548 | 0.016667 |
| 0.348877 | 0.198748 | 0.227654 | 0.48717948 | 0.15999998 | 0.188518 | 0.583333 |
| 0.1019 | 0.081377 | 0.097067 | 0 | 0.19999998 | 0.082234 | 0.016667 |
| 0.031088 | 0.046948 | 0.057961 | 0.10256412 | 0.19999998 | 0.15128 | 0.033333 |
| 0.208981 | 0.084507 | 0.039106 | 0.10256412 | 0.02 | 0.19007 | 0.15 |
| 0.033679 | 0.050078 | 0.009078 | 0 | 0.02 | 0.016292 | 0.133333 |
| 0.011226 | 0 | 0.028631 | 0 | 0 | 0.002327 | 0 |
| 0.027634 | 0.00939 | 0.006983 | 0.02564102 | 0 | 0.006206 | 0 |

| | | | | | | |
|---|---|---|---|---|---|---|
| 0 | 0.012075 | 0.028986 | 0.048077 | 0.058458 | 0.55799 | 0.190476 |
| 0 | 0.004391 | 0 | 0.001374 | 0 | 0.006443 | 0.063492 |
| 0.101852 | 0.008782 | 0.014493 | 0.006181 | 0.048507 | 0.052835 | 0.190476 |
| 0.055556 | 0.002195 | 0.024155 | 0.03022 | 0.050995 | 0.291237 | 0.111111 |
| 0 | 0 | 0 | 0 | 0 | 0 | 0 |
| 0 | 0.005488 | 0.024155 | 0 | 0.001244 | 0.006443 | 0 |
| 0.055556 | 0.072448 | 0.289855 | 0.400412 | 0.022388 | 0.061856 | 0.031746 |
| 0 | 0.069155 | 0.028986 | 0.068681 | 0.115672 | 0.014175 | 0.142857 |
| 0.231482 | 0.307355 | 0.289855 | 0.179945 | 0.243781 | 0.002577 | 0.063492 |
| 0.277778 | 0.255763 | 0.15942 | 0.083791 | 0.174129 | 0 | 0.079365 |
| 0.166667 | 0.04281 | 0.043478 | 0.018544 | 0.050995 | 0.006443 | 0.031746 |
| 0.101852 | 0.145993 | 0.028986 | 0.055632 | 0.080846 | 0 | 0.063492 |
| 0.009259 | 0.041712 | 0.028986 | 0.087225 | 0.034826 | 0 | 0.031746 |
| 0 | 0.002195 | 0 | 0.006868 | 0.097015 | 0 | 0 |
| 0 | 0.029638 | 0.038647 | 0.013049 | 0.021144 | 0 | 0 |

| | | | | | | |
|---|---|---|---|---|---|---|
| 0.018405 | 0.215813 | 0.143541 | 0.056575 | 0.180826 | 0.012295 | 0.095238 |
| 0.018405 | 0.007129 | 0.009569 | 0.461009 | 0.046765 | 0 | 0 |
| 0.030675 | 0.00324 | 0.043062 | 0.002294 | 0.021824 | 0.015369 | 0.042328 |
| 0.04908 | 0.020091 | 0.028708 | 0.014526 | 0.048324 | 0.033811 | 0.121693 |
| 0 | 0.002592 | 0 | 0 | 0 | 0 | 0.007937 |
| 0.006135 | 0.002592 | 0.014354 | 0.003058 | 0.004677 | 0.003074 | 0.026455 |
| 0.361963 | 0.006481 | 0.301435 | 0.229358 | 0.096648 | 0.417008 | 0.140212 |
| 0.177914 | 0.021387 | 0.172249 | 0.036697 | 0.007015 | 0.060451 | 0.108466 |
| 0.184049 | 0.2372 | 0.143541 | 0.112385 | 0.154326 | 0.155738 | 0.256614 |
| 0.092025 | 0.34932 | 0.076555 | 0.026758 | 0.073266 | 0.167008 | 0.074074 |
| 0.02454 | 0.0849 | 0.019139 | 0.0237 | 0.028839 | 0.067623 | 0.097884 |
| 0.02454 | 0.034349 | 0.009569 | 0.019113 | 0.118472 | 0.038934 | 0.029101 |
| 0.006135 | 0.012314 | 0.004785 | 0.012232 | 0.183164 | 0.010246 | 0 |
| 0 | 0 | 0 | 0.000765 | 0.003897 | 0.003074 | 0 |
| 0.006135 | 0.002592 | 0.033493 | 0.001529 | 0.031956 | 0.015369 | 0 |

0.306533
0.080402
0.0201
0.060302
0
0.005025
0.256281
0.0201
0.090452
0.025126
0.005025
0.085427
0.045226
0
0

**Subfigure:** **Figure 8G**

Plot Name: **Healthy fibroblast defined proportions fan QC**

| | **Healthy** | | | | |
|---|---|---|---|---|---|
| **PRG4** | 0.07874 | 0.022913 | 0.284722 | 0.433432 | 0.547497 |
| **PLIN2** | 0.228346 | 0.266776 | 0.001736 | 0.026627 | 0.066394 |
| **DKK3** | 0.098425 | 0.016367 | 0.388889 | 0.155325 | 0.022472 |
| **CXCL12** | 0.251969 | 0.423895 | 0.213542 | 0.232249 | 0.103166 |
| **CD34** | 0.11811 | 0.00982 | 0.043403 | 0.063609 | 0.012257 |
| **APOD** | 0.204724 | 0.250409 | 0.060764 | 0.065089 | 0.023493 |
| **MT** | 0.007874 | 0.00982 | 0 | 0.007396 | 0.003064 |
| **VCAM1** | 0.011811 | 0 | 0.006944 | 0.016272 | 0.221655 |

| | | | | | | |
|---|---|---|---|---|---|---|
| 0.342105 | 0.138047 | 0.041971 | 0.041637 | 0.627389 | 0.030423 | 0.704374 |
| 0 | 0.23569 | 0.259124 | 0.366267 | 0.067675 | 0.253968 | 0.095023 |
| 0.210526 | 0.087542 | 0.458029 | 0.146789 | 0.054936 | 0.345238 | 0.038462 |
| 0.236842 | 0.385522 | 0.021898 | 0.110797 | 0.152866 | 0.064815 | 0.128959 |
| 0.026316 | 0.070707 | 0.167883 | 0.154552 | 0.009554 | 0.234127 | 0.004525 |
| 0.131579 | 0.074074 | 0.036496 | 0.129852 | 0.021497 | 0.059524 | 0.006033 |
| 0 | 0.006734 | 0.014599 | 0.038814 | 0.001592 | 0.002646 | 0.000754 |
| 0.052632 | 0.001684 | 0 | 0.011291 | 0.06449 | 0.009259 | 0.02187 |

| | | | | | |
|---|---|---|---|---|---|
| | | | | **RA** | |
| 0.016708 | 0.01 | 0.225714 | 0.579203 | 0 | 0.342714 |
| 0.182122 | 0.385667 | 0.402857 | 0.050108 | 0.0625 | 0.017085 |
| 0.468672 | 0.215333 | 0.08 | 0.015625 | 0.357143 | 0.01206 |
| 0.012531 | 0.089333 | 0.054286 | 0.241379 | 0.464286 | 0.611055 |
| 0.299916 | 0.213333 | 0.08 | 0.01347 | 0.008929 | 0 |
| 0.010025 | 0.068667 | 0.134286 | 0.07597 | 0.080357 | 0.01005 |
| 0.010025 | 0.016333 | 0.014286 | 0.001078 | 0.008929 | 0.00201 |
| 0 | 0.001333 | 0.008571 | 0.023168 | 0.017857 | 0.005025 |

| | | | | | | |
|---|---|---|---|---|---|---|
| 0.461874 | 0.003546 | 0.591476 | 0.142857 | 0.111111 | 0.083333 | 0.142569 |
| 0.008715 | 0.021277 | 0 | 0 | 0 | 0.003205 | 0.023201 |
| 0.006536 | 0.212766 | 0.02079 | 0.178571 | 0.055556 | 0.051282 | 0.193006 |
| 0.48366 | 0.677305 | 0.323285 | 0.392857 | 0.305556 | 0.817308 | 0.612979 |
| 0 | 0.010638 | 0 | 0 | 0 | 0.003205 | 0.012777 |
| 0.021786 | 0.067376 | 0.050936 | 0.285714 | 0.5 | 0.016026 | 0.007061 |
| 0.004357 | 0.003546 | 0.004158 | 0 | 0 | 0.003205 | 0.001681 |
| 0.013072 | 0.003546 | 0.009356 | 0 | 0.027778 | 0.022436 | 0.006725 |

| | | | | | | |
|---|---|---|---|---|---|---|
| 0.239867 | 0.082278 | 0 | 0.057143 | 0.269231 | 0.014395 | 0.008696 |
| 0.022765 | 0.025316 | 0 | 0.028571 | 0 | 0.03387 | 0.026087 |
| 0.084398 | 0.21519 | 0.111111 | 0.114286 | 0.076923 | 0.300593 | 0.365217 |
| 0.625763 | 0.556962 | 0.777778 | 0.028571 | 0.461538 | 0.605419 | 0.556522 |
| 0.000555 | 0.012658 | 0 | 0.685714 | 0 | 0.020322 | 0 |
| 0.012771 | 0.088608 | 0.111111 | 0.085714 | 0.153846 | 0.024555 | 0.043478 |
| 0.001666 | 0 | 0 | 0 | 0 | 0.000847 | 0 |
| 0.012215 | 0.018987 | 0 | 0 | 0.038462 | 0 | 0 |

| | | | | | | |
|---|---|---|---|---|---|---|
| 0.852116 | 0.120975 | 0 | 0.015152 | 0.247967 | 0.363554 | 0 |
| 0.000951 | 0.006963 | 0.088889 | 0 | 0 | 0.011786 | 0.026667 |
| 0.005706 | 0.014795 | 0.288889 | 0.409091 | 0.308943 | 0.059837 | 0.426667 |
| 0.115549 | 0.826806 | 0.222222 | 0.090909 | 0.390244 | 0.504986 | 0.16 |
| 0 | 0 | 0.177778 | 0.318182 | 0.020325 | 0 | 0.32 |
| 0.004755 | 0.020017 | 0.155556 | 0.121212 | 0.020325 | 0.036265 | 0.066667 |
| 0.000476 | 0.002611 | 0.022222 | 0.045455 | 0.004065 | 0.016319 | 0 |
| 0.020447 | 0.007833 | 0.044444 | 0 | 0.00813 | 0.007253 | 0 |

| | | | OA | | |
|---|---|---|---|---|---|
| 0.008655 | 0.163265 | 0.114286 | 0.547463 | 0.67013 | 0.222222 |
| 0.039281 | 0 | 0.028571 | 0.00209 | 0.002597 | 0.006734 |
| 0.72237 | 0.122449 | 0.2 | 0.014925 | 0.054545 | 0.03367 |
| 0.096538 | 0.591837 | 0.571429 | 0.363284 | 0.18961 | 0.609428 |
| 0.081225 | 0 | 0 | 0 | 0.002597 | 0 |
| 0.046605 | 0.122449 | 0.085714 | 0.051642 | 0.05974 | 0.107744 |
| 0.005326 | 0 | 0 | 0.007761 | 0.018182 | 0.013468 |
| 0 | 0 | 0 | 0.012836 | 0.002597 | 0.006734 |

| | | | | | |
|---|---|---|---|---|---|
| 0.776 | 0.328904 | 0.273856 | 0.729342 | 0.089091 | 0.327078 |
| 0.008296 | 0.053156 | 0.007758 | 0.001897 | 0.028485 | 0.002681 |
| 0.004741 | 0.299003 | 0.147401 | 0.016798 | 0.052121 | 0.010724 |
| 0.184296 | 0.27907 | 0.462374 | 0.198591 | 0.807273 | 0.568365 |
| 0 | 0 | 0.04616 | 0 | 0 | 0 |
| 0.012148 | 0.0299 | 0.051202 | 0.036034 | 0.01697 | 0.075067 |
| 0.003852 | 0.006645 | 0.007758 | 0.013276 | 0.000606 | 0.005362 |
| 0.010667 | 0.003322 | 0.003491 | 0.004064 | 0.005455 | 0.010724 |

**REM**

| | | |
|---|---|---|
| 0.244344 | 0.207143 | 0.538987 |
| 0.150452 | 0.015179 | 0.002156 |
| 0.133484 | 0.048661 | 0.298599 |
| 0.21776 | 0.641518 | 0.098455 |
| 0.119344 | 0.035714 | 0.028027 |
| 0.076357 | 0.037054 | 0.033417 |
| 0.045814 | 0.003571 | 0.000359 |
| 0.012443 | 0.011161 | 0 |

**Subfigure:** Figure 8I Bulk Module Scores: GC users compared

Plot Name: **Cortisol score: Steroid users vs non-users**

| **Groups:** | Healthy Non-reported | Healthy Steroids |
|---|---|---|
| **Data Points:** | 0.098041 | 0.181085 |
| | 0.157596 | 0.194575 |
| | 0.224444 | 0.14452 |
| | 0.165339 | 0.156431 |
| | 0.164132 | 0.19136 |
| | 0.119049 | 0.158774 |
| | 0.220185 | |
| | 0.185089 | |
| | 0.175247 | |
| | 0.119264 | |

Plot Name: **FCM score: Steroid users vs non-users**

| **Groups:** | Healthy Non-reported | Healthy Steroids |
|---|---|---|
| **Data Points:** | 0.042612 | 0.088419 |
| | 0.079728 | 0.090581 |
| | 0.107215 | 0.063301 |
| | 0.130702 | 0.072227 |
| | 0.060483 | 0.090319 |
| | 0.097929 | 0.077223 |
| | 0.113681 | |
| | 0.088338 | |
| | 0.102388 | |
| | 0.05432 | |

Plot Name: FCM+GCRant score: Steroid users vs non-users

| **Groups:** | HealthyNon-reported | HealthySteroids |
|---|---|---|
| **Data Points:** | | 0.065281 |
| | | 0.052427 |
| | | 0.035768 |

| | |
|---|---|
| | 0.026965 |
| | 0.057242 |
| 0.03561 | |
| 0.053041 | |
| | 0.048659 |
| 0.05589 | |
| 0.097697 | |
| 0.027374 | |
| 0.085124 | |
| 0.083739 | |
| 0.050756 | |
| 0.096811 | |
| 0.037771 | |

**Subfigure:** Figure 9A

Plot Name: **Cortisol Activation Score**

| | | |
|---|---|---|
| **Groups:** | Ctl | Case |
| **Data Points:** | 0.017929 | 0.008785 |
| | 0.040384 | 0.040634 |
| | 0.00451 | -0.00902 |
| | 0.060579 | -0.04652 |
| | 0.046134 | 0.062749 |
| | -0.00653 | -0.06991 |
| | 0.015985 | 0.009098 |
| | 0.02774 | 0.037477 |
| | 0.048457 | 0.000284 |
| | 0.021236 | -0.02374 |
| | 0.021932 | 0.052071 |
| | 0.051994 | 0.0297 |
| | 0.037371 | 0.009376 |
| | 0.024053 | 0.049112 |
| | 0.029509 | 0.035566 |
| | 0.018414 | 0.059547 |
| | 0.035575 | 0.044234 |
| | 0.038064 | -0.0069 |
| | 0.050858 | 0.030525 |
| | 0.033904 | 0.02286 |
| | 0.018765 | -0.03712 |
| | 0.042637 | 0.013893 |
| | 0.021633 | 0.033718 |
| | 0.014151 | -0.01201 |
| | 0.050535 | 0.0463 |
| | 0.023416 | -0.05359 |
| | 0.04436 | -0.014 |
| | 0.091648 | -0.03792 |
| | 0.036041 | -0.02404 |
| | 0.005176 | -0.03063 |
| | 0.042144 | 0.066233 |
| | 0.057255 | 0.041722 |
| | 0.030723 | -0.05913 |
| | 0.054255 | -0.01842 |
| | 0.044703 | 0.020066 |
| | 0.047142 | 0.063475 |
| | 0.053933 | 0.049028 |
| | 0.040186 | 0.025153 |
| | -0.0146 | 0.026025 |
| | 0.009181 | 0.040048 |
| | 0.052238 | 0.016495 |
| | 0.093531 | -0.00923 |
| | 0.039191 | 0.003822 |

| | |
|---|---|
| -0.01318 | 0.021263 |
| -0.02118 | -0.02665 |
| 0.056135 | 0.052999 |
| 0.066172 | -0.00091 |
| 0.011813 | 0.065679 |
| 0.025304 | -0.0115 |
| 0.057751 | 0.027329 |
| 0.041263 | 0.012515 |
| 0.028841 | 0.031192 |
| 0.025112 | 0.037895 |
| 0.091108 | 0.013521 |
| 0.033356 | 0.019059 |
| -0.01466 | -0.02481 |
| 0.026031 | 0.024465 |
| 0.021755 | -0.02071 |
| 0.038725 | 0.050205 |
| 0.098204 | -0.00961 |
| 0.024437 | -0.05756 |
| 0.090225 | -0.00104 |
| 0.07228 | 0.025808 |
| 0.014111 | 0.013515 |
| 0.094748 | -0.01758 |
| 0.03842 | -0.06075 |
| 0.002699 | 0.023308 |
| 0.04504 | 0.021637 |
| 0.000669 | 0.024628 |
| -0.05389 | -0.0381 |
| 0.047512 | 0.014108 |
| 0.013329 | 0.017462 |
| 0.017413 | 0.022692 |
| 0.025772 | 0.010901 |
| 0.023434 | 0.026787 |
| -0.01796 | -0.03324 |
| 0.045973 | 0.017731 |
| 0.036056 | 0.011439 |
| 0.030869 | -0.02591 |
| 0.003812 | -0.01634 |
| 0.016501 | 0.015618 |
| 0.014444 | -0.03141 |
| 0.027036 | 0.005978 |
| 0.023861 | -0.00598 |
| 0.049136 | 0.015092 |
| 0.02767 | 0.020636 |
| 0.061263 | -0.01519 |
| 0.031116 | -0.0249 |
| 0.06182 | -0.00796 |
| 0.000906 | 0.010973 |

| | |
|---|---|
| 0.03016 | -0.00974 |
| -0.01057 | 0.079083 |
| 0.025201 | 0.016617 |
| 0.086798 | 0.078078 |
| 0.016966 | 0.075079 |
| 0.022933 | 0.012128 |
| 0.039739 | -0.03949 |
| 0.020808 | 0.106571 |
| 0.01973 | 0.01556 |
| 0.034947 | -0.02599 |
| -0.01042 | 0.0183 |
| 0.052733 | 0.046965 |
| 0.054128 | -0.00403 |
| 0.011321 | 0.008048 |
| 0.065255 | -0.03208 |
| 0.054535 | 0.043677 |
| 0.065667 | 0.01408 |
| 0.034562 | 0.044572 |
| 0.033765 | 0.075477 |
| 0.052568 | 0.011078 |
| 0.057154 | 0.055146 |
| 0.008387 | 0.036103 |
| 0.036677 | 0.023123 |
| 0.037919 | 0.038481 |
| 0.043657 | 0.023971 |
| -0.00512 | 0.007331 |
| 0.047293 | 0.040021 |
| 0.009691 | -0.03744 |
| 0.040732 | 0.071694 |
| 0.03506 | -0.0904 |
| 0.038761 | -0.01619 |
| 0.021974 | 0.053832 |
| 0.047372 | 0.025642 |
| 0.049012 | -0.01284 |
| 0.042421 | -0.07814 |
| 0.052284 | -0.02773 |
| 0.048808 | 0.031746 |
| 0.055581 | 0.041249 |
| 0.036192 | 0.03115 |
| 0.028399 | 0.044887 |
| 0.017389 | 0.001606 |
| 0.015398 | 0.001945 |
| 0.060872 | -0.00376 |
| 0.057593 | 0.029943 |
| 0.052121 | -0.03567 |
| 0.008507 | 0.050375 |
| 0.007272 | 0.066222 |

| | |
|---|---|
| 0.048858 | 0.009018 |
| 0.032906 | 0.040475 |
| -0.00821 | 0.013574 |
| 0.058304 | 0.072547 |
| 0.070886 | 0.004032 |
| 0.054139 | -0.00246 |
| 0.024031 | 0.021353 |
| 0.012326 | -0.02415 |
| 0.033798 | 0.004603 |
| 0.060737 | 0.068165 |
| 0.073794 | -0.00057 |
| -0.02063 | 0.027942 |
| 0.062262 | 0.014338 |
| 0.056407 | 0.05172 |
| 0.037123 | -0.01061 |
| 0.00868 | 0.07929 |
| 0.063805 | -0.00541 |
| 0.000691 | 0.048133 |
| 0.039944 | 0.10153 |
| 0.037043 | -0.0237 |
| 0.024535 | -0.00226 |
| 0.074613 | 0.003943 |
| 0.00167 | 0.038368 |
| 0.031114 | 0.058005 |
| 0.068081 | 0.034586 |
| 0.038202 | 0.037358 |
| 0.05535 | -0.03421 |
| 0.037632 | -0.00114 |
| 0.008774 | 0.014219 |
| 0.06529 | -0.02773 |
| 0.018222 | 0.02781 |
| 0.024289 | 0.057505 |
| 0.048286 | 0.024955 |
| 0.008186 | 0.031853 |
| 0.073202 | 0.06535 |
| 0.024447 | 0.01618 |
| 0.019285 | -0.00366 |
| 0.093876 | 0.01137 |
| 0.042815 | 0.028299 |
| -0.0149 | -0.04431 |
| 0.012888 | -0.00267 |
| -0.00682 | -0.02016 |
| 0.073483 | 0.04919 |
| 0.052416 | 0.027812 |
| 0.053476 | 0.026609 |
| 0.08646 | 0.008998 |
| 0.054488 | 0.007716 |

| | |
|---|---|
| 0.042405 | -0.00559 |
| 0.068295 | 0.062468 |
| 0.010419 | 0.054872 |
| 0.014071 | -0.01528 |
| 0.053054 | 0.05099 |
| 0.035733 | 0.009604 |
| 0.066679 | 0.013242 |
| 0.03983 | -0.00175 |
| 0.029449 | 0.07295 |
| 0.012881 | 0.075699 |
| 0.055279 | 0.087116 |
| 0.029761 | -0.02379 |
| 0.028671 | 0.001098 |
| 0.019916 | -0.04373 |
| 0.020919 | 0.036781 |
| 0.066692 | -0.0053 |
| 0.05783 | 0.023135 |
| 0.02247 | -0.04908 |
| 0.030383 | -0.01035 |
| 0.025173 | 0.042317 |
| 0.022226 | 0.029684 |
| 0.092394 | 0.029318 |
| 0.050278 | 0.026132 |
| 0.033975 | 0.014156 |
| 0.053603 | 0.020231 |
| 0.031695 | 0.033243 |
| 0.048553 | -0.04685 |
| 0.014112 | 0.016561 |
| 0.057818 | -0.06329 |
| -0.0051 | 0.052282 |
| 0.004418 | -0.03179 |
| 0.007124 | 0.035437 |
| -0.00128 | 0.027148 |
| 0.052104 | 0.049509 |
| 0.013366 | 0.019593 |
| 0.030757 | -0.00566 |
| 0.01449 | -0.01675 |
| -0.02197 | 0.025145 |
| 0.045319 | -0.02697 |
| -0.00819 | -0.02049 |
| 0.013759 | 0.01514 |
| 0.025594 | 0.013536 |
| 0.044288 | -0.07479 |
| 0.044368 | -0.01303 |
| 0.013593 | 0.006935 |
| -0.03635 | 0.009687 |
| 0.033262 | 0.000368 |

| | |
|---|---|
| 0.001825 | 0.003034 |
| 0.045385 | 0.016734 |
| 0.004974 | 0.083302 |
| 0.012455 | 0.050506 |
| 0.043995 | 0.011969 |
| 0.022738 | 0.008139 |
| 0.014918 | -0.00989 |
| 0.029392 | -0.05181 |
| 0.029394 | -0.02733 |
| 0.009477 | 0.041001 |
| 0.003988 | 0.014763 |
| 0.091122 | 0.053085 |
| 0.035649 | -0.05731 |
| 0.110527 | -0.02107 |
| 0.014212 | 0.032486 |
| 0.107321 | -0.0252 |
| 0.045425 | 0.075372 |
| 0.063259 | -0.01194 |
| 0.023938 | 0.032798 |
| 0.010042 | 0.007568 |
| 0.00611 | 0.008262 |
| 0.024024 | -0.05052 |
| 0.040838 | 0.062191 |
| 0.015879 | 0.075364 |
| 0.072483 | 0.035298 |
| 0.05003 | -0.00872 |
| 0.047145 | 0.039068 |
| 0.037135 | -0.01709 |
| 0.012858 | 0.086591 |
| 0.080779 | -0.01907 |
| 0.037045 | 0.032494 |
| 0.047509 | -0.0008 |
| 0.030902 | -0.0045 |
| 0.06469 | -0.00199 |
| 0.00392 | 0.067506 |
| 0.028418 | 0.049682 |
| 0.019567 | 0.003097 |
| 0.036226 | 0.063875 |
| 0.012742 | 0.013602 |
| 0.059906 | 0.020676 |
| 0.00396 | 0.007992 |
| 0.019705 | 0.029397 |
| 0.033474 | 0.04622 |
| -0.01206 | 0.044982 |
| 0.046525 | 0.010944 |
| 0.020511 | -0.00621 |
| 0.051847 | -0.02248 |

| | |
|---|---|
| 0.060297 | 0.02568 |
| 0.05948 | 0.000145 |
| 0.033602 | 0.019138 |
| -0.00924 | -0.0389 |
| 0.01179 | 0.025809 |
| 0.070521 | 0.025653 |
| 0.063994 | 0.03385 |
| 0.001346 | 0.035304 |
| -0.01606 | 0.009855 |
| 0.067957 | 0.050696 |
| 0.044063 | 0.017038 |
| 0.045452 | -0.01989 |
| 0.050212 | 0.012533 |
| 0.028318 | 0.05093 |
| 0.011327 | -0.02659 |
| 0.034799 | 0.024842 |
| 0.03687 | 0.016496 |
| -0.00722 | -0.01603 |
| 0.020267 | 0.033839 |
| 0.067834 | 0.038279 |
| 0.018274 | 0.038945 |
| 0.05959 | -0.0546 |
| 0.078465 | 0.023677 |
| 0.040536 | -0.00145 |
| 0.017554 | 0.041021 |
| -0.03068 | 0.00845 |
| 0.013747 | 0.026274 |
| 0.021176 | -0.01035 |
| 0.029752 | -0.03993 |
| 0.004134 | 0.007428 |
| 0.012607 | 0.059471 |
| 0.03978 | 0.02749 |
| 0.024746 | 0.036403 |
| -0.00564 | 0.081023 |
| 0.0291 | 0.022123 |
| -0.01402 | 0.003091 |
| 0.032833 | 0.063045 |
| 0.01965 | -0.00454 |
| 0.007758 | -0.0024 |
| 0.024482 | 0.018608 |
| 0.020469 | 0.070668 |
| 0.03008 | 0.029641 |
| 0.034558 | -0.03319 |
| 0.024303 | -0.06047 |
| 0.049189 | 0.025315 |
| 0.043471 | -0.0172 |
| 0.06613 | 0.000494 |

| | |
|---|---|
| 0.015953 | 0.035711 |
| 0.045595 | 0.01868 |
| 0.047157 | 0.032216 |
| 0.062846 | 0.014713 |
| 0.081636 | 0.072271 |
| 0.04218 | 0.014181 |
| 0.071708 | 0.060578 |
| 0.008191 | -0.05092 |
| 0.023088 | -0.05195 |
| 0.022602 | -0.01787 |
| 0.033056 | 0.04244 |
| 0.025567 | 0.047025 |
| 0.049961 | 0.060387 |
| 0.054753 | 0.050645 |
| 0.0885 | 0.008085 |
| 0.063005 | 0.04303 |
| 0.048561 | 0.030291 |
| 0.023023 | 0.014285 |
| 0.043959 | -0.0101 |
| 0.047582 | -0.00302 |
| -0.003 | 0.06801 |
| 0.042069 | 0.04415 |
| 0.060953 | -0.00293 |
| 0.001581 | -0.02951 |
| 0.069974 | -0.00756 |
| 0.052186 | -0.0129 |
| 0.022616 | -0.02922 |
| 0.009047 | -0.04312 |
| 0.055437 | 0.021866 |
| 0.017502 | 0.080226 |
| 0.019446 | 0.059787 |
| 0.042807 | -0.02351 |
| 0.005231 | 0.06138 |
| 0.065124 | 0.070661 |
| 0.021395 | 0.019156 |
| 0.049164 | 0.071145 |
| 0.02547 | -0.00708 |
| 0.044041 | 0.007119 |
| 0.068775 | 0.04251 |
| 0.084107 | 0.033356 |
| -0.01045 | 0.039151 |
| 0.000652 | -0.02657 |
| 0.037167 | -0.05025 |
| 0.05949 | 0.03369 |
| 0.007267 | 0.010966 |
| 0.021501 | 0.03092 |
| 0.016837 | -0.01404 |

| | |
|---|---|
| 0.040743 | 0.053426 |
| -0.00171 | 0.003318 |
| 0.015407 | -0.03183 |
| 0.018978 | 0.009788 |
| 0.023572 | 0.002115 |
| 0.058528 | 0.107259 |
| 0.063573 | -0.00197 |
| 0.073752 | 0.042276 |
| 0.071076 | 0.020982 |
| 0.039415 | -0.01882 |
| -0.00826 | 0.008291 |
| 0.028532 | 0.041823 |
| 0.08606 | 0.045716 |
| 0.011476 | -0.05621 |
| 0.067112 | 0.01766 |
| 0.019912 | -0.01377 |
| 0.022338 | 0.025213 |
| 0.026993 | 0.042277 |
| 0.069866 | 0.040521 |
| 0.04977 | 0.090608 |
| 0.052324 | 0.05097 |
| -0.01956 | 0.029547 |
| 0.03729 | -0.01501 |
| 0.044124 | -0.00916 |
| 0.028876 | 0.052133 |
| -0.01911 | -0.00319 |
| 0.046707 | 0.043674 |
| 0.017584 | -0.01166 |
| 0.052512 | -0.03192 |
| 0.038303 | 0.002555 |
| 0.010834 | 0.065789 |
| 0.015144 | 0.037082 |
| 0.002911 | 0.000199 |
| 0.040027 | 0.006789 |
| 0.037313 | -0.02111 |
| 0.032409 | 0.015853 |
| -0.00596 | 0.000192 |
| 0.033359 | 0.0312 |
| -0.00327 | 0.02264 |
| 0.015128 | 0.051826 |
| 0.01248 | 0.038705 |
| 0.012528 | 0.029527 |
| 0.00624 | -0.05366 |
| 0.06102 | -0.04989 |
| 0.049719 | 0.042678 |
| 0.059711 | -0.03073 |
| 0.059648 | 0.039617 |

| | |
|---|---|
| -0.0321 | 0.034448 |
| 0.064175 | 0.050247 |
| -0.00031 | 0.041901 |
| 0.031873 | 0.072778 |
| 0.080243 | -0.00864 |
| 0.015858 | -0.04828 |
| 0.042777 | 0.087079 |
| 0.050169 | 0.039641 |
| 0.062523 | 0.01054 |
| 0.061638 | -0.0106 |
| 0.024739 | 0.030116 |
| 0.075319 | -0.01413 |
| 0.050578 | 0.020922 |
| 0.064126 | 0.082436 |
| 0.043042 | 0.025336 |
| 0.019473 | 0.042799 |
| 0.048854 | 0.024531 |
| 0.034799 | 0.002163 |
| 0.034799 | 0.004422 |
| -0.00862 | 0.052901 |
| 0.013589 | -0.00204 |
| 0.092023 | 0.020972 |
| 0.028827 | 0.005916 |
| 0.019731 | 0.038406 |
| 0.046046 | 0.059167 |
| 0.054723 | -0.00455 |
| 0.045925 | 0.082896 |
| 0.049775 | 0.030259 |
| 0.047617 | 0.062837 |
| 0.02702 | -0.0137 |
| 0.049218 | 0.015972 |
| 0.047228 | -0.00139 |
| 0.030978 | 0.063297 |
| 0.039163 | 0.067502 |
| 0.072948 | 0.054937 |
| 0.056345 | -0.0079 |
| 0.035791 | 0.006939 |
| 0.020012 | 0.00491 |
| 0.025948 | 0.058255 |
| 0.045826 | 0.014302 |
| 0.034529 | -0.00357 |
| -0.01482 | -0.03516 |
| 0.042211 | -0.02273 |
| 0.056346 | 0.008037 |
| 0.022976 | -0.06914 |
| 0.035194 | -0.05395 |
| 0.019869 | 0.01883 |

| | |
|---|---|
| 0.025589 | 0.058652 |
| -0.00281 | 0.063738 |
| 0.021924 | 0.04031 |
| 0.0417 | 0.037108 |
| 0.003538 | -0.01595 |
| 0.030374 | 0.013707 |
| 0.060944 | 0.104765 |
| 0.020011 | -0.00595 |
| 0.026315 | 0.087798 |
| 0.040775 | 0.071307 |
| 0.05726 | 0.005912 |
| 0.054944 | 0.007198 |
| -0.03178 | 0.010001 |
| 0.024712 | -0.00377 |
| 0.084599 | 0.028813 |
| 0.058881 | 0.05289 |
| -0.01014 | -0.10077 |
| 0.043435 | -0.06179 |
| 0.003769 | 0.017194 |
| 0.066597 | 0.025908 |
| 0.009846 | 0.031771 |
| 0.033782 | -0.0164 |
| 0.039179 | -0.00388 |
| 0.010852 | 0.000529 |
| 0.05249 | -0.01876 |
| 0.001695 | 0.02611 |
| 0.015629 | 0.054993 |
| 0.026817 | |
| 0.066856 | |
| 0.021219 | |
| 0.015945 | |
| 0.033449 | |
| 0.007143 | |
| 0.055287 | |
| -0.03091 | |
| 0.044618 | |
| 0.077373 | |
| 0.074359 | |
| 0.037604 | |
| 0.020675 | |
| 0.067662 | |
| 0.030212 | |
| 0.029878 | |
| 0.008448 | |
| 0.04276 | |
| 0.00136 | |
| 0.039289 | |

| |
|---|
| 0.051368 |
| 0.037157 |
| 0.039482 |
| 0.047074 |
| 0.040887 |
| 0.030236 |
| 0.030696 |
| 0.010358 |
| 0.01284 |
| 0.056232 |
| 0.022289 |
| 0.0119 |
| 0.020401 |
| 0.055233 |
| 0.016317 |
| 0.009122 |
| 0.029674 |
| 0.038987 |
| 0.050234 |
| 0.004406 |
| 0.023806 |
| 0.039119 |
| 0.030049 |
| 0.046682 |
| 0.031967 |
| 0.056335 |
| 0.016739 |
| 0.024941 |
| 0.051678 |
| 0.025596 |
| 0.024553 |
| 0.021597 |
| 0.004688 |
| 0.033502 |
| 0.072457 |
| 0.024336 |
| 0.046319 |
| 0.019336 |
| 0.043571 |
| 0.065226 |
| 0.026239 |
| 0.028279 |
| 0.053191 |
| 0.000695 |
| 0.032813 |
| 0.054765 |
| 0.037546 |

0.047788
-0.03786
0.065541
-0.00597
0.020956
0.086391
-0.00533
-0.00353
0.04209
0.028711
0.096789
0.030407
-0.02914
-0.00261
0.023347
0.054347
-0.00688
0.021401
0.00168
0.010261
0.014408
0.016887
0.008932
0.038109
-0.01458
0.052272
0.054742
0.049419
-0.02593
0.071921
-0.00725
0.079871
0.035226
0.018957
0.040223
0.04132
0.014289
0.036797
0.00125
0.058962
0.044004
0.041434
0.021994
0.035538
0.044888
0.013116
0.030541

0.079057
-0.00281
-0.01664
0.054302
-0.00494
0.069849
0.068124
0.055608
0.035256
0.08692
0.017672
0.058565
0.028973
0.051075
0.032697
0.016567
0.023842
0.020028
0.034841
0.029997
0.049961
0.044458
-0.0147
-0.05421
0.051326
0.032461
0.040501
0.047744
0.021598
0.041186
0.04163
0.09759
0.001285
0.081609
0.045421
0.01098
0.021561
0.058385
0.057541
0.01989
0.057872
0.028465
0.040255
0.0534
0.036838
0.003949
0.021244

0.03983
0.057863
0.019805
0.034798
-0.00455
-0.01384
0.058089
0.014738
0.081852
0.059132
0.026151
0.020581
0.054657
0.053981
0.047527
0.03472
0.048011
0.078604
0.01625
0.028438
0.023806
0.048993
0.048946
0.059611
-0.00465
0.027286
0.033883
0.03933
0.014197
-0.0098
-0.02409
0.021324
0.040851
0.030588
0.053811
0.033519
0.059605
0.038542
0.082029
0.026143
0.033998
0.007559
0.026975
0.046279
0.035279
0.013113
0.03398

0.032712
0.055474
0.000737
0.009466
0.016522
-0.00038
0.05451
0.01685
0.0528
0.036124
-0.0078
0.033606
0.059934
0.072648
0.034323
0.056908
0.083029
0.02072
0.073355
0.065527
0.010468
0.074036
0.018968
0.03407
0.038398
0.011473
0.068477
0.030725
0.057834
0.051285
0.022739
0.0651
0.026387
0.015295
0.042867
0.06178
0.012442
0.0269
0.018129
-0.02242
-0.01053
0.023421
0.030852
-0.02315
0.087186
0.0336
0.040699

0.023692
0.061051
0.045351
0.014218
0.061093
0.026244
0.041178
0.030351
0.005652
0.061896
0.081404
0.0579
0.020868
0.042757
0.084141
0.019279
0.076979
0.023275
0.013665
0.083955
0.008205
0.00435
0.038431
0.010188
0.061156
0.03787
0.040215
0.032194
0.024391
-0.02003
0.016518
-0.00736
0.032439
0.03723
0.008305
0.016748
0.065669
0.038497
0.035038
0.03687
0.070627
0.048025
0.038196
0.043298
0.036573
0.05389
0.070113

0.020201
0.014903
0.012981
0.036919
0.023474
0.037455
0.007605
0.034221
-0.01577
0.063324
0.084259
0.049965
0.039947
-0.0006
0.104105
0.011602
0.041099
0.025463
0.038019
0.037326
0.053496
0.034
0.067013
0.0384
0.071177
0.020766
0.043153
0.03055
0.027715
0.02041
0.011318
0.020118
0.028564
0.033989
0.054183
0.04309
0.017032
0.033733
0.019014
0.077717
0.063733
0.008635
0.000703
0.012759
0.059999
0.031789
0.023238

0.031984
0.031966
0.018771
0.041853
0.020164
0.022886
0.053442
0.048512
0.037134
0.060941
0.018569
0.034571
0.048983
0.022916
0.076425
-0.00165
0.038984
0.021752
0.073757
-0.03965
0.033566
0.030763
0.058297
0.022697
0.055731
0.028137
0.016582
0.05005
-0.00301
0.026349
0.055609
-0.00533
0.071754
-0.01902
0.024606
0.079784
0.069092
-0.00783
0.037034
0.004906
0.032912
0.023872
0.036556
0.046975
-0.00571
0.045593
-0.01242

| |
|---|
| 0.036983 |
| 0.021508 |
| 0.043042 |
| 0.053324 |
| 0.075404 |
| -0.03224 |
| 0.066403 |
| -0.00155 |
| 0.035061 |
| 0.011661 |
| 0.024022 |
| 0.024313 |
| 0.032479 |
| 0.061531 |
| 0.057243 |
| -0.02411 |
| 0.069629 |
| 0.047782 |
| 0.052022 |
| -0.04242 |
| 0.041871 |
| 0.019409 |
| 0.024395 |
| -0.01687 |
| 0.049797 |
| 0.047018 |
| 0.016513 |
| -0.01597 |
| 0.032344 |
| 0.055559 |
| 0.032302 |
| 0.012337 |
| 0.000314 |
| 0.025688 |
| 0.025039 |
| 0.030979 |
| 0.018837 |
| 0.043317 |
| 0.015412 |
| -0.02051 |
| 0.02575 |
| 0.037604 |
| 0.083967 |
| 0.012245 |
| 0.008301 |
| 0.004087 |
| 0.017253 |

0.009686
0.044016
0.008978
0.087723
0.042167
0.098872
0.004098
0.034288
0.051509
0.016805
0.043386
0.042166
0.024622
0.026199
0.074679
0.01125
0.010584
0.034374
0.112583
0.090418
0.034552
0.028904
0.081927
0.01518
0.008667
0.020504
0.053853
0.054087
0.025138
0.005298
0.034725
0.013733
-0.04045
0.00221
0.054903
0.046795
0.075178
0.052575
0.018367
0.081068
0.068727
0.014135
0.03014
0.048203
-0.01228
0.070883
0.030884

0.084487
0.067269
0.026514
0.005725
0.0162
0.029556
0.036175
0.039405
0.017253
0.038891
0.000567
0.061915
0.020376
0.076735
0.02878
0.026994
0.059936
0.017076
0.054371
0.062683
0.011205
0.06654
-0.01762
0.004983
0.004709
0.011636
0.029907
0.034663
0.045382
0.000658
0.040937
0.033127
0.049868
-0.00492
0.047978
0.02137
0.036486
0.008018
0.056754
-0.01877
0.034949
-0.01191
0.005067
0.046073
0.050607
0.094714
0.036405

-0.01583
0.02868
0.059689
0.050721
0.047778
0.085878
0.013262
0.034283
0.004369
-0.01297
0.045697
0.012751
0.038947
0.059415
0.039869
0.016485
0.014836
0.001259
0.031971
0.061363
0.040325
0.055489
0.026074
0.038172
0.018347
0.049763
0.035847
0.022427
0.010347
0.047823
0.030019
-0.00649
0.044392
0.047271
-0.0057
0.028153
0.014141
0.02596
0.015131
0.044441
0.040267
0.019792
0.084949
0.029175
0.045343
0.047267
0.038932

-0.00878
0.037867
0.028518
0.0855
0.06465
0.0314
0.039829
0.059667
0.00027
0.036947
0.013497
-0.00926
0.043106
0.074034
0.062098
0.026384
0.046532
0.022727
0.034509
0.034119
0.067647
0.0316
0.042113
0.069268
0.027707
0.047665
0.010143
0.059472
0.044705
0.011434
0.011357
0.026964
0.032977
-0.00021
-0.00775
0.041277
0.003801
0.015374
0.012633
-0.00861
0.048751
0.041844
0.072423
0.059945
0.06657
0.021479
0.061985

-0.01425
0.025492
0.014694
0.083471
0.024565
0.076027
0.034651
0.044634
0.035047
0.01129
0.024457
0.054219
0.062593
0.02233

**Subfigure:** Figure 9B

Plot Name: **Fibroblast Cortisol Score**

| | | |
|---|---|---|
| **Groups:** | WT4wk | Tg4wk |
| **Data Points:** | 0.029995 | 0.014949 |
| | 0.017353 | -0.00491 |
| | -0.01033 | -0.02941 |
| | -0.01952 | -0.03019 |
| | -0.04132 | -0.0362 |
| | 0.00737 | -0.0237 |
| | 0.002511 | -0.03167 |
| | 0.036916 | 0.002821 |
| | -0.01281 | 0.050313 |
| | 0.007762 | -0.038 |
| | 0.003065 | 0.019279 |
| | -0.01512 | -0.00833 |
| | 0.052073 | -0.03876 |
| | -0.02172 | -0.03077 |
| | -0.00429 | -0.03918 |
| | -0.01763 | -0.08156 |
| | 0.05033 | 0.002204 |
| | -0.03786 | 0.025403 |
| | 0.03056 | -0.0287 |
| | 0.01569 | -0.04396 |
| | 0.02411 | -0.00505 |
| | 0.018042 | 0.007277 |
| | 0.020313 | -0.0338 |
| | 0.041382 | 0.04679 |
| | 0.011145 | 0.019767 |
| | -0.00331 | -0.06716 |
| | -0.01783 | 0.009746 |
| | -0.01794 | -0.06137 |
| | -0.025 | -0.06205 |
| | 0.038385 | -0.01707 |
| | 0.001865 | -0.07031 |
| | 0.044186 | 0.03115 |
| | 0.016546 | -0.02682 |
| | 0.049378 | 0.000793 |
| | 0.007539 | -0.09593 |
| | -0.01153 | -0.05514 |
| | 0.080979 | -0.05495 |
| | 0.015691 | -0.0416 |
| | 0.007006 | -0.00706 |
| | 0.030002 | -0.03206 |
| | 0.031505 | -0.08984 |
| | -0.00335 | 0.011932 |
| | -0.01963 | -0.00752 |

| | |
|---|---|
| 0.015819 | -0.03521 |
| -0.01577 | -0.01815 |
| 0.045138 | 0.000976 |
| 0.003694 | 0.001517 |
| 0.017935 | -0.05282 |
| 0.034219 | -0.04684 |
| -0.01151 | -0.00163 |
| 0.001361 | 0.008718 |
| 0.02367 | 0.016059 |
| 0.001719 | -0.00747 |
| -0.00454 | 0.035118 |
| -0.023 | -0.00443 |
| -0.01924 | 0.003407 |
| 0.053611 | -0.07113 |
| 0.062656 | -0.08237 |
| 0.014967 | -0.01461 |
| -0.00579 | -0.03184 |
| 0.009466 | -0.03138 |
| 0.018734 | 0.027421 |
| 0.091392 | 0.007336 |
| 0.026426 | -0.01154 |
| 0.005725 | -0.08853 |
| 0.024246 | -0.0542 |
| 0.005029 | -0.04162 |
| 0.028417 | 0.000246 |
| 0.026905 | -0.01254 |
| -0.02715 | -0.05057 |
| 0.013948 | 0.020687 |
| 0.007985 | -0.00514 |
| 0.009804 | -0.02873 |
| -0.00355 | 0.007802 |
| -0.01466 | -0.00283 |
| -0.02196 | 0.004678 |
| 0.006813 | -0.0032 |
| 0.007248 | -0.03366 |
| 0.052272 | 0.052245 |
| -0.0232 | -0.05566 |
| 0.017454 | -0.03337 |
| 0.038509 | -0.06595 |
| -0.00316 | 0.046676 |
| -0.05598 | -0.02325 |
| 0.013233 | -0.00409 |
| 0.067987 | -0.05239 |
| 0.011899 | 0.003373 |
| 0.019194 | 0.028673 |
| 0.0211 | 0.018822 |
| 0.010154 | 0.000928 |

| | |
|---|---|
| -0.0369 | -0.00835 |
| -0.02373 | -0.0564 |
| 0.018328 | -0.07317 |
| 0.002696 | -0.01701 |
| 0.007221 | 0.034149 |
| 0.009758 | 0.009695 |
| 0.014544 | -0.02852 |
| 0.110407 | -0.05449 |
| 0.048968 | 0.008769 |
| 0.025081 | 0.037699 |
| 0.016407 | 0.010387 |
| -0.00951 | -0.0186 |
| 0.027767 | -0.07312 |
| 0.024856 | -0.00275 |
| -0.06181 | -0.04736 |
| -0.02127 | -0.03393 |
| 0.020757 | 0.015815 |
| -0.04038 | -0.06324 |
| -0.0096 | -0.02571 |
| 0.054227 | -0.09527 |
| -0.00797 | -0.02173 |
| -0.00811 | 0.040873 |
| -0.02806 | -0.05764 |
| -0.00749 | -0.03355 |
| -0.01208 | -0.05522 |
| 0.030638 | -0.04354 |
| 0.015012 | 0.020426 |
| -0.00025 | 0.030527 |
| -0.03239 | 0.07133 |
| -0.00881 | 0.035098 |
| -0.03173 | 0.013961 |
| -0.02791 | 0.01407 |
| -0.01535 | 0.009507 |
| 0.029267 | 0.017553 |
| -0.03658 | -0.01555 |
| 0.00162 | -0.03205 |
| -0.01861 | 0.039842 |
| 0.030475 | -0.01652 |
| -0.01307 | -0.01962 |
| 0.009324 | 0.03145 |
| -0.02337 | -0.03716 |
| -0.00267 | 0.019801 |
| 0.020893 | 0.044332 |
| 0.026995 | 0.0143 |
| -0.02812 | -0.00615 |
| 0.02871 | 0.020144 |
| 0.019446 | 0.009742 |

| | |
|---|---|
| 0.02486 | 0.014228 |
| 0.005392 | -0.00181 |
| 0.023375 | -0.07494 |
| 0.008017 | -0.02555 |
| -0.00588 | 0.002824 |
| 0.014485 | -0.08274 |
| -0.00971 | 0.023561 |
| 0.013956 | 0.040373 |
| -0.00365 | -0.0311 |
| 0.024912 | -0.0781 |
| 0.036424 | -0.01672 |
| -0.01873 | -0.04374 |
| -0.00926 | -0.01689 |
| -0.00204 | -0.02535 |
| -0.03673 | -0.00955 |
| -0.00816 | -0.01844 |
| 0.016801 | 0.004716 |
| 0.030408 | 0.042975 |
| 0.027117 | -0.03364 |
| -0.00491 | 0.032966 |
| -0.00774 | 0.004077 |
| -0.00159 | 0.02909 |
| 0.03163 | -0.06574 |
| -0.01677 | 0.038231 |
| -0.011 | 0.012895 |
| 0.004393 | 0.019537 |
| 0.058409 | 0.01766 |
| 0.028768 | 0.017187 |
| -0.01123 | -0.06463 |
| 0.097986 | 0.018931 |
| -0.01261 | -0.01059 |
| -0.02608 | 0.012862 |
| 0.051611 | -0.0066 |
| -0.01415 | -0.04038 |
| 0.010637 | -0.0843 |
| -0.0132 | 0.014023 |
| 0.05014 | 0.063485 |
| 0.038719 | 0.004308 |
| -0.01534 | -0.00871 |
| 0.006878 | 0.055344 |
| -0.00213 | -0.05146 |
| -0.00649 | 0.035203 |
| -0.05337 | -0.08944 |
| -0.03167 | -0.08786 |
| 0.020654 | 0.047658 |
| 0.023498 | -0.04543 |
| 0.035919 | 0.020301 |

| | |
|---|---|
| 0.00631 | -0.02056 |
| 0.041156 | 0.005537 |
| -0.01451 | -0.0165 |
| 0.012638 | 0.030811 |
| -0.00981 | -0.02809 |
| -0.00375 | -0.03763 |
| 0.027621 | -0.00491 |
| 0.056638 | -0.07248 |
| 0.035703 | -0.06239 |
| 0.015022 | -0.01931 |
| -0.0065 | -0.00375 |
| -0.0116 | -0.01647 |
| 0.004773 | -0.06047 |
| -0.02998 | 0.033145 |
| -0.01811 | -0.06233 |
| 0.018783 | 0.003462 |
| 0.009396 | -0.02732 |
| 0.053313 | -0.00787 |
| -0.00186 | -0.05543 |
| 0.017608 | 0.006839 |
| -0.03237 | -0.01471 |
| -0.0259 | -0.0164 |
| -0.03395 | -0.04019 |
| 0.033468 | 0.011798 |
| 0.062307 | 0.003144 |
| 0.006154 | 0.023079 |
| 0.047317 | -0.04124 |
| 0.016429 | -0.07353 |
| -0.01189 | -0.02376 |
| -0.04568 | -0.00106 |
| 0.025085 | 0.051511 |
| 0.021249 | 0.03179 |
| 0.008422 | -0.00082 |
| 0.043068 | 0.020861 |
| -0.0016 | 0.053695 |
| 0.054572 | 0.023558 |
| -0.00811 | -0.04994 |
| 0.00289 | -0.06704 |
| 0.014521 | 0.027691 |
| -0.01518 | -0.03494 |
| 0.040789 | 0.01629 |
| 0.031628 | -0.03736 |
| 0.014408 | 0.016318 |
| -0.00413 | 0.036011 |
| -0.04129 | -0.03988 |
| -0.00963 | 0.052142 |
| 0.029757 | 0.041604 |

| | |
|---|---|
| 0.005181 | 0.062163 |
| 0.034223 | 0.05614 |
| -0.00999 | -0.02453 |
| 0.002788 | 0.022635 |
| -0.01435 | -0.00638 |
| 0.043259 | -0.00734 |
| 0.035752 | 0.017966 |
| -0.03344 | -0.07193 |
| 0.029273 | -0.0215 |
| -0.02025 | -0.00855 |
| -0.00276 | -0.00683 |
| -0.0277 | 0.017441 |
| -0.03052 | -0.01031 |
| 0.020917 | 0.038305 |
| -0.01567 | -0.03488 |
| -0.0017 | -0.02467 |
| 0.043002 | 0.043179 |
| 0.022803 | 0.023684 |
| -0.00023 | -0.00689 |
| 0.016331 | 0.024032 |
| 0.054447 | 0.044026 |
| 0.003531 | -0.03518 |
| -0.02131 | -0.00442 |
| -0.03741 | -0.05636 |
| -0.03042 | 0.006745 |
| 0.0072 | 0.029229 |
| -0.01324 | 0.000823 |
| -0.00438 | -0.01869 |
| 0.012718 | 0.024713 |
| 0.01243 | -0.00631 |
| -0.04549 | 0.013541 |
| -0.00616 | 0.034882 |
| 0.019781 | 0.025559 |
| -0.01257 | 0.0325 |
| -0.0081 | 0.066353 |
| 0.016341 | -0.07824 |
| 0.025334 | -0.03924 |
| 0.018583 | -0.085 |
| -0.01817 | -0.01034 |
| -0.01407 | 0.022069 |
| -0.01979 | 0.072926 |
| 0.058106 | 0.019981 |
| 0.007885 | 0.000517 |
| 0.012964 | 0.023295 |
| 0.031002 | 0.030538 |
| 0.044239 | -0.0164 |
| 0.017165 | -0.01728 |

| | |
|---|---|
| -0.0197 | -0.00524 |
| 0.069837 | 0.040916 |
| -0.01224 | -0.02519 |
| 0.044143 | 0.03625 |
| -0.01043 | -0.01632 |
| -0.03455 | 0.005468 |
| 0.020118 | -0.02532 |
| -0.00989 | -0.01954 |
| -0.01032 | -0.0209 |
| 0.018585 | 0.013245 |
| 0.030088 | -0.10281 |
| -0.00753 | -0.05279 |
| 0.011286 | 0.043692 |
| 0.06263 | -0.00873 |
| 0.047554 | -0.04135 |
| -0.01723 | -0.00399 |
| 0.061116 | 0.010799 |
| 0.02003 | -0.02687 |
| 0.015856 | -0.10747 |
| 0.030967 | -0.02765 |
| -0.02057 | -0.00325 |
| 0.004485 | 0.00325 |
| -0.01345 | 0.03971 |
| 0.057751 | -0.00087 |
| -0.01826 | 0.029057 |
| 0.004174 | 0.029119 |
| 0.033789 | 0.014466 |
| 0.010389 | -0.02119 |
| 0.001821 | -0.02151 |
| -0.00274 | -0.03127 |
| 0.06305 | 0.009136 |
| 0.026057 | 0.003321 |
| 0.027791 | -0.0094 |
| 0.017617 | -0.00053 |
| -0.00665 | 0.030105 |
| 0.004975 | 0.002672 |
| 0.001253 | -7.63E-06 |
| 0.003881 | -0.02817 |
| -0.03637 | -0.0101 |
| 0.014235 | 0.004511 |
| 0.031372 | 0.007977 |
| 0.012781 | 0.011857 |
| -0.00815 | -0.07926 |
| -0.02522 | 0.001552 |
| 0.029339 | -0.02351 |
| -0.02166 | -0.01158 |
| -0.00318 | -0.05819 |

| | |
|---|---|
| -0.00377 | -0.03781 |
| 0.064833 | -0.02738 |
| 0.002342 | -0.02151 |
| 0.002519 | -0.01925 |
| -0.00482 | 0.011226 |
| 0.064444 | -0.06966 |
| -0.01443 | 0.01536 |
| -0.01059 | 0.000745 |
| -0.0209 | 0.021662 |
| 0.022038 | 0.0319 |
| -0.01265 | -0.0108 |
| -0.04902 | 0.04 |
| 0.014221 | 0.006528 |
| 0.005315 | -0.00171 |
| 0.027083 | 0.023677 |
| -0.02973 | 0.032684 |
| -0.04122 | -0.05484 |
| 0.009272 | 0.013434 |
| -0.03003 | 0.061366 |
| -0.01041 | -0.05036 |
| 0.00367 | -0.07212 |
| 0.03797 | 0.021723 |
| -0.00255 | -0.00239 |
| -0.01336 | 0.01539 |
| 0.065202 | 0.022558 |
| 0.006102 | 0.012751 |
| -0.00899 | -0.000074 |
| 0.033725 | 0.058732 |
| 0.011482 | 0.022314 |
| 0.016853 | -0.00682 |
| -0.02833 | -0.0716 |
| 0.002688 | 0.030449 |
| -0.02283 | -0.00787 |
| -0.01359 | 0.058807 |
| 0.017522 | 0.058563 |
| 0.018436 | 0.080096 |
| 0.019174 | -0.05769 |
| -0.02756 | -0.00936 |
| -0.00472 | -0.00314 |
| 0.000534 | 0.040353 |
| -0.01278 | -0.01738 |
| 0.001194 | -0.0753 |
| 0.06517 | 0.002667 |
| 0.008629 | 0.008689 |
| -0.0207 | -0.01 |
| -0.06045 | -0.0258 |
| 0.00091 | -0.02409 |

| | |
|---|---|
| 0.00327 | -0.01567 |
| 0.053912 | -0.00928 |
| 0.006748 | 0.022306 |
| 0.022835 | -0.02561 |
| 0.022739 | -0.01026 |
| -0.00605 | -0.05607 |
| -0.02982 | -0.0063 |
| -0.03151 | -0.04258 |
| 0.006906 | 0.009938 |
| -0.05414 | -0.05632 |
| -0.00942 | -0.06425 |
| 0.028667 | -0.00553 |
| -0.02185 | -0.01355 |
| 0.015023 | -0.00961 |
| 0.073789 | 0.006401 |
| 0.006555 | 0.028888 |
| 0.012591 | -0.00765 |
| -0.01089 | 0.006238 |
| 0.033311 | -0.05507 |
| 0.072566 | -0.08305 |
| 0.014845 | -0.04403 |
| -0.00572 | -0.00757 |
| -0.04772 | -0.11101 |
| 0.022002 | 0.003563 |
| 0.046629 | -0.01889 |
| -0.04033 | 0.029145 |
| 0.063282 | -0.07093 |
| 0.033975 | -0.06163 |
| 0.031709 | -0.00453 |
| 0.028027 | 0.026264 |
| 0.023969 | -0.03569 |
| 0.000866 | 0.046637 |
| 0.002911 | -0.01182 |
| -0.00528 | -0.02786 |
| 0.046101 | 0.001456 |
| 0.02703 | -0.02899 |
| -0.0414 | 0.047758 |
| 0.012343 | -0.0713 |
| -0.01063 | -0.0413 |
| 0.001269 | -0.00026 |
| -0.03285 | -0.01888 |
| -0.0291 | -0.04749 |
| -0.0508 | -0.07461 |
| 0.000356 | 0.030796 |
| 0.011235 | 0.002348 |
| 0.0000481 | -0.02276 |
| 0.011041 | 0.034391 |

| | |
|---|---|
| 0.027224 | 0.015162 |
| 0.010448 | 0.038466 |
| -0.00479 | -0.03549 |
| 0.004155 | -0.06784 |
| -0.03366 | -0.03456 |
| 0.017199 | 0.006003 |
| 0.02504 | -0.05643 |
| -0.00043 | 0.034024 |
| -0.0426 | -0.03744 |
| 0.009111 | 0.031357 |
| 0.040965 | -0.009 |
| 0.023393 | 0.030074 |
| 0.041199 | -0.0012 |
| 0.01596 | 0.038454 |
| -0.02794 | -0.0789 |
| -0.01796 | 0.013862 |
| 0.037165 | -0.03532 |
| 0.030206 | 0.025574 |
| 0.015738 | 0.053379 |
| 0.06794 | 0.024217 |
| 0.01941 | -0.02426 |
| -0.00685 | -0.0525 |
| 0.017227 | -0.02126 |
| 0.031425 | 0.047419 |
| 0.041524 | -0.04373 |
| 0.056121 | -0.01405 |
| -0.06855 | 0.006152 |
| 0.021904 | -0.02568 |
| 0.053771 | -0.05191 |
| -0.00852 | 0.048791 |
| 0.008621 | 0.006251 |
| 0.004846 | -0.01792 |
| 0.007434 | -0.01492 |
| 0.013043 | 0.014582 |
| -0.00197 | -0.04956 |
| 0.012943 | -0.06889 |
| -0.02141 | -0.02752 |
| -0.01786 | -0.03543 |
| -0.02723 | 0.07616 |
| 0.049021 | 0.057532 |
| 0.004301 | -0.01338 |
| 0.034286 | 0.016098 |
| 0.029037 | -0.0036 |
| -0.02799 | -0.03484 |
| -0.01281 | 0.02836 |
| -0.02101 | 0.033816 |
| 0.007948 | -0.07766 |

| | |
|---|---|
| -0.01851 | -0.01384 |
| -0.0352 | 0.027177 |
| 0.001245 | -0.09328 |
| -0.01366 | 0.016361 |
| -0.01222 | -0.05098 |
| 0.006458 | 0.004257 |
| 0.012845 | 0.027474 |
| 0.011994 | 0.021322 |
| -0.01089 | -0.04269 |
| 0.01398 | -0.00555 |
| 0.006668 | 0.057122 |
| -0.01055 | -0.04999 |
| 0.007826 | -0.06137 |
| -0.02844 | -0.04436 |
| 0.023698 | 0.065492 |
| 0.019088 | -0.03039 |
| 0.013978 | -0.02291 |
| -0.04061 | -0.00702 |
| -0.00709 | 0.012641 |
| 0.001755 | -0.02558 |
| 0.031963 | 0.002816 |
| 0.0361 | -0.01663 |
| 0.012934 | 0.036301 |
| 0.006157 | 0.025211 |
| 0.035928 | -0.05425 |
| -0.02349 | 0.013858 |
| -0.01008 | 0.026276 |
| -0.02897 | -0.0327 |
| -0.03574 | -0.00798 |
| 0.034211 | 0.004571 |
| -0.05348 | 0.019316 |
| -0.00993 | -0.01926 |
| -0.01618 | 0.036421 |
| 0.020605 | 0.030308 |
| 0.031305 | -0.04417 |
| 0.017575 | 0.034207 |
| -0.05344 | -0.00437 |
| 0.025337 | -0.02931 |
| 0.044906 | -0.00927 |
| 0.034679 | -0.06874 |
| 0.062973 | -0.06212 |
| 0.030486 | -0.04862 |
| 0.023081 | 0.052007 |
| 0.001378 | 0.027369 |
| 0.0032 | -0.01011 |
| 0.018536 | 0.066338 |
| 0.043353 | -0.06445 |

| | |
|---|---|
| -0.01634 | -0.08701 |
| -0.00232 | 0.036128 |
| 0.024459 | -0.01502 |
| 0.019671 | -0.02428 |
| -0.04817 | -0.00782 |
| 0.043441 | 0.005208 |
| 0.026693 | -0.01964 |
| -0.01741 | 0.020761 |
| 0.025606 | -0.03514 |
| -0.0046 | -0.00339 |
| -0.01208 | 0.023176 |
| -0.02849 | -0.07806 |
| -0.00726 | -0.00803 |
| -0.01834 | 0.01819 |
| 0.057469 | 0.005724 |
| 0.020111 | -0.06536 |
| -0.01831 | -0.00716 |
| 0.048206 | 0.036661 |
| 0.007909 | -0.00419 |
| -0.03311 | 0.032557 |
| -0.02511 | 0.004274 |
| 0.023686 | 0.007384 |
| 0.032317 | -0.0252 |
| 0.02024 | -0.04209 |
| 0.031491 | -0.05528 |
| -0.02162 | -0.03425 |
| -0.0042 | -0.06252 |
| -0.06273 | -0.04696 |
| 0.007503 | -0.02567 |
| -0.02236 | 0.034197 |
| 0.003633 | -0.03018 |
| 0.069263 | -0.02316 |
| 0.023142 | -0.06392 |
| -0.05798 | -0.03186 |
| -0.01064 | -0.0429 |
| -0.00311 | 0.019211 |
| 0.030261 | -0.00165 |
| -0.02028 | -0.02312 |
| 0.022821 | 0.04043 |
| 0.010497 | 0.008816 |
| -0.01205 | -0.00643 |
| 0.049926 | -0.04444 |
| 0.022935 | -0.09791 |
| -0.00231 | -0.01181 |
| 0.036927 | -0.00205 |
| -0.011 | 0.008798 |
| 0.011689 | 0.016484 |

| | |
|---|---|
| 0.040333 | 0.029571 |
| 0.009916 | 0.030438 |
| 0.039929 | 0.014812 |
| 0.041596 | -0.01632 |
| 0.011576 | -0.02021 |
| 0.041226 | 0.018574 |
| 0.012034 | -0.07288 |
| -0.01371 | -0.02314 |
| 0.014716 | 0.015926 |
| 0.041147 | -0.03494 |
| 0.033874 | 0.002268 |
| 0.022147 | -0.09092 |
| -0.0028 | -0.03088 |
| 0.040301 | 0.015742 |
| -0.01785 | -0.04086 |
| 0.036138 | 0.046026 |
| -0.03484 | 0.024666 |
| -0.02595 | 0.021991 |
| 0.016337 | -0.06401 |
| 0.042366 | 0.005377 |
| -0.04208 | 0.003339 |
| 0.013886 | -0.00503 |
| 0.026487 | -0.03196 |
| 0.011804 | -0.05748 |
| 0.005769 | -0.04588 |
| -0.01401 | 0.051491 |
| 0.027214 | -0.03379 |
| 0.015079 | 0.035764 |
| 0.035705 | -0.0162 |
| 0.023055 | -0.0044 |
| 0.022983 | -0.03974 |
| -0.00314 | -0.00674 |
| -0.00971 | -0.00324 |
| -0.00958 | -0.02751 |
| 0.002907 | 0.039123 |
| 0.035457 | 0.006002 |
| -0.03796 | -0.01647 |
| -0.0369 | -0.05469 |
| -0.00703 | -0.05313 |
| -0.01416 | 0.031379 |
| -0.00883 | -0.0227 |
| -0.00556 | 0.014577 |
| 0.033597 | -0.04916 |
| 0.024277 | -0.00978 |
| -0.00607 | 0.00782 |
| -0.00314 | 0.018094 |
| -0.01372 | -0.06826 |

| | |
|---|---|
| -0.03245 | -0.0773 |
| -0.02895 | -0.01238 |
| 0.023599 | 0.04529 |
| 0.028494 | 0.000726 |
| -0.02873 | -0.00981 |
| 0.012078 | 0.023705 |
| 0.047109 | 0.047875 |
| 0.043448 | -0.05085 |
| -0.02389 | 0.020094 |
| 0.011002 | 0.01925 |
| -0.02231 | -0.01106 |
| -0.00753 | 0.007912 |
| -0.00402 | -0.02139 |
| -0.00411 | 0.019506 |
| -0.03284 | -0.02255 |
| -0.05331 | -0.05648 |
| -0.03908 | 0.023849 |
| 0.050463 | -0.00935 |
| -0.00152 | 0.041389 |
| 0.006602 | 0.04048 |
| -0.01144 | 0.01806 |
| -0.00351 | 0.035283 |
| 0.005818 | -0.02232 |
| -0.00259 | -0.02523 |
| -0.04658 | 0.045208 |
| -0.01352 | 0.018072 |
| 0.005313 | -0.03932 |
| 0.041587 | -0.01546 |
| -0.00873 | -0.05448 |
| 0.050216 | -0.01307 |
| 0.02548 | -0.04467 |
| 0.031505 | 0.028704 |
| 0.039681 | 0.018647 |
| -0.01455 | 0.033046 |
| 0.007428 | -0.01125 |
| 0.074891 | -0.02563 |
| 0.001165 | 0.051425 |
| 0.023611 | -0.03015 |
| 0.036962 | 0.026572 |
| 0.047388 | 0.065035 |
| -0.02676 | -0.01821 |
| -0.01809 | 0.016794 |
| 0.00985 | -0.04883 |
| 0.030228 | 0.013436 |
| 0.017579 | -0.06533 |
| 0.024796 | -0.01487 |
| -0.04146 | -0.10443 |

| | |
|---|---|
| 0.030415 | -0.07008 |
| 0.026625 | -0.00021 |
| 0.006716 | 0.055229 |
| 0.001173 | 0.010304 |
| 0.031019 | 0.004706 |
| -0.0103 | -0.00349 |
| -0.02205 | 0.01854 |
| 0.033329 | -0.0171 |
| -0.0265 | -0.04287 |
| -0.0002 | -0.00351 |
| 0.018427 | 0.055589 |
| -0.01218 | 0.029715 |
| 0.010292 | 0.065171 |
| 0.036503 | 0.049923 |
| -0.02077 | -0.064 |
| -0.04694 | -0.06783 |
| 0.014961 | 0.041747 |
| -0.00573 | 0.034224 |
| 0.03475 | 0.022103 |
| -0.04007 | 0.032275 |
| 0.009017 | -0.09022 |
| 0.002332 | -0.01845 |
| 0.037114 | 0.039869 |
| 0.001519 | 0.062485 |
| 0.035912 | 0.050827 |
| 0.016666 | 0.004223 |
| -0.00399 | 0.05294 |
| 0.044697 | -0.01809 |
| -0.04747 | -0.00618 |
| -0.04186 | 0.021753 |
| 0.009632 | 0.003318 |
| 0.01954 | 0.010779 |
| 0.024804 | -0.02866 |
| 0.038021 | -0.03162 |
| -0.02469 | 0.026905 |
| -0.02572 | -0.00883 |
| -0.02901 | 0.024951 |
| -0.03958 | -0.08045 |
| -0.00675 | -0.0328 |
| 0.051701 | -0.03892 |
| 0.0502 | -0.00836 |
| 0.044702 | -0.0439 |
| 0.01101 | 0.05463 |
| 0.016101 | 0.054106 |
| 0.036509 | -0.04736 |
| 0.013977 | -0.06619 |
| 0.058782 | -0.04772 |

| | |
|---|---|
| 0.038486 | -0.0495 |
| -0.0061 | -0.00211 |
| -0.01393 | -0.00393 |
| 0.007297 | -0.02238 |
| 0.024314 | 0.023915 |
| 0.019153 | 0.038917 |
| 0.042183 | -0.03954 |
| 0.06795 | -0.03359 |
| 0.001466 | 0.071853 |
| -0.00651 | -0.02337 |
| -0.02087 | -0.0244 |
| -0.0045 | -0.03947 |
| -0.02043 | -0.01126 |
| -0.04326 | 0.058232 |
| 0.022167 | 0.017614 |
| 0.061454 | 0.008177 |
| 0.027412 | -0.03982 |
| -0.04199 | -0.02121 |
| 0.021084 | -0.04116 |
| -0.05104 | -0.0173 |
| -0.00852 | -0.02334 |
| 0.036058 | -0.0154 |
| 0.036808 | 0.040496 |
| -0.00697 | -0.02953 |
| 0.012015 | -0.05066 |
| 0.029479 | 0.017062 |
| 0.071305 | -0.01806 |
| -0.02744 | -0.0033 |
| 0.045823 | 0.013976 |
| -0.00999 | 0.01134 |
| 0.030385 | 0.042351 |
| -0.0459 | 0.030053 |
| -0.0381 | -0.0332 |
| -0.03018 | 0.013675 |
| -0.02364 | 0.017289 |
| -0.00393 | 0.004993 |
| -0.05871 | 0.020516 |
| 0.020506 | 0.046201 |
| 0.003006 | 0.011425 |
| 0.004227 | -0.00388 |
| 0.00735 | 0.011284 |
| 0.033544 | -0.00636 |
| 0.000711 | -0.02113 |
| -0.02088 | 0.032116 |
| 0.018583 | -0.01142 |
| 0.007909 | 0.011813 |
| 0.015964 | -0.00609 |

| | |
|---|---|
| 0.02528 | -0.00576 |
| -0.00937 | -0.05322 |
| 0.045968 | -0.04433 |
| 0.024794 | -0.02913 |
| -0.05402 | -0.04193 |
| 0.024244 | 0.042727 |
| 0.092068 | 0.031438 |
| 0.001749 | -0.00677 |
| -0.00149 | 0.00126 |
| 0.007851 | -0.01238 |
| -0.04306 | 0.002519 |
| 0.0000361 | -0.05491 |
| 0.001303 | -0.07203 |
| 0.046565 | -0.04555 |
| 0.015182 | 0.054919 |
| 0.035259 | -0.08453 |
| -0.01514 | 0.017874 |
| 0.008319 | 0.046372 |
| 0.016276 | 0.022253 |
| -0.02142 | 0.020907 |
| -0.04729 | 0.071051 |
| -0.02785 | -0.04873 |
| -0.00655 | -0.00617 |
| -0.00379 | 0.036908 |
| -0.01845 | 0.042499 |
| 0.022664 | -0.03864 |
| 0.000179 | -0.02853 |
| -0.00305 | -0.07578 |
| 0.027171 | 0.048993 |
| 0.009219 | -0.03058 |
| -0.01974 | 0.01449 |
| -0.02352 | -0.00483 |
| 0.036536 | 0.05393 |
| -0.03902 | 0.072538 |
| 0.043562 | -0.04849 |
| 0.017717 | 0.023867 |
| -0.05467 | 0.024224 |
| -0.01157 | -0.02003 |
| 0.009339 | -0.03676 |
| 0.01289 | -0.0011 |
| 0.034973 | -0.00168 |
| 0.043563 | -0.06152 |
| -0.00453 | 0.007593 |
| 0.020932 | -0.01145 |
| -0.01522 | 0.039684 |
| -0.00642 | 0.037798 |
| -0.03381 | -0.0177 |

| | |
|---|---|
| 0.002679 | -0.00972 |
| -0.02031 | 0.009285 |
| 0.026642 | 0.035947 |
| -0.00313 | 0.036704 |
| 0.026785 | -0.04028 |
| 0.014906 | 0.027025 |
| -0.02357 | 0.035171 |
| -0.04603 | -0.00868 |
| -0.00178 | 0.000355 |
| 0.029597 | 0.015272 |
| -0.00426 | 0.001123 |
| -0.00558 | -0.03229 |
| 0.000345 | -0.02881 |
| 0.027885 | 0.040284 |
| -0.00949 | -0.01277 |
| 0.039293 | 0.036946 |
| 0.007202 | -0.07021 |
| 0.006277 | 0.004212 |
| -0.00413 | 0.003763 |
| -0.04144 | -0.04905 |
| -0.02978 | -0.02097 |
| 0.042774 | -0.01638 |
| -0.01249 | -0.05424 |
| -0.08473 | 0.007486 |
| 0.004899 | -0.07078 |
| 0.010496 | 0.003536 |
| -0.01582 | -0.00621 |
| -0.00714 | -0.00276 |
| 0.016723 | -0.00577 |
| 0.005754 | 0.006319 |
| -0.00562 | 0.010003 |
| -0.0126 | -0.00815 |
| -0.0119 | 0.055567 |
| 0.024624 | 0.014448 |
| 0.00036 | 0.031633 |
| 0.01529 | 0.020866 |
| -0.03388 | -0.01573 |
| 0.041045 | 0.037093 |
| 0.012297 | -0.01257 |
| 0.002902 | -0.12593 |
| 0.014117 | -0.00888 |
| -0.04599 | -0.01748 |
| 0.020458 | -0.0926 |
| -0.0029 | -0.0363 |
| -0.03413 | -0.00602 |
| 0.051314 | 0.020361 |
| -0.02111 | 0.036783 |

| | |
|---|---|
| -0.00388 | -0.03705 |
| 0.01866 | -0.01477 |
| 0.021719 | 0.024319 |
| 0.017732 | -0.04475 |
| -0.02485 | 0.033029 |
| 0.015181 | 0.050182 |
| 0.03576 | 0.011095 |
| -0.00248 | 0.0114 |
| 0.042928 | 0.030579 |
| 0.005251 | -0.01334 |
| 0.000806 | -0.07418 |
| -0.01204 | -0.0595 |
| 0.024279 | 0.020561 |
| 0.006078 | -0.02216 |
| -0.0367 | 0.021617 |
| -0.01974 | -0.05799 |
| -0.0082 | 0.002697 |
| -0.01932 | -0.00734 |
| 0.06542 | 0.027847 |
| 0.022601 | -0.08643 |
| 0.01809 | -0.00508 |
| -0.00713 | 0.010128 |
| -0.01113 | -0.01285 |
| -0.01633 | 0.015881 |
| 0.005498 | -0.02203 |
| -0.00175 | 0.009808 |
| -0.00324 | -0.012 |
| 0.000942 | -0.02712 |
| 0.016243 | 0.009076 |
| 0.033712 | 0.010022 |
| 0.028864 | -0.02302 |
| 0.006464 | 0.073 |
| -0.0494 | -0.07252 |
| -0.01241 | 0.007898 |
| 0.007831 | -0.01053 |
| -0.01805 | -0.0236 |
| 0.013439 | 0.029938 |
| 0.012009 | 0.017924 |
| -0.00659 | 0.071531 |
| -0.00549 | -0.017 |
| 0.022585 | -0.06242 |
| 0.003878 | -0.07835 |
| 0.075135 | -0.00739 |
| 0.024866 | 0.001725 |
| 0.041648 | 0.019641 |
| -0.00802 | -0.03978 |
| 0.018843 | -0.01395 |

| | |
|---|---|
| 0.000782 | 0.009296 |
| -0.02066 | 0.06601 |
| -0.00132 | 0.039275 |
| -0.02606 | -0.0037 |
| 0.035986 | -0.02107 |
| 0.031009 | -0.00439 |
| -0.00668 | 0.057552 |
| 0.005593 | 0.021455 |
| 0.011868 | 0.018181 |
| 0.033583 | -0.02296 |
| -0.02785 | 0.091117 |
| -0.01859 | 0.009752 |
| -0.00017 | -0.02679 |
| 0.037147 | -0.05897 |
| 0.00259 | -0.00669 |
| 0.028699 | -0.04514 |
| -0.03493 | 0.002239 |
| -0.00178 | -0.07529 |
| 0.02579 | 0.02746 |
| -0.0181 | -0.00881 |
| 0.011531 | 0.013153 |
| 0.004907 | -0.09008 |
| -0.01719 | -0.08947 |
| 0.003834 | -0.02105 |
| 0.013653 | 0.011761 |
| 0.05841 | -0.03736 |
| 0.011039 | -0.02417 |
| 0.043794 | -0.02736 |
| 0.027624 | 0.013545 |
| 0.035723 | -0.00494 |
| 0.020608 | -0.00434 |
| 0.006581 | 0.000551 |
| -0.01102 | 0.009445 |
| 0.007342 | 0.005686 |
| -0.00225 | -0.00279 |
| 0.031854 | 0.034872 |
| 0.047297 | -0.00739 |
| 0.014968 | 0.003677 |
| 0.041121 | -0.04427 |
| 0.014419 | -0.04425 |
| 0.047583 | -0.0158 |
| -0.00235 | 0.00422 |
| 0.025669 | -0.0141 |
| -0.03779 | 0.033257 |
| 0.00124 | 0.015646 |
| -0.02542 | -0.00061 |
| -0.00308 | 0.018966 |

| | |
|---|---|
| 0.028461 | -0.05421 |
| -0.02088 | 0.063467 |
| 0.010183 | 0.047339 |
| -0.00341 | -0.04831 |
| -0.02121 | -0.03765 |
| 0.014973 | 0.018604 |
| 0.080386 | -0.07334 |
| -0.00183 | -0.04271 |
| -0.01944 | 0.010155 |
| -0.02063 | -0.00164 |
| 0.023653 | -0.01349 |
| -0.00836 | 0.010023 |
| 0.050661 | -0.01661 |
| -0.06132 | 0.048491 |
| -0.05596 | -0.05957 |
| -0.0024 | -0.02904 |
| 0.013441 | -0.09607 |
| 0.058319 | -0.04981 |
| 0.018184 | -0.00922 |
| -0.00049 | -0.02696 |
| -0.00953 | 0.0312 |
| 0.011955 | 0.018562 |
| 0.006857 | 0.008162 |
| 0.034048 | -0.06952 |
| 0.003775 | -0.01806 |
| 0.022561 | 0.002624 |
| 0.003121 | -0.01944 |
| -0.01212 | 0.023773 |
| 0.022607 | -0.04846 |
| -0.01168 | 0.012819 |
| 0.026397 | 0.036265 |
| -0.03145 | 0.011712 |
| 0.021733 | 0.026383 |
| 0.034852 | -0.04441 |
| 0.015625 | -0.00723 |
| 0.002847 | -0.00252 |
| 0.005556 | -0.02477 |
| 0.021511 | 0.010862 |
| -0.03294 | -0.04187 |
| -0.01217 | -0.08153 |
| -0.04649 | 0.019395 |
| 0.000532 | -0.06518 |
| -0.00084 | 0.008753 |
| 0.035572 | 0.032091 |
| -0.00372 | 0.027129 |
| -0.04344 | -0.03165 |
| 0.022516 | -0.04118 |

| | |
|---|---|
| -0.03515 | -0.0088 |
| -0.00616 | -0.01876 |
| -0.04798 | 0.024724 |
| -0.0048 | -0.06235 |
| -0.01993 | -0.05154 |
| 0.054064 | -0.02385 |
| 0.033728 | -0.04285 |
| 0.00997 | -0.0849 |
| 0.0108 | 0.009774 |
| 0.005285 | -0.00976 |
| 0.022851 | 0.035401 |
| 0.035914 | -0.05705 |
| -0.00333 | -0.01297 |
| 0.04685 | -0.0643 |
| 0.008077 | -0.02431 |
| -0.00365 | -0.00581 |
| 0.012461 | -0.05484 |
| 0.008182 | 0.025167 |
| -0.00692 | -0.01413 |
| 0.01934 | -0.03124 |
| 0.020717 | 0.026868 |
| 0.012341 | -0.04777 |
| -0.03068 | 0.022975 |
| 0.020264 | 0.011103 |
| 0.03499 | 0.03759 |
| 0.007126 | 0.005535 |
| 0.034781 | -0.05367 |
| -0.02866 | -0.02315 |
| 0.026531 | -0.00319 |
| 0.044521 | -0.03669 |
| -0.01042 | 0.066314 |
| -0.03628 | -0.04164 |
| -0.01376 | -0.0181 |
| -0.03298 | -0.0324 |
| -0.01172 | -0.02433 |
| 0.027647 | 0.006389 |
| -0.00975 | -0.06437 |
| 0.027146 | 0.004291 |
| 0.0029 | 0.019 |
| 0.062591 | -0.00791 |
| -0.0497 | -0.0375 |
| 0.021728 | 0.021876 |
| 0.02223 | -0.01859 |
| 0.008142 | 0.024749 |
| -0.00559 | 0.043461 |
| 0.012494 | 0.035344 |
| 0.050838 | 0.002041 |

| | |
|---|---|
| 0.016408 | 0.0583 |
| 0.013474 | 0.010909 |
| -0.02723 | 0.038059 |
| 0.038212 | 0.001858 |
| 0.023859 | 0.019283 |
| 0.012384 | 0.005566 |
| 0.03977 | -0.0239 |
| 0.002833 | 0.039695 |
| -0.00831 | -0.03738 |
| -0.02656 | -0.01756 |
| -0.02308 | -0.02321 |
| -0.00719 | -0.01965 |
| -0.01058 | -0.06956 |
| -0.00355 | 0.049957 |
| -0.02463 | 0.037702 |
| -0.05514 | 0.021681 |
| 0.009493 | -0.01202 |
| 0.056129 | 0.030593 |
| 0.005062 | -0.0965 |
| -0.02802 | 0.014408 |
| -0.0161 | -0.00076 |
| -0.01629 | 0.01444 |
| -0.00069 | -0.0193 |
| -0.01321 | -0.02329 |
| 0.013612 | -0.03519 |
| 0.015812 | -0.0521 |
| 0.013871 | 0.067587 |
| 0.03262 | -0.00413 |
| 0.029793 | -0.02785 |
| 0.007338 | 0.046817 |
| 0.014957 | -0.0555 |
| 0.017025 | -0.00702 |
| -0.05446 | -0.01206 |
| -0.00801 | -0.03424 |
| 0.032344 | 0.020339 |
| -0.00221 | -0.01315 |
| 0.01558 | 0.022958 |
| 0.023383 | -0.04248 |
| -0.00251 | 0.021274 |
| -0.04307 | -0.01736 |
| 0.02597 | -0.04361 |
| 0.023978 | 0.005083 |
| -0.00801 | 0.047125 |
| 0.016304 | -0.0396 |
| 0.036257 | -0.02193 |
| 0.014051 | -0.02064 |
| -0.04981 | -0.00463 |

| | |
|---|---|
| -0.02107 | 0.012625 |
| -0.02323 | -0.06197 |
| -0.02428 | -0.05429 |
| -0.02159 | 0.033235 |
| 0.034374 | -0.04783 |
| -0.02926 | -0.03534 |
| -0.00551 | 0.027614 |
| -0.01953 | -0.00416 |
| 0.025371 | 0.012332 |
| 0.011723 | 0.019335 |
| 0.028582 | -0.07058 |
| -0.01142 | -0.01922 |
| 0.020714 | 0.013177 |
| -0.0047 | -0.03876 |
| 0.023327 | 0.002915 |
| -0.00808 | -0.06841 |
| 0.043209 | -0.02308 |
| 0.051556 | -0.0034 |
| 0.006377 | 0.038996 |
| -0.03434 | 0.010075 |
| 0.004181 | 0.007065 |
| -0.03092 | 0.055886 |
| -0.00465 | 0.0392 |
| -0.04181 | -0.01265 |
| 0.075285 | -0.0436 |
| 0.000694 | 0.041086 |
| -0.02675 | -0.01273 |
| -0.00308 | -0.0282 |
| 0.009932 | -0.01848 |
| 0.016511 | 0.039298 |
| 0.061532 | 0.03613 |
| 0.0356 | 0.040365 |
| 0.005335 | 0.022516 |
| 0.021752 | -0.00917 |
| 0.059801 | -0.00949 |
| 0.009344 | 0.007614 |
| -0.04339 | 0.021309 |
| -0.0037 | -0.06266 |
| -0.01566 | 0.001887 |
| 0.03118 | -0.06055 |
| 0.049646 | -0.09352 |
| 0.018734 | 0.023983 |
| -0.02463 | 0.026034 |
| -0.00567 | -0.05935 |
| 0.019596 | 0.124267 |
| -0.03793 | -0.02768 |
| 0.020205 | -0.05178 |

| | |
|---|---|
| 0.004241 | -0.00172 |
| -0.03141 | -0.04372 |
| 0.029892 | -0.02221 |
| -0.00526 | -0.04692 |
| -0.01086 | -0.02487 |
| -0.03185 | 0.000387 |
| 0.029188 | -0.03265 |
| 0.020361 | -0.00329 |
| 0.020893 | 0.025457 |
| 0.024886 | 0.042166 |
| -0.01551 | -0.05031 |
| 0.041471 | -0.0768 |
| 0.028686 | 0.014023 |
| 0.033223 | -0.06761 |
| -0.00108 | -0.03972 |
| -0.01281 | -0.02937 |
| 0.012633 | 0.019733 |
| 0.017859 | 0.027861 |
| -0.00334 | 0.008097 |
| 0.005547 | 0.055025 |
| 0.012508 | 0.002836 |
| 0.085699 | 0.002207 |
| 0.000371 | -0.02311 |
| 0.028224 | 0.006956 |
| -0.00498 | -0.06213 |
| 0.029322 | 0.000274 |
| -0.04513 | -0.01472 |
| -0.00626 | -0.01951 |
| 0.02241 | -0.02166 |
| 0.064718 | 0.008902 |
| -0.00599 | -0.01368 |
| 0.00915 | -0.04561 |
| -0.00931 | -0.01139 |
| 0.041292 | -0.02008 |
| 0.016246 | -0.01329 |
| 0.043505 | 0.017821 |
| 0.025264 | 0.025633 |
| -0.04079 | 0.032763 |
| -0.01596 | 0.025659 |
| 0.036194 | 0.006367 |
| 0.027113 | -0.034 |
| -0.02933 | 0.029444 |
| 0.023557 | -0.0768 |
| 0.02347 | 0.062881 |
| -0.02818 | -0.092 |
| -0.01746 | -0.03715 |
| -0.0096 | -0.05881 |

| | |
|---|---|
| 0.06746 | -0.04322 |
| -0.01955 | 0.006451 |
| 0.015416 | 0.03373 |
| 0.04181 | 0.012493 |
| 0.010754 | 0.005147 |
| 0.017184 | 0.031762 |
| 0.057892 | -0.01811 |
| 0.0217 | 0.027881 |
| 0.038944 | -0.01546 |
| 0.022888 | -0.0654 |
| 0.001702 | -0.03342 |
| 0.028936 | -0.0183 |
| 0.000427 | -0.02381 |
| 0.003627 | -0.0334 |
| 0.013509 | -0.00387 |
| 0.038014 | -0.07481 |
| 0.029778 | 0.006765 |
| 0.023562 | 0.011504 |
| 0.039428 | -0.01595 |
| -0.01298 | -0.05572 |
| 0.042243 | -0.00016 |
| 0.052948 | -0.03015 |
| -0.03395 | -0.03504 |
| -0.04204 | 0.013521 |
| 0.011742 | -0.03712 |
| 0.055789 | -0.02901 |
| 0.089194 | -0.08932 |
| -0.0133 | -0.07403 |
| -0.04527 | 0.016797 |
| -0.0154 | 0.057309 |
| -0.03442 | -0.05658 |
| -0.01076 | 0.011464 |
| 0.000241 | -0.01598 |
| -0.00422 | 0.01929 |
| 0.029294 | -0.0241 |
| 0.081383 | -0.07416 |
| -0.00473 | -0.04808 |
| -0.01639 | 0.044479 |
| -0.01781 | -0.0576 |
| -0.0603 | -0.0533 |
| 0.003491 | -0.06027 |
| 0.041996 | -0.04989 |
| 0.073025 | -0.05724 |
| 0.02691 | -0.04175 |
| 0.005519 | 0.051934 |
| 0.012323 | -0.07914 |
| 0.025708 | 0.021403 |

| | |
|---|---|
| -0.0194 | 0.000848 |
| 0.001493 | 0.019162 |
| 0.015159 | 0.02909 |
| -0.00143 | -0.06956 |
| -0.04838 | 0.030517 |
| -0.05619 | -0.03501 |
| -0.01056 | -0.07998 |
| -0.00441 | 0.019623 |
| 0.012732 | 0.006548 |
| 0.025575 | 0.010124 |
| 0.01611 | 0.009513 |
| -0.04847 | 0.0266 |
| 0.006525 | 0.011407 |
| 0.004014 | 0.049384 |
| 0.009786 | -0.01921 |
| -0.00116 | 0.048176 |
| 0.002641 | -0.03744 |
| 0.040727 | 0.010742 |
| -0.02665 | 0.038817 |
| 0.011999 | -0.00856 |
| 0.0412 | 0.006291 |
| 0.015429 | 0.006077 |
| -0.02461 | -0.01958 |
| 0.04054 | 0.004128 |
| 0.009356 | -0.01466 |
| -0.03036 | -0.02709 |
| 0.006084 | -0.029 |
| -0.02821 | 0.041304 |
| 0.027968 | -0.00744 |
| -0.00998 | 0.007505 |
| -0.02817 | -0.06838 |
| 0.00421 | -0.074 |
| -0.00631 | -0.02346 |
| 0.009504 | -0.03159 |
| 0.015791 | 0.041526 |
| -0.00355 | -0.00641 |
| -0.02757 | 0.011481 |
| 0.058751 | -0.02866 |
| -0.00891 | -0.05018 |
| 0.031873 | -0.00815 |
| 0.015836 | -0.01337 |
| 0.034653 | 0.041972 |
| 0.041121 | -0.01258 |
| 0.04958 | -0.03802 |
| 0.025279 | 0.004541 |
| 0.00843 | 0.028153 |
| 0.003491 | -0.03753 |

| | |
|---|---|
| 0.027085 | -0.0292 |
| -0.00584 | 0.015928 |
| -0.02602 | -0.00111 |
| 0.055785 | 0.003163 |
| -0.00069 | -0.03729 |
| -0.00375 | 0.045044 |
| 0.062662 | -0.01302 |
| 0.03683 | 0.060433 |
| 0.030817 | 0.02249 |
| 0.034769 | 0.023096 |
| 0.004566 | 0.009869 |
| 0.021431 | 0.01383 |
| -0.00124 | -0.06368 |
| 0.036102 | 0.015571 |
| -0.01971 | 0.035845 |
| 0.031301 | -0.01031 |
| -0.00363 | -0.07248 |
| -0.00444 | 0.003809 |
| 0.046571 | 0.031326 |
| -0.01359 | -0.00554 |
| 0.009341 | 0.027616 |
| 0.018694 | -0.07771 |
| -0.02493 | 0.020911 |
| -0.03491 | -0.04125 |
| -0.00597 | -0.05078 |
| 0.017154 | -0.00317 |
| -0.00994 | 0.028636 |
| 0.030322 | 0.013798 |
| 0.001784 | 0.007468 |
| -0.01539 | -0.09143 |
| 0.051097 | -0.0384 |
| 0.016593 | -0.0679 |
| -0.01061 | 0.023323 |
| 0.001551 | -0.0083 |
| 0.019055 | -0.01748 |
| -0.02544 | 0.032826 |
| -0.01077 | 0.00062 |
| 0.008464 | -0.03847 |
| -0.0271 | 0.026363 |
| -0.02128 | 0.01338 |
| -0.01971 | -0.06903 |
| 0.020313 | 0.022517 |
| -0.00712 | -0.02033 |
| -0.0096 | 0.014143 |
| 0.04043 | 0.013541 |
| 0.019794 | 0.001276 |
| -0.0146 | -0.03413 |

| | |
|---|---|
| -0.02461 | -0.01342 |
| 0.046174 | -0.05027 |
| 0.050757 | -0.04199 |
| -0.0655 | -0.00332 |
| 0.035192 | -0.01619 |
| 0.051109 | -0.00694 |
| -0.00381 | -0.00744 |
| -0.02262 | 0.012389 |
| 0.007875 | -0.01517 |
| -0.01601 | 0.011837 |
| 0.006896 | -0.00564 |
| -0.00185 | 0.056667 |
| -0.04291 | -0.04184 |
| 0.025991 | 0.041786 |
| 0.029845 | 0.040191 |
| -0.02428 | -0.01821 |
| -0.02667 | -0.00969 |
| 0.002231 | -0.0571 |
| 0.017955 | -0.01099 |
| 0.001534 | -0.05076 |
| -0.01407 | -0.04487 |
| -0.04323 | -0.04418 |
| -0.01883 | 0.001901 |
| 0.060725 | 0.044066 |
| 0.004469 | -0.01695 |
| 0.030217 | -0.05194 |
| 0.028405 | 0.010476 |
| -0.00393 | -0.02566 |
| 0.0000397 | -0.05902 |
| 0.010552 | 0.061982 |
| -0.02179 | -0.0236 |
| 0.02969 | 0.009436 |
| 0.011851 | -0.02852 |
| 0.03274 | 0.012437 |
| 0.00366 | 0.033812 |
| -0.00249 | 0.002474 |
| 0.040237 | 0.011294 |
| -0.0141 | -0.10386 |
| -0.02508 | 0.057986 |
| -0.04937 | 0.018419 |
| -0.00548 | -0.04625 |
| 0.014478 | 0.0017 |
| -0.05554 | -0.04192 |
| 0.000474 | 0.042173 |
| 0.015061 | 0.00416 |
| -0.02482 | -0.01184 |
| -0.00624 | 0.003281 |

| | |
|---|---|
| 0.011711 | -0.00962 |
| 0.033235 | -0.0397 |
| 0.012887 | 0.004795 |
| -0.00942 | -0.07248 |
| 0.022324 | 0.0301 |
| -0.01267 | 0.036719 |
| 0.018417 | -0.0466 |
| 0.018224 | -0.06413 |
| -0.01211 | -0.0497 |
| 0.002131 | -0.01927 |
| 0.004752 | 0.031229 |
| -0.01457 | -0.01168 |
| 0.051036 | 0.003044 |
| 0.010872 | -0.03908 |
| -0.00033 | 0.054272 |
| -0.00347 | -0.01317 |
| 0.030809 | 0.017877 |
| 0.00471 | -0.02126 |
| 0.025778 | 0.029313 |
| 0.026716 | 0.000449 |
| 0.001443 | 0.017026 |
| -0.0123 | 0.074781 |
| -0.02683 | -0.01305 |
| 0.003073 | -0.00384 |
| -0.0243 | 0.032023 |
| 0.024956 | -0.03449 |
| 0.016058 | -0.04488 |
| -0.00663 | -0.00468 |
| -0.00933 | 0.010179 |
| 0.016309 | 0.00718 |
| 0.003589 | -0.02805 |
| -0.00574 | 0.085495 |
| -0.00433 | -0.00417 |
| -0.0022 | -0.02625 |
| 0.020907 | -0.03037 |
| 0.001015 | -0.00186 |
| 0.029961 | 0.025224 |
| -0.0184 | -0.05106 |
| 0.018595 | 0.030641 |
| 0.035337 | -0.06551 |
| 0.049579 | -0.01141 |
| 0.0144 | 0.008006 |
| 0.034979 | 0.028871 |
| -0.014 | 0.008215 |
| 0.026727 | 0.007278 |
| 0.008312 | -0.02177 |
| 0.001521 | 0.013401 |

| | |
|---|---|
| 0.038427 | 0.027294 |
| 0.026961 | -0.01713 |
| -0.01329 | 0.008076 |
| 0.036618 | -0.01197 |
| -0.02481 | -0.00533 |
| -0.04197 | -0.0000587 |
| 0.022175 | 0.008873 |
| -0.0172 | -0.01037 |
| 0.052946 | -0.04327 |
| -0.06476 | 0.022994 |
| 0.018031 | 0.050429 |
| 0.041083 | -0.05406 |
| 0.010054 | -0.05418 |
| 0.019409 | 0.025035 |
| -0.00658 | -0.01952 |
| -0.01634 | -0.01582 |
| -0.00866 | 0.017052 |
| 0.016636 | -0.05524 |
| 0.027273 | 0.030268 |
| 0.007452 | 0.036562 |
| 0.018622 | 0.018064 |
| 0.04391 | 0.040037 |
| -0.01245 | -0.0432 |
| 0.015107 | -0.00658 |
| -0.00857 | 0.004644 |
| 0.003322 | 0.014728 |
| -0.00931 | -0.04418 |
| 0.026273 | 0.013039 |
| 0.035653 | -0.08436 |
| -0.00546 | -0.02309 |
| -0.05015 | 0.01801 |
| -0.00678 | -0.02269 |
| -0.00357 | -0.00271 |
| -0.00425 | 0.037191 |
| 0.018228 | -0.01947 |
| 0.022912 | 0.011907 |
| 0.016474 | 0.025121 |
| 0.030462 | -0.03471 |
| 0.016349 | 0.007267 |
| 0.003498 | 0.04269 |
| 0.030826 | -0.03861 |
| 0.06782 | -0.05221 |
| 0.010093 | 0.008387 |
| -0.01515 | 0.010658 |
| -0.00287 | -0.0401 |
| 0.03232 | -0.02815 |
| -0.01179 | -0.01217 |

| | |
|---|---|
| -0.01206 | 0.010596 |
| 0.011429 | 0.009657 |
| -0.02606 | 0.014988 |
| 0.033098 | -0.03424 |
| 0.024609 | 0.00455 |
| 0.043278 | -0.02143 |
| -0.02307 | 0.008032 |
| -0.02042 | -0.06301 |
| -0.01451 | -0.0196 |
| -0.02274 | 0.040555 |
| 0.006451 | 0.005885 |
| -0.00586 | -0.10324 |
| 0.010733 | 0.039401 |
| -0.02722 | 0.020728 |
| -0.00583 | -0.06017 |
| -0.01881 | -0.04102 |
| -0.00083 | 0.008868 |
| -0.03878 | 0.000849 |
| -0.05508 | -0.02909 |
| 0.00488 | 0.033451 |
| 0.002463 | -0.03998 |
| 0.020271 | -0.03558 |
| -0.01227 | -0.02673 |
| -0.06455 | -0.02496 |
| 0.022505 | 0.014346 |
| 0.004899 | -0.06801 |
| 0.014623 | -0.01148 |
| 0.003267 | 0.009784 |
| -0.00415 | -0.00579 |
| 0.02777 | -0.01396 |
| 0.012724 | -0.02965 |
| 0.036424 | -0.00138 |
| -0.04769 | -0.02885 |
| -0.02679 | -0.08769 |
| 0.035647 | -0.05385 |
| 0.001749 | 0.003191 |
| -0.02124 | -0.0271 |
| 0.013595 | -0.05554 |
| 0.010118 | 0.001337 |
| 0.038664 | 0.06578 |
| 0.000391 | -0.02804 |
| -0.00903 | 0.05292 |
| -0.00809 | -0.0065 |
| -0.02623 | -0.02686 |
| -0.0018 | 0.037762 |
| -0.02497 | 0.013063 |
| -0.00085 | 0.043014 |

| | |
|---|---|
| 0.038109 | -0.06443 |
| 0.02678 | 0.010051 |
| 0.022217 | -0.02138 |
| -0.03619 | -0.0503 |
| 0.003271 | -0.03123 |
| 0.004031 | -0.04105 |
| -0.02341 | -0.04594 |
| -0.00096 | -0.01632 |
| -0.03187 | -0.03017 |
| 0.017072 | -0.03065 |
| -0.02382 | -0.0285 |
| 0.017056 | -0.08279 |
| -0.01064 | 0.021327 |
| -0.02236 | 0.03259 |
| -0.05417 | 0.027229 |
| -0.02756 | -0.01786 |
| 0.065645 | -0.0486 |
| -0.01415 | -0.07257 |
| 0.035055 | 0.029056 |
| -0.01148 | 0.056253 |
| 0.011816 | -0.00601 |
| -0.00693 | 0.006831 |
| 0.027657 | 0.011761 |
| 0.093822 | -0.00047 |
| 0.009036 | -0.04835 |
| -0.00237 | 0.009709 |
| -0.03662 | -0.02003 |
| 0.024214 | -0.05938 |
| 0.019615 | 0.042991 |
| -0.03576 | -0.01492 |
| -0.01494 | -0.03278 |
| 0.05066 | -0.03876 |
| 0.033339 | 0.014116 |
| 0.012453 | -0.03615 |
| -0.01778 | 0.029374 |
| 0.041571 | 0.007041 |
| -0.02116 | 0.00485 |
| -0.01352 | 0.022765 |
| 0.072344 | 0.004948 |
| 0.009306 | -0.04037 |
| 0.033484 | 0.019259 |
| -0.03214 | -0.02102 |
| -0.05615 | 0.025444 |
| -0.05016 | 0.027645 |
| -0.00046 | -0.03516 |
| 0.039698 | -0.10087 |
| 0.002218 | 0.036499 |

| | |
|---|---|
| -0.04858 | -0.06749 |
| 0.007768 | 0.002054 |
| 0.030337 | -0.03808 |
| -0.01815 | 0.009022 |
| 0.005574 | 0.012902 |
| 0.030629 | 0.009931 |
| 0.003943 | |
| 0.009849 | |
| -0.00263 | |
| -0.00171 | |
| 0.020496 | |
| -0.00178 | |
| 0.061003 | |
| 0.019642 | |
| -0.032 | |
| -0.0322 | |
| -0.03433 | |
| 0.038785 | |
| -0.01156 | |
| -0.01873 | |
| 0.011608 | |
| -0.01251 | |
| 0.023518 | |
| -0.02165 | |
| -0.01048 | |
| -0.02592 | |
| -0.00726 | |
| -0.03164 | |
| -0.01798 | |
| 0.001405 | |
| -0.0119 | |
| -0.00465 | |
| -0.00574 | |
| -0.0000184 | |
| 0.000239 | |
| 0.005111 | |
| 0.042466 | |
| -0.05023 | |
| 0.01698 | |
| 0.004736 | |
| -0.00059 | |
| -0.00123 | |
| -0.00281 | |
| 0.021151 | |
| 0.042024 | |
| 0.038694 | |
| -0.0008 | |

0.013513
0.016979
-0.01588
-0.01625
0.000906
-0.00744
0.006511
0.023581
-0.03486
0.017132
0.042499
0.01345
0.049614
-0.00347
-0.0151
-0.00318
0.009969
0.029508
0.009945
-0.01124
0.040429
0.013759
0.027642
0.028184
-0.00371
0.028226
-0.00928
-0.01069
0.025102
-0.05446
0.006065
0.005315
0.060597
-0.01116
0.012121
0.039508
-0.01796
0.016557
-0.0066
-0.00507
0.016386
0.021513
-0.01046
0.005422
-0.01114
0.012645
0.052238

-0.01324
0.021425
0.036142
-0.0000963
0.032561
0.005627
0.059325
0.007134
0.008141
-0.02218
-0.00714
0.031404
-0.01518
0.018696
0.040661
0.04403
0.005595
0.005191
0.035051
0.020026
0.060544
-0.04492
0.054226
0.01973
0.015253
-0.01963
-0.01497
-0.0155
-0.0075
0.006105
0.011936
0.000735
0.012108
-0.03229
0.013701
-0.00696
-0.00014
0.000358
-0.07412
0.030032
-0.03092
0.007
-0.00696
0.023131
0.014067
0.022159
-0.00346

0.015151
0.029484
0.002165
0.013689
0.027703
0.022219
-0.00374
-0.0362
0.017974
-0.01221
0.030093
0.036567
0.021601
0.016743
0.048045
-0.01429
0.000465
-0.02209
0.00989
0.051587
0.002546
0.00895
0.01095
0.035237
-0.00097
-0.00938
0.045249
-0.01283
0.034772
0.018519
-0.024
-0.02601
-0.01284
0.035381
0.048963
0.028535
0.040816
-0.05
0.028721
-0.00411
-0.00154
0.007535
0.003543
0.038424
0.014189
0.038675
0.023714

0.029174
-0.02457
-0.00306
0.038564
-0.01456
0.008643
0.019473
-0.00258
0.035713
0.0188
0.013634
-0.01955
-0.01682
-0.00835
0.020363
0.037826
0.015312
0.017612
0.010981
0.019049
-0.02122
0.015957
-0.06186
-0.01229
0.002579
-0.0178
0.053895
0.061691
0.019573
-0.01091
0.023111
-0.00744
0.006033
0.024857
-0.05132
-0.01414
-0.00396
-0.00309
-0.0000613
-0.01174
-0.03202
-0.03925
0.00472
-0.01162
0.033587
0.022388
-0.00792

0.019959
0.028597
0.004552
-0.00761
0.036046
0.051466
-0.02072
0.014931
-0.01672
0.006269
-0.04842
-0.0423
-0.02361
0.032065
-0.00861
0.002976
0.015968
-0.0187
-0.00296
-0.02569
0.014882
0.039072
-0.01024
-0.01099
0.031588
0.007034
0.017334
0.005147
-0.02655
-0.03277
-0.04316
0.025271
0.016245
0.023873
0.000387
0.047595
-0.01107
0.005041
-0.0419
0.000309
-0.03836
-0.02983
0.010228
0.024296
0.004692
-0.0449
-0.00337

0.028905
-0.01778
-0.03261
-0.00905
0.003936
0.049078
0.040924
0.012361
-0.00632
-0.01437
0.006715
0.040592
0.027535
-0.01949
-0.00436
0.015169
-0.01243
0.04483
0.058561
0.0286
0.006285
-0.01636
0.024375
-0.00476
-0.01472
0.007366
0.002875
-0.0257
-0.01731
-0.04831
-0.01543
-0.04081
-0.00976
0.038066
-0.02184
0.023757
0.018859
-0.01543
0.000209
0.036467
0.018352
-0.01232
-0.00427
-0.05341
0.014627
-0.02514
0.009349

0.025731
0.028942
0.015674
0.001961
-0.02235
0.04436
0.0305
-0.02817
0.00949
0.022488
-0.00596
0.02649
-0.00088
-0.00933
0.059311
-0.00148
0.019727
-0.0113
-0.00627
-0.00886
-0.02045
0.012368
-0.03121
0.006415
0.034372
-0.01705
0.027611
0.02114
0.028227
-0.02278
0.031743
0.013099
-0.02212
-0.03775
-0.02765
0.002586
0.004721
-0.02312
-0.01921
0.035784
-0.0132
0.041981
-0.03502
0.041026
-0.03259
0.03509
-0.02291

-0.01406
0.008825
0.008006
-0.01276
-0.00755
-0.00243
0.020833
-0.00385
-0.02453
-0.00992
-0.00101
-0.0001
-0.01545
0.019745
-0.00738
0.005476
0.032768
0.017305
0.010507
0.009289
-0.0051
0.031506
0.002706
0.022393
0.013167
-0.02336
-0.01523
0.010241
-0.025
0.010519
-0.00329
-0.01158
0.034171
0.030687
0.043039
-0.02185
-0.00308
0.016014
-0.02638
0.004281
0.002152
-0.04471
0.027339
0.000271
0.001124
0.038471
-0.0161

0.010474
0.048796
0.019617
0.062758
0.020428
0.071442
0.040399
0.014784
0.006763
0.037237
0.008281
0.032899
-0.05007
0.003458
-0.01364
0.001945
-0.03481
-0.05335
-0.01676
0.008119
-0.00468
-0.02377
-0.01026
-0.01136
-0.00277
-0.00181
0.044005
-0.01894
0.008972
-0.00068
0.010805
0.008536
-0.00763
0.028744
-0.01185
0.065445
0.016523
0.009092
0.00409
0.023058
0.012473
-0.00624
-0.03292
-0.00242
0.055838
0.003876
0.035508

0.027344
-0.00385
-0.01832
-0.01519
-0.00378
0.01073
-0.01635
-0.05448
-0.00272
0.038034
-0.01094
0.040081
0.017689
-0.02931
0.02245
0.004374
0.028119
0.021047
-0.03156
-0.01137
-0.0183
0.014843
0.003017
0.025633
-0.01987
0.007058
-0.01997
-0.00162
0.03332
-0.0321
-0.03628
0.022478
0.022754
-0.02473
-0.04594
0.024747
0.015138
-0.0143
0.02268
-0.02087
-0.04128
-0.04797
0.034636
-0.01479
0.043234
0.043935
0.019865

0.035075
0.021777
-0.04324
-0.0019
0.011896
-0.00323
0.007126
0.021063
-0.00354
-0.01754
-0.0234
-0.00591
-0.03361
0.02582
-0.00504
-0.01657
0.053105
0.060806
-0.01488
-0.00119
-0.02541
0.026174
-0.00601
-0.00681
-0.04846
0.00427
0.002156
0.037634
0.006188
-0.01903
-0.02584
0.017671
0.03067
-0.02632
-0.01749
-0.04465
0.000738
-0.02394
-0.00485
0.009679
0.081458
0.044751
0.014416
-0.03477
-0.00395
-0.01461
-3.44E-06

0.05583
-0.01456
-0.00899
-0.00084
0.021215
-0.02243
0.00865
0.043675
-0.02132
0.003515
0.071505
0.04361
-0.00478
0.015565
0.052264
-0.00659
-0.02344
0.030565
0.024205
-0.04574
-0.00815
-0.02774
0.057275
0.015954
0.025488
0.037243
0.000289
0.058933
0.037115
0.007427
0.022013
-0.02766
-0.00123
0.006086
0.020209
-0.00591
0.018811
0.027199
-0.01607
0.02208
0.019585
0.017776
0.024376
-0.00442
0.017835
0.007275
0.005128

0.016802
-0.01655
-0.00605
0.048834
0.029914
0.014181
-0.01588
-0.00264
-0.03352
0.065987
-0.03144
0.009983
0.034787
-0.00933
-0.01534
0.000499
0.045144
0.052771
0.006848
-0.00689
0.017346
0.026448
0.034483
0.000762
0.003349
0.044354
0.011328
0.030296
0.003897
0.012721
-0.02455
0.016598
0.032157
0.053693
0.042118
0.061022
-0.00255
-0.02071
0.003213
0.020683
-0.01517
0.005066
-0.00688
-0.01012
-0.00258
0.029409
0.04206

-0.00713
0.024754
0.01499
0.001797
0.046023
-0.03044
0.023974
0.004033
0.004015
-0.02145
-0.02084
0.04664
0.03826
0.001914
0.071468
-0.02112
0.043343
-0.00372
0.019969
-0.03789
-0.00973
-0.01869
0.022561
0.022718
-0.06225
-0.01344
0.010409
0.029838
0.00918
-0.00285
0.038698
-0.01686
0.030257
-0.05855
0.044569
0.004245
-0.00539
0.001448
0.033973
0.008932
-0.02182
0.03502
-0.02187
0.013812
-0.03034
0.001778
0.000919

-0.00847
0.03305
-0.02088
0.010915
0.021242
0.015055
-0.01568
-0.00306
-0.04767
0.007309
0.0461
-0.00735
0.042881
-0.01687
-0.0002
0.024562
0.038087
0.003719
0.024745
-0.0908
0.016507
-0.00559
0.029924
-0.08399
0.002342
0.049875
-0.00729
0.015271
0.012141
-0.01247
-0.02806
0.067102
0.015155
0.008355
0.019885
-0.00285
-0.01373
-0.00411
0.001622
0.030319
0.036767
0.01069
-0.0399
-0.03548
0.027345
0.027257
-0.01495

0.02181
-0.01022
-0.0197
0.016547
-0.0489
0.054473
-0.00741
0.002772
0.038335
0.0234
0.001706
0.011801
-0.0204
0.002394
-0.00194
0.019961
-0.00462
-0.02841
-0.03097
0.054341
-0.01827
0.010568
-0.02541
0.026662
0.020445
0.009161
0.042364
0.016006
0.029715
0.010428
0.024349
0.013922
-0.03009
0.034589
-0.03124
-0.01503
0.025962
0.009985
0.030932
-0.03326
0.027658
0.003197
0.017922
0.038591
0.029316
0.014589
0.044645

0.032078
0.073341
-0.0061
-0.03282
-0.00259
0.044837
0.004105
-0.03778
0.018521
0.016222
0.023429
-0.03086
-0.02001
0.044494
-0.00945
0.000218
0.02547
0.016993
0.052866
-0.01418
-0.00285
-0.00937
0.012556
0.040177
-0.00954
0.022557
-0.0072
0.021687
0.020548
-0.03152
0.005713
0.020848
-0.02013
0.026024
-0.02278
0.002294
0.030912
0.012805
-0.01952
0.032613
-0.03022
-0.00358
-0.03086
-0.00536
0.002905
-0.02068
-0.01793

0.028709
-0.01421
0.014171
0.028613
-0.01649
0.01342
-0.04265
-0.0376
0.025632
-0.01761
0.030169
0.01487
0.014805
-0.01456
-0.00956
0.021735
0.0134
0.016852
-0.00868
-0.00514
0.04572
-0.03807
-0.01076
-0.02468
-0.00165
-0.03628
-0.01259
-0.01878
-0.01107
0.007812
0.063053
0.015968
0.04299
-0.00959
0.010832
-0.0101
0.053077
-0.04334
0.023768
-0.00286
-0.04806
-0.0069
0.024949
-0.04819
0.02257
-0.00379
-0.03143

0.030983
-0.05375
0.019352
0.03163
-0.03007
0.020708
-0.00237
-0.02728
0.033828
-0.02736
-0.04348
-0.00017
-0.00845
0.045418
0.015553
-0.02278
0.020384
0.016608
0.047216
0.015819
-0.00714
-0.01348
-0.01294
0.051183
-0.00892
0.016741
0.057096
0.027351
0.044985
-0.00402
0.032176
-0.01073
-0.03145
0.031034
-0.02043
-0.00404
-0.02653
0.005918
-0.00475
-0.01817
0.023111
-0.01867
0.004777
0.022778
0.029664
0.003153
0.001223

| |
|---|
| -0.00116 |
| -0.02794 |
| 0.016645 |
| -0.02196 |
| 0.011004 |
| 0.05722 |
| -0.00698 |
| 0.034285 |
| 0.000431 |
| -0.01632 |
| 0.005826 |
| 0.050485 |
| 0.035705 |
| 0.026312 |
| 0.023027 |
| -0.00853 |
| -0.00486 |
| -0.02335 |
| 0.015228 |
| -0.00127 |
| -0.03741 |
| 0.023015 |
| -0.01219 |
| -0.02465 |
| 0.047277 |
| 0.016631 |
| 0.059172 |
| 0.016827 |
| 0.035112 |
| -0.04478 |
| 0.035387 |
| 0.010846 |
| 0.015216 |
| -0.03308 |
| -0.00169 |
| -0.02166 |
| -0.02406 |
| -0.00605 |
| 0.002913 |
| 0.025465 |
| 0.001198 |
| 0.007338 |
| 0.034853 |
| -0.01302 |
| -0.01348 |
| -0.01674 |
| 0.032074 |

0.024486
0.036945
-0.02714
0.014249
0.012805
0.016156
-0.02084
-0.00272
0.025044
0.038253
-0.02532
0.008848
0.000442
-0.00874
0.034819
-0.01594
0.010561
-0.02838
0.034769
-0.01835
0.023219
-0.00519
-0.0302
0.032082
0.008115
0.021379
0.000956
-0.04299
0.045176
-0.02018
0.027397
-0.028
-0.02635
0.027805
-0.00689
0.020834
0.056897
-0.05282
0.060306
0.029479
0.032989
-0.03819
-0.01238
-0.00879
0.028546
0.002322
-0.02936

-0.00473
-0.01234
0.014683
0.011952
0.041223
0.005219
0.023778
0.038352
-0.02058
0.009232
-0.02932
-0.0208
0.032767
0.015658
0.043609
-0.0026
-0.01739
-0.00689
0.022955
0.013325
0.019041
-0.02979
-0.00485
-0.01192
0.03735
0.001154
0.026806
-0.01033
-0.01008
0.042115
0.015795
0.012652
-0.0043
0.016879
0.020051
0.080191
-0.01877
-0.01316
0.038152
0.004989
-0.02741
-0.01696
0.028578
-0.01819
0.013011
0.041193
0.021861

0.041926
0.032881
0.019139
0.006943
0.008989
0.020419
0.009099
-0.01233
0.026895
0.043118
0.022566
0.012153
-0.01825
0.003201
-0.01284
-0.01543
-0.00746
0.014058
-0.00217
-0.08044
-0.02748
-0.00041
0.01505
0.015347
-0.01137
0.025734
0.006967
-0.00906
-0.03654
0.005229
0.019536
0.051491
0.028316
-0.0283
0.013506
0.007822
-0.04621
-0.01524
-0.05224
0.015682
-0.00738
-0.01151
0.011523
-0.01659
0.022517
0.012613
-0.04604

0.033826
-0.02547
0.025126
-0.02837
-0.04598
0.008303
-0.01311
-0.03691
-0.00979
0.043792
-0.03012
0.004406
0.027965
-0.00642
-0.0472
-0.00227
0.064134
-0.0007
-0.0492
-0.00629
0.009751
0.02198
0.015617
-0.00407
-0.02306
0.017411
0.038353
-0.03045
0.046474
0.046689
-0.00046
0.014228
-0.01061
0.021766
-0.01406
0.015265
0.008516
0.016904
0.037217
0.031101
0.051696
-0.05854
0.025543
0.029058
-0.00194
0.00281
0.019811

0.004841
0.004204
-0.02164
-0.00275
0.017848
0.046699
-0.03599
0.056553
-0.02174
0.006481
-0.01743
-0.02338
-0.00054
-0.0198
0.001135
0.022078
-0.01119
0.002779
-0.0654
0.034794
-0.0089
0.022319
0.027324
0.046755
0.029941
0.029711
0.021612
0.0125
0.029807
-0.05905
0.00874
0.038075
0.052379
0.007459
0.007543
0.031596
-0.02634
-0.00211
0.048569
0.01861
0.003175
-0.0005
0.002579
-0.04783
0.021659
-0.00566
0.018688

0.012248
0.011098
-0.03153
0.016951
0.035036
0.01753
-0.02269
0.032635
-0.02483
-0.01254
0.032254
0.017409
0.006993
-0.00463
-0.01021
0.015397
0.025287
0.008435
0.039783
0.013699
-0.03206
-0.0049
0.003204
0.012445
0.004344
0.020961
0.006515
0.002719
-0.01119
0.008083
-0.02563
-0.00803
0.031639
0.001312
-0.02572
-0.01247
-0.02679
0.044103
-0.03327
0.034418
-0.00717
0.081688
-0.02861
0.049334
-0.00785
0.00269
0.004767

| |
|---|
| -0.01396 |
| 0.022658 |
| 0.01063 |
| 0.047897 |
| 0.0299 |
| 0.004507 |
| 0.025599 |
| -0.02277 |
| -0.00773 |
| -0.01072 |
| 0.03099 |
| 0.008898 |
| 0.020287 |
| -0.00937 |
| -0.02768 |
| -0.02231 |
| -0.00382 |
| -0.01475 |
| 0.03304 |
| -0.02371 |
| 0.023343 |
| 0.066006 |
| -0.01548 |
| 0.015847 |
| -0.03045 |
| -0.01001 |
| -0.00336 |
| 0.018061 |
| -0.00461 |
| -0.03394 |
| 0.015439 |
| 0.024148 |
| -0.00194 |
| -0.01097 |
| -0.00491 |
| 0.043056 |

**Subfigure:** Figure 11B

Plot Name: **Adipose cell proportions by BMI**

| | **BMI 20-40** | | | | |
|---|---|---|---|---|---|
| **DPP4+ Progenitors** | 0.06879195 | 0.757515 | 0.289118 | 0.471191 | 0.179245 |
| **CEBPD+ PreAd** | 0.01971477 | 0.104208 | 0.034771 | 0.145967 | 0.051887 |
| **VIT+Areg** | 0.8091443 | 0.072144 | 0.060528 | 0.06274 | 0.15566 |
| **FABP4+ PreAd** | 0.10234899 | 0.066132 | 0.615583 | 0.320102 | 0.613208 |

| | | | | | | | |
|---|---|---|---|---|---|---|---|
| | | | | | | **BMI 40-60** | |
| 0.339382 | 0.113333 | 0.36002994 | 0.137427 | 0.26776 | | 0.252101 | 0.328729 |
| 0.391801 | 0.62 | 0.1489521 | 0.02924 | 0.420765 | | 0.420168 | 0.185083 |
| 0.06586 | 0.08 | 0.42327844 | 0.716374 | 0.054645 | | 0 | 0.008287 |
| 0.202957 | 0.186667 | 0.06773952 | 0.116959 | 0.256831 | | 0.327731 | 0.477901 |

| | | | | | | | |
|---|---|---|---|---|---|---|---|
| 0.264085 | 0.652568 | 0.409091 | 0.017857 | 0.051724 | 0.136 | 0.378731 | 0.222222 |
| 0.204225 | 0.072508 | 0.318182 | 0.339286 | 0.586207 | 0.128 | 0.104478 | 0.148148 |
| 0.038732 | 0.057402 | 0 | 0 | 0.034483 | 0.048 | 0.018657 | 0.037037 |
| 0.492958 | 0.217523 | 0.272727 | 0.642857 | 0.327586 | 0.688 | 0.498134 | 0.592593 |

| | | | | | | | |
|---|---|---|---|---|---|---|---|
| 0.652655 | 0.501393 | 0.284153 | 0.112782 | 0.12963 | 0.035714 | 0.093884 | 0.080371 |
| 0.028761 | 0.069638 | 0.311475 | 0.067669 | 0.259259 | 0.321429 | 0.829936 | 0.770802 |
| 0.013274 | 0.066852 | 0.019126 | 0.022556 | 0 | 0.035714 | 0.018777 | 0.024395 |
| 0.30531 | 0.362117 | 0.385246 | 0.796992 | 0.611111 | 0.607143 | 0.057403 | 0.124433 |

**Subfigure:** Figure 12E

| | | |
|---|---|---|
| Plot Name: | **PDPN Staining** | |
| **Groups:** | TNFa+IFNy | FCM+TNFa+IFNy |
| **Data Points:** | 58.4 | 50.8 |
| | 44.1 | 47.9 |
| | 50.7 | 36.3 |
| | 65.1 | 43.2 |
| | 52.2 | 38.5 |
| | 43 | 46.6 |
| | 44 | 37.1 |
| | 58.5 | 29.2 |
| | | 33.3 |
| | | 36.8 |

**Subfigure:** Figure 12G

| | | | | |
|---|---|---|---|---|
| Plot Name: | **IL-1B** | | | |
| **Groups:** | Basal | TNFa5ng/mL | TNFa+FCM | TNFa+Cortisol1uM |
| **Data Points:** | 1.726166 | 204.644 | 16.56426 | 9.141897 |
| | 0.579319 | 164.1089 | 8.084131 | 15.08419 |
| | | 88.02391 | 13.52677 | 2.320771 |

| | | | | |
|---|---|---|---|---|
| Plot Name: | **IL6** | | | |
| **Groups:** | Basal | TNFa5ng/mL | TNFa+FCM | TNFa+Cortisol1uM |
| **Data Points:** | 1.680314 | 92.93387 | 15.46743 | 13.72423 |
| | 0.546612 | 109.1123 | 12.96233 | 15.14873 |
| | 1.088755 | 106.9642 | 20.44826 | 13.93619 |
